# Strain-level diversity of giant viruses infecting chlorarachniophyte algae in the subtropical North Pacific

**DOI:** 10.1101/2025.11.19.688996

**Authors:** Max Emil Schön, Christopher R. Schvarcz, Silja V. Malkewitz, Fanny C. Hinner, Anna Koslova, Ulrike Mersdorf, Fiona Schimm, Sebastian Rickert, Nadiia Pozhydaieva, Kelsey McBeain, Thomas Hackl, Alina Cosima Schneider, Karina Barenhoff, Katharina Höfer, Kyle F. Edwards, Grieg F. Steward, Matthias G. Fischer

**Affiliations:** Max Planck Institute for Medical Research, Heidelberg, Germany; University of Hawaiʻi at Mānoa, Honolulu, HI, USA; Institute of Molecular Genetics, Czech Academy of Sciences, Prague, Czech Republic; Max Planck Institute for Terrestrial Microbiology, Marburg, Germany; Groningen University, Groningen, The Netherlands; Department of Pharmacy, Institute of Pharmaceutica Biology and Biotechnology, Philipps Universität Marburg, Marburg, Germany; Center for Synthetic Microbiology (SYNMIKRO), Philipps Universität Marburg, Marburg, Germany

**Author notes:** Max Planck Institute for Marine Microbiology, Bremen, Germany.

## Abstract

Giant DNA viruses are ubiquitous among unicellular eukaryotes and occur in marine, freshwater, and terrestrial environments. Despite intense metagenomic data mining, their strain-level diversity remains largely unexplored. Here we introduce a model system comprising four isolates of a giant virus called ChlorV, which infects marine microalgae of the class Chlorarachniophyceae (Rhizaria) from station ALOHA, Hawai’i. The ChlorV genomes are 469 kbp to 493 kbp long and encode approximately 400 proteins, at least 106 of which are present n purified virions. Although the four viral genomes are highly syntenic, they differ by several insertions and deletions that often encode methyltransferases. Interestingly, we found that some of these methyltransferase genes correlated with specific DNA methylation patterns in the ChlorV strain Our study describes the first giant viruses infecting the eukaryotic supergroup Rhizaria and demonstrates how viral strain-level variation in gene content and epigenetic features may affect eco-evolutionary processes in marine microalgae.

## Introduction

Double-stranded DNA viruses of the phylum *Nucleocytoviricota* are the giants of the viral world, with particle sizes of up to 1.5 µm and genome lengths of up to 2.5 megabases^1–5^ When the first such virus, Acanthamoeba polyphaga mimivirus, was isolated and analyzed, it showed remarkable genetic make-up previously unknown from any other virus^6^ Since then, a large number of additional giant virus isolates and thousands of metagenome-assembled genomes have been published. The phylum *Nucleocytoviricota* currently comprises five orders and 15 families^7^ among them, the family *Mimiviridae* (order *Imitervirales*) is the best-studied. The *Mimiviridae* comprises three subfamilies: *Megamimivirinae Klosneuvirinae* and *Aliimimivirinae* The *Megamimivirinae* include the original mimivirus and relatives such tupanvirus, moumouvirus, megavirus etc., all of which have been isolated on Amoebozoa hosts such as *Acanthamoeba* or *Vermamoeba*^1,8–11^ The *Klosneuvirinae* subfamily contains the isolates Bodo saltans virus, Fadolivirus and Yasminevirus^12–15^ The third subfamily, *Aliimimivirinae* contains the well-characterised Cafeteria roenbergensis virus (CroV) that infects the marine heterotrophic stramenopile *Cafeteria burkhardae* and Chlorella Virus XW01 that was isolated recently on the freshwater green alga *Chlorella sp*^16,17^ Nevertheless, the abundance of mimivirid-like sequence fragments found in eukaryotic genomes as wel as the inferred co-evolutionary history of giant viruses and protists suggests that most microbial eukaryotes are potential hosts for giant viruses^18,19^

The marine alga *Bigelowiella natans* belongs to the chlorarachniophytes, a mixotrophic lineage within the otherwise heterotrophic Rhizaria whose members acquired their plastid from a green alga by secondary endosymbiosis^20,21^ In addition to the nuclear and plastid genomes they retain a remnant of the green algal nucleus, called the nucleomorph^22,23^ *B natans* has been studied in the context of plastid evolution and was the first sequenced genome of any rhizarian species^24^ Blanc et al.^25^ identified several endogenous viral elements in the *B natans* genome assembly, suggesting that this alga frequently interacts with both giant viruses and virophages, the latter being smaller dsDNA viruses that depend on giant viruses for their ow replication^26^ However, no virus infecting chlorarachniophyte has been described to date, limiting insight into virus–host interactions in this clade.

Here, we characterize four isolated strains of Chlorarachniophyte virus (ChlorV), a new species of virulent giant virus isolated from Pacific waters of Hawai’i. Phylogenetically, ChlorVs cluster with members of the subfamily *Aliimimivirinae* thus expanding the confirmed host range of giant viruses to include chlorarachniophytes (Rhizaria). We describe the infection dynamics of ChlorV infection in its algal host and present a comparative genome analysis of the four ChlorV strains including methylation pattern and associated methyltransferase genes.

## Results & Discussion

In search of new algal viruses from tropical marine environments, we collected samples from surface waters at Station ALOHA, long term oceanographic monitoring site located 100 km north of the Hawaiian Island of Oʻahu (Fig. S1). Several strains of chlorarachniophyte algae (Fig. S2) and viruses infecting them were isolated between 2010 and 2012 (Table S1). As these viruses replicate in members of the Chlorarachniophyceae, we refer to them collectively as ‘ChlorVs’ (for **chlor**arachniophyte **v**iruses).

All viruses replicated in chlorarachniophyte host strains, albeit with notable differences in strain specificity. Whereas host strains AL-TEMP06, AL-TEMP07 and AL-DI0 were permissive to all four ChlorV strains, host strain AL-FL05 supported only ChlorV-1 replication and was resistant to the remaining three ChlorV strains. In contrast, host strain AL-FL10 could only be infected by ChlorV-2, but not by the other virus strains. All host strains belong to *Bigelowiella natans* except for strain AL-FL05, which appears to represent a distinct chlorarachniophyte genus (Fig. S2). Because algal strains AL-FL05 and AL-TEMP06 grew most favorably in our culture conditions, we used them as production strains for ChlorV-1 and ChlorV-2 to ChlorV-4, respectively.

Extracellular ChlorV particles were readily detectable by flow cytometry after DNA staining with SYBR Gold (Fig. S3). By infecting several iters of algal cultures, we collected and concentrated approximately 1E+1 virus particles for each of the four ChlorV strains. Imaging virus particles by negative staining transmission electron microscopy revealed projected pentagonal or hexagonal outlines typical of icosahedral capsids (Fig. 1A-D). Similar to the two other isolated viruses of the subfamily *Aliimimivirinae* (CroV and Chlorella virus XW01), their capsids appear to be lacking external fibrils. The capsid diameter varied from 200 to 230 across the four virus strains. Occasionally, small protrusions (13 x 5 nm) at one of the capsid corners were visible, which suggests a modified vertex that could function in host recognition or genome release (Fig. 1A,C)^27,28^

**Figure 1:**
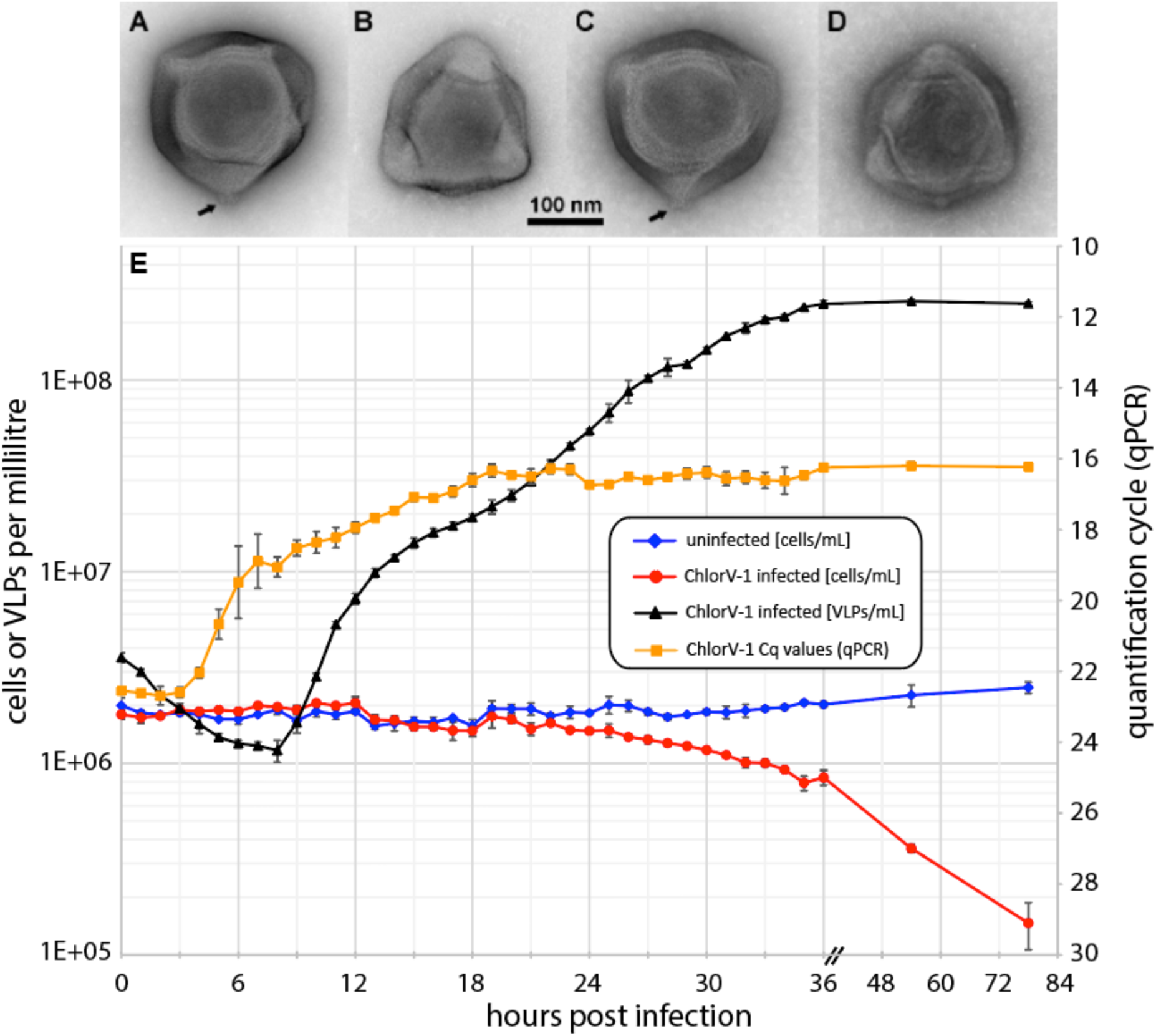
ChlorV particle morphology and infection dynamics. A-D: Negative stain transmission electron micrographs of ChlorV particles. A) ChlorV-1. B) ChlorV-2. C) ChlorV-3. D) ChlorV-4. Arrows indicate potential unique capsid structures such as vertex portals. E) ChlorV- infection dynamics n host strain AL-FL05. Densities of host cells and extracellular virus-like particles (VLPs) of uninfected and ChlorV-1 infected AL-FL05 cultures were measured by flow cytometry. Quantitative PCR was used to monitor viral DNA replication. VLP concentrations and Cq values of uninfected cultures are not plotted, as these were without exception below the limit of detection. Error bars show standard deviations between the three biological replicates and may be smaller than the data symbols.

### Infection dynamics

We analyzed the infection dynamics of ChlorV- in host strain AL-FL05, because this combination resulted consistently in the highest virus titers when compared to any other ChlorV host strain combinations. We exposed the algal cultures to a 12 h light 12 h dark cycle and infected AL-FL05 with ChlorV-1 at the onset of the light cycle. Triplicate cultures of AL-FL05 were infected with freshly prepared ChlorV-1 at a virus-to-host ratio of two and monitored hourly for the first 36 hours by flow cytometry and quantitative PCR using ChlorV-specific primers (Fig. 1E). The qPCR signal remained constant from the time of infection until 4 h post infection (hpi); thereafter, the quantification cycle (Cq) decreased from 22.5 to 16.0 by 19 hpi, indicating the onset of viral DNA replication around 3-4 hpi and a peak in viral DNA by ∼19 hpi. Using flow cytometry, we observed a continuous decrease in extracellular VLPs from 0 hours post infection (hpi) to 8 hpi, most likely due to the uptake of ChlorV-1 particles by host cells. Following this eclipse phase, VLP concentrations in the culture medium ncreased steadily from 9 hpi to 36 hpi. The slope of VLP increase was steeper towards the end of the first light cycle (8-12 hpi) than during the following dark and light cycles (13-36 hpi). Furthermore, we observed a delayed decline of host cells in the infected cultures, which occurs only after extracellular VLP concentrations have reached their maximum at 36 hpi. These data are not compatible with a one-step growth curve and instead suggest a constant release of virus particles. In summary, our infection experiments show that ChlorV-1 is a lytic virus; however, the release of progeny virus particles is not immediately linked to host lysis, suggesting a continuous and, at least initially, non-destructive exit strategy, such as budding from host membranes or exocytosis^29^

### Genome features

After purifying the virus particles through iodixanol density gradients and isolation of viral DNA, we generated high-quality genome assemblies using ultra long MinION sequencing reads at 31x to 700x coverage (Table 1). For ChlorV-1 and ChlorV-4, we also generated short-read Illumina sequencing data to further polish the assemblies. The genomes appear to have a linear topology, as they contained short (1.5 kb to 2.5 kb) terminal inverted repeats (TIR) at both ends, and ranged in size from 469 kb (ChlorV-3) to 493 kb (ChlorV-4). While the genomes of ChlorV-2 and ChlorV-4 were virtually identical with an average nucleotide identity (ANI) exceeding 99.8%, ANIs between other pairs of genomes ranged between 93.2% (ChlorV-1 vs ChlorV-4) and 97.3% (ChlorV-3 vs ChlorV-4), see Fig. S4. The differences (insertions/deletions) between genomes were distributed over their complete lengths, and none of the observed differences were longer than 10,000 bp. n comparison, the ANI between any of the four ChlorV genomes and their most closely related giant virus isolates, Cafeteria roenbergensis virus (CroV) and Chlorella virus XW01, was ∼70%. However, the alignments between Chlorella Virus XW01 and ChlorVs covered about 10-12% of their respective genomes, whereas those between CroV and the ChlorVs covered only ∼4–6% (within ChlorVs, the coverage ranged from 75% to 100%, Fig. S4).

**Table 1:** Genome statistics of ChlorV strains 1-4. In all cases, ONT was used for the initial assembly which was then polished using Illumina data if available. PacBio data for ChlorV-1 was only used to validate the ONT-based assembly.

| Strain | Genome length (bp) | % GC | # ORFs | Coverage (ONT) | Data source | Accession |
| --- | --- | --- | --- | --- | --- | --- |
| ChlorV-1 | 487,585 | 19.05 | 402 | 263 | ONT+PacBio +Illumina | GCA_974392635 |
| ChlorV-2 | 490,841 | 18.54 | 409 | 639 | ONT | GCA_975145565 |
| ChlorV-3 | 468,809 | 18.82 | 391 | 700 | ONT | GCA_976281805 |
| ChlorV-4 | 493,487 | 18.59 | 394 | 31 | ONT+Illumina | GCA_975141545 |

In each of the four ChlorV genomes, we identified ≈400 open reading frames (ORFs) encoding proteins >100 amino acids, with 56 to 72 sequences per genome not matching any known cellular or viral genes (ORFans). Accounting for ORFs that were split n some assemblies, 396 non-redundant genes were identified across all virus strains. A core genome of 307 genes was shared across all strains, while 38 were shared between ChlorV-2 and ChlorV-4, 336 between ChlorV-2, ChlorV-3 and ChlorV-4 and 316 between ChlorV-1 and ChlorV-3.

### ChlorV phylogenetic position

To establish the phylogenetic position of these viruses, we identified putative homologs of established marker genes specific for viruses of the phylum *Nucleocytoviricota*^30^ Each ChlorV genome contained a full complement of these conserved genes, further validating the quality of the assemblies. For each marker gene we reconstructed a phylogenetic tree with broad taxonomic sampling across the *Nucleocytoviricota* In each of these trees, the four ChlorV genomes formed a tight cluster in the *Mimiviridae* family, with CroV and Chlorella Virus XW01 being the closest isolated relatives (Supplementary File 1).

We then reconstructed a phylogenetic tree based on a concatenated alignment of seve marker proteins, focusing on genomes of *Mimiviridae* members (isolates and environmental genomes) from the giant virus database (GVDB). Selected representatives of other *Imitervirales* lineages were chosen as an outgroup. The ChlorVs formed a well-supported monophyletic group with Chlorella Virus XW01, CroV and several environmental genomes in the subfamily *Aliimimivirinae* (Fig. 2).

**Figure 2:**
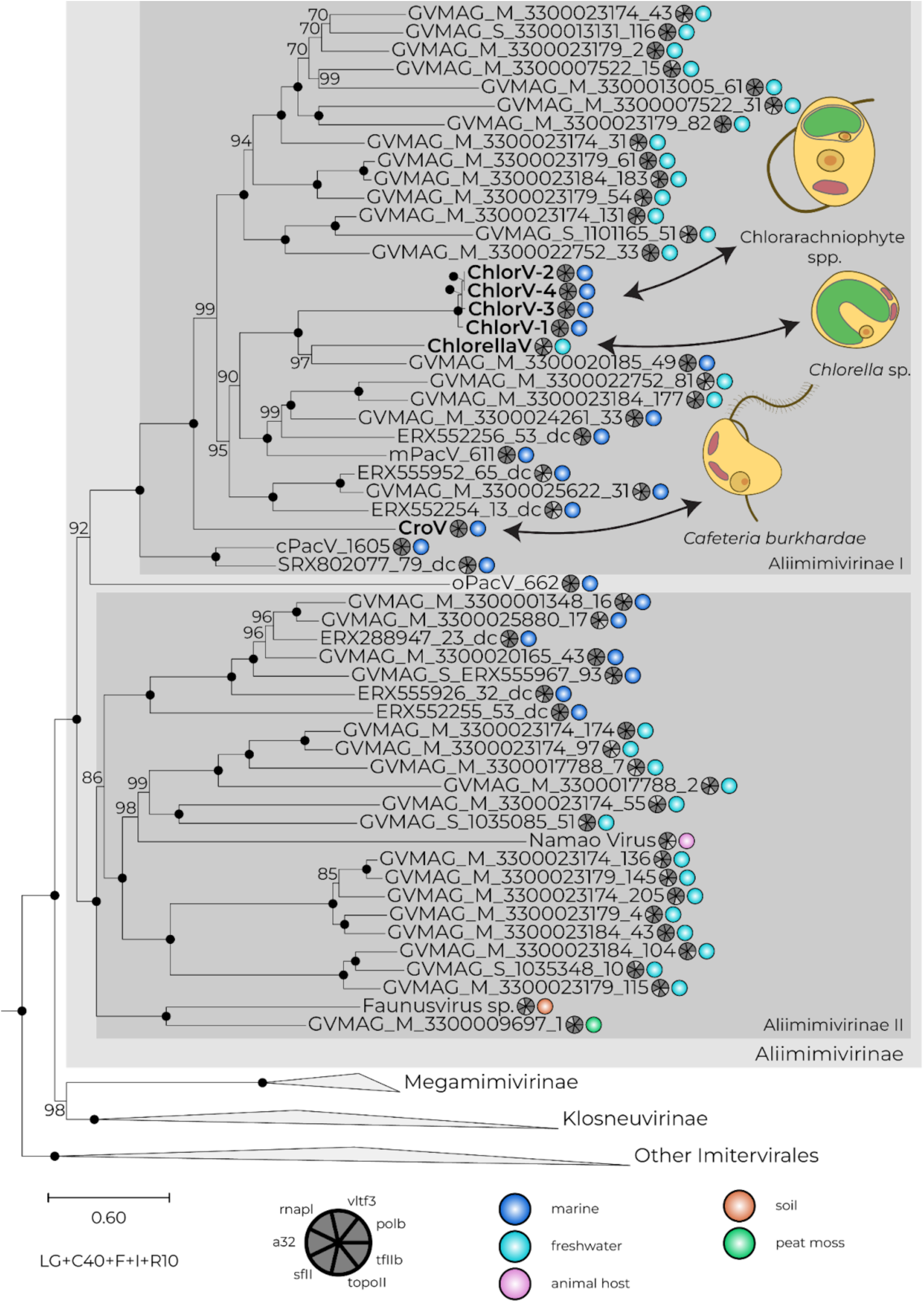
Phylogenetic position of ChlorV isolates. ChlorV 1-4 form a well-supported and closely related group within the *Mimiviridae* subfamily *Aliimimivirinae* Their closest relative isolates are Chlorella Virus XW01 and the Cafeteria roenbergensis virus (CroV) in the same subfamily. The tree is based on the concatenated alignment of 7 marker genes and was inferred under the best fitting evolutionary model LG+C40+F+I+R10, while support was estimated using 1000 ultrafast bootstrap replicates. Besides the phylogenetic position, the distribution of marker genes per genome is drawn as a coulson plot and the source environment as a colored circle (dark blue: marine, light blue: freshwater, pink: animal host, orange: soil, green: peat moss).

The addition of ChlorVs makes the *Aliimimivirinae* one of the most diverse groups of *Nucleocytoviricota* with respect to host range. Three different eukaryotic supergroups (Rhizaria: Chlorarachniophytes; Stramenopiles: *Cafeteria* Archaeplastida: *Chlorella*) are infected by members of this subfamily. In fact, the ChlorVs, CroV and Chlorella Virus XW01 cluster in one group within the *Aliimimivirinae* (*Aliimimivirinae* I), whereas the lake sturgeon-associated Namao virus and various metagenome-assembled genomes form a second group (*Aliimimivirinae* II), indicating two well-supported clades within *Aliimimivirinae* (here referred to as clades I and II; Fig. 2, Fig. S5).

### Genome annotation & Comparative genomics

We annotated the predicted protein sequences using multiple strategies and databases, including EggNOG, InterPro (incorporating PFAM) and the giant virus orthologous groups (GVOGs)^30–33^ Besides ORFans with o matches to any known sequences, 128-146 ORFs per genome only matched poorly characterized reference sequences, often predicted from other giant viruses. Of the 173-190 sequences with a robust annotation, most were categorized as ‘hydrolases’ (e.g. peptidases or nucleases), ‘replication, transcription and DNA repair’ (e.g. helicases, replication factors, polymerases) or ‘transferases’ (e.g. methyl-and glycosyltransferases) (Table S2). The ChlorV genomes encode a full complement of DNA replication genes, including family B DNA polymerase (ChlorV-1..065), a DNA clamp (ChlorV-1..016) and clamp loader (replication factor C, e.g. ChlorV-1..164). They further encode a DNA ligase (ChlorV-1..107) and a ribonuclease H (ChlorV-1..329) as well as multiple helicases including a D5-like helicase-primase (ChlorV-1..361) and two subunits of a ribonucleotide reductase (ChlorV-1..059 and ChlorV-1..370). Transcription-related enzymes are also encoded in the genomes, including several subunits of the RNA polymerase (Rpb1,2,3/11,5,6,7, ChlorV-1..202,039,325,253,359,251), an mRNA capping enzyme (ChlorV-1..311), a poly(A) polymerase (ChlorV-1..312) and several transcription factors including homologs of the Poxvirus VLTF2 (ChlorV-1..152) and VLTF3 (ChlorV-1..384). Several translation initiation (e.g. eIF4E, ChlorV-1..017) and elongation (EF-Tu, ChlorV-1..328) factors are also present. A homolog of the DNA polymerase beta (ChlorV-1..031) that is involved in base excision repair (BER), a putative DNA mismatch repair protein MutS (ChlorV-1..354) and two DNA photolyases (ChlorV-1..224 and ChlorV-1..336) that are potentially involved in DNA repair outside of the host are encoded as well Additionally encoded proteins include many peptidases (e.g. Chlorv-1..086) and several ankyrin repeat proteins (e.g. Chlorv-1..071). Further noteworthy is the presence of a putative YqaJ-like viral recombinase (ChlorV-1..348) which is involved in cleaving genomic concatemers in some bacteriophages^34^ A putative tubulin–tyrosine ligase (Chlorv-1..043) could be involved in manipulation of host cells^35^ Several glycosyltransferases (e.g. Chlorv-1..212) and glycosylases (e.g. ChlorV-1..397) are encoded in the genomes of all strains and could be involved in capsid modifications and host interactions^36^ At least one acetyltransferase (ChlorV-1..005) and one aminotransferase (ChlorV-1..145) were also present.

Each ChlorV genome encoded multiple predicted methyltransferase genes (12 in ChlorV-3, 11 in the other strains), and these were often located in strain-specific insertions or deletions (Fig. 3 A-G). In contrast, most other groups of genes were usually conserved across all strains. Several types of nucleases (both RNases and DNases) were also commo throughout the genomes, being either shared among strains or strain specific (e.g. the DNA endonuclease ChlorV-1..079, Fig. 3 D, F).

**Figure 3:**
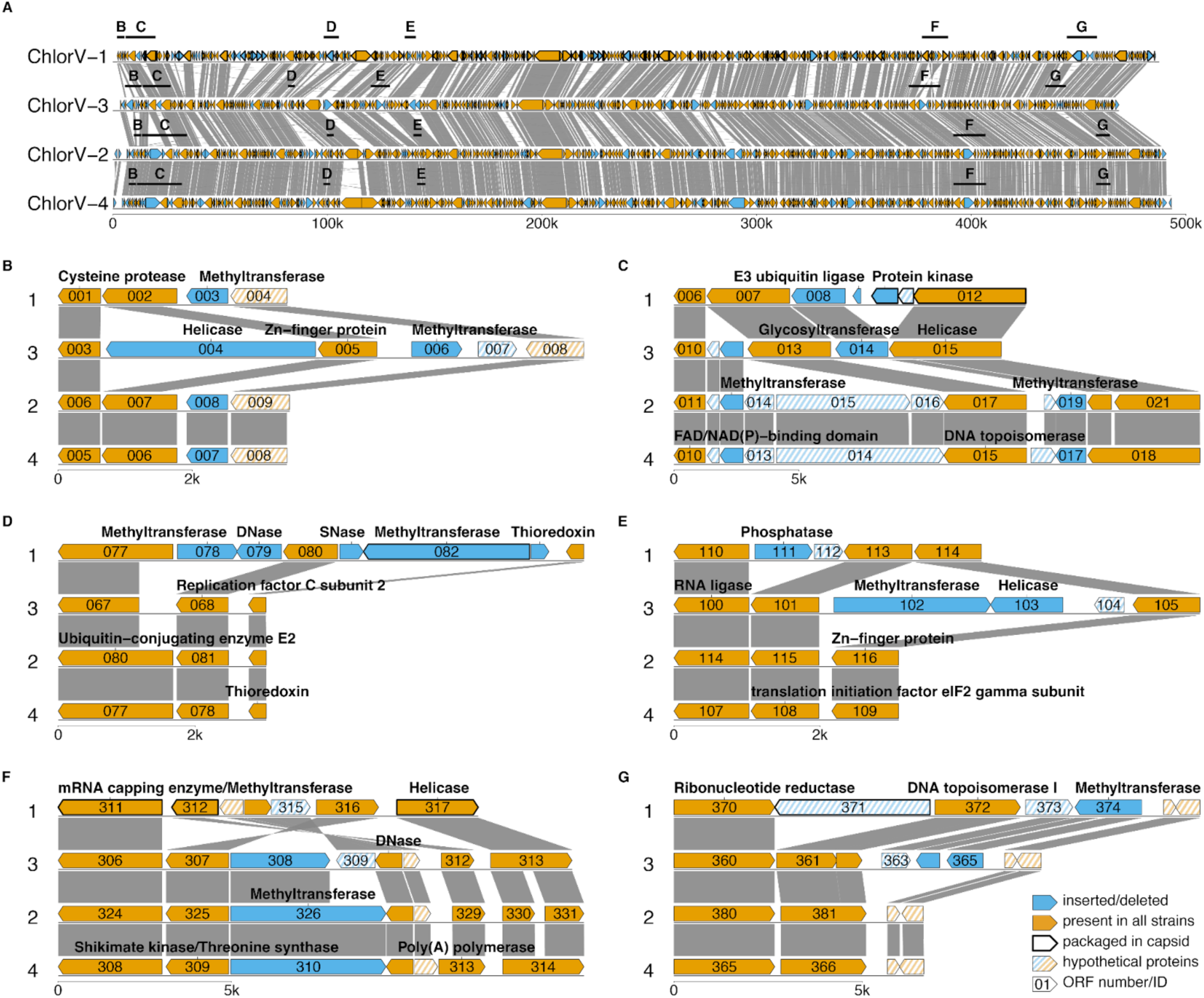
ChlorV strain level genomics. C: Overview of the complete genomes from four strains of Chlorarachniophyte viruses (ChlorVs). Genes are colored according to whether they wer identified in all strains (orange) or absent in at least one strain (blue). Genes with significant sequence matches are connected by dark grey ribbons. A,B,D-G: Details of genomic loci where putative insertions/deletions and an inversion (F) can be observed. DNase: Deoxyribonuclease, SNase: Staphylococcal nuclease (DNA and RNA endo-exonuclease)

We then predicted the 3D structure of all ChlorV-1 ORFs using AlphaFold 3 and compared them to experimentally derived structures in the protein database to improve the genome annotation (Supplementary File 2)^37,38^ For most ORFs, the sequence-based and structure-based annotations (where available) agreed well. However, occasionally the structural comparison provided annotations for ORFs with no sequence-based annotation, e.g. for the endonuclease ChlorV-1..079 or the putative penton protein Chlorv-1..095 (Table S3). In other cases, sequence-based annotation could not be substantiated by the structural comparison, e.g. the predicted homing endonuclease Chlorv-1..154.

Whereas CroV encodes many FNIP/IP22 repeat proteins, these repeats are absent from ChlorVs, Chlorella virus XW01, and *Aliimimivirinae* environmental genomes. Besides the protein-coding genes, ChlorV strains encode up to 7 tRNA genes, in contrast to the high number of 48 that are encoded in the genome of CroV, whereas Chlorella Virus XW01 encodes none.

We also investigated putative promoter sequences of ChlorV genes based the phylogenetic position close to CroV and previous work on gene promoters in giant viruses generally^39,16,40^ The early motif (AAAAATTGA) and the late motif (TCTA) appear conserved between CroV and ChlorVs in both sequence and positional context (Fig. S6). Unlike the promoters, gene content seems to be more dynamic over larger phylogenetic distances (Fig. S5), which has also been observed in other groups of giant viruses.

#### Genome modifications

For all ChlorV genomes, nanopore sequencing data was used to analyze DNA base modification patterns. In all assemblies, the dominant modification was N6-methyladenine (6mA), followed by C5-methylcytosine (5mC) and N4-methylcytosine (4mC). The latter two were rare, except in ChlorV-1 where 5mC was comparatively frequent (Fig. S7). This pattern confirms previous observations that giant viruses display prokaryotic-like DNA modifications with a majority of 6mA^41^ We identified sequence motifs that were highly enriched (all were modified in >90% of occurrences) in methylated bases across the four genomes. We identified three distinct motifs, including two motifs that were shared by two genomes each (Fig. 4A). All motifs were palindromic; accordingly, we did not detect strand-specific differences. ChlorV-1 and ChlorV-3 display a 10 base pair motif (C**A**NNNNNNTG) that resembles a bipartite recognition sequence, consistent with Type or certain Type I systems^42,43^ We identified one gene that is present in ChlorV-1 and ChlorV-3 but not the other strains as a candidate methyltransferase targeting this sequence (Fig. 3G). It possesses a 6-adenine-specific methyltransferase domain as well as a single target specificity domain similar to other type modification genes (Fig. 4B). However, this enzyme could also function as a homooligomer, recognizing and modifying the same DNA sequence on opposite strands, like the type IIB system HaeIV^44^ This is supported by the high confidence scores (ipTM=0.8, pTM=0.85) of a protein-DNA complex containing two copies of this protein together with a DNA sequence containing the motif C**A**NNNNNNTG, but lower confidences (ipTM 0.8) if using only a single copy or another DNA sequence (Table S4, Fig. S8). ChlorV-3 (but not ChlorV-1) encodes another candidate methyltransferase with similar DNA methyltransferase domain and two specificity domains, more similar to typical type modification genes^45^ (Fig. 4E). We could additionally identify a motif for 5mC methylation (R**C**GY) in ChlorV-1. This motif could be the target of a putative 5-cytosine-specific methyltransferase encoded only in the ChlorV-1 genome (Fig. 3D, Fig. 4C). Right next to this methyltransferase endonuclease (ChlorV-1..079) encoded, suggesting that this is indeed a functioning restriction modification system. Finally, we identified a different 6mA motif (TGC**A**) in ChlorV-2 and ChlorV-4. This could again be putatively linked to a 6-adenine-specific methyltransferase gene that resembles type IV restriction modification methyltransferases and was encoded only in these two strains (Fig. 3C, Fig. 4A, D).

**Figure 4:**
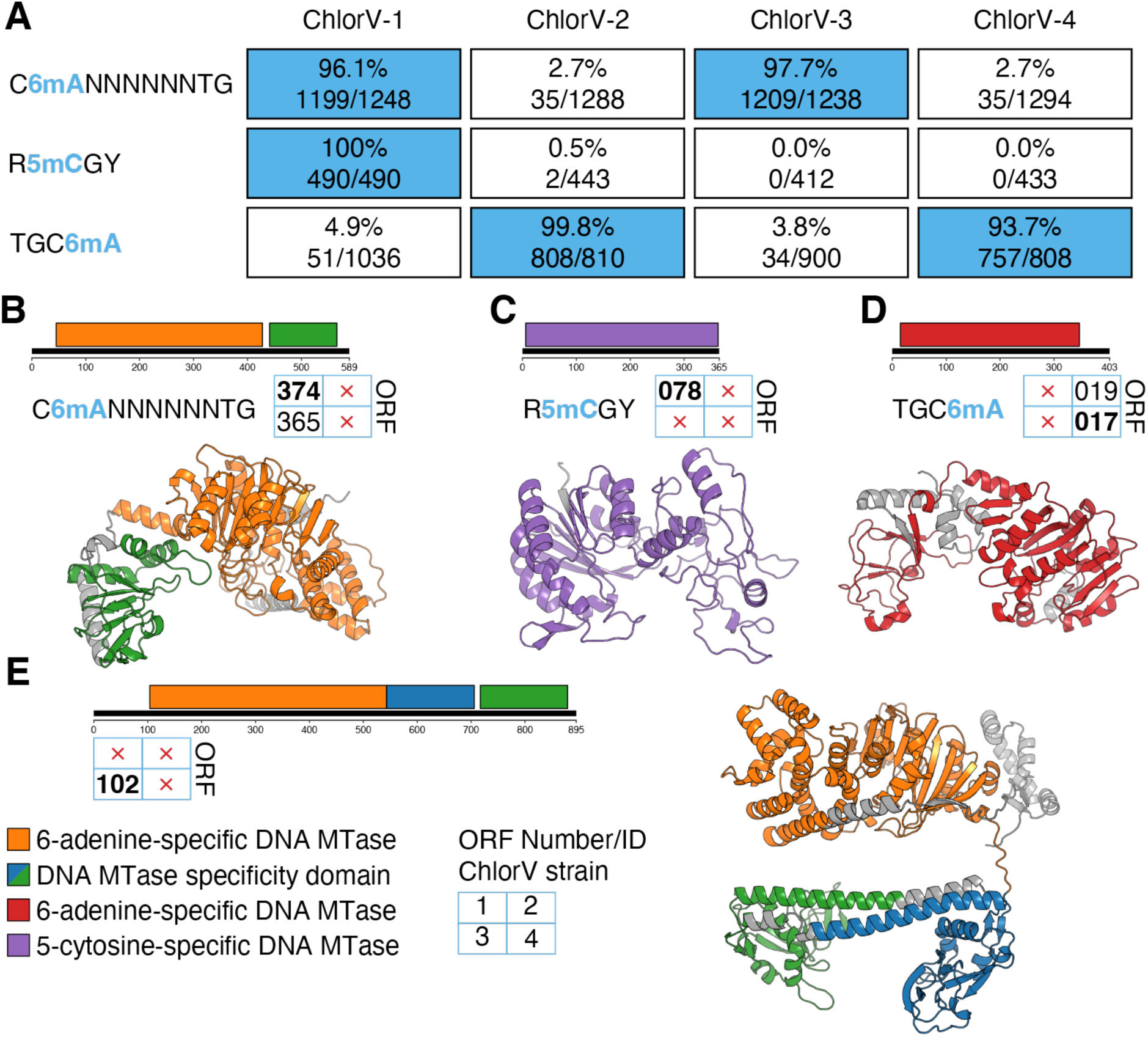
DNA modifications in ChlorVs. A: Detected sequence motifs with strong (>99%) modification signal and putative associated methyltransferases. ChlorV-3..102 is a candidate besides ChlorV-3..365 for methylation of the CANNNNNNTG motif, although no homologue of ChlorV-3..102 could be identified in ChlorV-1. For each methyltransferase gene, domain-level annotations are shown. Three genes represent 6-adenine-specific DNA methyltransferases (with ChlorV-4..019 having a different domain than e.g. ChlorV-1..374), while ChlorV-1..078 was inferred to modify cytosine. B-E: Structural models of the four presented methyltransferases. n each model, the sequence domain annotations are visualized using the same colors.

#### Virion proteomics

We analyzed the protein composition of purified ChlorV-1 virions by mass spectrometry and identified 106 vira proteins, each represented by at least two peptides (Table S5). These virion proteins were encoded by genes that are spread fairly evenly across the viral genome (Fig. S9). However, nine of the ten most abundant virion proteins were encoded by genes that located within 90 kb central region of the genome, indicating non-random distribution of important structural genes The most abundant protein was the double jelly roll (DJR) major capsid protein ChlorV-1..153, followed by the major core protein ChlorV-1..167 which is homologous to the core proteins of poxvirus (P4B) and CroV (Crov332). It is noteworthy that the third-most abundant virion protein was ChlorV-1..175, a 709 amino acid long hypothetical protein that lacked homologs in other ChlorV strains. Peptides for all three additional DJR capsid paralogs, well for two of the four single jelly roll (SJR) penton protein candidates, were also found. We then generated structural predictions for three of the four DJR proteins as trimers and for the four SJR proteins as pentamers (Fig. S10, S11). The structures were generally predicted with high confidence, except for some surface protrusions that might interact with other capsid proteins. The peculiarly structured SJR protein ChlorV-1..087 had a lower confidence value, although the core penton-like structure was still resolved with good support (Fig. S11). We did not predict hetero-multimers for pseudo-hexameric capsomers involving more than one DJR protein, although it is quite possible that such capsomers exist in ChlorVs, similar to e.g. Paramecium bursaria Chlorella virus 1 (PBCV-1)^46^

Among virion proteins with predicted enzymatic functions, we identified the two predicted DNA photolyases ChlorV-1..224 and ChlorV-1..336. As is typical for mimivirids, ChlorV-1 also packages its own transcription system that is represented here by five RNA polymerase subunits, mRNA capping enzyme, poly(A) polymerase, and two transcription factors. All of these transcription proteins are also encoded and packaged by CroV^16,47^

## Methods

### Isolation of hosts and viruses

Strains of diverse eukaryotic phytoplankton were isolated from seawater samples collected from an oligotrophic open-ocean site (Station ALOHA, 22°45’ N, 158°00’ W)^48^ located approximately 100 km north of O‘ahu. For collection dates, see Table S1. Seawater samples were typically enriched with artificial seawater media, F/2 medium or Keller (K) medium^49,50^ and allowed to incubate in bottles or multiwell plates until cells of interest were identified. These cells were then isolated either by micropipette or by serial dilution. All phytoplankton cultures were maintained at 24–26 °C on 12:12 light:dark cycle with approximately 30–100 µmol photons m-2 s-1 of photosynthetically active radiation (PAR).

Small subunit ribosomal RNA (18S rRNA) gene sequencing was used initially to identify algal species. For this purpose, cells were pelleted (4000 RCF for 10 min) and DNA extracted and purified (MasterPure Complete DNA and RNA Purification Kit; LGC Biosearch Technologies) or ZymoBIOMICS DNA Mini Kit; Zymo Research). The near-full length gene was amplified by PCR (Expand High Fidelity PCR System; Roche) with forward (5ʻ- ACC TGG TTG ATC CTG CC AG -3ʻ) and reverse (5ʻ- TGA TCC TTC YGC AGG TTC AC -3ʻ) primers targeting the 5’ and 3’ ends of the gene^51^ PCR products were cloned prior to sequencing (TOPO TA Cloning Kit for Sequencing; Invitrogen), and the insert was sequenced using four separate primers (M13f, M13r, 502f, and 1174r)^52^ using a fluorescence-based Sanger technique (BigDye Terminator; Applied Biosystems) and capillary electrophoresis (3730XL DNA Analyzer; Applied Biosystems) at the University of Hawai‘i at Mānoa’s facility for Advanced Studies in Genomics, Proteomics, and Bioinformatics (ASGPB).

Viruses were isolated from seawater samples collected from the open-ocean Station ALOHA sampling site. Large volumes of seawater were pre-filtered (using either 0.8 µm or 0.45 µm pore size filters) to partially remove cells, and the virus community was concentrated by tangential flow filtration (TFF; Millipore Pellicon 2 Mini System) using 30 kDa nominal molecular weight limit (NMWL) filters. The viral concentrates were amended with media (f/2 or K) and added to healthy phytoplankton cultures. The virus-challenged cultures were observed for 1–2 weeks for signs of cell lysis, and lysates that continued to produce lytic effect after multiple transfers to healthy cells (putatively virus-containing) were stored at 4 °C and propagated at least once per month by challenging new cells (1–10% v/v of lysate added per challenge).

#### Infection experiments

Chlorarachniophyte cultures were grown in K medium in sterile polycarbonate flasks, either in 250 mL Erlenmeyer flasks, or in 3 L Fernbach flasks. Cultures were grown in an incubator at 24 °C, 50 rpm shaking, and in 12 h light (at 1300 to 1500 lux) 12 h dark conditions. Immediately prior to inoculation, host cultures were diluted with K medium to a density of 1.0E+06 cells per mL and aliquoted in 50 ml portions in 250 ml PC Erlenmeyer flasks. Triplicate cultures were inoculated at the onset of the 12 h light cycle with a fresh (less than 2 weeks old, 1.2 µm filtered and stored at 4 °C) ChlorV-1 suspension at a virus-to-host ratio of two. Three flasks with uninfected host cultures were incubated as a negative control. The six flasks were sampled every hour for the first 36 hours, and again at 3 dpi and 5 dpi. At each timepoint, the densities of cells and free virus particles were analyzed immediately by flow cytometry, and 25 µl of each suspension culture were frozen for later qPCR analysis.

#### Flow cytometry

For cell counts, 200 µl of suspension culture were transferred to a 1.5 ml microfuge tube and analyzed o an Attune^TM^ NxT Acoustic Focusing Flow Cytometer (ThermoFisher Scientific, Waltham, MA, USA) and proprietary software, with the following instrument settings: Red Laser (RL1-H, voltage 320, threshold 1000) for chlorophyll autofluorescence over side scatter (SSC-H, voltage 350, threshold 1000); blue laser filter slot 1 488/10+OD2; 50 µl acquisition volume, 25 µl/min flow rate, 25 µl total acquisition. For virus particle counts, 10 µl of suspension culture were pipetted in a 1.5 ml microfuge tube containing 2 µl of 25% EM-grade glutaraldehyde solution (Sigma Aldrich) and 88 µl of 0.1 µm filtered K medium. After incubation at 4 C for 10 min, 860 µl of 0.1 µm filtered Tris-EDTA buffer and 40 µl of 100x SYBR Gold solution (Life Technologies Corporation, Eugene, OR, USA) added. The mixture was incubated at 80 C for 10 min in the dark, allowed to cool to room temperature, and analyzed on an Attune^TM^ NxT Flow Cytometer with the following settings: Blue Laser (BL1-H, voltage 490, threshold 5000) for SYBR fluorescence over side scatter (SSC-H, voltage 340, no threshold); blue laser filter slot 1: 488/10 (small particle filter); 50 µl acquisition volume, 25 µl/min flow rate, 25 µl total acquisition. Gates for cells and virus particles were set manually, and concentrations derived directly from the Attune^TM^ software.

#### Quantitative PCR analysis

Sample lysis for qPCR use was performed by adding 50 µL of 2x lysis buffer (20 mM Tris-HCl, pH 8.0, 2 mM EDTA, 0.002% Triton X-100, 0.0002% SDS, 2 mg/ml proteinase K) and 25 µL of distilled H_2_O to 25 µl of sample (stored at -20 °C). The tubes were mixed by vortexing and briefly centrifuged, then incubated for 60 minutes at 58 °C and for 10 minutes at 95 °C. Each 20 µl qPCR reaction consisted of 10 µl PowerTrack SYBR Green Master Mix (ThermoFisher), 7 µl nuclease-free water, 0.5 µl of 100 µM forward primer, 0.5 µl of 100 µM reverse primer, and 2 µl of the lysed sample. For ChlorV-1 detection, primers MCP_ChlorV-1_F (5’- CAG CAA TAC TCC ATC CGA CGG -3’) and MCP_ChlorV-1_R (5’-ACT GTT AGC ACC GCC AAT GT -3’) were used, for detection of ChlorV-2, ChlorV-3, and ChlorV-4 we used primers MCP_ChlorV-2-4_F (5’ GTG CTG CAC CTA CTG TTC CT -3’) and MCP_ChlorV-2-4_R (5’ TAG CAG CTT CCC AAT CAC CG- 3’). Samples were analyzed in technical duplicates per 96-well plate, using Mx3005P Real-Time PCR System (Stratagene) with the following cycling conditions: cycle of 7 min at 95 °C, 40 cycles of 15 sec at 95 °C and min at 60 °C, cycle of min at 72 °C, and a 55-95 °C ramping for dissociation curve analysis.

#### Sequencing of ChlorV genomes

ChlorV-1 samples for Illumina and PacBio sequencing were prepared using 0.45 µm filtration (Millipore Durapore 142mm filter, overlaid with Whatman GF/C filter) followed by tangential flow filtration (30 kDa NMWL; Millipore Pellicon 2 Mini system). Viral particles were purified using centrifugation onto CsCl density cushions (6.5 mL 1.6 g/mL CsCl solution, overlaid with 2 mL 1.1 g/mL CsCl solution; 25,000 rpm for 50 min, using SW28 rotor) followed by centrifugation on a CsCl continuous gradient and 0.45 µm filtration (Millipore Sterivex) followed by another round on a CsCl continuous gradient. CsCl in the virus-containing fraction was exchanged with 1X TE buffer by three rounds of concentration and dilution in a centrifugal ultrafilter (30 kDa NMWL; Millipore Amicon Ultra-0.5) followed by DNA extraction and purification (MasterPure Complete DNA and RNA Purification Kit). Sequencing runs for ChlorV-1 were performed on an Illumina MiSeq Instrument (250bp paired-end) using the Nextera XT library preparation kit at the Georgia Genomics and Bioinformatics Core. PacBio sequencing was performed o a PacBio RS I with P6-C4 chemistry at the University of Washington PacBio Sequencing Services.

For Nanopore MinION sequencing, several liters of ChlorV suspension containing at least 1E+08 virus particles per millilitre were centrifuged in Nalgene L centrifuge bottles in a Fiberlite™ F9-6 1000 LEX rotor (ThermoFisher) for 20 minutes at 6000 g, 18 °C. The supernatant was concentrated by tangential flow filtration in a Vivaflow 200 0.2 µm PES unit (Sartorius) to 30-50 ml final volume. The concentrate was loaded on a 10-40% (w/v) linear Optiprep gradient (in 50 mM Tris-HCl, pH 8.0, 250 mM NaCl, 2 mM MgCl_2_) and centrifuged in 14 ml Ultra-Clear SW40 tubes and an SW40 rotor (Beckman) for 2 h at 100,000 x g, 18°C. The visible virus band extracted through the tube wall with syringe and needle, diluted 3-fold with 50 mM Tris-HCl, pH 8.0, 2 mM MgCl_2_ and the virus particles were pelleted by centrifugation (30 min at 20,000 g, 18 °C) in 1.5 m microfuge tubes, pellets were pooled, washed, and pelleted again. The final virus sample was resuspended in 200 µl of 50 mM Tris-HCl, pH 8.0, 2 mM MgCl_2_ for DNA extraction with the QIAamp DNA Mini Kit (Qiagen) following the manufacturer’s instructions, but eluting in 50 µl of nuclease-free water. The genomes of ChlorV-1, ChlorV-2 and ChlorV-3 were sequenced using the SQK-LSK114 kit and a MinION FLO-MIN114 flow cell; the genome of ChlorV-4 was sequenced with an SQK-LSK110 kit and a FLO-MIN106 flow cell (Oxford Nanopore Technologies). For sequencing details, see Table S6.

In addition, a ChlorV-4 DNA sample was sent to Eurofins Genomics Germany GmbH, where it was sequenced on an Illumina NovaSeq 6000 S4 instrument using the TruSeq DNA PCR-Free library preparation kit.

#### Proteomics sample preparation

A virus suspension with a concentration exceeding 10^10 PFU/ml was used for the determination of structural proteomics. Virus particles were purified using a linear idioxanol (Optiprep) gradient according to the protocol above for Nanopore sequencing. The purified virions were resuspended in lysis buffer (50 mM Tris-HCl, pH 7.5, 1% SLS, 2 mM TCEP) and heated to 95 °C, 10 min. A sonication step (10 sec, 20% amplitude, 0.5 pulse) performed to degrade nucleic acids in the samples. Following this, iodoacetamide was added to the final concentration of 4 mM, and the samples were incubated for 30 minutes under light protection.

A total of 20 µl each of Sera-Mag carboxylated magnetic beads (Cytivia) and Sera-Mag carboxylated magnetic beads with 0.05% Azide (Cytivia) were mixed, followed by two rounds of washing with 200 µl water and resuspension in 100 µl water. Subsequently, 4 µl of the bead mixture was used for protein purification. To this, a protein sample adjusted to a final acetonitrile (ACN) concentration of 70% (v/v) was added. The mixture was vortexed before the supernatant was discarded after the magnetic separation of the beads. The beads were washed twice with 300 µl of 70% Ethanol, followed by a wash with 200 µl of ACN, and were subsequently dried. For protein digestion, 100 µl of digestion buffer (50 mM ammonium bicarbonate with µg of trypsin) was added to the beads. The mixture was incubated overnight at 30 °C and 1200 rpm. The supernatant was collected and the beads were further washed with 100 µl of water, which was then combined with the initially collected supernatant. To acidify the final sample, containing purified and digested proteins, 30 µl of 5% Trifluoroacetic acid (TFA) were added.

The resulting supernatant was desalted for mass spectrometric analysis using C18 solid-phase columns (Chromabond C18 spin columns, Macherey Nagel, Düren, Germany). After desalting, the solvent was removed by evaporation and dried peptides were stored at-20 °C until further use.

#### Peptide analysis using liquid chromatography and mass spectrometry

Dried peptides were reconstituted in 0.1% trifluoroacetic acid and then analyzed using liquid-chromatography-mass spectrometry carried out on a Exploris 480 instrument connected to an Ultimate 3000 RSLC nano and a nanospray flex ion source (all Thermo Scientific). Peptide separation performed phase HPLC column (75 µm 42 cm) packed in-house with C18 resin (2.4 µm; Dr. Maisch). The following separating gradient was used: 98% solvent A (0.15% formic acid) and 5% solvent B (99.85% acetonitrile, 0.15% formic acid) to 30% solvent B over 45 minutes at a flow rate of 300 nl/min.

The data acquisition mode set to obtain high-resolution MS at resolution of 60,000 full width at half maximum (at m/z 200) followed by MS/MS scans of the most intense ions within s (cycle 1s). To increase the efficiency of MS/MS attempts, the charged state screening modus was enabled to exclude unassigned and singly charged ions. The dynamic exclusion duration was set to 14 sec The ion accumulation time was set to 50 ms (MS) and 50 at 17,500 resolution (MS/MS). The automatic gain control (AGC) set to 3x106 for MS survey scan and 2x105 for MS/MS scans.

For spectral-based assessment, MS raw files searches were carried out using MSFragger embedded within Scaffold 4 (Proteome Software) with 20 ppm peptide and fragment tolerance with Carbamidomethylation (C) as fixed, and oxidation (M) as variable modification using a uniprot protein database.

#### Host sequencing and 18S+28S trees

We extracted genomic DNA from uninfected host cultures Each sample then sequenced o an ONT MinION flow cell and basecalled (for specific flow cell and model versions, see Table S7). Sequences were assembled as described below for viral genomes. The minimal read length was 1000 bp, minimal Phred quality score 5 except for AL-FL05 where we used a minimum of 10.

We extracted the full 18S and 28S sequences from the initial assemblies using barrnap^53^ Together with a selection of chlorarachniophytes and an outgroup from Rhizaria (Table S7), each gene was aligned separately using MAFFT-G-INS-i v.7.505^54^ and trimmed using trimAl -gt 0.1 v.1.4.rev15^55^ Both trimmed alignments were then concatenated and used to reconstruct a phylogenetic tree using IQ-TREE v2.0.3^56^ (--ufboot 1000 --bnni -m MFP^57^), selecting TIM3+F+R3 as the best fitting model.

### Assembly

Raw Nanopore signals were basecalled with Dorado v0.5.0 (ChlorV-1; duplex mode) using the highest-accuracy model dna_r10.4.1_e8.2_400bps_sup@v4.1.0 (ChlorV-1). For a full list of all basecalling models and versions of dorado see Table S6.

The Nanopore reads were assembled using MetaFlye v.2.9.1-b1780^58^ with options ‘--iterations 3 --nano-hq’ after filtering reads with varying thresholds for length (ChlorV-1: 10000, ChlorV-2: 1000, ChlorV-3: 1000, ChlorV-4 3000) and basecalling Phred quality score (ChlorV-1: 20, ChlorV-2: 5, ChlorV-3: 15, ChlorV-4: 15). Assemblies were further polished using racon v.1.5.0 with options ‘-m 8 -x -6 -g -8 -w 500’ and medaka v.1.11.3 consensus If available (i.e. ChlorV-1 and ChlorV-4), Illumina reads were used to polish assemblies using polypolish v.0.6.0^59^

Average nucleotide identity (ANI) and the corresponding alignment coverage between assemblies and related genomes of isolated viruses was assessed with pyani v0.3.0-alpha^60^ (average_nucleotide_identity.py -m ANIb --fragsize 500).

Open reading frames (ORFs) were identified using prodigal v.2.6.3^61^ and GeneMark v.4.32 Both sets of ORFs were then merged, picking longer ORFs in case of overlapping features and disregarding ORFs coding for less than 100 amino acids.

#### Phylogenetic analyses

The seven marker genes A32 genome packaging ATPase, protein-primed DNA polymerase B, large subunit of the RNA polymerase, superfamily helicase, transcription factor II B, topoisomerase II and the poxvirus late transcription factor VLTF3 were used for phylogenetic reconstruction. For each marker, DIAMOND protein similarity searches (v.2.0.15.153^62^ BLASTp mode) between the predicted protein sequences and the reference sequences from the GVOG database were computed. Alignments were then computed per marker (MAFFT-E-INS-i v.7.505^54^ trimAl -gt 0.1 v.1.4.rev15^55^) for a taxon sampling including all *Mimiviridae* sequences and other *Imitervirales* from the GVDB^30^ as an outgroup. Single gene trees were reconstructed using IQ-TREE v2.0.3^56^ (--ufboot 1000 -m MFP^57^ -mset LG) and inspected for duplicates, false positives etc. Curated marker sets were re-aligned as above and concatenated. Final tree reconstruction was performed with IQ-TREE (-m MFP --mset LG,LG+C10,LG+C20,LG+C30,LG+C40,LG4X --ufboot 1000), selecting LG+C40+F+I+R10 as the best fitting model.

#### Annotation & comparative genomics

Predicted protein sequences were functionally annotated using the eggnog-mapper v.2.1.12^32^ using the eggNOG database v.5.0.2^31^ the GVOG database (using HMMer v.3.3.2^63^) and InterProScan v.5.62_94.0^33^ tRNAs were annotated using tRNAscan-SE v.2.0.9^64^ Taxonomic annotation of the proteins was performed with diamond v.2.0.15.153^62^ against the NR database. Transposable elements were identified using various tools as implemented in the reasonaTE pipeline v.1.0^65^

The MEME tool v.5.5.0^66^ was used for predicting candidate promoter motifs. First, the upstream sequences of each CDS were partitioned based on the presence of the subsequence ‘TCTA’. The tool was then run on the sequences not containing ‘TCTA’ and spanning 50 bases upstream and the sequences that do contain this subsequence and spanning 40 bases upstream of a CDS. This partitioning was necessary to improve motif signals, as o transcriptomic data for predicting ‘early’ and ‘late’ genes^16^ was present. The two candidate motifs were then tested on all upstream sequences using the tools CENTRIMO and FIMO from the MEME suite v.5.5.0.

For each predicted protein sequence of ChlorV-1, we applied AlphaFold v.3.0.1^37^ to infer its 3D structure. Each predicted structure was then used as a query to search the PDB database using foldseek v.8.ef4e960^38^ ‘easy-search’. Alignments were visualized with Open-Source PyMOL v.3.1.0^67^ For the four identified penton proteins and 3 DJR capsid proteins we additionally performed pentameric or trimeric structural predictions using AlphaFold v.3.0.1, respectively.

All predicted proteins from all isolate genomes and MAGs that were classified as *Aliimimivirinae* in GVDB^30^ were subjected to ortholog identification using OrthoFinder^68^ v.2.5.4. For the different strains of ChlorVs, CroV and Chlorella Virus XW01, all-vs-all proteins searches performed using MMseqs2^69^ v.14.7e284. Based these searches, genes in ChlorV genomes with at least 70% sequence identity were manually grouped together.

#### Modifications

For ChlorV-2,3 and 4, basecalling of modified bases using dorado v.0.7.0 performed using the context-free models dna_r10.4.1_e8.2_400bps_sup@v5.0.0_4mC_5mC@v3 for 4mC and 5mC methylations and dna_r10.4.1_e8.2_400bps_sup@v5.0.0_6mA@v3 for 6mA methylations. For ChlorV-1, the rerio^70^ models res_dna_r10.4.1_e8.2_400bps_sup@v4.0.1_6mA@v2 and res_dna_r10.4.1_e8.2_400bps_sup@v4.3.0_4mC_5mC@v1 used. The basecalled reads were mapped to the respective genome using minimap2^71^ v.2.24-r1122 and summarized per-base using modkit^72^ pileup v.0.4.2. Modkit motif was used to find short sequence motifs that were enriched in methylations.

For each of the four DNA methyltransferases putatively linked to a methylation motif, we performed additionally AlphaFold 3 structural predictions including DNA Sequences either including the motif or not, and including the protein both as a monomer or as a monodimer. The genomes additionally scanned for restriction-modification enzymes and systems using DefenseFinder^73^ v.2.0.0 and diamond v.2.0.15.153 searches against the Restriction Enzyme Database^43^

### Data and code availability

All sequence data, including raw nanopore, PacBio and Illumina reads, assemblies and host rRNA genes (OZ264022-OZ264049) have been deposited in the European Nucleotide Archive under project PRJEB90372 Further data such as multiple sequence alignments, phylogenetic trees, AlphaFold 3 models and other relevant data have been deposited in Edmond (https://doi.org/10.17617/3.CC4JL7). Custom code and scripts are available at https://github.com/maxemil/chlorv-genomes.

## Acknowledgements

We would like to thank Chris Roome for IT-support.

## Supplements

Table S1 Isolation times and location of hosts and viruses.

Table S2 Sequence-based annotation for all predicted proteins from all four ChlorV strains

Table S3 ChlorV gene clusters and sequence/structure-based selected annotations.

Table S4 Summary of ipTM and pTM values for all structural prediction of dimeric or monomeric methyltransferases and different DNA sequences.

Table S5 Experimentally detected proteins using mass spectrometry.

Table S6 Sequencing read accessions and basecalling models.

Table S7 Chlorarachniophyte hosts 18S and 28S gene accessions.

Supplementary File 1: Single gene trees for giant virus phylogenetic markers.

Supplementary File 2: Protein structure predictions using AlphaFold 3 and structure-based annotation.

Datashare: Trees and alignments, AlphaFold 3 models, tabular methylation calls, defense finder result files.

---

**Figure S1:**
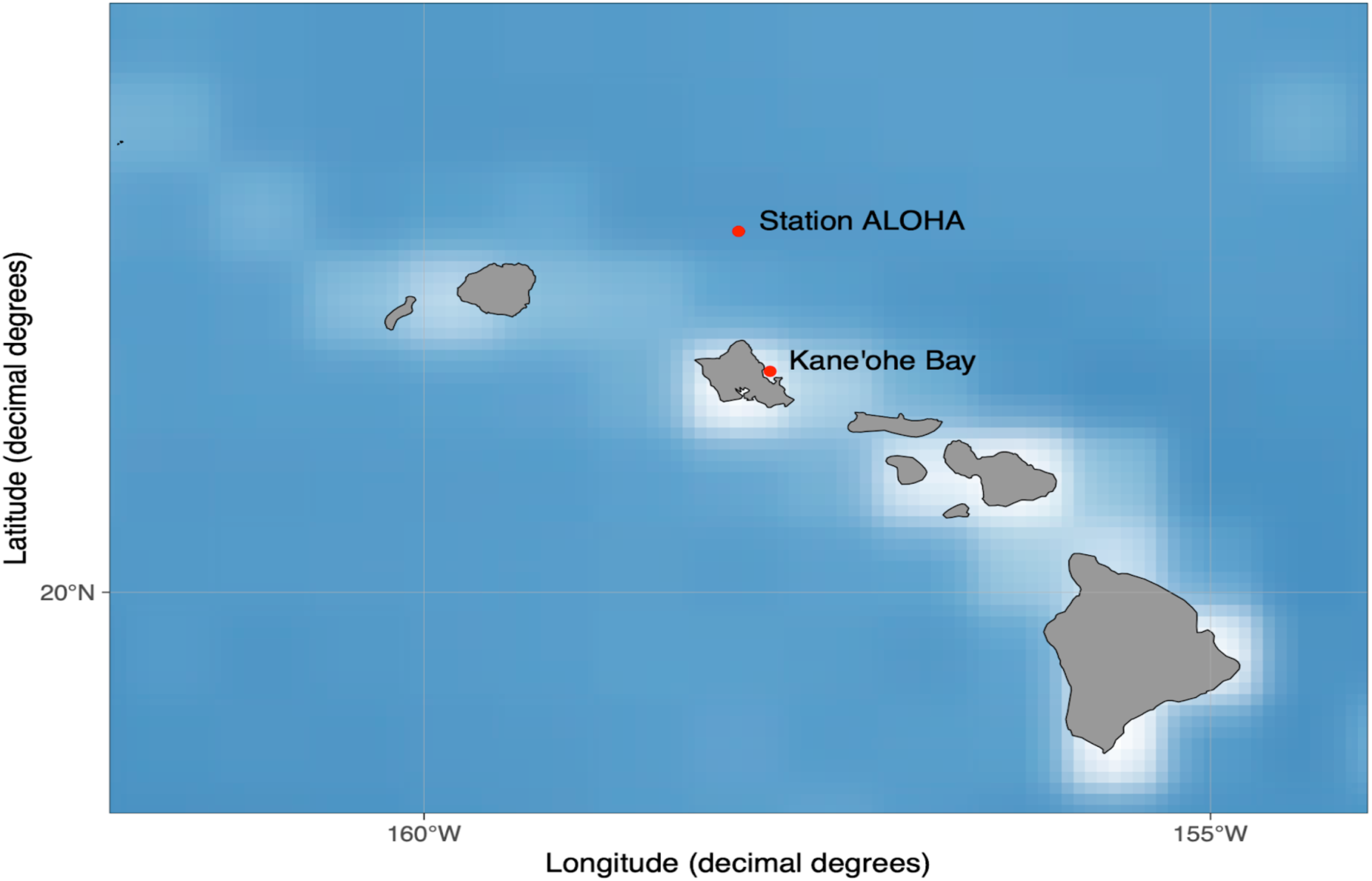
Location of the open-ocean collection point Station ALOHA approximately 100 km north of O‘ahu.

**Figure S2:**
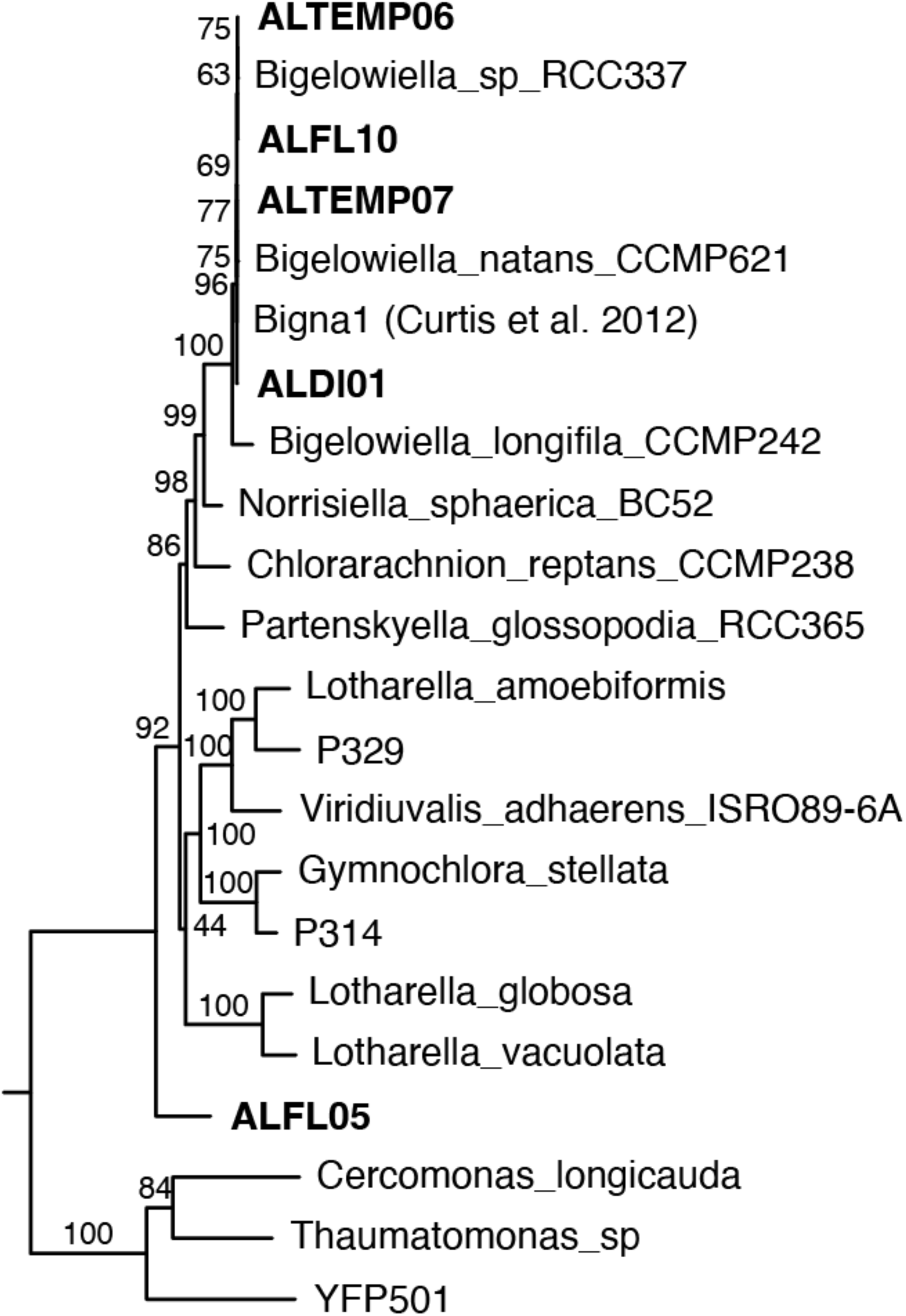
Phylogenetic tree of chlorarachniophyte host strains and references. 18S and 28S rRNA gene sequence alignments concatenated and used to reconstruct phylogenetic tree under the TIM3+F+R3 model (selected by automatic model selection) in IQ-TREE with 1000 ultrafast bootstraps.

**Figure S3:**
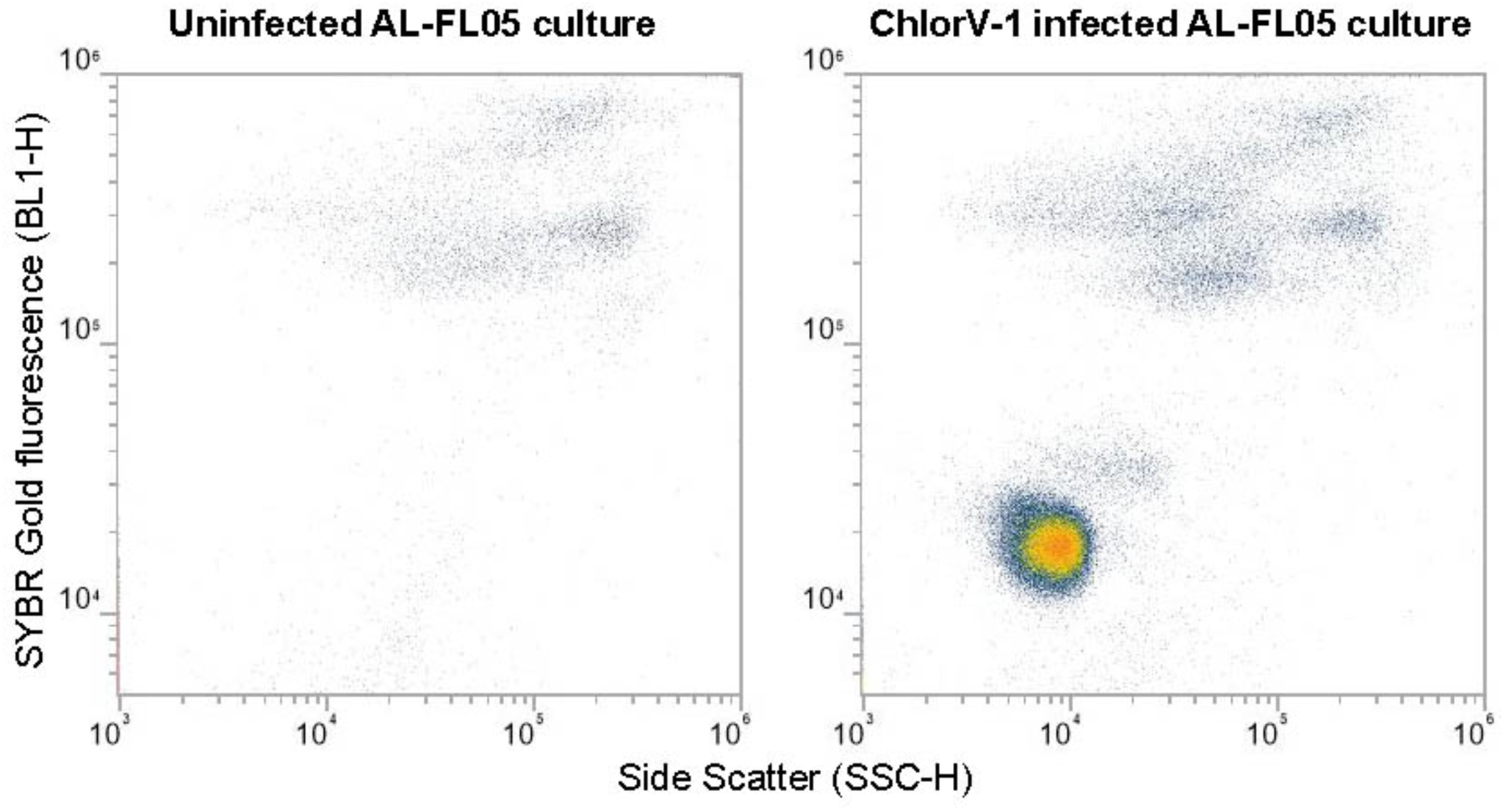
Flow cytometric detection of ChlorV-1 particles. Viral particles are clearly detectable after glutaraldehyde fixation and staining with SYBR Gold Attune NxT™ flow cytometer equipped with a small particle SSC filter (488/10). The viral population is the circular cloud of dots with the yellow/red center; diffuse populations of dots at the top represent bacteria that are present in the cultures.

**Figure S4:**
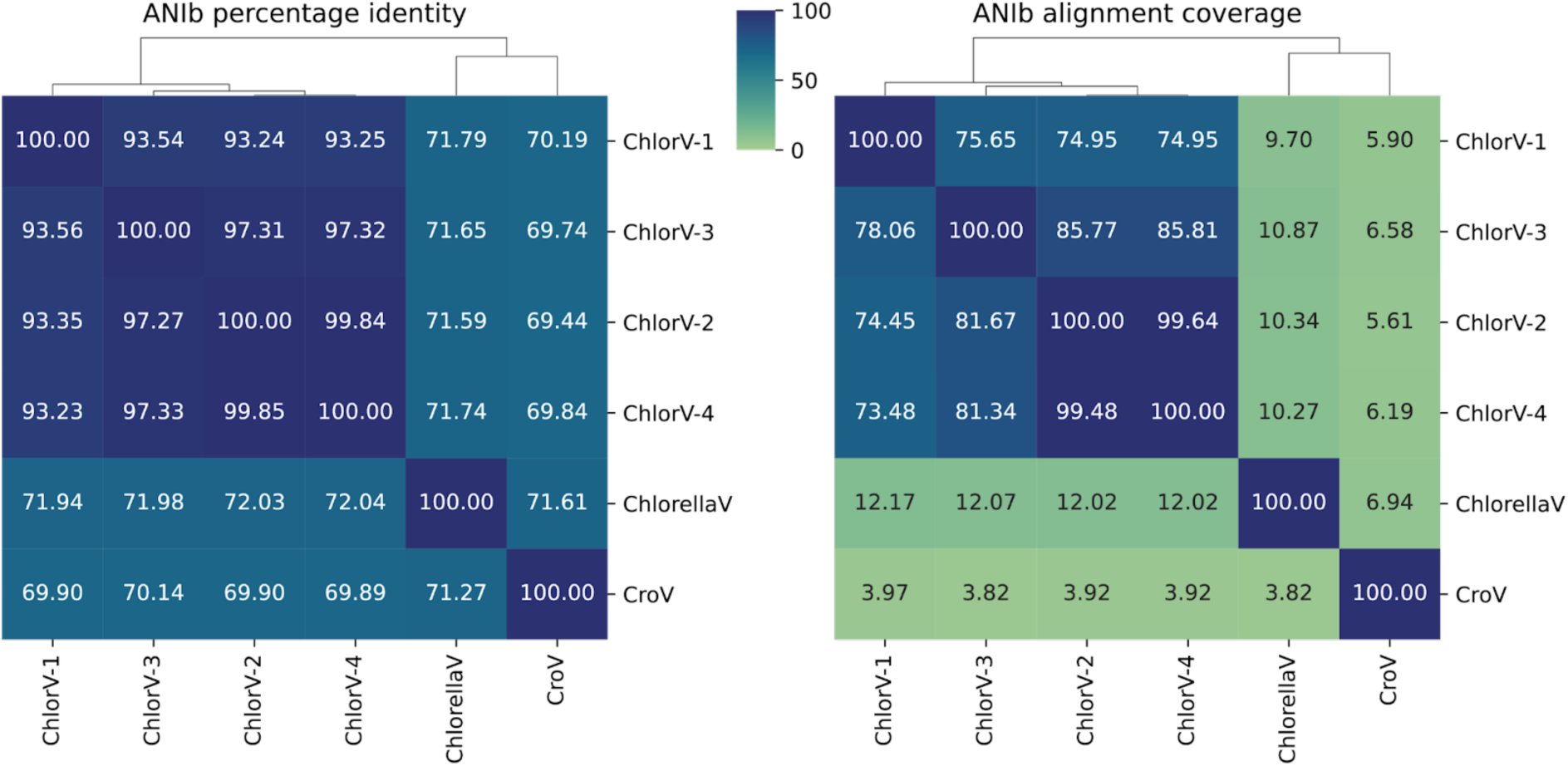
ANI and pairwise alignment coverage of ChlorVs and isolated virus genomes in *Aliimimivirinae*

**Figure S5:**
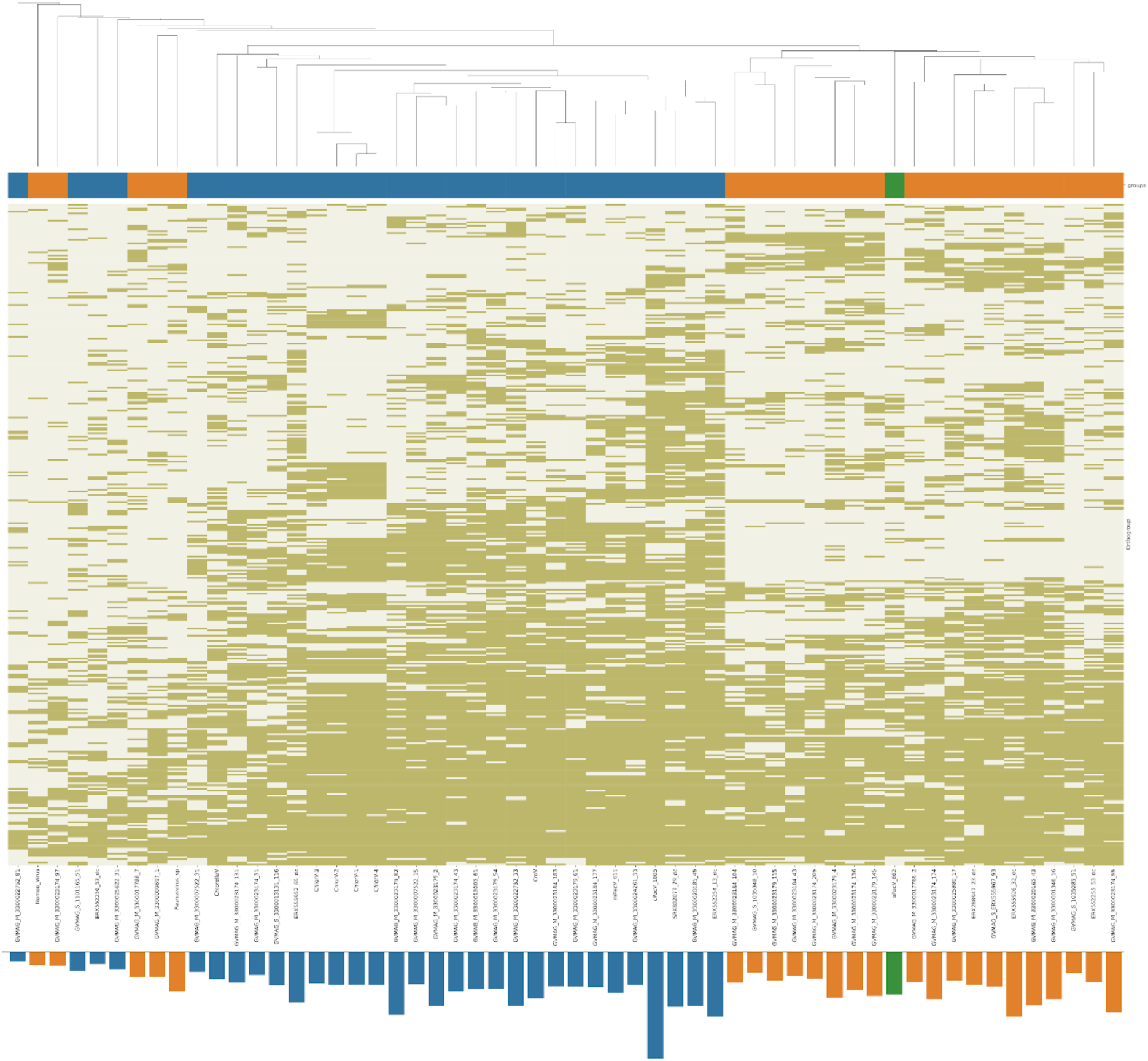
*Aliimimivirinae* gene sharing network and genome sizes. All predicted proteins of isolate and cultivation-independently acquired genomes within the subfamily *Aliimimivirinae* were subjected to orthogroup prediction. The orthogroups were then used to cluster the genomes. Genomes in the cluster *Aliimimivirinae* are labelled blue, while members of *Aliimimivirinae* II are labelled orange. The genome oPacV_662 was not confidently placed in either of these groups and thus labelled green.

**Figure S6:**
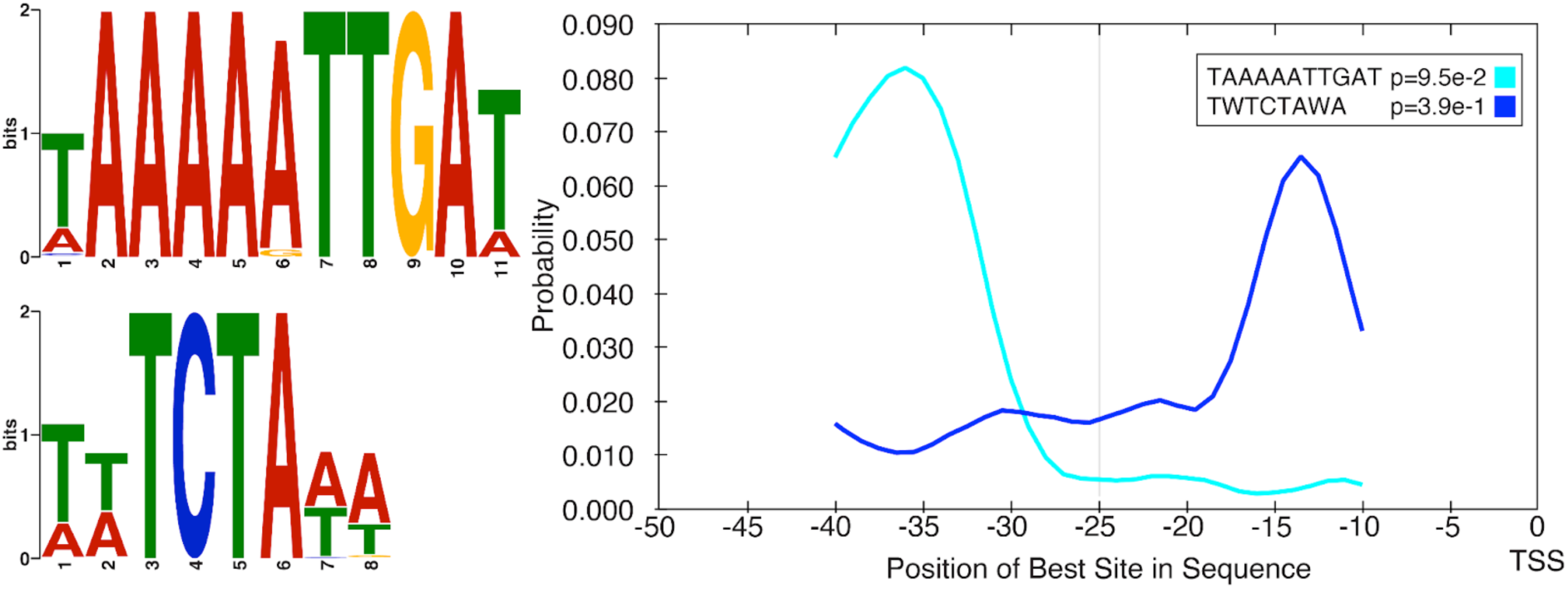
Promoter motifs in ChlorV-1. Two significant motifs were found, likely representing early (AAAAATTGA) and late (TCTA) promoter, similar to the related Cafeteria roenbergensis virus. The position relative to the transcription start site (TSS) differed between these two motifs, with AAAAATTGA showing a peak around -34 and TCTA around-14

**Figure S7:**
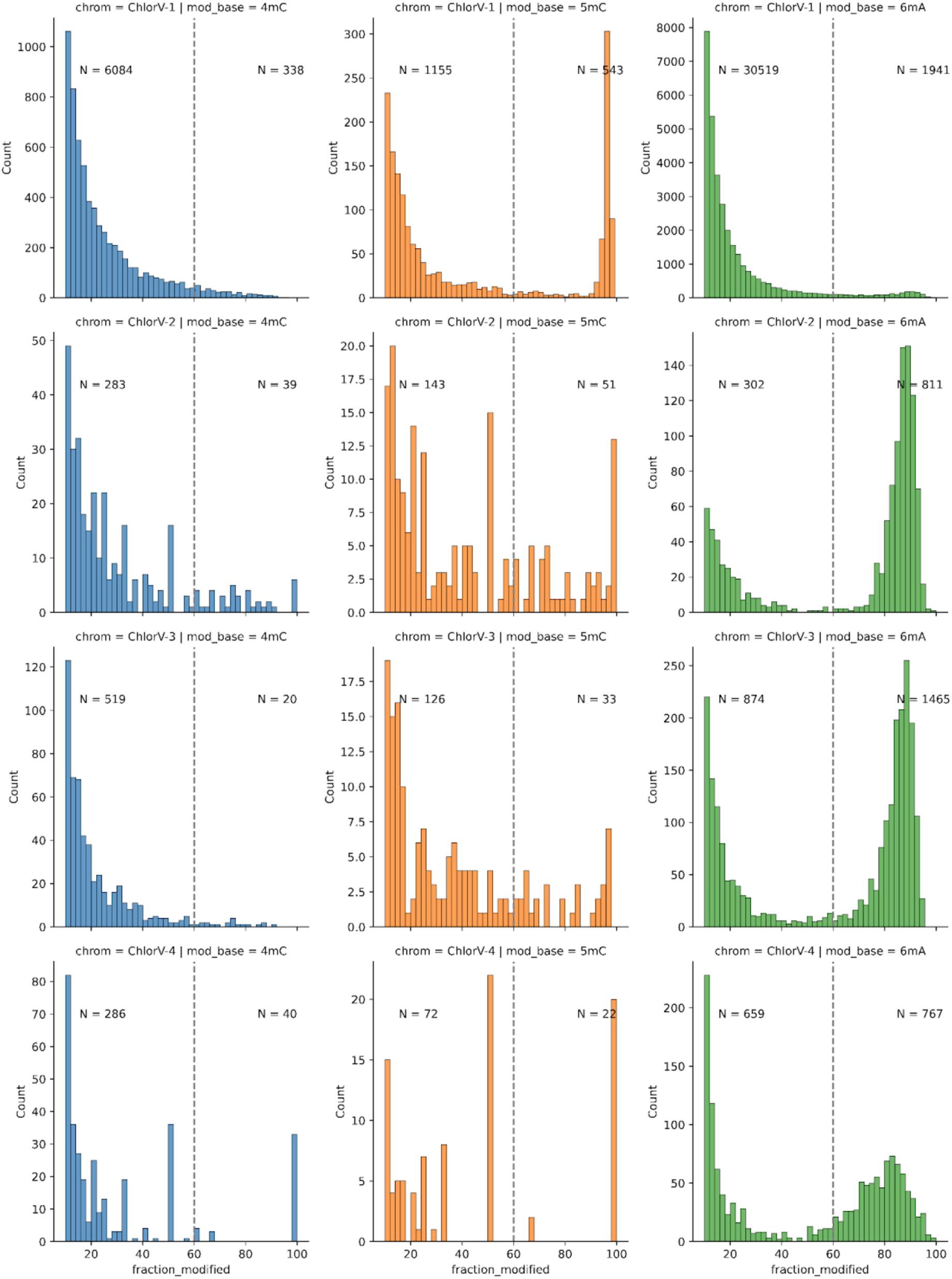
Modification frequency in ChlorVs for N4-methylcytosine (4mC), C5-methylcytosine (5mC) and N6-methyladenine (6mA). The ‘fraction modified’ is the ratio of reads corroborating a specific modification per site and the total coverage. For ChlorV-2,3 and 4 slightly different models (dna_r10.4.1_e8.2_400bps_sup@v5.0.0_4mC_5mC@v3 for 4mC and 5mC methylations and dna_r10.4.1_e8.2_400bps_sup@v5.0.0_6mA@v3 for 6mA) were used to basecall the data than for ChlorV-1 (rerio models res_dna_r10.4.1_e8.2_400bps_sup@v4.0.1_6mA@v2 and res_dna_r10.4.1_e8.2_400bps_sup@v4.3.0_4mC_5mC@v1) due to delays between sequencing runs Depth of coverage for ChlorV-1 was 2328, while ChlorV-2 had an average depth of 434, ChlorV-3 983 and ChlorV-4 81.

**Figure S8:**
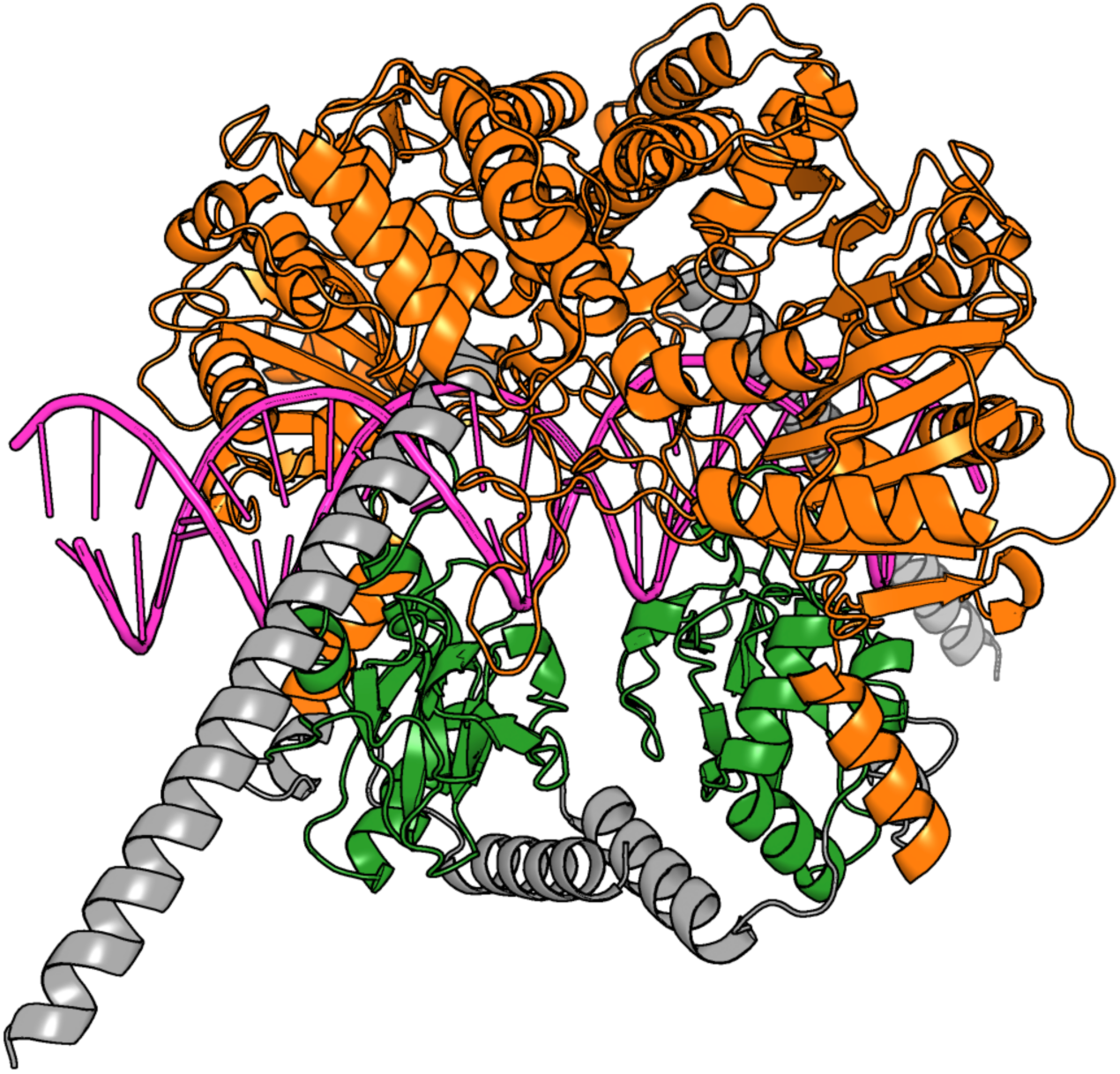
AlphaFold 3 prediction of a dimer of ChlorV-1..374 with the DNA sequence ATTAACAT**CAATTGTATG**AAAATT. Protein domains colored in Figure 4. pTM=0.85, ipTM=0.8.

**Figure S9:**
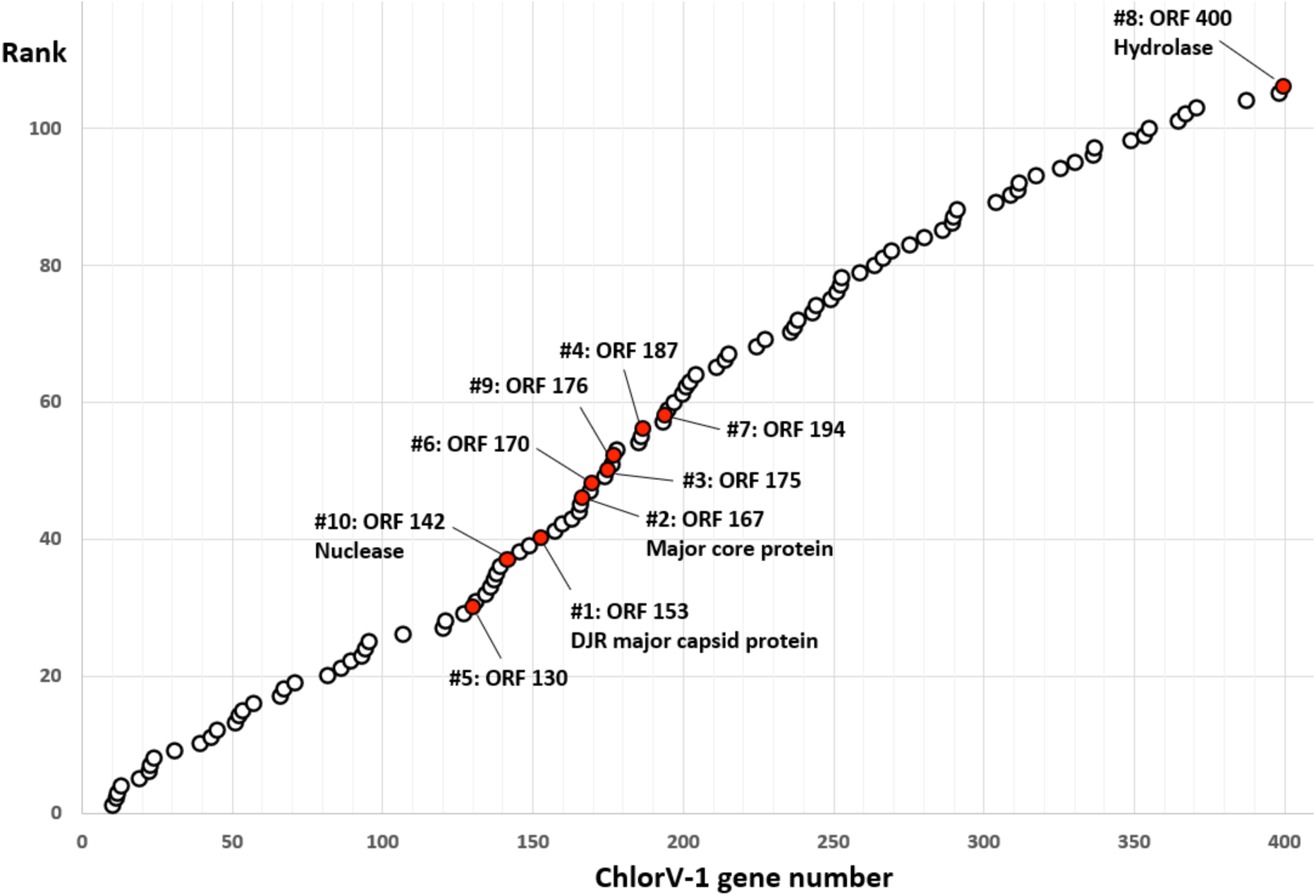
Genome position plot of ChlorV-1 virion proteins. Genes encoding proteins that detected in purified virions by spectrometry shown circles in ascending order of their respective gene number. The 10 most abundant virion proteins are marked in red and labeled with their gene number.

**Figure S10:**
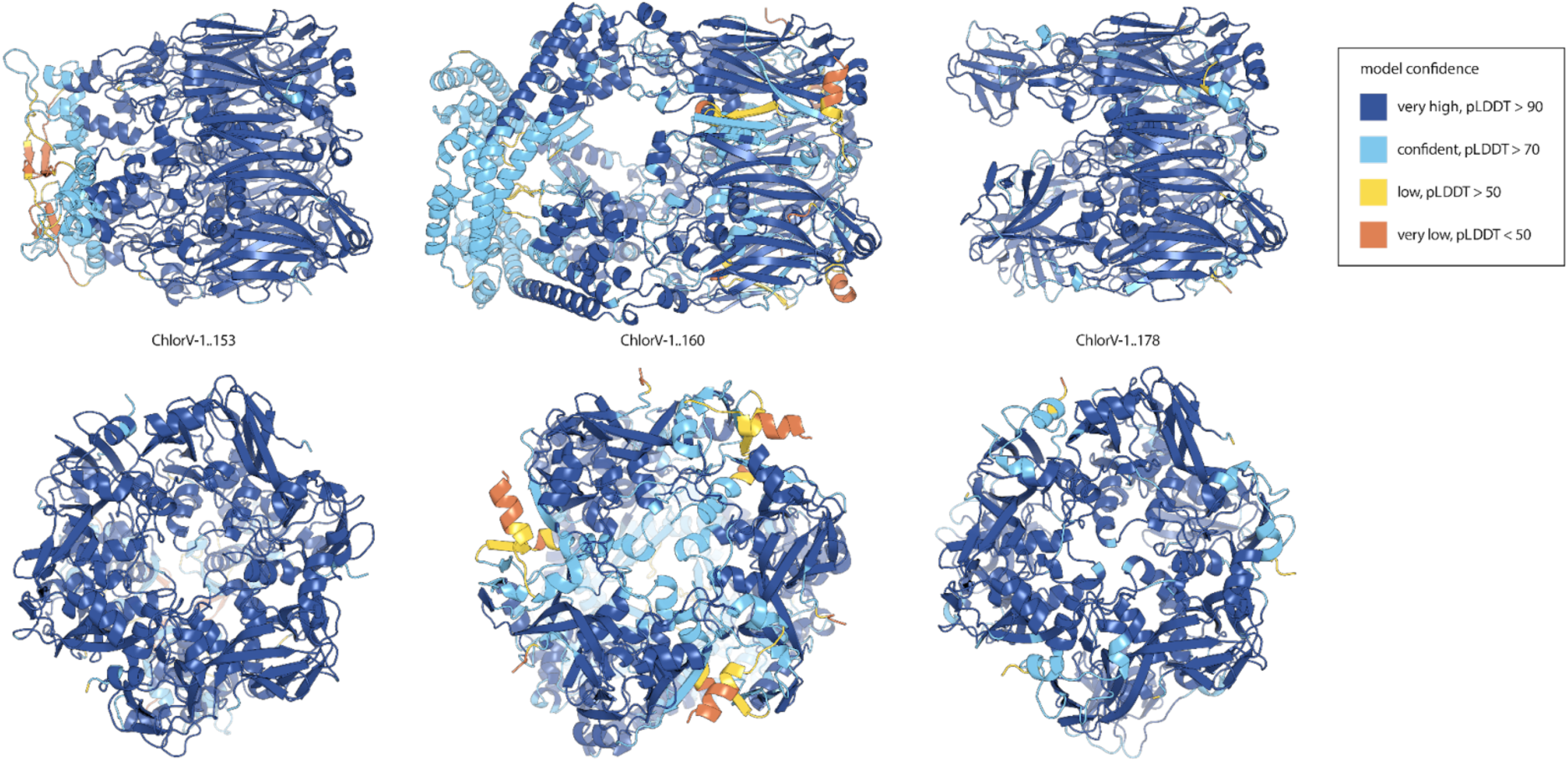
Trimeric structure predictions of the putative ChlorV- DJR capsid proteins 153, 160, and 178 with AlphaFold 3. All proteins are experimentally verified virion components.

**Figure S11:**
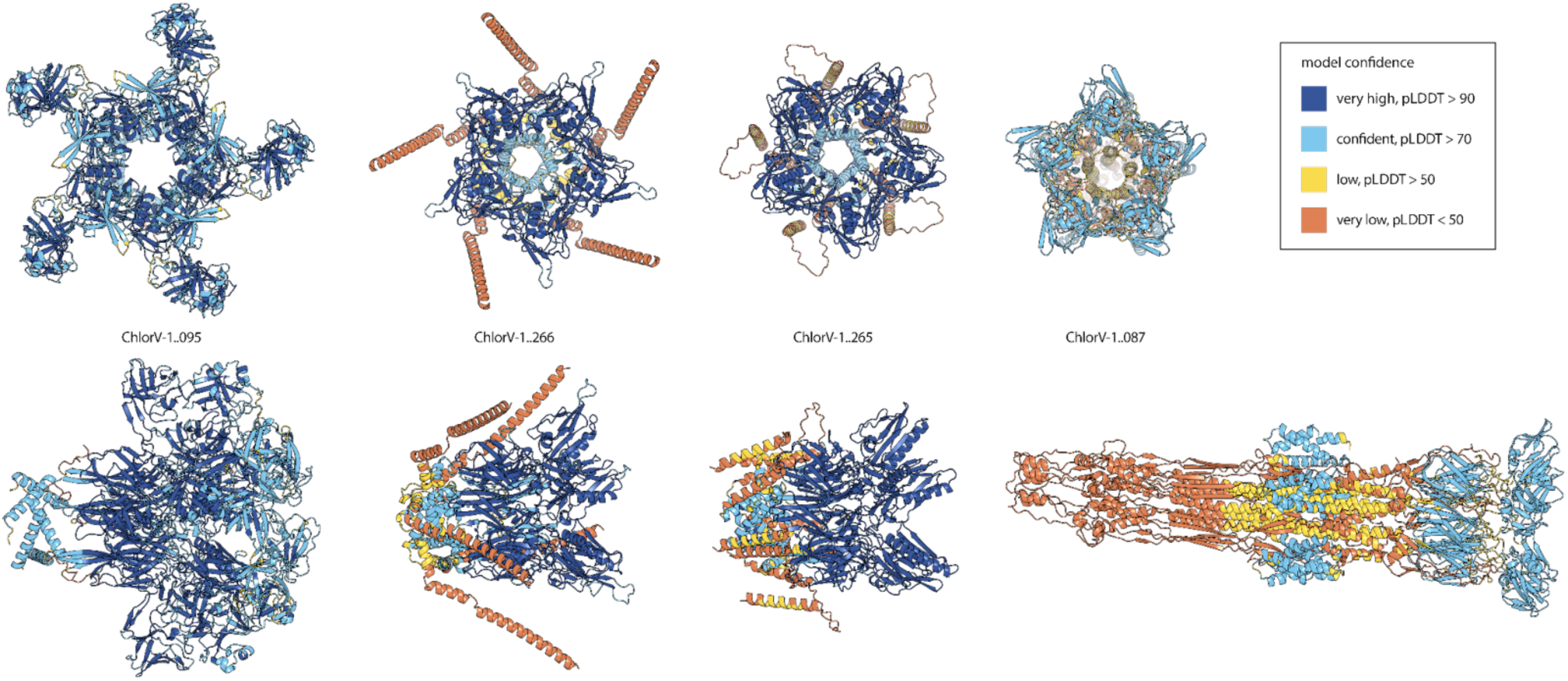
Pentameric structure predictions of the putative ChlorV-1 penton proteins 087, 095, 265 and 266 with AlphaFold 3 ChlorV-1..087 and ChlorV-1..266 experimentally verified virion proteins.

