## supplementary_material for "Strain-level diversity of giant viruses infecting chlorarachniophyte algae in the subtropical North Pacific": Supplementary_File2_ChlorV_structural_alignments.pdf

### chlorv-1..001

- Sequence-based annotation for chlorv-1..001 is putative cysteine protease
- Best hit was 2jjq chain A: UNCHARACTERIZED RNA METHYLTRANSFERASE PYRAB10780

| target | prob | fident | alnlen | evalue | theader |
|---|---|---|---|---|---|
| 2jjq-assembly1.cif.gz_A | 1 | 0.158 | 221 | 6.58e-10 | The crystal structure of Pyrococcus abyssi tRNA (uracil-54, C5)- methyltransferase in complex with S-adenosyl-L-homocysteine |
| 5zq8-assembly1.cif.gz_B | 1 | 0.162 | 216 | 8.932e-10 | Crystal structure of spRlmCD with U747 stemloop RNA |
| 1uwv-assembly1.cif.gz_A | 1 | 0.134 | 208 | 1.009e-09 | Crystal Structure of RumA, the iron-sulfur cluster containing E. coli 23S Ribosomal RNA 5-Methyluridine Methyltransferase |

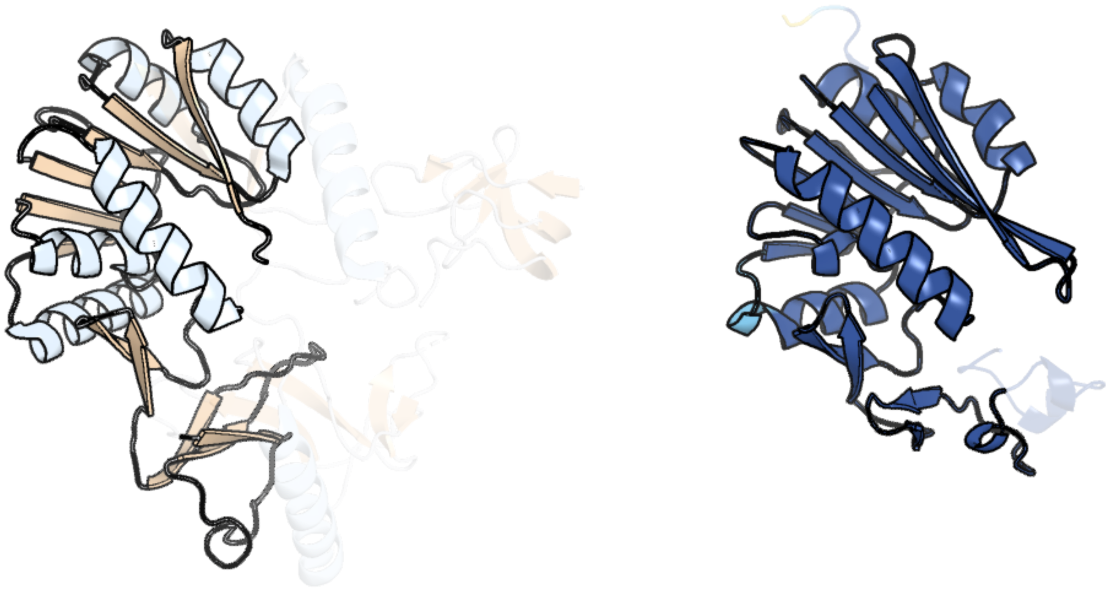

Figure 1: left: reference structure of 2jjq chain A. right: predicted structure of chlorv-1..001, unaligned sequences are shown as transparent

#### chlorv-1..002

- Sequence-based annotation for chlorv-1..002 is putative Zinc finger
- No significant structural hit found

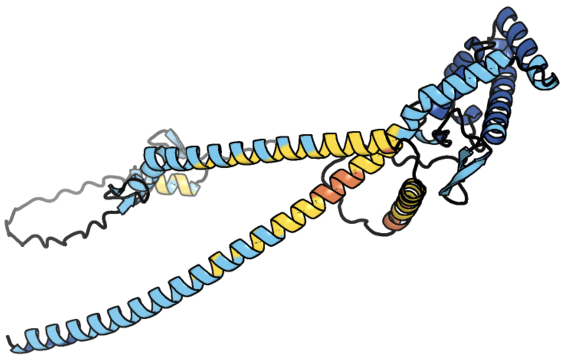

Figure 2: predicted structure of chlorv-1..002

#### chlorv-1..003

- Sequence-based annotation for chlorv-1..003 is putative Methyltransferase
- Best hit was 2py6 chain A: Methyltransferase FkbM

| target | prob | fident | alnlen | evalue | theader |
|---|---|---|---|---|---|
| 2py6-assembly1.cif.gz_A | 1 | 0.122 | 221 | 4.785e-06 | Crystal structure of Methyltransferase FkbM (YP_546752.1) from Methylobacillus flagellatus KT at 2.20 A resolution |
| 6rxx-assembly1.cif.gz_CB | 1 | 0.091 | 218 | 0.0001177 | Cryo-EM structure of the 90S pre-ribosome (Kre33-Noc4) from Chaetomium thermophilum, state C, Poly-Ala |
| 3e05-assembly2.cif.gz_C | 1 | 0.109 | 219 | 0.0001177 | CRYSTAL STRUCTURE OF Precorrin-6y C5,15-methyltransferase FROM Geobacter metallireducens GS-15 |

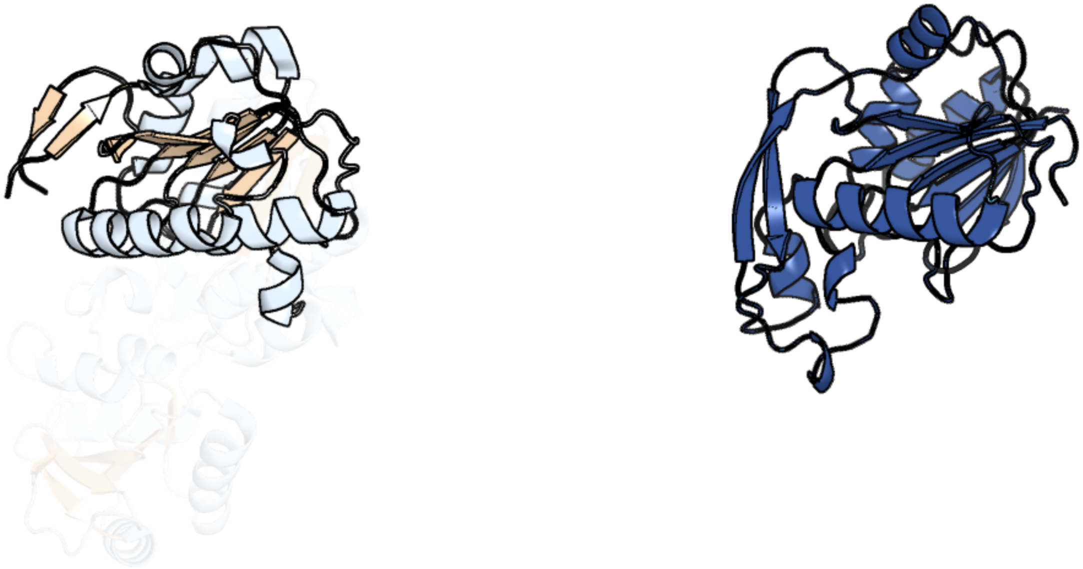

Figure 3: left: reference structure of 2py6 chain A. right: predicted structure of chlorv-1..003, unaligned sequences are shown as transparent

#### chlorv-1..004

- Sequence-based annotation for chlorv-1..004 is hypothetical protein
- No significant structural hit found

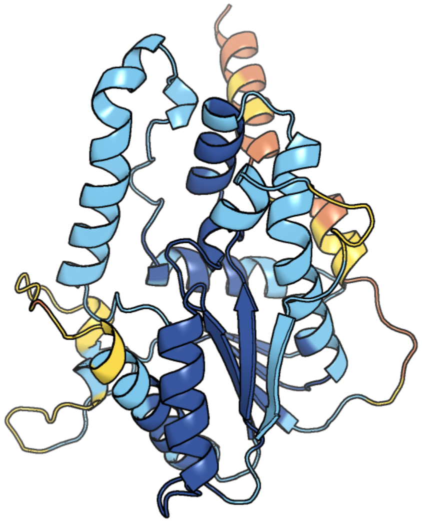

Figure 4: predicted structure of chlorv-1..004

#### chlorv-1..005

- Sequence-based annotation for chlorv-1..005 is putative Acetyltransferase
- Best hit was 1yvk chain D: hypothetical protein BSU33890

| target | prob | fident | alnlen | evalue | theader |
|---|---|---|---|---|---|
| 1yvk-assembly1.cif.gz_D | 1 | 0.114 | 149 | 3.843e-06 | Crystal Structure of the Bacillis subtilis Acetyltransferase in complex with CoA, Northeast Structural Genomics Target SR237. |
| 3owc-assembly2.cif.gz_B | 1 | 0.181 | 110 | 3.843e-06 | Crystal structure of GNAT superfamily protein PA2578 from Pseudomonas aeruginosa |
| 5ix3-assembly1.cif.gz_A | 1 | 0.238 | 113 | 6.661e-06 | Crystal structure of N-acetyltransferase from Staphylococcus aureus. |

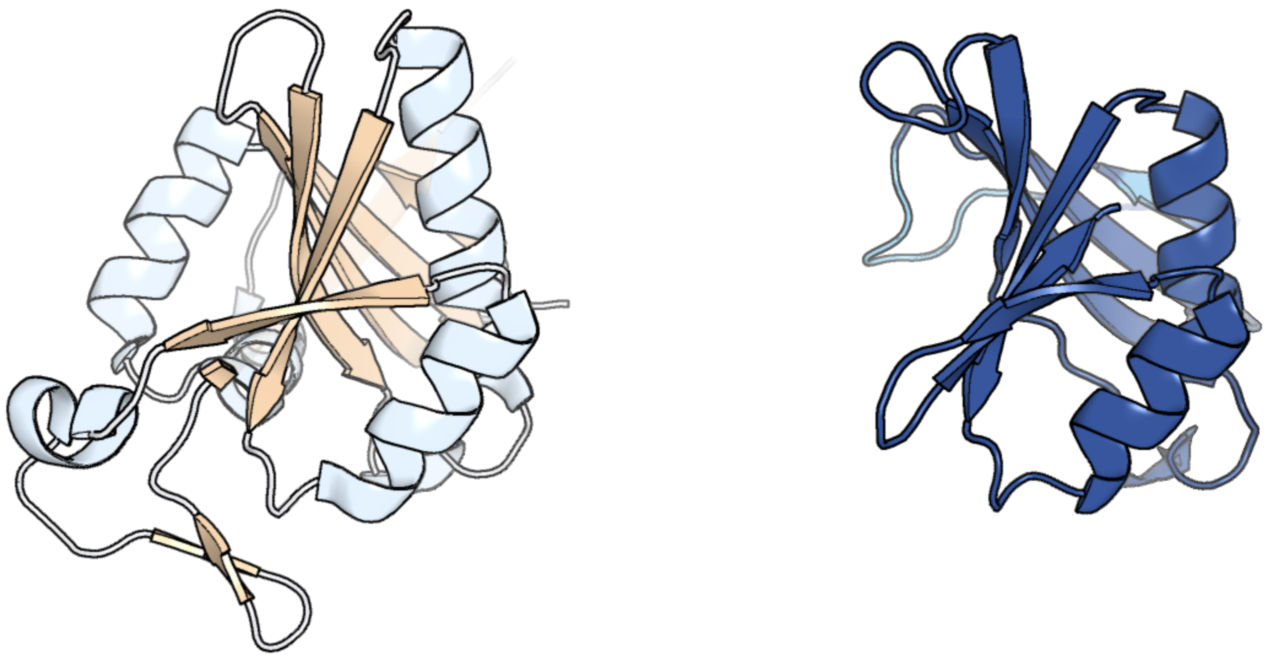

Figure 5: left: reference structure of 1yvk chain D. right: predicted structure of chlorv-1..005, unaligned sequences are shown as transparent

#### chlorv-1..006

- Sequence-based annotation for chlorv-1..006 is putative FAD/NAD(P)-binding domain
- Best hit was 6f7l chain B: Amine oxidase LkcE

| target | prob | fident | alnlen | evalue | theader |
|---|---|---|---|---|---|
| 6f7l-assembly1.cif.gz_B | 1 | 0.118 | 412 | 1.371e-17 | Crystal structure of LkcE R326Q mutant in complex with its substrate |
| 6f7l-assembly1.cif.gz_A | 1 | 0.115 | 416 | 1.449e-17 | Crystal structure of LkcE R326Q mutant in complex with its substrate |
| 6f32-assembly1.cif.gz_A | 1 | 0.113 | 423 | 2.806e-17 | Crystal structure of a dual function amine oxidase/cyclase in complex with substrate analogues |

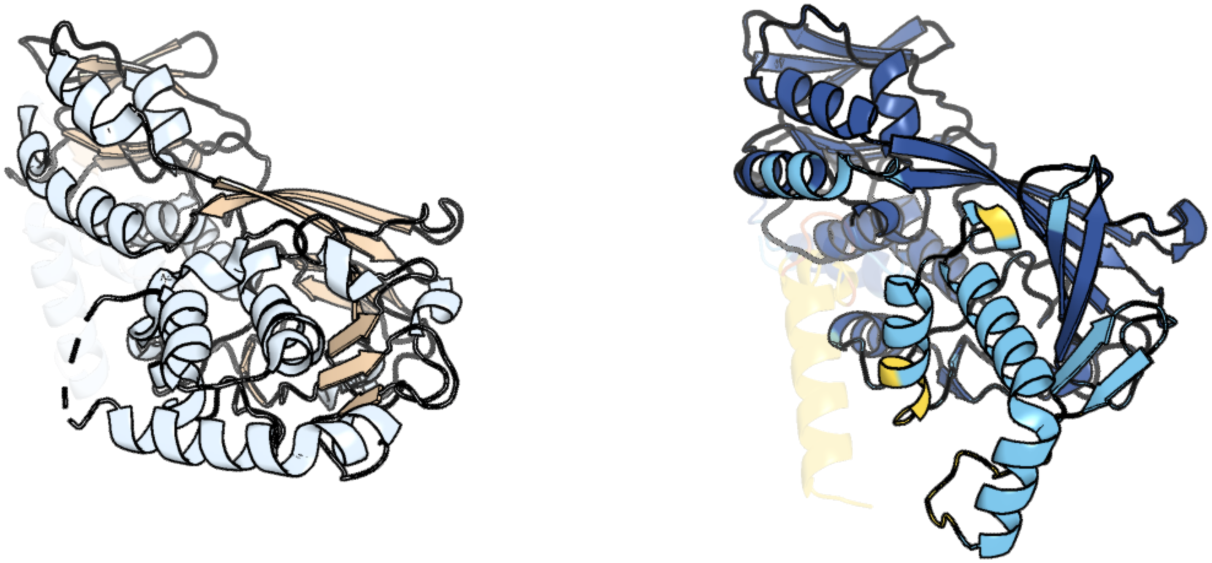

Figure 6: left: reference structure of 6f7l chain B. right: predicted structure of chlorv-1..006, unaligned sequences are shown as transparent

#### chlorv-1..007

- Sequence-based annotation for chlorv-1..007 is putative DNA topoisomerase/gyrase
- Best hit was 4gfh chain F: DNA topoisomerase 2

| target | prob | fident | alnlen | evalue | theader |
|---|---|---|---|---|---|
| 4gfh-assembly1.cif.gz_F | 1 | 0.421 | 1123 | 0 | Topoisomerase II-DNA-AMPPNP complex |
| 6zy8-assembly1.cif.gz_A | 1 | 0.422 | 1142 | 0 | Cryo-EM structure of the entire Human topoisomerase II alpha in State 2 |
| 6zy7-assembly1.cif.gz_B | 1 | 0.42 | 1139 | 0 | Cryo-EM structure of the entire Human topoisomerase II alpha in State 1 |

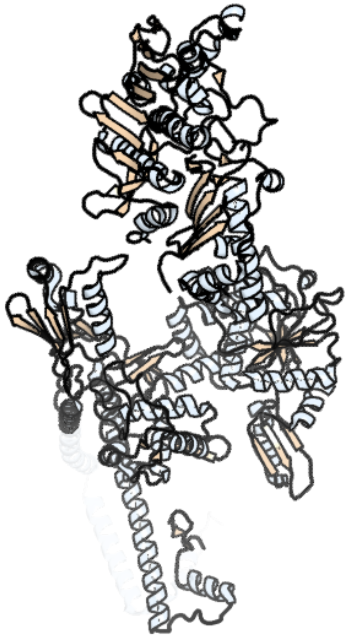

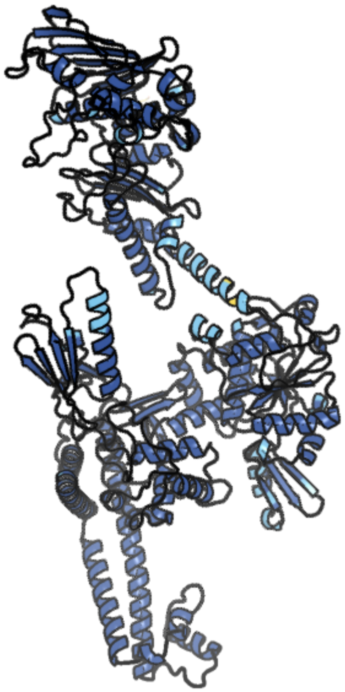

Figure 7: left: reference structure of 4gfh chain F. right: predicted structure of chlorv-1..007, unaligned sequences are shown as transparent

#### chlorv-1..008

- Sequence-based annotation for chlorv-1..008 is putative Glycosyltransferase
- Best hit was 2z87 chain A: Chondroitin synthase

| target | prob | fident | alnlen | evalue | theader |
|---|---|---|---|---|---|
| 2z87-assembly2.cif.gz_A | 1 | 0.102 | 596 | 2.551e-12 | Crystal structure of chondroitin polymerase from Escherichia coli strain K4 (K4CP) complexed with UDP-GalNAc and UDP |
| 2z86-assembly1.cif.gz_A | 1 | 0.1 | 585 | 2.692e-12 | Crystal structure of chondroitin polymerase from Escherichia coli strain K4 (K4CP) complexed with UDP-GlcUA and UDP |
| 2z86-assembly1.cif.gz_B | 1 | 0.096 | 611 | 5.423e-12 | Crystal structure of chondroitin polymerase from Escherichia coli strain K4 (K4CP) complexed with UDP-GlcUA and UDP |

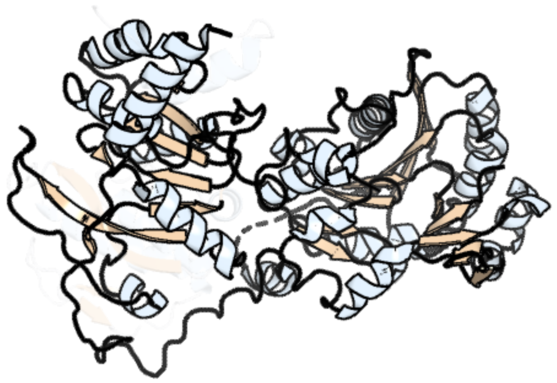

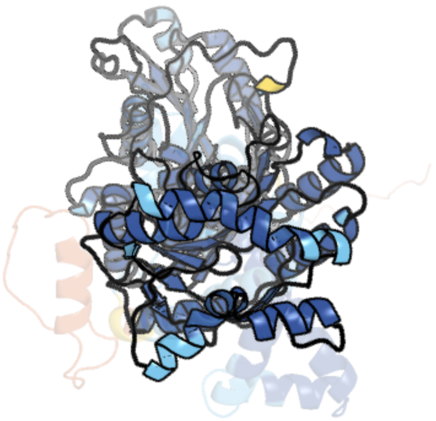

Figure 8: left: reference structure of 2z87 chain A. right: predicted structure of chlorv-1..008, unaligned sequences are shown as transparent

#### chlorv-1..009

- Sequence-based annotation for chlorv-1..009 is putative RING-finger-containing E3 ubiquitin ligase
- No significant structural hit found

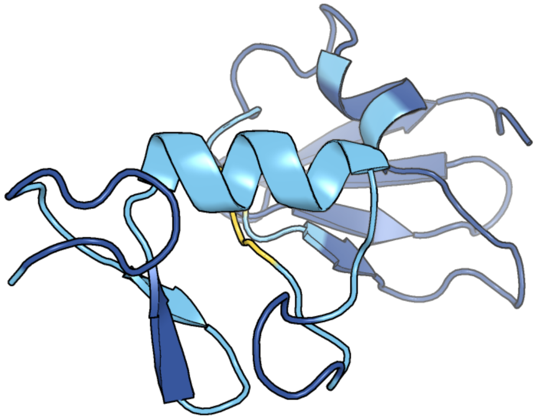

Figure 9: predicted structure of chlorv-1..009

#### chlorv-1..010

- Sequence-based annotation for chlorv-1..010 is putative Protein kinase
- Best hit was 5jzn chain A: Serine/threonine-protein kinase DCLK1

| target | prob | fident | alnlen | evalue | theader |
|---|---|---|---|---|---|
| 5jzn-assembly1.cif.gz_A | 1 | 0.166 | 265 | 1.214e-11 | Crystal structure of DCLK1-KD in complex with NVP-TAE684 |
| 5jzj-assembly1.cif.gz_A | 1 | 0.16 | 287 | 1.214e-11 | Crystal structure of DCLK1-KD in complex with AMPPN |
| 7kx6-assembly1.cif.gz_A | 1 | 0.171 | 274 | 1.284e-11 | Crystal structure of DCLK1-KD in complex with XMD8-85 |

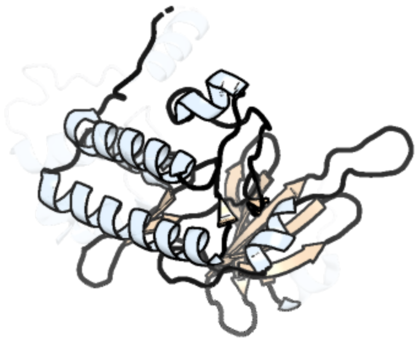

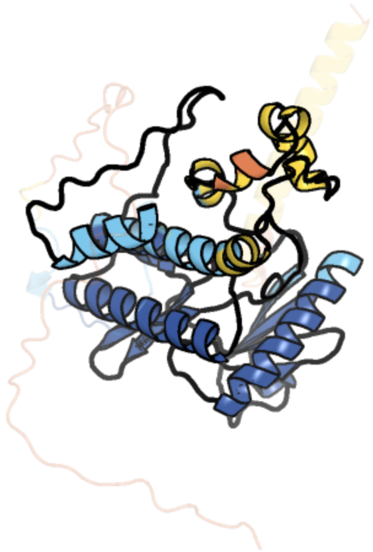

Figure 10: left: reference structure of 5jzn chain A. right: predicted structure of chlorv-1..010, unaligned sequences are shown as transparent

#### chlorv-1..011

- Sequence-based annotation for chlorv-1..011 is hypothetical protein
- No significant structural hit found

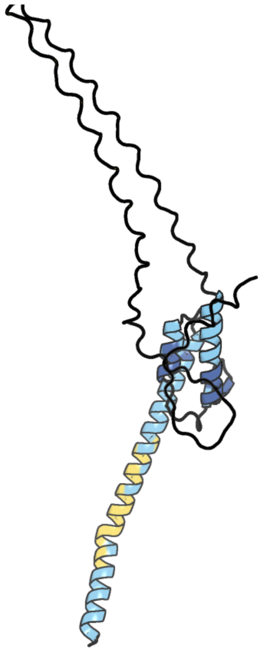

Figure 11: predicted structure of chlorv-1..011

#### chlorv-1..012

- Sequence-based annotation for chlorv-1..012 is putative Helicase
- Best hit was 5lta chain A: Pre-mRNA-splicing factor ATP-dependent RNA helicase PRP43

| target | prob | fident | alnlen | evalue | theader |
|---|---|---|---|---|---|
| 5lta-assembly1.cif.gz_A | 1 | 0.202 | 549 | 2.423e-19 | Crystal structure of the Prp43-ADP-BeF3-U7-RNA complex |
| 6zww-assembly2.cif.gz_C | 1 | 0.2 | 534 | 5.952e-19 | Crystal structure of E. coli RNA helicase HrpA in complex with RNA |
| 5vhd-assembly1.cif.gz_D | 1 | 0.169 | 620 | 1.337e-18 | DHX36 with an N-terminal truncation bound to ADP-AlF4 |

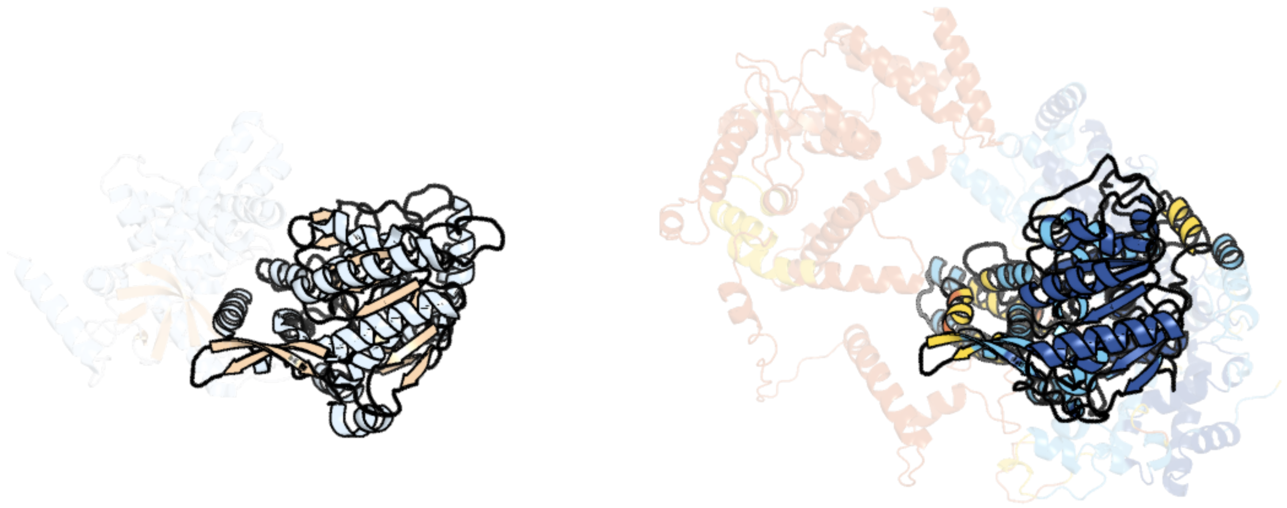

Figure 12: left: reference structure of 5lta chain A. right: predicted structure of chlorv-1..012, unaligned sequences are shown as transparent

#### chlorv-1..013

- Sequence-based annotation for chlorv-1..013 is putative Phosphoglycerate mutase
- Best hit was 2yn2 chain A: UNCHARACTERIZED PROTEIN YNL108C

| target | prob | fident | alnlen | evalue | theader |
|---|---|---|---|---|---|
| 2yn2-assembly1.cif.gz_A | 1 | 0.216 | 226 | 2.573e-11 | Huf protein - paralogue of the tau55 histidine phosphatase domain |
| 2yn0-assembly1.cif.gz_A | 1 | 0.205 | 243 | 3.114e-11 | tau55 histidine phosphatase domain |
| 4ij6-assembly1.cif.gz_B | 1 | 0.163 | 208 | 3.114e-11 | Crystal Structure of a Novel-type Phosphoserine Phosphatase Mutant (H9A) from Hydrogenobacter thermophilus TK-6 in Complex with L-phosphoserine |

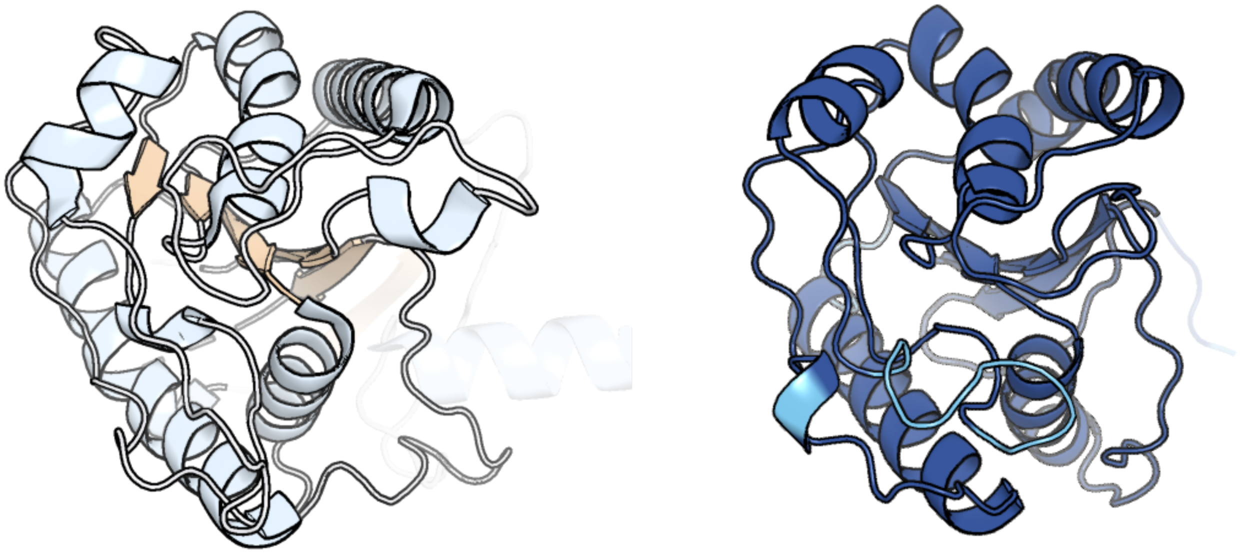

Figure 13: left: reference structure of 2yn2 chain A. right: predicted structure of chlorv-1..013, unaligned sequences are shown as transparent

#### chlorv-1..014

- Sequence-based annotation for chlorv-1..014 is hypothetical protein
- No significant structural hit found

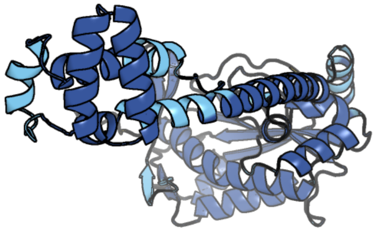

Figure 14: predicted structure of chlorv-1..014

#### chlorv-1..015

- Sequence-based annotation for chlorv-1..015 is putative DnaJ domain
- Best hit was 7zhs chain A: Ubiquitin-like protein SMT3,DnaJ homolog subfamily A member 2

| target | prob | fident | alnlen | evalue | theader |
|---|---|---|---|---|---|
| 7zhs-assembly1.cif.gz_A | 1 | 0.283 | 243 | 1.888e-24 | 3D reconstruction of the cylindrical assembly of DnaJA2 delta G/F by imposing D5 symmetry |
| 1nlt-assembly1.cif.gz_A | 1 | 0.246 | 227 | 4.557e-19 | The crystal structure of Hsp40 Ydj1 |
| 6jzb-assembly1.cif.gz_A | 1 | 0.175 | 245 | 2.865e-18 | Structural characterization of DnaJ from Streptococcus pneumonia presents a new tetramer of Hsp40 family |

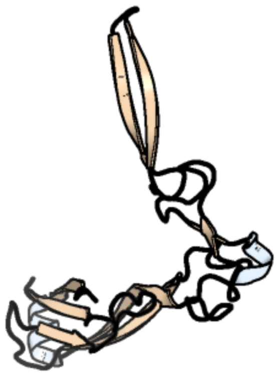

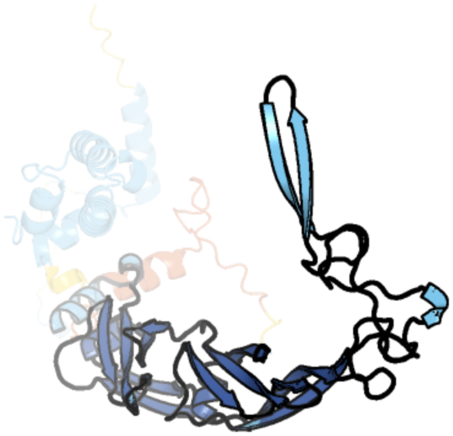

Figure 15: left: reference structure of 7zhs chain A. right: predicted structure of chlorv-1..015, unaligned sequences are shown as transparent

#### chlorv-1..016

- Sequence-based annotation for chlorv-1..016 is putative DNA clamp
- Best hit was 5yd8 chain X: Proliferating cell nuclear antigen

| target | prob | fident | alnlen | evalue | theader |
|---|---|---|---|---|---|
| 5yd8-assembly2.cif.gz_X | 1 | 0.261 | 264 | 5.365e-26 | Crystal structure of human PCNA in complex with APIM of human ZRANB3 |
| 3tbl-assembly1.cif.gz_A | 1 | 0.26 | 265 | 9.252e-26 | Structure of Mono-ubiquitinated PCNA: Implications for DNA Polymerase Switching and Okazaki Fragment Maturation |
| 6gis-assembly1.cif.gz_B | 1 | 0.261 | 264 | 1.596e-25 | Structural basis of human clamp sliding on DNA |

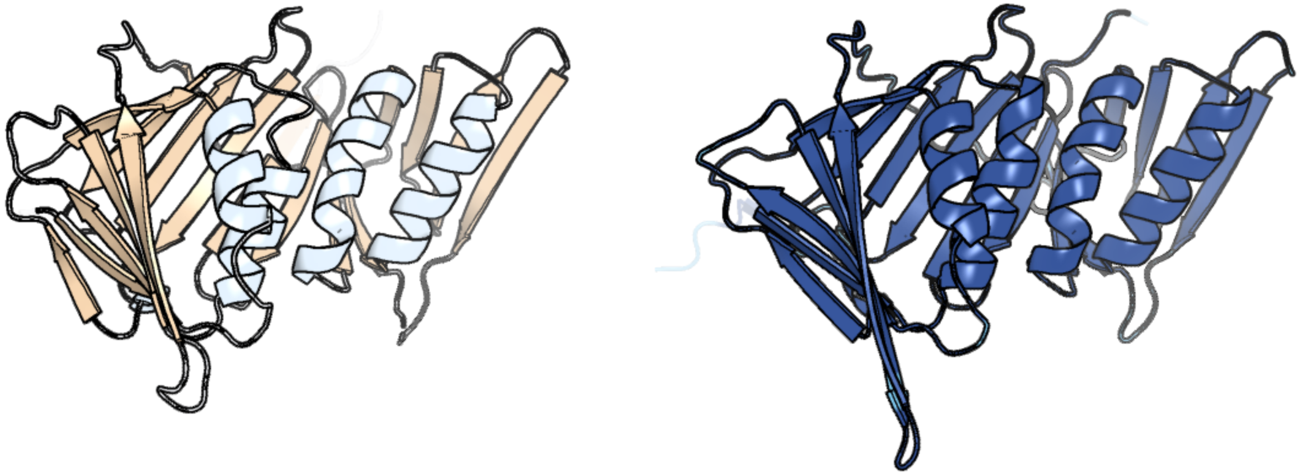

Figure 16: left: reference structure of 5yd8 chain X. right: predicted structure of chlorv-1..016, unaligned sequences are shown as transparent

#### chlorv-1..017

- Sequence-based annotation for chlorv-1..017 is putative eukaryotic translation initiation factor 4E
- Best hit was 4axg chain A: EUKARYOTIC TRANSLATION INITIATION FACTOR 4E

| target | prob | fident | alnlen | evalue | theader |
|---|---|---|---|---|---|
| 4axg-assembly1.cif.gz_A | 1 | 0.246 | 162 | 7.028e-12 | Structure of eIF4E-Cup complex |
| 4ueb-assembly1.cif.gz_C | 1 | 0.236 | 169 | 2.121e-11 | Complex of D. melanogaster eIF4E with a designed 4E-binding protein (Form II) |
| 5abv-assembly4.cif.gz_G | 1 | 0.217 | 170 | 2.884e-11 | Complex of D. melanogaster eIF4E with the 4E-binding protein Mextli |

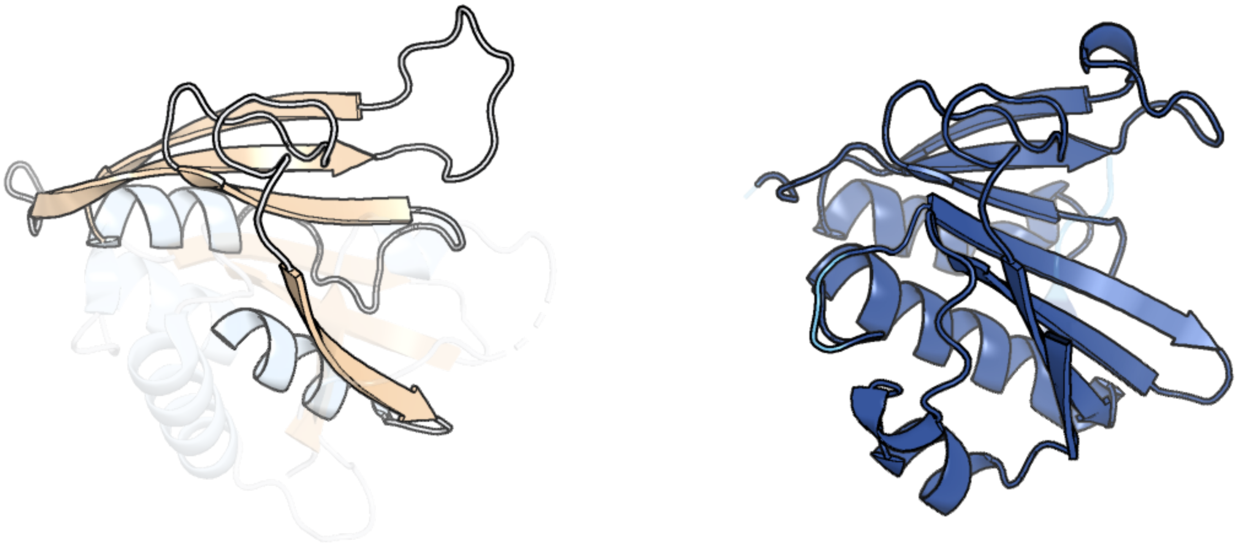

Figure 17: left: reference structure of 4axg chain A. right: predicted structure of chlorv-1..017, unaligned sequences are shown as transparent

#### chlorv-1..018

- Sequence-based annotation for chlorv-1..018 is hypothetical protein
- No significant structural hit found

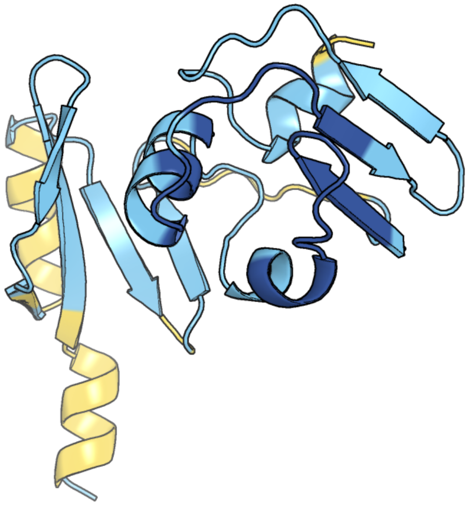

Figure 18: predicted structure of chlorv-1..018

#### chlorv-1..019

- Sequence-based annotation for chlorv-1..019 is hypothetical protein
- No significant structural hit found

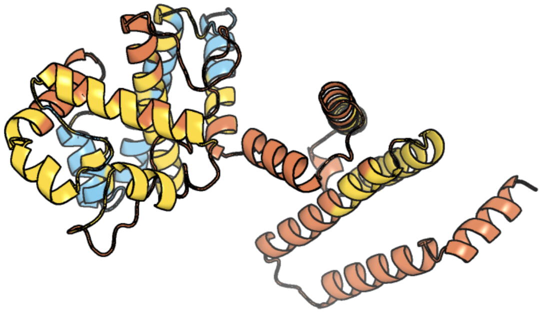

Figure 19: predicted structure of chlorv-1..019

#### chlorv-1..020

- Sequence-based annotation for chlorv-1..020 is hypothetical protein
- No significant structural hit found

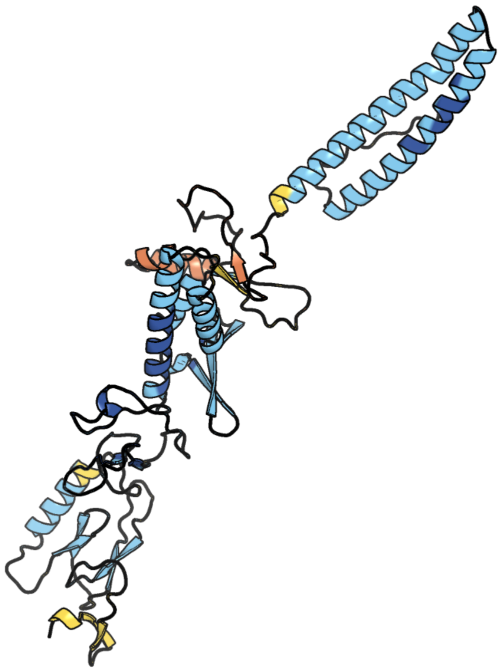

Figure 20: predicted structure of chlorv-1..020

### chlorv-1..021

- Sequence-based annotation for chlorv-1..021 is hypothetical protein
- No significant structural hit found

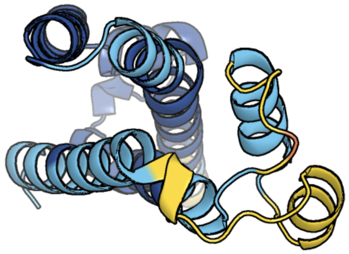

Figure 21: predicted structure of chlorv-1..021

#### chlorv-1..022

- Sequence-based annotation for chlorv-1..022 is putative Methyltransferase
- Best hit was 3gdh chain C: Trimethylguanosine synthase homolog

| target | prob | fident | alnlen | evalue | theader |
|---|---|---|---|---|---|
| 3gdh-assembly3.cif.gz_C | 1 | 0.216 | 203 | 2.148e-09 | Methyltransferase domain of human Trimethylguanosine Synthase 1 (TGS1) bound to m7GTP and adenosyl-homocysteine (active form) |
| 3gdh-assembly2.cif.gz_B | 1 | 0.221 | 203 | 3.956e-09 | Methyltransferase domain of human Trimethylguanosine Synthase 1 (TGS1) bound to m7GTP and adenosyl-homocysteine (active form) |
| 3egi-assembly1.cif.gz_A | 1 | 0.216 | 189 | 1.235e-07 | Methyltransferase domain of human trimethylguanosine synthase TGS1 bound to m7GpppA (inactive form) |

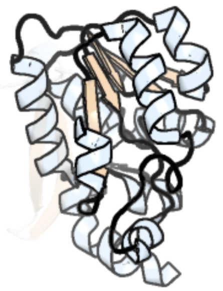

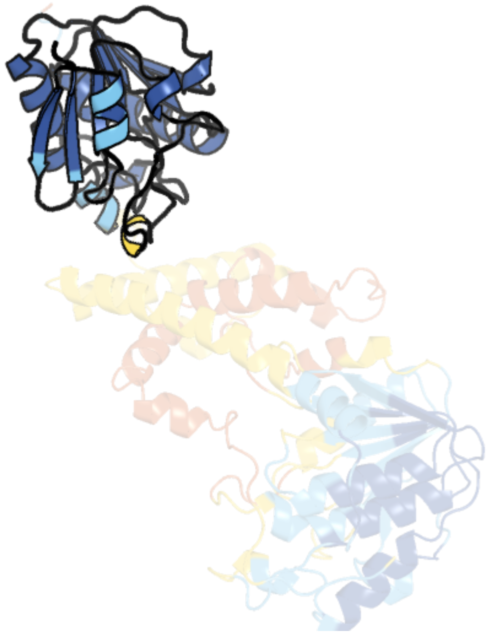

Figure 22: left: reference structure of 3gdh chain C. right: predicted structure of chlorv-1..022, unaligned sequences are shown as transparent

#### chlorv-1..023

- Sequence-based annotation for chlorv-1..023 is hypothetical protein
- No significant structural hit found

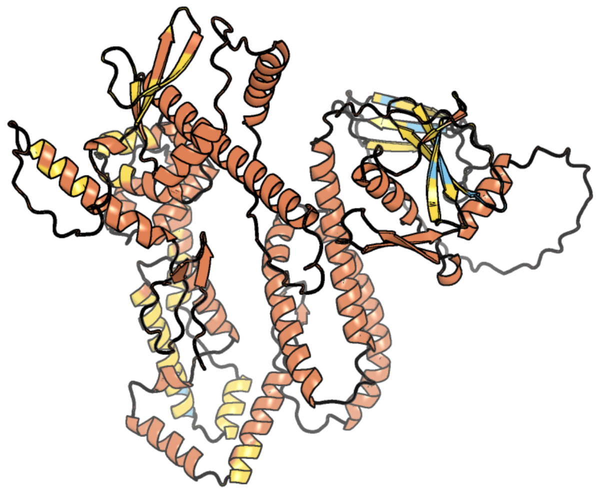

Figure 23: predicted structure of chlorv-1..023

### chlorv-1..024

- Sequence-based annotation for chlorv-1..024 is putative Helicase
- Best hit was 7amv chain W: ATP-dependent helicase VETFS

| target | prob | fident | alnlen | evalue | theader |
|---|---|---|---|---|---|
| 7amv-assembly1.cif.gz_W | 1 | 0.165 | 858 | 2.792e-22 | Atomic structure of the poxvirus transcription pre-initiation complex in the initially melted state |
| 6rfl-assembly1.cif.gz_Y | 1 | 0.163 | 702 | 3.652e-15 | Structure of the complete Vaccinia DNA-dependent RNA polymerase complex |
| 7aoh-assembly1.cif.gz_Y | 1 | 0.157 | 724 | 9.751e-15 | Atomic structure of the poxvirus late initially transcribing complex |

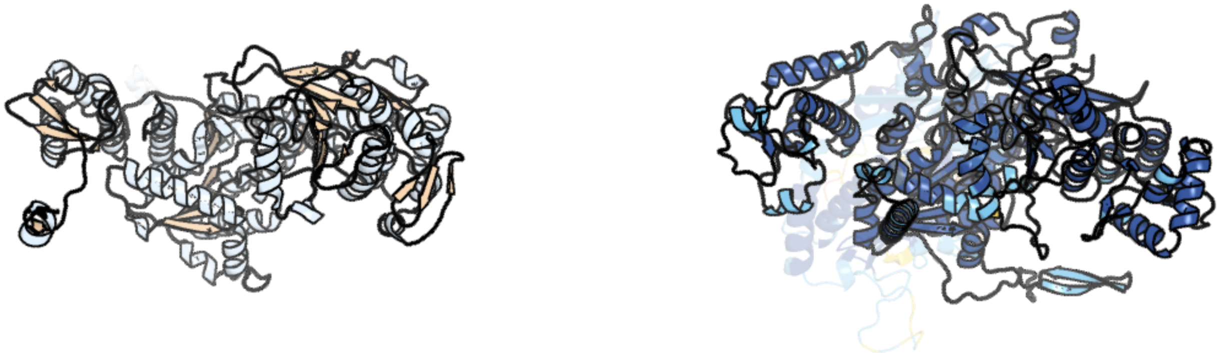

Figure 24: left: reference structure of 7amv chain W. right: predicted structure of chlorv-1..024, unaligned sequences are shown as transparent

#### chlorv-1..025

- Sequence-based annotation for chlorv-1..025 is hypothetical protein
- No significant structural hit found

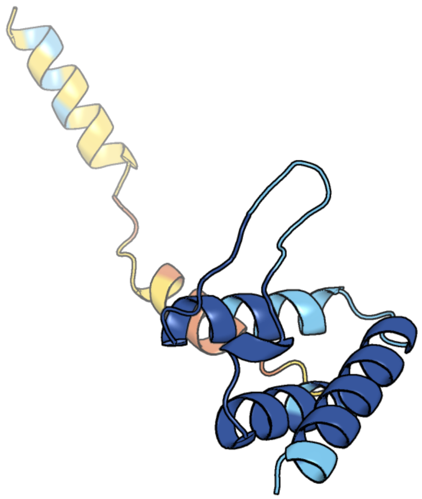

Figure 25: predicted structure of chlorv-1..025

#### chlorv-1..026

- Sequence-based annotation for chlorv-1..026 is hypothetical protein
- No significant structural hit found

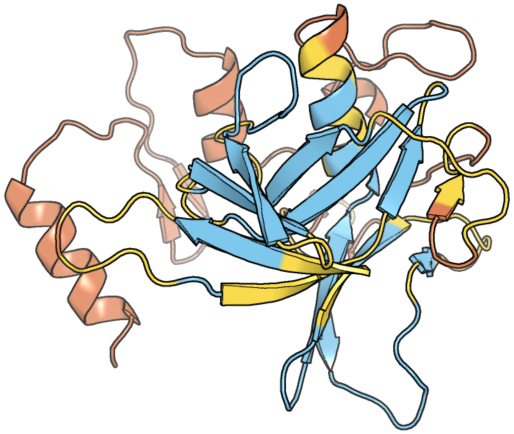

Figure 26: predicted structure of chlorv-1..026

#### chlorv-1..027

- Sequence-based annotation for chlorv-1..027 is putative POZ (Pox virus and Zinc finger) domain
- Best hit was 8h37 chain A: Kelch repeat and BTB domain-containing protein 2

| target | prob | fident | alnlen | evalue | theader |
|---|---|---|---|---|---|
| 8h37-assembly1.cif.gz_A | 1 | 0.145 | 213 | 8.206e-07 | Cryo-EM Structure of the KBTBD2-CUL3-Rbx1-p85a tetrameric complex |
| 8h37-assembly1.cif.gz_P | 1 | 0.147 | 217 | 1.543e-06 | Cryo-EM Structure of the KBTBD2-CUL3-Rbx1-p85a tetrameric complex |
| 6i2m-assembly1.cif.gz_A-2 | 1 | 0.155 | 200 | 2.178e-06 | Crystal structure of vaccinia virus protein A55 BTB-Back domain in complex with human Cullin-3 N-terminus |

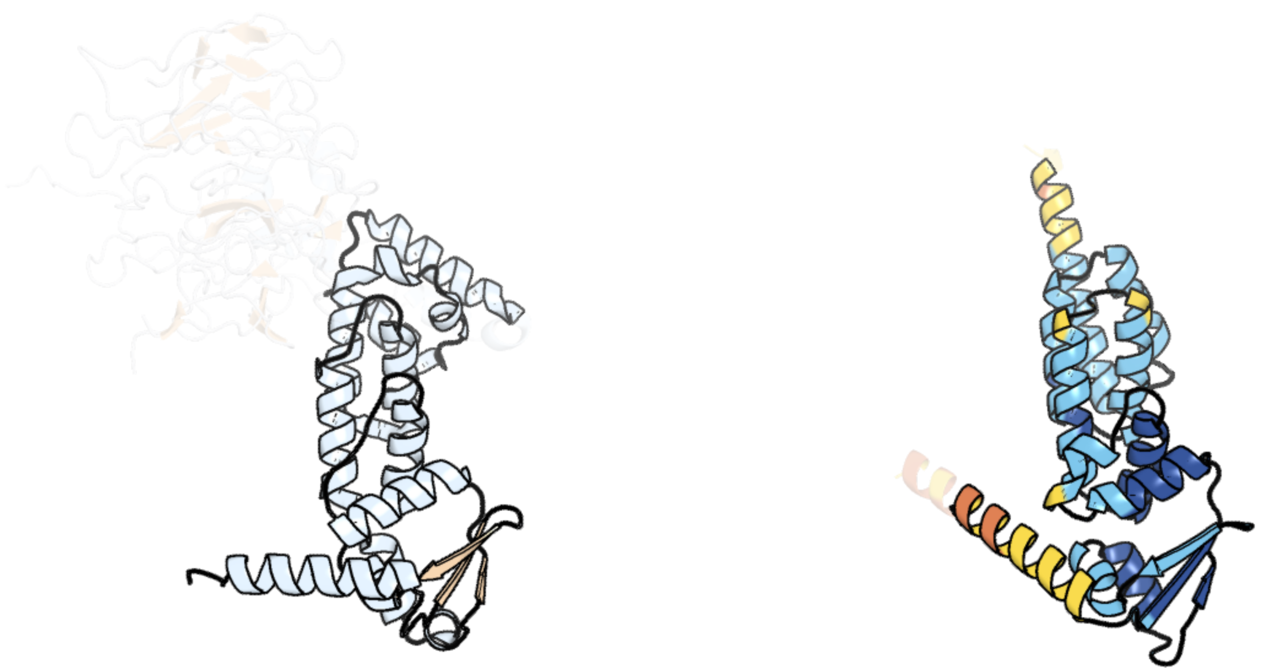

Figure 27: left: reference structure of 8h37 chain A. right: predicted structure of chlorv-1..027, unaligned sequences are shown as transparent

#### chlorv-1..028

- Sequence-based annotation for chlorv-1..028 is hypothetical protein
- No significant structural hit found

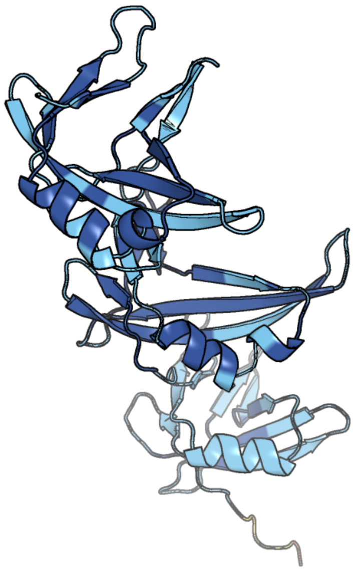

Figure 28: predicted structure of chlorv-1..028

#### chlorv-1..029

- Sequence-based annotation for chlorv-1..029 is hypothetical protein
- No significant structural hit found

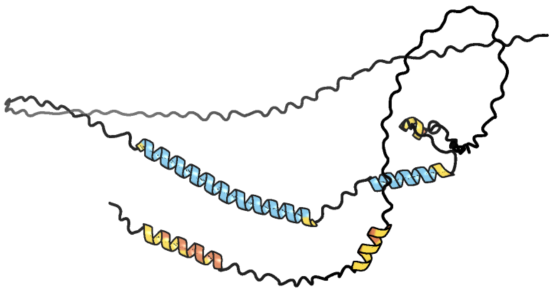

Figure 29: predicted structure of chlorv-1..029

#### chlorv-1..030

- Sequence-based annotation for chlorv-1..030 is hypothetical protein
- No significant structural hit found

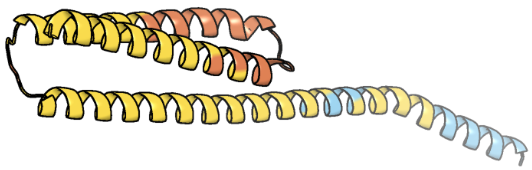

Figure 30: predicted structure of chlorv-1..030

#### chlorv-1..031

- Sequence-based annotation for chlorv-1..031 is putative DNA polymerase beta
- Best hit was 4m9h chain A: DNA polymerase beta

| target | prob | fident | alnlen | evalue | theader |
|---|---|---|---|---|---|
| 4m9h-assembly1.cif.gz_A | 1 | 0.258 | 356 | 3.476e-22 | DNA Polymerase Beta E295K Soaked with dTTP |
| 6uok-assembly2.cif.gz_A | 1 | 0.264 | 351 | 3.891e-22 | Y271G DNA polymerase beta substrate complex with templating cytosine and incoming r8-oxo-GTP |
| 4mfc-assembly1.cif.gz_A | 1 | 0.262 | 358 | 5.769e-22 | Structure of human DNA polymerase beta complexed with O6MG in the template base paired with incoming non-hydrolyzable CTP |

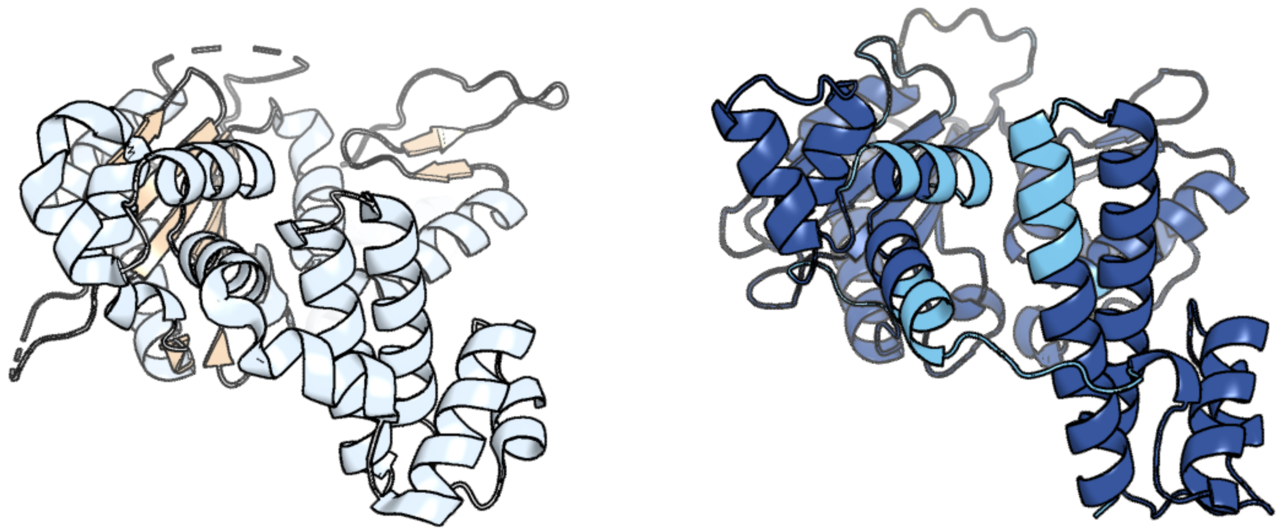

Figure 31: left: reference structure of 4m9h chain A. right: predicted structure of chlorv-1..031, unaligned sequences are shown as transparent

#### chlorv-1..032

- Sequence-based annotation for chlorv-1..032 is putative Ubiquitin Conjugating Enzyme
- Best hit was 3ptf chain A: Ubiquitin-conjugating enzyme E2 D1

| target | prob | fident | alnlen | evalue | theader |
|---|---|---|---|---|---|
| 3ptf-assembly1.cif.gz_A | 1 | 0.458 | 144 | 6.684e-19 | X-ray structure of the non-covalent complex between UbcH5A and Ubiquitin |
| 2oxq-assembly1.cif.gz_B | 1 | 0.479 | 144 | 7.104e-19 | Structure of the UbcH5 :CHIP U-box complex |
| 4wz3-assembly1.cif.gz_A | 1 | 0.465 | 144 | 7.104e-19 | Crystal structure of the complex between LubX/LegU2/Lpp2887 U-box 1 and Homo sapiens UBE2D2 |

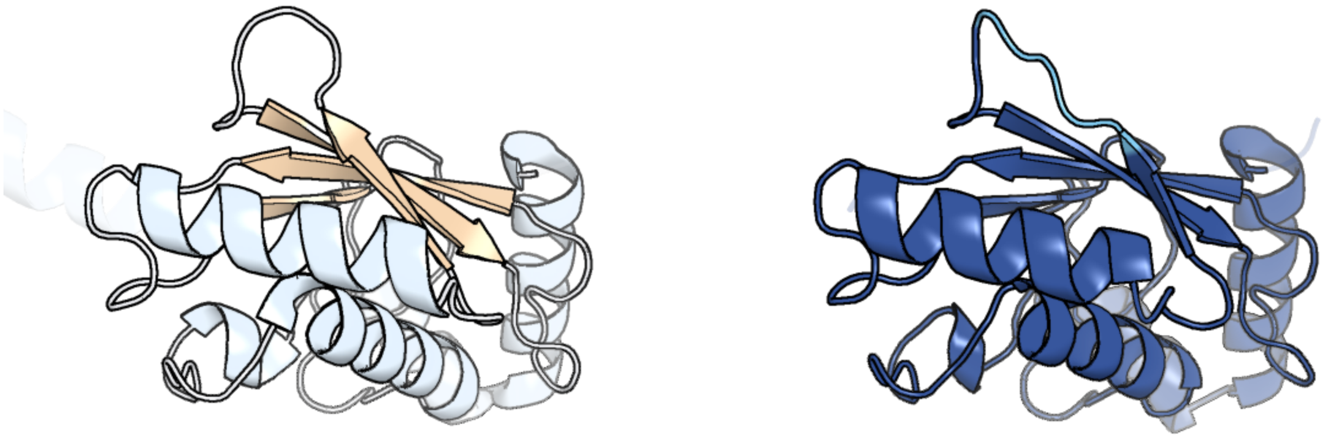

Figure 32: left: reference structure of 3ptf chain A. right: predicted structure of chlorv-1..032, unaligned sequences are shown as transparent

#### chlorv-1..033

- Sequence-based annotation for chlorv-1..033 is hypothetical protein
- No significant structural hit found

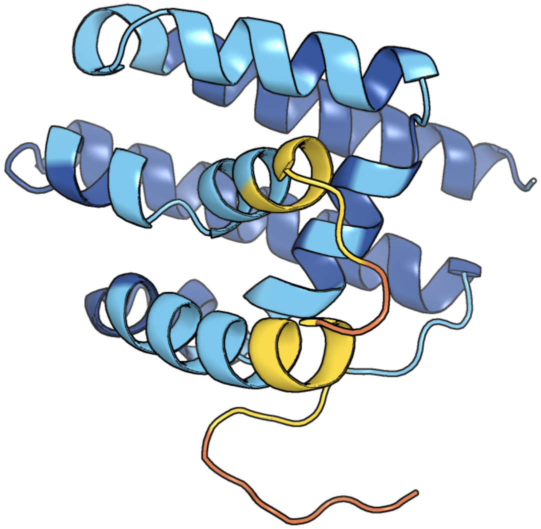

Figure 33: predicted structure of chlorv-1..033

#### chlorv-1..034

- Sequence-based annotation for chlorv-1..034 is hypothetical protein
- No significant structural hit found

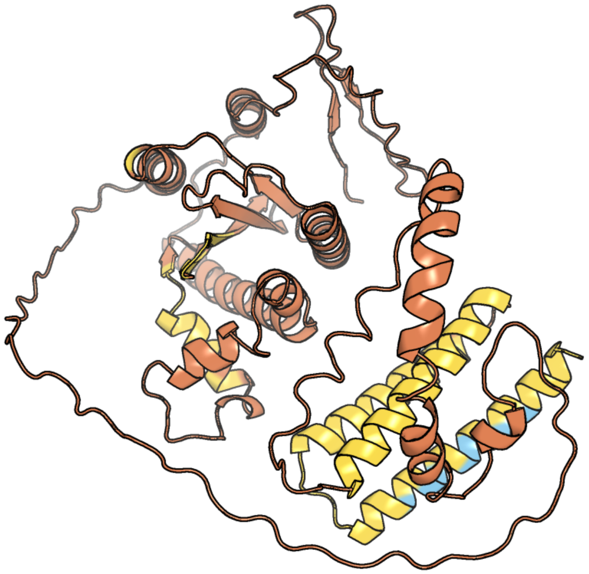

Figure 34: predicted structure of chlorv-1..034

#### chlorv-1..035

- Sequence-based annotation for chlorv-1..035 is hypothetical protein
- No significant structural hit found

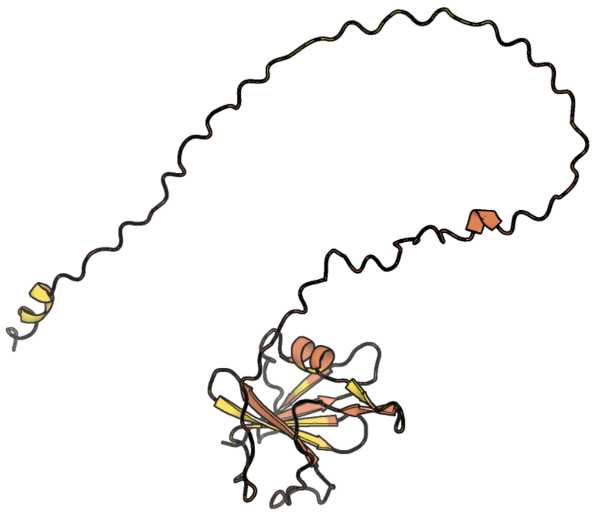

Figure 35: predicted structure of chlorv-1..035

#### chlorv-1..036

- Sequence-based annotation for chlorv-1..036 is putative Peptidase
- Best hit was 1cb5 chain A: BLEOMYCIN HYDROLASE

| target | prob | fident | alnlen | evalue | theader |
|---|---|---|---|---|---|
| 1cb5-assembly1.cif.gz_A | 1 | 0.336 | 443 | 4.469e-45 | HUMAN BLEOMYCIN HYDROLASE. |
| 2cb5-assembly1.cif.gz_A | 1 | 0.338 | 437 | 1.537e-44 | HUMAN BLEOMYCIN HYDROLASE, C73S/DELE455 MUTANT |
| 7v5l-assembly1.cif.gz_A | 1 | 0.338 | 443 | 3.373e-44 | Crystal structure of human bleomycin hydrolase |

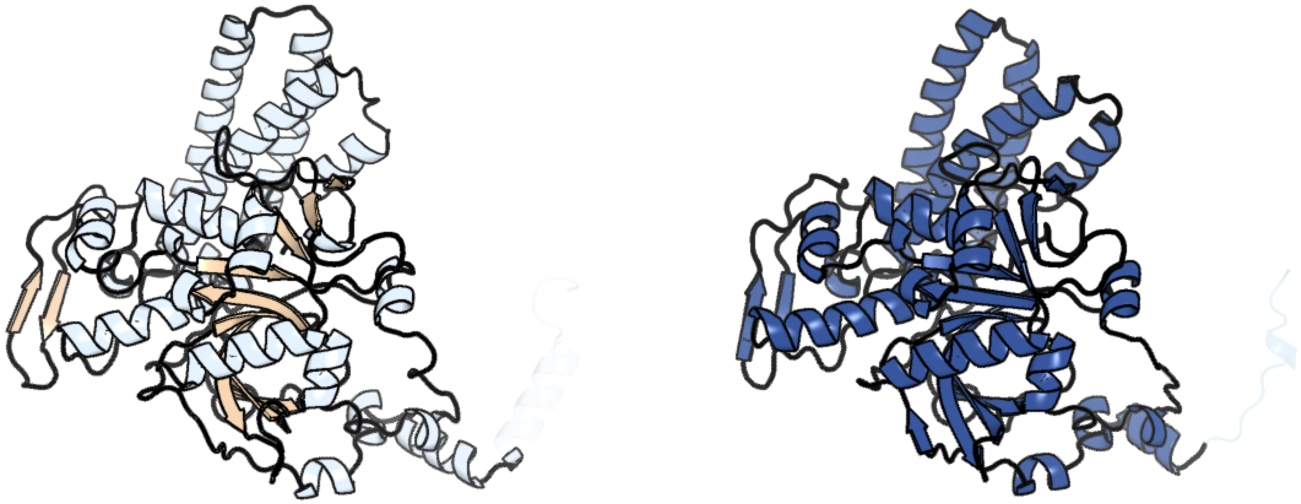

Figure 36: left: reference structure of 1cb5 chain A. right: predicted structure of chlorv-1..036, unaligned sequences are shown as transparent

#### chlorv-1..037

- Sequence-based annotation for chlorv-1..037 is putative Methyltransferase
- Best hit was 3gdh chain C: Trimethylguanosine synthase homolog

| target | prob | fident | alnlen | evalue | theader |
|---|---|---|---|---|---|
| 3gdh-assembly3.cif.gz_C | 1 | 0.212 | 207 | 6.271e-13 | Methyltransferase domain of human Trimethylguanosine Synthase 1 (TGS1) bound to m7GTP and adenosyl-homocysteine (active form) |
| 3gdh-assembly2.cif.gz_B | 1 | 0.216 | 203 | 3.528e-12 | Methyltransferase domain of human Trimethylguanosine Synthase 1 (TGS1) bound to m7GTP and adenosyl-homocysteine (active form) |
| 3egi-assembly1.cif.gz_A | 1 | 0.197 | 197 | 6.284e-10 | Methyltransferase domain of human trimethylguanosine synthase TGS1 bound to m7GpppA (inactive form) |

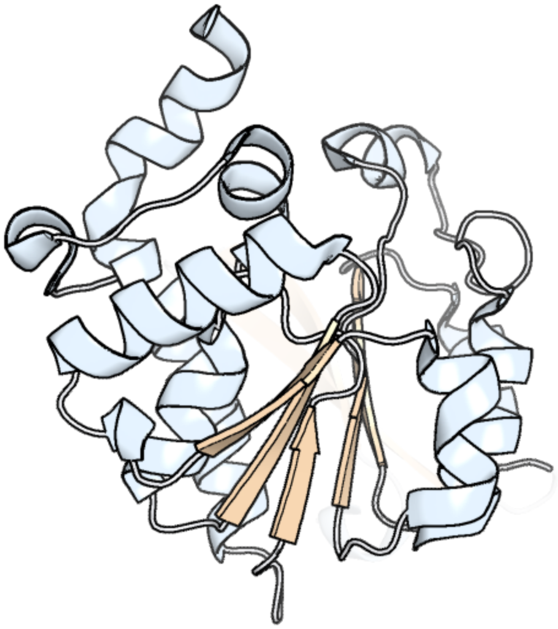

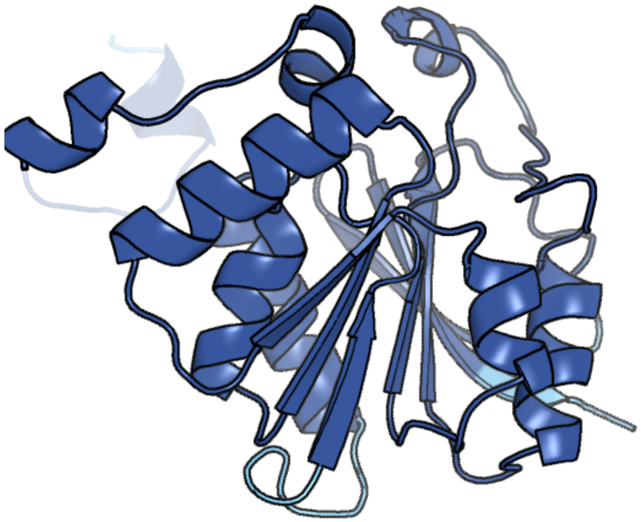

Figure 37: left: reference structure of 3gdh chain C. right: predicted structure of chlorv-1..037, unaligned sequences are shown as transparent

#### chlorv-1..038

- Sequence-based annotation for chlorv-1..038 is putative Peptidase
- Best hit was 5xu8 chain A: Ubiquitin carboxyl-terminal hydrolase 2

| target | prob | fident | alnlen | evalue | theader |
|---|---|---|---|---|---|
| 5xu8-assembly1.cif.gz_A | 1 | 0.182 | 434 | 8.761e-25 | Crystal structure of human USP2 in complex with ubiquitin and 6-thioguanine |
| 2ibi-assembly1.cif.gz_A | 1 | 0.168 | 439 | 1.039e-24 | Covalent Ubiquitin-USP2 Complex |
| 3v6e-assembly1.cif.gz_A | 1 | 0.18 | 438 | 1.461e-24 | Crystal Structure of USP2 and a mutant form of Ubiquitin |

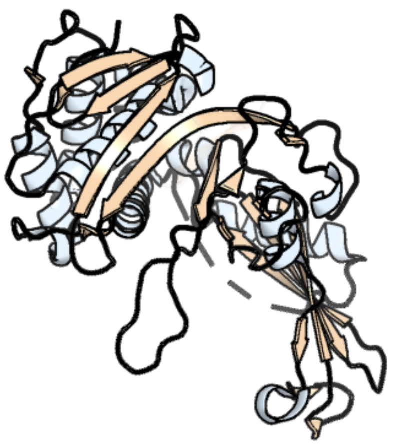

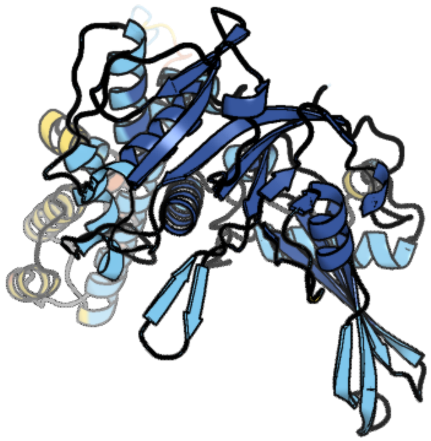

Figure 38: left: reference structure of 5xu8 chain A. right: predicted structure of chlorv-1..038, unaligned sequences are shown as transparent

#### chlorv-1..039

- Sequence-based annotation for chlorv-1..039 is putative RNA polymerase beta subunit
- Best hit was 8oev chain B: DNA-directed RNA polymerase subunit beta

| target | prob | fident | alnlen | evalue | theader |
|---|---|---|---|---|---|
| 8oev-assembly1.cif.gz_B | 1 | 0.356 | 1205 | 2.803e-102 | Structure of the mammalian Pol II-SPT6-Elongin complex, lacking ELOA latch (composite structure, structure 3) |
| 8was-assembly1.cif.gz_p | 1 | 0.355 | 1220 | 2.81e-101 | Structure of transcribing complex 9 (TC9), the initially transcribing complex with Pol II positioned 9nt downstream of TSS. |
| 7pks-assembly1.cif.gz_B | 1 | 0.36 | 1197 | 9.796e-101 | Structural basis of Integrator-mediated transcription regulation |

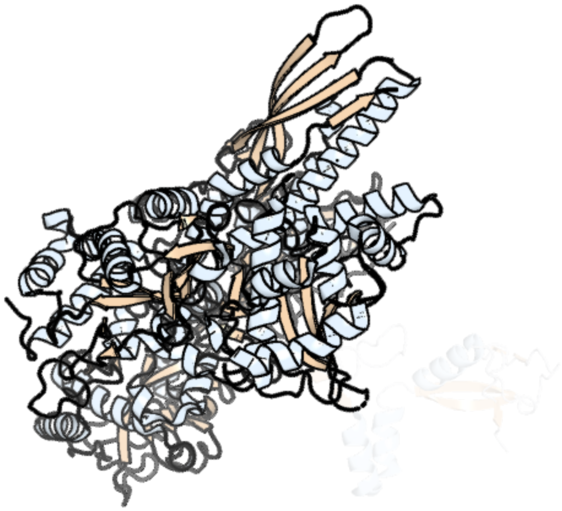

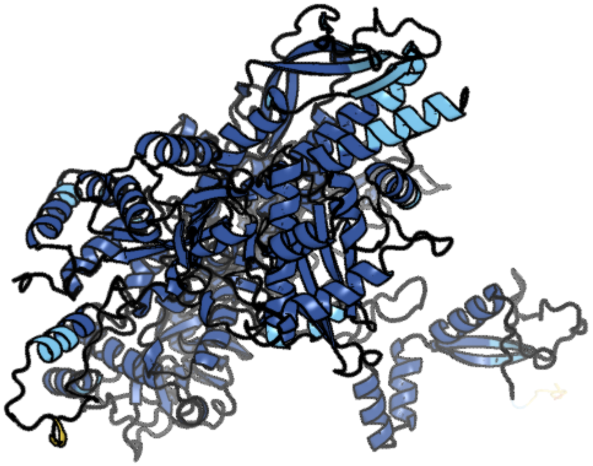

Figure 39: left: reference structure of 8oev chain B. right: predicted structure of chlorv-1..039, unaligned sequences are shown as transparent

### chlorv-1..040

- Sequence-based annotation for chlorv-1..040 is putative Translation initiation factor
- No significant structural hit found

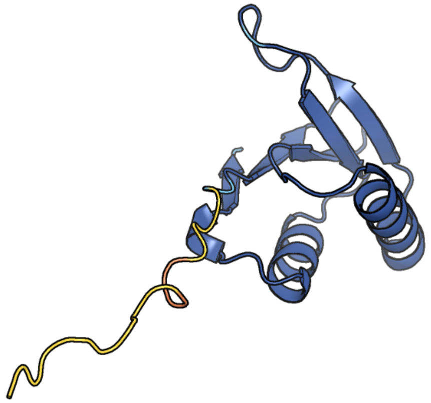

Figure 40: predicted structure of chlorv-1..040

#### chlorv-1..041

- Sequence-based annotation for chlorv-1..041 is hypothetical protein
- No significant structural hit found

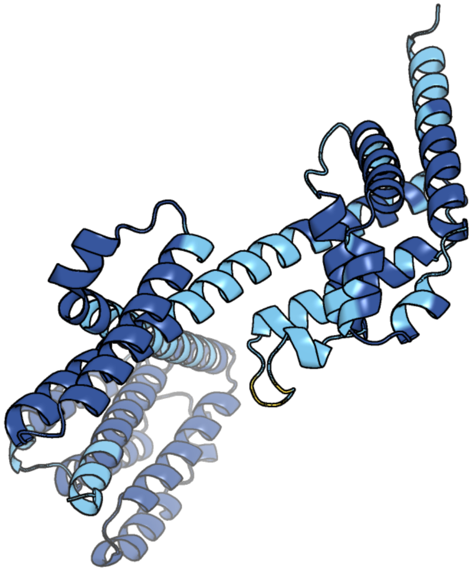

Figure 41: predicted structure of chlorv-1..041

#### chlorv-1..042

- Sequence-based annotation for chlorv-1..042 is hypothetical protein
- Best hit was 3wr7 chain A: Spermidine N1-acetyltransferase

| target | prob | fident | alnlen | evalue | theader |
|---|---|---|---|---|---|
| 3wr7-assembly1.cif.gz_A-3 | 1 | 0.161 | 161 | 6.269e-07 | Crystal Structure of Spermidine Acetyltransferase from Escherichia coli |
| 8fv0-assembly1.cif.gz_C | 1 | 0.198 | 161 | 7.418e-07 | SpeG spermidine N-acetyltransferase from Staphylococcus aureus in complex with spermine |
| 6vfn-assembly1.cif.gz_D | 1 | 0.16 | 162 | 1.098e-06 | Crystal structure of SpeG allosteric polyamine acetyltransferase from Bacillus thuringiensis in complex with spermine |

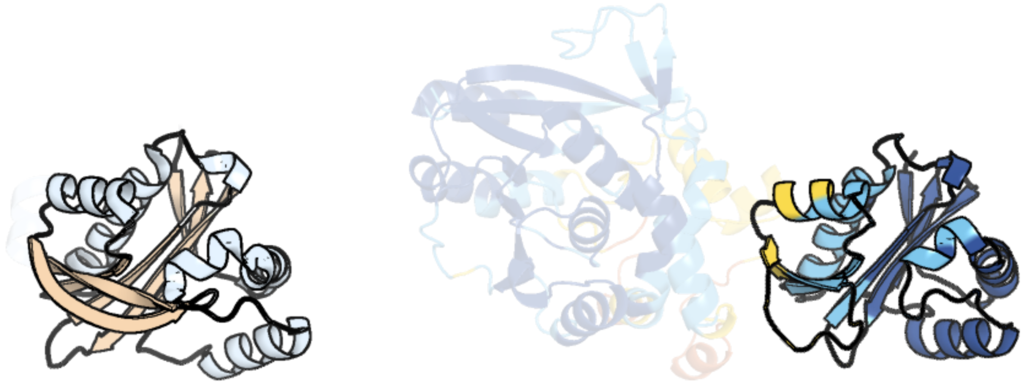

Figure 42: left: reference structure of 3wr7 chain A. right: predicted structure of chlorv-1..042, unaligned sequences are shown as transparent

### chlorv-1..043

- Sequence-based annotation for chlorv-1..043 is hypothetical protein
- Best hit was 8asn chain G: Tubulin–tyrosine ligase

| target | prob | fident | alnlen | evalue | theader |
|---|---|---|---|---|---|
| 8asn-assembly1.cif.gz_G | 1 | 0.154 | 356 | 2.528e-16 | Crystal structure of the apo human TTL in complex with tubulin-stathmin |
| 8asn-assembly1.cif.gz_I | 1 | 0.152 | 360 | 3.6e-16 | Crystal structure of the apo human TTL in complex with tubulin-stathmin |
| 8asn-assembly1.cif.gz_H | 1 | 0.147 | 359 | 8.214e-16 | Crystal structure of the apo human TTL in complex with tubulin-stathmin |

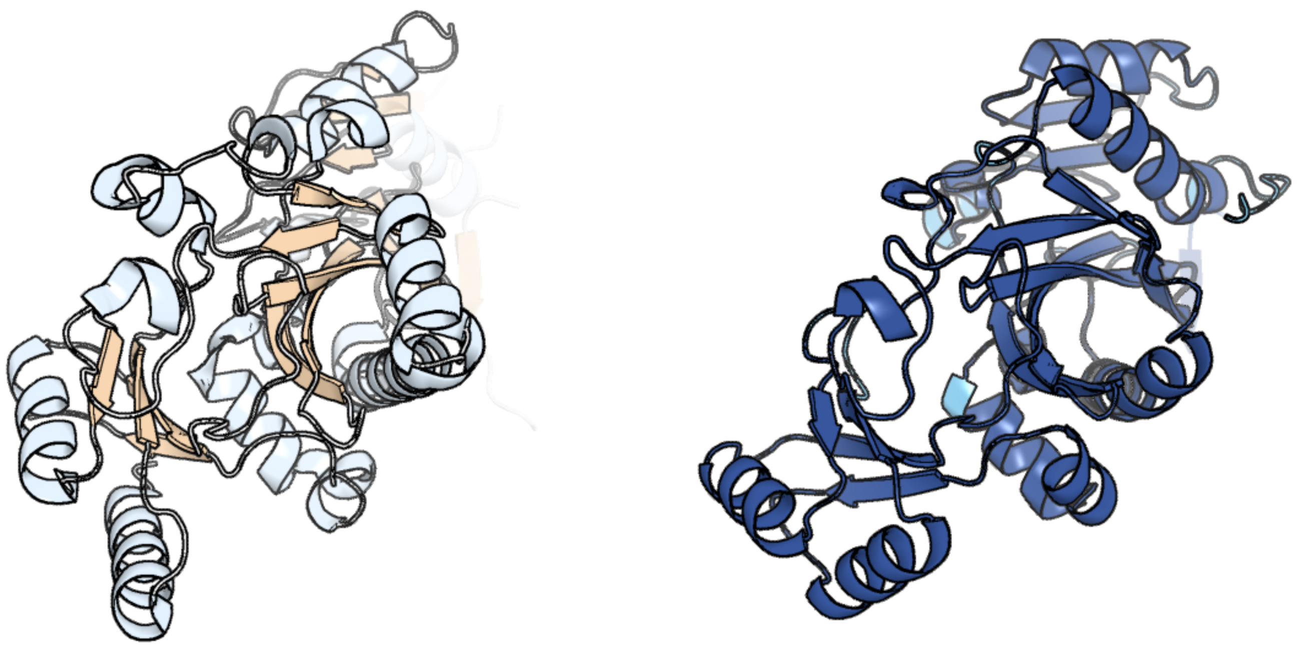

Figure 43: left: reference structure of 8asn chain G. right: predicted structure of chlorv-1..043, unaligned sequences are shown as transparent

#### chlorv-1..044

- Sequence-based annotation for chlorv-1..044 is hypothetical protein
- Best hit was 4n4a chain A: Cap-specific mRNA (nucleoside-2'-O-)-methyltransferase 1

| target | prob | fident | alnlen | evalue | theader |
|---|---|---|---|---|---|
| 4n4a-assembly1.cif.gz_A | 1 | 0.118 | 370 | 1.193e-08 | Cystal structure of Cap-specific mRNA (nucleoside-2'-O-)-methyltransferase 1 |
| 4n49-assembly1.cif.gz_A | 1 | 0.114 | 374 | 2.444e-07 | Cap-specific mRNA (nucleoside-2'-O-)-methyltransferase 1 Protein in complex with m7GpppG and SAM |
| 4n48-assembly1.cif.gz_B | 1 | 0.12 | 375 | 3.918e-07 | Cap-specific mRNA (nucleoside-2'-O-)-methyltransferase 1 Protein in complex with capped RNA fragment |

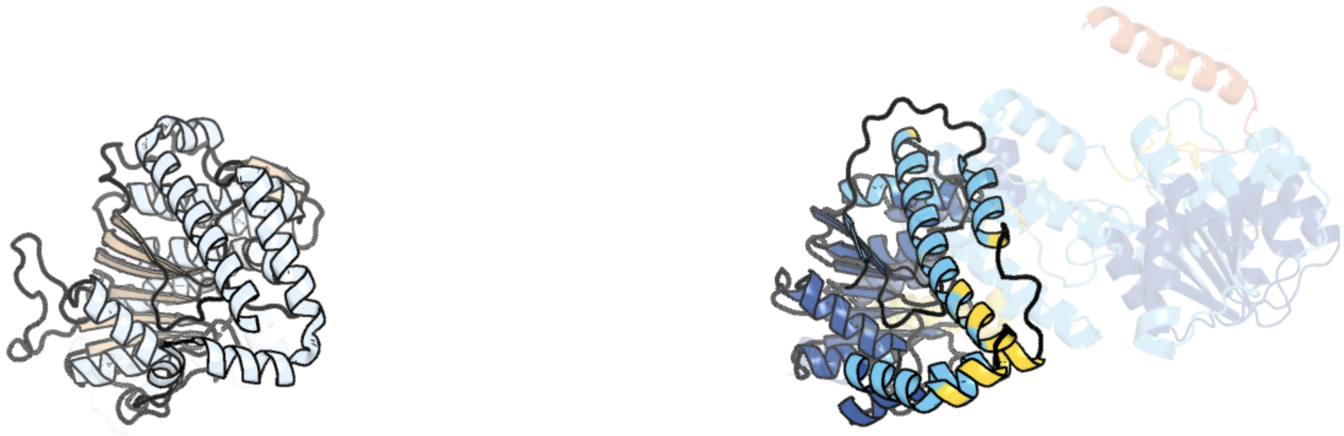

Figure 44: left: reference structure of 4n4a chain A. right: predicted structure of chlorv-1..044, unaligned sequences are shown as transparent

#### chlorv-1..045

- Sequence-based annotation for chlorv-1..045 is hypothetical protein
- No significant structural hit found

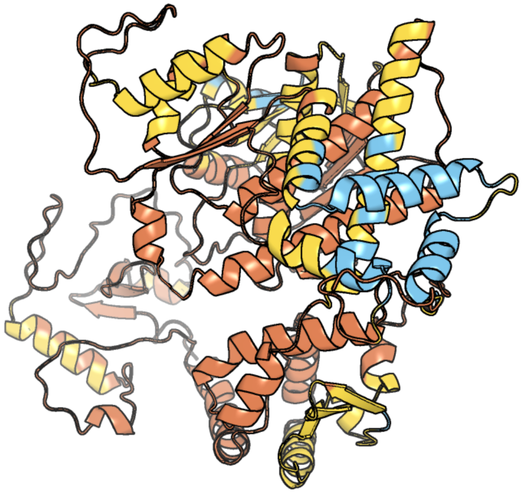

Figure 45: predicted structure of chlorv-1..045

#### chlorv-1..046

- Sequence-based annotation for chlorv-1..046 is putative Kinase
- Best hit was 1del chain A: DEOXYNUCLEOSIDE MONOPHOSPHATE KINASE

| target | prob | fident | alnlen | evalue | theader |
|---|---|---|---|---|---|
| 1del-assembly1.cif.gz_A | 1 | 0.197 | 243 | 1.807e-07 | DEOXYNUCLEOSIDE MONOPHOSPHATE KINASE COMPLEXED WITH DEOXY-GMP AND AMP |
| 2grj-assembly2.cif.gz_B | 1 | 0.177 | 197 | 5.028e-06 | Crystal structure of Dephospho-CoA kinase (EC 2.7.1.24) (Dephosphocoenzyme A kinase) (tm1387) from THERMOTOGA MARITIMA at 2.60 A resolution |
| 1del-assembly1.cif.gz_B | 1 | 0.16 | 256 | 2.737e-05 | DEOXYNUCLEOSIDE MONOPHOSPHATE KINASE COMPLEXED WITH DEOXY-GMP AND AMP |

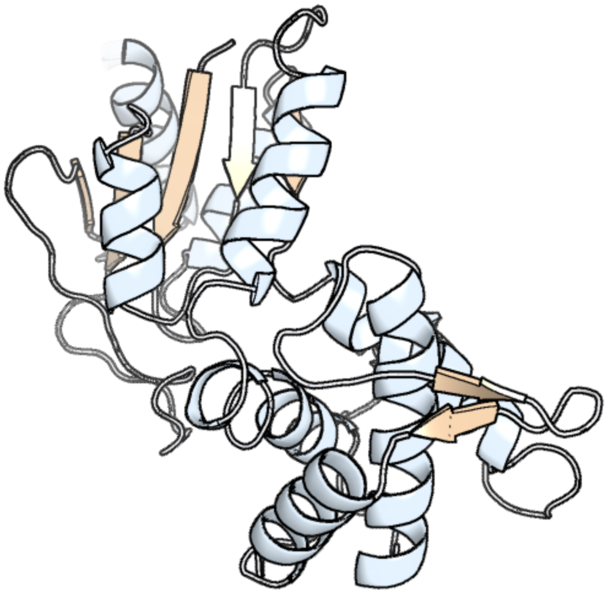

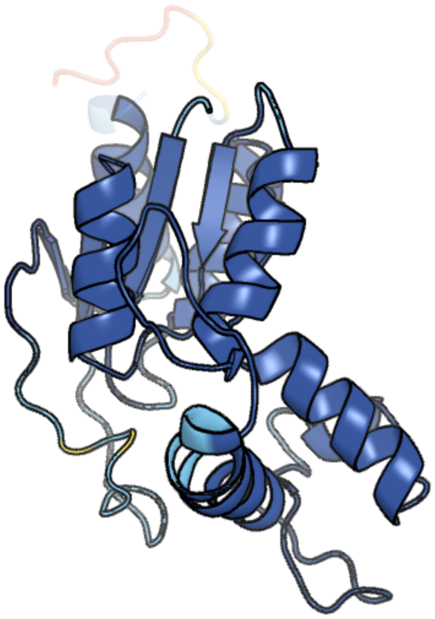

Figure 46: left: reference structure of 1del chain A. right: predicted structure of chlorv-1..046, unaligned sequences are shown as transparent

#### chlorv-1..047

- Sequence-based annotation for chlorv-1..047 is putative Protein kinase
- Best hit was 5ezv chain A: 5'-AMP-activated protein kinase catalytic subunit alpha-2/alpha-1 RIM SWAP chimera

| target | prob | fident | alnlen | evalue | theader |
|---|---|---|---|---|---|
| 5ezv-assembly1.cif.gz_A | 1 | 0.216 | 332 | 6.039e-16 | X-ray crystal structure of AMP-activated protein kinase alpha-2/alpha-1 RIM chimaera (alpha-2(1-347)/alpha-1(349-401)/alpha-2(397-end) beta-1 gamma-1) co-crystallized with C2 (5-(5-hydroxyl-isoxazol-3-yl)-furan-2-phosphonic acid) |
| 3iec-assembly4.cif.gz_D | 1 | 0.227 | 321 | 6.349e-16 | Helicobacter pylori CagA Inhibits PAR1/MARK Family Kinases by Mimicking Host Substrates |
| 6c9d-assembly1.cif.gz_A | 1 | 0.236 | 322 | 1.218e-15 | Crystal structure of KA1-autoinhibited MARK1 kinase |

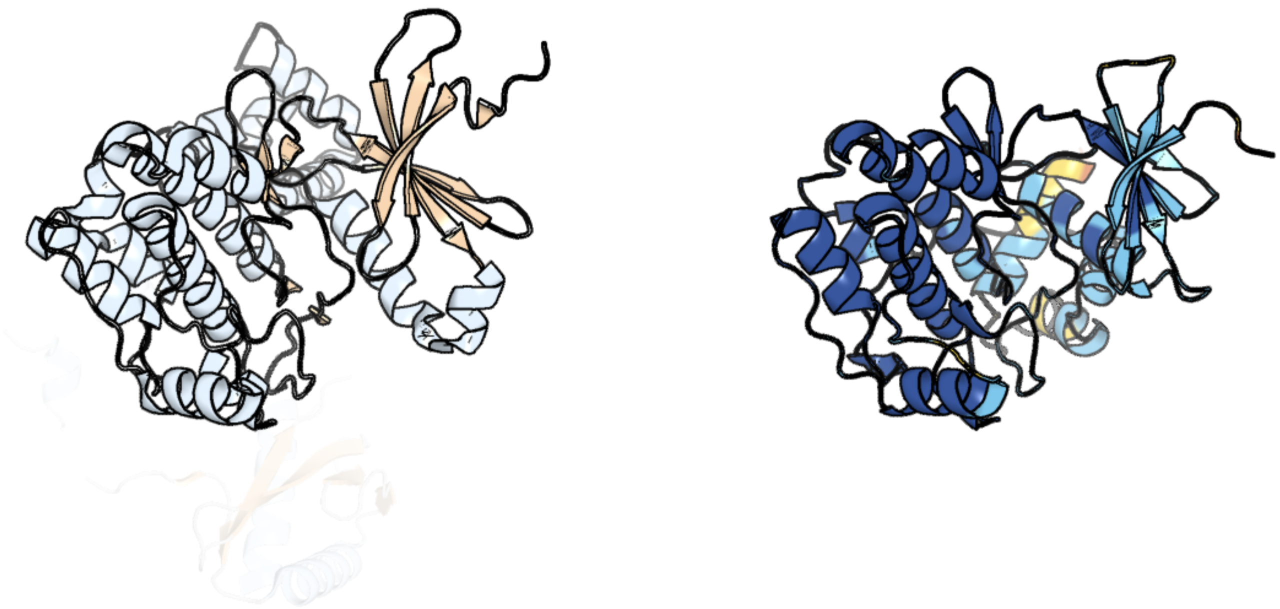

Figure 47: left: reference structure of 5ezv chain A. right: predicted structure of chlorv-1..047, unaligned sequences are shown as transparent

#### chlorv-1..048

- Sequence-based annotation for chlorv-1..048 is hypothetical protein
- No significant structural hit found

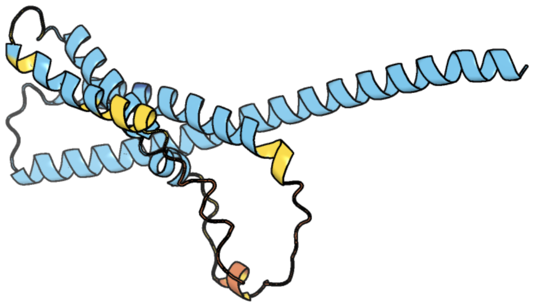

Figure 48: predicted structure of chlorv-1..048

#### chlorv-1..049

- Sequence-based annotation for chlorv-1..049 is putative dUTPase
- Best hit was 3t64 chain A: Deoxyuridine 5'-triphosphate nucleotidohydrolase, putative

| target | prob | fident | alnlen | evalue | theader |
|---|---|---|---|---|---|
| 3t64-assembly2.cif.gz_A | 1 | 0.311 | 183 | 1.075e-12 | 5'-Diphenyl Nucleoside Inhibitors of Plasmodium falciparum dUTPase |
| 1vyq-assembly1.cif.gz_A | 1 | 0.307 | 166 | 1.724e-12 | Novel inhibitors of Plasmodium Falciparum dUTPase provide a platform for anti-malarial drug design |
| 3t64-assembly2.cif.gz_C | 1 | 0.331 | 178 | 3.3e-12 | 5'-Diphenyl Nucleoside Inhibitors of Plasmodium falciparum dUTPase |

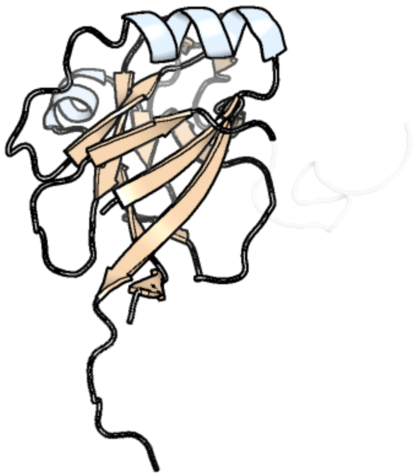

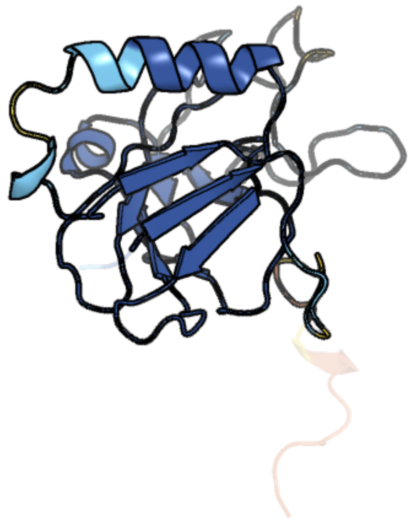

Figure 49: left: reference structure of 3t64 chain A. right: predicted structure of chlorv-1..049, unaligned sequences are shown as transparent

#### chlorv-1..050

- Sequence-based annotation for chlorv-1..050 is hypothetical protein
- No significant structural hit found

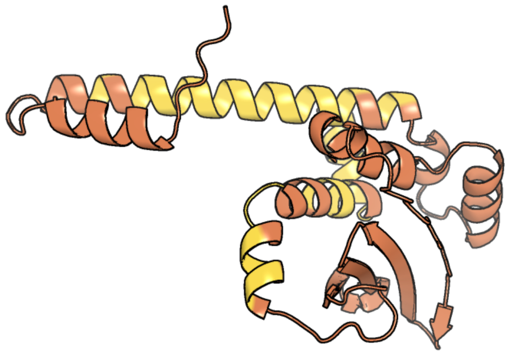

Figure 50: predicted structure of chlorv-1..050

#### chlorv-1..051

- Sequence-based annotation for chlorv-1..051 is hypothetical protein
- No significant structural hit found

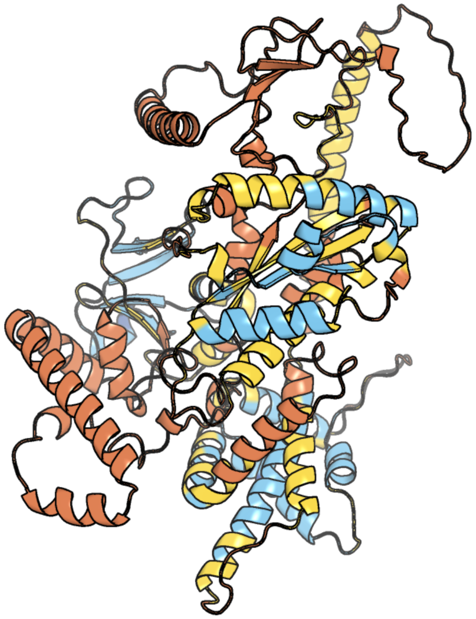

Figure 51: predicted structure of chlorv-1..051

#### chlorv-1..052

- Sequence-based annotation for chlorv-1..052 is putative Methyltransferase
- Best hit was 4n48 chain B: Cap-specific mRNA (nucleoside-2'-O-)-methyltransferase 1

| target | prob | fident | alnlen | evalue | theader |
|---|---|---|---|---|---|
| 4n48-assembly1.cif.gz_B | 1 | 0.145 | 418 | 1.196e-10 | Cap-specific mRNA (nucleoside-2'-O-)-methyltransferase 1 Protein in complex with capped RNA fragment |
| 4n48-assembly2.cif.gz_A | 1 | 0.146 | 409 | 1.314e-10 | Cap-specific mRNA (nucleoside-2'-O-)-methyltransferase 1 Protein in complex with capped RNA fragment |
| 4n49-assembly1.cif.gz_A | 1 | 0.154 | 415 | 1.662e-10 | Cap-specific mRNA (nucleoside-2'-O-)-methyltransferase 1 Protein in complex with m7GpppG and SAM |

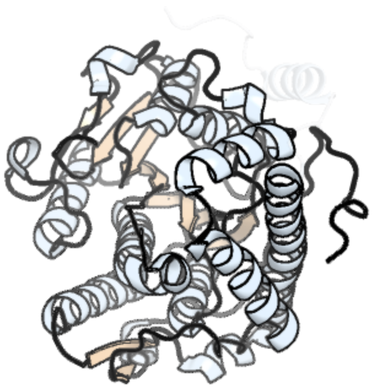

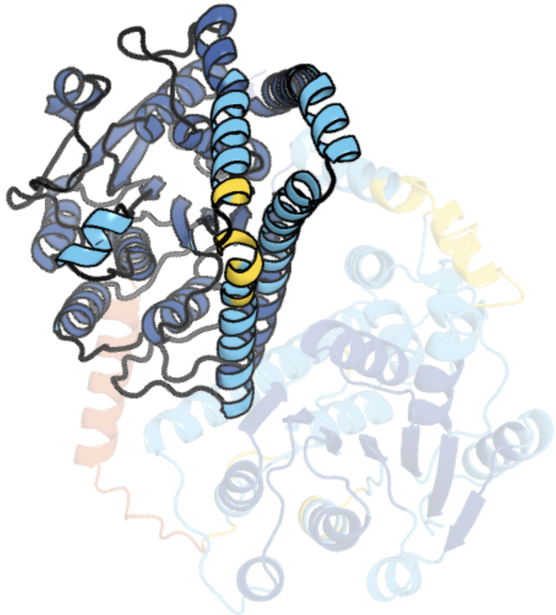

Figure 52: left: reference structure of 4n48 chain B. right: predicted structure of chlorv-1..052, unaligned sequences are shown as transparent

#### chlorv-1..053

- Sequence-based annotation for chlorv-1..053 is putative Methyltransferase
- Best hit was 4n48 chain A: Cap-specific mRNA (nucleoside-2'-O-)-methyltransferase 1

| target | prob | fident | alnlen | evalue | theader |
|---|---|---|---|---|---|
| 4n48-assembly2.cif.gz_A | 1 | 0.137 | 416 | 1.075e-09 | Cap-specific mRNA (nucleoside-2'-O-)-methyltransferase 1 Protein in complex with capped RNA fragment |
| 4n48-assembly1.cif.gz_B | 1 | 0.14 | 413 | 1.075e-09 | Cap-specific mRNA (nucleoside-2'-O-)-methyltransferase 1 Protein in complex with capped RNA fragment |
| 4n4a-assembly1.cif.gz_A | 1 | 0.139 | 401 | 1.348e-09 | Cystal structure of Cap-specific mRNA (nucleoside-2'-O-)-methyltransferase 1 |

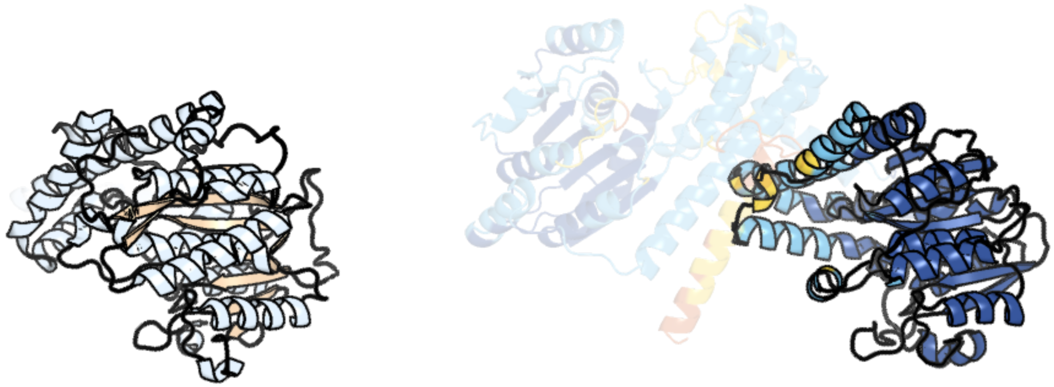

Figure 53: left: reference structure of 4n48 chain A. right: predicted structure of chlorv-1..053, unaligned sequences are shown as transparent

### chlorv-1..054

- Sequence-based annotation for chlorv-1..054 is putative Protein kinase
- Best hit was 3ckx chain A: Serine/threonine-protein kinase 24

| target | prob | fident | alnlen | evalue | theader |
|---|---|---|---|---|---|
| 3ckx-assembly1.cif.gz_A | 1 | 0.269 | 252 | 3.415e-20 | Crystal structure of sterile 20-like kinase 3 (MST3, STK24) in complex with staurosporine |
| 4zy4-assembly1.cif.gz_A | 1 | 0.274 | 237 | 3.622e-20 | Crystal structure of P21 activated kinase 1 in complex with an inhibitor compound 4 |
| 5dew-assembly1.cif.gz_A | 1 | 0.256 | 242 | 4.858e-20 | Crystal structure of PAK1 in complex with an inhibitor compound 5 |

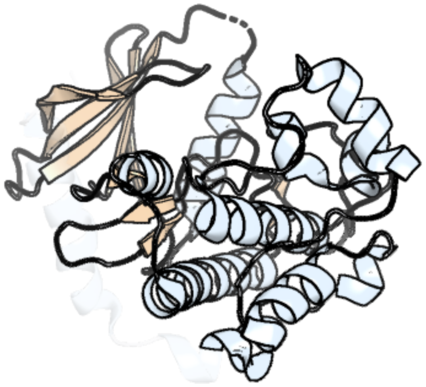

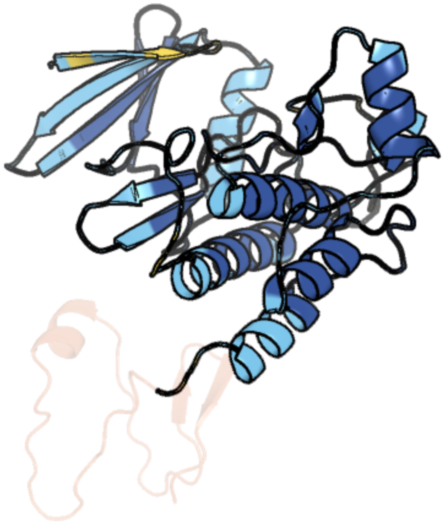

Figure 54: left: reference structure of 3ckx chain A. right: predicted structure of chlorv-1..054, unaligned sequences are shown as transparent

#### chlorv-1..055

- Sequence-based annotation for chlorv-1..055 is hypothetical protein
- No significant structural hit found

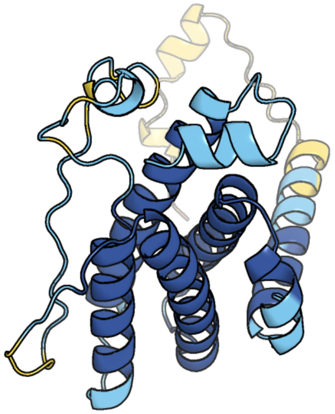

Figure 55: predicted structure of chlorv-1..055

#### chlorv-1..056

- Sequence-based annotation for chlorv-1..056 is hypothetical protein
- No significant structural hit found

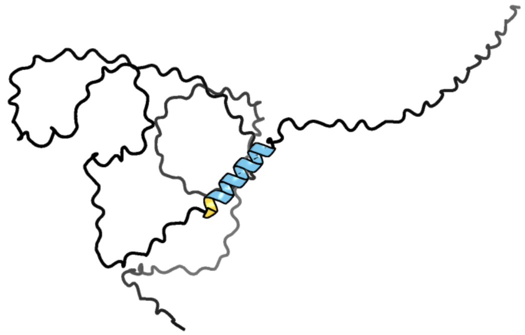

Figure 56: predicted structure of chlorv-1..056

#### chlorv-1..057

- Sequence-based annotation for chlorv-1..057 is hypothetical protein
- No significant structural hit found

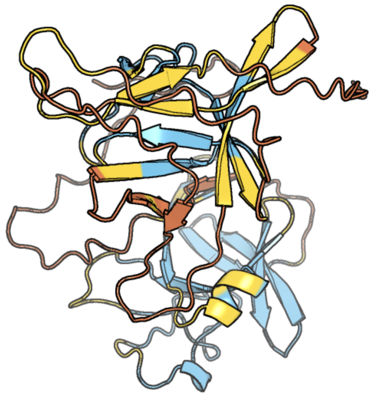

Figure 57: predicted structure of chlorv-1..057

#### chlorv-1..058

- Sequence-based annotation for chlorv-1..058 is hypothetical protein
- No significant structural hit found

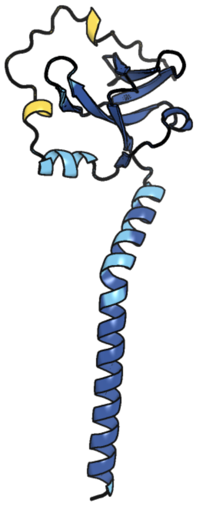

Figure 58: predicted structure of chlorv-1..058

#### chlorv-1..059

- Sequence-based annotation for chlorv-1..059 is putative Ribonucleotide reductase small subunit
- Best hit was 3vpm chain B: Ribonucleoside-diphosphate reductase subunit M2

| target | prob | fident | alnlen | evalue | theader |
|---|---|---|---|---|---|
| 3vpm-assembly1.cif.gz_B | 1 | 0.516 | 281 | 5.282e-23 | Crystal structure of human ribonucleotide reductase subunit M2 (hRRM2) mutant |
| 3vpo-assembly1.cif.gz_B | 1 | 0.516 | 281 | 7.401e-23 | Crystal structure of human ribonucleotide reductase subunit M2 (hRRM2) mutant |
| 3olj-assembly1.cif.gz_D | 1 | 0.519 | 281 | 1.28e-22 | Crystal structure of human ribonucleotide reductase subunit M2 (hRRM2) |

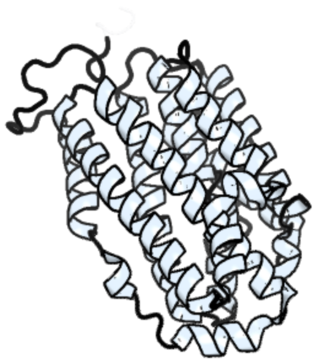

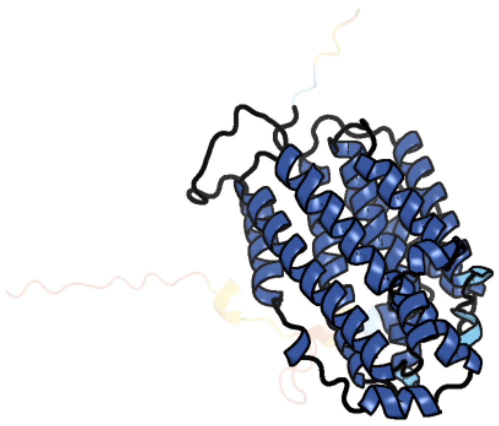

Figure 59: left: reference structure of 3vpm chain B. right: predicted structure of chlorv-1..059, unaligned sequences are shown as transparent

#### chlorv-1..060

- Sequence-based annotation for chlorv-1..060 is hypothetical protein
- No significant structural hit found

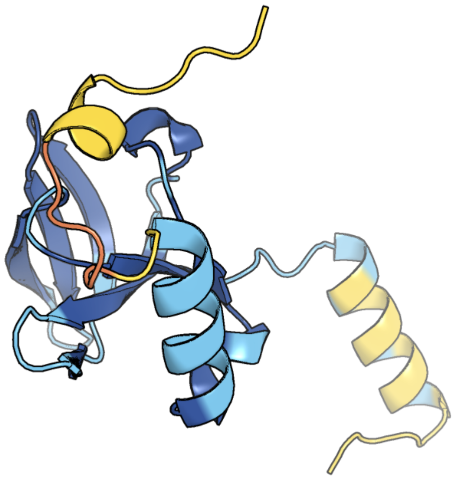

Figure 60: predicted structure of chlorv-1..060

### chlorv-1..061

- Sequence-based annotation for chlorv-1..061 is hypothetical protein
- Best hit was 6u05 chain A: tRNA ligase

| target | prob | fident | alnlen | evalue | theader |
|---|---|---|---|---|---|
| 6u05-assembly1.cif.gz_A | 1 | 0.126 | 449 | 1.103e-08 | Crystal Structure of Fungal RNA Kinase |
| 4jt2-assembly1.cif.gz_A | 1 | 0.257 | 140 | 9.69e-07 | Structure of Clostridium thermocellum polynucleotide kinase bound to CTP |
| 4gp6-assembly1.cif.gz_B-2 | 1 | 0.25 | 140 | 1.154e-06 | Polynucleotide kinase |

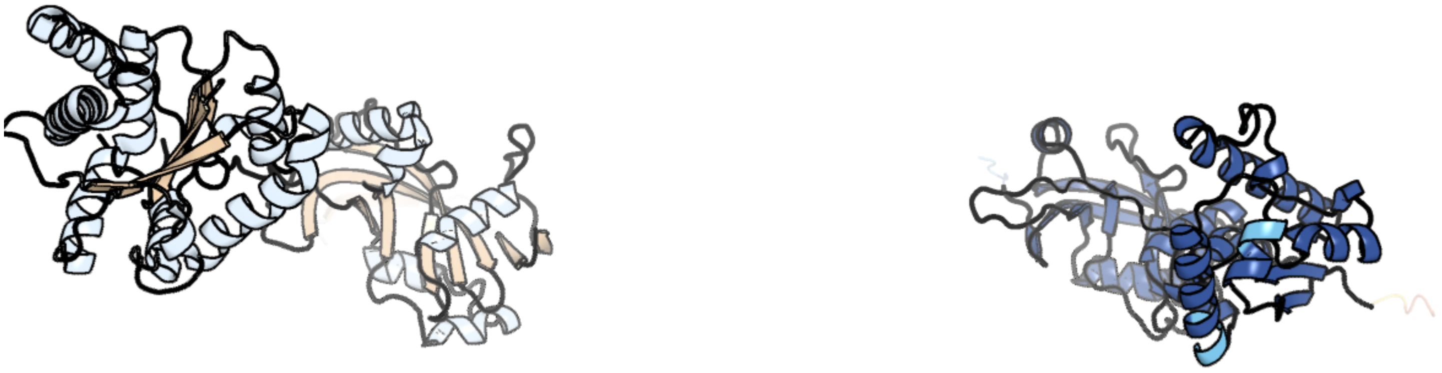

Figure 61: left: reference structure of 6u05 chain A. right: predicted structure of chlorv-1..061, unaligned sequences are shown as transparent

#### chlorv-1..062

- Sequence-based annotation for chlorv-1..062 is hypothetical protein
- No significant structural hit found

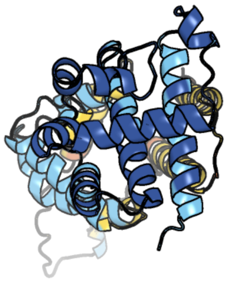

Figure 62: predicted structure of chlorv-1..062

#### chlorv-1..063

- Sequence-based annotation for chlorv-1..063 is hypothetical protein
- No significant structural hit found

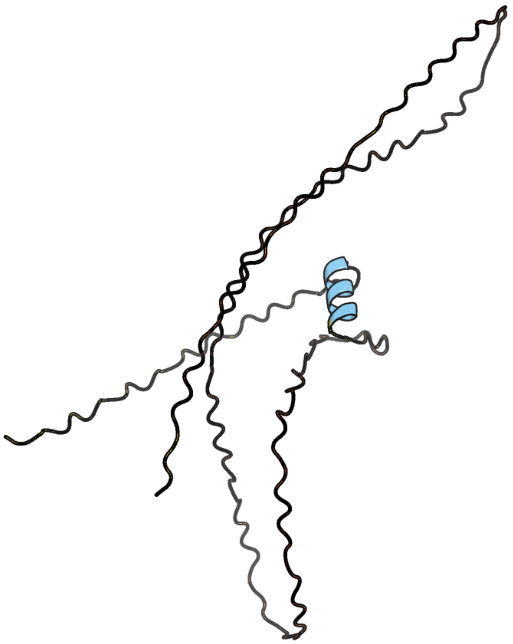

Figure 63: predicted structure of chlorv-1..063

#### chlorv-1..064

- Sequence-based annotation for chlorv-1..064 is putative eukaryotic translation initiation factor 5
- Best hit was 6fyx chain m: Eukaryotic translation initiation factor 5

| target | prob | fident | alnlen | evalue | theader |
|---|---|---|---|---|---|
| 6fyx-assembly1.cif.gz_m | 1 | 0.307 | 140 | 5.861e-10 | Structure of a partial yeast 48S preinitiation complex with eIF5 N-terminal domain (model C1) |
| 8cas-assembly1.cif.gz_m | 1 | 0.285 | 140 | 1.178e-09 | Cryo-EM structure of native Otu2-bound ubiquitinated 48S initiation complex (partial) |
| 2e9h-assembly1.cif.gz_A | 1 | 0.274 | 142 | 1.249e-09 | Solution structure of the eIF-5_eIF-2B domain from human Eukaryotic translation initiation factor 5 |

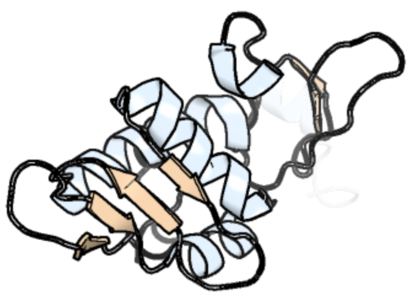

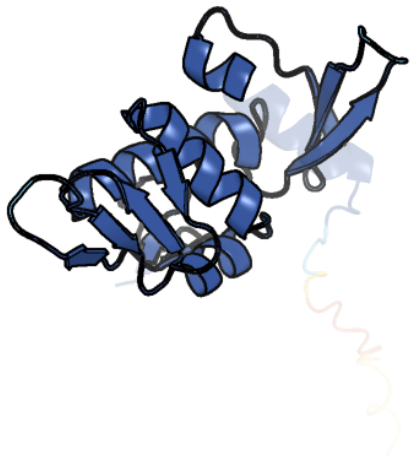

Figure 64: left: reference structure of 6fyx chain m. right: predicted structure of chlorv-1..064, unaligned sequences are shown as transparent

#### chlorv-1..065

- Sequence-based annotation for chlorv-1..065 is putative DNA polymerase family B
- Best hit was 6p1h chain A: DNA polymerase delta catalytic subunit

| target | prob | fident | alnlen | evalue | theader |
|---|---|---|---|---|---|
| 6p1h-assembly1.cif.gz_A | 1 | 0.197 | 1330 | 4.259e-56 | Cryo-EM Structure of DNA Polymerase Delta Holoenzyme |
| 3iay-assembly1.cif.gz_A | 1 | 0.205 | 1311 | 4.399e-55 | Ternary complex of DNA polymerase delta |
| 7kc0-assembly1.cif.gz_A | 1 | 0.201 | 1313 | 1.189e-51 | Structure of the Saccharomyces cerevisiae replicative polymerase delta in complex with a primer/template and the PCNA clamp |

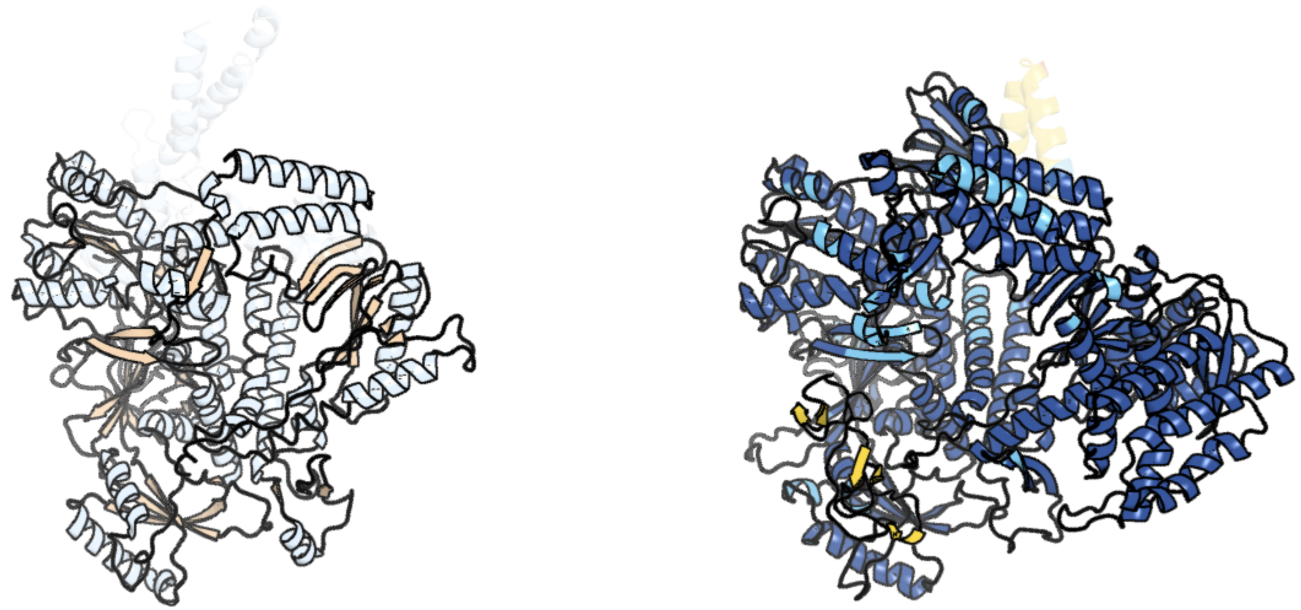

Figure 65: left: reference structure of 6p1h chain A. right: predicted structure of chlorv-1..065, unaligned sequences are shown as transparent

#### chlorv-1..066

- Sequence-based annotation for chlorv-1..066 is hypothetical protein
- No significant structural hit found

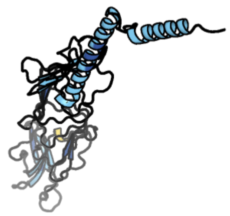

Figure 66: predicted structure of chlorv-1..066

#### chlorv-1..067

- Sequence-based annotation for chlorv-1..067 is putative Protein kinase
- No significant structural hit found

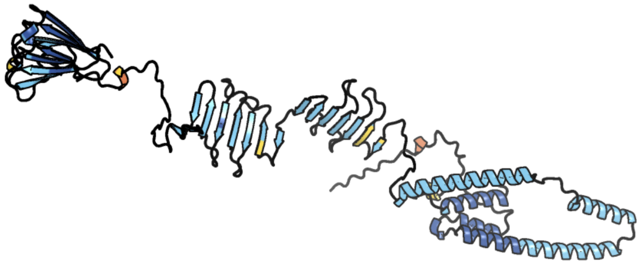

Figure 67: predicted structure of chlorv-1..067

#### chlorv-1..068

- Sequence-based annotation for chlorv-1..068 is putative Arginase
- Best hit was 4q3r chain D: Arginase

| target | prob | fident | alnlen | evalue | theader |
|---|---|---|---|---|---|
| 4q3r-assembly4.cif.gz_D | 1 | 0.284 | 299 | 3.526e-22 | Crystal structure of Schistosoma mansoni arginase in complex with inhibitor ABHDP |
| 4q3v-assembly3.cif.gz_C | 1 | 0.277 | 303 | 3.734e-22 | Crystal structure of Schistosoma mansoni arginase in complex with inhibitor BEC |
| 4q3v-assembly1.cif.gz_A | 1 | 0.284 | 302 | 4.698e-22 | Crystal structure of Schistosoma mansoni arginase in complex with inhibitor BEC |

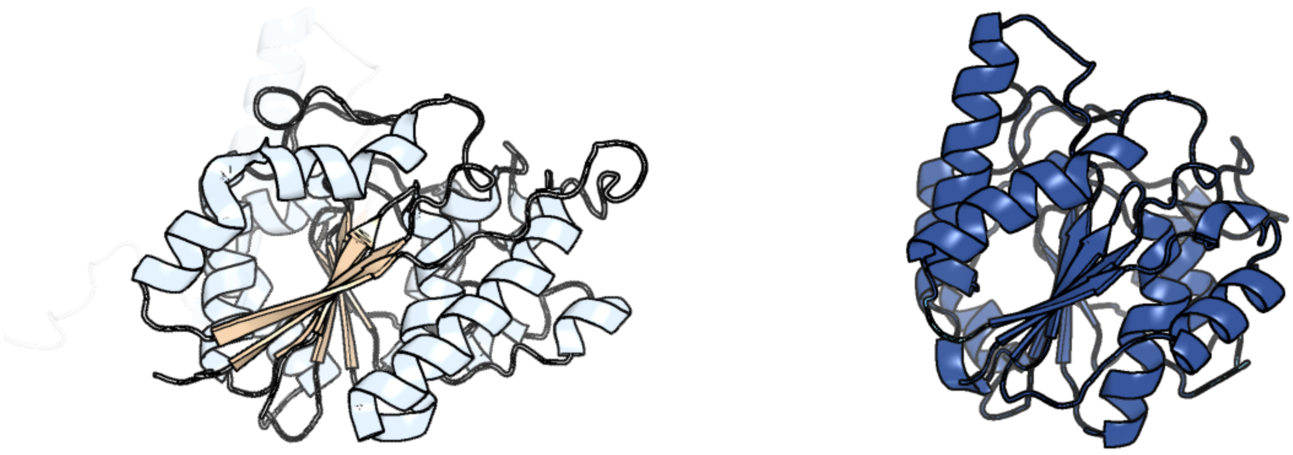

Figure 68: left: reference structure of 4q3r chain D. right: predicted structure of chlorv-1..068, unaligned sequences are shown as transparent

#### chlorv-1..069

- Sequence-based annotation for chlorv-1..069 is putative Ankyrin repeat protein
- Best hit was 5op1 chain A: DARPin A4

| target | prob | fident | alnlen | evalue | theader |
|---|---|---|---|---|---|
| 5op1-assembly1.cif.gz_A | 1 | 0.205 | 170 | 2.926e-07 | Designed Ankyrin Repeat Protein (DARPin) A4 in complex with Lysozyme |
| 6h46-assembly1.cif.gz_B | 1 | 0.195 | 174 | 3.791e-07 | Human KRAS in complex with darpin K13 |
| 4hna-assembly1.cif.gz_D | 1 | 0.184 | 179 | 6.042e-07 | Kinesin motor domain in the ADP-MG-ALFX state in complex with tubulin and a DARPIN |

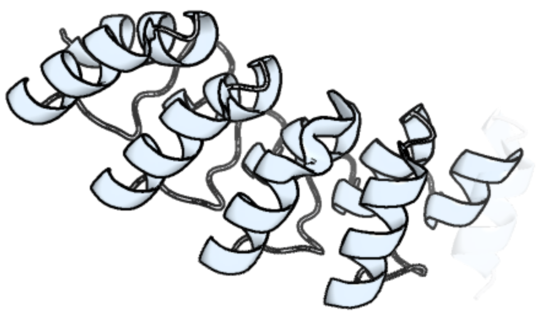

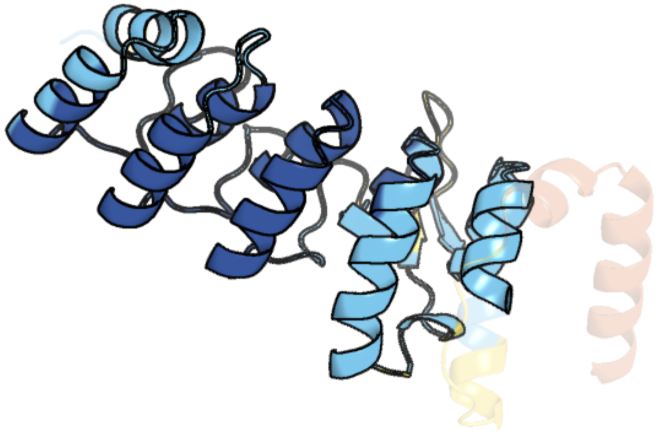

Figure 69: left: reference structure of 5op1 chain A. right: predicted structure of chlorv-1..069, unaligned sequences are shown as transparent

#### chlorv-1..070

- Sequence-based annotation for chlorv-1..070 is hypothetical protein
- No significant structural hit found

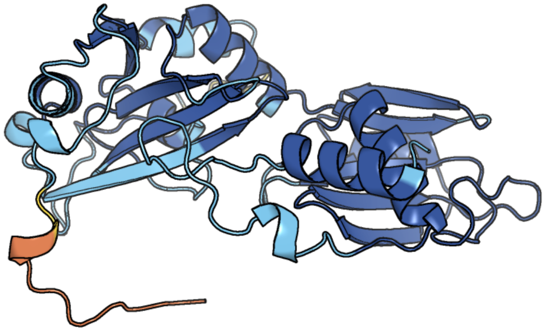

Figure 70: predicted structure of chlorv-1..070

#### chlorv-1..071

- Sequence-based annotation for chlorv-1..071 is putative Ankyrin repeat protein
- Best hit was 1n11 chain A: Ankyrin

| target | prob | fident | alnlen | evalue | theader |
|---|---|---|---|---|---|
| 8cs9-assembly1.cif.gz_A | 1 | 0.126 | 459 | 1.177e-06 | Composite reconstruction of Class 1 of the erythrocyte ankyrin-1 complex |
| 7v0x-assembly1.cif.gz_J | 1 | 0.115 | 397 | 2.308e-06 | Local refinement of ankyrin-1 (C-terminal half), class 1 of erythrocyte ankyrin-1 complex |
| 1n11-assembly1.cif.gz_A | 1 | 0.106 | 402 | 4.11e-06 | D34 REGION OF HUMAN ANKYRIN-R AND LINKER |

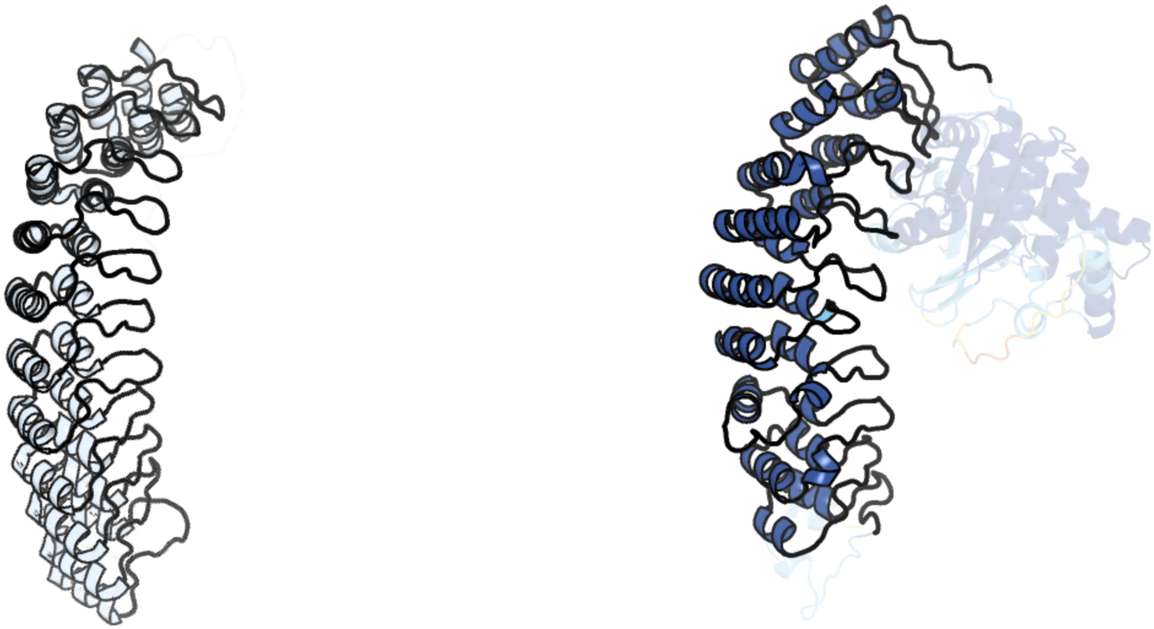

Figure 71: left: reference structure of 1n11 chain A. right: predicted structure of chlorv-1..071, unaligned sequences are shown as transparent

#### chlorv-1..072

- Sequence-based annotation for chlorv-1..072 is hypothetical protein
- Best hit was 8ppl chain Iq: Eukaryotic translation initiation factor 1A, X-chromosomal

| target | prob | fident | alnlen | evalue | theader |
|---|---|---|---|---|---|
| 8ppl-assembly1.cif.gz_Iq | 1 | 0.232 | 112 | 3.688e-06 | MERS-CoV Nsp1 bound to the human 43S pre-initiation complex |
| 6fyy-assembly1.cif.gz_i | 1 | 0.243 | 111 | 6.684e-06 | Structure of a partial yeast 48S preinitiation complex with eIF5 N-terminal domain (model C2) |
| 2oqk-assembly1.cif.gz_A | 1 | 0.26 | 100 | 1.364e-05 | Crystal structure of putative Cryptosporidium parvum translation initiation factor eIF-1A |

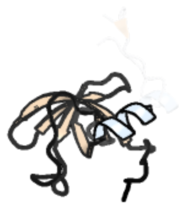

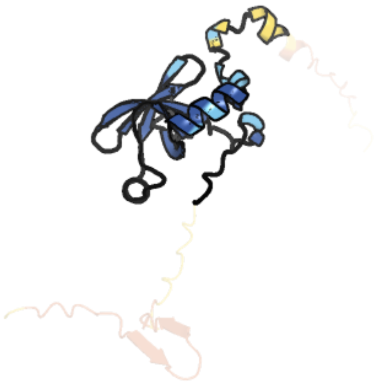

Figure 72: left: reference structure of 8ppl chain Iq. right: predicted structure of chlorv-1..072, unaligned sequences are shown as transparent

#### chlorv-1..073

- Sequence-based annotation for chlorv-1..073 is hypothetical protein
- No significant structural hit found

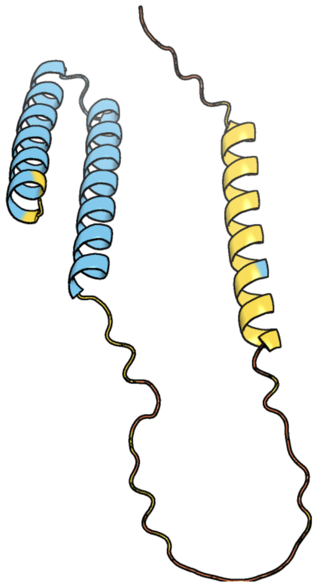

Figure 73: predicted structure of chlorv-1..073

#### chlorv-1..074

- Sequence-based annotation for chlorv-1..074 is hypothetical protein
- No significant structural hit found

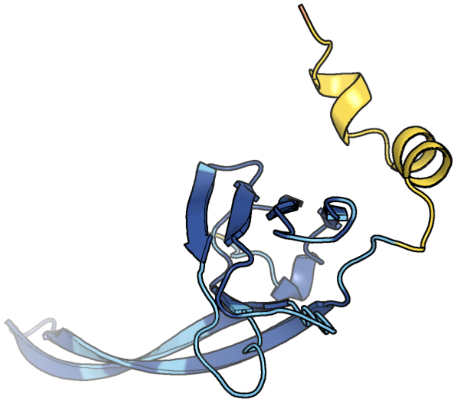

Figure 74: predicted structure of chlorv-1..074

#### chlorv-1..075

- Sequence-based annotation for chlorv-1..075 is putative Peptidase
- Best hit was 4rgh chain A: Protein DDI1 homolog 2

| target | prob | fident | alnlen | evalue | theader |
|---|---|---|---|---|---|
| 4rgh-assembly1.cif.gz_A | 1 | 0.303 | 122 | 4.185e-13 | Human DNA Damage-Inducible Protein: From Protein Chemistry and 3D Structure to Deciphering its Cellular Role |
| 4z2z-assembly1.cif.gz_B | 1 | 0.336 | 122 | 5.881e-13 | New crystal structure of yeast Ddi1 aspartyl protease reveals substrate engagement mode |
| 5yq8-assembly2.cif.gz_D | 1 | 0.333 | 120 | 8.938e-12 | Crystal structure of retroviral protease-like domain of Ddi1 from Leishmania major |

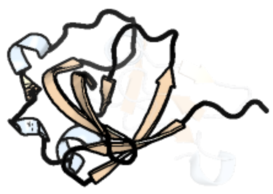

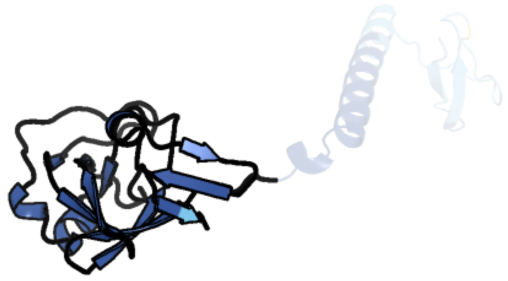

Figure 75: left: reference structure of 4rgh chain A. right: predicted structure of chlorv-1..075, unaligned sequences are shown as transparent

#### chlorv-1..076

- Sequence-based annotation for chlorv-1..076 is hypothetical protein
- No significant structural hit found

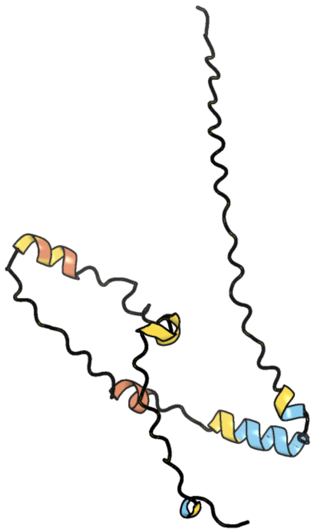

Figure 76: predicted structure of chlorv-1..076

#### chlorv-1..077

- Sequence-based annotation for chlorv-1..077 is putative Ubiquitin Conjugating Enzyme
- Best hit was 3ceg chain A: Baculoviral IAP repeat-containing protein 6

| target | prob | fident | alnlen | evalue | theader |
|---|---|---|---|---|---|
| 3ceg-assembly1.cif.gz_A | 1 | 0.385 | 285 | 1.257e-23 | Crystal structure of the UBC domain of baculoviral IAP repeat-containing protein 6 |
| 3ceg-assembly2.cif.gz_B | 1 | 0.383 | 284 | 6.597e-23 | Crystal structure of the UBC domain of baculoviral IAP repeat-containing protein 6 |
| 8gxr-assembly1.cif.gz_A | 1 | 0.266 | 274 | 1.517e-15 | crystal structure of UBC domain of UBE2O |

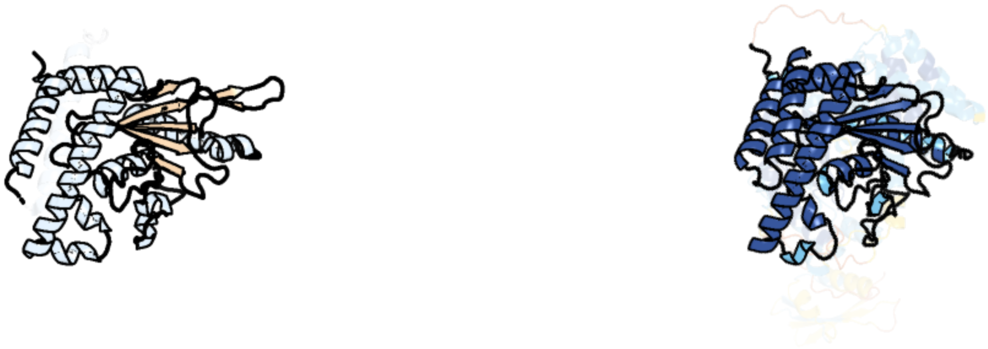

Figure 77: left: reference structure of 3ceg chain A. right: predicted structure of chlorv-1..077, unaligned sequences are shown as transparent

#### chlorv-1..078

- Sequence-based annotation for chlorv-1..078 is putative Methyltransferase
- Best hit was 3ubt chain B: Modification methylase HaeIII

| target | prob | fident | alnlen | evalue | theader |
|---|---|---|---|---|---|
| 3ubt-assembly3.cif.gz_B | 1 | 0.215 | 385 | 6.425e-20 | Crystal Structure of C71S Mutant of DNA Cytosine-5 Methyltransferase M.HaeIII Bound to DNA |
| 3g7u-assembly1.cif.gz_A | 1 | 0.185 | 398 | 2.36e-19 | Crystal structure of putative DNA modification methyltransferase encoded within prophage Cp-933R (E.coli) |
| 1dct-assembly2.cif.gz_B | 1 | 0.223 | 381 | 2.671e-19 | DNA (CYTOSINE-5) METHYLASE FROM HAEIII COVALENTLY BOUND TO DNA |

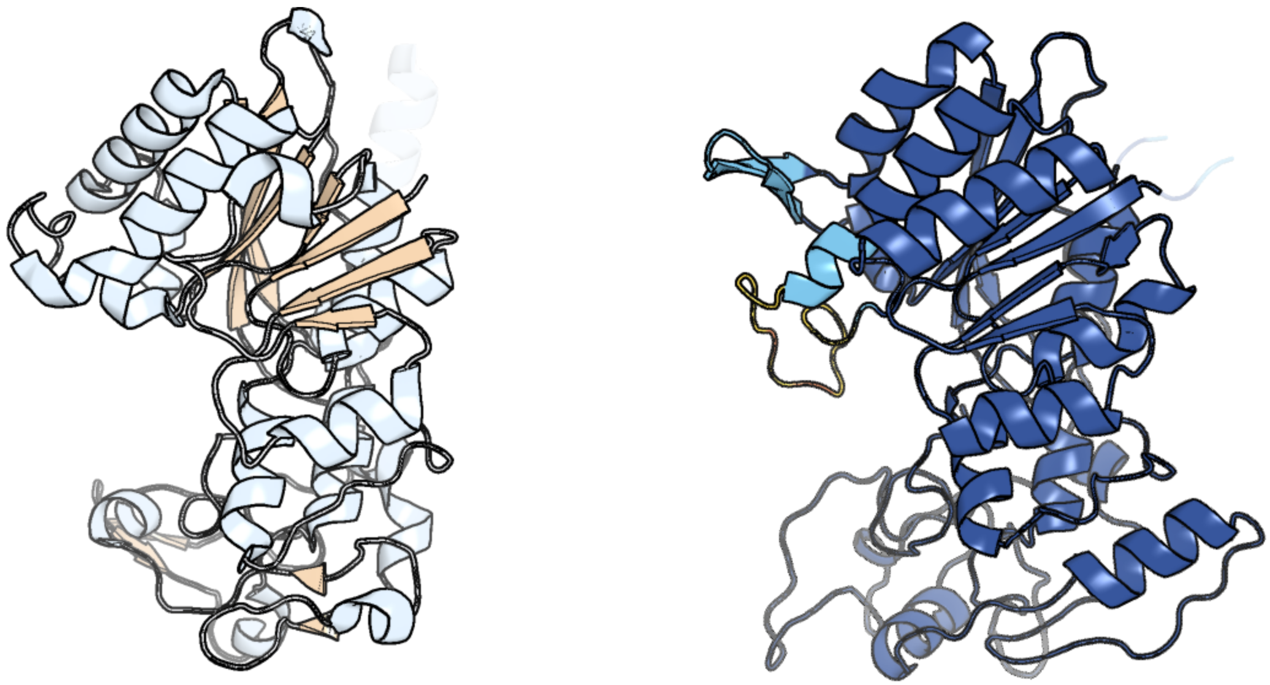

Figure 78: left: reference structure of 3ubt chain B. right: predicted structure of chlorv-1..078, unaligned sequences are shown as transparent

#### chlorv-1..079

- Sequence-based annotation for chlorv-1..079 is hypothetical protein
- Best hit was 2fkc chain B: R.HinP1I restriction endonuclease

| target | prob | fident | alnlen | evalue | theader |
|---|---|---|---|---|---|
| 2fkc-assembly2.cif.gz_B | 1 | 0.141 | 283 | 2.765e-08 | Crystal Form I of Pre-Reactive Complex of Restriction Endonuclease HinP1I with Cognate DNA and Calcium Ion |
| 1yfi-assembly2.cif.gz_B | 1 | 0.11 | 235 | 5.088e-05 | Crystal Structure of restriction endonuclease MspI in complex with its cognate DNA in P212121 space group |
| 4x28-assembly1.cif.gz_B | 0.029 | 0.094 | 85 | 0.6603 | Crystal structure of the ChsE4-ChsE5 complex from Mycobacterium tuberculosis |

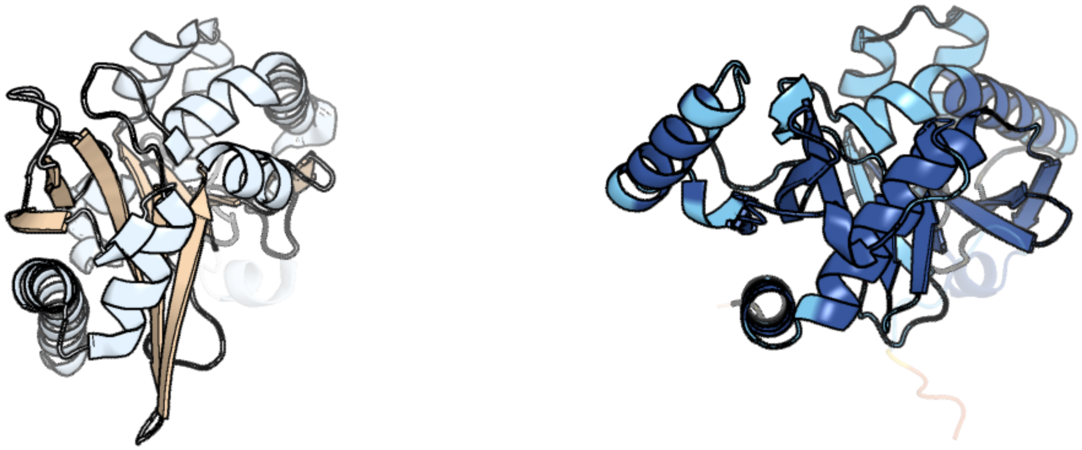

Figure 79: left: reference structure of 2fkc chain B. right: predicted structure of chlorv-1..079, unaligned sequences are shown as transparent

#### chlorv-1..080

- Sequence-based annotation for chlorv-1..080 is putative ATPase family protein/replication factor C small subunit 2
- Best hit was 8dqx chain C: Replication factor C subunit 3

| target | prob | fident | alnlen | evalue | theader |
|---|---|---|---|---|---|
| 8dqx-assembly1.cif.gz_C | 1 | 0.329 | 325 | 2.859e-22 | Open state of RFC:PCNA bound to a 3' ss/dsDNA junction |
| 6vvo-assembly1.cif.gz_C | 1 | 0.33 | 312 | 1.449e-21 | Structure of the human clamp loader (Replication Factor C, RFC) bound to the sliding clamp (Proliferating Cell Nuclear Antigen, PCNA) |
| 1sxj-assembly1.cif.gz_C | 1 | 0.33 | 321 | 5.015e-21 | Crystal Structure of the Eukaryotic Clamp Loader (Replication Factor C, RFC) Bound to the DNA Sliding Clamp (Proliferating Cell Nuclear Antigen, PCNA) |

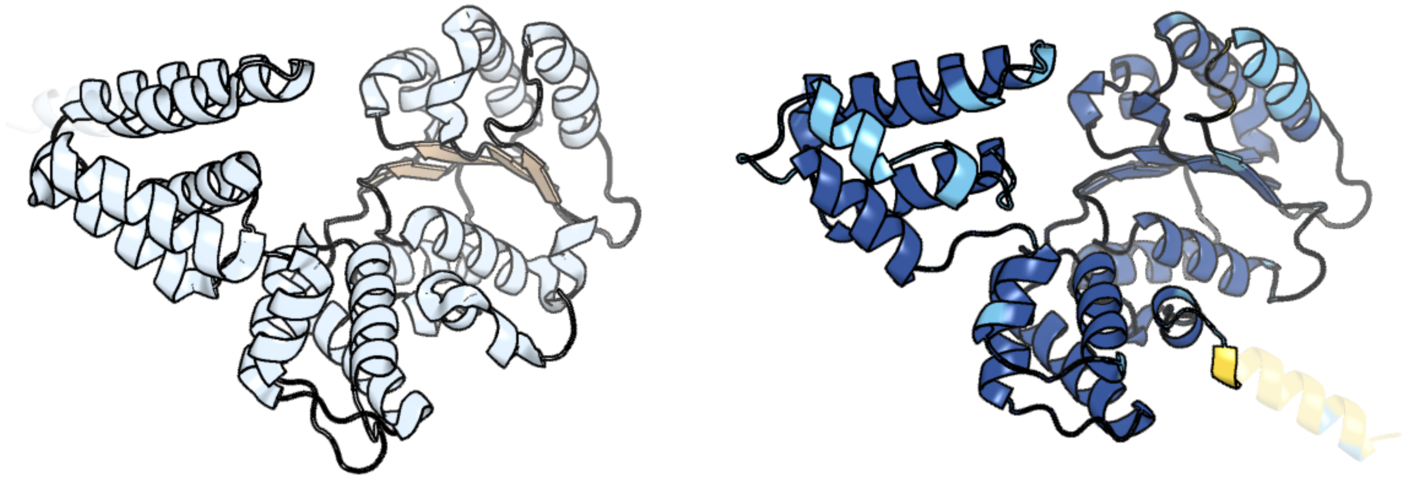

Figure 80: left: reference structure of 8dqx chain C. right: predicted structure of chlorv-1..080, unaligned sequences are shown as transparent

#### chlorv-1..081

- Sequence-based annotation for chlorv-1..081 is putative Nuclease
- Best hit was 4qmg chain C: Staphylococcal nuclease domain-containing protein 1

| target | prob | fident | alnlen | evalue | theader |
|---|---|---|---|---|---|
| 4qmg-assembly3.cif.gz_C | 1 | 0.191 | 115 | 3.611e-06 | The Structure of MTDH-SND1 Complex Reveals Novel Cancer-Promoting Interactions |
| 2f0w-assembly1.cif.gz_A | 1 | 0.26 | 123 | 5.059e-06 | Crystal structure of Staphylococcal nuclease mutant V23I/L25I/V66L/I72L |
| 4qmg-assembly5.cif.gz_E | 1 | 0.189 | 116 | 6.701e-06 | The Structure of MTDH-SND1 Complex Reveals Novel Cancer-Promoting Interactions |

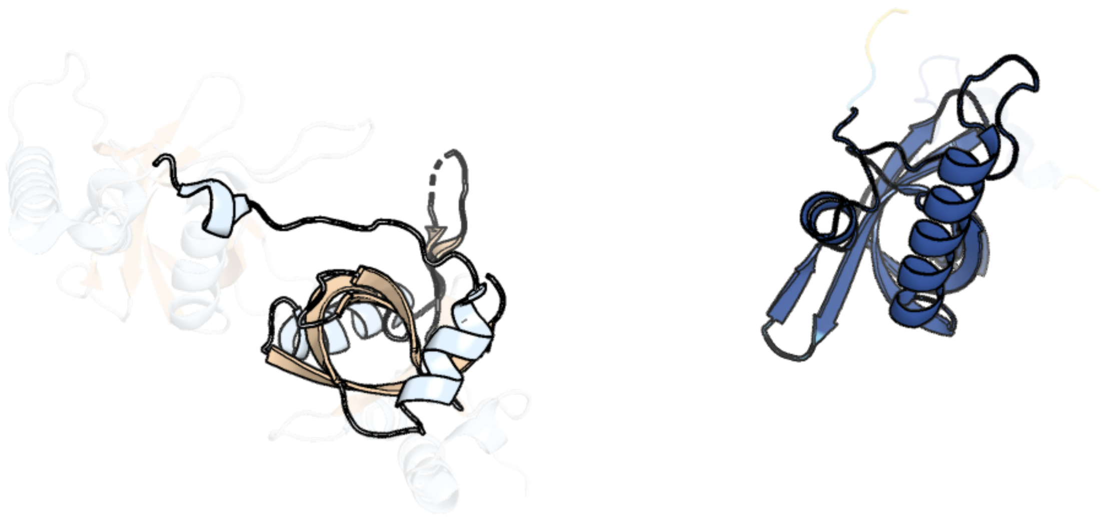

Figure 81: left: reference structure of 4qmg chain C. right: predicted structure of chlorv-1..081, unaligned sequences are shown as transparent

#### chlorv-1..082

- Sequence-based annotation for chlorv-1..082 is putative Methyltransferases
- Best hit was 1j0a chain A: 1-aminocyclopropane-1-carboxylate deaminase

| target | prob | fident | alnlen | evalue | theader |
|---|---|---|---|---|---|
| 1j0a-assembly1.cif.gz_A | 1 | 0.139 | 302 | 5.058e-08 | Crystal Structure Analysis of the ACC deaminase homologue |
| 3iau-assembly1.cif.gz_A | 1 | 0.132 | 279 | 9.878e-08 | The structure of the processed form of threonine deaminase isoform 2 from Solanum lycopersicum |
| 2egu-assembly1.cif.gz_A | 1 | 0.144 | 276 | 1.825e-07 | Crystal structure of O-acetylserine sulfhydrase from Geobacillus kaustophilus HTA426 |

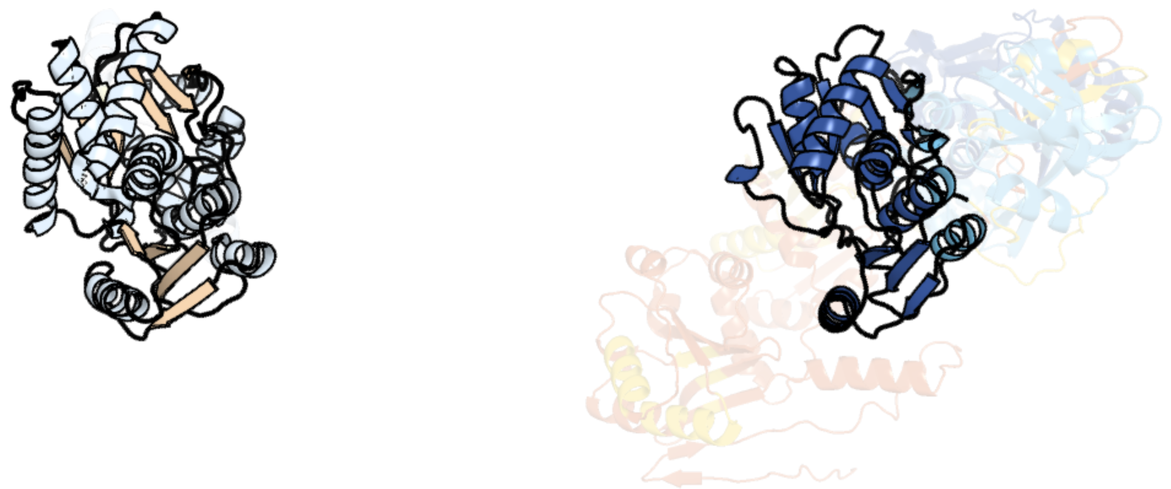

Figure 82: left: reference structure of 1j0a chain A. right: predicted structure of chlorv-1..082, unaligned sequences are shown as transparent

#### chlorv-1..083

- Sequence-based annotation for chlorv-1..083 is putative Thioredoxin
- No significant structural hit found

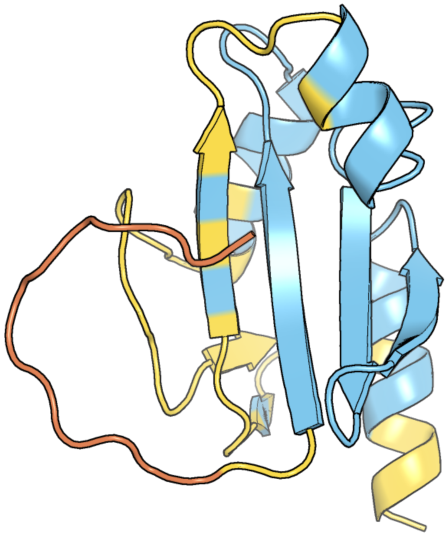

Figure 83: predicted structure of chlorv-1..083

#### chlorv-1..084

- Sequence-based annotation for chlorv-1..084 is putative Thioredoxin
- Best hit was 2vim chain A: THIOREDOXIN

| target | prob | fident | alnlen | evalue | theader |
|---|---|---|---|---|---|
| 2vim-assembly1.cif.gz_A | 1 | 0.152 | 105 | 1.792e-06 | X-ray structure of Fasciola hepatica thioredoxin |
| 2f51-assembly2.cif.gz_B | 1 | 0.205 | 107 | 3.683e-06 | Structure of Trichomonas vaginalis thioredoxin |
| 6z7o-assembly1.cif.gz_A-2 | 1 | 0.23 | 113 | 4.198e-06 | Crystal structure of Thioredoxin T from Drosophila melanogaster |

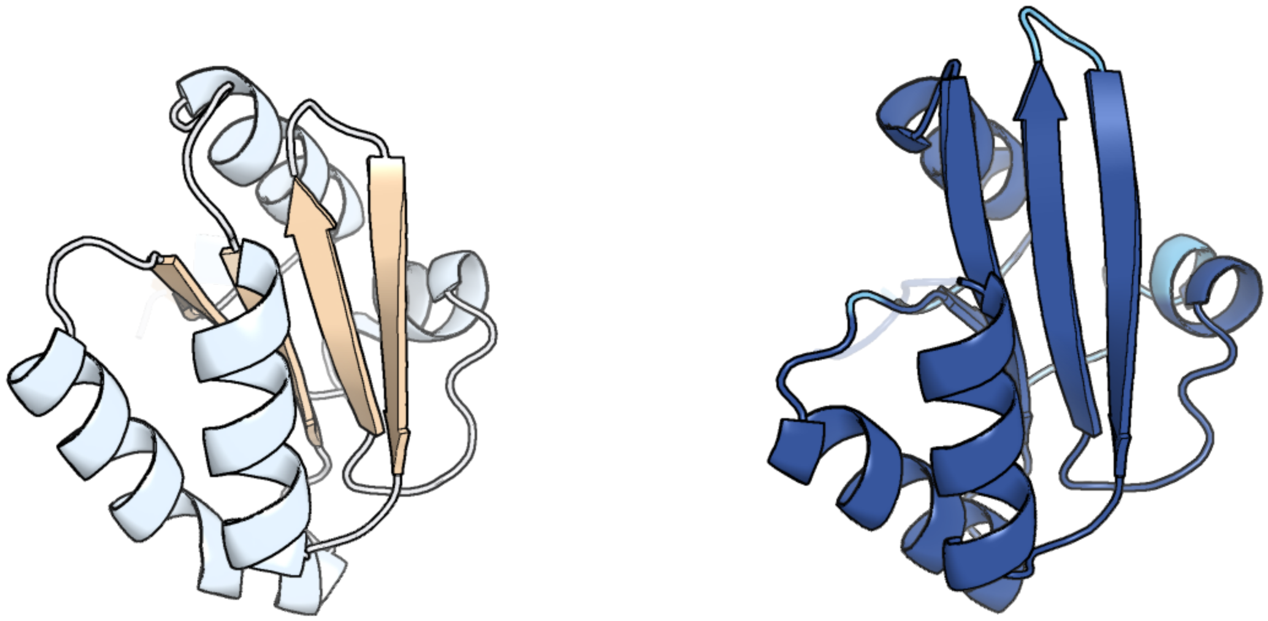

Figure 84: left: reference structure of 2vim chain A. right: predicted structure of chlorv-1..084, unaligned sequences are shown as transparent

#### chlorv-1..085

- Sequence-based annotation for chlorv-1..085 is hypothetical protein
- No significant structural hit found

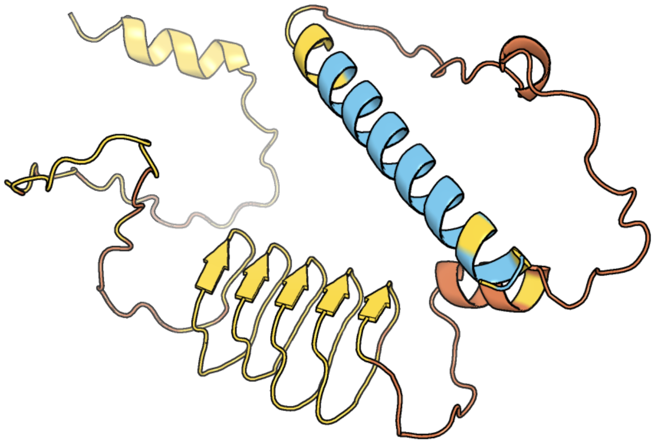

Figure 85: predicted structure of chlorv-1..085

### chlorv-1..086

- Sequence-based annotation for chlorv-1..086 is putative Peptidase
- Best hit was 6lj9 chain A: Cysteine protease S273R

| target | prob | fident | alnlen | evalue | theader |
|---|---|---|---|---|---|
| 6lj9-assembly1.cif.gz_A | 1 | 0.24 | 250 | 6.465e-14 | Crystal Structure of Se-Met ASFV pS273R protease |
| 6ljb-assembly1.cif.gz_A | 1 | 0.226 | 256 | 1.219e-13 | Crystal Structure of ASFV pS273R protease |
| 6lj9-assembly1.cif.gz_B | 1 | 0.222 | 252 | 8.748e-11 | Crystal Structure of Se-Met ASFV pS273R protease |

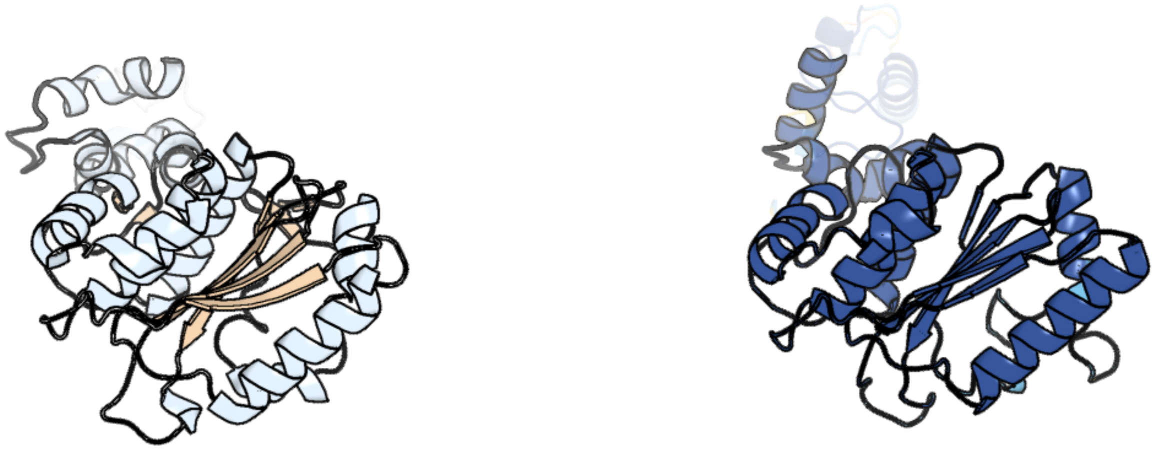

Figure 86: left: reference structure of 6lj9 chain A. right: predicted structure of chlorv-1..086, unaligned sequences are shown as transparent

#### chlorv-1..087

- Sequence-based annotation for chlorv-1..087 is hypothetical protein
- Best hit was 8h2i chain bG: P1v1

| target | prob | fident | alnlen | evalue | theader |
|---|---|---|---|---|---|
| 8h2i-assembly1.cif.gz_bG | 1 | 0.089 | 460 | 7.516e-06 | Near-atomic structure of five-fold averaged PBCV-1 capsid |
| 8rbs-assembly1.cif.gz_K | 0.993 | 0.111 | 268 | 0.0002393 | Emiliania huxleyi virus 201 (EhV-201) asymmetrical unit of capsid proteins predicted by AlphaFold2 fitted into the cryo-EM density of EhV-201 virion composite map. |
| 6yba-assembly1.cif.gz_M | 0.992 | 0.075 | 398 | 0.001474 | HAdV-F41 Capsid |

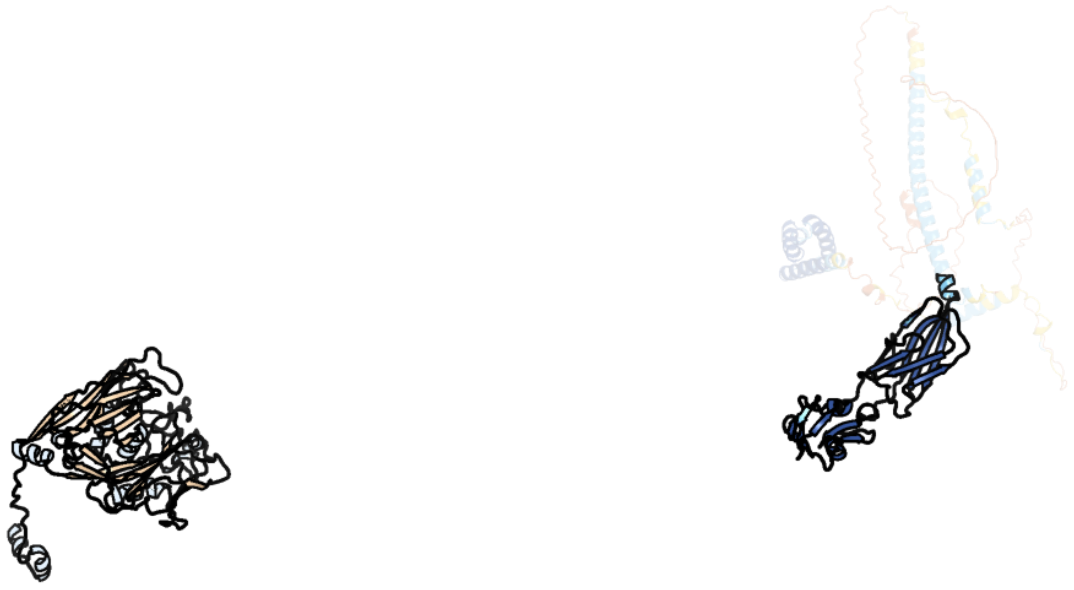

Figure 87: left: reference structure of 8h2i chain bG. right: predicted structure of chlorv-1..087, unaligned sequences are shown as transparent

#### chlorv-1..088

- Sequence-based annotation for chlorv-1..088 is hypothetical protein
- No significant structural hit found

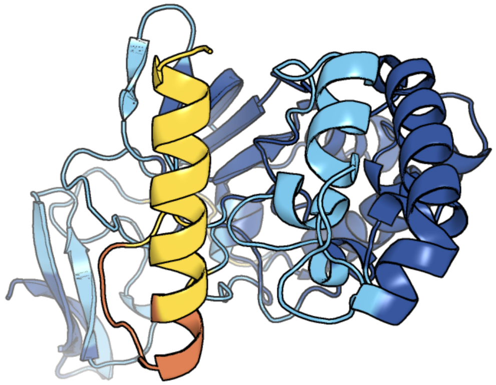

Figure 88: predicted structure of chlorv-1..088

### chlorv-1..089

- Sequence-based annotation for chlorv-1..089 is hypothetical protein
- No significant structural hit found

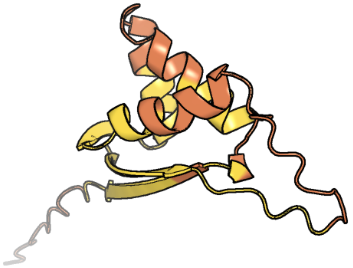

Figure 89: predicted structure of chlorv-1..089

#### chlorv-1..090

- Sequence-based annotation for chlorv-1..090 is putative GTP binding protein
- Best hit was 5wfs chain z: Elongation factor Tu 2

| target | prob | fident | alnlen | evalue | theader |
|---|---|---|---|---|---|
| 5wfs-assembly1.cif.gz_z | 1 | 0.11 | 398 | 1.693e-15 | 70S ribosome-EF-Tu H84A complex with GTP and near-cognate tRNA (Complex C4) |
| 5we4-assembly1.cif.gz_z | 1 | 0.105 | 399 | 1.897e-15 | 70S ribosome-EF-Tu wt complex with GppNHp |
| 4cxg-assembly1.cif.gz_A | 1 | 0.107 | 428 | 6.285e-15 | Regulation of the mammalian elongation cycle by 40S subunit rolling: a eukaryotic-specific ribosome rearrangement |

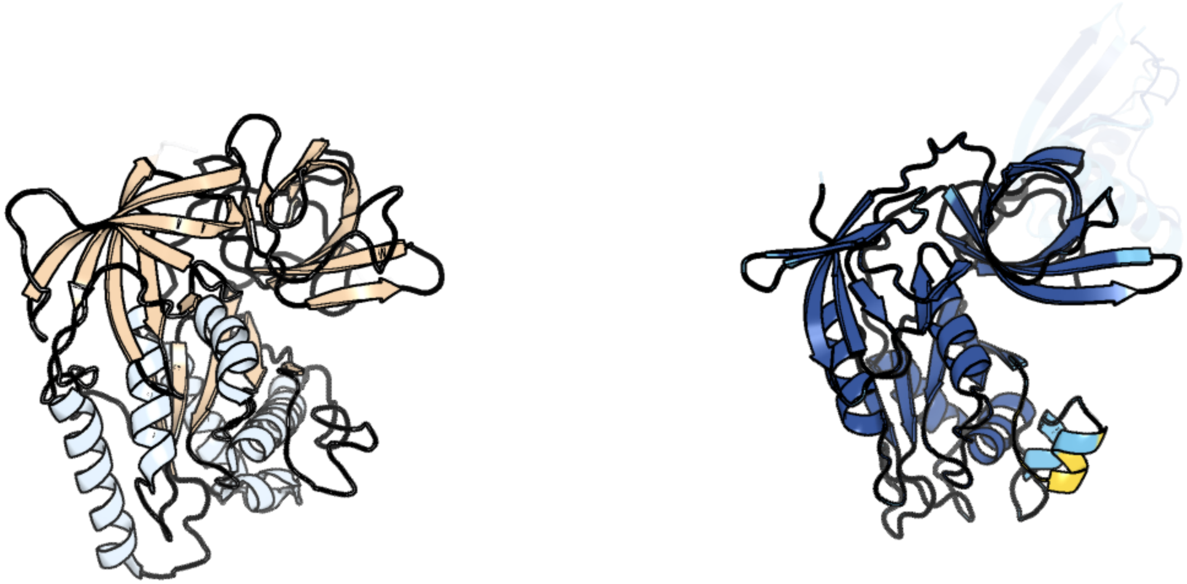

Figure 90: left: reference structure of 5wfs chain z. right: predicted structure of chlorv-1..090, unaligned sequences are shown as transparent

#### chlorv-1..091

- Sequence-based annotation for chlorv-1..091 is hypothetical protein
- No significant structural hit found

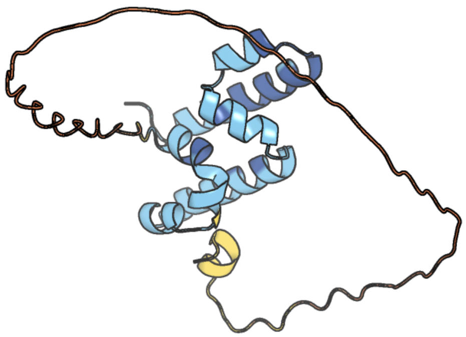

Figure 91: predicted structure of chlorv-1..091

#### chlorv-1..092

- Sequence-based annotation for chlorv-1..092 is hypothetical protein
- No significant structural hit found

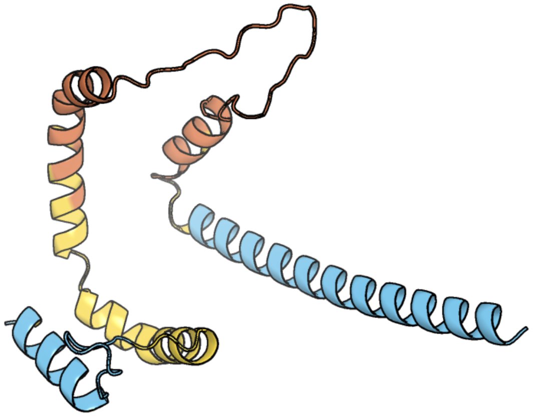

Figure 92: predicted structure of chlorv-1..092

#### chlorv-1..093

- Sequence-based annotation for chlorv-1..093 is hypothetical protein
- No significant structural hit found

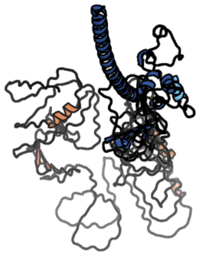

Figure 93: predicted structure of chlorv-1..093

#### chlorv-1..094

- Sequence-based annotation for chlorv-1..094 is putative Ankyrin repeat protein
- Best hit was 4hqd chain B: Engineered Protein OR265

| target | prob | fident | alnlen | evalue | theader |
|---|---|---|---|---|---|
| 4hqd-assembly2.cif.gz_B | 1 | 0.269 | 115 | 9.919e-06 | Crystal Structure of Engineered Protein. Northeast Structural Genomics Consortium Target OR265. |
| 7ujj-assembly1.cif.gz_G | 1 | 0.238 | 113 | 9.919e-06 | Stx2a and DARPin complex |
| 2xzt-assembly2.cif.gz_G | 1 | 0.23 | 117 | 1.277e-05 | Caspase-3 in Complex with DARPin-3.4_I78S |

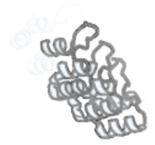

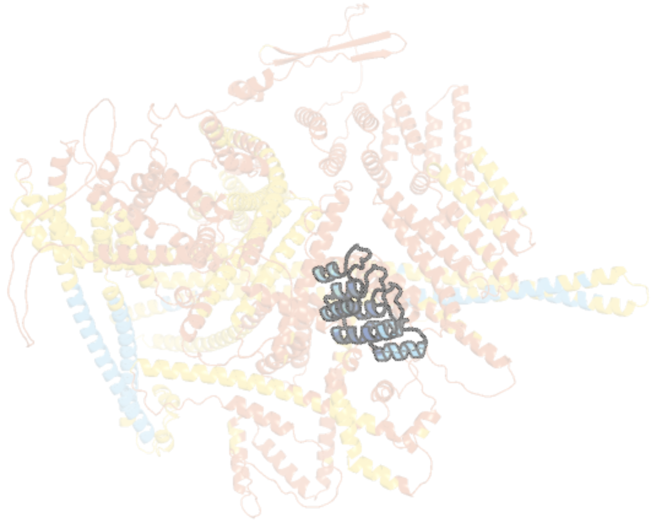

Figure 94: left: reference structure of 4hqd chain B. right: predicted structure of chlorv-1..094, unaligned sequences are shown as transparent

#### chlorv-1..095

- Sequence-based annotation for chlorv-1..095 is hypothetical protein
- Best hit was 8rbs chain K: Penton protein

| target | prob | fident | alnlen | evalue | theader |
|---|---|---|---|---|---|
| 8rbs-assembly1.cif.gz_K | 1 | 0.103 | 799 | 1.186e-10 | Emiliania huxleyi virus 201 (EhV-201) asymmetrical unit of capsid proteins predicted by AlphaFold2 fitted into the cryo-EM density of EhV-201 virion composite map. |
| 8h2i-assembly1.cif.gz_bG | 1 | 0.135 | 503 | 3.282e-08 | Near-atomic structure of five-fold averaged PBCV-1 capsid |
| 6g41-assembly1.cif.gz_B | 1 | 0.103 | 396 | 2.489e-05 | Crystal structure of SeMet-labeled mavirus penton protein |

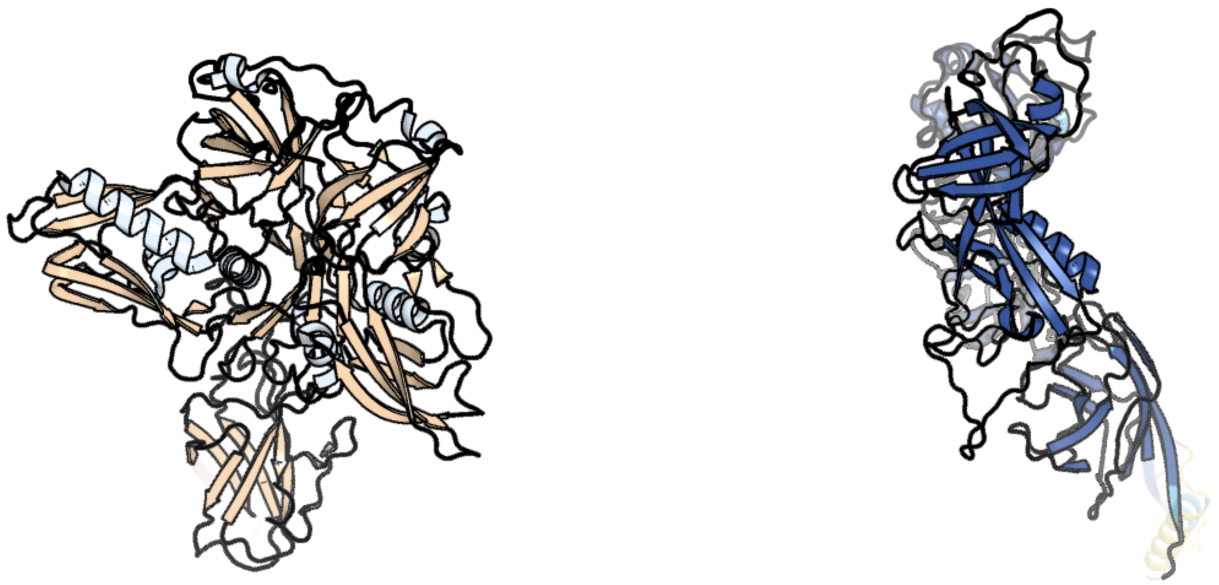

Figure 95: left: reference structure of 8rbs chain K. right: predicted structure of chlorv-1..095, unaligned sequences are shown as transparent

#### chlorv-1..096

- Sequence-based annotation for chlorv-1..096 is hypothetical protein
- No significant structural hit found

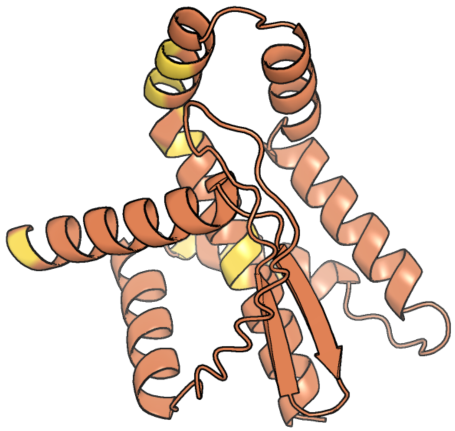

Figure 96: predicted structure of chlorv-1..096

#### chlorv-1..097

- Sequence-based annotation for chlorv-1..097 is hypothetical protein
- No significant structural hit found

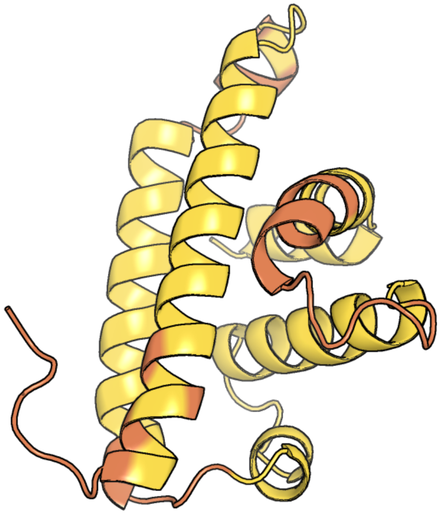

Figure 97: predicted structure of chlorv-1..097

#### chlorv-1..098

- Sequence-based annotation for chlorv-1..098 is hypothetical protein
- No significant structural hit found

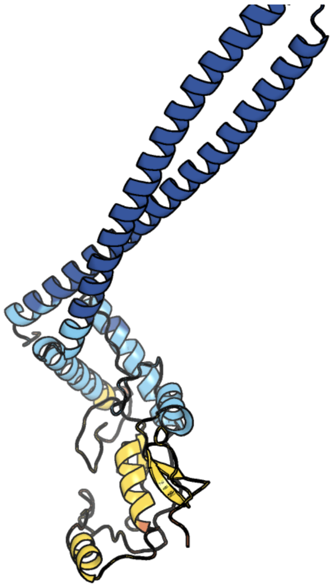

Figure 98: predicted structure of chlorv-1..098

#### chlorv-1..099

- Sequence-based annotation for chlorv-1..099 is putative Ceramidase
- Best hit was 6yxh chain A: Alkaline ceramidase 3

| target | prob | fident | alnlen | evalue | theader |
|---|---|---|---|---|---|
| 6yxh-assembly1.cif.gz_A | 1 | 0.2 | 230 | 3.071e-07 | Cryogenic human alkaline ceramidase 3 (ACER3) at 2.6 A resolution determined by Serial Crystallography (SSX) using CrystalDirect |
| 7y69-assembly1.cif.gz_B | 1 | 0.134 | 291 | 0.02441 | ApoSIDT2-pH5.5 |
| 7y68-assembly1.cif.gz_B | 0.988 | 0.119 | 293 | 0.06607 | SIDT2-pH5.5 plus miRNA |

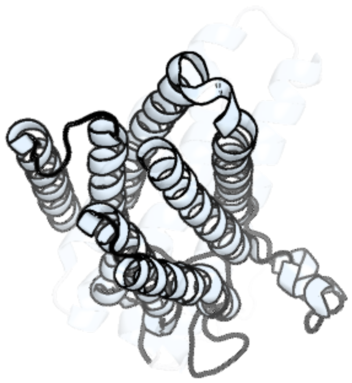

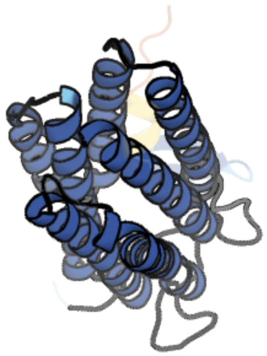

Figure 99: left: reference structure of 6yxh chain A. right: predicted structure of chlorv-1..099, unaligned sequences are shown as transparent

#### chlorv-1..100

- Sequence-based annotation for chlorv-1..100 is hypothetical protein
- No significant structural hit found

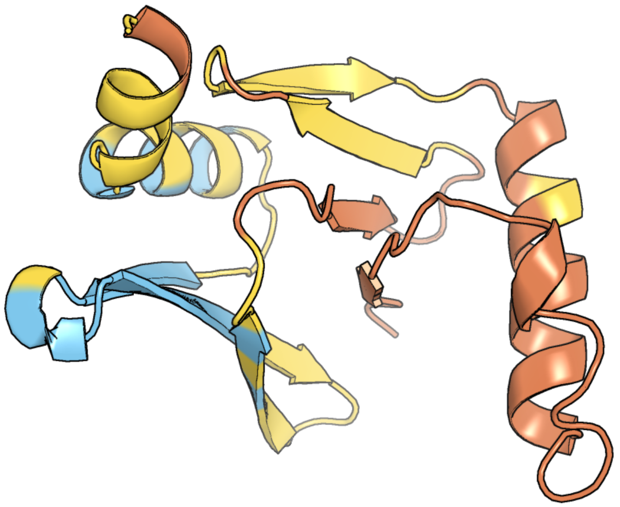

Figure 100: predicted structure of chlorv-1..100

### chlorv-1..101

- Sequence-based annotation for chlorv-1..101 is hypothetical protein
- No significant structural hit found

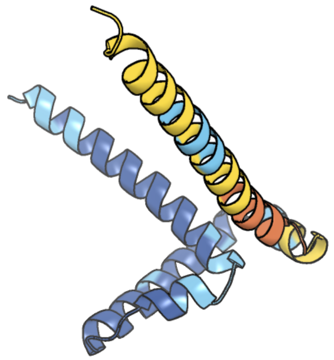

Figure 101: predicted structure of chlorv-1..101

#### chlorv-1..102

- Sequence-based annotation for chlorv-1..102 is hypothetical protein
- No significant structural hit found

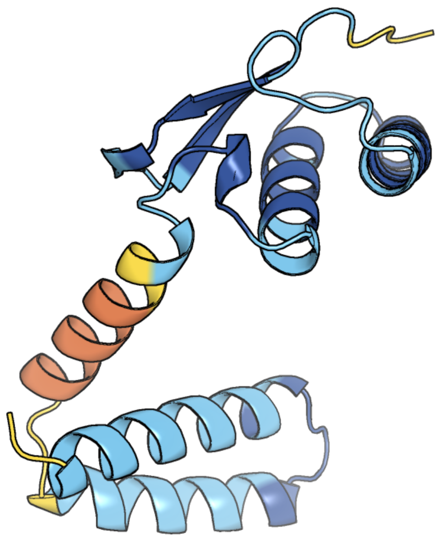

Figure 102: predicted structure of chlorv-1..102

#### chlorv-1..103

- Sequence-based annotation for chlorv-1..103 is putative Ribonuclease III
- Best hit was 1o0w chain A: Ribonuclease III

| target | prob | fident | alnlen | evalue | theader |
|---|---|---|---|---|---|
| 1o0w-assembly1.cif.gz_A | 1 | 0.239 | 271 | 4.342e-13 | Crystal structure of Ribonuclease III (TM1102) from Thermotoga maritima at 2.0 A resolution |
| 7r97-assembly1.cif.gz_B | 1 | 0.213 | 262 | 2.069e-12 | Crystal structure of postcleavge complex of Escherichia coli RNase III |
| 1rc7-assembly1.cif.gz_A | 1 | 0.217 | 258 | 1.163e-10 | Crystal structure of RNase III Mutant E110K from Aquifex Aeolicus complexed with ds-RNA at 2.15 Angstrom Resolution |

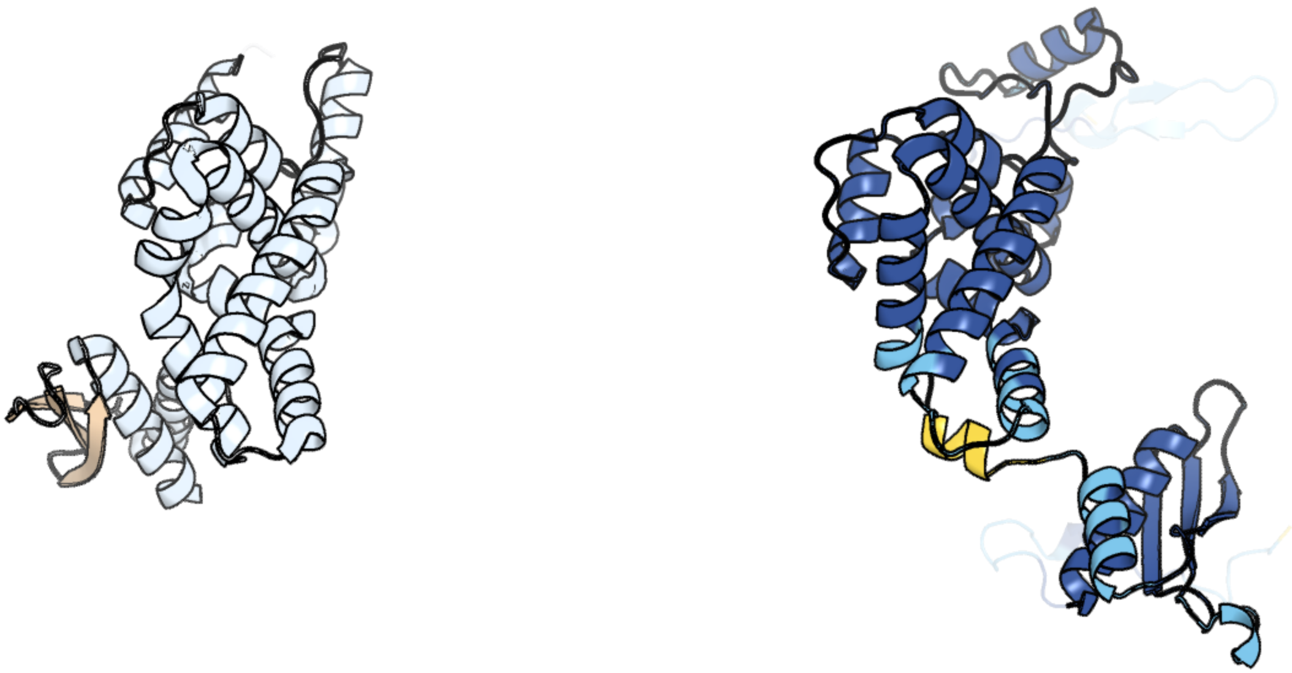

Figure 103: left: reference structure of 1o0w chain A. right: predicted structure of chlorv-1..103, unaligned sequences are shown as transparent

### chlorv-1..104

- Sequence-based annotation for chlorv-1..104 is putative Nuclease
- Best hit was 3q8l chain A: Flap endonuclease 1

| target | prob | fident | alnlen | evalue | theader |
|---|---|---|---|---|---|
| 3q8l-assembly1.cif.gz_A | 1 | 0.302 | 340 | 4.379e-25 | Crystal Structure of Human Flap Endonuclease FEN1 (WT) in complex with substrate 5'-flap DNA, SM3+, and K+ |
| 5um9-assembly1.cif.gz_A | 1 | 0.294 | 340 | 7.788e-25 | Flap endonuclease 1 (FEN1) D86N with 5'-flap substrate DNA and Sm3+ |
| 5k97-assembly1.cif.gz_A | 1 | 0.292 | 339 | 9.602e-25 | Flap endonuclease 1 (FEN1) D233N with cleaved product fragment and Sm3+ |

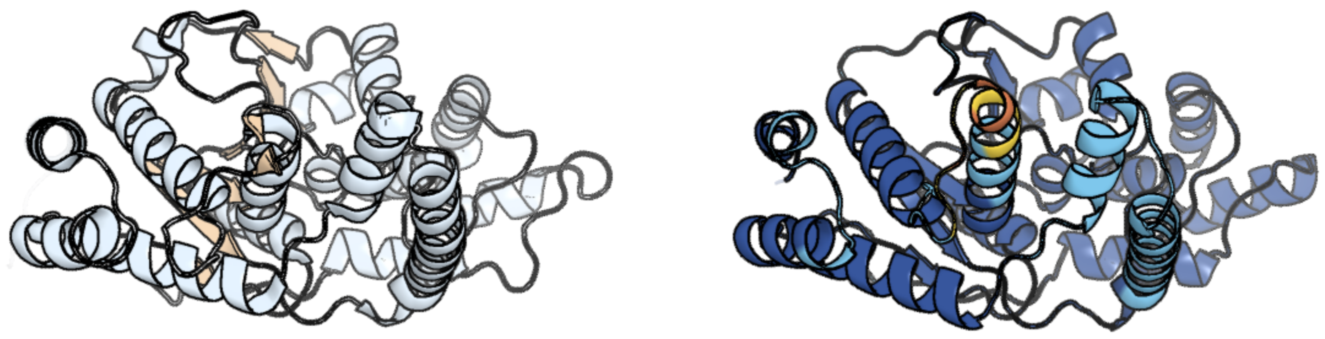

Figure 104: left: reference structure of 3q8l chain A. right: predicted structure of chlorv-1..104, unaligned sequences are shown as transparent

#### chlorv-1..105

- Sequence-based annotation for chlorv-1..105 is putative Nuclease
- Best hit was 3pif chain B: 5'->3' EXORIBONUCLEASE (xrn1)

| target | prob | fident | alnlen | evalue | theader |
|---|---|---|---|---|---|
| 3pif-assembly2.cif.gz_B | 1 | 0.187 | 681 | 6.256e-14 | Crystal structure of the 5'->3' exoribonuclease Xrn1, E178Q mutant in Complex with Manganese |
| 3fqd-assembly1.cif.gz_A | 1 | 0.151 | 711 | 6.54e-14 | Crystal Structure of the S. pombe Rat1-Rai1 Complex |
| 7opk-assembly1.cif.gz_A | 1 | 0.161 | 669 | 2.17e-13 | Crystal structure of C. thermophilum Xrn2 |

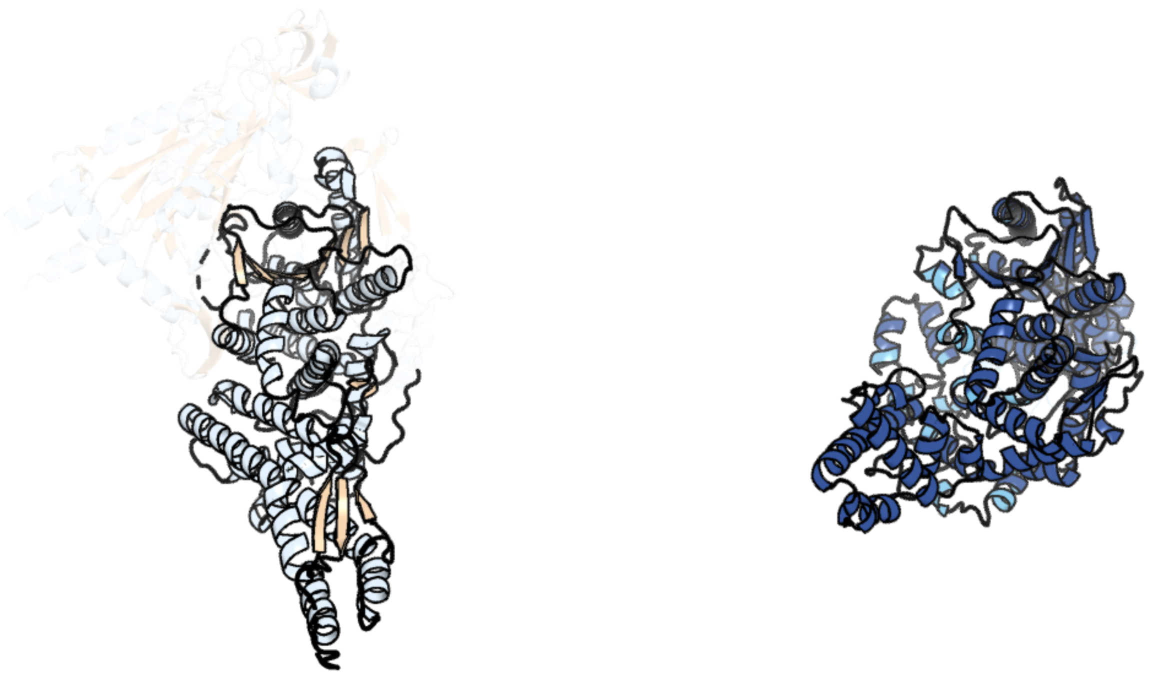

Figure 105: left: reference structure of 3pif chain B. right: predicted structure of chlorv-1..105, unaligned sequences are shown as transparent

### chlorv-1..106

- Sequence-based annotation for chlorv-1..106 is putative Peptidase
- Best hit was 6on2 chain C: ATP-dependent protease La

| target | prob | fident | alnlen | evalue | theader |
|---|---|---|---|---|---|
| 6on2-assembly1.cif.gz_C | 1 | 0.288 | 540 | 1.273e-38 | Lon Protease from Yersinia pestis with Y2853 substrate |
| 7p0m-assembly1.cif.gz_B | 1 | 0.27 | 554 | 1.805e-38 | Human mitochondrial Lon protease with substrate in the ATPase and protease domains |
| 7krz-assembly1.cif.gz_D | 1 | 0.271 | 557 | 1.994e-38 | Human mitochondrial LONP1 in complex with Bortezomib |

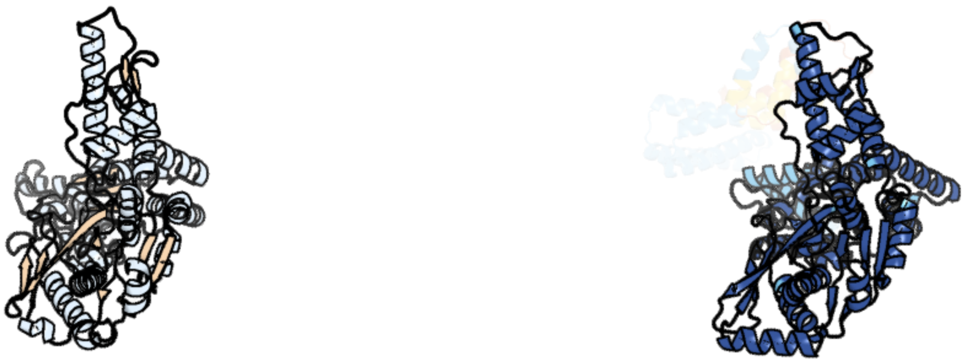

Figure 106: left: reference structure of 6on2 chain C. right: predicted structure of chlorv-1..106, unaligned sequences are shown as transparent

#### chlorv-1..107

- Sequence-based annotation for chlorv-1..107 is putative DNA ligase
- Best hit was 5tt5 chain A: DNA ligase

| target | prob | fident | alnlen | evalue | theader |
|---|---|---|---|---|---|
| 5tt5-assembly1.cif.gz_A | 1 | 0.176 | 605 | 1.772e-30 | Escherichia coli LigA (K115M) in complex with NAD+ |
| 2owo-assembly1.cif.gz_A | 1 | 0.178 | 609 | 5.476e-30 | Last Stop on the Road to Repair: Structure of E.coli DNA Ligase Bound to Nicked DNA-Adenylate |
| 4glx-assembly1.cif.gz_A | 1 | 0.177 | 597 | 8.881e-30 | DNA ligase A in complex with inhibitor |

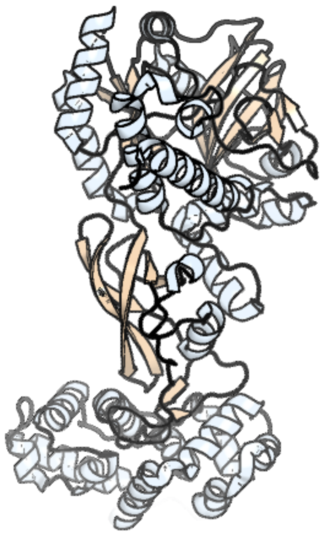

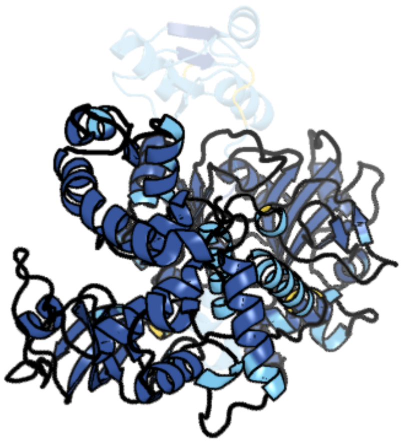

Figure 107: left: reference structure of 5tt5 chain A. right: predicted structure of chlorv-1..107, unaligned sequences are shown as transparent

#### chlorv-1..108

- Sequence-based annotation for chlorv-1..108 is putative Cytidine deaminase
- Best hit was 7zob chain E: Metagenomic cytidine deaminase Cdd

| target | prob | fident | alnlen | evalue | theader |
|---|---|---|---|---|---|
| 7zob-assembly2.cif.gz_E | 1 | 0.409 | 127 | 2.885e-15 | Metagenomic cytidine deaminase Cdd |
| 1r5t-assembly1.cif.gz_D | 1 | 0.333 | 132 | 2.791e-14 | The Crystal Structure of Cytidine Deaminase CDD1, an Orphan C to U editase from Yeast |
| 3dmo-assembly1.cif.gz_C | 1 | 0.33 | 127 | 4.394e-14 | 1.6 A crystal structure of cytidine deaminase from Burkholderia pseudomallei |

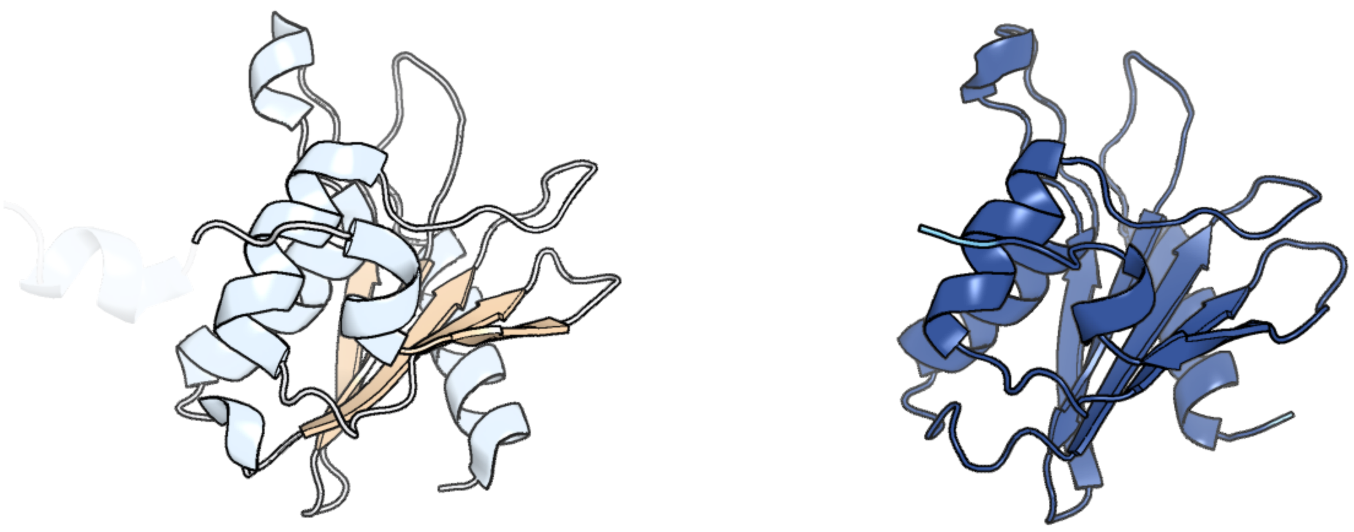

Figure 108: left: reference structure of 7zob chain E. right: predicted structure of chlorv-1..108, unaligned sequences are shown as transparent

#### chlorv-1..109

- Sequence-based annotation for chlorv-1..109 is hypothetical protein
- No significant structural hit found

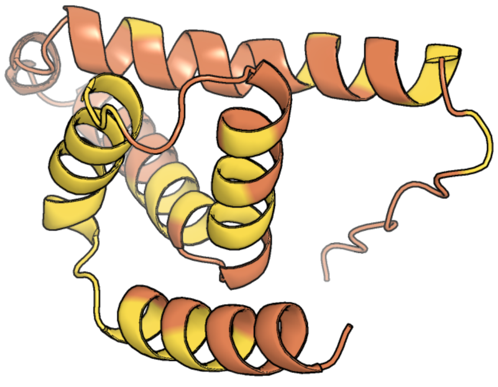

Figure 109: predicted structure of chlorv-1..109

#### chlorv-1..110

- Sequence-based annotation for chlorv-1..110 is putative RNA ligase
- Best hit was 2c5u chain B: RNA LIGASE

| target | prob | fident | alnlen | evalue | theader |
|---|---|---|---|---|---|
| 2c5u-assembly2.cif.gz_B | 1 | 0.148 | 405 | 2.005e-10 | T4 RNA Ligase (Rnl1) Crystal Structure |
| 5tt6-assembly1.cif.gz_A | 1 | 0.154 | 448 | 3.89e-09 | T4 RNA Ligase 1 (K99M) |
| 6n67-assembly1.cif.gz_A | 1 | 0.109 | 422 | 1.184e-06 | Crystal structure of the ligase domain of fungal tRNA ligase Trl1 |

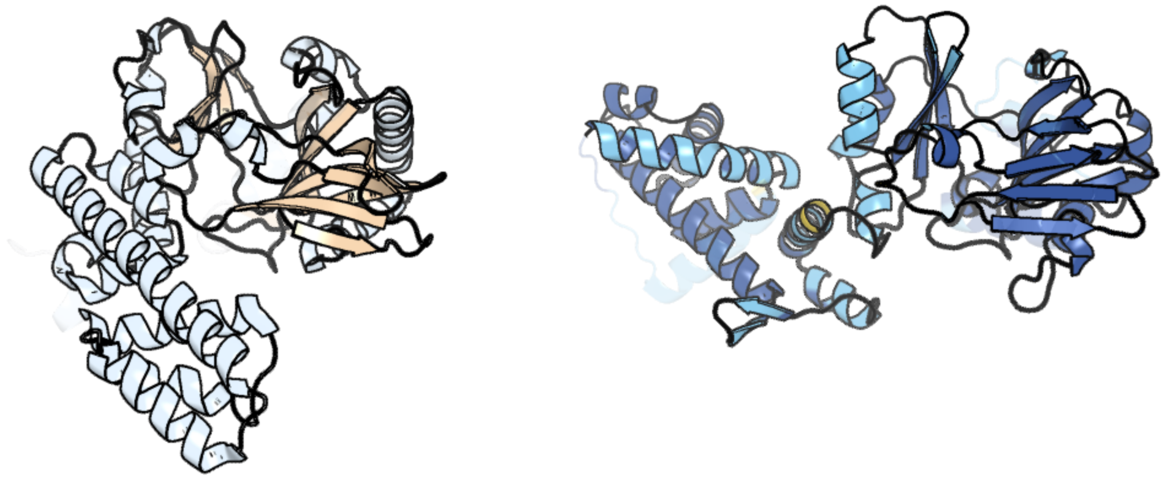

Figure 110: left: reference structure of 2c5u chain B. right: predicted structure of chlorv-1..110, unaligned sequences are shown as transparent

#### chlorv-1..111

- Sequence-based annotation for chlorv-1..111 is putative Phosphatase
- Best hit was 2iq1 chain A: Protein phosphatase 2C kappa, PPM1K

| target | prob | fident | alnlen | evalue | theader |
|---|---|---|---|---|---|
| 2iq1-assembly1.cif.gz_A | 1 | 0.191 | 324 | 3.039e-19 | Crystal structure of human PPM1K |
| 6ak7-assembly1.cif.gz_A | 1 | 0.19 | 320 | 5.325e-18 | Crystal structure of PPM1K-N94K |
| 4da1-assembly1.cif.gz_A | 1 | 0.19 | 326 | 2.511e-17 | Crystal structure of branched-chain alpha-ketoacid dehydrogenase phosphatase with Mg (II) ions at the active site |

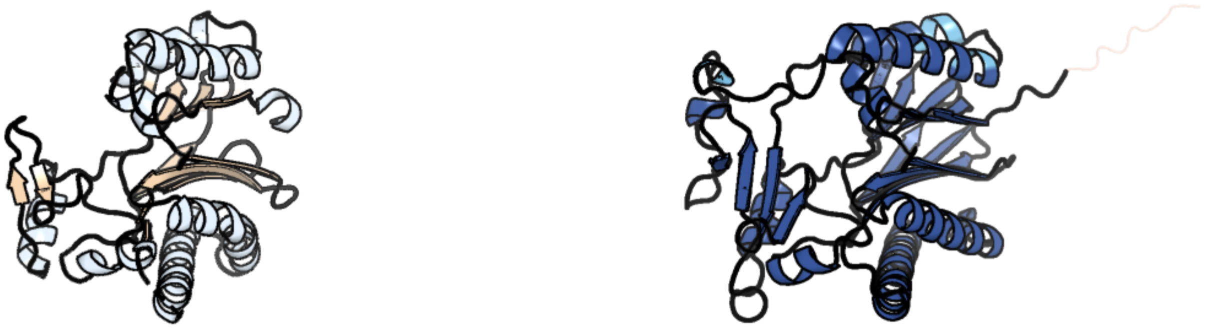

Figure 111: left: reference structure of 2iq1 chain A. right: predicted structure of chlorv-1..111, unaligned sequences are shown as transparent

### chlorv-1..112

- Sequence-based annotation for chlorv-1..112 is hypothetical protein
- No significant structural hit found

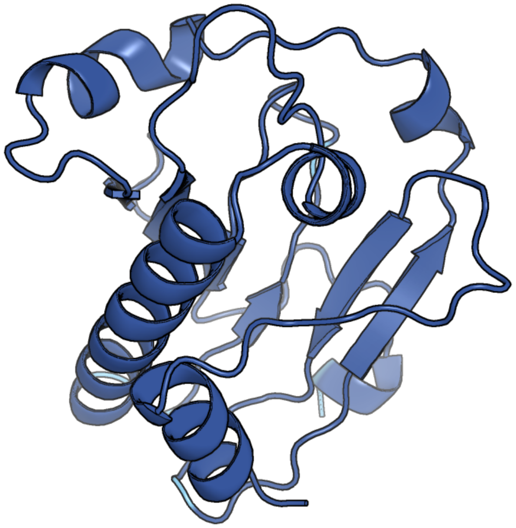

Figure 112: predicted structure of chlorv-1..112

### chlorv-1..113

- Sequence-based annotation for chlorv-1..113 is putative translation initiation factor eIF2 gamma subunit
- Best hit was 6r8s chain A: Translation initiation factor 2 subunit gamma

| target | prob | fident | alnlen | evalue | theader |
|---|---|---|---|---|---|
| 6r8s-assembly1.cif.gz_A | 1 | 0.33 | 411 | 4.892e-46 | Crystal structure of aIF2gamma subunit I181K from archaeon Sulfolobus solfataricus complexed with GDPCP |
| 4rd0-assembly1.cif.gz_A | 1 | 0.327 | 409 | 1.474e-45 | Structure of aIF2-gamma D19A variant from Sulfolobus solfataricus bound to GDP |
| 4qfm-assembly1.cif.gz_A | 1 | 0.33 | 411 | 1.971e-45 | The structure of aIF2gamma subunit D152A from archaeon Sulfolobus solfataricus complexed with GDPCP |

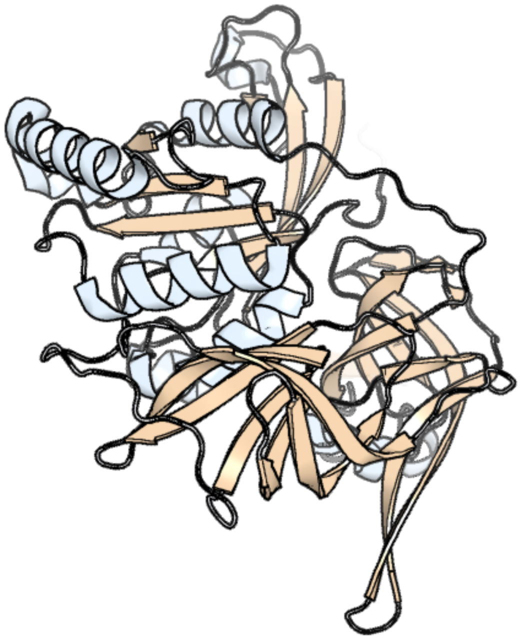

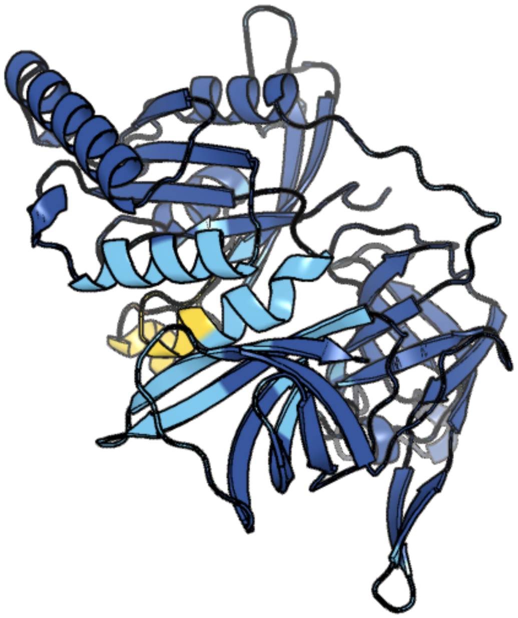

Figure 113: left: reference structure of 6r8s chain A. right: predicted structure of chlorv-1..113, unaligned sequences are shown as transparent

#### chlorv-1..114

- Sequence-based annotation for chlorv-1..114 is hypothetical protein
- No significant structural hit found

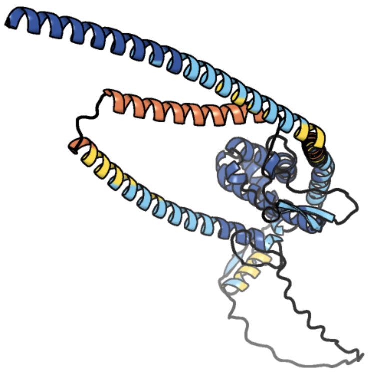

Figure 114: predicted structure of chlorv-1..114

#### chlorv-1..115

- Sequence-based annotation for chlorv-1..115 is putative DUF285 domain-containing protein
- No significant structural hit found

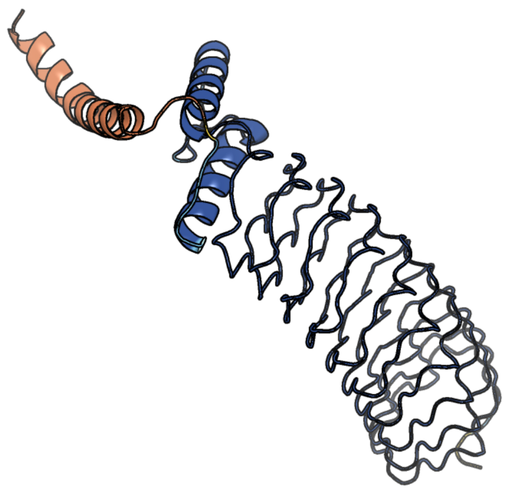

Figure 115: predicted structure of chlorv-1..115

#### chlorv-1..116

- Sequence-based annotation for chlorv-1..116 is putative DUF285 domain-containing protein
- No significant structural hit found

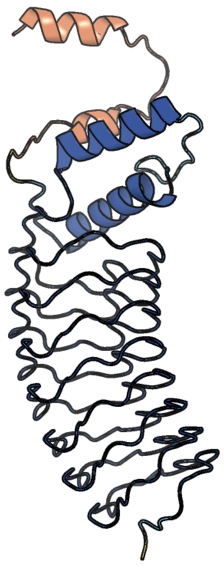

Figure 116: predicted structure of chlorv-1..116

#### chlorv-1..117

- Sequence-based annotation for chlorv-1..117 is hypothetical protein
- No significant structural hit found

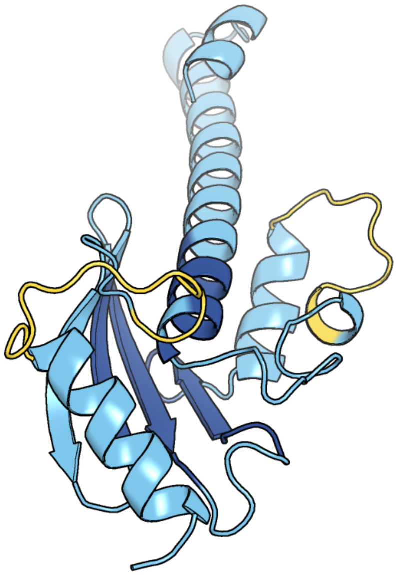

Figure 117: predicted structure of chlorv-1..117

#### chlorv-1..118

- Sequence-based annotation for chlorv-1..118 is putative IMP dehydrogenase / GMP reductase
- Best hit was 6jig chain A: GMP reductase

| target | prob | fident | alnlen | evalue | theader |
|---|---|---|---|---|---|
| 6jig-assembly1.cif.gz_A | 1 | 0.398 | 487 | 2.217e-63 | Crystal structure of GMP reductase C318A from Trypanosoma brucei in complex with guanosine 5'-monophosphate |
| 3tsb-assembly1.cif.gz_A | 1 | 0.354 | 483 | 4.499e-57 | Crystal Structure of Inosine-5'-monophosphate Dehydrogenase from Bacillus anthracis str. Ames |
| 6jl8-assembly1.cif.gz_A | 1 | 0.386 | 481 | 1.008e-56 | Crystal structure of GMP reductase C318A from Trypanosoma brucei |

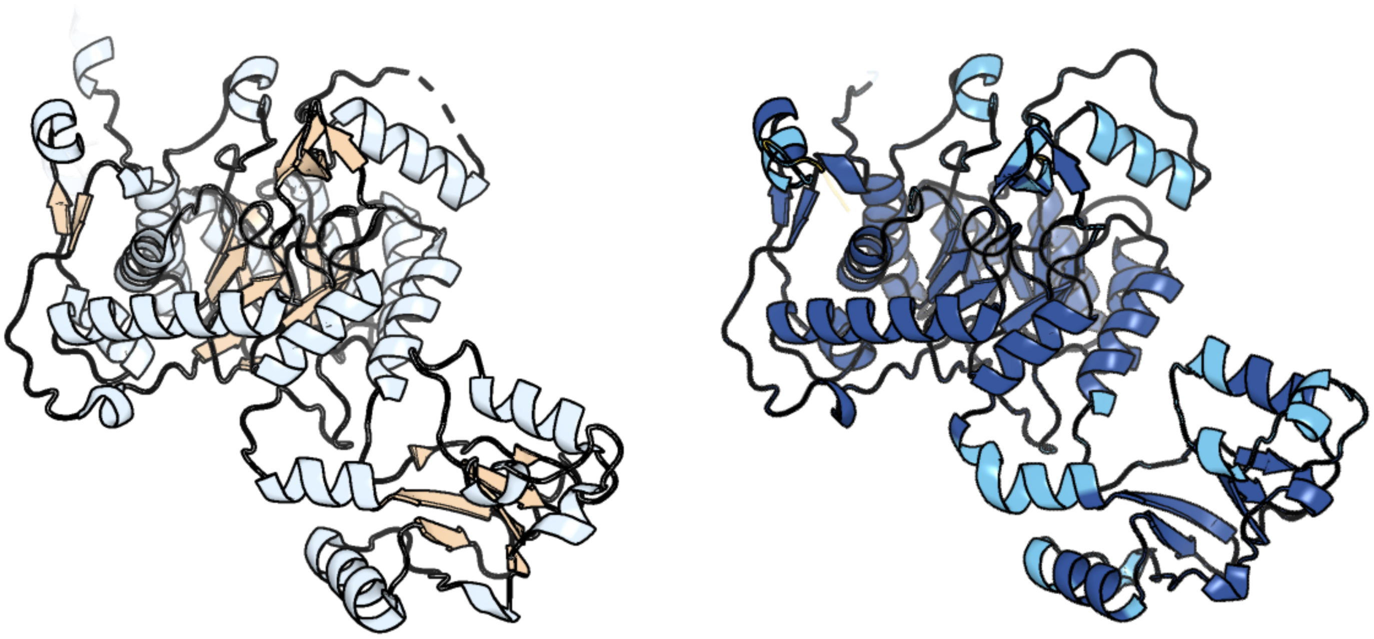

Figure 118: left: reference structure of 6jig chain A. right: predicted structure of chlorv-1..118, unaligned sequences are shown as transparent

#### chlorv-1..119

- Sequence-based annotation for chlorv-1..119 is putative Peptidase
- Best hit was 6on2 chain C: ATP-dependent protease La

| target | prob | fident | alnlen | evalue | theader |
|---|---|---|---|---|---|
| 7p0m-assembly1.cif.gz_B | 1 | 0.263 | 619 | 9.47e-37 | Human mitochondrial Lon protease with substrate in the ATPase and protease domains |
| 6on2-assembly1.cif.gz_C | 1 | 0.276 | 597 | 1.518e-36 | Lon Protease from Yersinia pestis with Y2853 substrate |
| 7p0m-assembly1.cif.gz_A | 1 | 0.271 | 594 | 1.834e-36 | Human mitochondrial Lon protease with substrate in the ATPase and protease domains |

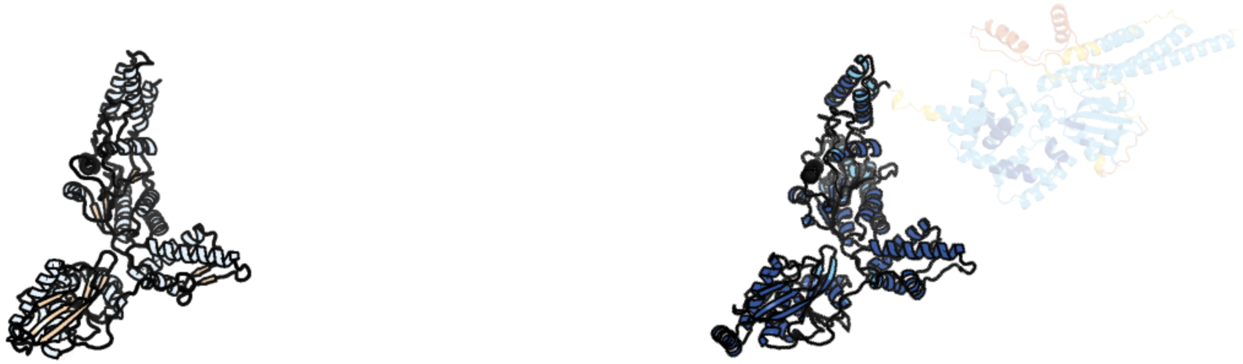

Figure 119: left: reference structure of 6on2 chain C. right: predicted structure of chlorv-1..119, unaligned sequences are shown as transparent

#### chlorv-1..120

- Sequence-based annotation for chlorv-1..120 is putative Rhodanese-like domain
- Best hit was 3tp9 chain B: BETA-LACTAMASE and RHODANESE DOMAIN PROTEIN

| target | prob | fident | alnlen | evalue | theader |
|---|---|---|---|---|---|
| 3tp9-assembly1.cif.gz_B | 1 | 0.231 | 95 | 3.534e-05 | Crystal structure of Alicyclobacillus acidocaldarius protein with beta-lactamase and rhodanese domains |
| 6mxv-assembly1.cif.gz_A | 1 | 0.277 | 108 | 4.299e-05 | The crystal structure of a rhodanese-like family protein from Francisella tularensis subsp. tularensis SCHU S4 |
| 3o3w-assembly2.cif.gz_D | 1 | 0.238 | 105 | 6.361e-05 | Crystal Structure of BH2092 protein (residues 14-131) from Bacillus halodurans, Northeast Structural Genomics Consortium Target BhR228A |

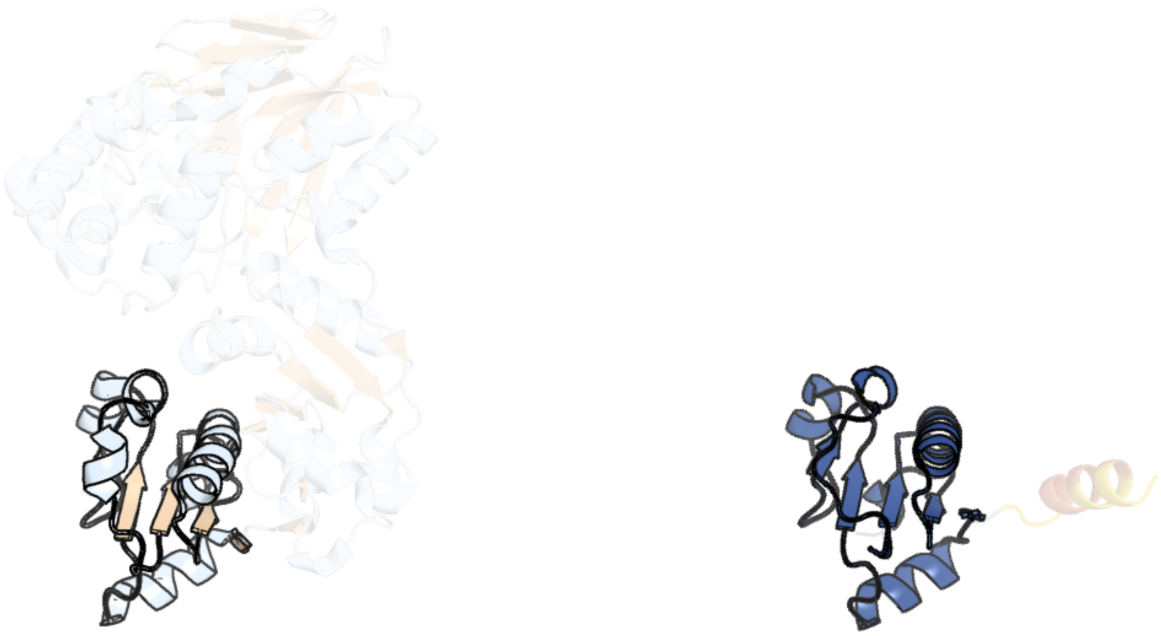

Figure 120: left: reference structure of 3tp9 chain B. right: predicted structure of chlorv-1..120, unaligned sequences are shown as transparent

#### chlorv-1..121

- Sequence-based annotation for chlorv-1..121 is hypothetical protein
- Best hit was 4gf1 chain B: Putative ADP-ribosyltransferase Certhrax

| target | prob | fident | alnlen | evalue | theader |
|---|---|---|---|---|---|
| 4gf1-assembly2.cif.gz_B | 1 | 0.164 | 419 | 1.677e-08 | Crystal Structure of Certhrax |
| 4fxq-assembly2.cif.gz_B | 1 | 0.167 | 419 | 1.86e-08 | Full-length Certhrax toxin from Bacillus cereus in complex with Inhibitor P6 |
| 4gf1-assembly1.cif.gz_A | 1 | 0.181 | 396 | 4.499e-08 | Crystal Structure of Certhrax |

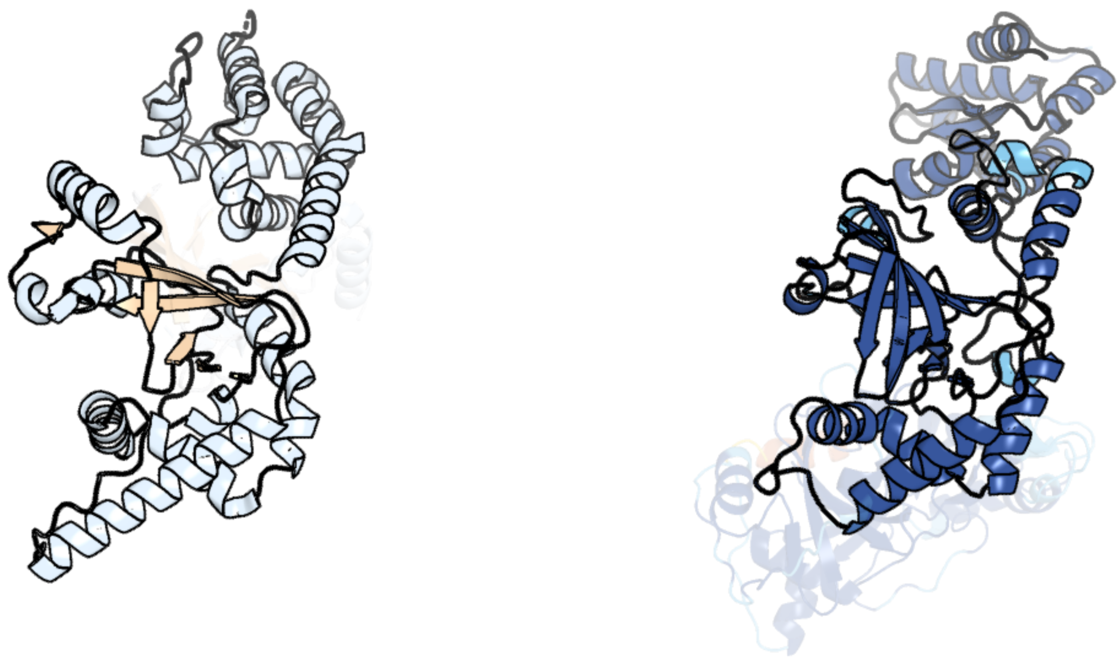

Figure 121: left: reference structure of 4gf1 chain B. right: predicted structure of chlorv-1..121, unaligned sequences are shown as transparent

#### chlorv-1..122

- Sequence-based annotation for chlorv-1..122 is hypothetical protein
- Best hit was 5gm6 chain e: Protein HSH49

| target | prob | fident | alnlen | evalue | theader |
|---|---|---|---|---|---|
| 5gm6-assembly1.cif.gz_e | 1 | 0.137 | 160 | 6.54e-06 | Cryo-EM structure of the activated spliceosome (Bact complex) at 3.5 angstrom resolution |
| 7abi-assembly1.cif.gz_B | 1 | 0.15 | 180 | 7.819e-06 | Human pre-Bact-2 spliceosome |
| 7uk1-assembly1.cif.gz_B | 1 | 0.125 | 175 | 1.186e-05 | Complex Structure of Human Polypyrimidine Splicing Factor (PSF/SFPQ) with Murine Virus-like 30S Transcript-1 (VS30-1) Reveals Cooperative Binding of RNA |

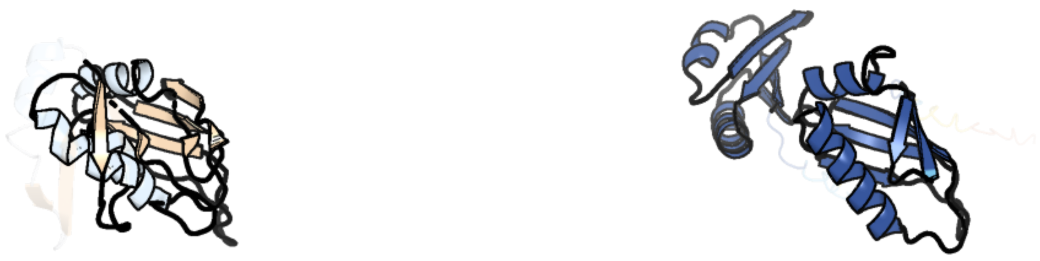

Figure 122: left: reference structure of 5gm6 chain e. right: predicted structure of chlorv-1..122, unaligned sequences are shown as transparent

#### chlorv-1..123

- Sequence-based annotation for chlorv-1..123 is putative Ribonuclease
- Best hit was 7uwe chain C: Ribonuclease HII

| target | prob | fident | alnlen | evalue | theader |
|---|---|---|---|---|---|
| 7uwe-assembly1.cif.gz_C | 1 | 0.364 | 192 | 6.732e-19 | CryoEM Structure of E. coli Transcription-Coupled Ribonucleotide Excision Repair (TC-RER) complex |
| 3o3g-assembly1.cif.gz_A | 1 | 0.384 | 190 | 1.817e-18 | T. maritima RNase H2 in complex with nucleic acid substrate and calcium ions |
| 5y9p-assembly1.cif.gz_A | 1 | 0.354 | 189 | 2.636e-18 | Staphylococcus aureus RNase HII |

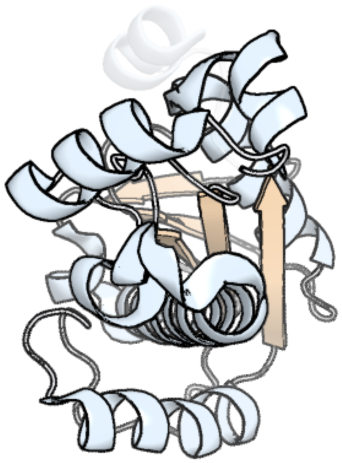

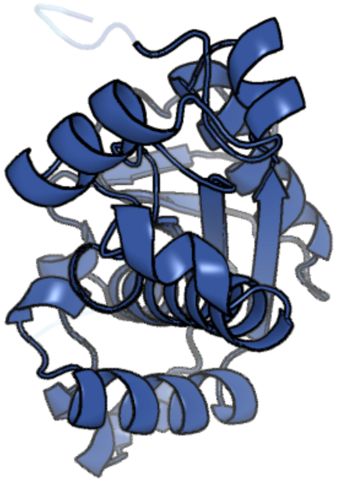

Figure 123: left: reference structure of 7uwe chain C. right: predicted structure of chlorv-1..123, unaligned sequences are shown as transparent

#### chlorv-1..124

- Sequence-based annotation for chlorv-1..124 is putative Exonuclease
- Best hit was 7opk chain A: 5'-3' exoribonuclease

| target | prob | fident | alnlen | evalue | theader |
|---|---|---|---|---|---|
| 7opk-assembly1.cif.gz_A | 1 | 0.301 | 574 | 8.139e-35 | Crystal structure of C. thermophilum Xrn2 |
| 3fqd-assembly1.cif.gz_A | 1 | 0.298 | 600 | 2.205e-34 | Crystal Structure of the S. pombe Rat1-Rai1 Complex |
| 5fir-assembly4.cif.gz_G | 1 | 0.261 | 566 | 9.262e-33 | Crystal structure of C. elegans XRN2 in complex with the XRN2-binding domain of PAXT-1 |

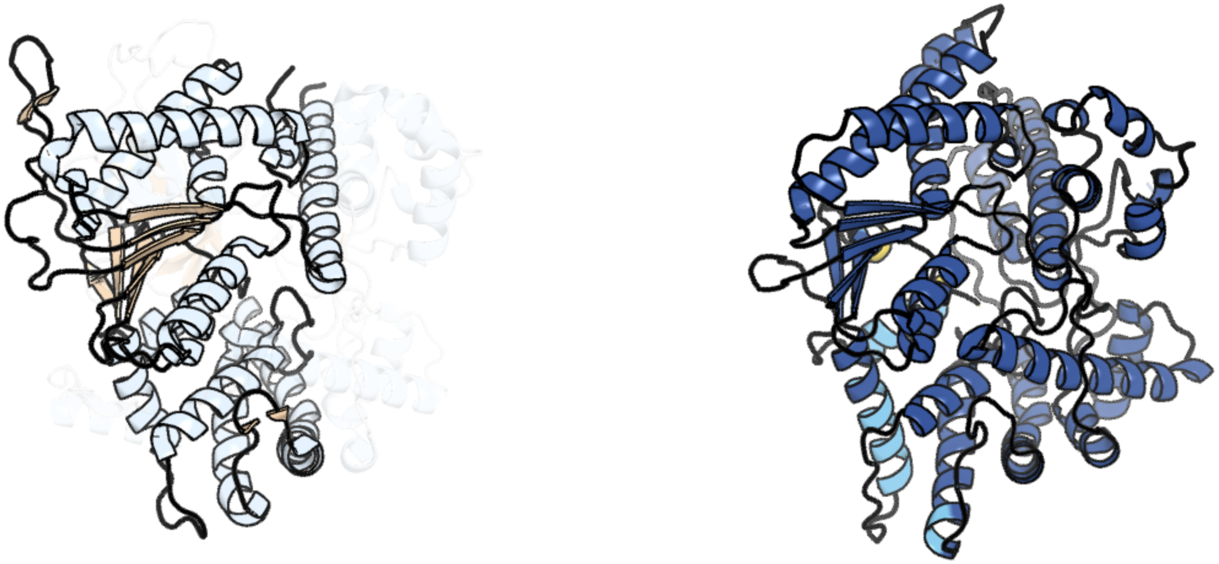

Figure 124: left: reference structure of 7opk chain A. right: predicted structure of chlorv-1..124, unaligned sequences are shown as transparent

#### chlorv-1..125

- Sequence-based annotation for chlorv-1..125 is putative Protein kinase
- Best hit was 2wb8 chain A: SERINE/THREONINE-PROTEIN KINASE HASPIN

| target | prob | fident | alnlen | evalue | theader |
|---|---|---|---|---|---|
| 2wb8-assembly1.cif.gz_A | 1 | 0.162 | 370 | 2.937e-11 | Crystal structure of Haspin kinase |
| 6g3a-assembly1.cif.gz_A | 1 | 0.163 | 367 | 1.079e-10 | Crystal structure of haspin F605T mutant in complex with 5-iodotubercidin |
| 6d3k-assembly1.cif.gz_A | 1 | 0.146 | 347 | 2.656e-07 | Crystal structure of unphosphorylated human PKR kinase domain in complex with ADP |

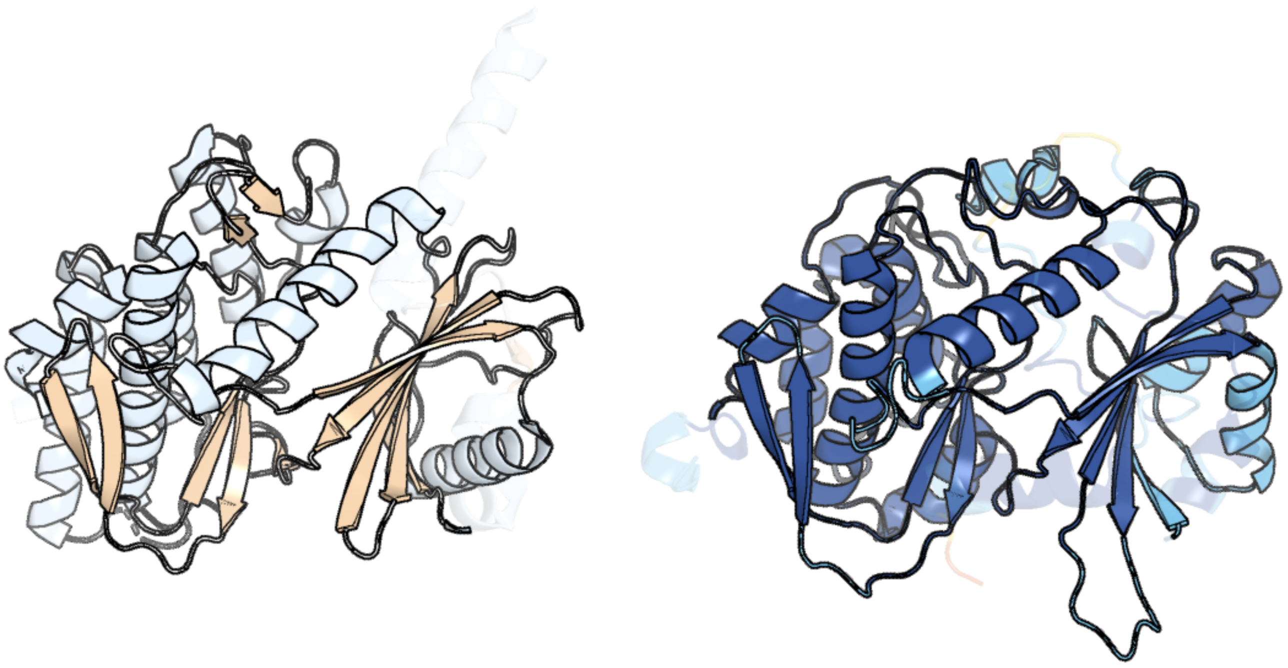

Figure 125: left: reference structure of 2wb8 chain A. right: predicted structure of chlorv-1..125, unaligned sequences are shown as transparent

#### chlorv-1..126

- Sequence-based annotation for chlorv-1..126 is hypothetical protein
- No significant structural hit found

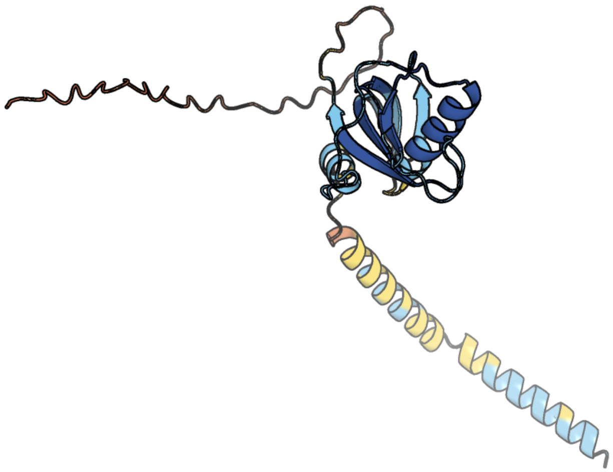

Figure 126: predicted structure of chlorv-1..126

#### chlorv-1..127

- Sequence-based annotation for chlorv-1..127 is hypothetical protein
- No significant structural hit found

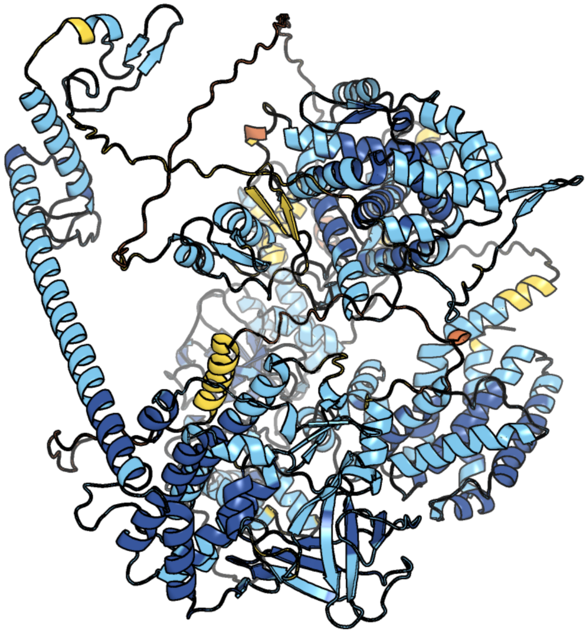

Figure 127: predicted structure of chlorv-1..127

#### chlorv-1..128

- Sequence-based annotation for chlorv-1..128 is hypothetical protein
- No significant structural hit found

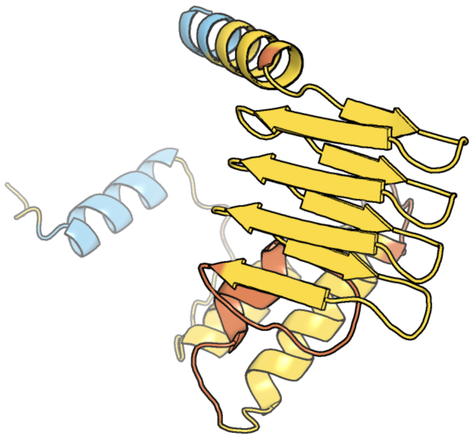

Figure 128: predicted structure of chlorv-1..128

#### chlorv-1..129

- Sequence-based annotation for chlorv-1..129 is putative Helicase
- Best hit was 7zsb chain 7: General transcription and DNA repair factor IIH helicase subunit XPB

| target | prob | fident | alnlen | evalue | theader |
|---|---|---|---|---|---|
| 7zsb-assembly1.cif.gz_7 | 1 | 0.155 | 503 | 7.849e-19 | Yeast RNA polymerase II transcription pre-initiation complex with the +1 nucleosome and NTP, complex C |
| 8bvw-assembly1.cif.gz_0 | 1 | 0.167 | 506 | 1.852e-18 | RNA polymerase II pre-initiation complex with the distal +1 nucleosome (PIC-Nuc18W) |
| 6p4w-assembly2.cif.gz_C | 1 | 0.222 | 440 | 2.061e-18 | XPB helicase in a complex with truncated Bax1 from Sulfurisphaera tokodaii at 2.96 Angstrom resolution |

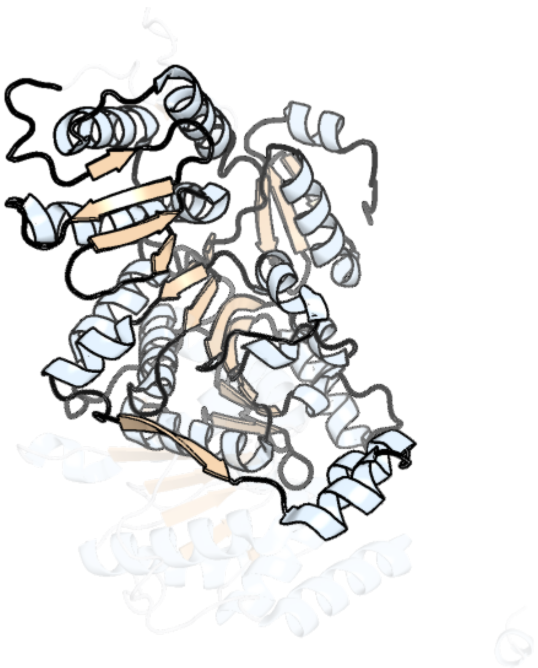

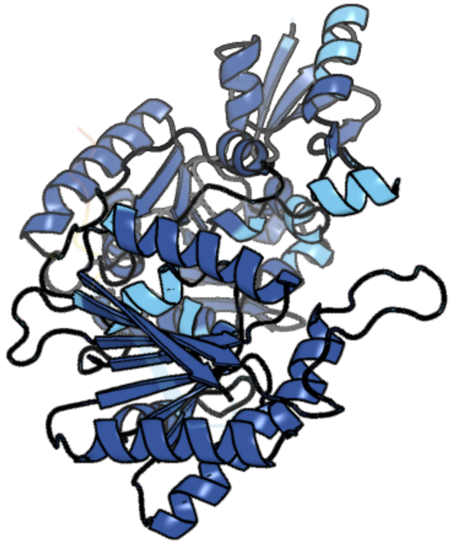

Figure 129: left: reference structure of 7zsb chain 7. right: predicted structure of chlorv-1..129, unaligned sequences are shown as transparent

#### chlorv-1..130

- Sequence-based annotation for chlorv-1..130 is hypothetical protein
- No significant structural hit found

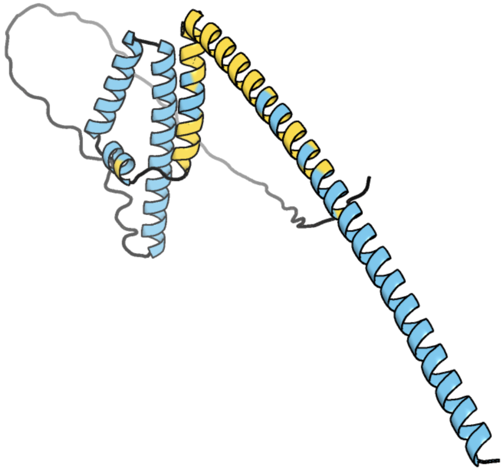

Figure 130: predicted structure of chlorv-1..130

#### chlorv-1..131

- Sequence-based annotation for chlorv-1..131 is putative Calcineurin-like phosphoesterase
- Best hit was 1v73 chain A: psychrophilic phosphatase I

| target | prob | fident | alnlen | evalue | theader |
|---|---|---|---|---|---|
| 1v73-assembly1.cif.gz_A | 1 | 0.217 | 373 | 9.711e-17 | Crystal Structure of Cold-Active Protein-Tyrosine Phosphatase of a Psychrophile Shewanella SP. |
| 2zbm-assembly1.cif.gz_A | 1 | 0.217 | 372 | 1.837e-16 | Crystal Structure of I115M Mutant Cold-Active Protein Tyrosine Phosphatase |
| 2z72-assembly1.cif.gz_A | 1 | 0.214 | 359 | 3.281e-16 | New Structure Of Cold-Active Protein Tyrosine Phosphatase At 1.1 Angstrom |

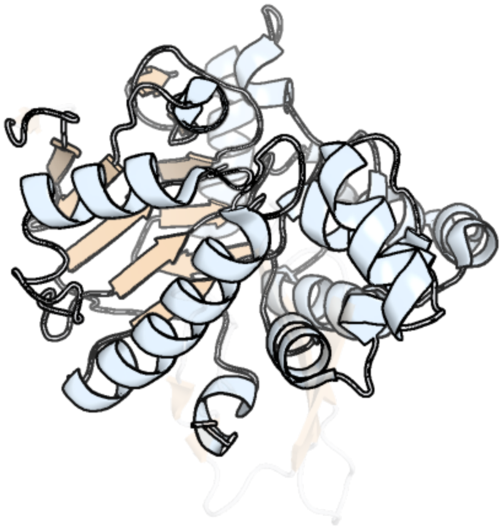

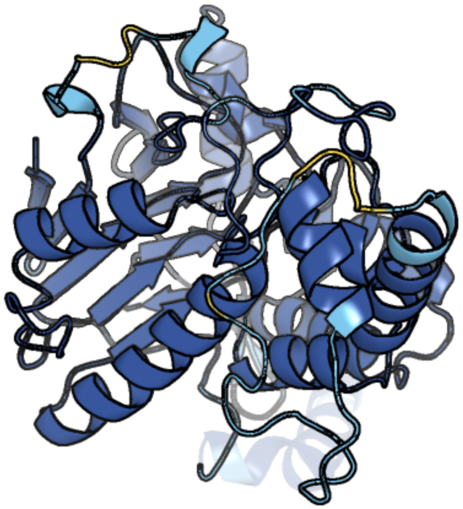

Figure 131: left: reference structure of 1v73 chain A. right: predicted structure of chlorv-1..131, unaligned sequences are shown as transparent

#### chlorv-1..132

- Sequence-based annotation for chlorv-1..132 is putative AAA family ATPase
- Best hit was 6sh3 chain A: Mitochondrial chaperone BCS1

| target | prob | fident | alnlen | evalue | theader |
|---|---|---|---|---|---|
| 6sh3-assembly1.cif.gz_A | 1 | 0.269 | 375 | 8.27e-25 | Structure of the ADP state of the heptameric Bcs1 AAA-ATPase |
| 6uks-assembly1.cif.gz_A | 1 | 0.251 | 389 | 8.774e-25 | ATPgammaS bound mBcs1 |
| 6uko-assembly1.cif.gz_C | 1 | 0.241 | 381 | 2.4e-24 | Structure analysis of full-length mouse bcs1 complex |

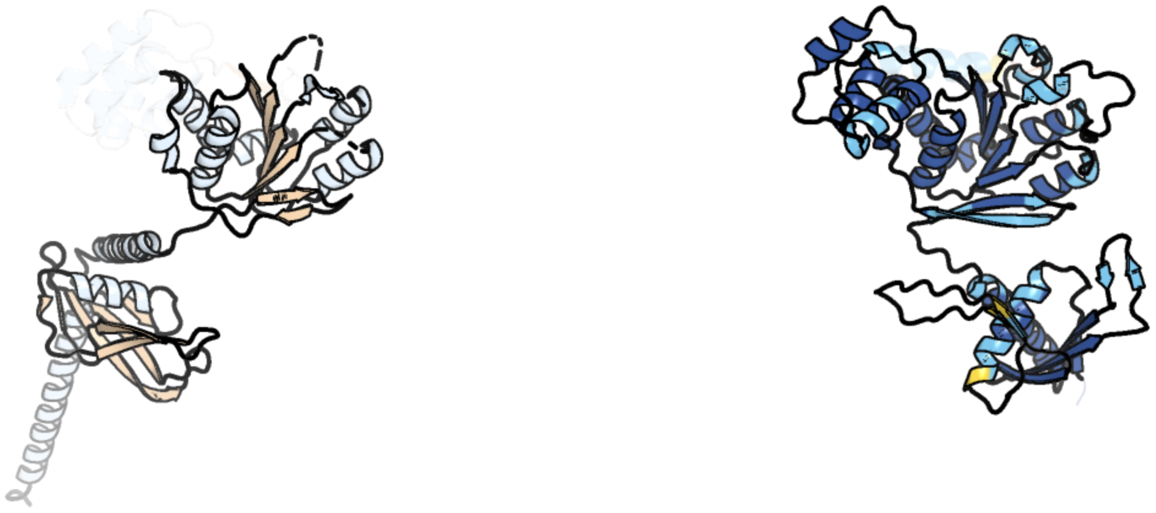

Figure 132: left: reference structure of 6sh3 chain A. right: predicted structure of chlorv-1..132, unaligned sequences are shown as transparent

#### chlorv-1..133

- Sequence-based annotation for chlorv-1..133 is putative eukaryotic translation initiation factor 5
- Best hit was 6sw9 chain 8: Translation initiation factor 2 subunit beta

| target | prob | fident | alnlen | evalue | theader |
|---|---|---|---|---|---|
| 8ppl-assembly1.cif.gz_Is | 1 | 0.188 | 143 | 1.615e-07 | MERS-CoV Nsp1 bound to the human 43S pre-initiation complex |
| 6sw9-assembly1.cif.gz_8 | 1 | 0.196 | 127 | 4.562e-07 | IC2A model of cryo-EM structure of a full archaeal ribosomal translation initiation complex devoid of aIF1 in P. abyssi |
| 6fyy-assembly1.cif.gz_l | 1 | 0.209 | 129 | 6.583e-07 | Structure of a partial yeast 48S preinitiation complex with eIF5 N-terminal domain (model C2) |

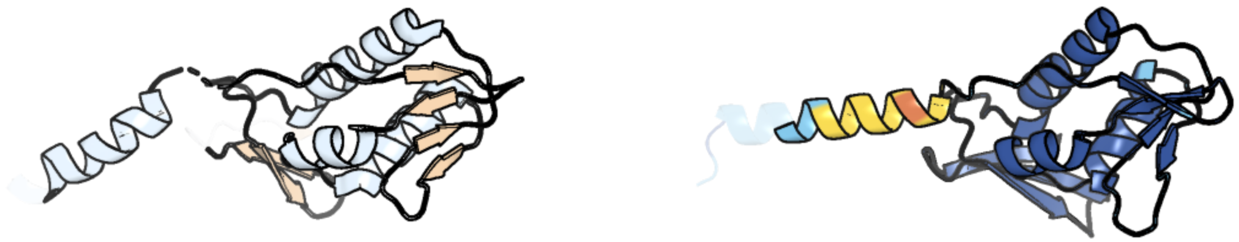

Figure 133: left: reference structure of 6sw9 chain 8. right: predicted structure of chlorv-1..133, unaligned sequences are shown as transparent

#### chlorv-1..134

- Sequence-based annotation for chlorv-1..134 is hypothetical protein
- No significant structural hit found

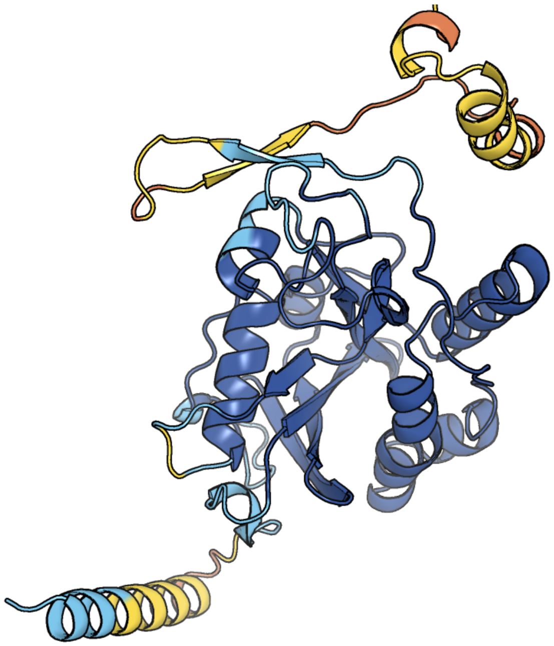

Figure 134: predicted structure of chlorv-1..134

#### chlorv-1..135

- Sequence-based annotation for chlorv-1..135 is hypothetical protein
- Best hit was 4d0q chain A: HYALURONATE LYASE

| target | prob | fident | alnlen | evalue | theader |
|---|---|---|---|---|---|
| 4d0q-assembly1.cif.gz_A | 1 | 0.161 | 118 | 3.842e-05 | Hyaluronan Binding Module of the Streptococcal Pneumoniae Hyaluronate Lyase |
| 6xwv-assembly1.cif.gz_D | 1 | 0.115 | 113 | 7.897e-05 | Crystal structure of drosophila melanogaster CENP-C bound to CAL1 |
| 2xon-assembly1.cif.gz_A | 0.998 | 0.175 | 131 | 8.347e-05 | Structure of TmCBM61 in complex with beta-1,4-galactotriose at 1.4 A resolution |

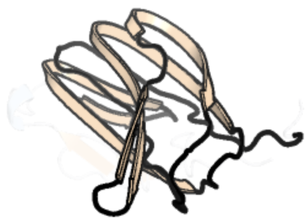

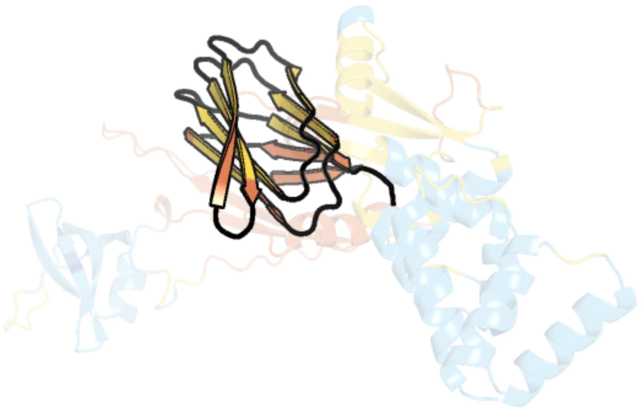

Figure 135: left: reference structure of 4d0q chain A. right: predicted structure of chlorv-1..135, unaligned sequences are shown as transparent

### chlorv-1..136

- Sequence-based annotation for chlorv-1..136 is hypothetical protein
- No significant structural hit found

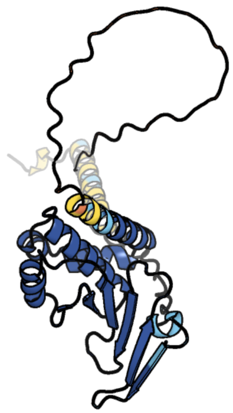

Figure 136: predicted structure of chlorv-1..136

#### chlorv-1..137

- Sequence-based annotation for chlorv-1..137 is hypothetical protein
- Best hit was 6tbn chain B: Probable cytosolic iron-sulfur protein assembly protein Ciao1

| target | prob | fident | alnlen | evalue | theader |
|---|---|---|---|---|---|
| 6tbn-assembly1.cif.gz_B | 1 | 0.102 | 460 | 1.97e-13 | Crystal structure of CIAO1-CIAO2B CIA core complex |
| 5yzv-assembly1.cif.gz_B | 1 | 0.13 | 436 | 2.857e-13 | Biophysical and structural characterization of the thermostable WD40 domain of a prokaryotic protein, Thermomonospora curvata PkwA |
| 7mqa-assembly1.cif.gz_LQ | 1 | 0.128 | 443 | 3.726e-13 | Cryo-EM structure of the human SSU processome, state post-A1 |

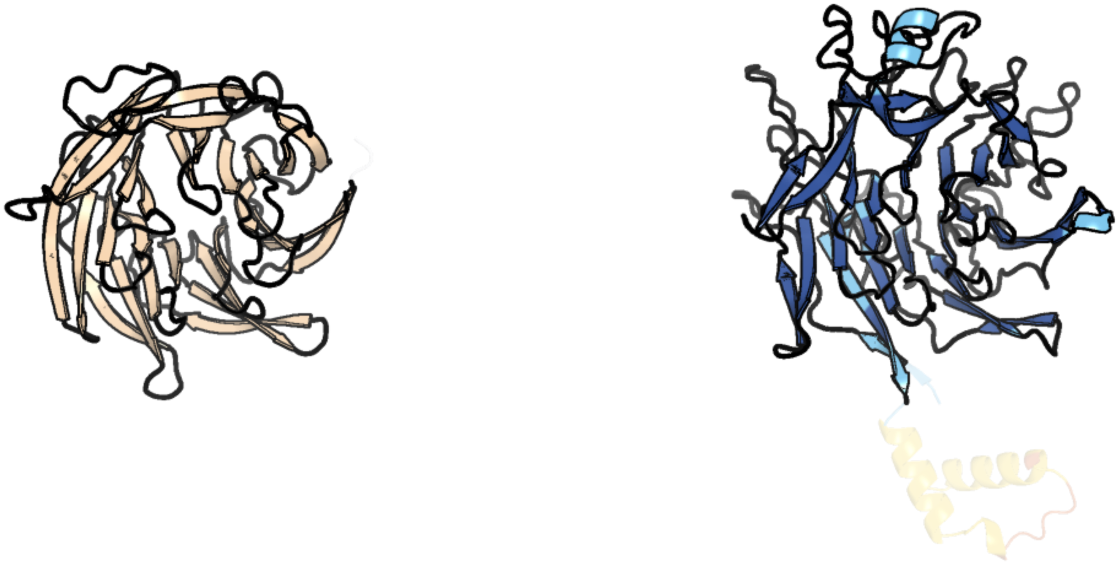

Figure 137: left: reference structure of 6tbn chain B. right: predicted structure of chlorv-1..137, unaligned sequences are shown as transparent

#### chlorv-1..138

- Sequence-based annotation for chlorv-1..138 is hypothetical protein
- Best hit was 7qi3 chain C: Arylamine N-acetyltransferase

| target | prob | fident | alnlen | evalue | theader |
|---|---|---|---|---|---|
| 7qi3-assembly1.cif.gz_C | 1 | 0.099 | 291 | 7.615e-08 | Structure of Fusarium verticillioides NAT1 (FDB2) N-malonyltransferase |
| 4dmo-assembly2.cif.gz_B | 1 | 0.142 | 273 | 1.185e-07 | Crystal structure of the (BACCR)NAT3 arylamine N-acetyltransferase from Bacillus cereus reveals a unique Cys-His-Glu catalytic triad |
| 4b55-assembly1.cif.gz_A | 1 | 0.093 | 289 | 4.726e-07 | Crystal Structure of the Covalent Adduct Formed between Mycobacterium marinum Aryalamine N-acetyltransferase and Phenyl vinyl ketone a derivative of Piperidinols |

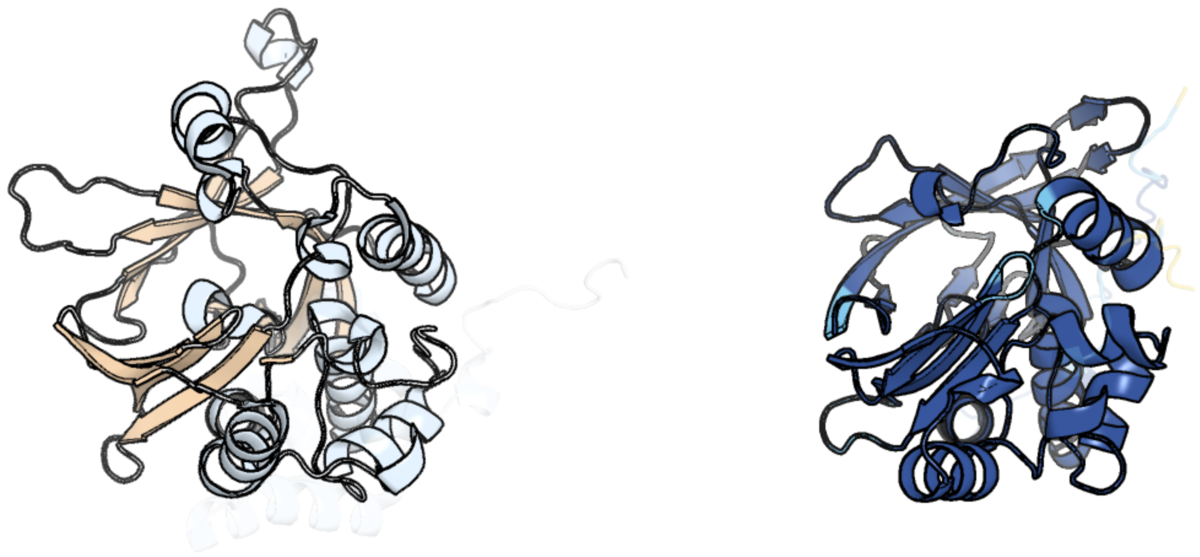

Figure 138: left: reference structure of 7qi3 chain C. right: predicted structure of chlorv-1..138, unaligned sequences are shown as transparent

#### chlorv-1..139

- Sequence-based annotation for chlorv-1..139 is putative Peptidase
- Best hit was 5ilb chain A: Protease Do-like 2, chloroplastic,Protease Do-like 9

| target | prob | fident | alnlen | evalue | theader |
|---|---|---|---|---|---|
| 5ilb-assembly1.cif.gz_A | 1 | 0.207 | 473 | 3.657e-32 | Crystal structure of protease domain of Deg2 linked with the PDZ domain of Deg9 |
| 5il9-assembly1.cif.gz_A | 1 | 0.208 | 466 | 9.676e-31 | Crystal structure of Deg9 |
| 5il9-assembly1.cif.gz_B | 1 | 0.204 | 459 | 1.343e-30 | Crystal structure of Deg9 |

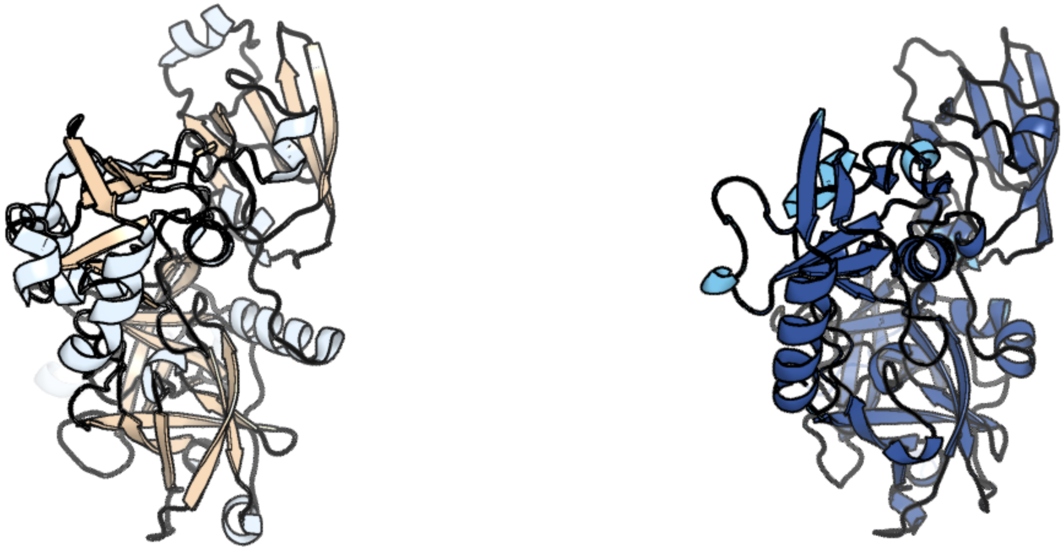

Figure 139: left: reference structure of 5ilb chain A. right: predicted structure of chlorv-1..139, unaligned sequences are shown as transparent

### chlorv-1..140

- Sequence-based annotation for chlorv-1..140 is putative Methyltransferases
- Best hit was 4ckc chain D: MRNA-CAPPING ENZYME CATALYTIC SUBUNIT

| target | prob | fident | alnlen | evalue | theader |
|---|---|---|---|---|---|
| 4ckc-assembly2.cif.gz_D | 1 | 0.132 | 696 | 2.412e-12 | Vaccinia virus capping enzyme complexed with SAH (monoclinic form) |
| 4ckc-assembly1.cif.gz_A | 1 | 0.134 | 656 | 3.261e-12 | Vaccinia virus capping enzyme complexed with SAH (monoclinic form) |
| 4ckb-assembly2.cif.gz_A | 1 | 0.132 | 679 | 9.858e-12 | Vaccinia virus capping enzyme complexed with GTP and SAH |

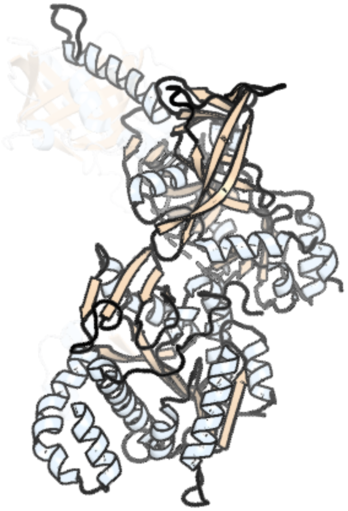

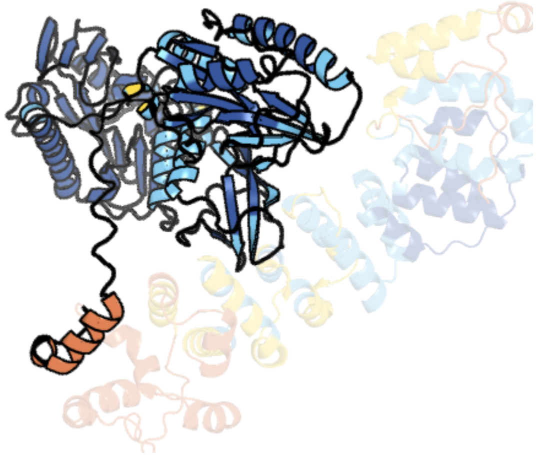

Figure 140: left: reference structure of 4ckc chain D. right: predicted structure of chlorv-1..140, unaligned sequences are shown as transparent

#### chlorv-1..141

- Sequence-based annotation for chlorv-1..141 is hypothetical protein
- No significant structural hit found

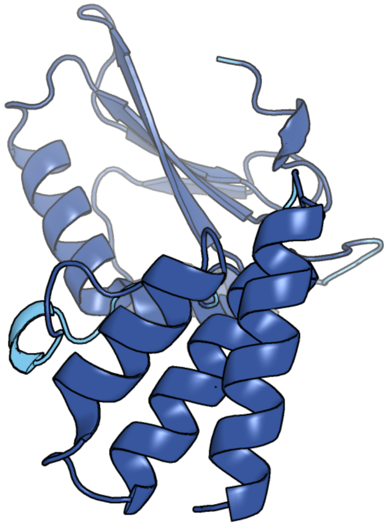

Figure 141: predicted structure of chlorv-1..141

#### chlorv-1..142

- Sequence-based annotation for chlorv-1..142 is putative Nuclease
- Best hit was 6bt1 chain A: Nocturnin

| target | prob | fident | alnlen | evalue | theader |
|---|---|---|---|---|---|
| 6bt1-assembly1.cif.gz_A | 1 | 0.156 | 345 | 4.806e-13 | Structure of the human Nocturnin catalytic domain |
| 6bt2-assembly2.cif.gz_B | 1 | 0.134 | 342 | 2.259e-12 | Structure of the human Nocturnin catalytic domain with bound sulfate anion |
| 6mal-assembly1.cif.gz_A | 1 | 0.154 | 337 | 6.214e-12 | Structure of human Nocturnin C-terminal domain |

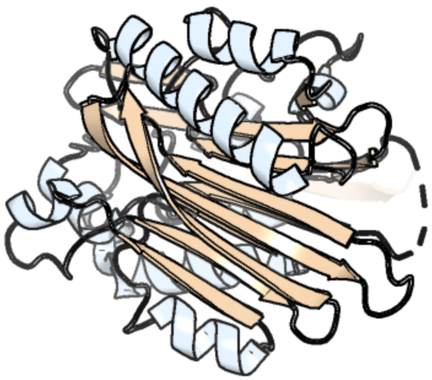

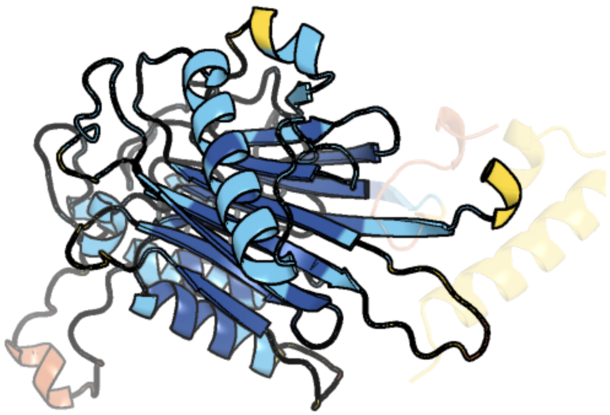

Figure 142: left: reference structure of 6bt1 chain A. right: predicted structure of chlorv-1..142, unaligned sequences are shown as transparent

### chlorv-1..143

- Sequence-based annotation for chlorv-1..143 is hypothetical protein
- No significant structural hit found

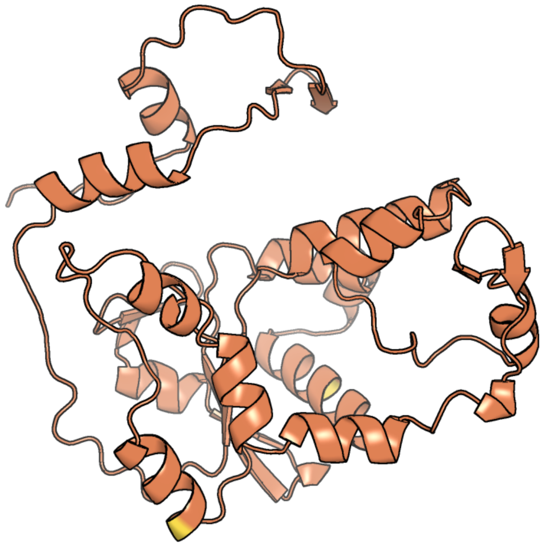

Figure 143: predicted structure of chlorv-1..143

#### chlorv-1..144

- Sequence-based annotation for chlorv-1..144 is hypothetical protein
- No significant structural hit found

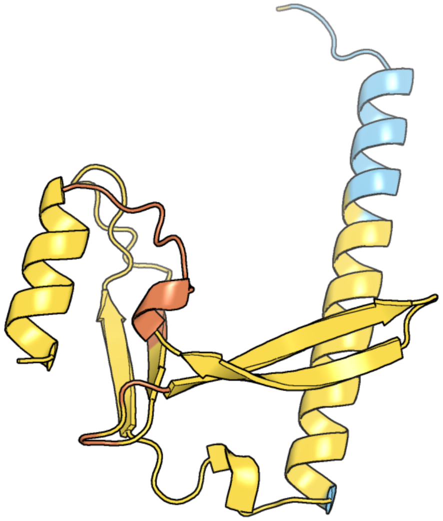

Figure 144: predicted structure of chlorv-1..144

#### chlorv-1..145

- Sequence-based annotation for chlorv-1..145 is putative Aminotransferase
- Best hit was 7lk1 chain A: Ornithine aminotransferase, mitochondrial

| target | prob | fident | alnlen | evalue | theader |
|---|---|---|---|---|---|
| 7lk1-assembly1.cif.gz_A | 1 | 0.517 | 402 | 2.951e-55 | Ornithine Aminotransferase (OAT) with its potent inhibitor - (S)-3-amino-4,4-difluorocyclopent-1-enecarboxylic acid (SS-1-148) - 1 Hour Soaking |
| 2byj-assembly2.cif.gz_C-2 | 1 | 0.504 | 406 | 4.632e-55 | Ornithine aminotransferase mutant Y85I |
| 6hx7-assembly3.cif.gz_C-3 | 1 | 0.512 | 400 | 6.14e-55 | Crystal structure of human R180T variant of ORNITHINE AMINOTRANSFERASE at 1.8 Angstrom |

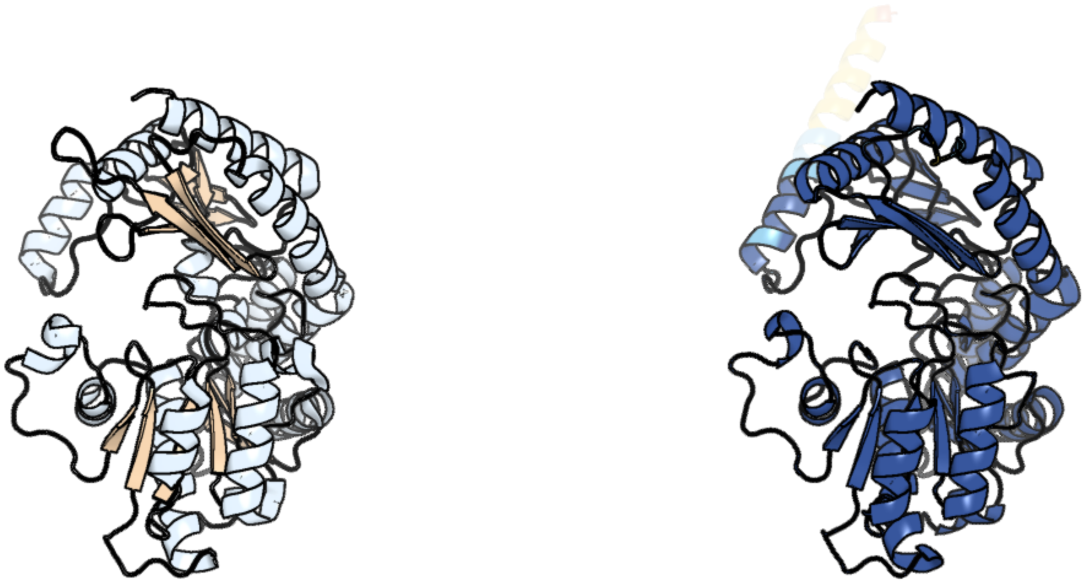

Figure 145: left: reference structure of 7lk1 chain A. right: predicted structure of chlorv-1..145, unaligned sequences are shown as transparent

#### chlorv-1..146

- Sequence-based annotation for chlorv-1..146 is hypothetical protein
- No significant structural hit found

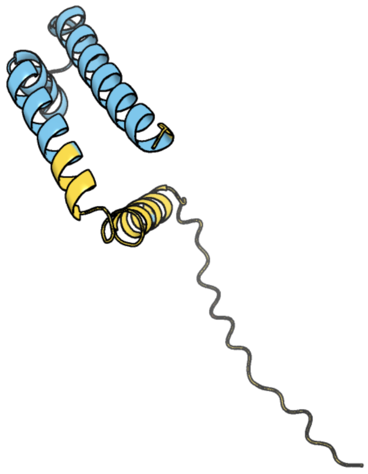

Figure 146: predicted structure of chlorv-1..146

#### chlorv-1..147

- Sequence-based annotation for chlorv-1..147 is putative DUF5760 domain-containing protein
- No significant structural hit found

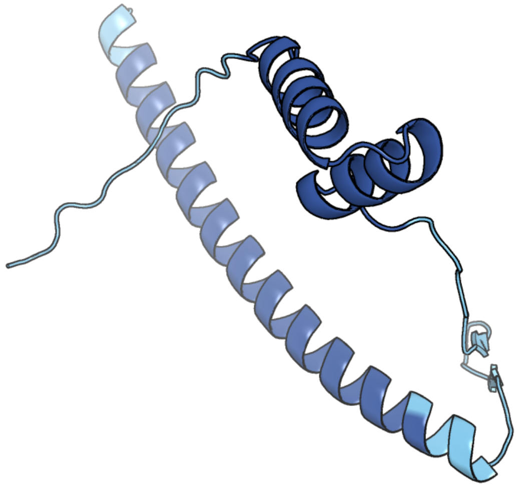

Figure 147: predicted structure of chlorv-1..147

### chlorv-1..148

- Sequence-based annotation for chlorv-1..148 is hypothetical protein
- Best hit was 7eyr chain B: 2-oxoglutarate/Fe(II)-dependent dioxygenase SptF

| target | prob | fident | alnlen | evalue | theader |
|---|---|---|---|---|---|
| 7eyr-assembly2.cif.gz_B-3 | 1 | 0.124 | 258 | 3.615e-07 | Fe(II)/(alpha)ketoglutarate-dependent dioxygenase SptF apo |
| 7eyw-assembly1.cif.gz_B | 1 | 0.121 | 205 | 4.091e-07 | Fe(II)/(alpha)ketoglutarate-dependent dioxygenase SptF with terretonin C |
| 7eyr-assembly1.cif.gz_D-2 | 1 | 0.125 | 240 | 4.629e-07 | Fe(II)/(alpha)ketoglutarate-dependent dioxygenase SptF apo |

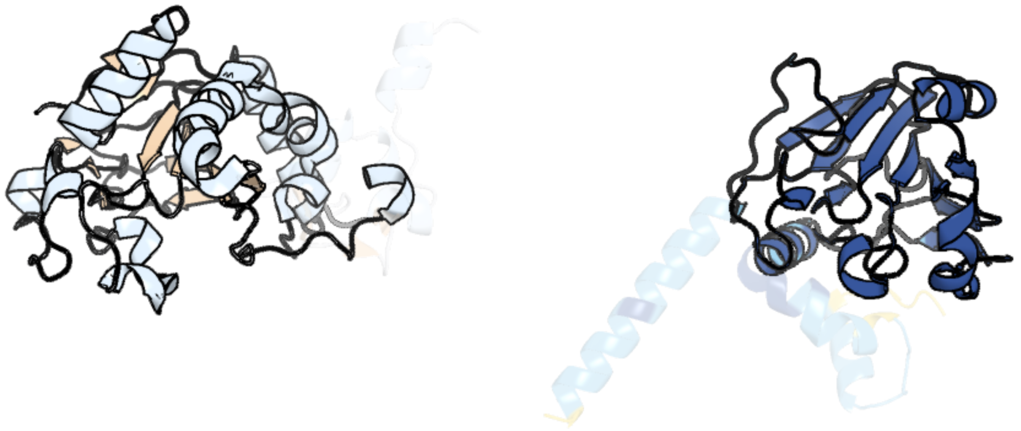

Figure 148: left: reference structure of 7eyr chain B. right: predicted structure of chlorv-1..148, unaligned sequences are shown as transparent

#### chlorv-1..149

- Sequence-based annotation for chlorv-1..149 is hypothetical protein
- No significant structural hit found

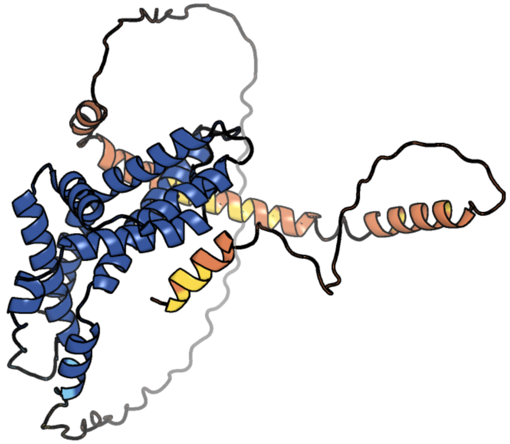

Figure 149: predicted structure of chlorv-1..149

#### chlorv-1..150

- Sequence-based annotation for chlorv-1..150 is putative packaging ATPase/Poxvirus A32 protein
- Best hit was 4kfs chain B: Genome packaging NTPase B204

| target | prob | fident | alnlen | evalue | theader |
|---|---|---|---|---|---|
| 4kfs-assembly2.cif.gz_B | 1 | 0.168 | 219 | 2.915e-09 | Structure of the genome packaging NTPase B204 from Sulfolobus turreted icosahedral virus 2 in complex with AMP |
| 4kfs-assembly1.cif.gz_A | 1 | 0.147 | 237 | 4.497e-09 | Structure of the genome packaging NTPase B204 from Sulfolobus turreted icosahedral virus 2 in complex with AMP |
| 4kfr-assembly3.cif.gz_C | 1 | 0.152 | 236 | 6.129e-09 | Structure of the genome packaging NTPase B204 from Sulfolobus turreted icosahedral virus 2 in complex with sulfate |

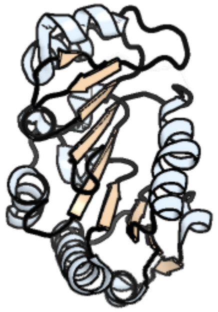

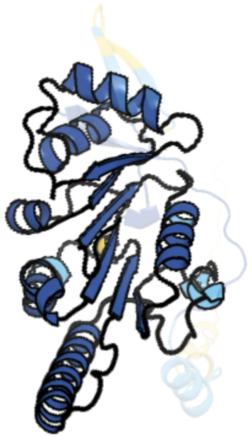

Figure 150: left: reference structure of 4kfs chain B. right: predicted structure of chlorv-1..150, unaligned sequences are shown as transparent

#### chlorv-1..151

- Sequence-based annotation for chlorv-1..151 is hypothetical protein
- No significant structural hit found

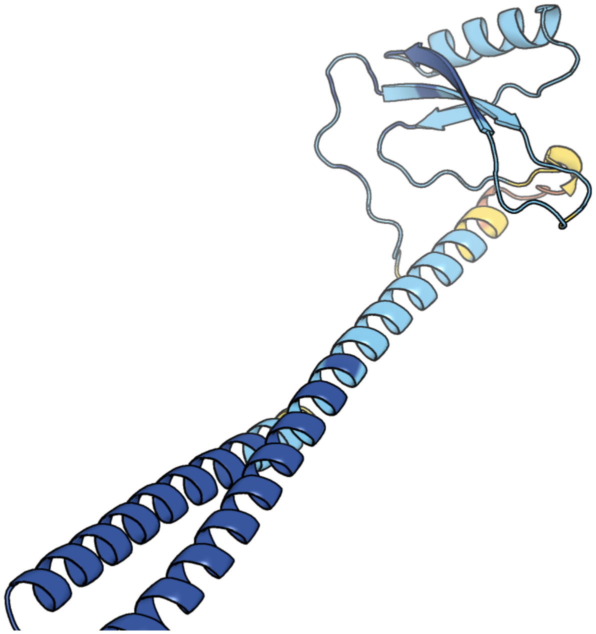

Figure 151: predicted structure of chlorv-1..151

#### chlorv-1..152

- Sequence-based annotation for chlorv-1..152 is putative Poxvirus Late Transcription Factor VLTF3
- No significant structural hit found

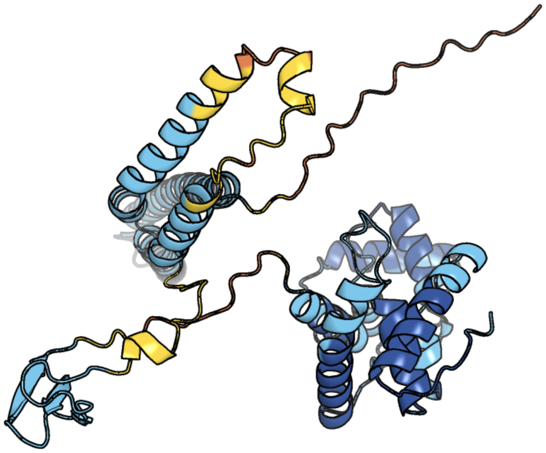

Figure 152: predicted structure of chlorv-1..152

#### chlorv-1..153

- Sequence-based annotation for chlorv-1..153 is putative Major capsid protein
- Best hit was 8h2i chain ab: Major capsid protein (MCP)

| target | prob | fident | alnlen | evalue | theader |
|---|---|---|---|---|---|
| 8h2i-assembly1.cif.gz_ab-5 | 1 | 0.35 | 550 | 4.038e-46 | Near-atomic structure of five-fold averaged PBCV-1 capsid |
| 6ncl-assembly1.cif.gz_d0 | 1 | 0.352 | 550 | 1.603e-45 | Near-atomic structure of icosahedrally averaged PBCV-1 capsid |
| 3kk5-assembly1.cif.gz_A | 1 | 0.328 | 532 | 3.621e-42 | Crystal structure of PBCV-1 VP54 fitted into a cryo-EM reconstruction of the virophage Sputnik |

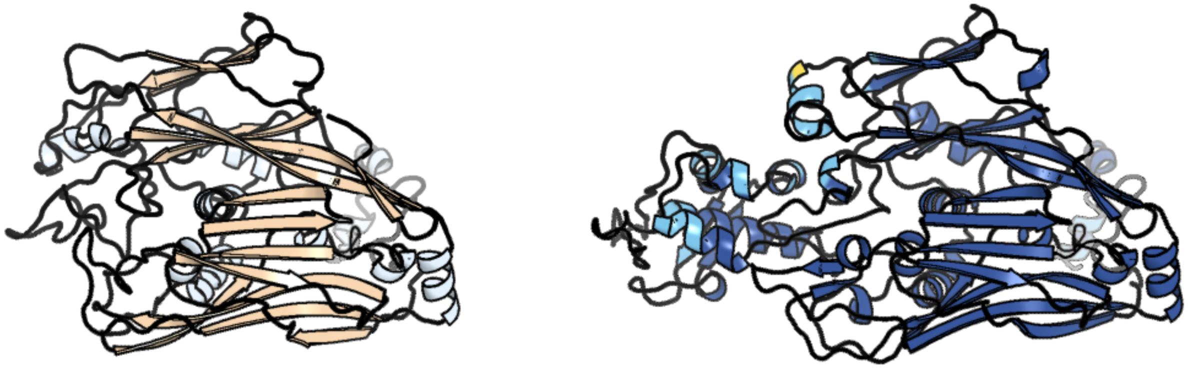

Figure 153: left: reference structure of 8h2i chain ab. right: predicted structure of chlorv-1..153, unaligned sequences are shown as transparent

#### chlorv-1..154

- Sequence-based annotation for chlorv-1..154 is putative HNH endonuclease
- No significant structural hit found

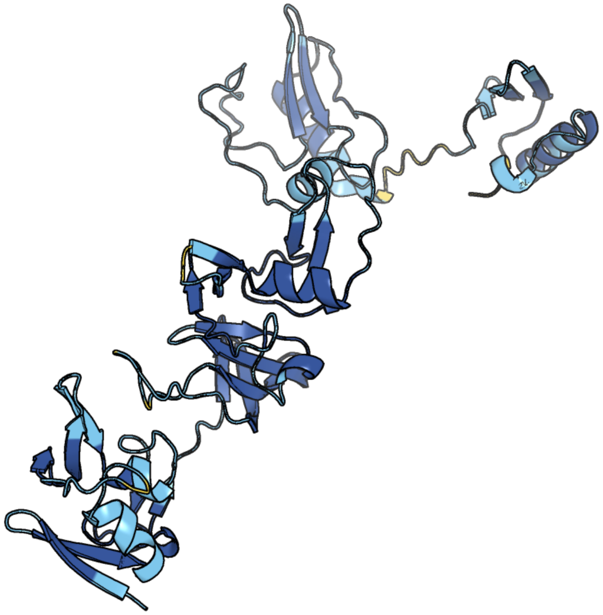

Figure 154: predicted structure of chlorv-1..154

#### chlorv-1..155

- Sequence-based annotation for chlorv-1..155 is hypothetical protein
- Best hit was 1qma chain A: NUCLEAR TRANSPORT FACTOR 2

| target | prob | fident | alnlen | evalue | theader |
| --- | --- | --- | --- | --- | --- |
| 1qma-assembly1.cif.gz_A | 1 | 0.136 | 117 | 2.886e-05 | Nuclear Transport Factor 2 (NTF2) W7A mutant |
| 2r4i-assembly1.cif.gz_B | 1 | 0.139 | 122 | 3.882e-05 | CRYSTAL STRUCTURE OF A NTF2-LIKE PROTEIN (CHU_1428) FROM CYTOPHAGA HUTCHINSONII ATCC 33406 AT 1.60 A RESOLUTION |
| 3nv0-assembly1.cif.gz_B | 1 | 0.108 | 138 | 4.371e-05 | Crystal structure and mutational analysis of the NXF2/NXT1 heterodimeric complex from caenorhabditis elegans at 1.84 A resolution |

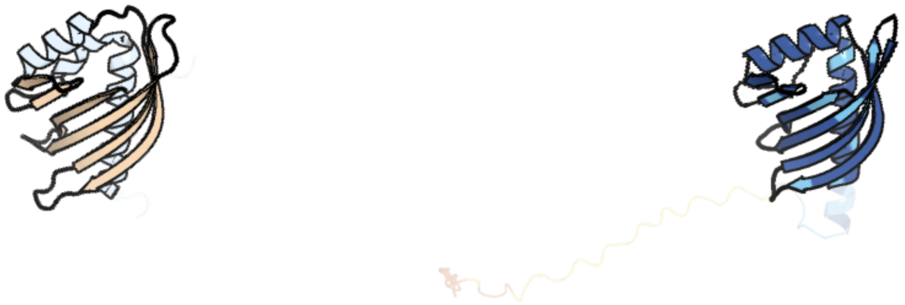

Figure 155: left: reference structure of 1qma chain A. right: predicted structure of chlorv-1..155, unaligned sequences are shown as transparent

#### chlorv-1..156

- Sequence-based annotation for chlorv-1..156 is putative Dioxygenase
- Best hit was 4z82 chain A: Cysteine dioxygenase type 1

| target | prob | fident | alnlen | evalue | theader |
|---|---|---|---|---|---|
| 4z82-assembly1.cif.gz_A | 1 | 0.271 | 114 | 2.365e-08 | Cysteine bound rat cysteine dioxygenase C164S variant at pH 8.1 |
| 5i0u-assembly1.cif.gz_A | 1 | 0.274 | 113 | 8.477e-08 | Incompletely interpreted D-cysteine soak of Cysteine Dioxygenase at pH 7.0 |
| 6u4l-assembly1.cif.gz_A | 1 | 0.25 | 116 | 9.944e-08 | cysteine dioxygenase variant - C93E |

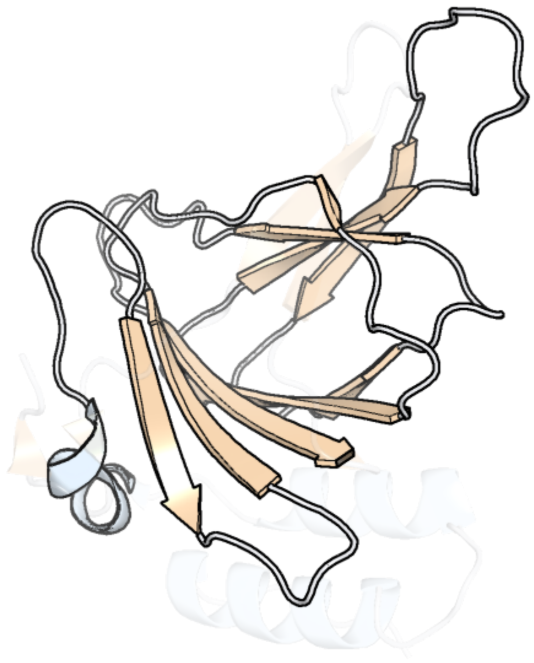

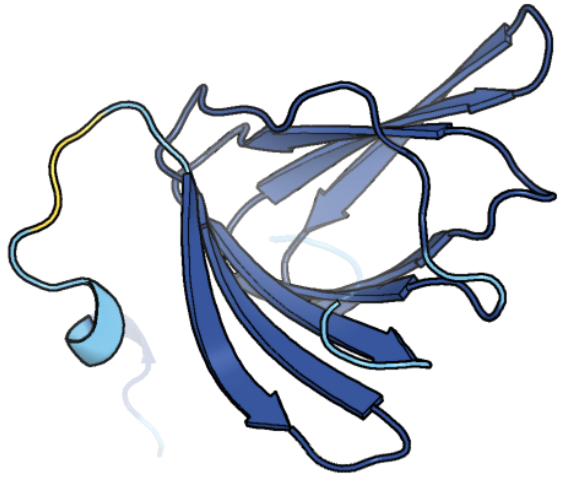

Figure 156: left: reference structure of 4z82 chain A. right: predicted structure of chlorv-1..156, unaligned sequences are shown as transparent

#### chlorv-1..157

- Sequence-based annotation for chlorv-1..157 is putative DUF2177 domain-containing membrane protein
- No significant structural hit found

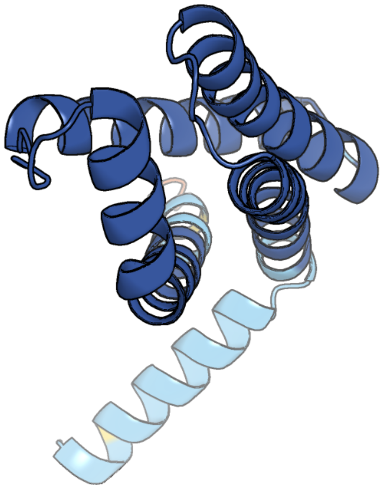

Figure 157: predicted structure of chlorv-1..157

### chlorv-1..158

- Sequence-based annotation for chlorv-1..158 is hypothetical protein
- No significant structural hit found

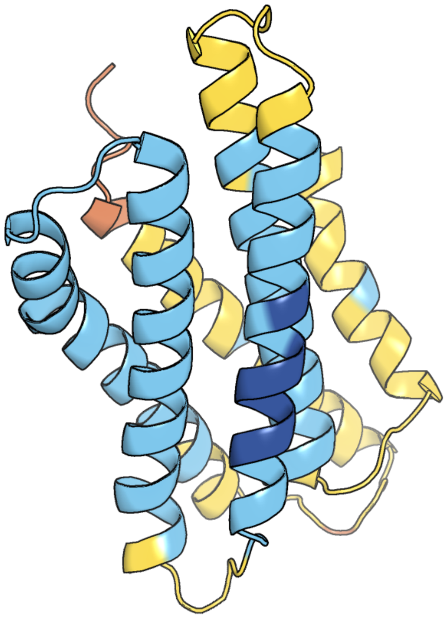

Figure 158: predicted structure of chlorv-1..158

#### chlorv-1..159

- Sequence-based annotation for chlorv-1..159 is hypothetical protein
- No significant structural hit found

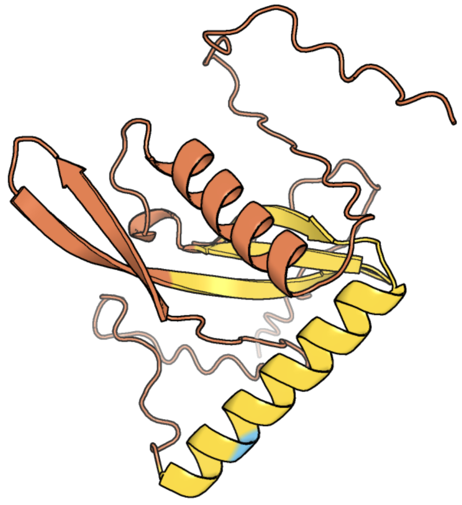

Figure 159: predicted structure of chlorv-1..159

#### chlorv-1..160

- Sequence-based annotation for chlorv-1..160 is putative Major capsid protein
- Best hit was 8rbs chain A1: Major capsid protein

| target | prob | fident | alnlen | evalue | theader |
|---|---|---|---|---|---|
| 8rbs-assembly1.cif.gz__A1 | 1 | 0.145 | 681 | 3.083e-29 | Emiliania huxleyi virus 201 (EhV-201) asymmetrical unit of capsid proteins predicted by AlphaFold2 fitted into the cryo-EM density of EhV-201 virion composite map. |
| 8h2i-assembly1.cif.gz__ab-5 | 1 | 0.147 | 704 | 6.205e-26 | Near-atomic structure of five-fold averaged PBCV-1 capsid |
| 6ncl-assembly1.cif.gz__d0 | 1 | 0.14 | 705 | 2.533e-25 | Near-atomic structure of icosahedrally averaged PBCV-1 capsid |

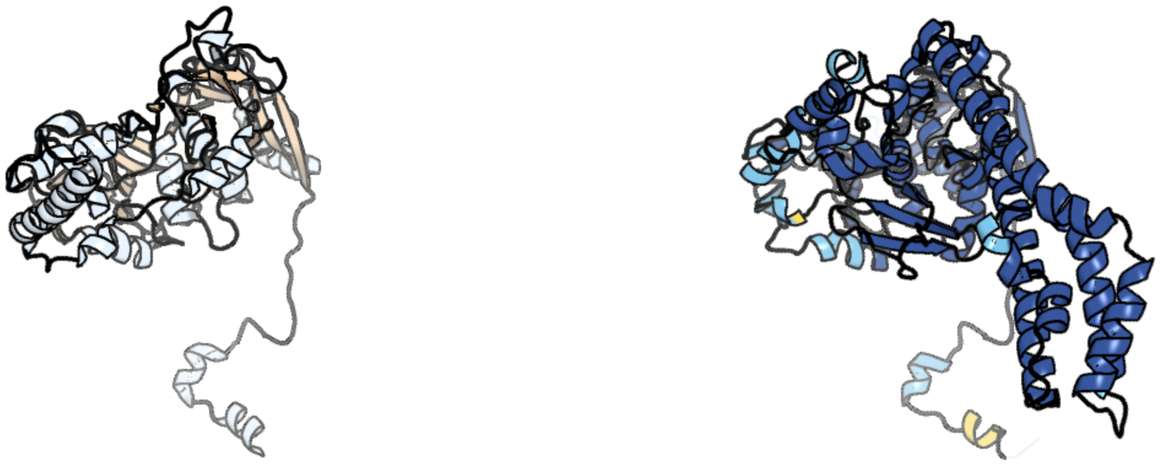

Figure 160: left: reference structure of 8rbs chain A1. right: predicted structure of chlorv-1..160, unaligned sequences are shown as transparent

### chlorv-1..161

- Sequence-based annotation for chlorv-1..161 is hypothetical protein
- No significant structural hit found

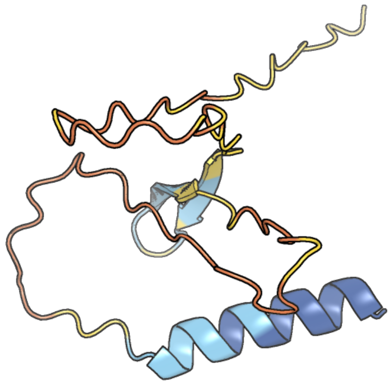

Figure 161: predicted structure of chlorv-1..161

#### chlorv-1..162

- Sequence-based annotation for chlorv-1..162 is hypothetical protein
- No significant structural hit found

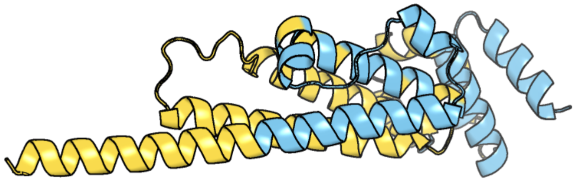

Figure 162: predicted structure of chlorv-1..162

#### chlorv-1..163

- Sequence-based annotation for chlorv-1..163 is putative Capsid protein
- Best hit was 1m4x chain A: PBCV-1 virus capsid

| target | prob | fident | alnlen | evalue | theader |
|---|---|---|---|---|---|
| 3kk5-assembly1.cif.gz_A | 1 | 0.18 | 433 | 2.876e-18 | Crystal structure of PBCV-1 VP54 fitted into a cryo-EM reconstruction of the virophage Sputnik |
| 1m4x-assembly1.cif.gz_A | 1 | 0.182 | 427 | 3.619e-18 | PBCV-1 virus capsid, quasi-atomic model |
| 8h2i-assembly1.cif.gz_ab-5 | 1 | 0.188 | 429 | 4.554e-18 | Near-atomic structure of five-fold averaged PBCV-1 capsid |

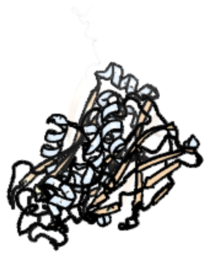

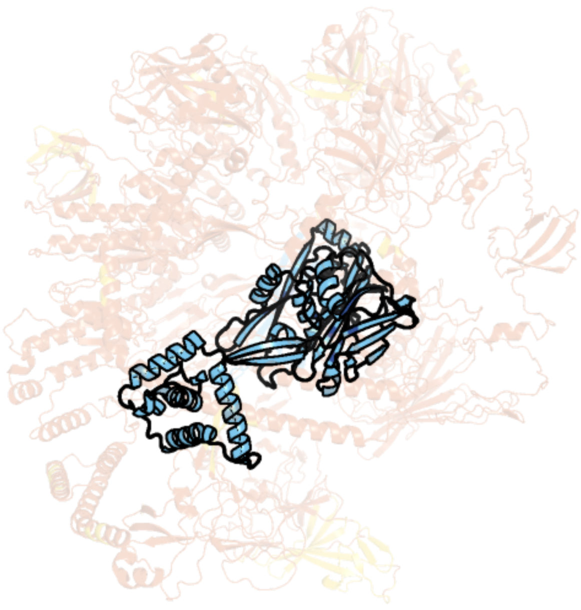

Figure 163: left: reference structure of 1m4x chain A. right: predicted structure of chlorv-1..163, unaligned sequences are shown as transparent

#### chlorv-1..164

- Sequence-based annotation for chlorv-1..164 is putative replication factor C small subunit 2
- Best hit was 7tku chain E: Replication factor C subunit 5

| target | prob | fident | alnlen | evalue | theader |
|---|---|---|---|---|---|
| 7tku-assembly1.cif.gz_E | 1 | 0.233 | 343 | 4.754e-20 | Structure of the yeast clamp loader (Replication Factor C RFC) bound to the open sliding clamp (Proliferating Cell Nuclear Antigen PCNA) |
| 8dr5-assembly1.cif.gz_E | 1 | 0.261 | 329 | 1.268e-19 | Open state of RFC:PCNA bound to a 3' ss/dsDNA junction (DNA2) with NTD |
| 7thv-assembly1.cif.gz_E | 1 | 0.258 | 325 | 3.748e-19 | Structure of the yeast clamp loader (Replication Factor C RFC) bound to the sliding clamp (Proliferating Cell Nuclear Antigen PCNA) in an autoinhibited conformation |

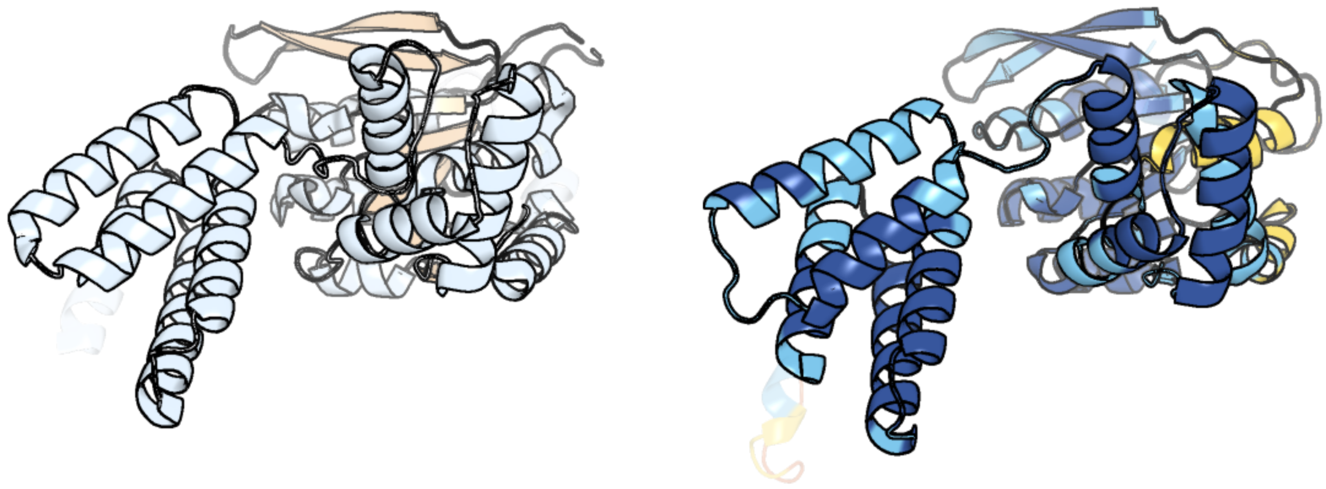

Figure 164: left: reference structure of 7tku chain E. right: predicted structure of chlorv-1..164, unaligned sequences are shown as transparent

#### chlorv-1..165

- Sequence-based annotation for chlorv-1..165 is putative Protein kinase
- No significant structural hit found

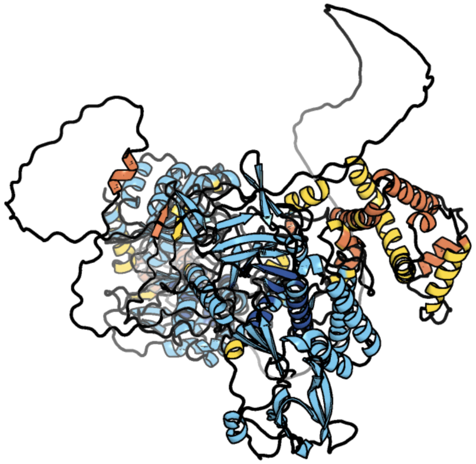

Figure 165: predicted structure of chlorv-1..165

#### chlorv-1..166

- Sequence-based annotation for chlorv-1..166 is putative DUF5761 domain-containing protein
- Best hit was 6ncl chain c9: P8

| target | prob | fident | alnlen | evalue | theader |
|---|---|---|---|---|---|
| 6ncl-assembly1.cif.gz_c9 | 1 | 0.25 | 136 | 5.349e-05 | Near-atomic structure of icosahedrally averaged PBCV-1 capsid |
| 5ku9-assembly2.cif.gz_B | 0.196 | 0.139 | 136 | 0.701 | Crystal structure of MCL1 with compound 1 |
| 5f3q-assembly1.cif.gz_B | 0.063 | 0.115 | 147 | 1.201 | Crystal structure of a noncanonical Dicer protein from Entamoeba histolytica |

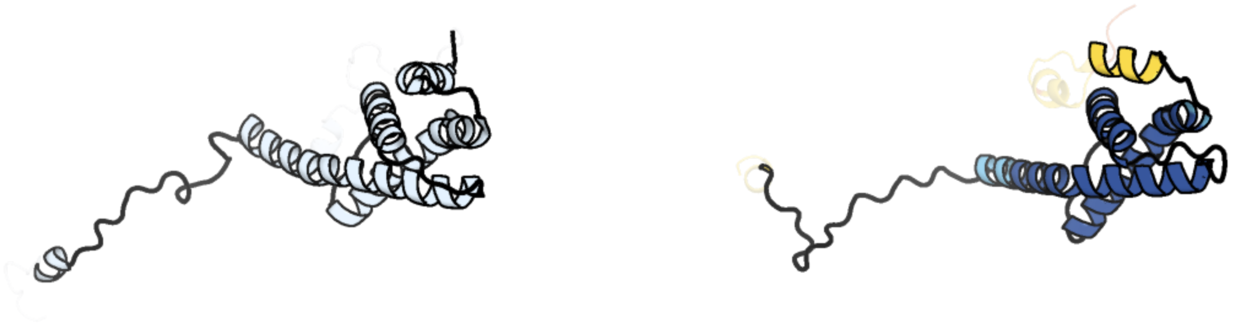

Figure 166: left: reference structure of 6ncl chain c9. right: predicted structure of chlorv-1..166, unaligned sequences are shown as transparent

#### chlorv-1..167

- Sequence-based annotation for chlorv-1..167 is putative Poxvirus P4B major core protein
- Best hit was 5chv chain B: Ubl carboxyl-terminal hydrolase 18

| target | prob | fident | alnlen | evalue | theader |
|---|---|---|---|---|---|
| 5chv-assembly2.cif.gz_B | 1 | 0.085 | 373 | 3.998e-08 | Crystal structure of USP18-ISG15 complex |
| 2y6e-assembly1.cif.gz_C | 1 | 0.128 | 404 | 4.881e-08 | Structure of the D1D2 domain of USP4, the conserved catalytic domain |
| 5l8h-assembly1.cif.gz_A | 1 | 0.084 | 380 | 5.131e-08 | Structure of USP46-UbVME |

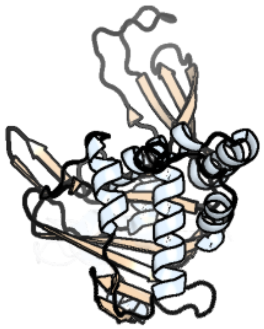

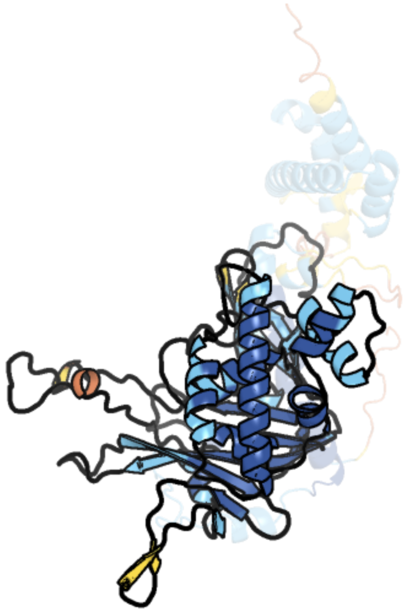

Figure 167: left: reference structure of 5chv chain B. right: predicted structure of chlorv-1..167, unaligned sequences are shown as transparent

#### chlorv-1..168

- Sequence-based annotation for chlorv-1..168 is hypothetical protein
- Best hit was 8dr5 chain A: Replication factor C subunit 1

| target | prob | fident | alnlen | evalue | theader |
|---|---|---|---|---|---|
| 8dr5-assembly1.cif.gz_A | 1 | 0.197 | 491 | 4.046e-17 | Open state of RFC:PCNA bound to a 3' ss/dsDNA junction (DNA2) with NTD |
| 8dr0-assembly1.cif.gz_A | 1 | 0.189 | 491 | 5.47e-17 | Closed state of RFC:PCNA bound to a 3' ss/dsDNA junction |
| 8dr3-assembly1.cif.gz_A | 1 | 0.203 | 482 | 9.508e-17 | Closed state of RFC:PCNA bound to a 3' ss/dsDNA junction (DNA2) with NTD |

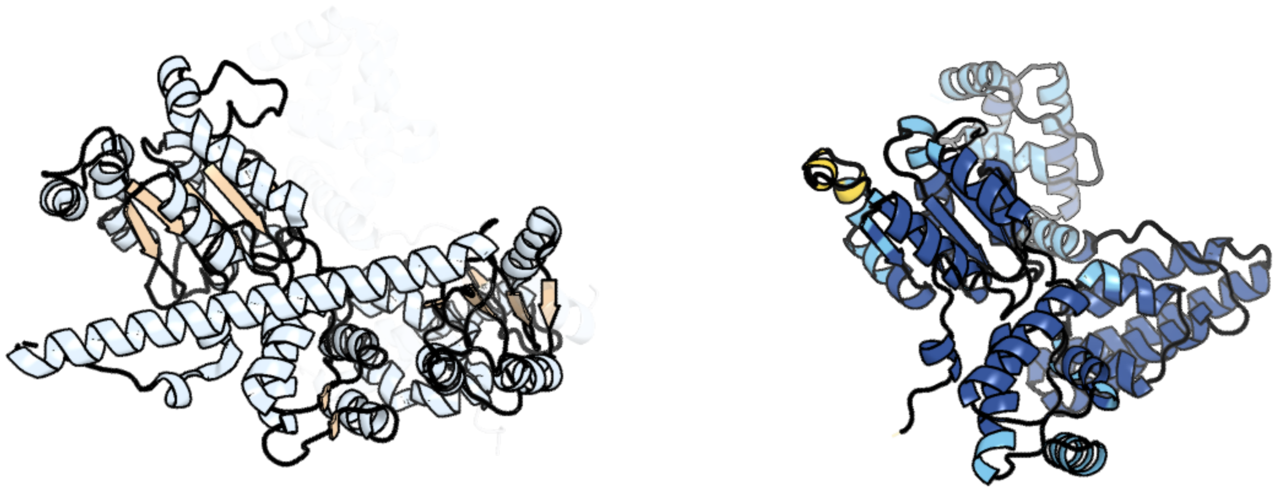

Figure 168: left: reference structure of 8dr5 chain A. right: predicted structure of chlorv-1..168, unaligned sequences are shown as transparent

#### chlorv-1..169

- Sequence-based annotation for chlorv-1..169 is hypothetical protein
- No significant structural hit found

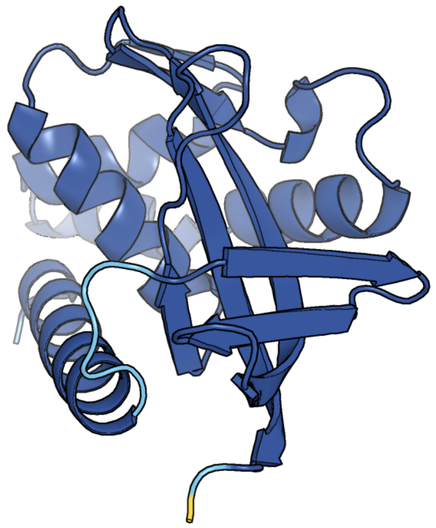

Figure 169: predicted structure of chlorv-1..169

#### chlorv-1..170

- Sequence-based annotation for chlorv-1..170 is hypothetical protein
- No significant structural hit found

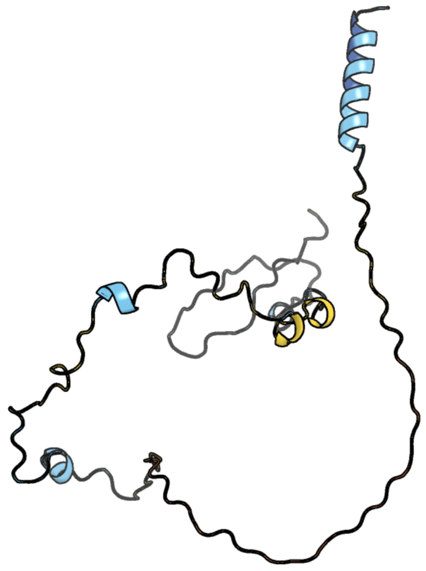

Figure 170: predicted structure of chlorv-1..170

#### chlorv-1..171

- Sequence-based annotation for chlorv-1..171 is putative ELP3-like acetyltransferase
- Best hit was 6iad chain A: Histone acetyltransferase, ELP3 family

| target | prob | fident | alnlen | evalue | theader |
|---|---|---|---|---|---|
| 6iad-assembly1.cif.gz_A | 1 | 0.36 | 452 | 4.67e-37 | Apo crystal structure of archaeal Methanocaldococcus infernus Elp3 (del1-54) |
| 8asw-assembly1.cif.gz_C | 1 | 0.32 | 453 | 2.651e-36 | Cryo-EM structure of yeast Elp123 in complex with alanine tRNA |
| 8asv-assembly1.cif.gz_C | 1 | 0.272 | 587 | 5.807e-36 | Cryo-EM structure of yeast Elongator complex |

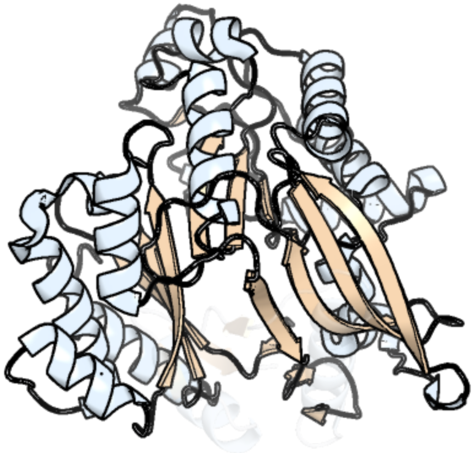

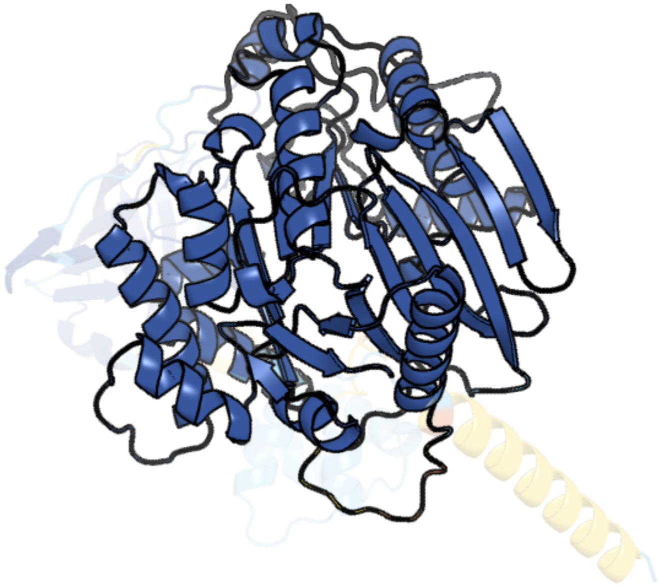

Figure 171: left: reference structure of 6iad chain A. right: predicted structure of chlorv-1..171, unaligned sequences are shown as transparent

#### chlorv-1..172

- Sequence-based annotation for chlorv-1..172 is putative Patatin-like phospholipase
- Best hit was 5fya chain A: PATATIN-LIKE PROTEIN, PLPD

| target | prob | fident | alnlen | evalue | theader |
|---|---|---|---|---|---|
| 5fya-assembly1.cif.gz_A | 1 | 0.187 | 283 | 3.529e-14 | Cubic crystal of the native PlpD |
| 5fqu-assembly1.cif.gz_A | 1 | 0.184 | 276 | 1.252e-13 | Orthorhombic crystal structure of of PlpD (selenomethionine derivative) |
| 5fya-assembly1.cif.gz_B | 1 | 0.187 | 283 | 1.669e-13 | Cubic crystal of the native PlpD |

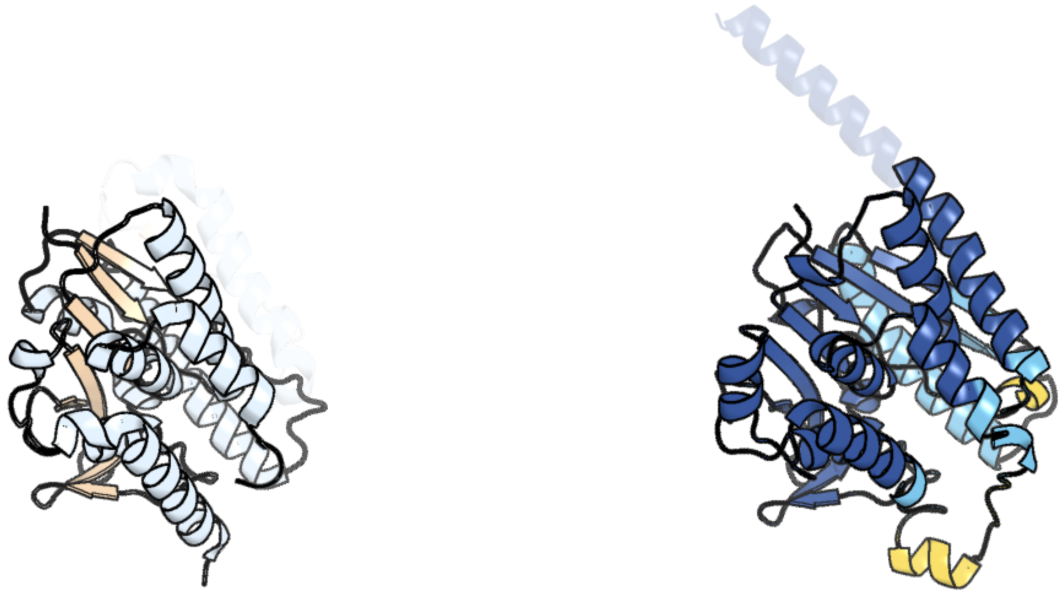

Figure 172: left: reference structure of 5fya chain A. right: predicted structure of chlorv-1..172, unaligned sequences are shown as transparent

### chlorv-1..173

- Sequence-based annotation for chlorv-1..173 is putative DnaJ domain
- No significant structural hit found

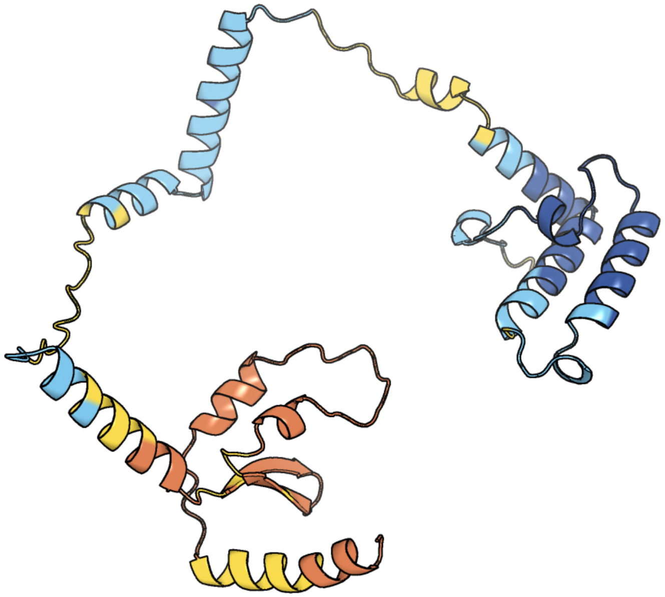

Figure 173: predicted structure of chlorv-1..173

### chlorv-1..174

- Sequence-based annotation for chlorv-1..174 is putative ADP-ribosylglycohydrolase
- Best hit was 3hfw chain A: Protein ADP-ribosylarginine hydrolase

| target | prob | fident | alnlen | evalue | theader |
|---|---|---|---|---|---|
| 3hfw-assembly1.cif.gz_A | 1 | 0.231 | 371 | 2.68e-14 | Crystal Structure of human ADP-ribosylhydrolase 1 (hARH1) |
| 7aks-assembly2.cif.gz_CCC | 1 | 0.167 | 376 | 7.6e-11 | Human ADP-ribosylserine hydrolase ARH3 mutant E41A in complex with H2B-S7-mar peptide |
| 6g1p-assembly1.cif.gz_A | 1 | 0.169 | 377 | 3.123e-10 | Apo form of ADP-ribosylserine hydrolase ARH3 of Latimeria chalumnae |

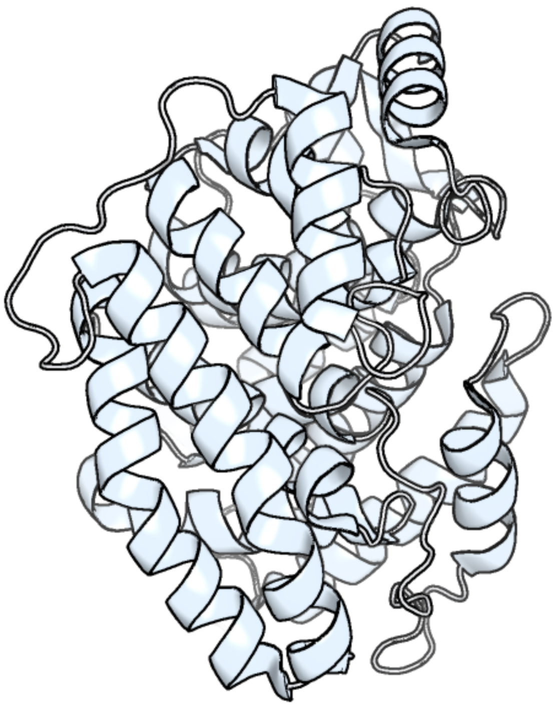

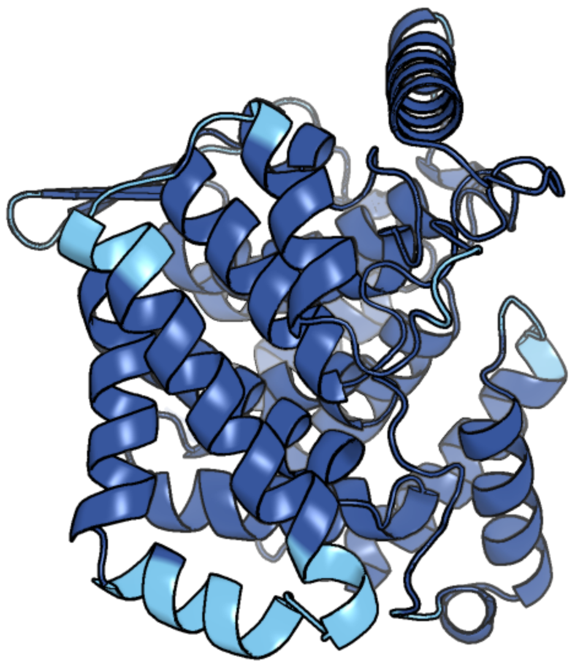

Figure 174: left: reference structure of 3hfw chain A. right: predicted structure of chlorv-1..174, unaligned sequences are shown as transparent

#### chlorv-1..175

- Sequence-based annotation for chlorv-1..175 is hypothetical protein
- No significant structural hit found

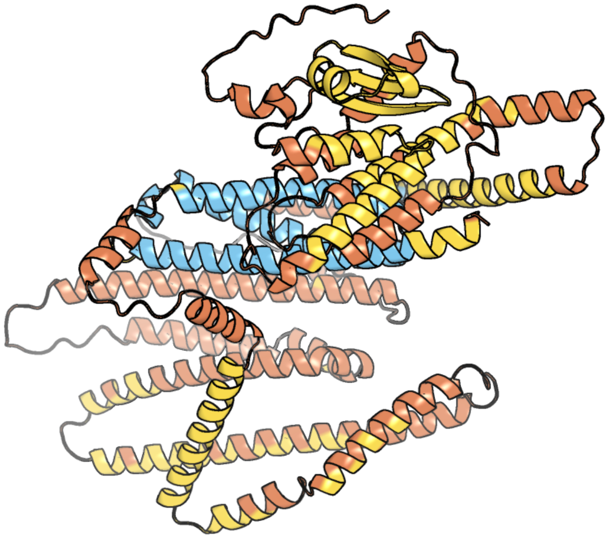

Figure 175: predicted structure of chlorv-1..175

#### chlorv-1..176

- Sequence-based annotation for chlorv-1..176 is hypothetical protein
- No significant structural hit found

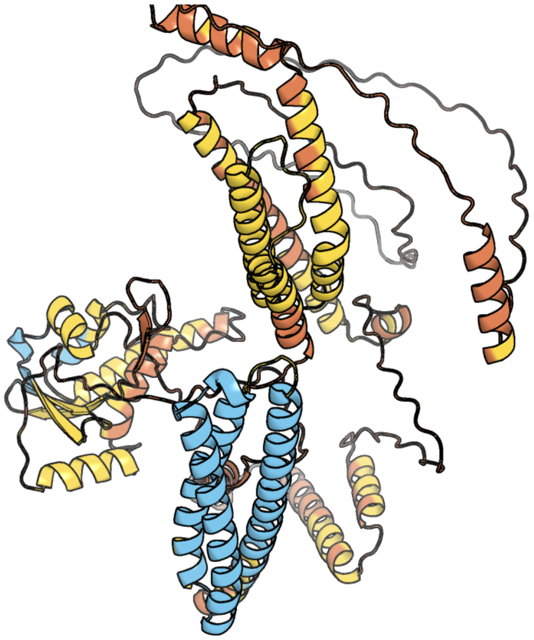

Figure 176: predicted structure of chlorv-1..176

### chlorv-1..177

- Sequence-based annotation for chlorv-1..177 is hypothetical protein
- No significant structural hit found

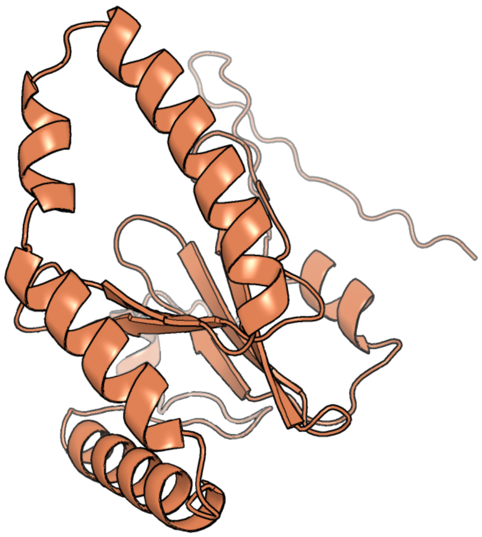

Figure 177: predicted structure of chlorv-1..177

#### chlorv-1..178

- Sequence-based annotation for chlorv-1..178 is putative Major capsid protein
- Best hit was 8rbs chain A1: Major capsid protein

| target | prob | fident | alnlen | evalue | theader |
|---|---|---|---|---|---|
| 8rbs-assembly1.cif.gz__A1 | 1 | 0.206 | 561 | 7.748e-36 | Emiliania huxleyi virus 201 (EhV-201) asymmetrical unit of capsid proteins predicted by AlphaFold2 fitted into the cryo-EM density of EhV-201 virion composite map. |
| 8h2i-assembly1.cif.gz__ab-5 | 1 | 0.235 | 530 | 1.655e-35 | Near-atomic structure of five-fold averaged PBCV-1 capsid |
| 8h2i-assembly1.cif.gz__bu | 1 | 0.229 | 593 | 2.554e-35 | Near-atomic structure of five-fold averaged PBCV-1 capsid |

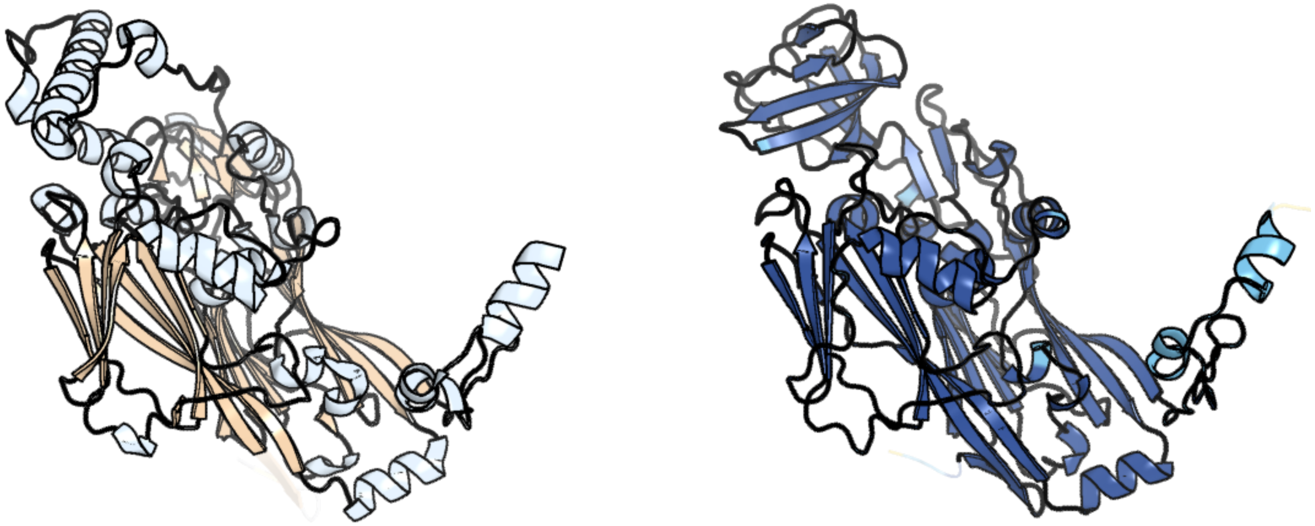

Figure 178: left: reference structure of 8rbs chain A1. right: predicted structure of chlorv-1..178, unaligned sequences are shown as transparent

#### chlorv-1..179

- Sequence-based annotation for chlorv-1..179 is putative Oxygenase
- Best hit was 3btz chain A: Alpha-ketoglutarate-dependent dioxygenase alkB homolog 2

| target | prob | fident | alnlen | evalue | theader |
|---|---|---|---|---|---|
| 3btz-assembly1.cif.gz_A | 1 | 0.211 | 199 | 4.043e-09 | Crystal structure of human ABH2 cross-linked to dsDNA |
| 3rzl-assembly1.cif.gz_A | 1 | 0.208 | 182 | 1.04e-08 | Duplex Interrogation by a Direct DNA Repair Protein in the Search of Damage |
| 3buc-assembly1.cif.gz_A | 1 | 0.194 | 190 | 1.162e-08 | X-ray structure of human ABH2 bound to dsDNA with Mn(II) and 2KG |

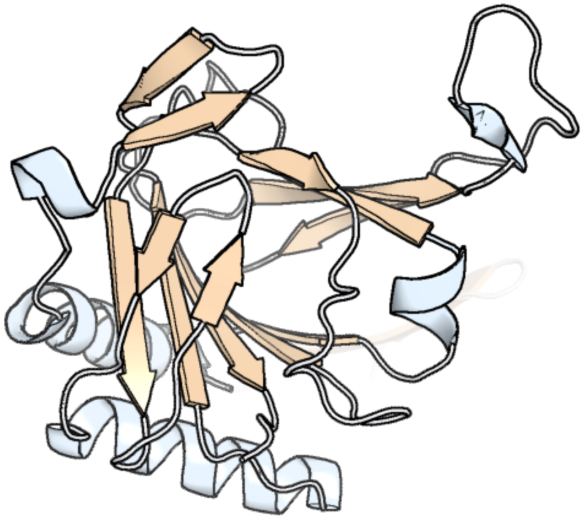

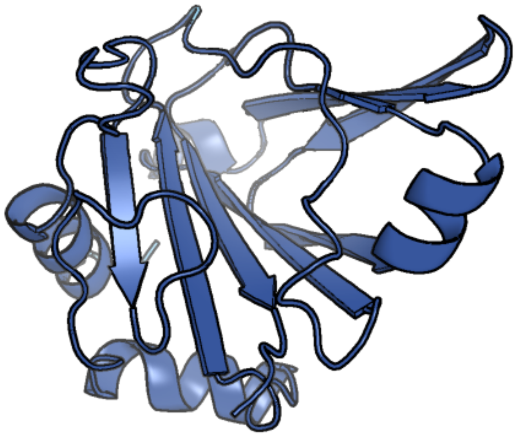

Figure 179: left: reference structure of 3btz chain A. right: predicted structure of chlorv-1..179, unaligned sequences are shown as transparent

#### chlorv-1..180

- Sequence-based annotation for chlorv-1..180 is putative Dihydrofolate reductase
- Best hit was 6n1s chain A: Bifunctional dihydrofolate reductase-thymidylate synthase

| target | prob | fident | alnlen | evalue | theader |
|---|---|---|---|---|---|
| 6n1s-assembly1.cif.gz_A | 1 | 0.335 | 515 | 2.806e-51 | Toxoplasma gondii TS-DHFR in complex with selective inhibitor 29 |
| 6kp2-assembly1.cif.gz_A | 1 | 0.326 | 515 | 3.355e-51 | Quadruple mutant plasmodium falciparum dihydrofolate reductase complexed with B10042 |
| 6kot-assembly1.cif.gz_A | 1 | 0.329 | 519 | 4.521e-51 | Quadruple mutant (N51I+C59R+S108N+I164L) plasmodium falciparum dihydrofolate reductase-thymidylate synthase (PfDHFR-TS) complexed with B12128 and NADPH |

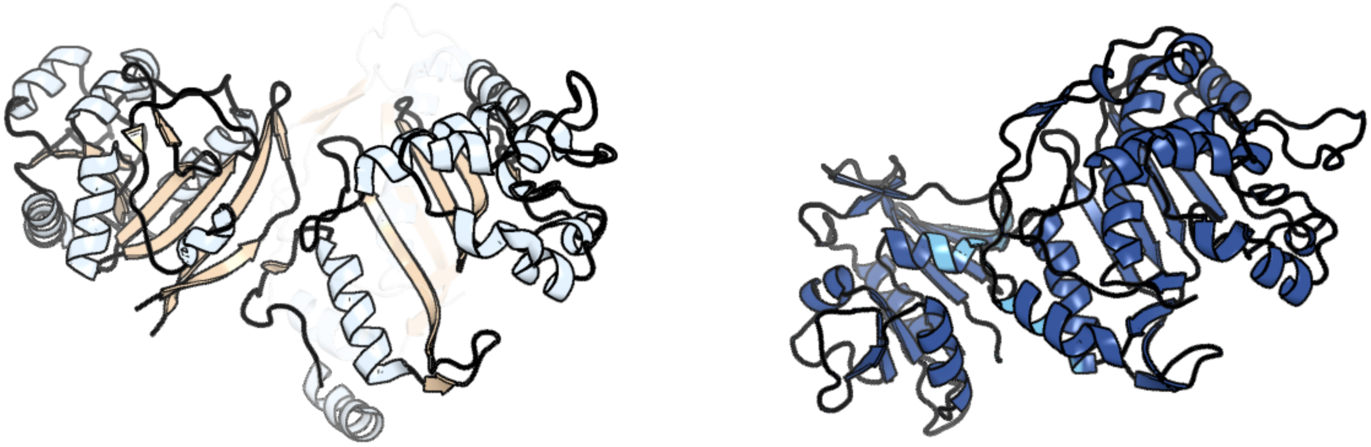

Figure 180: left: reference structure of 6n1s chain A. right: predicted structure of chlorv-1..180, unaligned sequences are shown as transparent

#### chlorv-1..181

- Sequence-based annotation for chlorv-1..181 is putative Cytidine deaminase
- No significant structural hit found

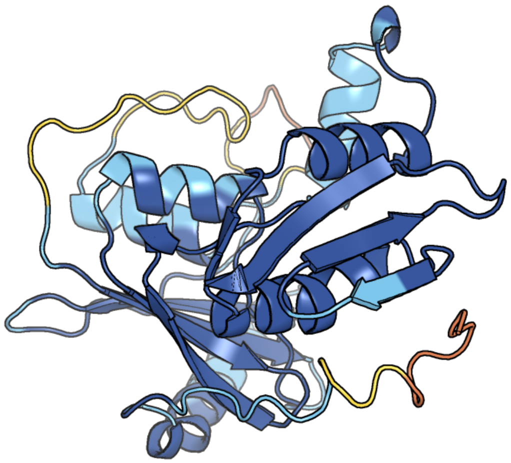

Figure 181: predicted structure of chlorv-1..181

#### chlorv-1..182

- Sequence-based annotation for chlorv-1..182 is putative Transcription initiation factor IIB
- Best hit was 7nvu chain M: Transcription initiation factor IIB

| target | prob | fident | alnlen | evalue | theader |
|---|---|---|---|---|---|
| 7nvu-assembly1.cif.gz_M | 1 | 0.238 | 298 | 6.766e-12 | RNA polymerase II core pre-initiation complex with open promoter DNA |
| 8wak-assembly1.cif.gz_R | 1 | 0.226 | 292 | 8.65e-11 | Structure of transcribing complex 2 (TC2), the initially transcribing complex with Pol II positioned 2nt downstream of TSS. |
| 8bvw-assembly1.cif.gz_M | 1 | 0.208 | 288 | 1.54e-10 | RNA polymerase II pre-initiation complex with the distal +1 nucleosome (PIC-Nuc18W) |

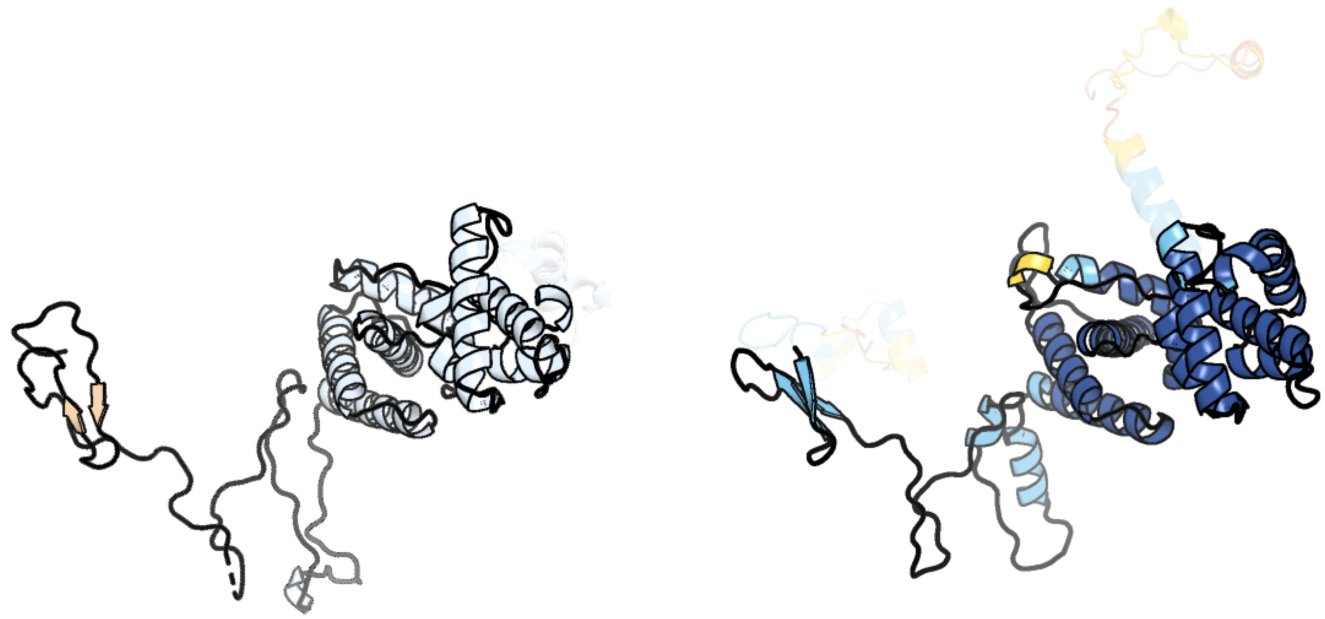

Figure 182: left: reference structure of 7nvu chain M. right: predicted structure of chlorv-1..182, unaligned sequences are shown as transparent

#### chlorv-1..183

- Sequence-based annotation for chlorv-1..183 is putative ATPase
- Best hit was 3syk chain A: Protein CbbX

| target | prob | fident | alnlen | evalue | theader |
|---|---|---|---|---|---|
| 3syk-assembly1.cif.gz_A | 1 | 0.272 | 283 | 2.024e-17 | Crystal structure of the AAA+ protein CbbX, selenomethionine structure |
| 3syl-assembly1.cif.gz_B | 1 | 0.267 | 284 | 1.165e-16 | Crystal structure of the AAA+ protein CbbX, native structure |
| 1iy0-assembly1.cif.gz_A | 1 | 0.189 | 264 | 2.255e-10 | Crystal structure of the FtsH ATPase domain with AMP-PNP from Thermus thermophilus |

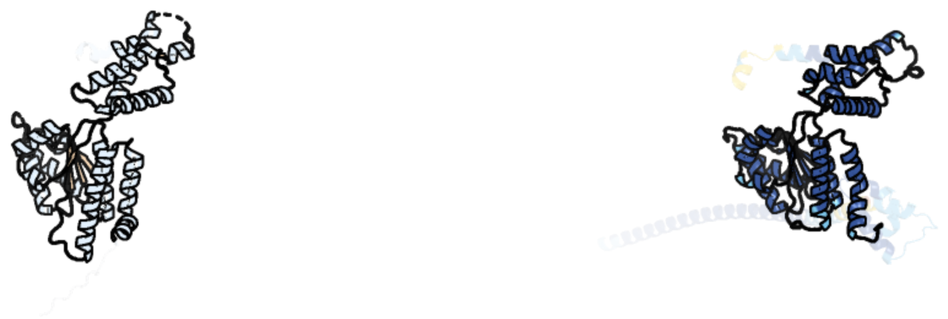

Figure 183: left: reference structure of 3syk chain A. right: predicted structure of chlorv-1..183, unaligned sequences are shown as transparent

#### chlorv-1..184

- Sequence-based annotation for chlorv-1..184 is putative Exonuclease
- Best hit was 6o3f chain B: Lysine–tRNA ligase

| target | prob | fident | alnlen | evalue | theader |
|---|---|---|---|---|---|
| 6o3f-assembly1.cif.gz__B | 1 | 0.164 | 201 | 2.793e-06 | Crystal Structure of Lysyl-tRNA Synthetase from Chlamydia trachomatis with complexed with L-lysine and a difluoro cyclohexyl chromone ligand |
| 3a74-assembly1.cif.gz__A | 1 | 0.134 | 171 | 5.04e-06 | Lysyl-tRNA synthetase from Bacillus stearothermophilus complexed with Diadenosine Tetraphosphate (AP4A) |
| 4ex5-assembly1.cif.gz__B | 1 | 0.135 | 170 | 8.081e-06 | Crystal structure of lysyl-tRNA synthetase LysRS from Burkholderia thailandensis bound to lysine |

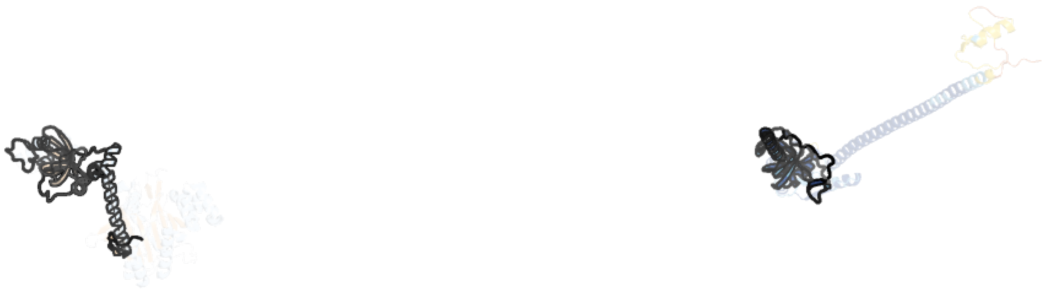

Figure 184: left: reference structure of 6o3f chain B. right: predicted structure of chlorv-1..184, unaligned sequences are shown as transparent

### chlorv-1..185

- Sequence-based annotation for chlorv-1..185 is hypothetical protein
- No significant structural hit found

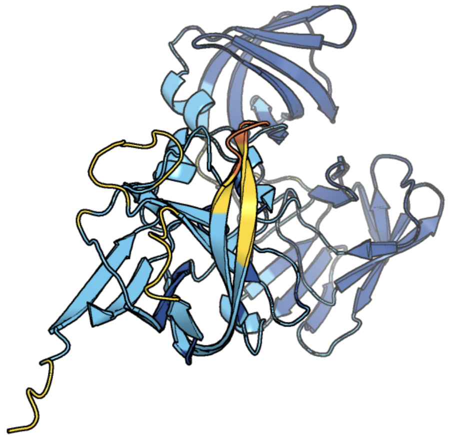

Figure 185: predicted structure of chlorv-1..185

#### chlorv-1..186

- Sequence-based annotation for chlorv-1..186 is hypothetical protein
- No significant structural hit found

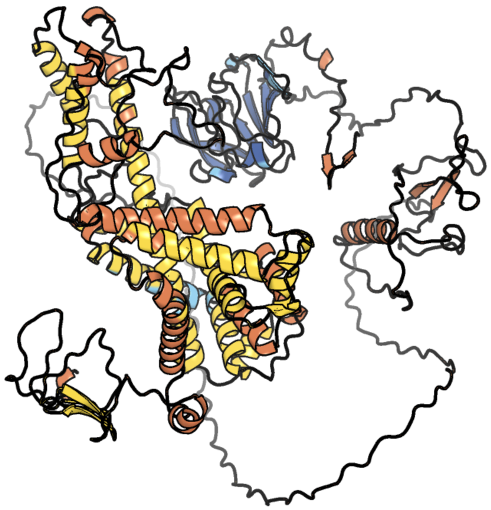

Figure 186: predicted structure of chlorv-1..186

#### chlorv-1..187

- Sequence-based annotation for chlorv-1..187 is hypothetical protein
- No significant structural hit found

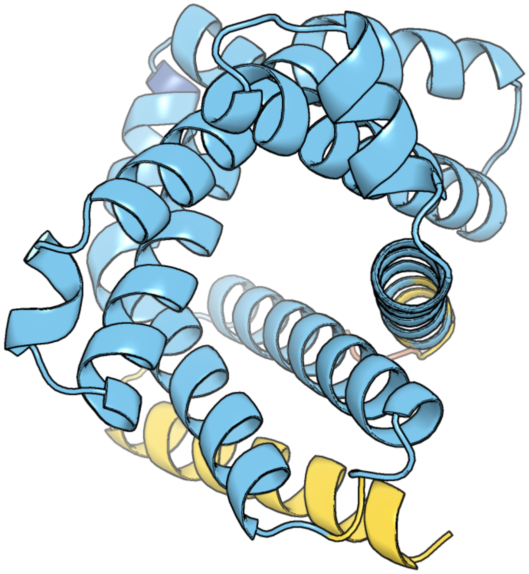

Figure 187: predicted structure of chlorv-1..187

#### chlorv-1..188

- Sequence-based annotation for chlorv-1..188 is hypothetical protein
- No significant structural hit found

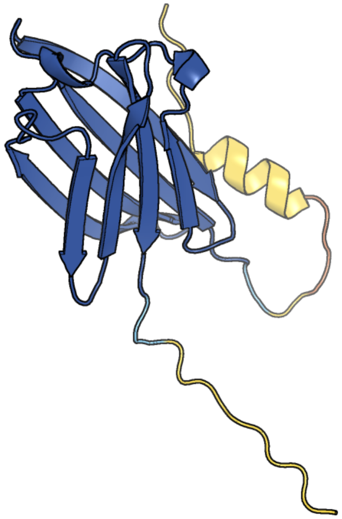

Figure 188: predicted structure of chlorv-1..188

#### chlorv-1..189

- Sequence-based annotation for chlorv-1..189 is hypothetical protein
- No significant structural hit found

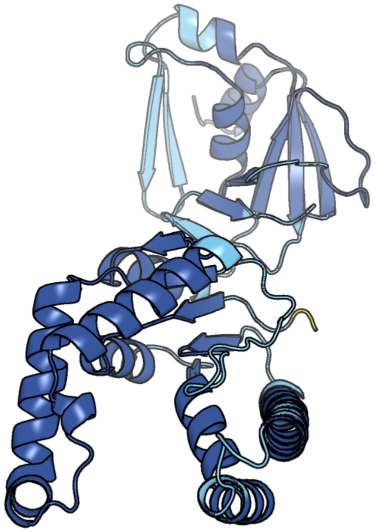

Figure 189: predicted structure of chlorv-1..189

#### chlorv-1..190

- Sequence-based annotation for chlorv-1..190 is putative Transcription factor S-II
- Best hit was 8a40 chain U: Transcription elongation factor A protein 1

| target | prob | fident | alnlen | evalue | theader |
|---|---|---|---|---|---|
| 8a40-assembly1.cif.gz_U | 1 | 0.247 | 178 | 5.394e-07 | Structure of mammalian Pol II-TFIIS elongation complex |
| 7und-assembly1.cif.gz_O | 1 | 0.224 | 178 | 1.022e-06 | Pol II-DSIF-SPT6-PAF1c-TFIIS-nucleosome complex (stalled at +38) |
| 6o9l-assembly1.cif.gz_U | 1 | 0.241 | 178 | 1.449e-06 | Human holo-PIC in the closed state |

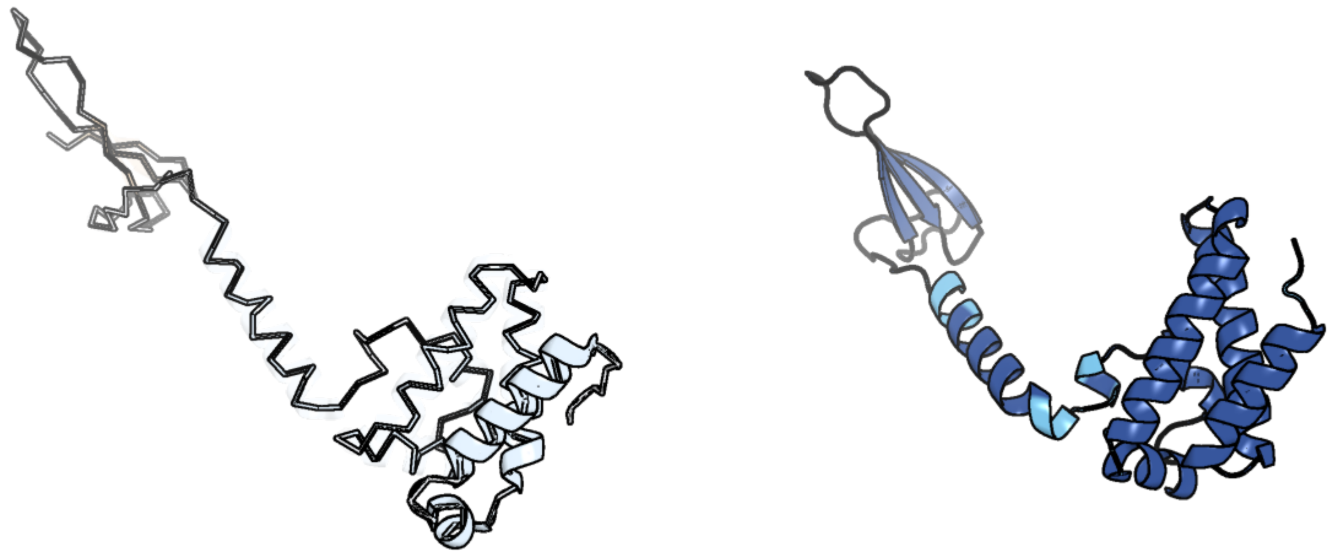

Figure 190: left: reference structure of 8a40 chain U. right: predicted structure of chlorv-1..190, unaligned sequences are shown as transparent

#### chlorv-1..191

- Sequence-based annotation for chlorv-1..191 is putative RING-finger-containing E3 ubiquitin ligase
- No significant structural hit found

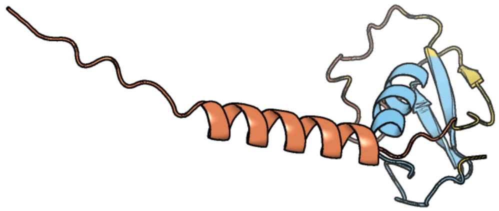

Figure 191: predicted structure of chlorv-1..191

#### chlorv-1..192

- Sequence-based annotation for chlorv-1..192 is putative Threonyl-tRNA synthetase
- Best hit was 4hwt chain A: Threonine–tRNA ligase, cytoplasmic

| target | prob | fident | alnlen | evalue | theader |
|---|---|---|---|---|---|
| 4hwt-assembly1.cif.gz_A | 1 | 0.53 | 390 | 5.12e-61 | Crystal structure of human Threonyl-tRNA synthetase bound to a novel inhibitor |
| 4p3n-assembly1.cif.gz_B | 1 | 0.533 | 390 | 1.504e-59 | Structural Basis for Full-Spectrum Inhibition of Threonyl-tRNA Synthetase by Borrelidin 1 |
| 7l3o-assembly2.cif.gz_B | 1 | 0.526 | 393 | 1.898e-58 | Crystal Structure of the RNA binding domain of Threonyl-tRNA synthetase from Cryptosporidium parvum Iowa II |

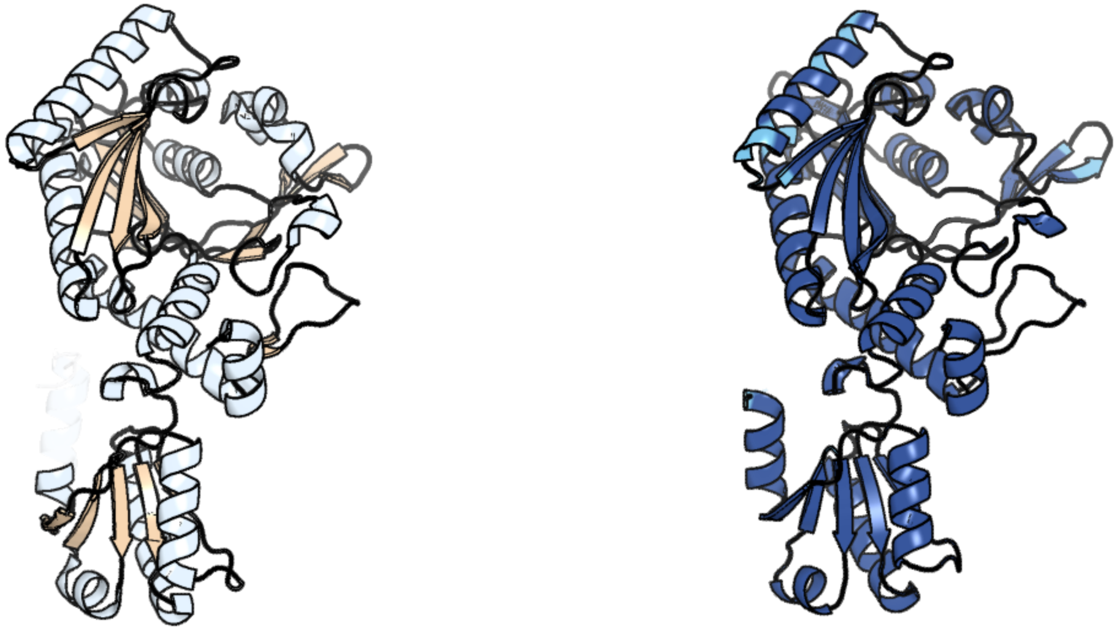

Figure 192: left: reference structure of 4hwt chain A. right: predicted structure of chlorv-1..192, unaligned sequences are shown as transparent

#### chlorv-1..193

- Sequence-based annotation for chlorv-1..193 is hypothetical protein
- No significant structural hit found

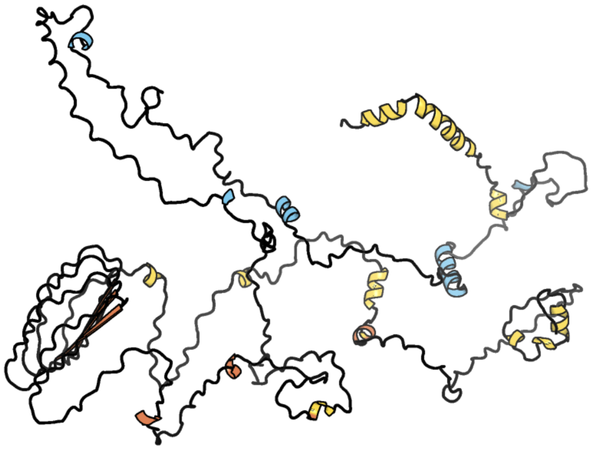

Figure 193: predicted structure of chlorv-1..193

#### chlorv-1..194

- Sequence-based annotation for chlorv-1..194 is hypothetical protein
- No significant structural hit found

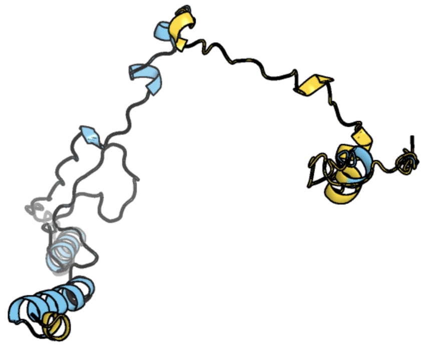

Figure 194: predicted structure of chlorv-1..194

### chlorv-1..195

- Sequence-based annotation for chlorv-1..195 is hypothetical protein
- No significant structural hit found

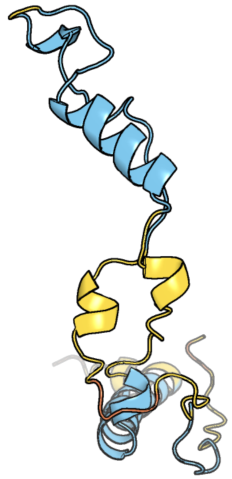

Figure 195: predicted structure of chlorv-1..195

#### chlorv-1..196

- Sequence-based annotation for chlorv-1..196 is hypothetical protein
- No significant structural hit found

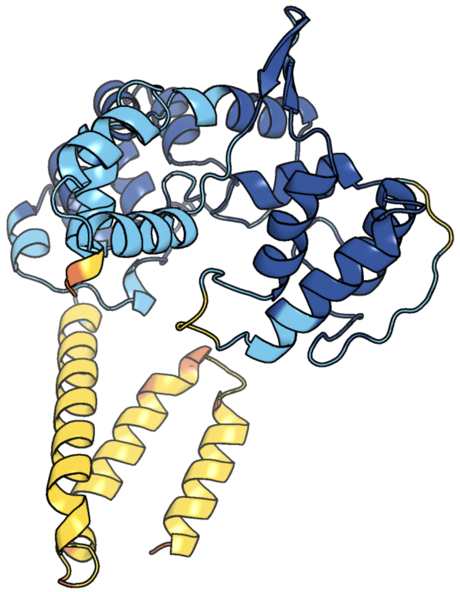

Figure 196: predicted structure of chlorv-1..196

### chlorv-1..197

- Sequence-based annotation for chlorv-1..197 is hypothetical protein
- No significant structural hit found

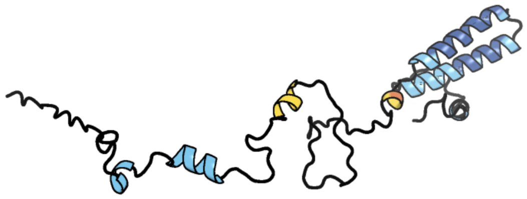

Figure 197: predicted structure of chlorv-1..197

#### chlorv-1..198

- Sequence-based annotation for chlorv-1..198 is putative Replication factor C subunit 2
- Best hit was 7z6h chain D: Replication factor C subunit 4

| target | prob | fident | alnlen | evalue | theader |
|---|---|---|---|---|---|
| 7z6h-assembly1.cif.gz_D | 1 | 0.386 | 321 | 4.777e-26 | Structure of DNA-bound human RAD17-RFC clamp loader and 9-1-1 checkpoint clamp |
| 8dqx-assembly1.cif.gz_C | 1 | 0.36 | 325 | 2.663e-24 | Open state of RFC:PCNA bound to a 3' ss/dsDNA junction |
| 1sxj-assembly1.cif.gz_C | 1 | 0.354 | 327 | 3.786e-24 | Crystal Structure of the Eukaryotic Clamp Loader (Replication Factor C, RFC) Bound to the DNA Sliding Clamp (Proliferating Cell Nuclear Antigen, PCNA) |

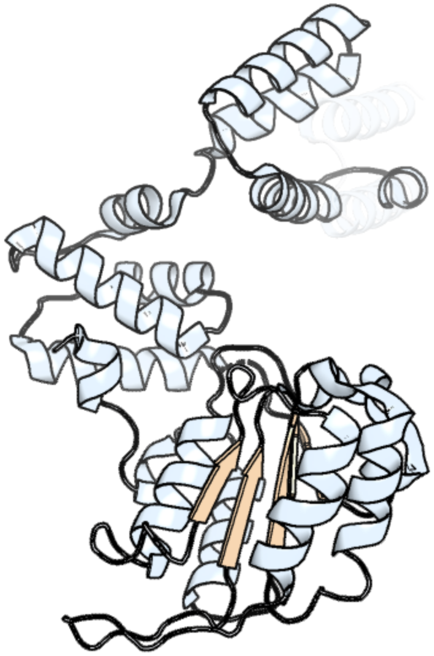

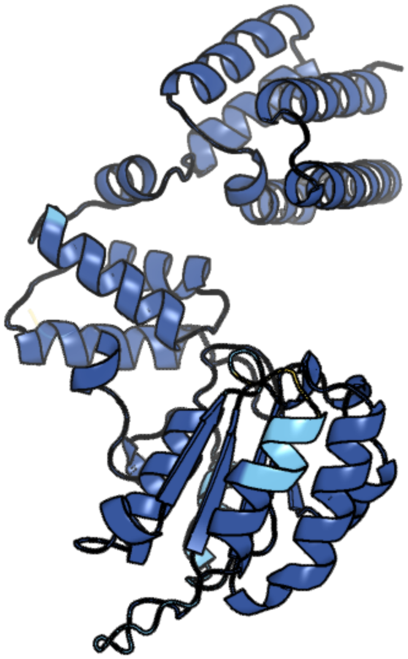

Figure 198: left: reference structure of 7z6h chain D. right: predicted structure of chlorv-1..198, unaligned sequences are shown as transparent

#### chlorv-1..199

- Sequence-based annotation for chlorv-1..199 is hypothetical protein
- No significant structural hit found

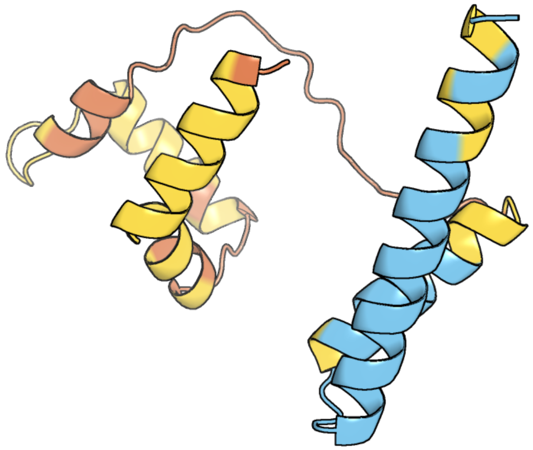

Figure 199: predicted structure of chlorv-1..199

#### chlorv-1..200

- Sequence-based annotation for chlorv-1..200 is putative DNA topoisomerase I
- Best hit was 1k4t chain A: DNA topoisomerase I

| target | prob | fident | alnlen | evalue | theader |
|---|---|---|---|---|---|
| 1k4t-assembly1.cif.gz_A | 1 | 0.365 | 563 | 3.318e-50 | HUMAN DNA TOPOISOMERASE I (70 KDA) IN COMPLEX WITH THE POISON TOPOTECAN AND COVALENT COMPLEX WITH A 22 BASE PAIR DNA DUPLEX |
| 1seu-assembly1.cif.gz_A | 1 | 0.367 | 563 | 2.532e-48 | Human DNA Topoisomerase I (70 Kda) In Complex With The Indolocarbazole SA315F and Covalent Complex With A 22 Base Pair DNA Duplex |
| 1lpq-assembly1.cif.gz_A | 1 | 0.371 | 560 | 2.064e-45 | Human DNA Topoisomerase I (70 Kda) In Non-Covalent Complex With A 22 Base Pair DNA Duplex Containing an 8-oxoG Lesion |

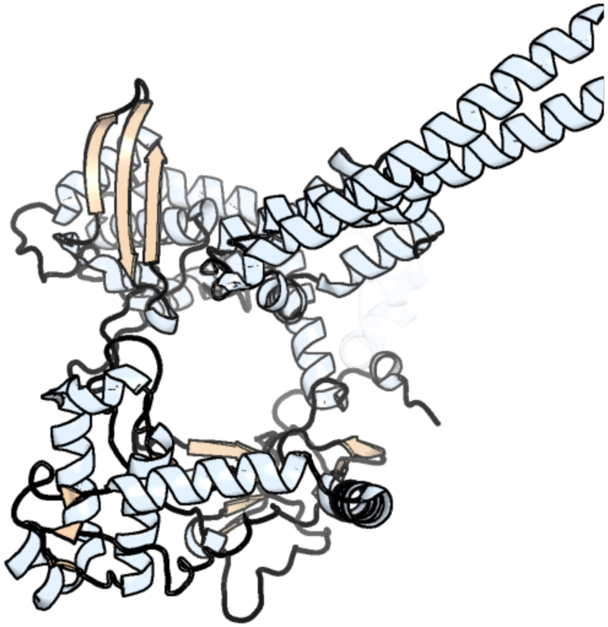

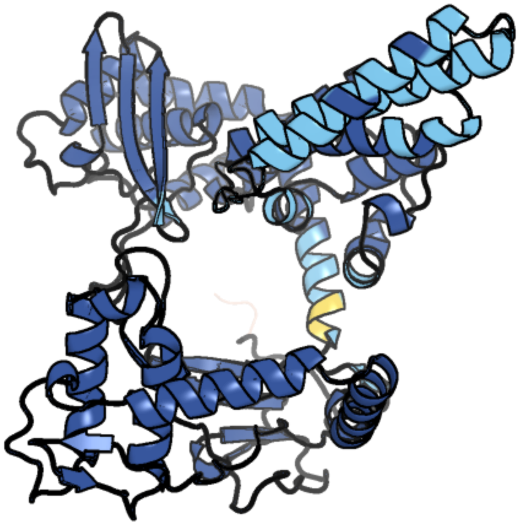

Figure 200: left: reference structure of 1k4t chain A. right: predicted structure of chlorv-1..200, unaligned sequences are shown as transparent

#### chlorv-1..201

- Sequence-based annotation for chlorv-1..201 is hypothetical protein
- No significant structural hit found

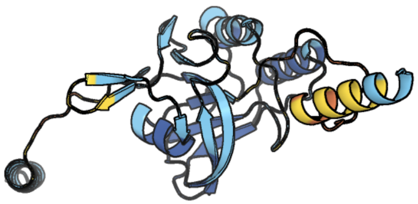

Figure 201: predicted structure of chlorv-1..201

#### chlorv-1..202

- Sequence-based annotation for chlorv-1..202 is putative RNA polymerase subunit alpha
- Best hit was 7okx chain A: DNA-directed RNA polymerase II subunit RPB1

| target | prob | fident | alnlen | evalue | theader |
|---|---|---|---|---|---|
| 8jh4-assembly1.cif.gz_A | 1 | 0.289 | 1499 | 0 | RNA polymerase II elongation complex containing 60 bp upstream DNA loop, stalled at SHL(-1) of the nucleosome |
| 8h0v-assembly1.cif.gz_A | 1 | 0.289 | 1511 | 0 | RNA polymerase II transcribing a chromatosome (type I) |
| 7okx-assembly1.cif.gz_A | 1 | 0.295 | 1516 | 0 | Structure of active transcription elongation complex Pol II-DSIF (SPT5-KOW5)-ELL2-EAF1 (composite structure) |

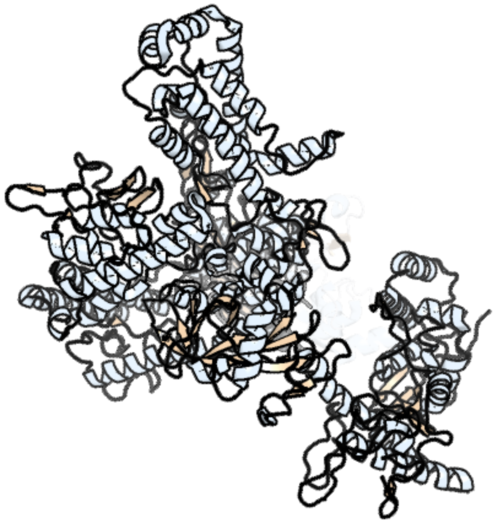

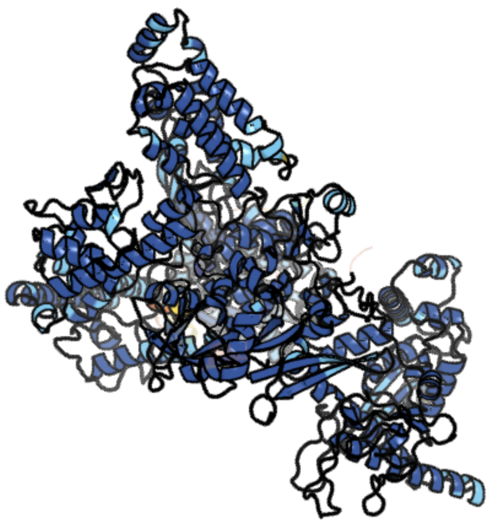

Figure 202: left: reference structure of 7okx chain A. right: predicted structure of chlorv-1..202, unaligned sequences are shown as transparent

#### chlorv-1..203

- Sequence-based annotation for chlorv-1..203 is hypothetical protein
- No significant structural hit found

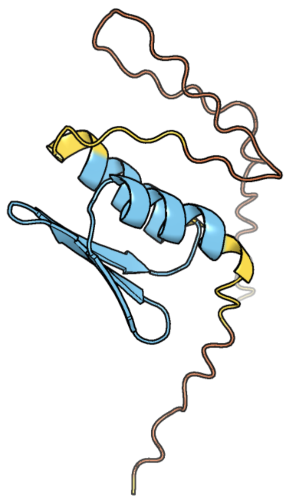

Figure 203: predicted structure of chlorv-1..203

#### chlorv-1..204

- Sequence-based annotation for chlorv-1..204 is putative Peptidase
- Best hit was 4pf9 chain A: Insulin-degrading enzyme

| target | prob | fident | alnlen | evalue | theader |
|---|---|---|---|---|---|
| 4pf9-assembly1.cif.gz_A | 1 | 0.202 | 935 | 6.53e-52 | Crystal structure of insulin degrading enzyme complexed with inhibitor |
| 7k1f-assembly1.cif.gz_B | 1 | 0.203 | 929 | 1.28e-51 | Crystal structure of human insulin degrading enzyme (IDE) in complex with compound BDM_88558 |
| 4nxo-assembly1.cif.gz_A | 1 | 0.2 | 932 | 1.88e-51 | Crystal Structure of Insulin Degrading Enzyme in complex with BDM44768 |

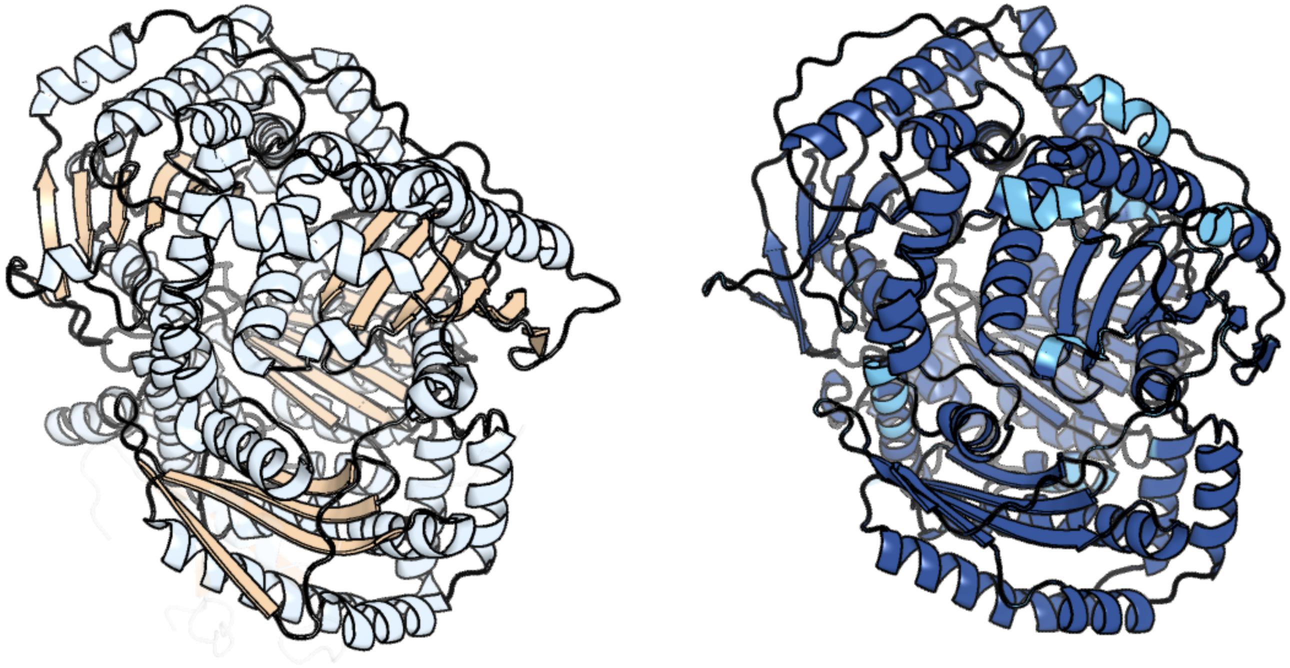

Figure 204: left: reference structure of 4pf9 chain A. right: predicted structure of chlorv-1..204, unaligned sequences are shown as transparent

#### chlorv-1..205

- Sequence-based annotation for chlorv-1..205 is hypothetical protein
- No significant structural hit found

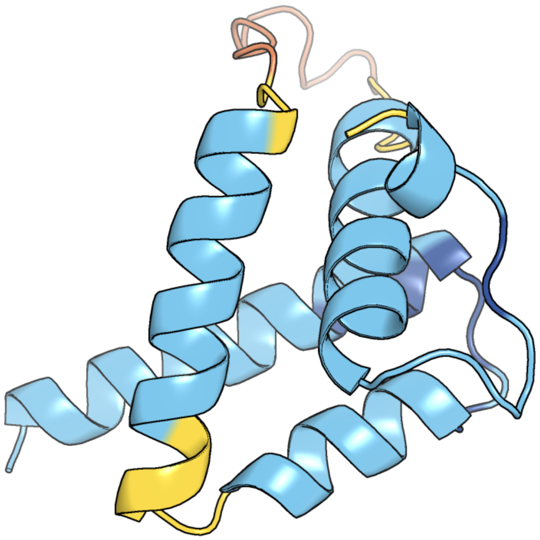

Figure 205: predicted structure of chlorv-1..205

#### chlorv-1..206

- Sequence-based annotation for chlorv-1..206 is putative Hsp70 protein
- Best hit was 7n1r chain A: Endoplasmic reticulum chaperone BiP

| target | prob | fident | alnlen | evalue | theader |
|---|---|---|---|---|---|
| 7n1r-assembly1.cif.gz_A | 1 | 0.162 | 597 | 4.163e-25 | A novel and unique ATP hydrolysis to AMP by a human Hsp70 BiP |
| 4b9q-assembly2.cif.gz_B | 1 | 0.17 | 651 | 2.349e-24 | Open conformation of ATP-bound Hsp70 homolog DnaK |
| 4b9q-assembly4.cif.gz_D | 1 | 0.176 | 639 | 3.573e-24 | Open conformation of ATP-bound Hsp70 homolog DnaK |

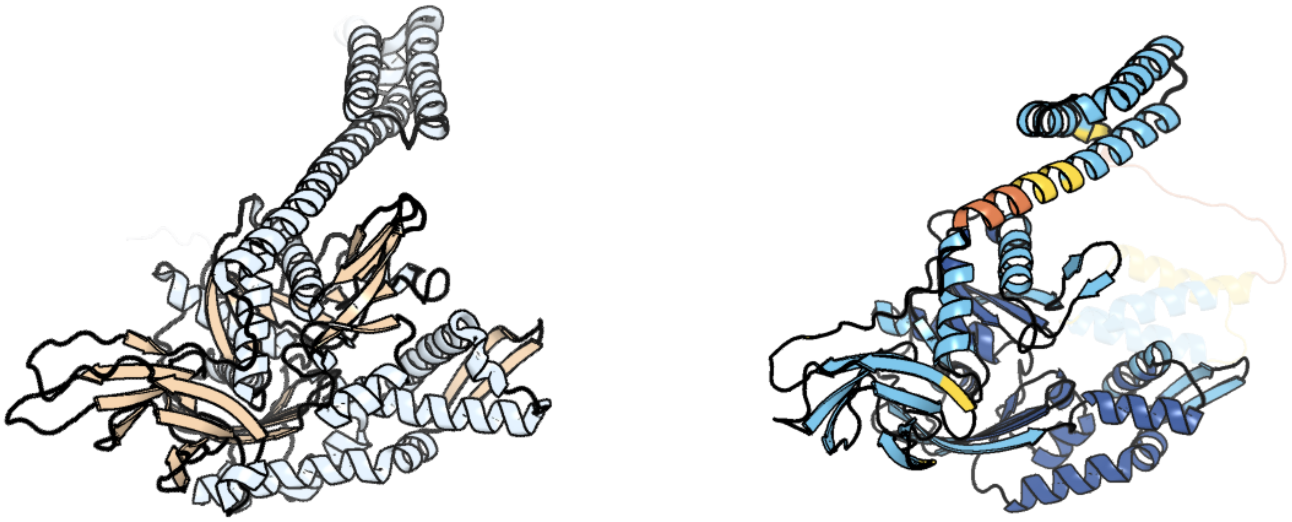

Figure 206: left: reference structure of 7n1r chain A. right: predicted structure of chlorv-1..206, unaligned sequences are shown as transparent

#### chlorv-1..207

- Sequence-based annotation for chlorv-1..207 is hypothetical protein
- No significant structural hit found

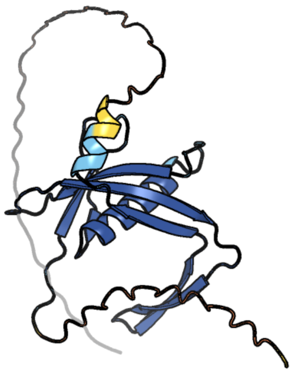

Figure 207: predicted structure of chlorv-1..207

#### chlorv-1..208

- Sequence-based annotation for chlorv-1..208 is putative Peptidase
- Best hit was 6on2 chain C: ATP-dependent protease La

| target | prob | fident | alnlen | evalue | theader |
|---|---|---|---|---|---|
| 6on2-assembly1.cif.gz_C | 1 | 0.279 | 551 | 4.504e-39 | Lon Protease from Yersinia pestis with Y2853 substrate |
| 7p0m-assembly1.cif.gz_B | 1 | 0.25 | 562 | 1.368e-38 | Human mitochondrial Lon protease with substrate in the ATPase and protease domains |
| 7p0m-assembly1.cif.gz_A | 1 | 0.255 | 555 | 1.433e-38 | Human mitochondrial Lon protease with substrate in the ATPase and protease domains |

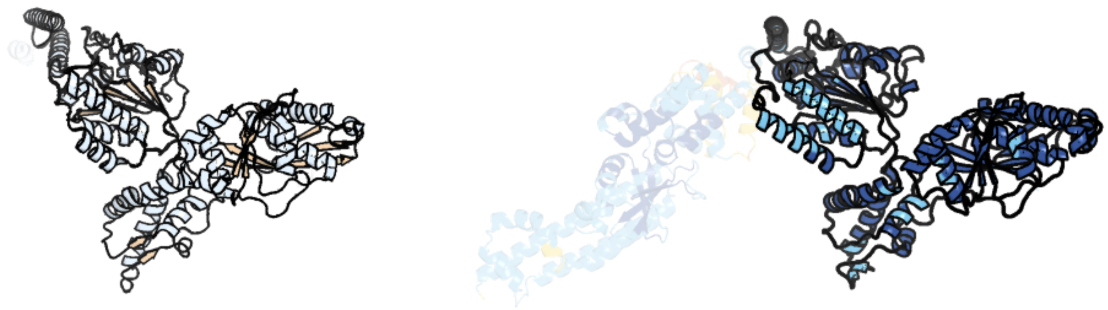

Figure 208: left: reference structure of 6on2 chain C. right: predicted structure of chlorv-1..208, unaligned sequences are shown as transparent

### chlorv-1..209

- Sequence-based annotation for chlorv-1..209 is hypothetical protein
- No significant structural hit found

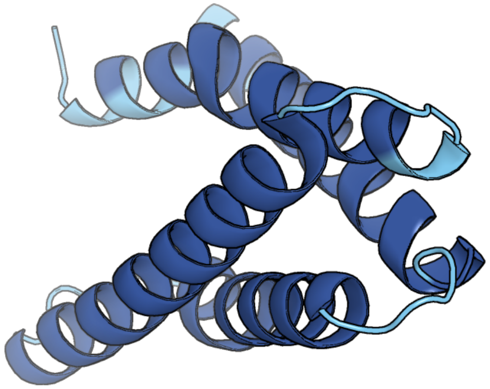

Figure 209: predicted structure of chlorv-1..209

### chlorv-1..210

- Sequence-based annotation for chlorv-1..210 is hypothetical protein
- No significant structural hit found

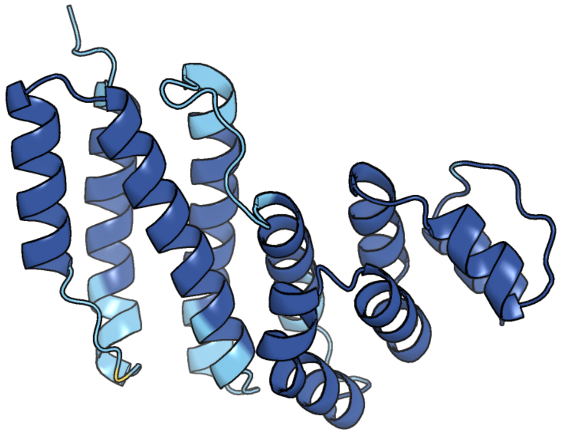

Figure 210: predicted structure of chlorv-1..210

#### chlorv-1..211

- Sequence-based annotation for chlorv-1..211 is hypothetical protein
- No significant structural hit found

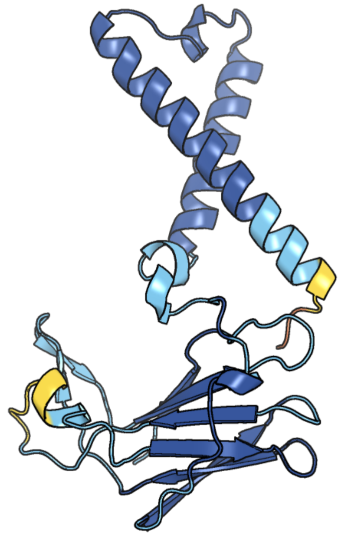

Figure 211: predicted structure of chlorv-1..211

### chlorv-1..212

- Sequence-based annotation for chlorv-1..212 is putative Glycosyltransferase
- Best hit was 7mi0 chain A: Glycosyltransferase

| target | prob | fident | alnlen | evalue | theader |
|---|---|---|---|---|---|
| 7mi0-assembly1.cif.gz_A | 1 | 0.132 | 453 | 3.668e-11 | Crystal Structure of Glycosyltransferase from Rickettsia africae ESF-5 |
| 3l01-assembly2.cif.gz_B | 1 | 0.094 | 496 | 6.756e-11 | Crystal structure of monomeric glycogen synthase from Pyrococcus abyssi |
| 3c4q-assembly1.cif.gz_A | 1 | 0.125 | 448 | 7.549e-11 | Structure of the retaining glycosyltransferase MshA : The first step in mycothiol biosynthesis. Organism : Corynebacterium glutamicum- Complex with UDP |

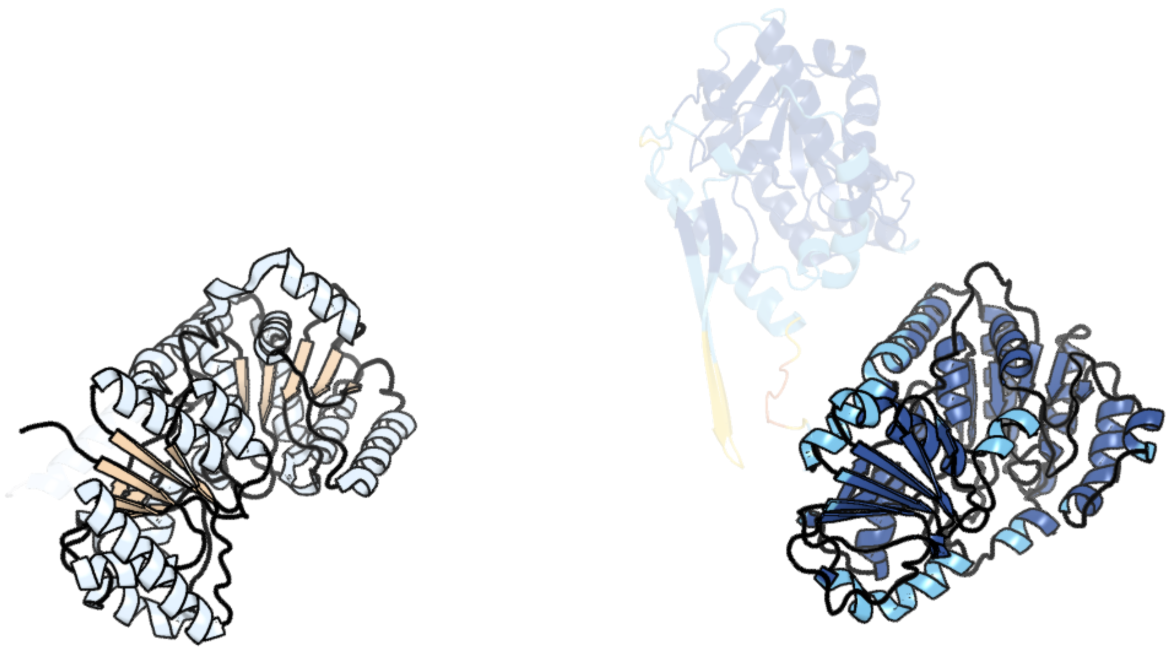

Figure 212: left: reference structure of 7mi0 chain A. right: predicted structure of chlorv-1..212, unaligned sequences are shown as transparent

#### chlorv-1..213

- Sequence-based annotation for chlorv-1..213 is hypothetical protein
- No significant structural hit found

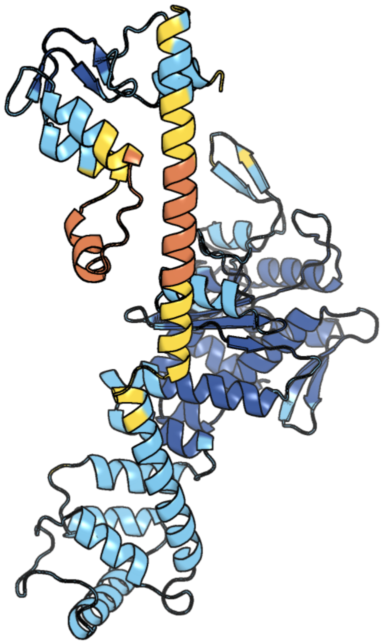

Figure 213: predicted structure of chlorv-1..213

#### chlorv-1..214

- Sequence-based annotation for chlorv-1..214 is hypothetical protein
- No significant structural hit found

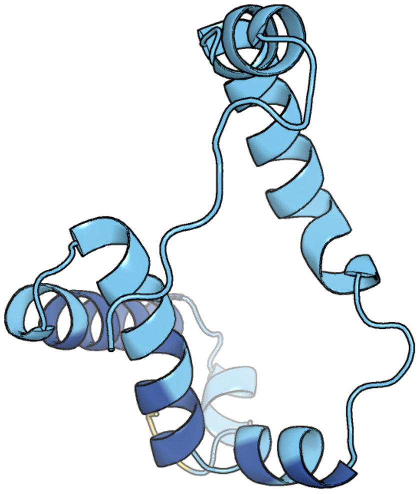

Figure 214: predicted structure of chlorv-1..214

#### chlorv-1..215

- Sequence-based annotation for chlorv-1..215 is putative Thioredoxin
- Best hit was 3vww chain A: Protein disulfide-isomerase A6

| target | prob | fident | alnlen | evalue | theader |
|---|---|---|---|---|---|
| 3vww-assembly1.cif.gz_A | 1 | 0.3 | 90 | 2.137e-07 | Crystal structure of a0-domain of P5 from H. sapiens |
| 1x5d-assembly1.cif.gz_A | 1 | 0.285 | 91 | 2.435e-07 | The solution structure of the second thioredoxin-like domain of human Protein disulfide-isomerase A6 |
| 3wge-assembly1.cif.gz_A | 1 | 0.275 | 87 | 6.062e-07 | Crystal structure of ERp46 Trx2 |

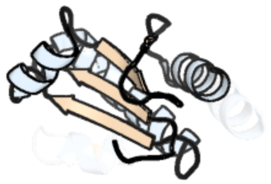

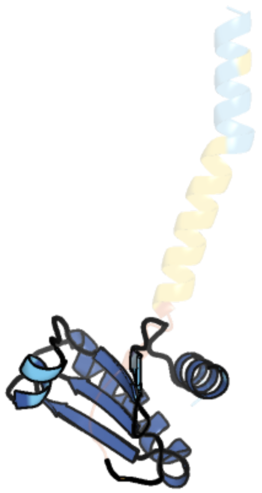

Figure 215: left: reference structure of 3vww chain A. right: predicted structure of chlorv-1..215, unaligned sequences are shown as transparent

#### chlorv-1..216

- Sequence-based annotation for chlorv-1..216 is hypothetical protein
- No significant structural hit found

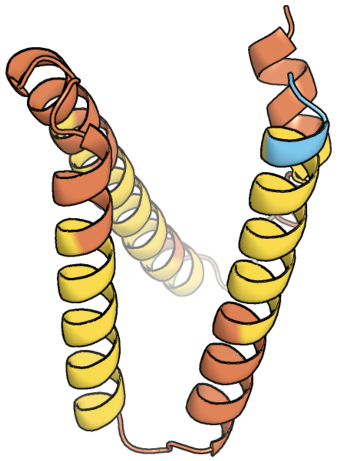

Figure 216: predicted structure of chlorv-1..216

#### chlorv-1..217

- Sequence-based annotation for chlorv-1..217 is putative Peptidase
- Best hit was 5cvm chain A: Ubiquitin carboxyl-terminal hydrolase 46

| target | prob | fident | alnlen | evalue | theader |
|---|---|---|---|---|---|
| 5cvm-assembly1.cif.gz_A | 1 | 0.239 | 309 | 1.671e-24 | USP46~ubiquitin BEA covalent complex |
| 5cvo-assembly2.cif.gz_E | 1 | 0.242 | 321 | 1.878e-24 | WDR48:USP46~ubiquitin ternary complex |
| 6dgf-assembly1.cif.gz_A | 1 | 0.229 | 327 | 5.384e-24 | Ubiquitin Variant bound to USP2 |

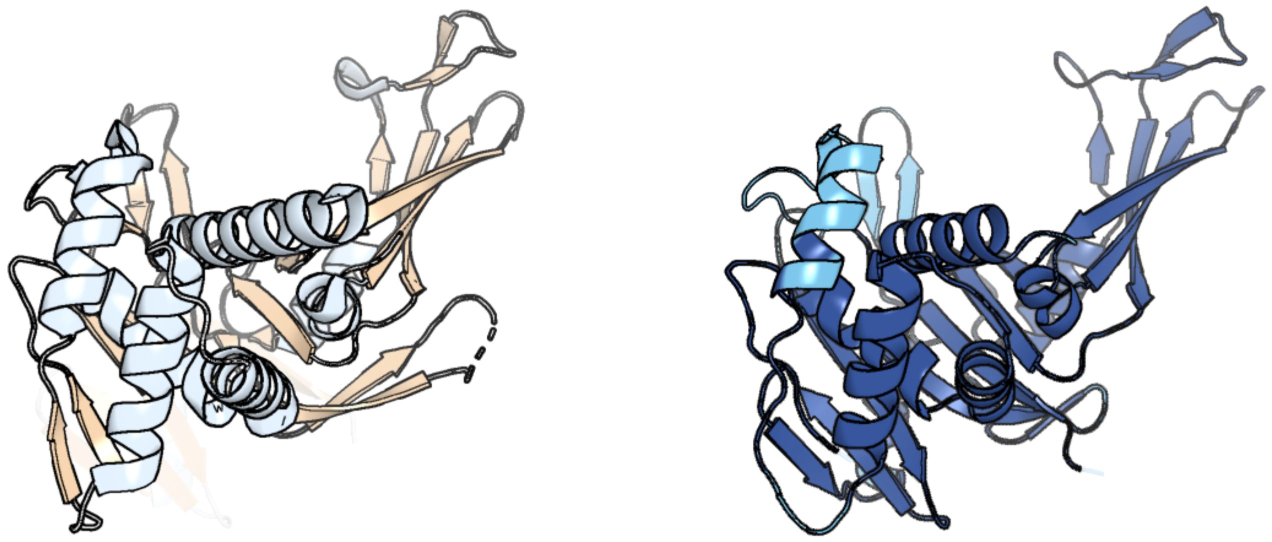

Figure 217: left: reference structure of 5cvm chain A. right: predicted structure of chlorv-1..217, unaligned sequences are shown as transparent

#### chlorv-1..218

- Sequence-based annotation for chlorv-1..218 is hypothetical protein
- No significant structural hit found

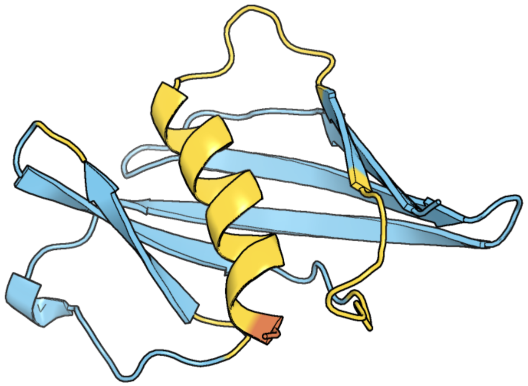

Figure 218: predicted structure of chlorv-1..218

#### chlorv-1..219

- Sequence-based annotation for chlorv-1..219 is putative Nuclease
- Best hit was 7dcy chain A: Ribonuclease R

| target | prob | fident | alnlen | evalue | theader |
|---|---|---|---|---|---|
| 7dcy-assembly1.cif.gz_A | 1 | 0.18 | 660 | 3.424e-25 | Apo form of Mycoplasma genitalium RNase R |
| 7did-assembly1.cif.gz_A | 1 | 0.175 | 665 | 1.639e-24 | Mycoplasma genitalium RNase R in complex with ribose methylated single-stranded RNA |
| 7tuv-assembly1.cif.gz_A | 1 | 0.149 | 582 | 5.873e-23 | Crystal structure of the exoribonucleolytic module of T. brucei RRP44 |

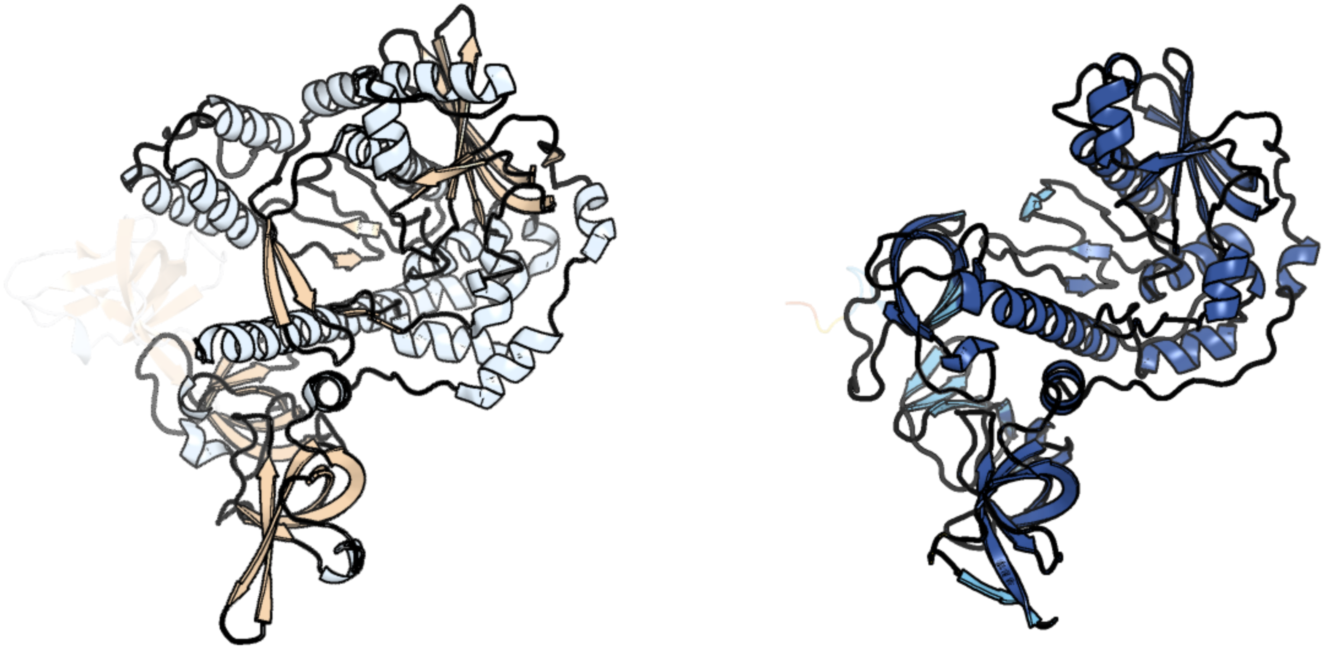

Figure 219: left: reference structure of 7dcy chain A. right: predicted structure of chlorv-1..219, unaligned sequences are shown as transparent

#### chlorv-1..220

- Sequence-based annotation for chlorv-1..220 is hypothetical protein
- No significant structural hit found

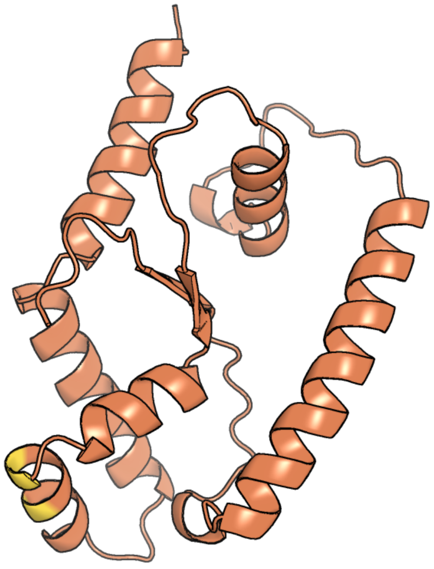

Figure 220: predicted structure of chlorv-1..220

#### chlorv-1..221

- Sequence-based annotation for chlorv-1..221 is hypothetical protein
- No significant structural hit found

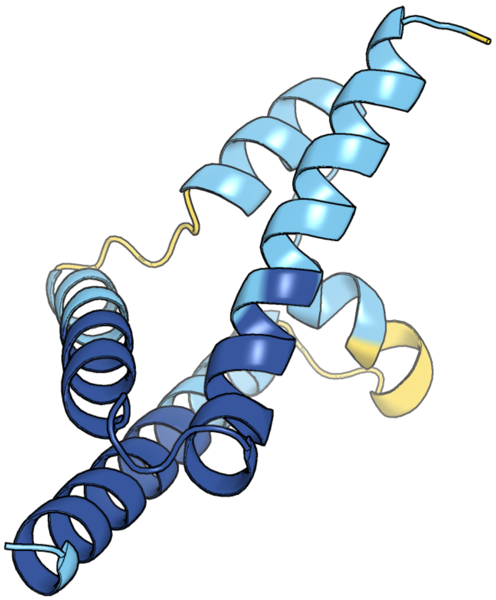

Figure 221: predicted structure of chlorv-1..221

### chlorv-1..222

- Sequence-based annotation for chlorv-1..222 is hypothetical protein
- No significant structural hit found

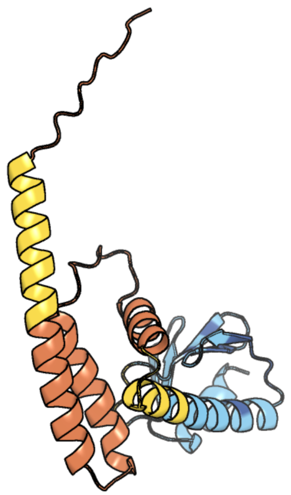

Figure 222: predicted structure of chlorv-1..222

### chlorv-1..223

- Sequence-based annotation for chlorv-1..223 is hypothetical protein
- No significant structural hit found

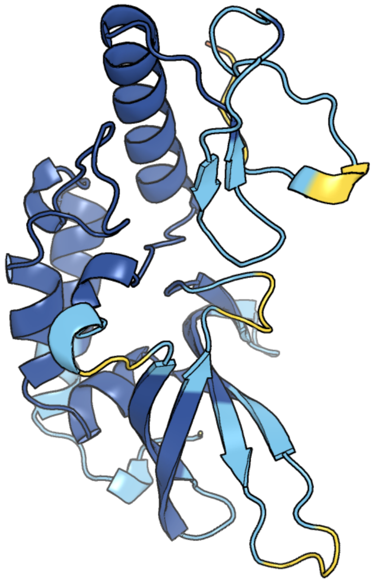

Figure 223: predicted structure of chlorv-1..223

### chlorv-1..224

- Sequence-based annotation for chlorv-1..224 is putative Photolyase
- Best hit was 4dja chain A: Photolyase

| target | prob | fident | alnlen | evalue | theader |
|---|---|---|---|---|---|
| 4dja-assembly1.cif.gz_A | 1 | 0.257 | 474 | 3.081e-27 | Crystal structure of a prokaryotic (6-4) photolyase PhrB from Agrobacterium Tumefaciens with an Fe-S cluster and a 6,7-dimethyl-8-ribityllumazine antenna chromophore at 1.45A resolution |
| 5kcm-assembly2.cif.gz_B | 1 | 0.256 | 479 | 5.129e-27 | Crystal structure of iron-sulfur cluster containing photolyase PhrB mutant I51W |
| 5lfa-assembly1.cif.gz_A | 1 | 0.253 | 469 | 6.618e-27 | Crystal structure of iron-sulfur cluster containing bacterial (6-4) photolyase PhrB - Y424F mutant with impaired DNA repair activity |

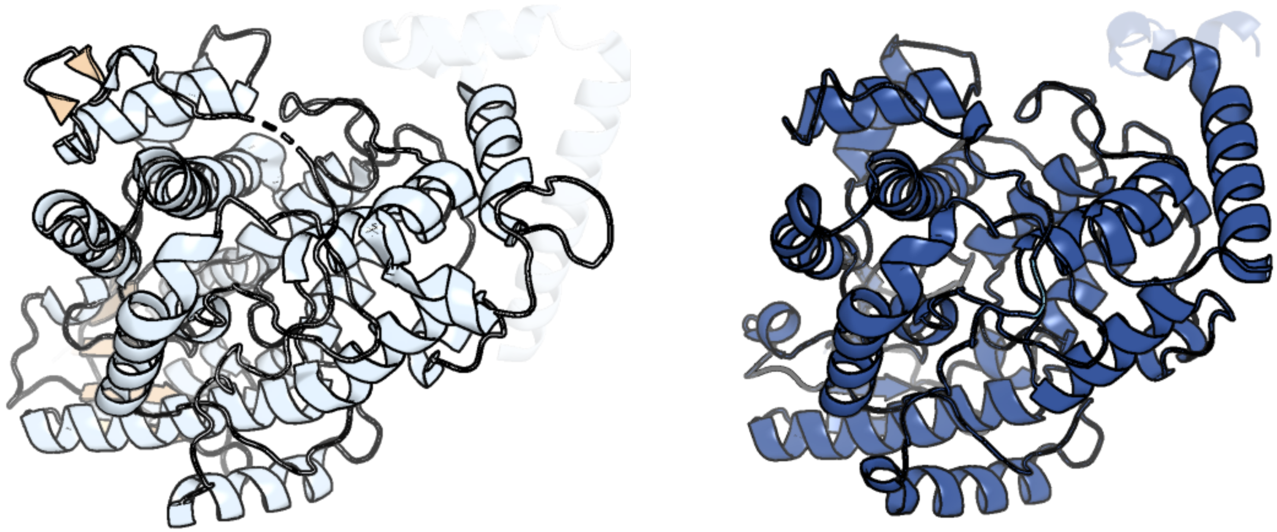

Figure 224: left: reference structure of 4dja chain A. right: predicted structure of chlorv-1..224, unaligned sequences are shown as transparent

#### chlorv-1..225

- Sequence-based annotation for chlorv-1..225 is putative ABC transporter
- Best hit was 5zxd chain B: ATP-binding cassette sub-family F member 1

| target | prob | fident | alnlen | evalue | theader |
|---|---|---|---|---|---|
| 5zxd-assembly2.cif.gz_B | 1 | 0.309 | 539 | 8.39e-41 | Crystal structure of ATP-bound human ABCF1 |
| 5zxd-assembly1.cif.gz_A | 1 | 0.284 | 545 | 6.451e-38 | Crystal structure of ATP-bound human ABCF1 |
| 4fin-assembly1.cif.gz_A | 1 | 0.24 | 562 | 8.462e-36 | Crystal Structure of EttA (formerly YjjK) - an E. coli ABC-type ATPase |

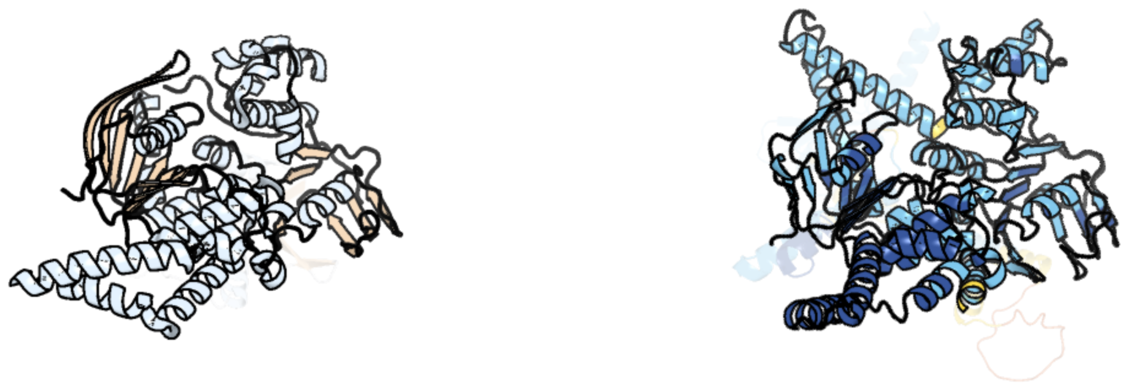

Figure 225: left: reference structure of 5zxd chain B. right: predicted structure of chlorv-1..225, unaligned sequences are shown as transparent

#### chlorv-1..226

- Sequence-based annotation for chlorv-1..226 is putative Helicase
- Best hit was 6ro1 chain A: Exosome RNA helicase MTR4

| target | prob | fident | alnlen | evalue | theader |
|---|---|---|---|---|---|
| 6ro1-assembly1.cif.gz_A | 1 | 0.252 | 923 | 2.082e-49 | X-ray crystal structure of the MTR4 NVL complex |
| 2xgj-assembly2.cif.gz_B | 1 | 0.279 | 806 | 2.512e-49 | Structure of Mtr4, a DExH helicase involved in nuclear RNA processing and surveillance |
| 4u4c-assembly1.cif.gz_A | 1 | 0.251 | 939 | 3.485e-47 | The molecular architecture of the TRAMP complex reveals the organization and interplay of its two catalytic activities |

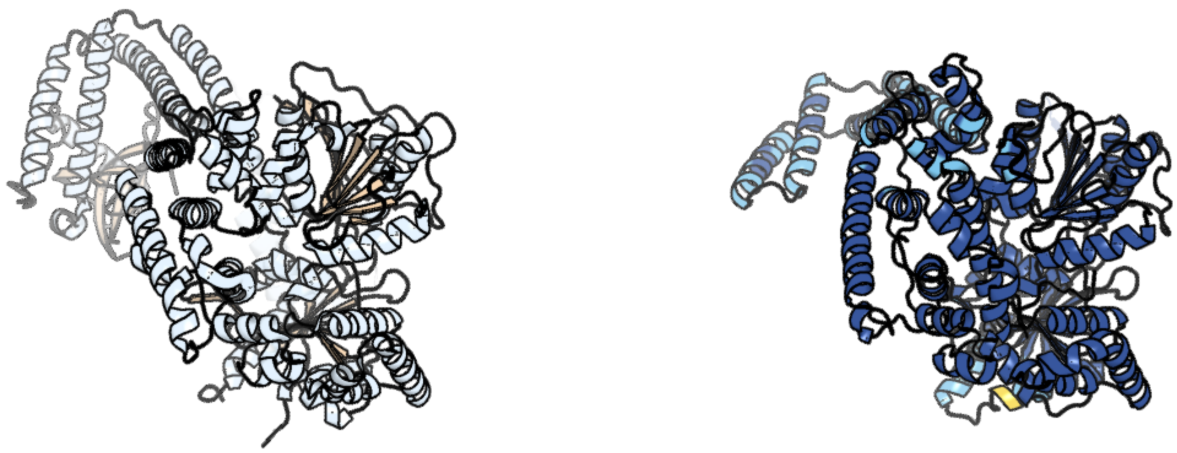

Figure 226: left: reference structure of 6ro1 chain A. right: predicted structure of chlorv-1..226, unaligned sequences are shown as transparent

#### chlorv-1..227

- Sequence-based annotation for chlorv-1..227 is hypothetical protein
- No significant structural hit found

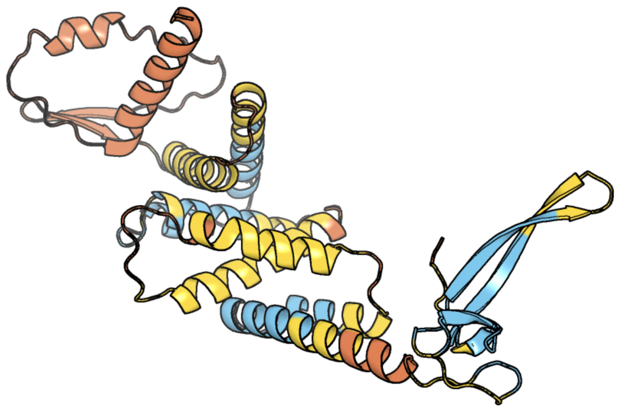

Figure 227: predicted structure of chlorv-1..227

#### chlorv-1..228

- Sequence-based annotation for chlorv-1..228 is putative Cysteine peptidase (DUF1796)
- No significant structural hit found

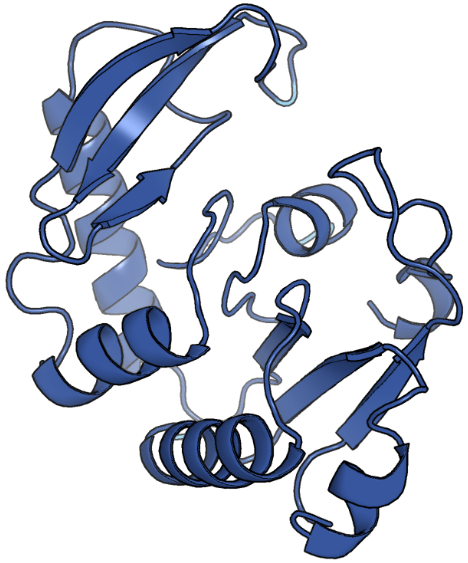

Figure 228: predicted structure of chlorv-1..228

#### chlorv-1..229

- Sequence-based annotation for chlorv-1..229 is hypothetical protein
- No significant structural hit found

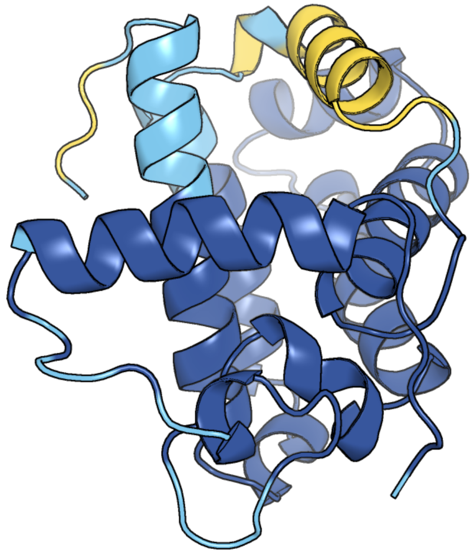

Figure 229: predicted structure of chlorv-1..229

### chlorv-1..230

- Sequence-based annotation for chlorv-1..230 is hypothetical protein
- No significant structural hit found

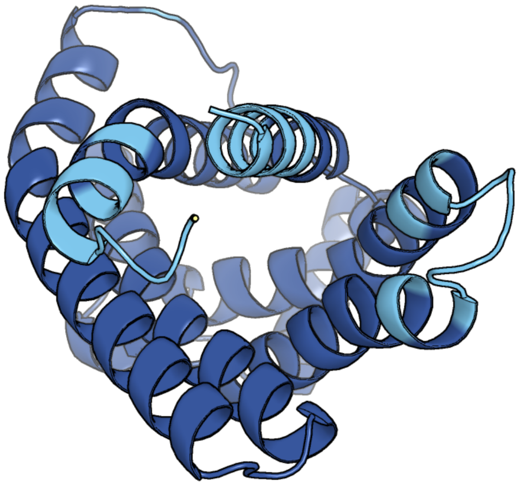

Figure 230: predicted structure of chlorv-1..230

#### chlorv-1..231

- Sequence-based annotation for chlorv-1..231 is hypothetical protein
- Best hit was 5dku chain B: Prex DNA polymerase

| target | prob | fident | alnlen | evalue | theader |
|---|---|---|---|---|---|
| 5dku-assembly2.cif.gz_B | 1 | 0.189 | 327 | 3.536e-06 | C-terminal His tagged apPOL exonuclease mutant |
| 5dkt-assembly1.cif.gz_A | 1 | 0.182 | 329 | 3.896e-06 | N-terminal His tagged apPOL exonuclease mutant |
| 5dku-assembly1.cif.gz_A | 1 | 0.195 | 302 | 6.641e-06 | C-terminal His tagged apPOL exonuclease mutant |

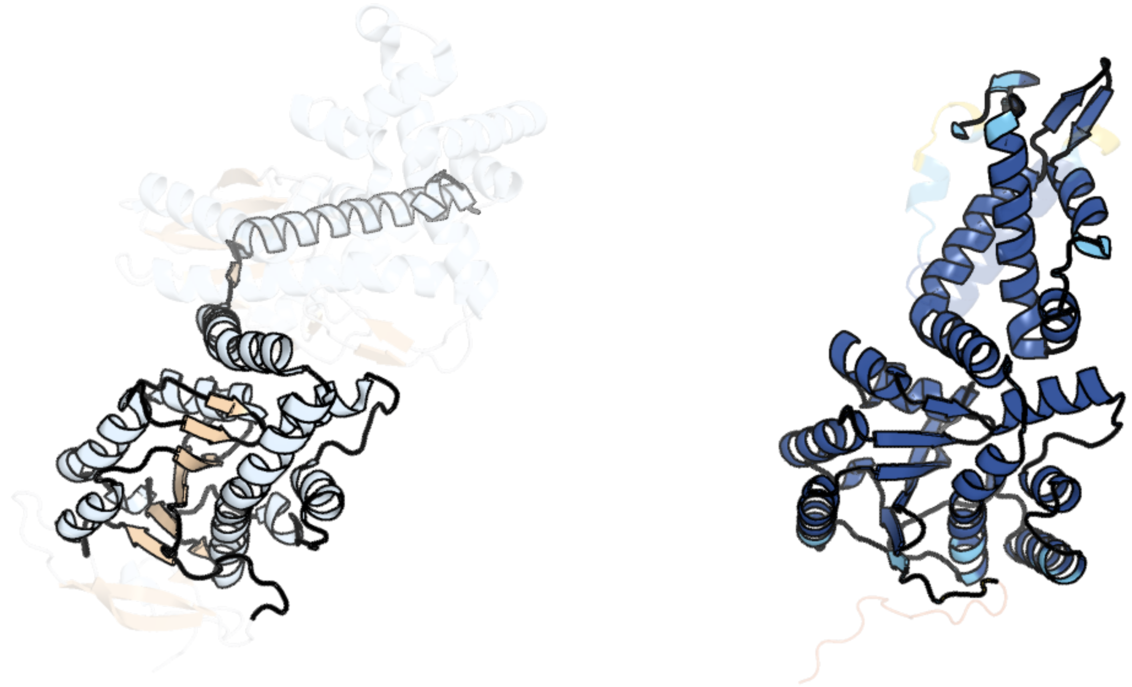

Figure 231: left: reference structure of 5dku chain B. right: predicted structure of chlorv-1..231, unaligned sequences are shown as transparent

#### chlorv-1..232

- Sequence-based annotation for chlorv-1..232 is hypothetical protein
- Best hit was 1yvp chain B: 60-kDa SS-A/Ro ribonucleoprotein

| target | prob | fident | alnlen | evalue | theader |
|---|---|---|---|---|---|
| 1yvp-assembly2.cif.gz_B | 1 | 0.13 | 588 | 1.146e-08 | Ro autoantigen complexed with RNAs |
| 8e59-assembly1.cif.gz_D | 1 | 0.129 | 217 | 0.004582 | Human L-type voltage-gated calcium channel Cav1.3 in the presence of Amiodarone at 3.1 Angstrom resolution |
| 7n1o-assembly1.cif.gz_A | 0.999 | 0.086 | 277 | 0.006035 | The von Willebrand factor A domain of human capillary morphogenesis gene II, flexibly fused to the 1TEL crystallization chaperone |

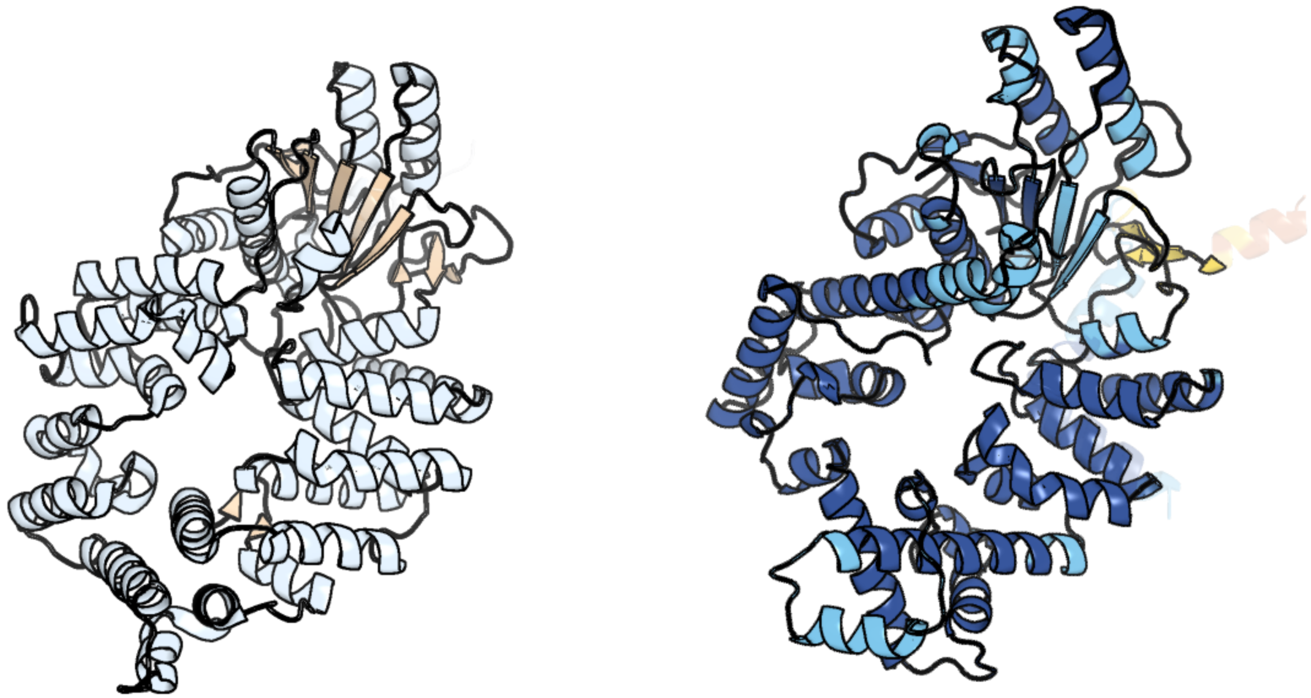

Figure 232: left: reference structure of 1yvp chain B. right: predicted structure of chlorv-1..232, unaligned sequences are shown as transparent

#### chlorv-1..233

- Sequence-based annotation for chlorv-1..233 is hypothetical protein
- Best hit was 6gci chain A: mitochondrial ADP/ATP carrier

| target | prob | fident | alnlen | evalue | theader |
|---|---|---|---|---|---|
| 6gci-assembly1.cif.gz_A | 1 | 0.169 | 284 | 3.96e-06 | Structure of the bongkrekic acid-inhibited mitochondrial ADP/ATP carrier |
| 8g8w-assembly1.cif.gz_A | 1 | 0.192 | 286 | 1.253e-05 | Molecular mechanism of nucleotide inhibition of human uncoupling protein 1 |
| 8gym-assembly1.cif.gz_m2 | 1 | 0.128 | 288 | 0.0003796 | Cryo-EM structure of Tetrahymena thermophila respiratory mega-complex MC IV2+(I+III2+II)2 |

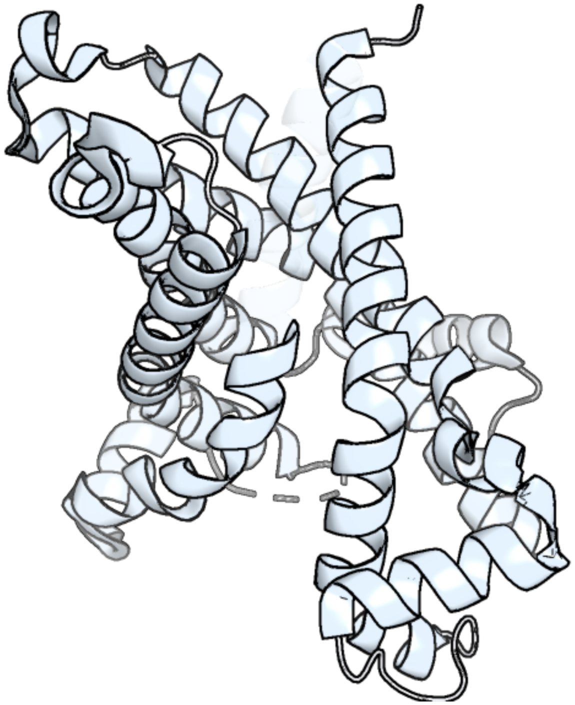

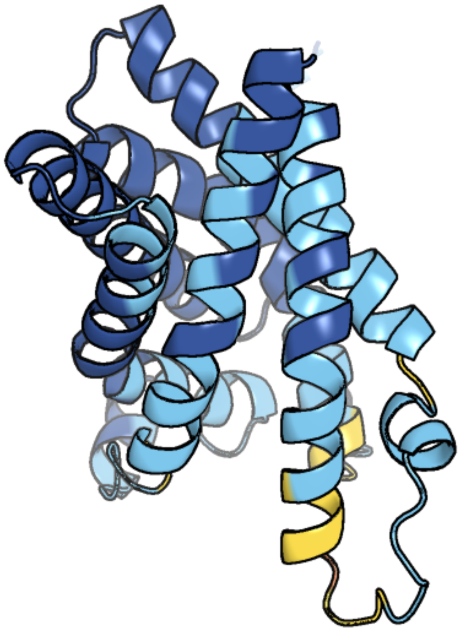

Figure 233: left: reference structure of 6gci chain A. right: predicted structure of chlorv-1..233, unaligned sequences are shown as transparent

#### chlorv-1..234

- Sequence-based annotation for chlorv-1..234 is hypothetical protein
- No significant structural hit found

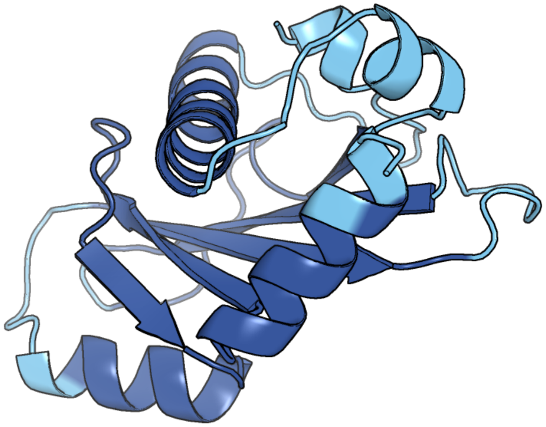

Figure 234: predicted structure of chlorv-1..234

#### chlorv-1..235

- Sequence-based annotation for chlorv-1..235 is hypothetical protein
- No significant structural hit found

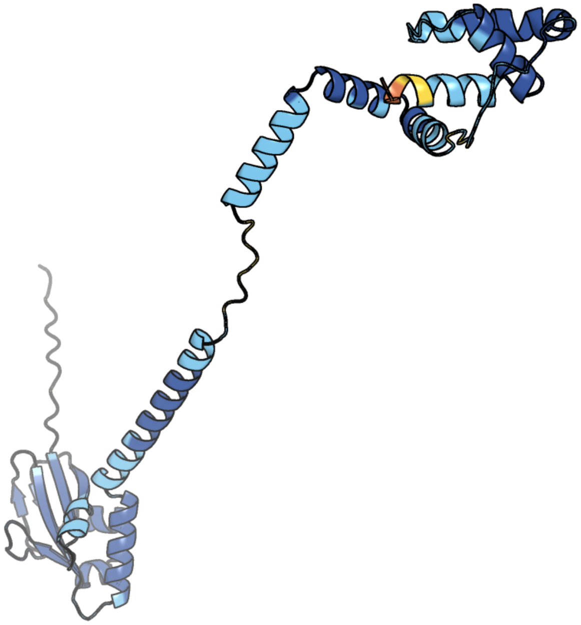

Figure 235: predicted structure of chlorv-1..235

#### chlorv-1..236

- Sequence-based annotation for chlorv-1..236 is hypothetical protein
- No significant structural hit found

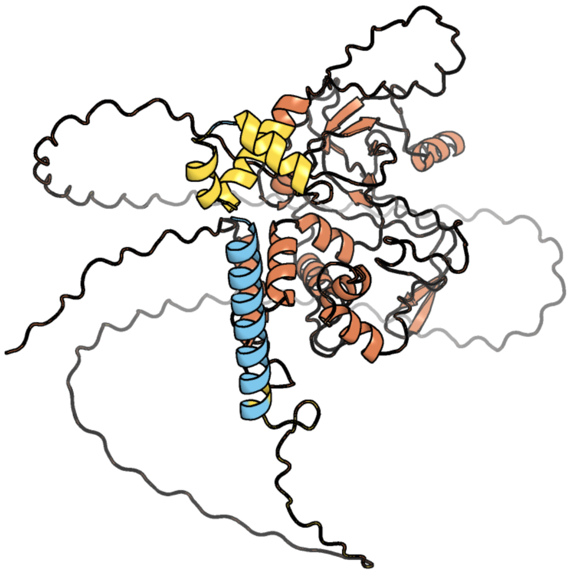

Figure 236: predicted structure of chlorv-1..236

#### chlorv-1..237

- Sequence-based annotation for chlorv-1..237 is hypothetical protein
- No significant structural hit found

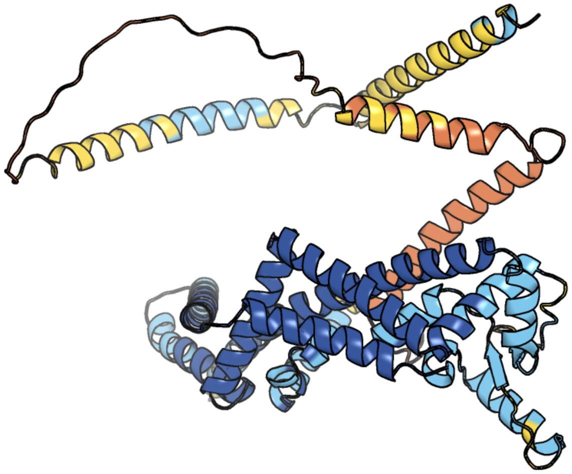

Figure 237: predicted structure of chlorv-1..237

### chlorv-1..238

- Sequence-based annotation for chlorv-1..238 is hypothetical protein
- No significant structural hit found

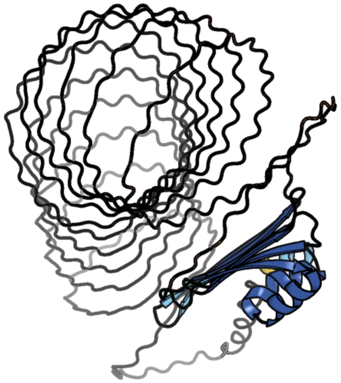

Figure 238: predicted structure of chlorv-1..238

#### chlorv-1..239

- Sequence-based annotation for chlorv-1..239 is hypothetical protein
- No significant structural hit found

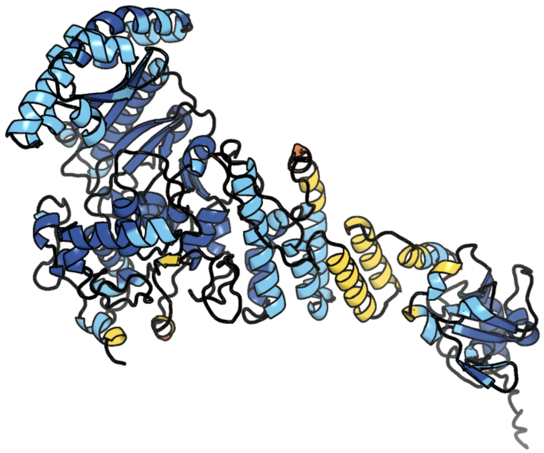

Figure 239: predicted structure of chlorv-1..239

#### chlorv-1..240

- Sequence-based annotation for chlorv-1..240 is hypothetical protein
- Best hit was 8cdj chain C: Cullin-1

| target | prob | fident | alnlen | evalue | theader |
|---|---|---|---|---|---|
| 8cdj-assembly1.cif.gz_C | 1 | 0.103 | 761 | 4.624e-06 | CAND1 b-hairpin++-SCF-SKP2 CAND1 rolling SCF engaged |
| 6v9i-assembly1.cif.gz_C | 1 | 0.117 | 783 | 5.717e-06 | cryo-EM structure of Cullin5 bound to RING-box protein 2 (Cul5-Rbx2) |
| 8or3-assembly1.cif.gz_A | 1 | 0.1 | 750 | 7.069e-06 | CAND1-CUL1-RBX1-SKP1-SKP2-DCNL1 |

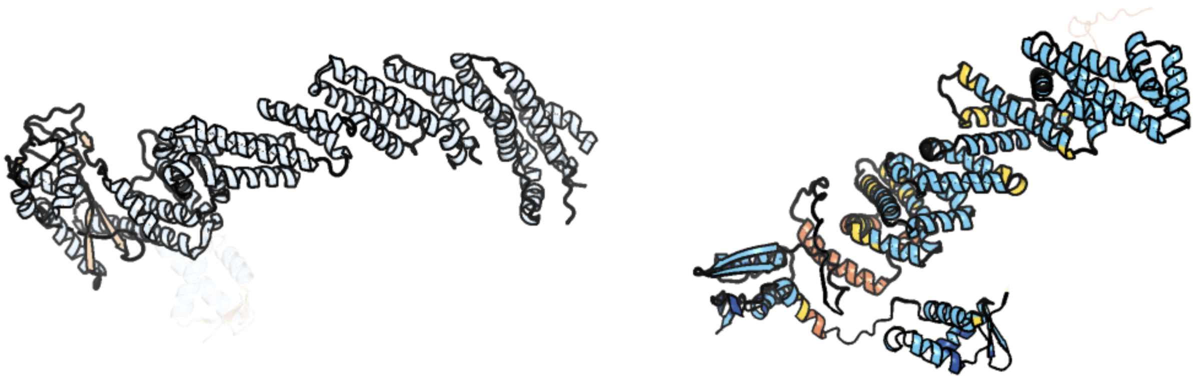

Figure 240: left: reference structure of 8cdj chain C. right: predicted structure of chlorv-1..240, unaligned sequences are shown as transparent

#### chlorv-1..241

- Sequence-based annotation for chlorv-1..241 is hypothetical protein
- No significant structural hit found

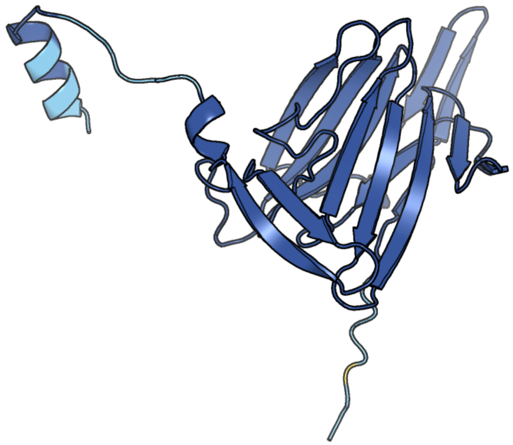

Figure 241: predicted structure of chlorv-1..241

#### chlorv-1..242

- Sequence-based annotation for chlorv-1..242 is hypothetical protein
- No significant structural hit found

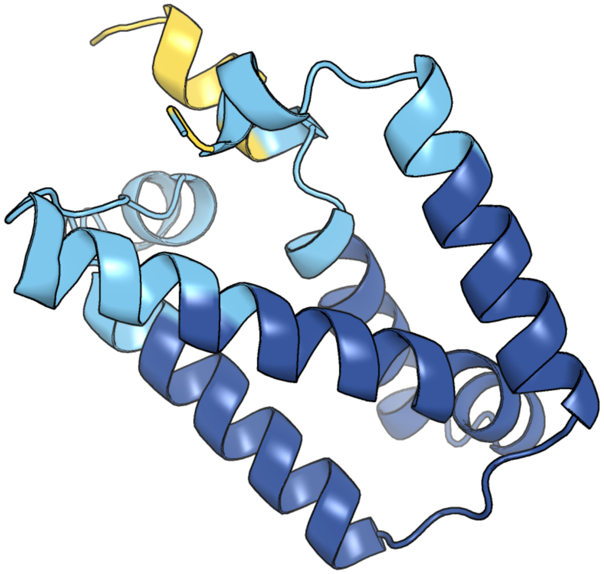

Figure 242: predicted structure of chlorv-1..242

#### chlorv-1..243

- Sequence-based annotation for chlorv-1..243 is putative DUF5755 domain-containing protein
- No significant structural hit found

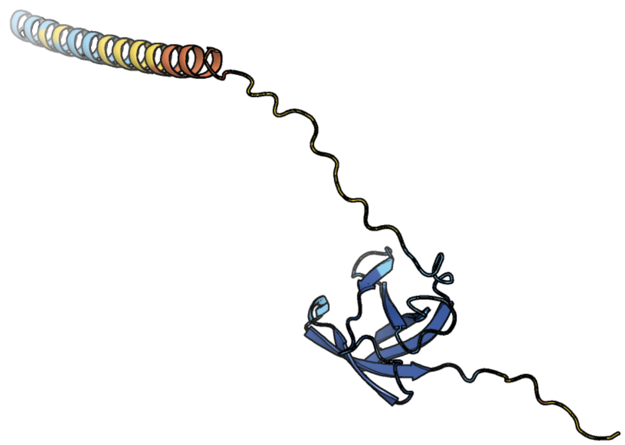

Figure 243: predicted structure of chlorv-1..243

#### chlorv-1..244

- Sequence-based annotation for chlorv-1..244 is hypothetical protein
- No significant structural hit found

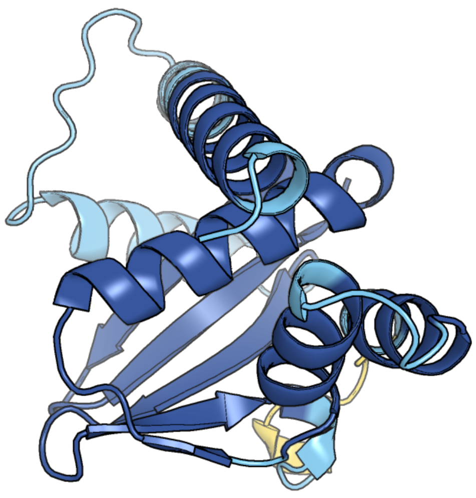

Figure 244: predicted structure of chlorv-1..244

#### chlorv-1..245

- Sequence-based annotation for chlorv-1..245 is putative mRNA decapping protein 2 NUDIX hydrolase domain
- Best hit was 7dnu chain A: mRNA-decapping protein g5R

| target | prob | fident | alnlen | evalue | theader |
|---|---|---|---|---|---|
| 7dnu-assembly1.cif.gz_A | 1 | 0.217 | 280 | 1.487e-10 | mRNA-decapping enzyme g5Rp with inhibitor insp6 complex |
| 5j3t-assembly1.cif.gz_B | 0.997 | 0.148 | 236 | 0.0002148 | Crystal structure of S. pombe Dcp2:Dcp1:Edc1 mRNA decapping complex |
| 4kg4-assembly2.cif.gz_B | 1 | 0.243 | 119 | 0.0003015 | Crystal structure of Saccharomyces cerevisiae Dcp2 Nudix domain (E198Q mutation) |

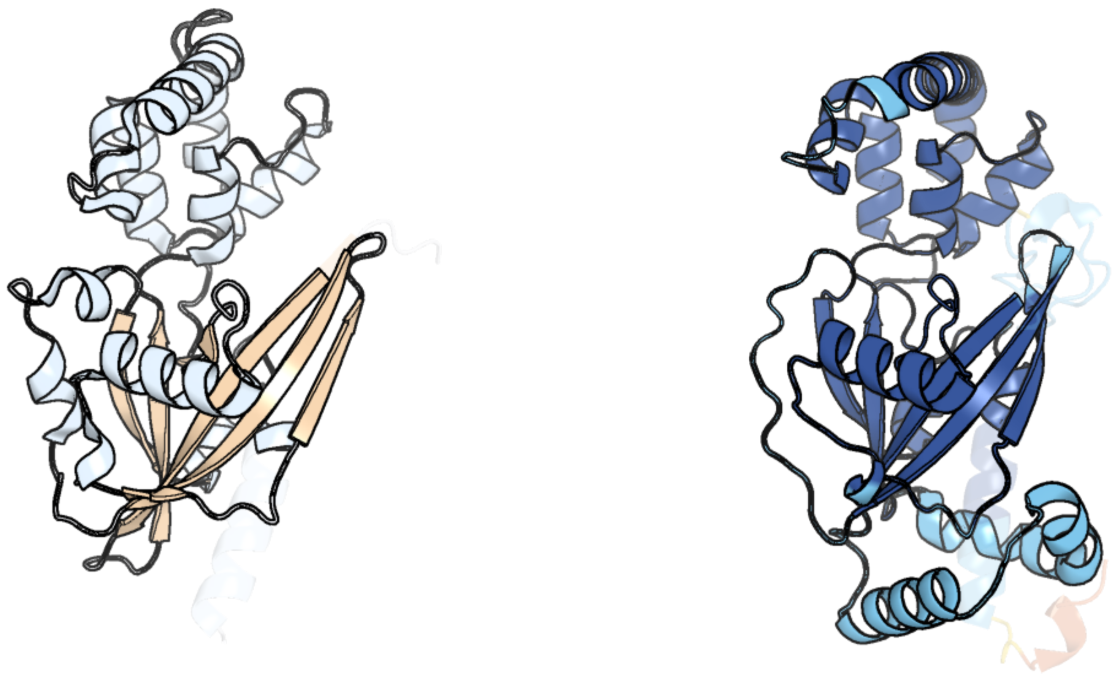

Figure 245: left: reference structure of 7dnu chain A. right: predicted structure of chlorv-1..245, unaligned sequences are shown as transparent

#### chlorv-1..246

- Sequence-based annotation for chlorv-1..246 is hypothetical protein
- No significant structural hit found

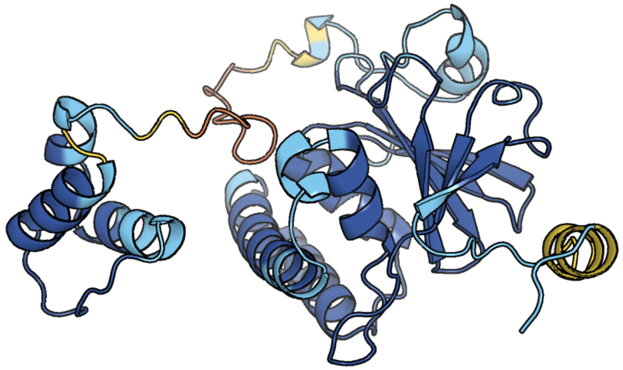

Figure 246: predicted structure of chlorv-1..246

#### chlorv-1..247

- Sequence-based annotation for chlorv-1..247 is putative Erv1/Alr family protein/Oxidoreductase
- Best hit was 3gwn chain B: Probable FAD-linked sulfhydryl oxidase R596

| target | prob | fident | alnlen | evalue | theader |
|---|---|---|---|---|---|
| 3gwn-assembly1.cif.gz_B | 1 | 0.372 | 110 | 3.74e-07 | Crystal structure of the FAD binding domain from mimivirus sulfhydryl oxidase R596 |
| 3td7-assembly1.cif.gz_A-2 | 1 | 0.371 | 113 | 5.545e-07 | Crysal structure of the mimivirus sulfhydryl oxidase R596 |
| 2hj3-assembly1.cif.gz_A | 1 | 0.265 | 98 | 3.972e-06 | Structure of the Arabidopsis Thaliana Erv1 Thiol Oxidase |

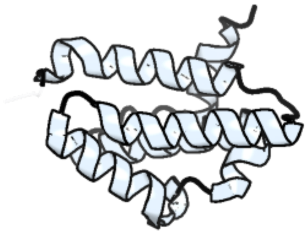

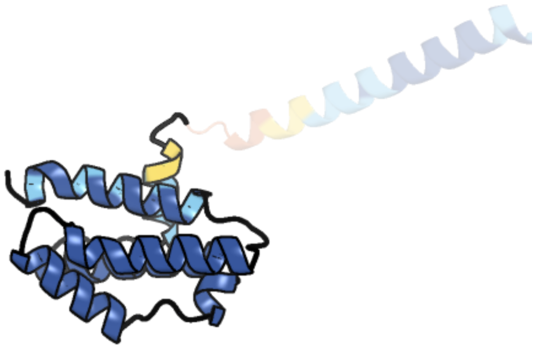

Figure 247: left: reference structure of 3gwn chain B. right: predicted structure of chlorv-1..247, unaligned sequences are shown as transparent

#### chlorv-1..248

- Sequence-based annotation for chlorv-1..248 is hypothetical protein
- No significant structural hit found

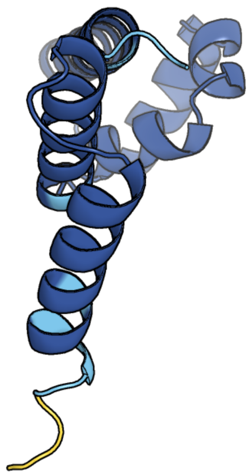

Figure 248: predicted structure of chlorv-1..248

#### chlorv-1..249

- Sequence-based annotation for chlorv-1..249 is putative Methyltransferase
- Best hit was 4n48 chain B: Cap-specific mRNA (nucleoside-2'-O-)-methyltransferase 1

| target | prob | fident | alnlen | evalue | theader |
|---|---|---|---|---|---|
| 4n48-assembly1.cif.gz_B | 1 | 0.178 | 442 | 2.201e-15 | Cap-specific mRNA (nucleoside-2'-O-)-methyltransferase 1 Protein in complex with capped RNA fragment |
| 8p4e-assembly1.cif.gz_O | 1 | 0.183 | 435 | 3.232e-15 | Structural insights into human co-transcriptional capping - structure 5 |
| 4n48-assembly2.cif.gz_A | 1 | 0.174 | 436 | 3.558e-15 | Cap-specific mRNA (nucleoside-2'-O-)-methyltransferase 1 Protein in complex with capped RNA fragment |

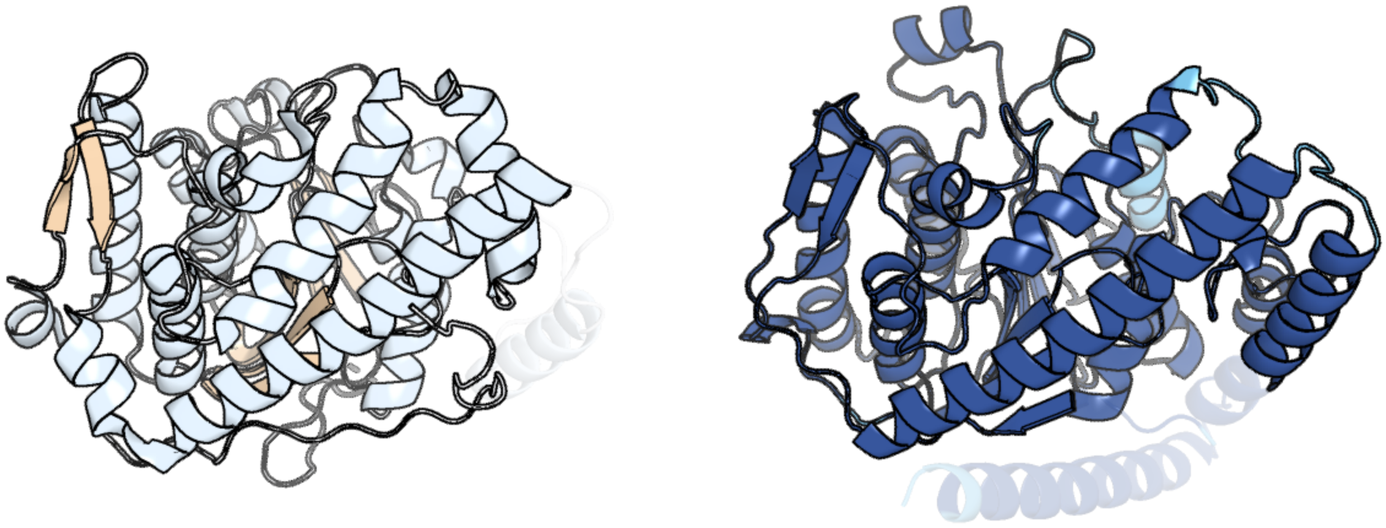

Figure 249: left: reference structure of 4n48 chain B. right: predicted structure of chlorv-1..249, unaligned sequences are shown as transparent

#### chlorv-1..250

- Sequence-based annotation for chlorv-1..250 is hypothetical protein
- No significant structural hit found

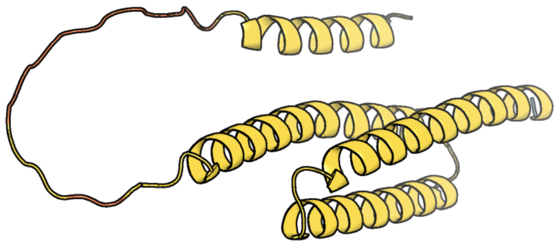

Figure 250: predicted structure of chlorv-1..250

#### chlorv-1..251

- Sequence-based annotation for chlorv-1..251 is hypothetical protein
- Best hit was 7nvu chain G: DNA-directed RNA polymerase II subunit RPB7

| target | prob | fident | alnlen | evalue | theader |
|---|---|---|---|---|---|
| 7nvu-assembly1.cif.gz_G | 1 | 0.218 | 169 | 6.014e-11 | RNA polymerase II core pre-initiation complex with open promoter DNA |
| 1go3-assembly1.cif.gz_E | 1 | 0.194 | 159 | 7.185e-10 | Structure of an archeal homolog of the eukaryotic RNA polymerase II RPB4/RPB7 complex |
| 3h0g-assembly1.cif.gz_G | 1 | 0.182 | 164 | 8.478e-10 | RNA Polymerase II from Schizosaccharomyces pombe |

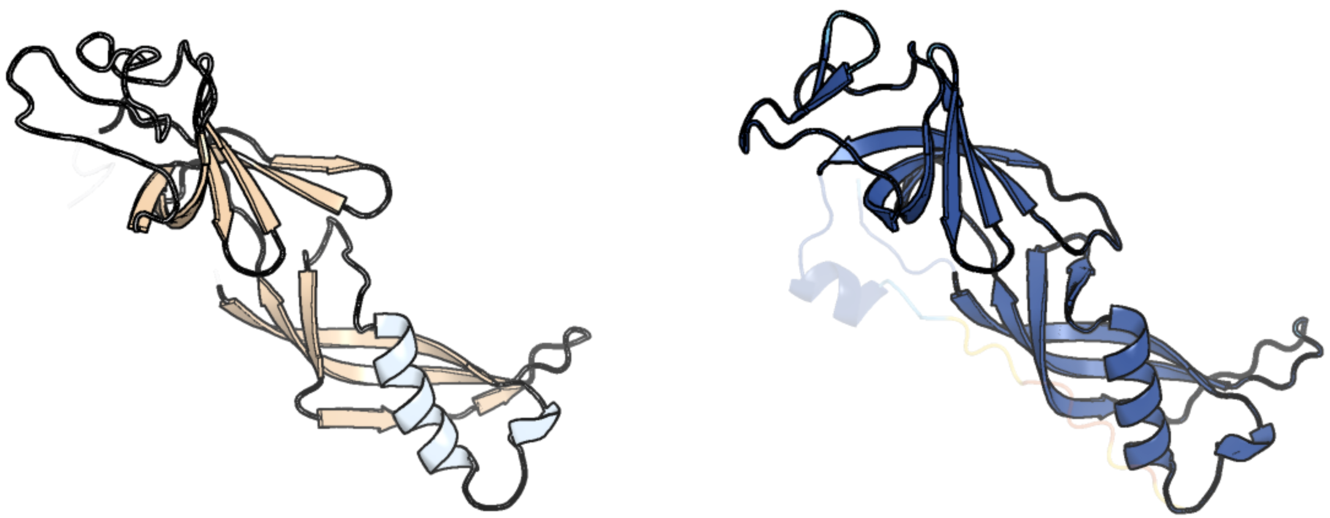

Figure 251: left: reference structure of 7nvu chain G. right: predicted structure of chlorv-1..251, unaligned sequences are shown as transparent

#### chlorv-1..252

- Sequence-based annotation for chlorv-1..252 is putative Ankyrin repeat protein
- Best hit was 5lee chain A: DDD_D12_12_D12_12_D12

| target | prob | fident | alnlen | evalue | theader |
|---|---|---|---|---|---|
| 5lee-assembly1.cif.gz_A | 1 | 0.185 | 367 | 1.335e-08 | Crystal structure of DARPin-DARPin rigid fusion, variant DDD_D12_12_D12_12_D12 |
| 5leb-assembly1.cif.gz_A | 1 | 0.17 | 369 | 1.673e-08 | Crystal structure of DARPin-DARPin rigid fusion, variant DDD_D12_06_D12_06_D12 |
| 5le8-assembly2.cif.gz_B | 1 | 0.178 | 341 | 2.294e-08 | Crystal structure of DARPin-DARPin rigid fusion, variant DD_D12_15_D12 |

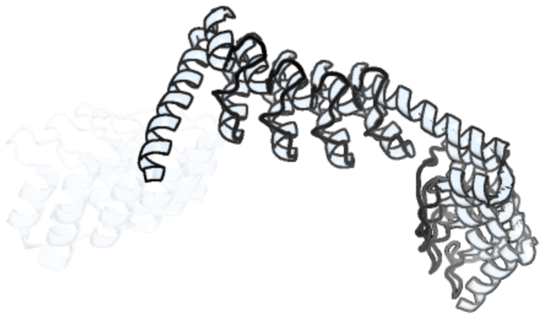

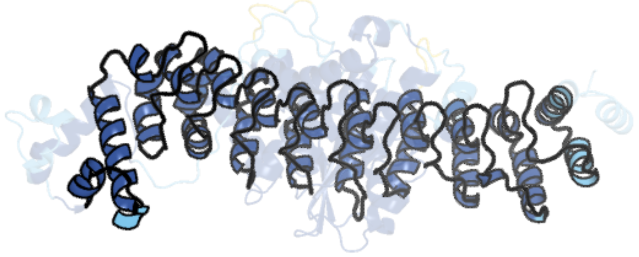

Figure 252: left: reference structure of 5lee chain A. right: predicted structure of chlorv-1..252, unaligned sequences are shown as transparent

#### chlorv-1..253

- Sequence-based annotation for chlorv-1..253 is putative RNA polymerase subunit 5
- Best hit was 7vba chain E: DNA-directed RNA polymerases I, II, and III subunit RPABC1

| target | prob | fident | alnlen | evalue | theader |
|---|---|---|---|---|---|
| 7vba-assembly1.cif.gz_E | 1 | 0.231 | 207 | 4.32e-13 | Structure of the pre state human RNA Polymerase I Elongation Complex |
| 5iy8-assembly1.cif.gz_E | 1 | 0.238 | 210 | 9.004e-13 | Human holo-PIC in the initial transcribing state |
| 8cen-assembly1.cif.gz_E | 1 | 0.215 | 209 | 2.255e-12 | Yeast RNA polymerase II transcription pre-initiation complex with core Mediator |

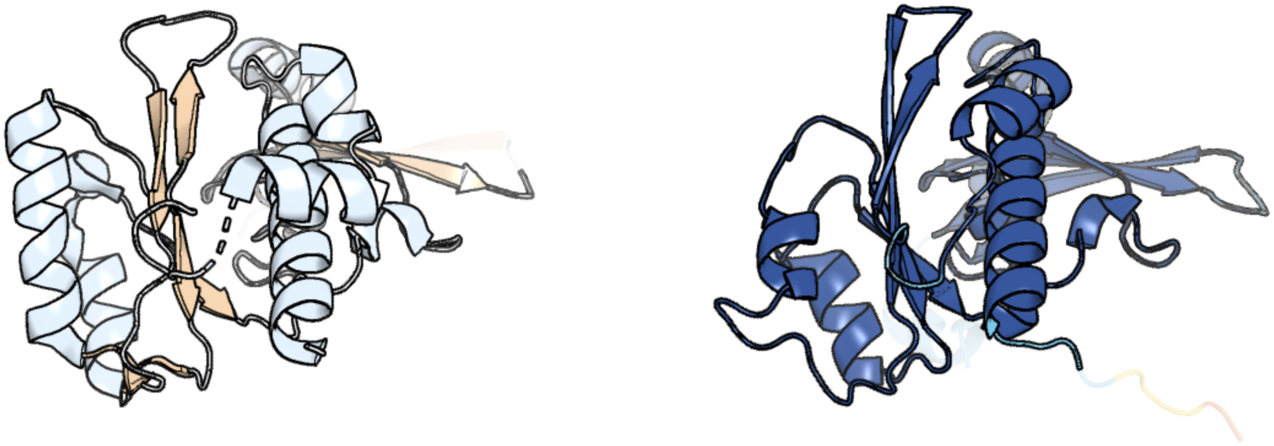

Figure 253: left: reference structure of 7vba chain E. right: predicted structure of chlorv-1..253, unaligned sequences are shown as transparent

#### chlorv-1..254

- Sequence-based annotation for chlorv-1..254 is hypothetical protein
- No significant structural hit found

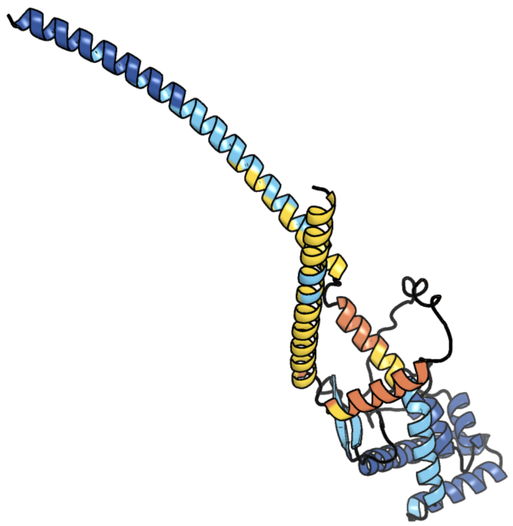

Figure 254: predicted structure of chlorv-1..254

### chlorv-1..255

- Sequence-based annotation for chlorv-1..255 is putative Ubiquitin-activating enzyme
- Best hit was 6dc6 chain A: Ubiquitin-like modifier-activating enzyme 1

| target | prob | fident | alnlen | evalue | theader |
|---|---|---|---|---|---|
| 6dc6-assembly1.cif.gz_A | 1 | 0.125 | 884 | 1.523e-32 | Crystal structure of human ubiquitin activating enzyme E1 (Uba1) in complex with ubiquitin |
| 6zhu-assembly4.cif.gz_G | 1 | 0.138 | 885 | 4.203e-32 | Yeast Uba1 in complex with Ubc3 and ATP |
| 6zhs-assembly1.cif.gz_A | 1 | 0.133 | 884 | 5.572e-32 | Uba1 bound to two E2 (Ubc13) molecules |

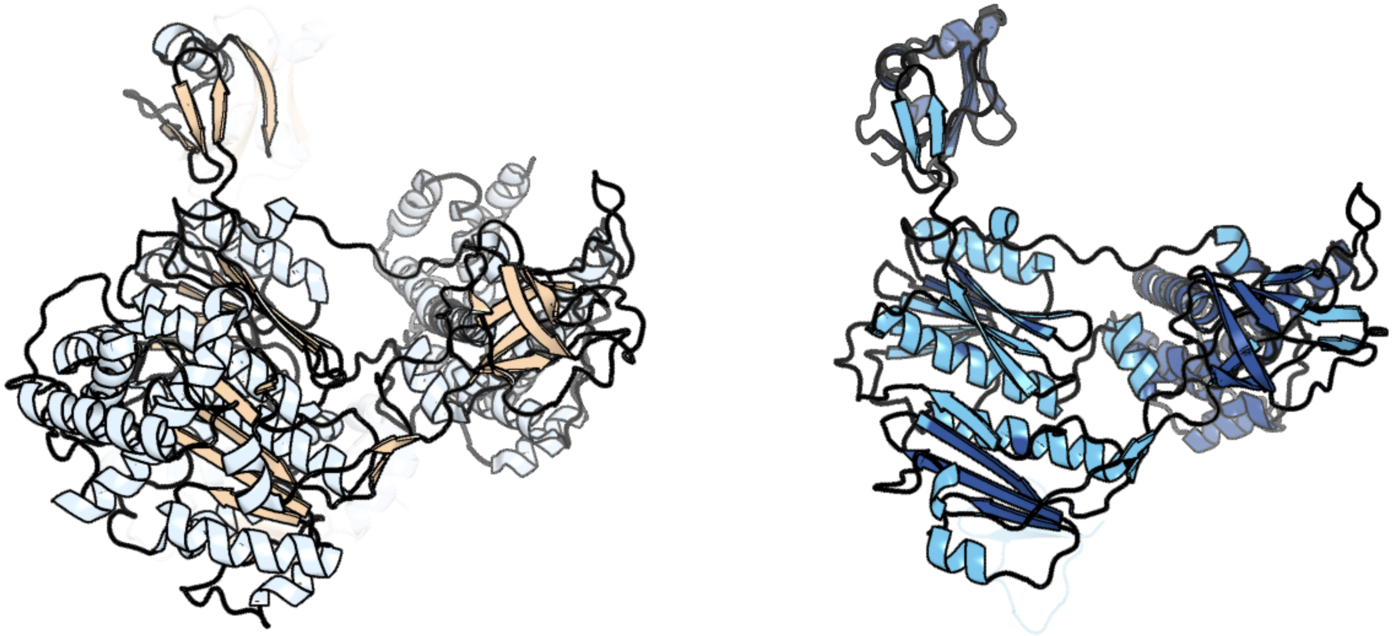

Figure 255: left: reference structure of 6dc6 chain A. right: predicted structure of chlorv-1..255, unaligned sequences are shown as transparent

#### chlorv-1..256

- Sequence-based annotation for chlorv-1..256 is putative Methionine aminopeptidase
- Best hit was 5lyx chain A: Methionine aminopeptidase 2

| target | prob | fident | alnlen | evalue | theader |
|---|---|---|---|---|---|
| 5lyx-assembly1.cif.gz_A | 1 | 0.366 | 311 | 3.003e-32 | CRYSTAL STRUCTURE OF HUMAN METHIONINE AMINOPEPTIDASE-2 IN COMPLEX; WITH AN INHIBITOR 5-((R)-1-[1,2,4]Triazolo[1,5-a]pyrimidin-7-yl-pyrrolidin-2-ylmethoxy)-isoquinoline |
| 1kq9-assembly1.cif.gz_A | 1 | 0.372 | 311 | 5.083e-32 | Human methionine aminopeptidase type II in complex with L-methionine |
| 1yw7-assembly1.cif.gz_A | 1 | 0.372 | 311 | 1.295e-31 | h-MetAP2 complexed with A444148 |

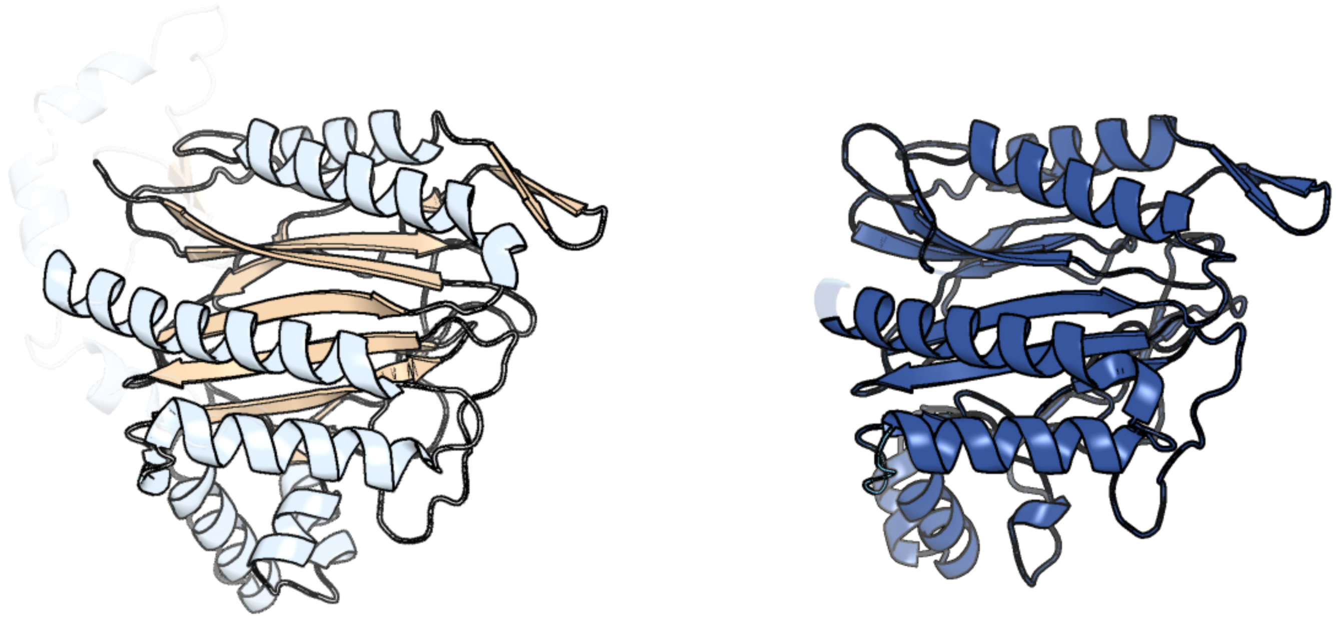

Figure 256: left: reference structure of 5lyx chain A. right: predicted structure of chlorv-1..256, unaligned sequences are shown as transparent

#### chlorv-1..257

- Sequence-based annotation for chlorv-1..257 is putative Ankyrin repeat protein
- Best hit was 5orm chain A: cPPR-Telo1

| target | prob | fident | alnlen | evalue | theader |
|---|---|---|---|---|---|
| 5orm-assembly1.cif.gz_A | 1 | 0.117 | 374 | 9.735e-09 | Crystal structure of designed cPPR-Telo1 |
| 5i9g-assembly1.cif.gz_C | 1 | 0.09 | 355 | 9.735e-09 | Crystal structure of designed pentatricopeptide repeat protein dPPR-U8C2 in complex with its target RNA U8C2 |
| 5i9f-assembly1.cif.gz_A | 1 | 0.082 | 387 | 3.267e-08 | Crystal structure of designed pentatricopeptide repeat protein dPPR-U10 in complex with its target RNA U10 |

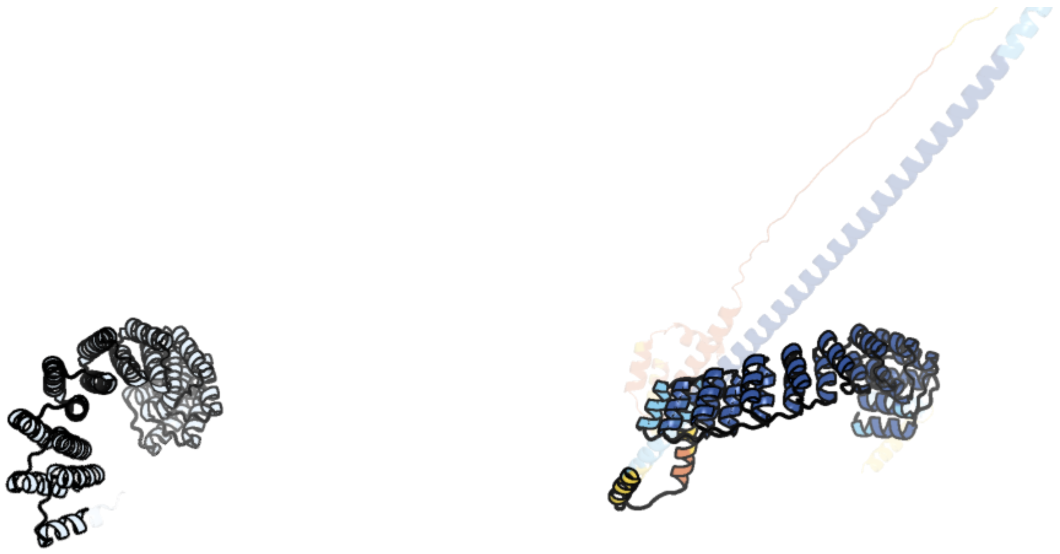

Figure 257: left: reference structure of 5orm chain A. right: predicted structure of chlorv-1..257, unaligned sequences are shown as transparent

### chlorv-1..258

- Sequence-based annotation for chlorv-1..258 is hypothetical protein
- No significant structural hit found

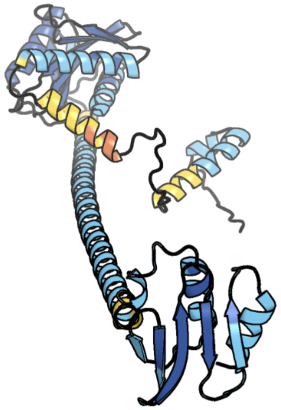

Figure 258: predicted structure of chlorv-1..258

#### chlorv-1..259

- Sequence-based annotation for chlorv-1..259 is hypothetical protein
- No significant structural hit found

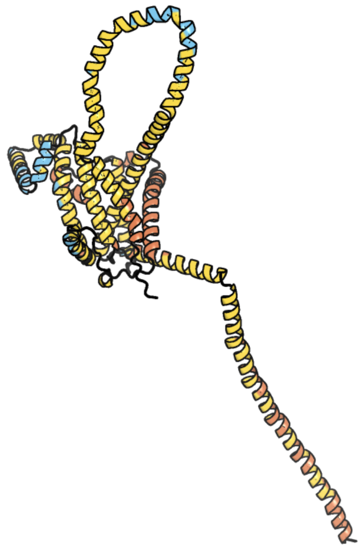

Figure 259: predicted structure of chlorv-1..259

### chlorv-1..260

- Sequence-based annotation for chlorv-1..260 is hypothetical protein
- Best hit was 2jrs chain A: RNA-binding protein 39

| target | prob | fident | alnlen | evalue | theader |
|---|---|---|---|---|---|
| 2jrs-assembly1.cif.gz_A | 1 | 0.163 | 98 | 8.287e-06 | Solution NMR Structure of CAPER RRM2 Domain. Northeast Structural Genomics Target HR4730A |
| 3hi9-assembly1.cif.gz_A | 1 | 0.215 | 79 | 1.074e-05 | The x-ray crystal structure of the first RNA recognition motif (RRM1) of the AU-rich element (ARE) binding protein HuR at 2.0 angstrom resolution |
| 8fle-assembly1.cif.gz_SH | 1 | 0.139 | 79 | 1.305e-05 | Human nuclear pre-60S ribosomal subunit (State L2) |

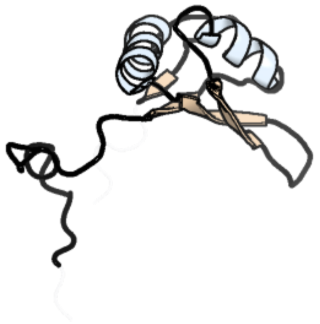

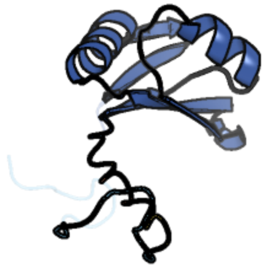

Figure 260: left: reference structure of 2jrs chain A. right: predicted structure of chlorv-1..260, unaligned sequences are shown as transparent

#### chlorv-1..261

- Sequence-based annotation for chlorv-1..261 is hypothetical protein
- No significant structural hit found

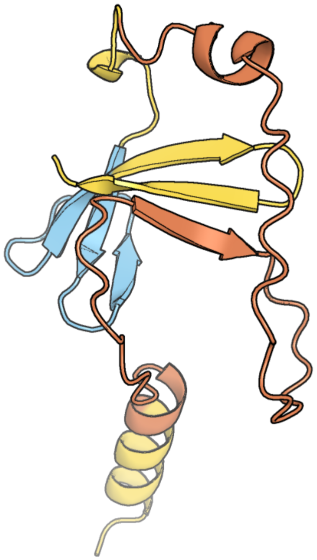

Figure 261: predicted structure of chlorv-1..261

#### chlorv-1..262

- Sequence-based annotation for chlorv-1..262 is hypothetical protein
- No significant structural hit found

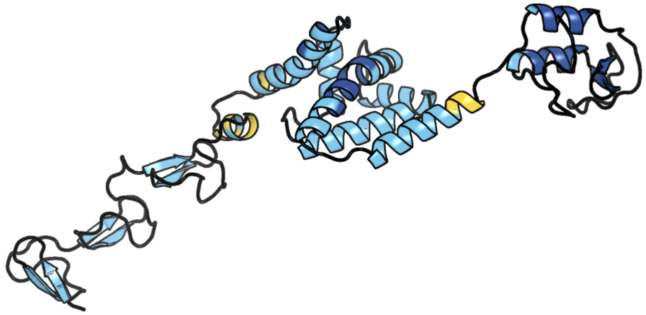

Figure 262: predicted structure of chlorv-1..262

#### chlorv-1..263

- Sequence-based annotation for chlorv-1..263 is hypothetical protein
- Best hit was 6er3 chain A: BNR/Asp-box repeat protein

| target | prob | fident | alnlen | evalue | theader |
|---|---|---|---|---|---|
| 6er3-assembly1.cif.gz_A | 1 | 0.132 | 188 | 1.434e-08 | Ruminococcus gnavus IT-sialidase CBM40 bound to alpha2,3 sialyllactose |
| 6er4-assembly1.cif.gz_B | 1 | 0.13 | 191 | 1.984e-08 | Ruminococcus gnavus IT-sialidase CBM40 bound to alpha2,6 sialyllactose |
| 6er2-assembly1.cif.gz_A | 1 | 0.132 | 188 | 2.094e-08 | Ruminococcus gnavus IT-sialidase CBM40 |

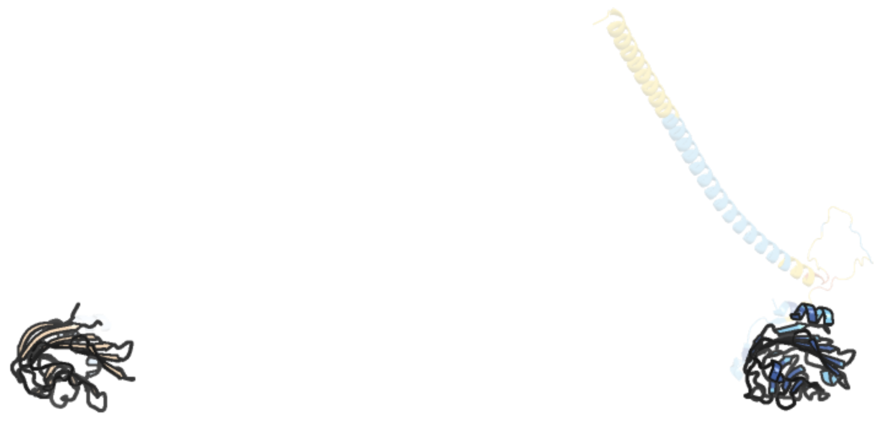

Figure 263: left: reference structure of 6er3 chain A. right: predicted structure of chlorv-1..263, unaligned sequences are shown as transparent

#### chlorv-1..264

- Sequence-based annotation for chlorv-1..264 is hypothetical protein
- No significant structural hit found

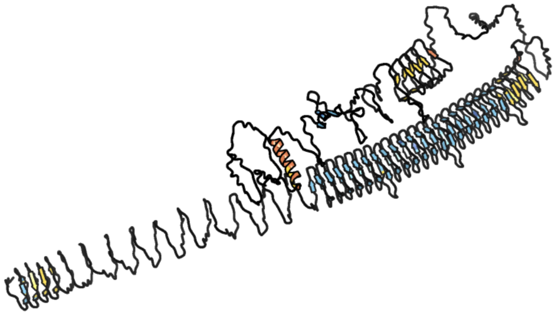

Figure 264: predicted structure of chlorv-1..264

#### chlorv-1..265

- Sequence-based annotation for chlorv-1..265 is hypothetical protein
- Best hit was 6g41 chain B: Minor capsid protein

| target | prob | fident | alnlen | evalue | theader |
|---|---|---|---|---|---|
| 6g41-assembly1.cif.gz_B | 1 | 0.106 | 310 | 2.75e-09 | Crystal structure of SeMet-labeled mavirus penton protein |
| 6g42-assembly1.cif.gz_E | 1 | 0.103 | 318 | 6.018e-09 | Crystal structure of mavirus penton protein |
| 6g41-assembly2.cif.gz_G | 1 | 0.121 | 314 | 9.415e-09 | Crystal structure of SeMet-labeled mavirus penton protein |

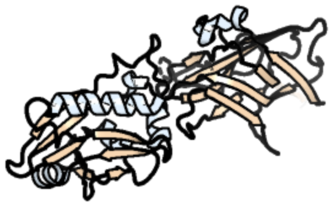

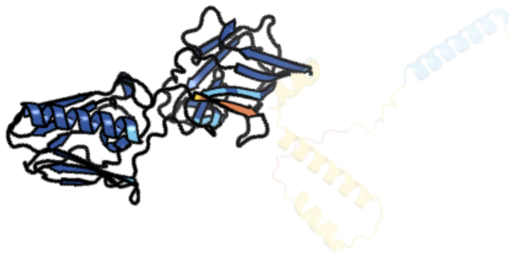

Figure 265: left: reference structure of 6g41 chain B. right: predicted structure of chlorv-1..265, unaligned sequences are shown as transparent

#### chlorv-1..266

- Sequence-based annotation for chlorv-1..266 is hypothetical protein
- Best hit was 6g41 chain G: Minor capsid protein

| target | prob | fident | alnlen | evalue | theader |
|---|---|---|---|---|---|
| 6g41-assembly2.cif.gz_G | 1 | 0.091 | 305 | 2.458e-08 | Crystal structure of SeMet-labeled mavirus penton protein |
| 6g41-assembly1.cif.gz_B | 1 | 0.1 | 310 | 2.596e-08 | Crystal structure of SeMet-labeled mavirus penton protein |
| 6g42-assembly1.cif.gz_E | 1 | 0.09 | 320 | 9.085e-08 | Crystal structure of mavirus penton protein |

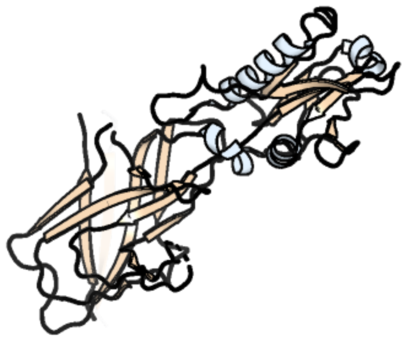

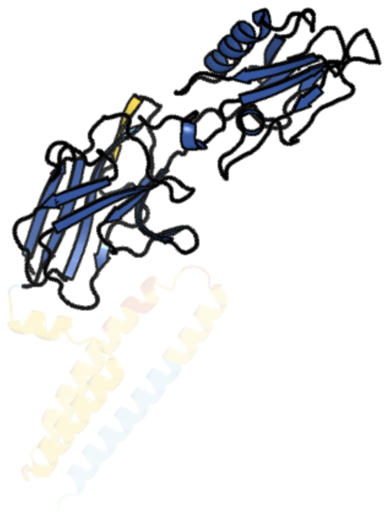

Figure 266: left: reference structure of 6g41 chain G. right: predicted structure of chlorv-1..266, unaligned sequences are shown as transparent

#### chlorv-1..267

- Sequence-based annotation for chlorv-1..267 is hypothetical protein
- No significant structural hit found

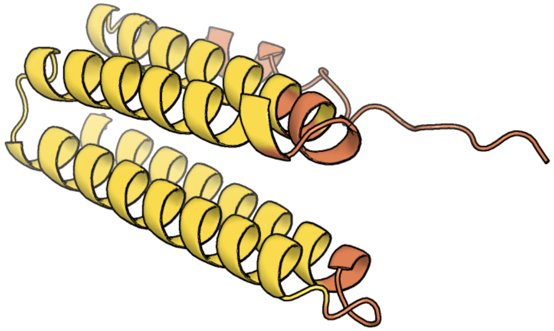

Figure 267: predicted structure of chlorv-1..267

#### chlorv-1..268

- Sequence-based annotation for chlorv-1..268 is hypothetical protein
- No significant structural hit found

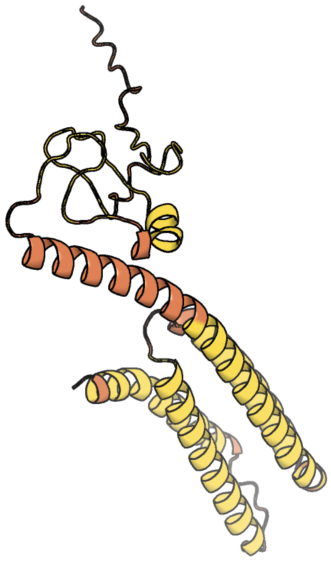

Figure 268: predicted structure of chlorv-1..268

#### chlorv-1..269

- Sequence-based annotation for chlorv-1..269 is hypothetical protein
- No significant structural hit found

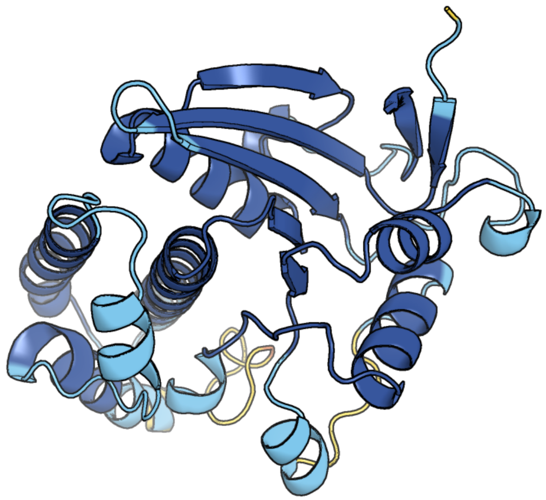

Figure 269: predicted structure of chlorv-1..269

#### chlorv-1..270

- Sequence-based annotation for chlorv-1..270 is putative Helcase
- Best hit was 8qca chain A: Antiviral helicase SKI2

| target | prob | fident | alnlen | evalue | theader |
|---|---|---|---|---|---|
| 8qca-assembly1.cif.gz_A | 1 | 0.157 | 827 | 1.889e-16 | CryoEM structure of a S. Cerevisiae Ski2387 complex in the closed state bound to RNA |
| 4buj-assembly1.cif.gz_A | 1 | 0.164 | 853 | 4.377e-16 | Crystal structure of the S. cerevisiae Ski2-3-8 complex |
| 4buj-assembly2.cif.gz_E | 1 | 0.157 | 856 | 7.12e-16 | Crystal structure of the S. cerevisiae Ski2-3-8 complex |

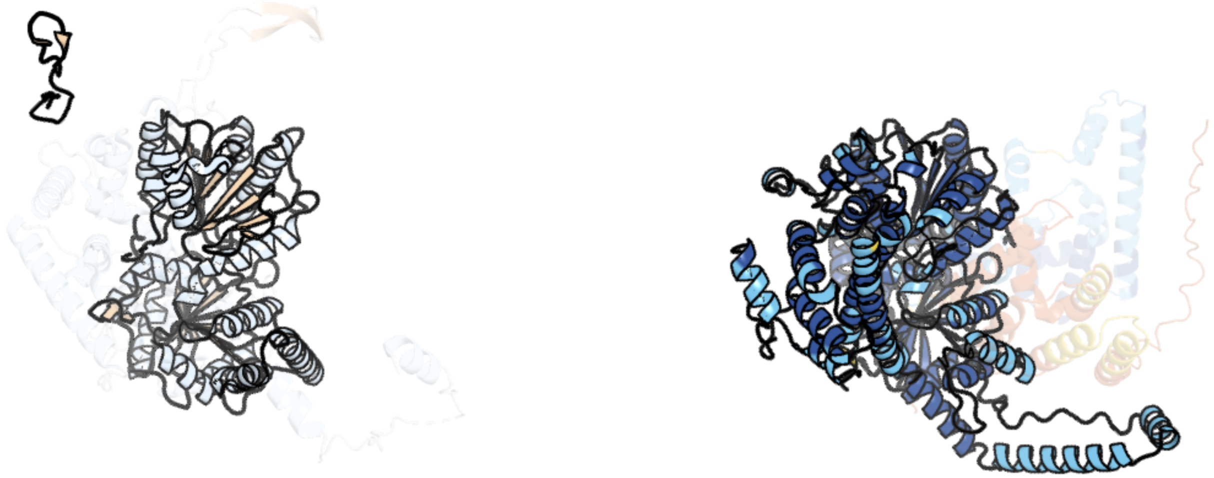

Figure 270: left: reference structure of 8qca chain A. right: predicted structure of chlorv-1..270, unaligned sequences are shown as transparent

#### chlorv-1..271

- Sequence-based annotation for chlorv-1..271 is hypothetical protein
- Best hit was 4ix3 chain A: MsStt7d protein

| target | prob | fident | alnlen | evalue | theader |
|---|---|---|---|---|---|
| 4ix3-assembly1.cif.gz_A | 1 | 0.141 | 389 | 5.45e-08 | Crystal structure of a Stt7 homolog from Micromonas algae |
| 4mvf-assembly1.cif.gz_A | 1 | 0.149 | 327 | 4.375e-07 | Crystal Structure of Plasmodium falciparum CDPK2 complexed with inhibitor staurosporine |
| 8u2o-assembly1.cif.gz_A | 1 | 0.153 | 346 | 5.64e-07 | Crystal Structure of Cdk-related protein kinase 6 (PK6) from Plasmodium falciparum in complex with inhibitor TCMDC-123995 |

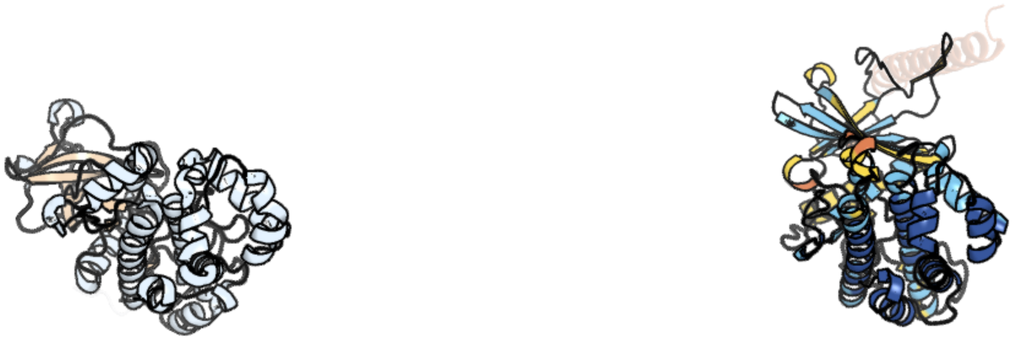

Figure 271: left: reference structure of 4ix3 chain A. right: predicted structure of chlorv-1..271, unaligned sequences are shown as transparent

#### chlorv-1..272

- Sequence-based annotation for chlorv-1..272 is hypothetical protein
- No significant structural hit found

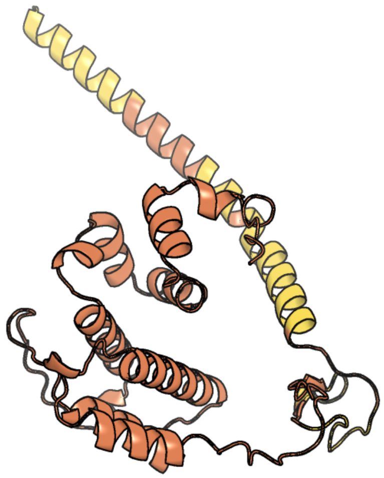

Figure 272: predicted structure of chlorv-1..272

#### chlorv-1..273

- Sequence-based annotation for chlorv-1..273 is hypothetical protein
- No significant structural hit found

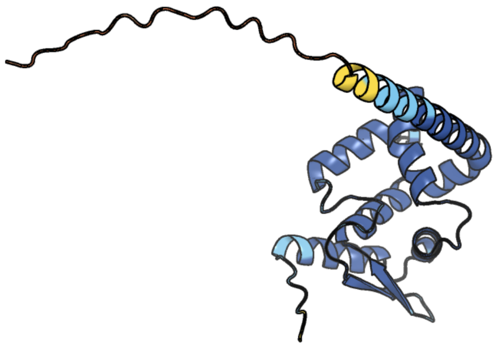

Figure 273: predicted structure of chlorv-1..273

#### chlorv-1..274

- Sequence-based annotation for chlorv-1..274 is hypothetical protein
- No significant structural hit found

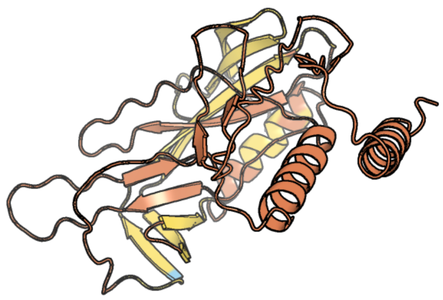

Figure 274: predicted structure of chlorv-1..274

#### chlorv-1..275

- Sequence-based annotation for chlorv-1..275 is hypothetical protein
- Best hit was 7yeq chain A: CP312R

| target | prob | fident | alnlen | evalue | theader |
|---|---|---|---|---|---|
| 7yeq-assembly1.cif.gz_A | 1 | 0.082 | 254 | 4.201e-06 | Structural insight into African Swine Fever Virus CP312R protein reveals it as a single-stranded DNA binding protein |
| 5odk-assembly1.cif.gz_A | 0.999 | 0.111 | 198 | 0.0003457 | Single-stranded DNA-binding protein from bacteriophage Enc34, C-terminal truncation |
| 5odj-assembly1.cif.gz_A | 0.999 | 0.131 | 198 | 0.0004677 | Single-stranded DNA-binding protein from bacteriophage Enc34 |

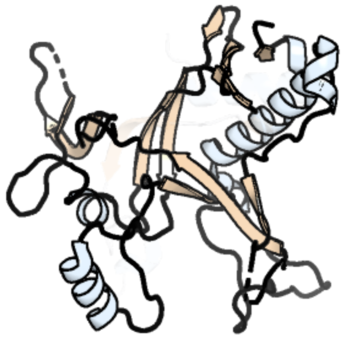

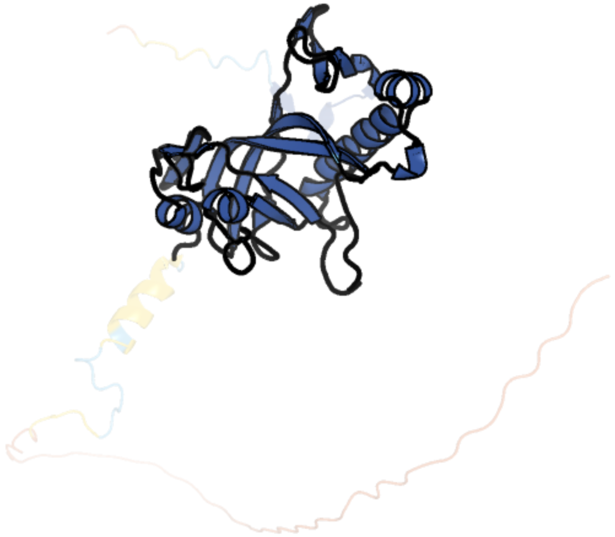

Figure 275: left: reference structure of 7yeq chain A. right: predicted structure of chlorv-1..275, unaligned sequences are shown as transparent

#### chlorv-1..276

- Sequence-based annotation for chlorv-1..276 is hypothetical protein
- No significant structural hit found

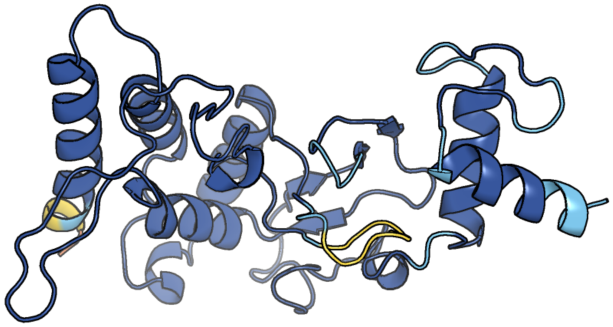

Figure 276: predicted structure of chlorv-1..276

#### chlorv-1..277

- Sequence-based annotation for chlorv-1..277 is putative Helicase
- Best hit was 2vsx chain A: ATP-DEPENDENT RNA HELICASE EIF4A

| target | prob | fident | alnlen | evalue | theader |
|---|---|---|---|---|---|
| 2vsx-assembly1.cif.gz_A | 1 | 0.363 | 380 | 2.047e-35 | Crystal Structure of a Translation Initiation Complex |
| 8c6j-assembly1.cif.gz_7 | 1 | 0.384 | 377 | 5.216e-35 | Human spliceosomal PM5 C* complex |
| 1fuu-assembly1.cif.gz_B | 1 | 0.375 | 376 | 6.204e-34 | YEAST INITIATION FACTOR 4A |

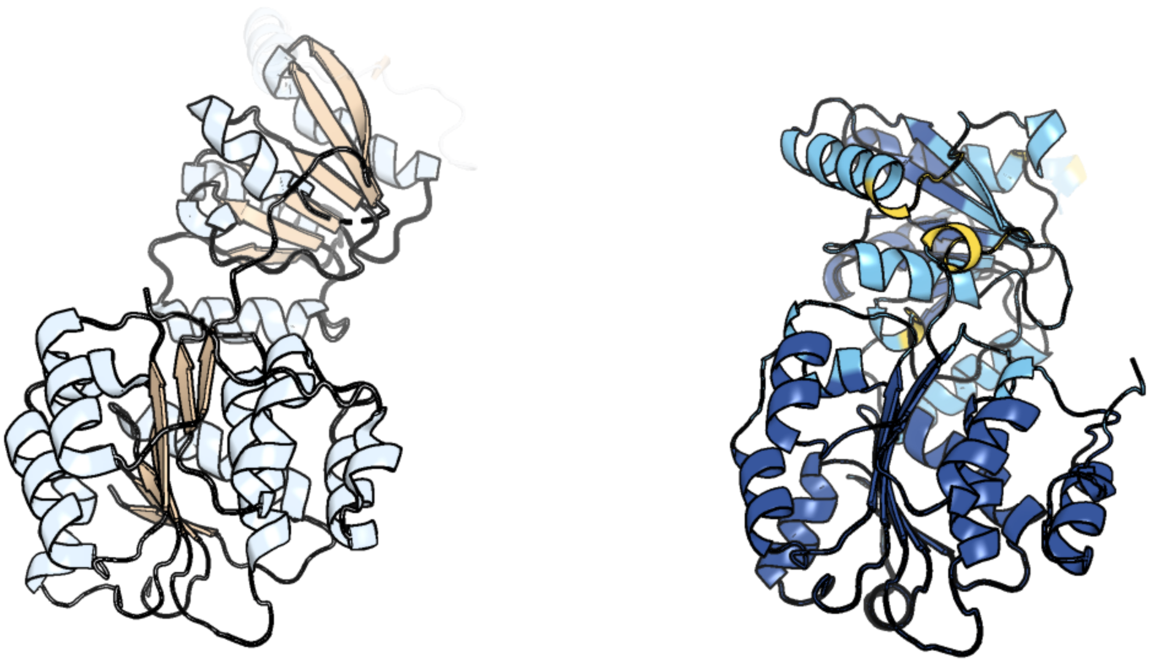

Figure 277: left: reference structure of 2vsx chain A. right: predicted structure of chlorv-1..277, unaligned sequences are shown as transparent

#### chlorv-1..278

- Sequence-based annotation for chlorv-1..278 is putative Erv1/Alr family protein/Oxidoreductase
- Best hit was 3gwn chain B: Probable FAD-linked sulfhydryl oxidase R596

| target | prob | fident | alnlen | evalue | theader |
|---|---|---|---|---|---|
| 3gwn-assembly1.cif.gz_B | 1 | 0.415 | 113 | 6.397e-09 | Crystal structure of the FAD binding domain from mimivirus sulfhydryl oxidase R596 |
| 3td7-assembly1.cif.gz_A-2 | 1 | 0.421 | 114 | 4.138e-08 | Crysal structure of the mimivirus sulfhydryl oxidase R596 |
| 2hj3-assembly1.cif.gz_A | 1 | 0.323 | 102 | 6.468e-06 | Structure of the Arabidopsis Thaliana Erv1 Thiol Oxidase |

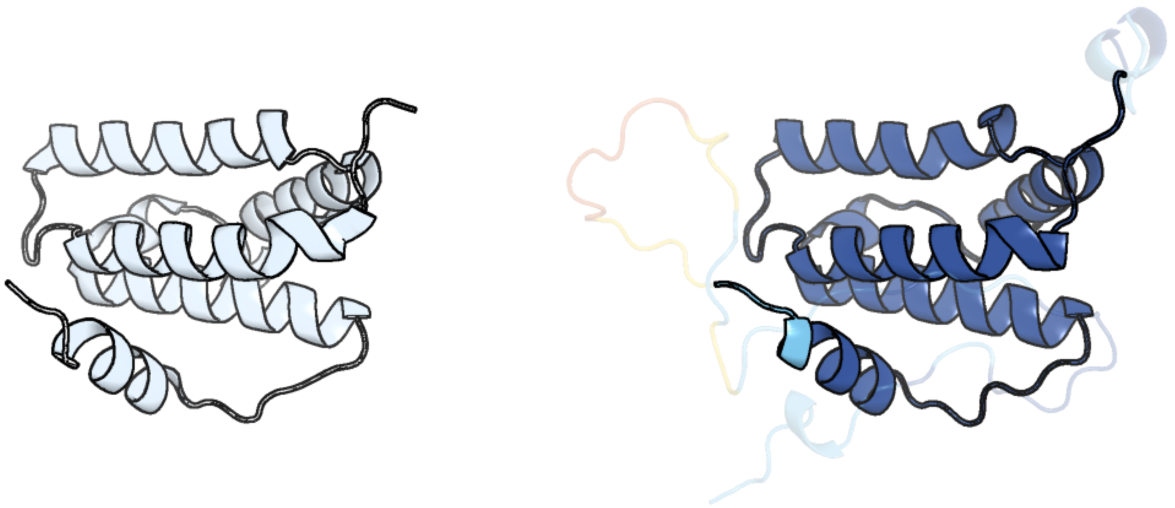

Figure 278: left: reference structure of 3gwn chain B. right: predicted structure of chlorv-1..278, unaligned sequences are shown as transparent

#### chlorv-1..279

- Sequence-based annotation for chlorv-1..279 is hypothetical protein
- Best hit was 8h2f chain A: DnaQ

| target | prob | fident | alnlen | evalue | theader |
|---|---|---|---|---|---|
| 8h2f-assembly1.cif.gz_A | 1 | 0.161 | 204 | 4.036e-09 | Crystal structure of DnaQ domain in complex witn TMP of Streptococcus thermophilus strain DGCC 7710 |
| 2f96-assembly1.cif.gz_B | 1 | 0.085 | 223 | 2.529e-07 | 2.1 A crystal structure of Pseudomonas aeruginosa rnase T (Ribonuclease T) |
| 3nh2-assembly1.cif.gz_A | 1 | 0.119 | 218 | 2.861e-07 | Crystal structure of RNase T in complex with a stem DNA with a 3' overhang |

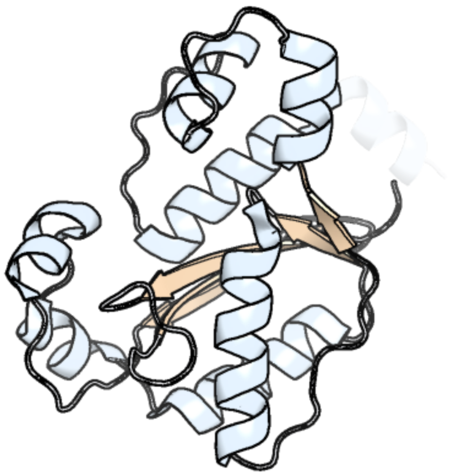

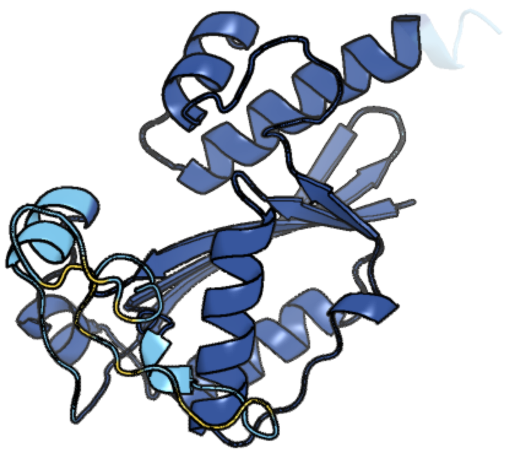

Figure 279: left: reference structure of 8h2f chain A. right: predicted structure of chlorv-1..279, unaligned sequences are shown as transparent

#### chlorv-1..280

- Sequence-based annotation for chlorv-1..280 is hypothetical protein
- Best hit was 3dkq chain B: PKHD-type hydroxylase Sbal_3634

| target | prob | fident | alnlen | evalue | theader |
|---|---|---|---|---|---|
| 3dkq-assembly1.cif.gz_B-2 | 1 | 0.167 | 221 | 1.075e-06 | Crystal structure of Putative Oxygenase (YP_001051978.1) from SHEWANELLA BALTICA OS155 at 2.26 A resolution |
| 3dkq-assembly1.cif.gz_C-2 | 1 | 0.148 | 235 | 2.239e-06 | Crystal structure of Putative Oxygenase (YP_001051978.1) from SHEWANELLA BALTICA OS155 at 2.26 A resolution |
| 4j25-assembly7.cif.gz_G | 1 | 0.138 | 195 | 3.312e-06 | Crystal structure of a Pseudomonas putida prolyl-4-hydroxylase (P4H) |

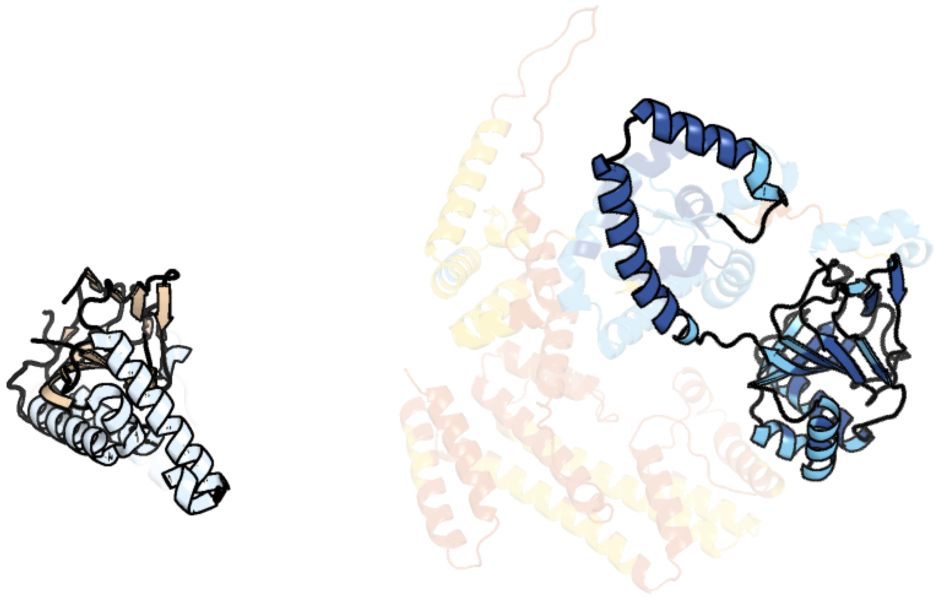

Figure 280: left: reference structure of 3dkq chain B. right: predicted structure of chlorv-1..280, unaligned sequences are shown as transparent

#### chlorv-1..281

- Sequence-based annotation for chlorv-1..281 is hypothetical protein
- No significant structural hit found

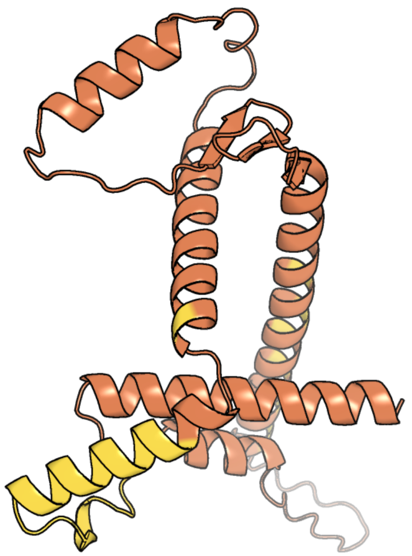

Figure 281: predicted structure of chlorv-1..281

#### chlorv-1..282

- Sequence-based annotation for chlorv-1..282 is putative HNH endonuclease
- No significant structural hit found

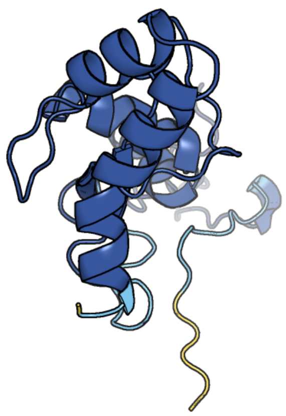

Figure 282: predicted structure of chlorv-1..282

#### chlorv-1..283

- Sequence-based annotation for chlorv-1..283 is hypothetical protein
- Best hit was 6hnq chain Q: Probable ss-1,3-N-acetylglucosaminyltransferase

| target | prob | fident | alnlen | evalue | theader |
|---|---|---|---|---|---|
| 6hnq-assembly4.cif.gz_Q | 1 | 0.158 | 366 | 3.115e-11 | TarP-6RboP-(CH2)6NH2 |
| 6h4m-assembly2.cif.gz_I | 1 | 0.168 | 367 | 3.52e-11 | TarP-UDP-GlcNAc-3RboP |
| 6hnq-assembly2.cif.gz_H | 1 | 0.164 | 370 | 4.23e-11 | TarP-6RboP-(CH2)6NH2 |

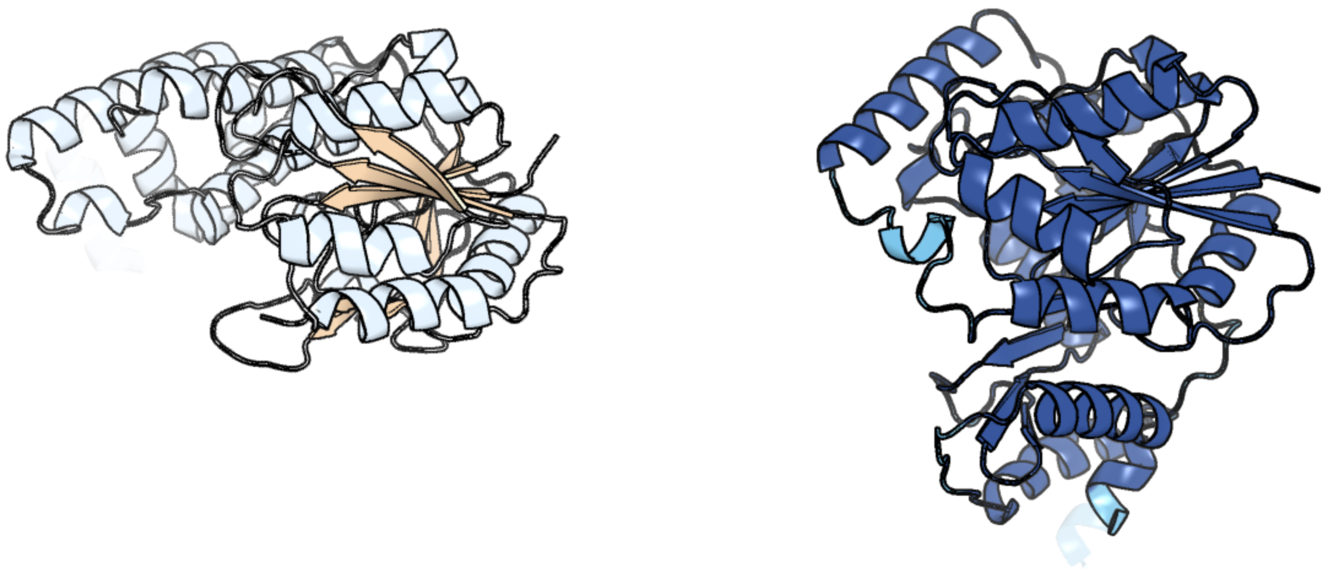

Figure 283: left: reference structure of 6hnq chain Q. right: predicted structure of chlorv-1..283, unaligned sequences are shown as transparent

#### chlorv-1..284

- Sequence-based annotation for chlorv-1..284 is hypothetical protein
- Best hit was 7opk chain A: 5'-3' exoribonuclease

| target | prob | fident | alnlen | evalue | theader |
|---|---|---|---|---|---|
| 7opk-assembly1.cif.gz_A | 1 | 0.135 | 511 | 2.306e-08 | Crystal structure of C. thermophilum Xrn2 |
| 6q8y-assembly1.cif.gz_z | 1 | 0.132 | 722 | 2.64e-08 | Cryo-EM structure of the mRNA translating and degrading yeast 80S ribosome-Xrn1 nuclease complex |
| 3pif-assembly2.cif.gz_B | 1 | 0.131 | 752 | 6.216e-08 | Crystal structure of the 5'->3' exoribonuclease Xrn1, E178Q mutant in Complex with Manganese |

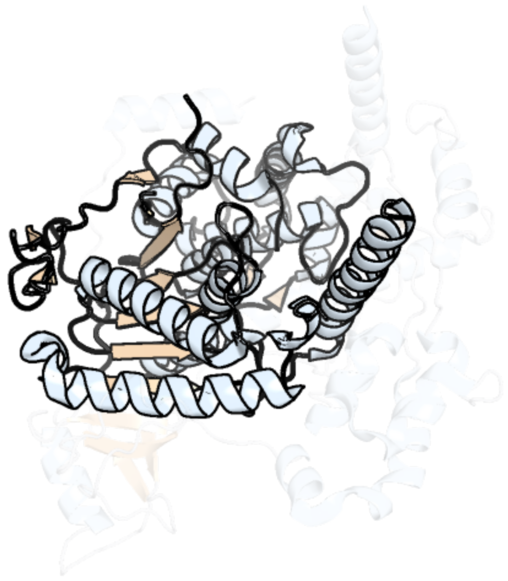

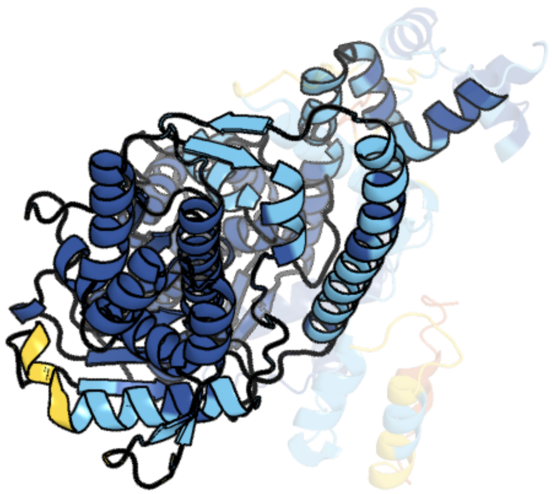

Figure 284: left: reference structure of 7opk chain A. right: predicted structure of chlorv-1..284, unaligned sequences are shown as transparent

#### chlorv-1..285

- Sequence-based annotation for chlorv-1..285 is hypothetical protein
- No significant structural hit found

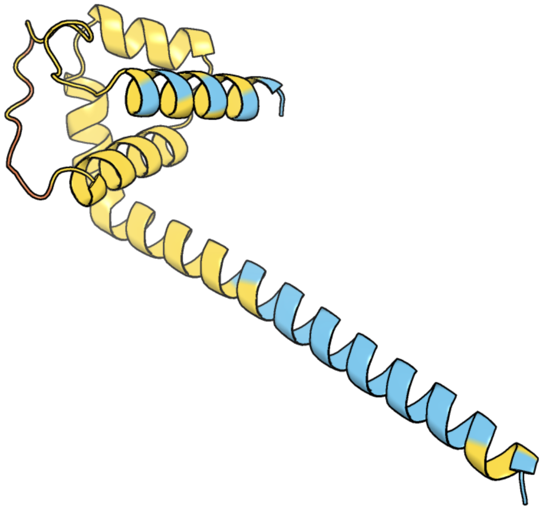

Figure 285: predicted structure of chlorv-1..285

#### chlorv-1..286

- Sequence-based annotation for chlorv-1..286 is hypothetical protein
- No significant structural hit found

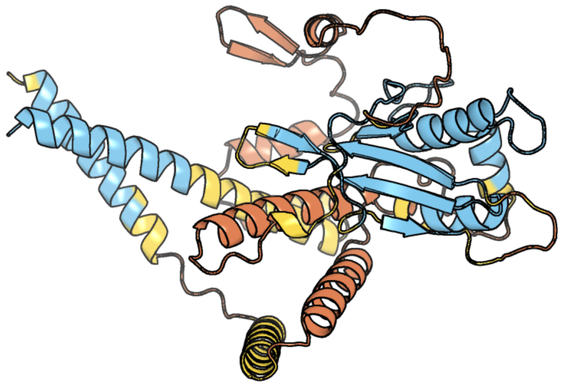

Figure 286: predicted structure of chlorv-1..286

#### chlorv-1..287

- Sequence-based annotation for chlorv-1..287 is hypothetical protein
- Best hit was 5mog chain C: Phytoene dehydrogenase, chloroplastic/chromoplastic

| target | prob | fident | alnlen | evalue | theader |
|---|---|---|---|---|---|
| 5mog-assembly1.cif.gz_C | 1 | 0.124 | 491 | 2.704e-19 | Oryza sativa phytoene desaturase inhibited by norflurazon |
| 3i6d-assembly1.cif.gz_A | 1 | 0.141 | 465 | 3.734e-16 | Crystal structure of PPO from bacillus subtilis with AF |
| 3i6d-assembly2.cif.gz_B | 1 | 0.139 | 465 | 5.635e-16 | Crystal structure of PPO from bacillus subtilis with AF |

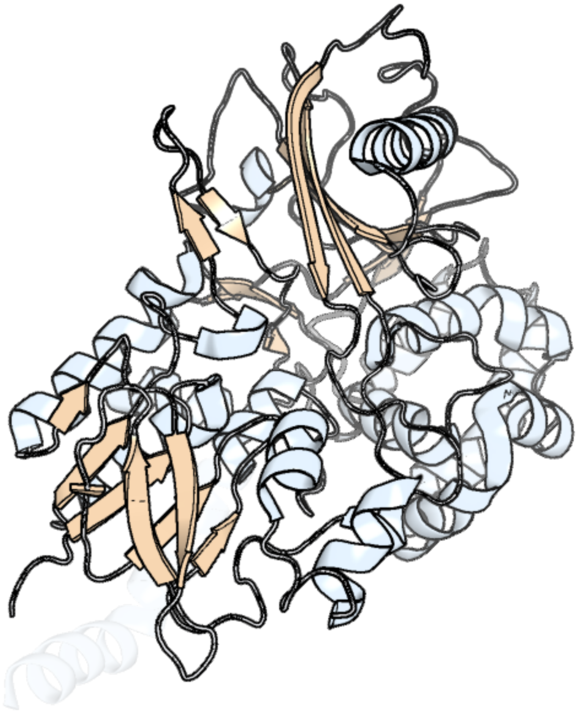

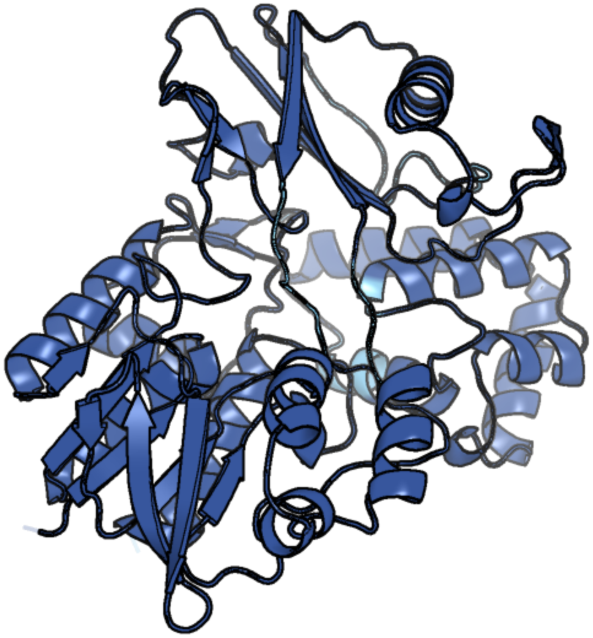

Figure 287: left: reference structure of 5mog chain C. right: predicted structure of chlorv-1..287, unaligned sequences are shown as transparent

#### chlorv-1..288

- Sequence-based annotation for chlorv-1..288 is putative Phosphatase
- No significant structural hit found

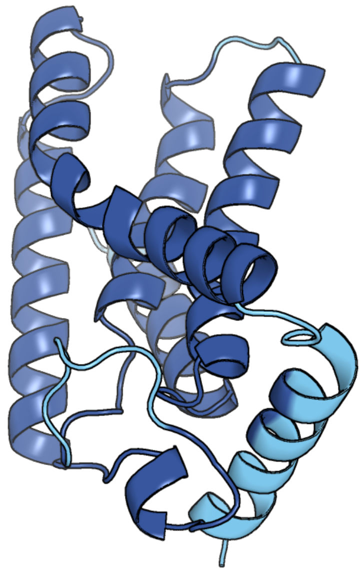

Figure 288: predicted structure of chlorv-1..288

#### chlorv-1..289

- Sequence-based annotation for chlorv-1..289 is putative Helicase
- Best hit was 7amv chain W: ATP-dependent helicase VETFS

| target | prob | fident | alnlen | evalue | theader |
|---|---|---|---|---|---|
| 7amv-assembly1.cif.gz_W | 1 | 0.189 | 592 | 4.272e-22 | Atomic structure of the poxvirus transcription pre-initiation complex in the initially melted state |
| 7aoh-assembly1.cif.gz_Y | 1 | 0.197 | 581 | 1.281e-21 | Atomic structure of the poxvirus late initially transcribing complex |
| 6rfl-assembly1.cif.gz_Y | 1 | 0.202 | 578 | 9.758e-21 | Structure of the complete Vaccinia DNA-dependent RNA polymerase complex |

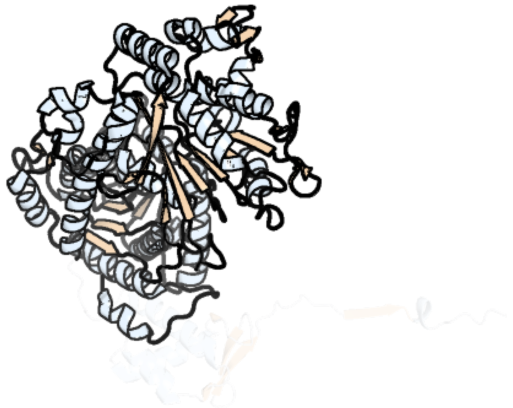

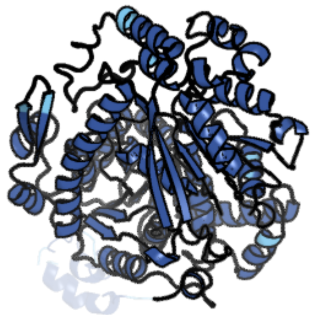

Figure 289: left: reference structure of 7amv chain W. right: predicted structure of chlorv-1..289, unaligned sequences are shown as transparent

#### chlorv-1..290

- Sequence-based annotation for chlorv-1..290 is hypothetical protein
- Best hit was 5mqp chain F: Glycoside hydrolase BT_1002

| target | prob | fident | alnlen | evalue | theader |
|---|---|---|---|---|---|
| 5mqp-assembly6.cif.gz_F | 1 | 0.11 | 753 | 3.281e-17 | Glycoside hydrolase BT_1002 |
| 3vsv-assembly1.cif.gz_B | 1 | 0.095 | 772 | 3.631e-08 | The complex structure of XylC with xylose |
| 4ru5-assembly1.cif.gz_B | 1 | 0.094 | 686 | 2.673e-07 | Crystal Structure of the Pseudomonas phage phi297 tailspike gp61 |

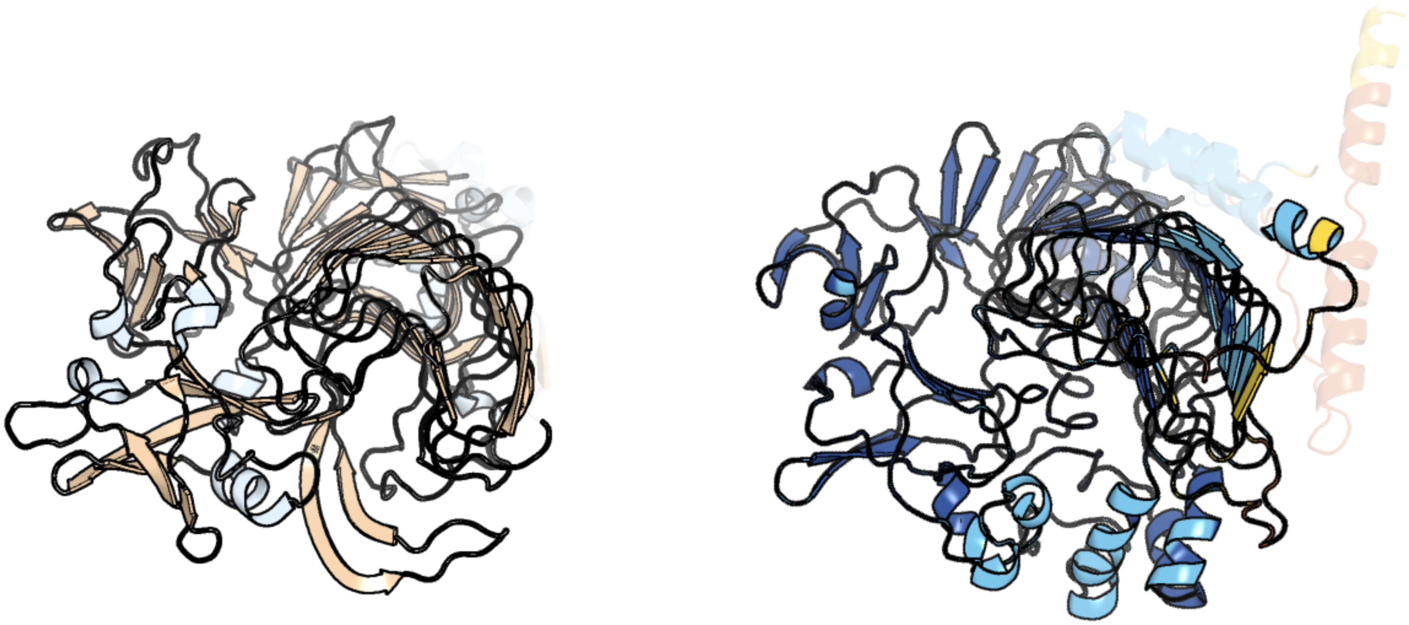

Figure 290: left: reference structure of 5mqp chain F. right: predicted structure of chlorv-1..290, unaligned sequences are shown as transparent

#### chlorv-1..291

- Sequence-based annotation for chlorv-1..291 is hypothetical protein
- No significant structural hit found

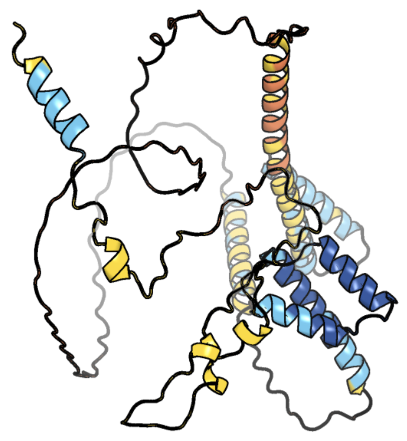

Figure 291: predicted structure of chlorv-1..291

#### chlorv-1..292

- Sequence-based annotation for chlorv-1..292 is putative DnaJ domain
- Best hit was 8j07 chain u: DnaJ homolog subfamily B member 13

| target | prob | fident | alnlen | evalue | theader |
|---|---|---|---|---|---|
| 8j07-assembly1.cif.gz_u | 1 | 0.149 | 361 | 3.479e-12 | 96nm repeat of human respiratory doublet microtubule and associated axonemal complexes |
| 8j07-assembly1.cif.gz_v | 1 | 0.144 | 361 | 1.164e-11 | 96nm repeat of human respiratory doublet microtubule and associated axonemal complexes |
| 4j80-assembly2.cif.gz_D | 1 | 0.117 | 349 | 2.858e-10 | Thermus thermophilus DnaJ |

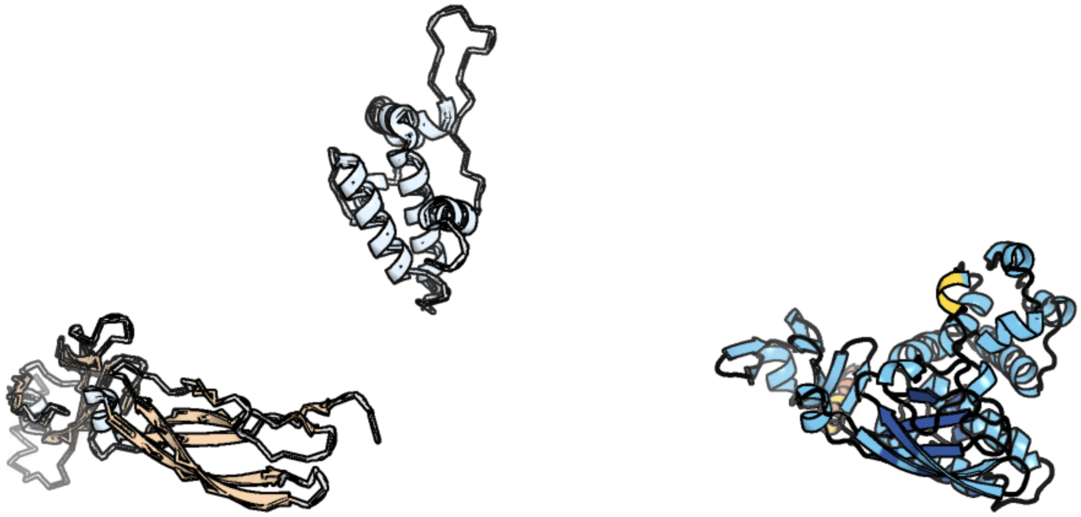

Figure 292: left: reference structure of 8j07 chain u. right: predicted structure of chlorv-1..292, unaligned sequences are shown as transparent

#### chlorv-1..293

- Sequence-based annotation for chlorv-1..293 is hypothetical protein
- No significant structural hit found

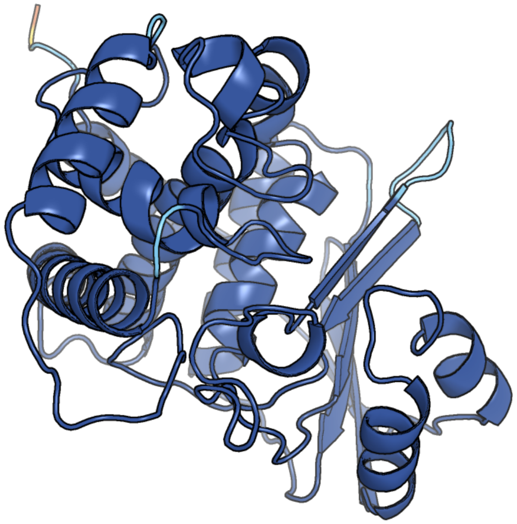

Figure 293: predicted structure of chlorv-1..293

#### chlorv-1..294

- Sequence-based annotation for chlorv-1..294 is hypothetical protein
- No significant structural hit found

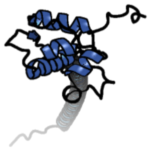

Figure 294: predicted structure of chlorv-1..294

### chlorv-1..295

- Sequence-based annotation for chlorv-1..295 is hypothetical protein
- No significant structural hit found

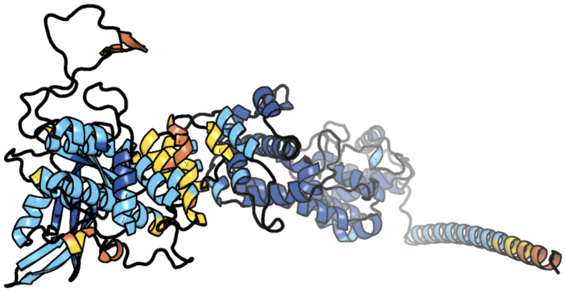

Figure 295: predicted structure of chlorv-1..295

#### chlorv-1..296

- Sequence-based annotation for chlorv-1..296 is hypothetical protein
- No significant structural hit found

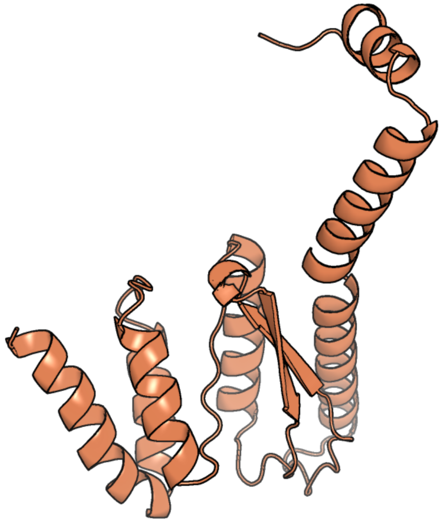

Figure 296: predicted structure of chlorv-1..296

#### chlorv-1..297

- Sequence-based annotation for chlorv-1..297 is hypothetical protein
- No significant structural hit found

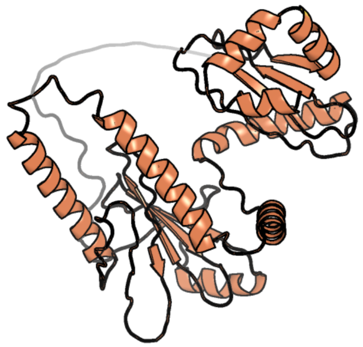

Figure 297: predicted structure of chlorv-1..297

### chlorv-1..298

- Sequence-based annotation for chlorv-1..298 is hypothetical protein
- No significant structural hit found

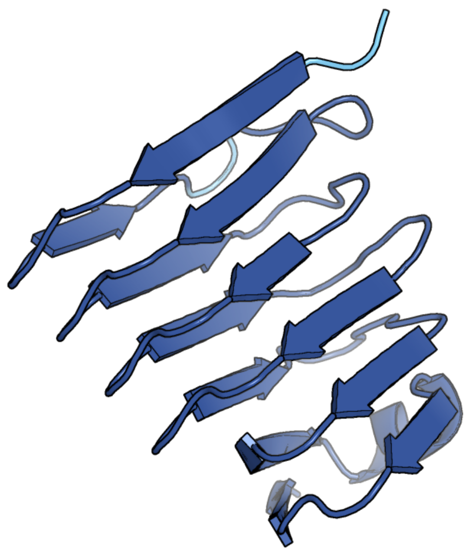

Figure 298: predicted structure of chlorv-1..298

#### chlorv-1..299

- Sequence-based annotation for chlorv-1..299 is hypothetical protein
- Best hit was 6irw chain A: Phosphorylated CTD-interacting factor 1

| target | prob | fident | alnlen | evalue | theader |
|---|---|---|---|---|---|
| 6irw-assembly1.cif.gz_A | 1 | 0.141 | 495 | 2.928e-14 | Crystal structure of the human cap-specific adenosine methyltransferase bound to SAH |
| 6iry-assembly1.cif.gz_A | 1 | 0.153 | 496 | 2.928e-14 | Crystal structure of the zebrafish cap-specific adenosine methyltransferase bound to SAH |
| 6irx-assembly1.cif.gz_A | 1 | 0.145 | 496 | 3.094e-14 | Crystal structure of the zebrafish cap-specific adenosine methyltransferase |

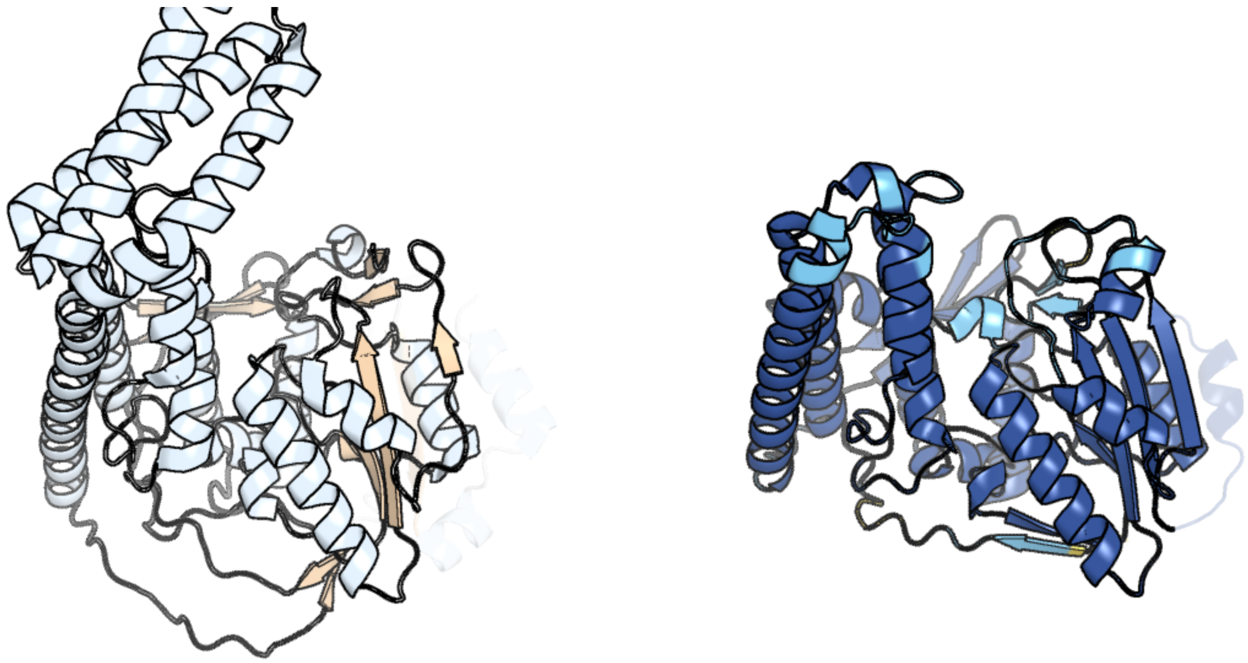

Figure 299: left: reference structure of 6irw chain A. right: predicted structure of chlorv-1..299, unaligned sequences are shown as transparent

#### chlorv-1..300

- Sequence-based annotation for chlorv-1..300 is putative Ubiquitin-conjugating enzyme
- Best hit was 5knl chain C: Ubiquitin-conjugating enzyme E2 15

| target | prob | fident | alnlen | evalue | theader |
|---|---|---|---|---|---|
| 5knl-assembly1.cif.gz_C | 1 | 0.375 | 165 | 5.562e-18 | Crystal structure of S. pombe ubiquitin E1 (Uba1) in complex with Ubc15 and ubiquitin |
| 5knl-assembly2.cif.gz_F | 1 | 0.353 | 164 | 1.306e-17 | Crystal structure of S. pombe ubiquitin E1 (Uba1) in complex with Ubc15 and ubiquitin |
| 3fsh-assembly2.cif.gz_B | 1 | 0.408 | 169 | 1.568e-17 | Crystal structure of the ubiquitin conjugating enzyme Ube2g2 bound to the G2BR domain of ubiquitin ligase gp78 |

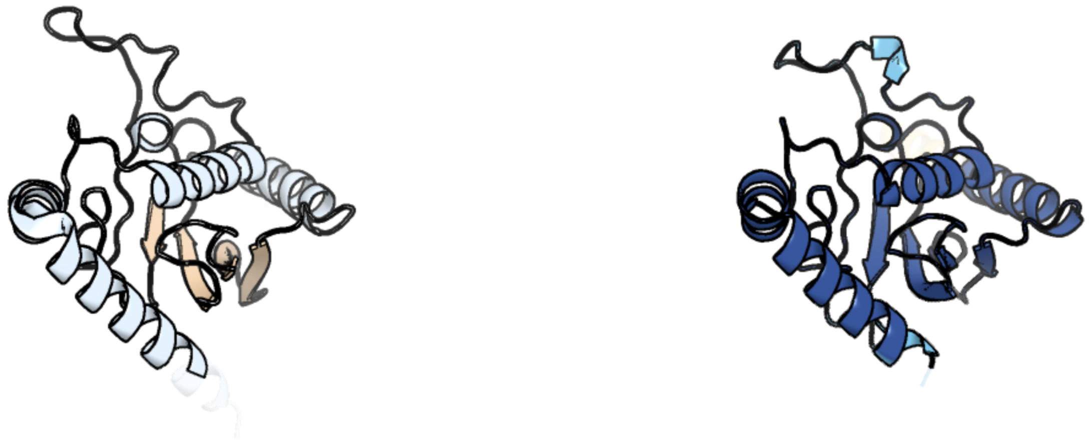

Figure 300: left: reference structure of 5knl chain C. right: predicted structure of chlorv-1..300, unaligned sequences are shown as transparent

#### chlorv-1..301

- Sequence-based annotation for chlorv-1..301 is hypothetical protein
- No significant structural hit found

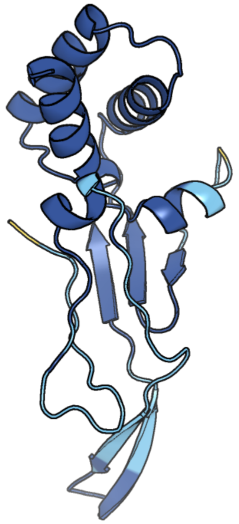

Figure 301: predicted structure of chlorv-1..301

#### chlorv-1..302

- Sequence-based annotation for chlorv-1..302 is putative Aidotransferase
- Best hit was 4zfl chain A: Amidohydrolase EgtC

| target | prob | fident | alnlen | evalue | theader |
|---|---|---|---|---|---|
| 4zfl-assembly1.cif.gz_A | 1 | 0.156 | 255 | 1.277e-13 | Ergothioneine-biosynthetic Ntn hydrolase variant EgtC_C2A with natural substrate |
| 4zfk-assembly1.cif.gz_A | 1 | 0.16 | 256 | 3.191e-13 | Ergothioneine-biosynthetic Ntn hydrolase EgtC with glutamine |
| 4zfj-assembly1.cif.gz_B | 1 | 0.149 | 254 | 1.993e-12 | Ergothioneine-biosynthetic Ntn hydrolase EgtC, apo form |

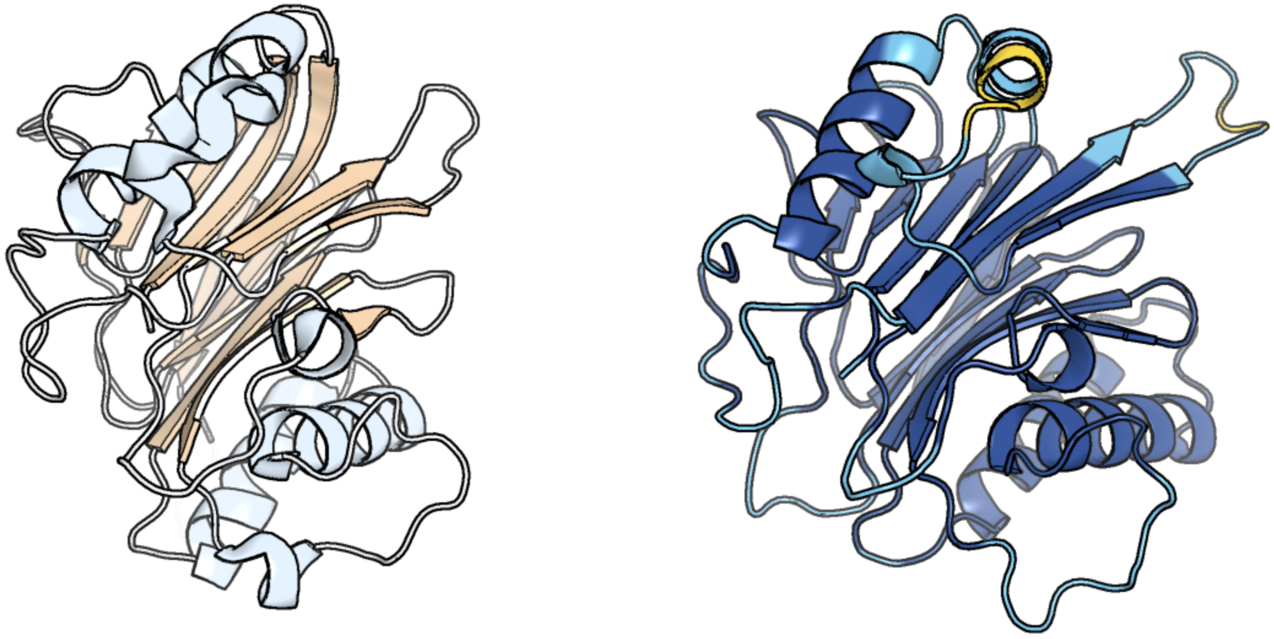

Figure 302: left: reference structure of 4zfl chain A. right: predicted structure of chlorv-1..302, unaligned sequences are shown as transparent

#### chlorv-1..303

- Sequence-based annotation for chlorv-1..303 is hypothetical protein
- No significant structural hit found

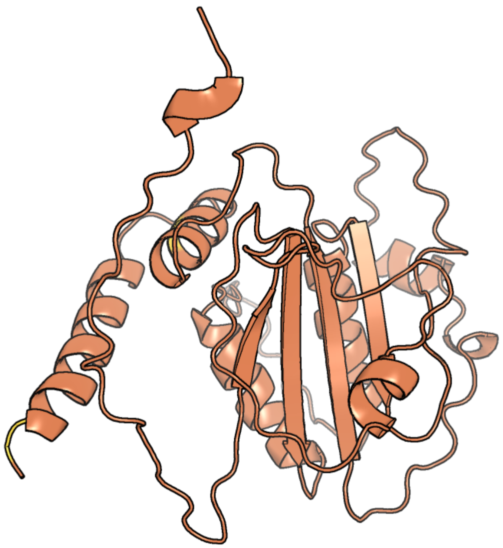

Figure 303: predicted structure of chlorv-1..303

#### chlorv-1..304

- Sequence-based annotation for chlorv-1..304 is hypothetical protein
- No significant structural hit found

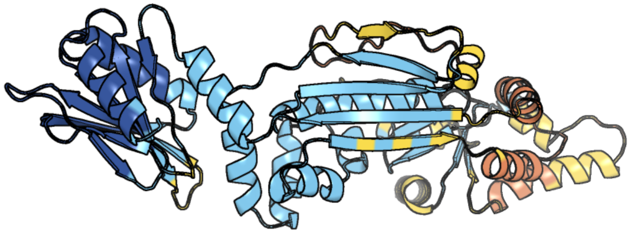

Figure 304: predicted structure of chlorv-1..304

#### chlorv-1..305

- Sequence-based annotation for chlorv-1..305 is hypothetical protein
- Best hit was 7x87 chain B: Beta-galactosidase

| target | prob | fident | alnlen | evalue | theader |
|---|---|---|---|---|---|
| 7x87-assembly1.cif.gz_B | 0.994 | 0.112 | 293 | 3.388e-05 | The complex structure of beta-1,2-glucosyltransferase from Ignavibacterium album with sophotetraose observed as sophorose |
| 7vkx-assembly1.cif.gz_A | 0.992 | 0.111 | 296 | 3.817e-05 | The complex structure of beta-1,2-glucosyltransferase from Ignavibacterium album with glucose |
| 7x87-assembly1.cif.gz_A | 0.992 | 0.116 | 293 | 4.844e-05 | The complex structure of beta-1,2-glucosyltransferase from Ignavibacterium album with sophotetraose observed as sophorose |

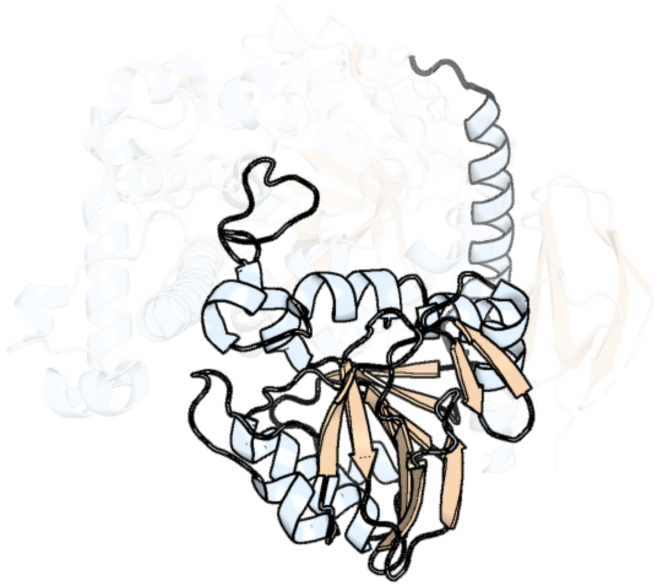

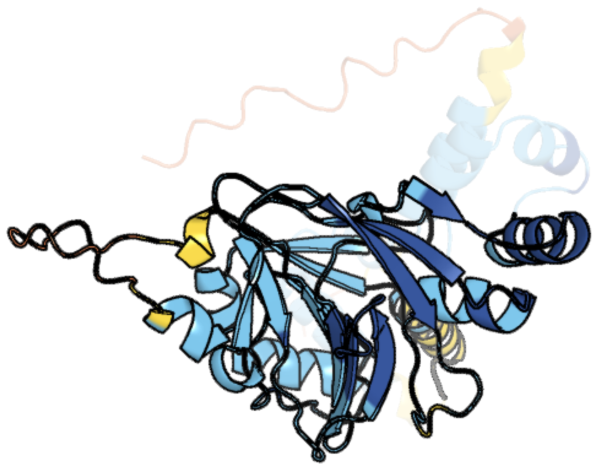

Figure 305: left: reference structure of 7x87 chain B. right: predicted structure of chlorv-1..305, unaligned sequences are shown as transparent

#### chlorv-1..306

- Sequence-based annotation for chlorv-1..306 is hypothetical protein
- No significant structural hit found

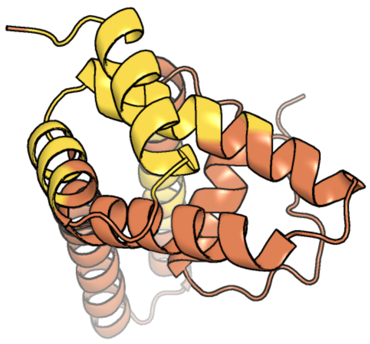

Figure 306: predicted structure of chlorv-1..306

#### chlorv-1..307

- Sequence-based annotation for chlorv-1..307 is hypothetical protein
- Best hit was 2iuw chain A: ALKYLATED REPAIR PROTEIN ALKB HOMOLOG 3

| target | prob | fident | alnlen | evalue | theader |
|---|---|---|---|---|---|
| 2iuw-assembly1.cif.gz_A-2 | 1 | 0.193 | 212 | 5.46e-09 | Crystal structure of human ABH3 in complex with iron ion and 2-oxoglutarate |
| 3rzg-assembly1.cif.gz_A | 1 | 0.183 | 185 | 8.343e-09 | Duplex Interrogation by a Direct DNA Repair Protein in the Search of Damage |
| 3btz-assembly1.cif.gz_A | 1 | 0.176 | 204 | 1.354e-08 | Crystal structure of human ABH2 cross-linked to dsDNA |

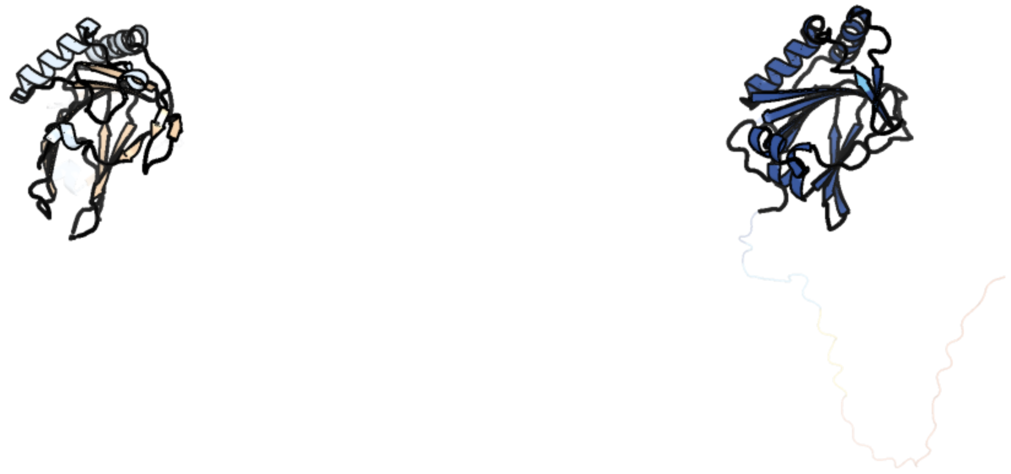

Figure 307: left: reference structure of 2iuw chain A. right: predicted structure of chlorv-1..307, unaligned sequences are shown as transparent

### chlorv-1..308

- Sequence-based annotation for chlorv-1..308 is putative Peptidase
- Best hit was 4boz chain A: UBIQUITIN THIOESTERASE OTU1

| target | prob | fident | alnlen | evalue | theader |
|---|---|---|---|---|---|
| 4boz-assembly1.cif.gz_A | 1 | 0.304 | 161 | 1.517e-15 | Structure of OTUD2 OTU domain in complex with K11-linked di ubiquitin |
| 4bos-assembly1.cif.gz_B | 1 | 0.302 | 162 | 5.371e-15 | Structure of OTUD2 OTU domain in complex with Ubiquitin K11-linked peptide |
| 4boq-assembly1.cif.gz_A-2 | 1 | 0.325 | 163 | 7.259e-15 | Structure of OTUD2 OTU domain |

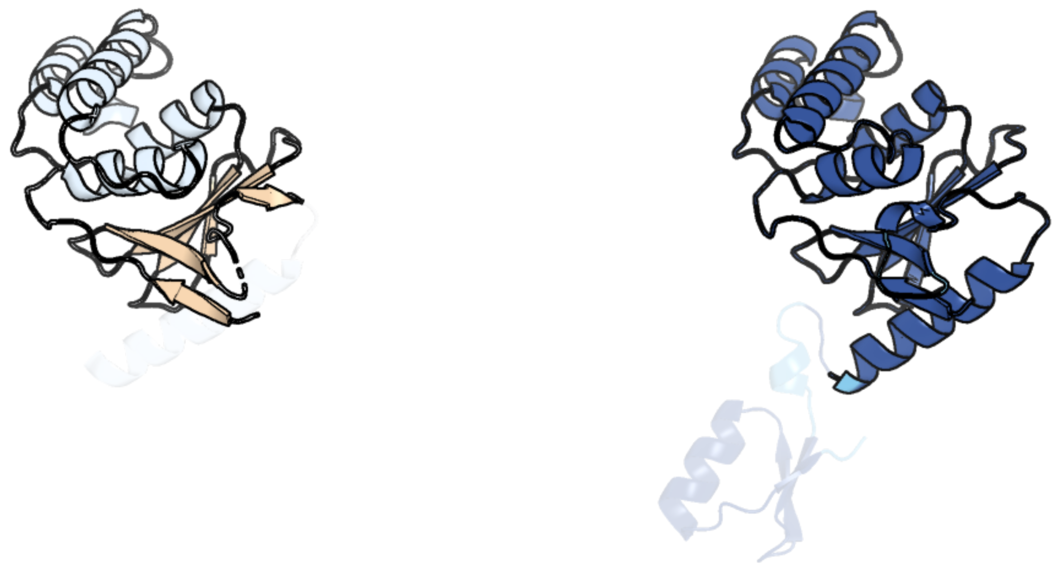

Figure 308: left: reference structure of 4boz chain A. right: predicted structure of chlorv-1..308, unaligned sequences are shown as transparent

#### chlorv-1..309

- Sequence-based annotation for chlorv-1..309 is hypothetical protein
- No significant structural hit found

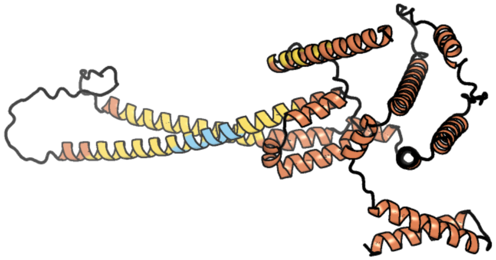

Figure 309: predicted structure of chlorv-1..309

#### chlorv-1..310

- Sequence-based annotation for chlorv-1..310 is hypothetical protein
- No significant structural hit found

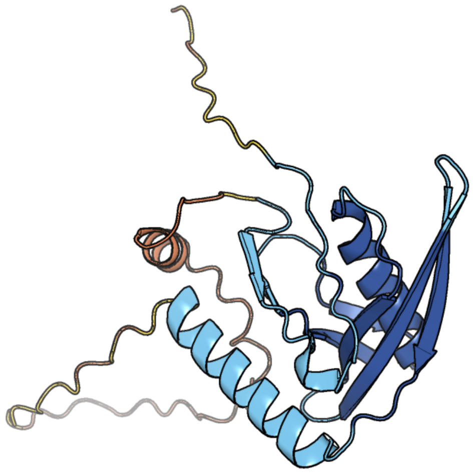

Figure 310: predicted structure of chlorv-1..310

#### chlorv-1..311

- Sequence-based annotation for chlorv-1..311 is putative mRNA capping enzyme
- Best hit was 4ckb chain D: MRNA-CAPPING ENZYME CATALYTIC SUBUNIT

| target | prob | fident | alnlen | evalue | theader |
|---|---|---|---|---|---|
| 4ckb-assembly1.cif.gz_D | 1 | 0.163 | 1047 | 2.578e-40 | Vaccinia virus capping enzyme complexed with GTP and SAH |
| 6rfl-assembly1.cif.gz_O | 1 | 0.17 | 1041 | 3.011e-40 | Structure of the complete Vaccinia DNA-dependent RNA polymerase complex |
| 4ckc-assembly2.cif.gz_D | 1 | 0.17 | 1046 | 4.105e-40 | Vaccinia virus capping enzyme complexed with SAH (monoclinic form) |

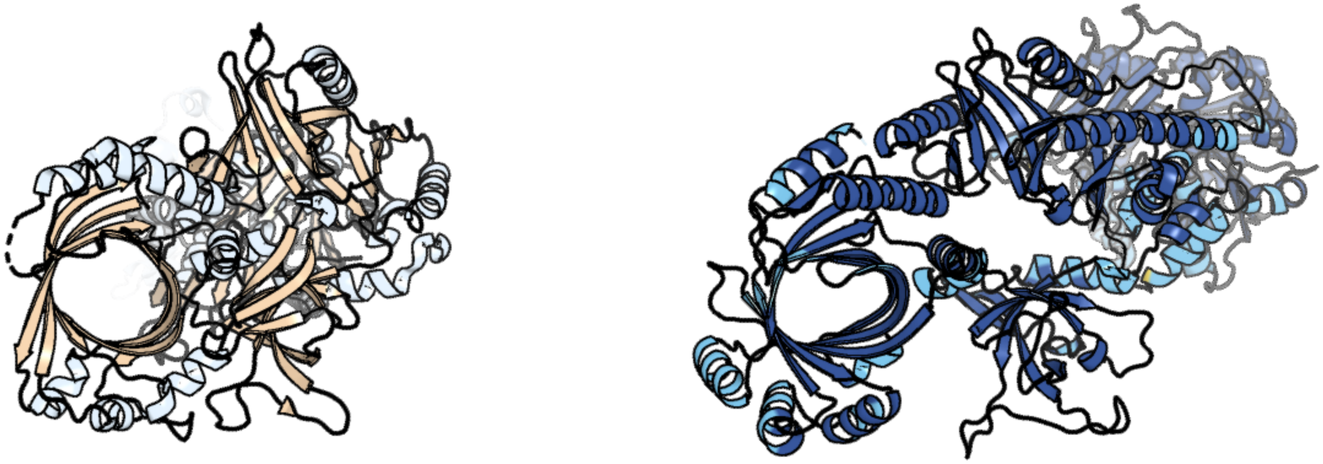

Figure 311: left: reference structure of 4ckb chain D. right: predicted structure of chlorv-1..311, unaligned sequences are shown as transparent

#### chlorv-1..312

- Sequence-based annotation for chlorv-1..312 is putative Poly(A) polymerase catalytic subunit
- Best hit was 4wse chain B: Putative poly(A) polymerase catalytic subunit

| target | prob | fident | alnlen | evalue | theader |
|---|---|---|---|---|---|
| 4wse-assembly1.cif.gz_B | 1 | 0.267 | 464 | 2.245e-37 | Crystal structure of the Mimivirus polyadenylate synthase |
| 4wse-assembly1.cif.gz_A | 1 | 0.263 | 467 | 3.645e-37 | Crystal structure of the Mimivirus polyadenylate synthase |
| 4p37-assembly1.cif.gz_A | 1 | 0.256 | 467 | 1.014e-36 | Crystal structure of the Megavirus polyadenylate synthase |

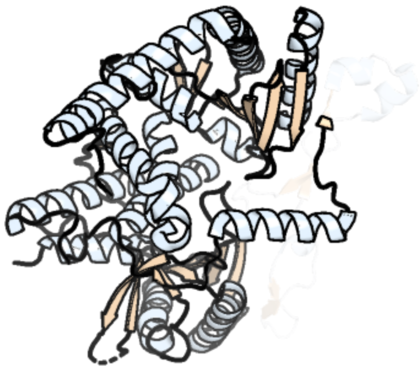

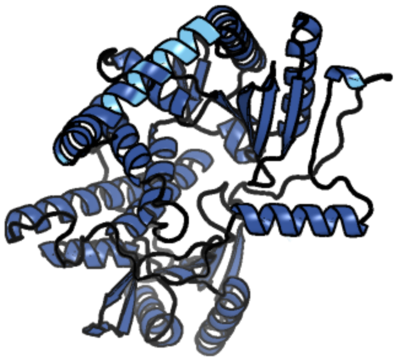

Figure 312: left: reference structure of 4wse chain B. right: predicted structure of chlorv-1..312, unaligned sequences are shown as transparent

#### chlorv-1..313

- Sequence-based annotation for chlorv-1..313 is hypothetical protein
- No significant structural hit found

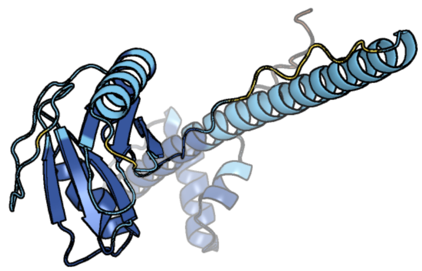

Figure 313: predicted structure of chlorv-1..313

#### chlorv-1..314

- Sequence-based annotation for chlorv-1..314 is putative Endonuclease/Exonuclease/phosphatase
- Best hit was 8igi chain B: Exodeoxyribonuclease (LexA)

| target | prob | fident | alnlen | evalue | theader |
|---|---|---|---|---|---|
| 8igi-assembly2.cif.gz_B | 1 | 0.467 | 261 | 1.679e-34 | Crystal structure of HP1526 (XthA)- a base excision DNA repair protein in Helicobacter pylori |
| 6bov-assembly1.cif.gz_A | 1 | 0.401 | 259 | 2.607e-34 | Human APE1 substrate complex with an A/G mismatch adjacent the THF |
| 2o3c-assembly1.cif.gz_C | 1 | 0.406 | 263 | 4.047e-34 | Crystal structure of zebrafish Ape |

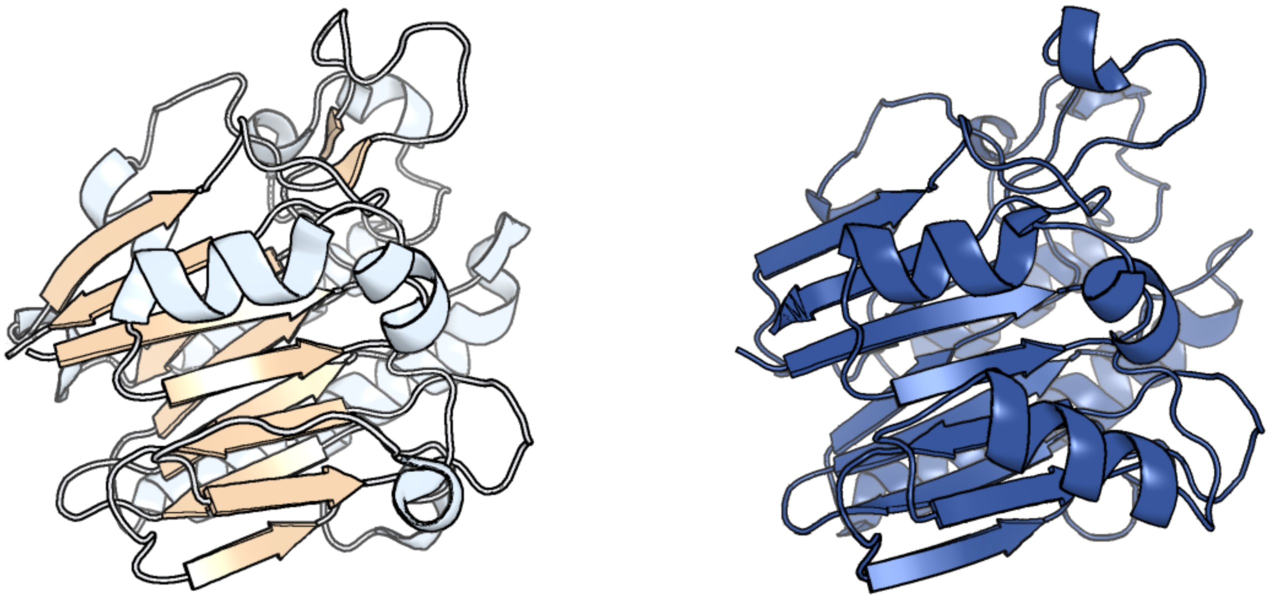

Figure 314: left: reference structure of 8igi chain B. right: predicted structure of chlorv-1..314, unaligned sequences are shown as transparent

### chlorv-1..315

- Sequence-based annotation for chlorv-1..315 is hypothetical protein
- No significant structural hit found

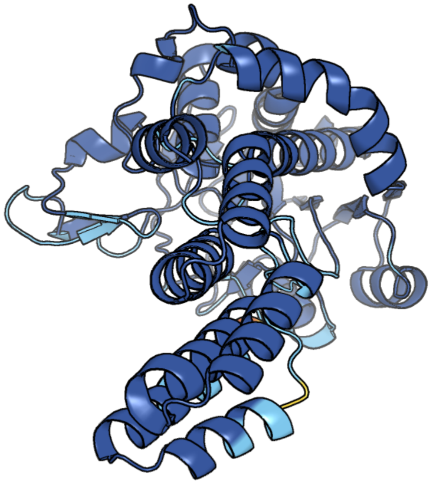

Figure 315: predicted structure of chlorv-1..315

#### chlorv-1..316

- Sequence-based annotation for chlorv-1..316 is putative Threonine synthase
- Best hit was 1vb3 chain A: Threonine synthase

| target | prob | fident | alnlen | evalue | theader |
|---|---|---|---|---|---|
| 1vb3-assembly1.cif.gz_A | 1 | 0.337 | 429 | 1.819e-37 | Crystal Structure of Threonine Synthase from Escherichia coli |
| 4f4f-assembly2.cif.gz_B | 1 | 0.297 | 461 | 5.304e-34 | X-Ray crystal structure of PLP bound Threonine synthase from Brucella melitensis |
| 1kl7-assembly2.cif.gz_B | 1 | 0.293 | 501 | 3.346e-30 | Crystal Structure of Threonine Synthase from Yeast |

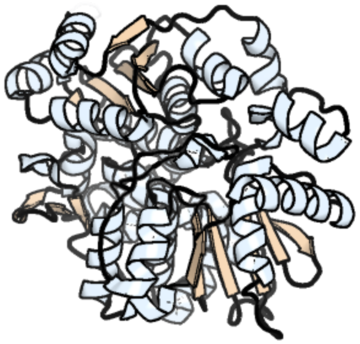

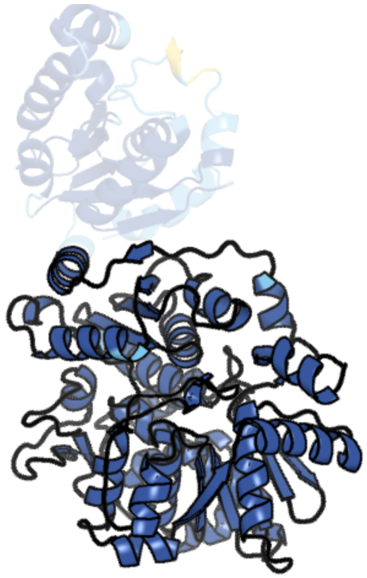

Figure 316: left: reference structure of 1vb3 chain A. right: predicted structure of chlorv-1..316, unaligned sequences are shown as transparent

#### chlorv-1..317

- Sequence-based annotation for chlorv-1..317 is putative Helicase
- Best hit was 6rfl chain Y: Nucleoside triphosphate phosphohydrolase-I

| target | prob | fident | alnlen | evalue | theader |
|---|---|---|---|---|---|
| 6rfl-assembly1.cif.gz_Y | 1 | 0.183 | 773 | 8.165e-31 | Structure of the complete Vaccinia DNA-dependent RNA polymerase complex |
| 7aoh-assembly1.cif.gz_Y | 1 | 0.188 | 778 | 9.101e-31 | Atomic structure of the poxvirus late initially transcribing complex |
| 7tn2-assembly1.cif.gz_W | 1 | 0.145 | 680 | 4.811e-16 | Composite model of a Chd1-nucleosome complex in the nucleotide-free state derived from 2.3A and 2.7A Cryo-EM maps |

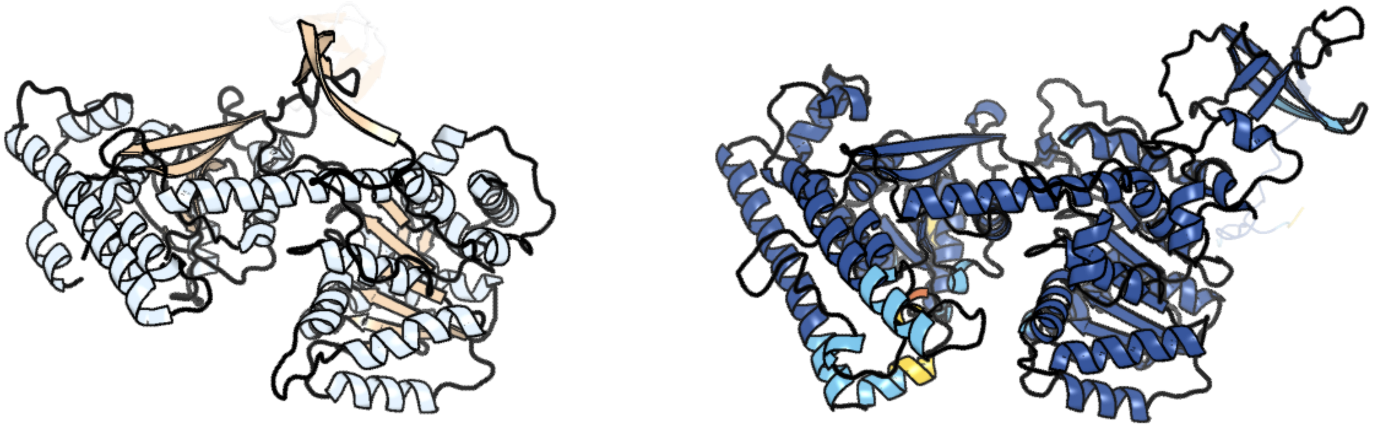

Figure 317: left: reference structure of 6rfl chain Y. right: predicted structure of chlorv-1..317, unaligned sequences are shown as transparent

#### chlorv-1..318

- Sequence-based annotation for chlorv-1..318 is putative Exonuclease/DNA polymerase III subunit epsilon
- Best hit was 8h2f chain A: DnaQ

| target | prob | fident | alnlen | evalue | theader |
|---|---|---|---|---|---|
| 8h2f-assembly1.cif.gz_A | 1 | 0.184 | 206 | 1.804e-09 | Crystal structure of DnaQ domain in complex witn TMP of Streptococcus thermophilus strain DGCC 7710 |
| 4js5-assembly2.cif.gz_B | 1 | 0.196 | 204 | 4.581e-08 | Crystal structure of E. coli Exonuclease I in complex with a dT13 oligonucleotide |
| 6a4b-assembly1.cif.gz_B | 1 | 0.149 | 228 | 9.4e-08 | Structure of TREX2 in complex with a duplex DNA with 2 nucleotide 3'-overhang |

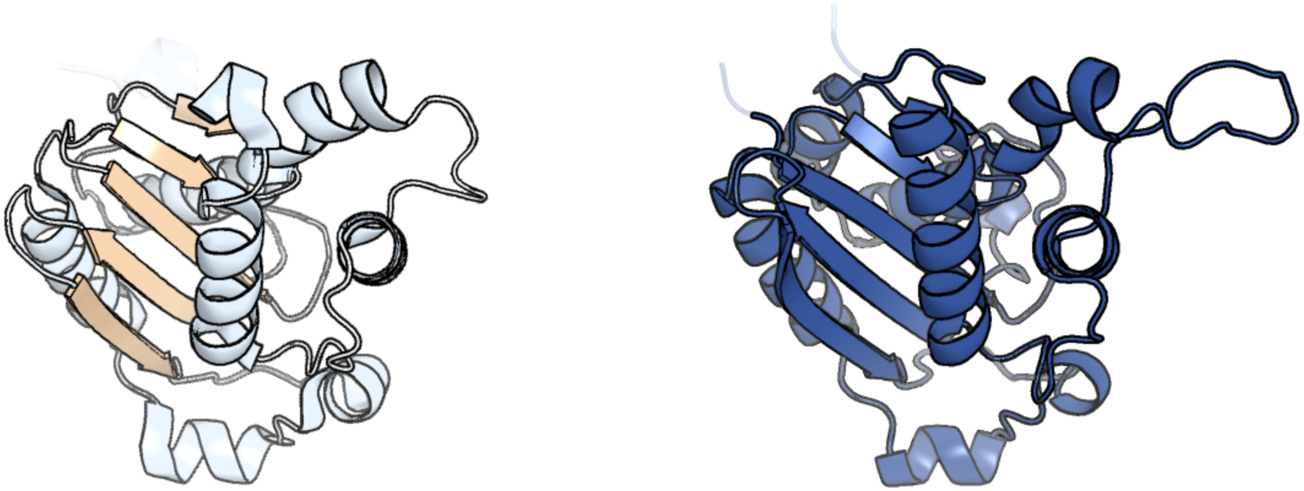

Figure 318: left: reference structure of 8h2f chain A. right: predicted structure of chlorv-1..318, unaligned sequences are shown as transparent

#### chlorv-1..319

- Sequence-based annotation for chlorv-1..319 is hypothetical protein
- No significant structural hit found

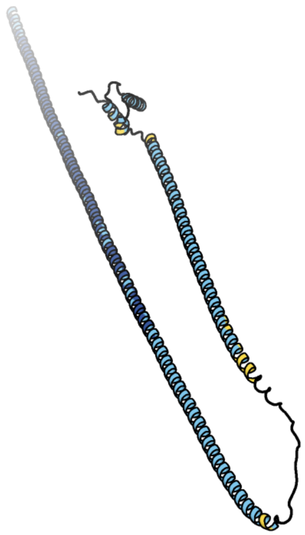

Figure 319: predicted structure of chlorv-1..319

#### chlorv-1..320

- Sequence-based annotation for chlorv-1..320 is putative Helicase
- Best hit was 2is6 chain B: DNA helicase II

| target | prob | fident | alnlen | evalue | theader |
|---|---|---|---|---|---|
| 2is6-assembly1.cif.gz_B | 1 | 0.197 | 639 | 1.594e-30 | Crystal structure of UvrD-DNA-ADPMgF3 ternary complex |
| 2is4-assembly1.cif.gz_B | 1 | 0.196 | 625 | 3.321e-30 | Crystal structure of UvrD-DNA-ADPNP ternary complex |
| 2is4-assembly1.cif.gz_A | 1 | 0.184 | 638 | 1.232e-29 | Crystal structure of UvrD-DNA-ADPNP ternary complex |

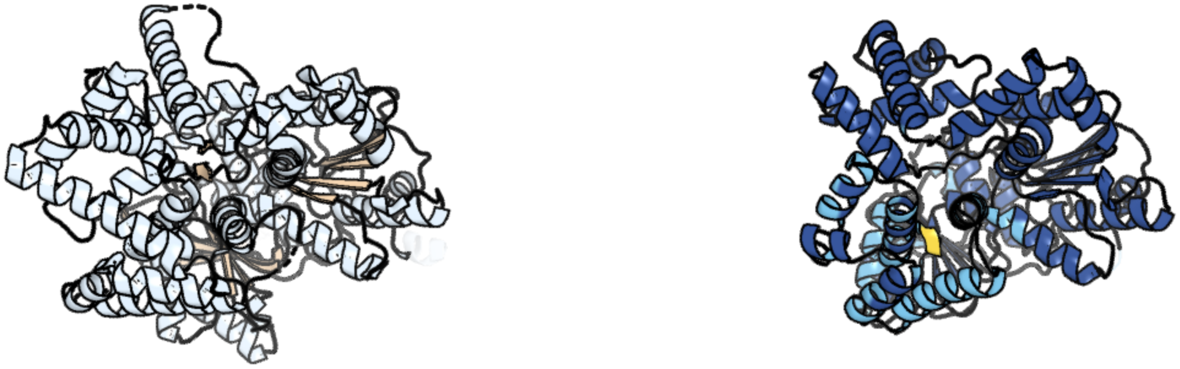

Figure 320: left: reference structure of 2is6 chain B. right: predicted structure of chlorv-1..320, unaligned sequences are shown as transparent

#### chlorv-1..321

- Sequence-based annotation for chlorv-1..321 is hypothetical protein
- No significant structural hit found

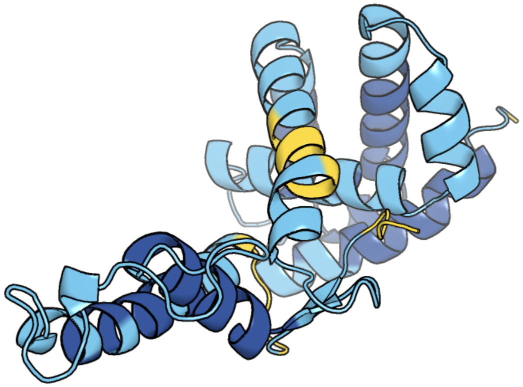

Figure 321: predicted structure of chlorv-1..321

#### chlorv-1..322

- Sequence-based annotation for chlorv-1..322 is hypothetical protein
- No significant structural hit found

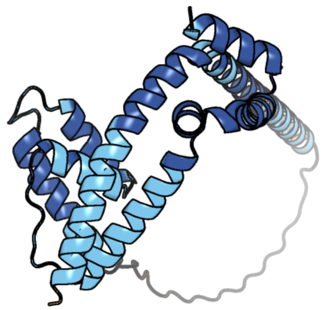

Figure 322: predicted structure of chlorv-1..322

#### chlorv-1..323

- Sequence-based annotation for chlorv-1..323 is putative DAHP synthetase I/aldolase
- Best hit was 5dcd chain B: Phospho-2-dehydro-3-deoxyheptonate aldolase

| target | prob | fident | alnlen | evalue | theader |
|---|---|---|---|---|---|
| 5dcd-assembly1.cif.gz_B | 1 | 0.468 | 344 | 1.708e-43 | Neisseria meningitidis 3-deoxy-D-arabino-heptulosonate 7-phosphate synthase regulated (Tyrosine) |
| 8e0y-assembly1.cif.gz_A | 1 | 0.497 | 340 | 3.3e-43 | DAHP (3-deoxy-D-arabinoheptulosonate-7-phosphate) Synthase complexed with DAHP oxime, Pr(III), and Pi in unbound:(bound)2:other Conformations |
| 1n8f-assembly1.cif.gz_A | 1 | 0.495 | 343 | 3.683e-43 | Crystal structure of E24Q mutant of phenylalanine-regulated 3-deoxy-D-arabino-heptulosonate-7-phosphate synthase (DAHP synthase) from Escherichia Coli in complex with Mn2+ and PEP |

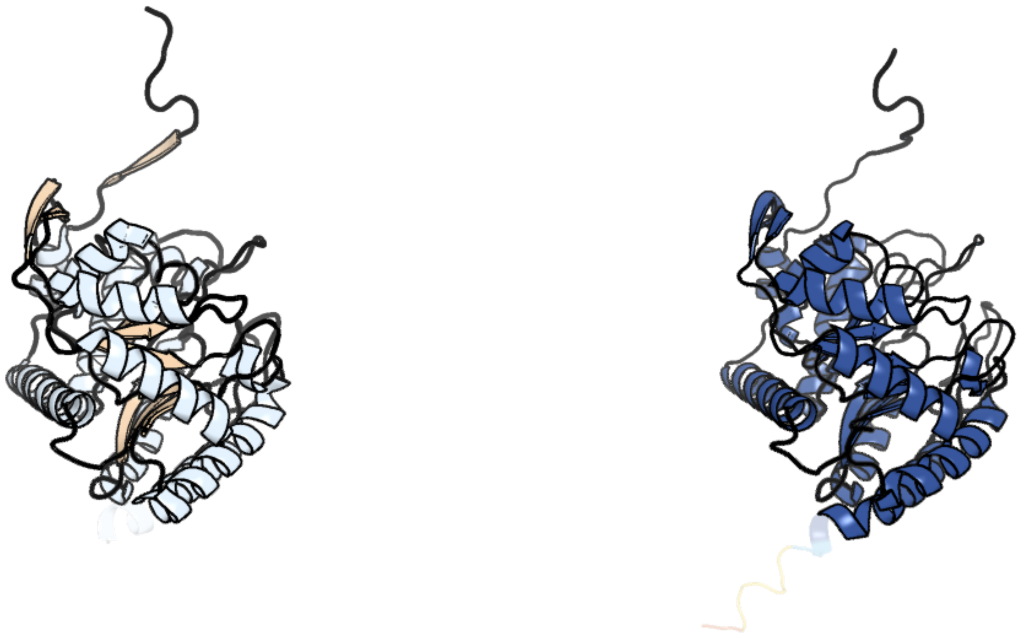

Figure 323: left: reference structure of 5dcd chain B. right: predicted structure of chlorv-1..323, unaligned sequences are shown as transparent

#### chlorv-1..324

- Sequence-based annotation for chlorv-1..324 is putative Dioxygenase
- Best hit was 6d0o chain C: (R)-phenoxypropionate/alpha-ketoglutarate-dioxygenase

| target | prob | fident | alnlen | evalue | theader |
|---|---|---|---|---|---|
| 6d0o-assembly1.cif.gz_C | 1 | 0.218 | 266 | 8.626e-22 | rdpA dioxygenase holoenzyme |
| 6d3i-assembly1.cif.gz_J-2 | 1 | 0.222 | 270 | 1.578e-21 | ftv7 dioxygenase with 2,4-D bound |
| 1gy9-assembly1.cif.gz_B | 1 | 0.257 | 264 | 2.409e-21 | Taurine/alpha-ketoglutarate Dioxygenase from Escherichia coli |

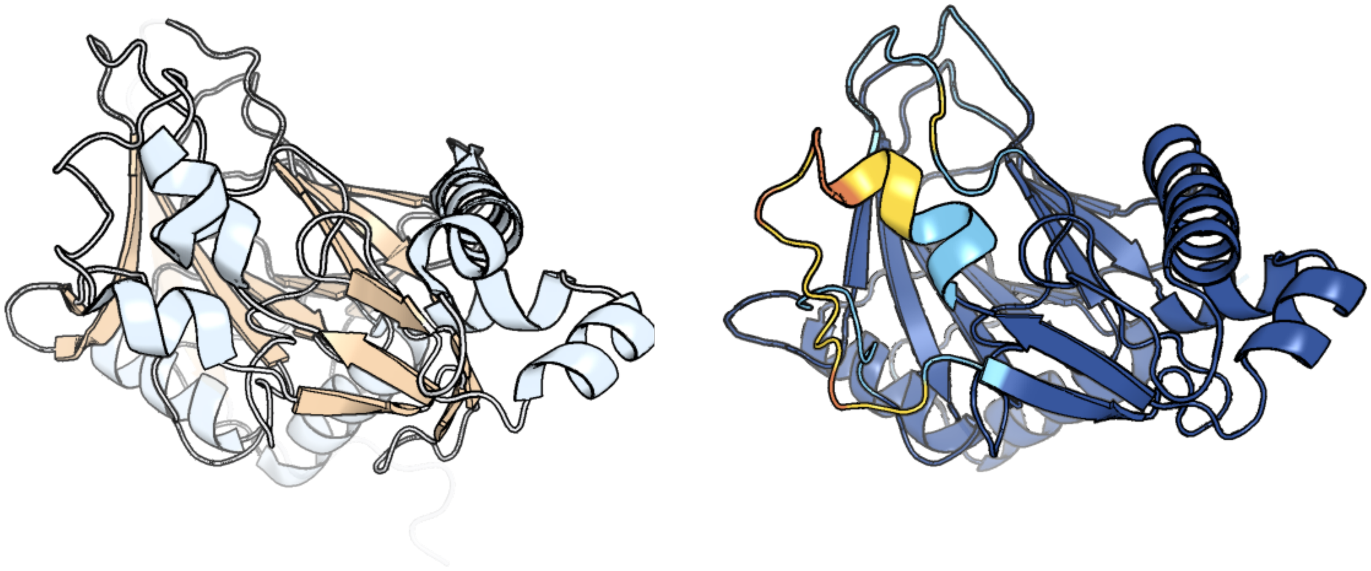

Figure 324: left: reference structure of 6d0o chain C. right: predicted structure of chlorv-1..324, unaligned sequences are shown as transparent

#### chlorv-1..325

- Sequence-based annotation for chlorv-1..325 is putative RNA polymerase Rpb3/Rpb11
- Best hit was 7ok0 chain D: DNA-directed RNA polymerase subunit D

| target | prob | fident | alnlen | evalue | theader |
|---|---|---|---|---|---|
| 7ok0-assembly1.cif.gz_D | 1 | 0.181 | 286 | 6.624e-13 | Cryo-EM structure of the Sulfolobus acidocaldarius RNA polymerase at 2.88 A |
| 4v8s-assembly2.cif.gz_BD | 1 | 0.188 | 286 | 1.227e-12 | Archaeal RNAP-DNA binary complex at 4.32Ang |
| 7oqy-assembly1.cif.gz_D | 1 | 0.169 | 289 | 1.451e-12 | Cryo-EM structure of the cellular negative regulator TFS4 bound to the archaeal RNA polymerase |

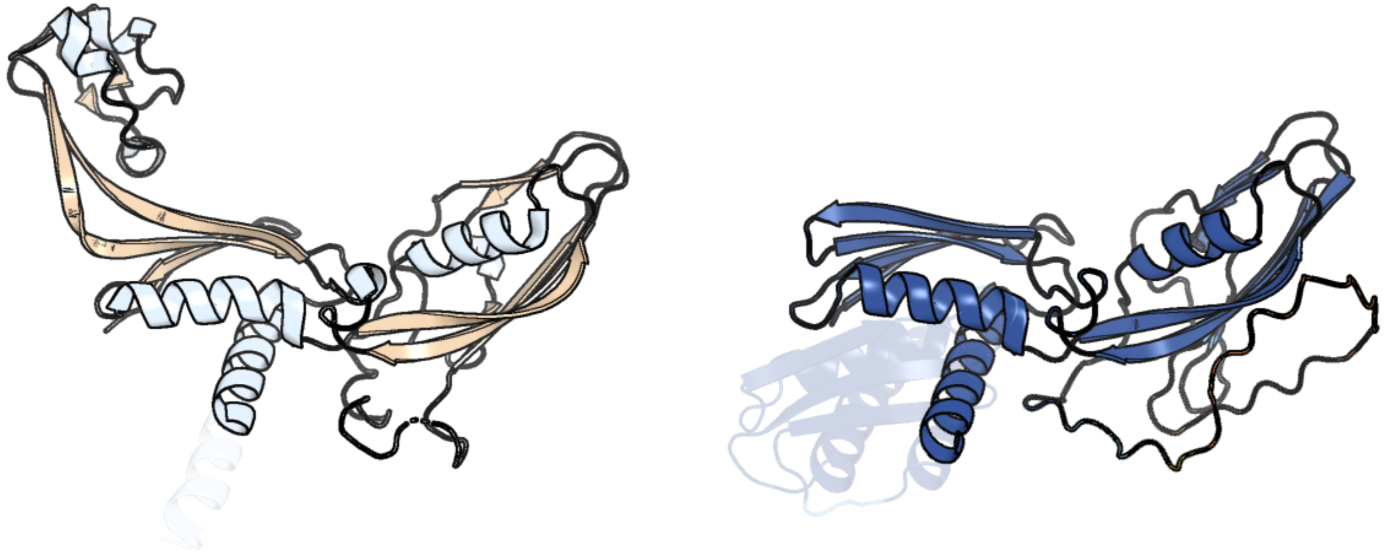

Figure 325: left: reference structure of 7ok0 chain D. right: predicted structure of chlorv-1..325, unaligned sequences are shown as transparent

### chlorv-1..326

- Sequence-based annotation for chlorv-1..326 is hypothetical protein
- No significant structural hit found

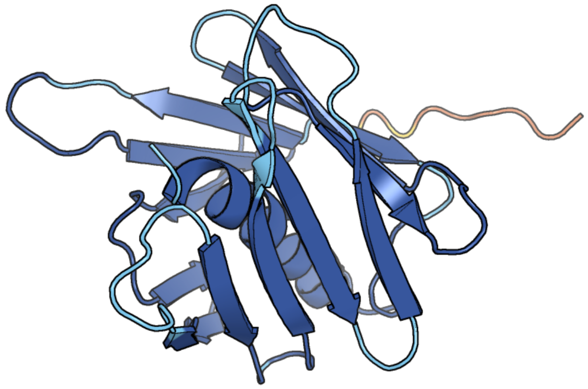

Figure 326: predicted structure of chlorv-1..326

### chlorv-1..327

- Sequence-based annotation for chlorv-1..327 is hypothetical protein
- No significant structural hit found

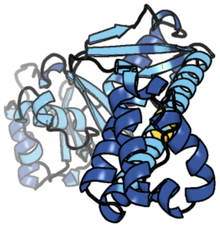

Figure 327: predicted structure of chlorv-1..327

#### chlorv-1..328

- Sequence-based annotation for chlorv-1..328 is putative GTP binding protein
- Best hit was 1eft chain A: ELONGATION FACTOR TU

| target | prob | fident | alnlen | evalue | theader |
|---|---|---|---|---|---|
| 1eft-assembly1.cif.gz_A | 1 | 0.18 | 487 | 2.543e-28 | THE CRYSTAL STRUCTURE OF ELONGATION FACTOR EF-TU FROM THERMUS AQUATICUS IN THE GTP CONFORMATION |
| 1exm-assembly1.cif.gz_A | 1 | 0.181 | 479 | 7.59e-28 | CRYSTAL STRUCTURE OF THERMUS THERMOPHILUS ELONGATION FACTOR TU (EF-TU) IN COMPLEX WITH THE GTP ANALOGUE GPPNHP. |
| 4lbv-assembly1.cif.gz_A | 1 | 0.178 | 481 | 2.399e-27 | Identifying ligand binding hot spots in proteins using brominated fragments |

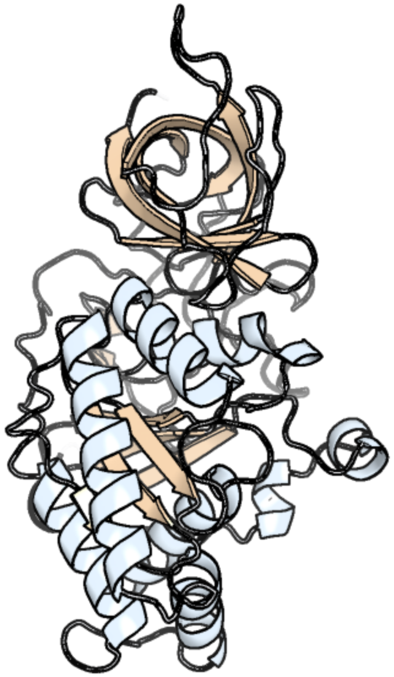

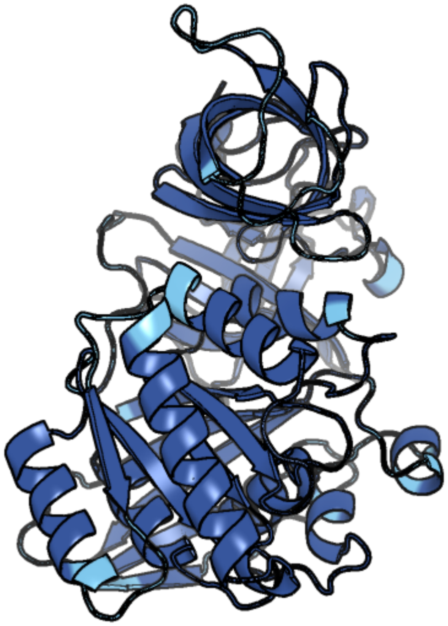

Figure 328: left: reference structure of 1eft chain A. right: predicted structure of chlorv-1..328, unaligned sequences are shown as transparent

#### chlorv-1..329

- Sequence-based annotation for chlorv-1..329 is putative Ribonuclease H
- Best hit was 2qkk chain M: Ribonuclease H1

| target | prob | fident | alnlen | evalue | theader |
|---|---|---|---|---|---|
| 2qkk-assembly4.cif.gz_M | 1 | 0.331 | 151 | 2.446e-11 | Human RNase H catalytic domain mutant D210N in complex with 14-mer RNA/DNA hybrid |
| 2qkk-assembly4.cif.gz_N | 1 | 0.302 | 152 | 6.185e-11 | Human RNase H catalytic domain mutant D210N in complex with 14-mer RNA/DNA hybrid |
| 2qkk-assembly3.cif.gz_J | 1 | 0.324 | 154 | 8.426e-11 | Human RNase H catalytic domain mutant D210N in complex with 14-mer RNA/DNA hybrid |

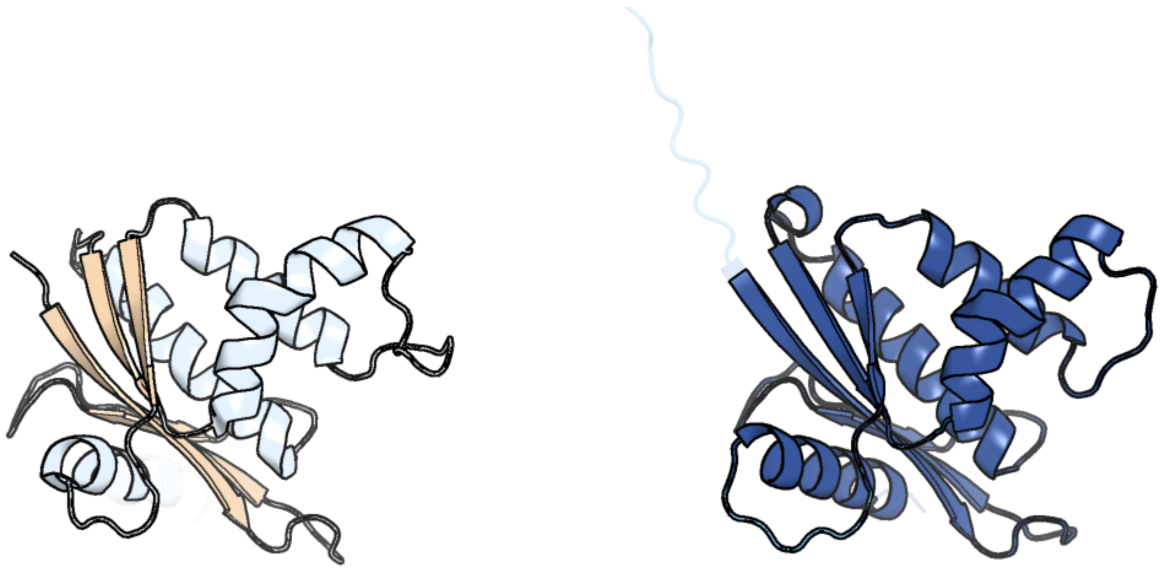

Figure 329: left: reference structure of 2qkk chain M. right: predicted structure of chlorv-1..329, unaligned sequences are shown as transparent

#### chlorv-1..330

- Sequence-based annotation for chlorv-1..330 is hypothetical protein
- No significant structural hit found

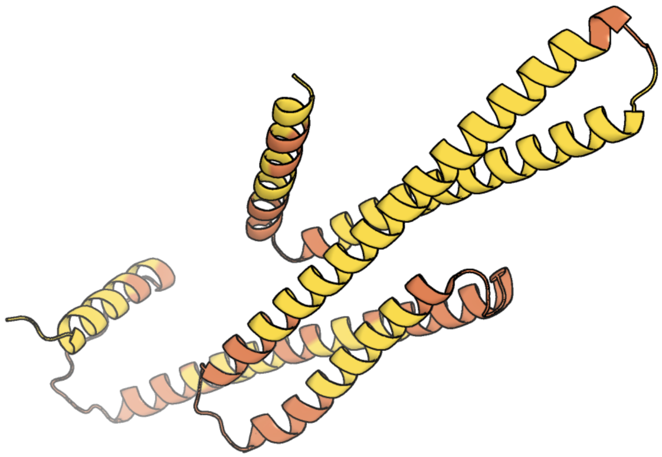

Figure 330: predicted structure of chlorv-1..330

#### chlorv-1..331

- Sequence-based annotation for chlorv-1..331 is putative replication factor C small subunit 2
- Best hit was 7tfh chain B: Replication factor C subunit 4

| target | prob | fident | alnlen | evalue | theader |
|---|---|---|---|---|---|
| 7tfh-assembly1.cif.gz_B | 1 | 0.273 | 318 | 1.618e-20 | Atomic model of the S. cerevisiae clamp-clamp loader complex PCNA-RFC bound to two DNA molecules, one at the 5'-recessed end and the other at the 3'-recessed end |
| 8fs4-assembly1.cif.gz_B | 1 | 0.273 | 318 | 6.574e-20 | Structure of S. cerevisiae Rad24-RFC loading the 9-1-1 clamp onto a 10-nt gapped DNA in step 2 (open 9-1-1 ring and flexibly bound chamber DNA) |
| 7z6h-assembly1.cif.gz_D | 1 | 0.263 | 323 | 3.402e-19 | Structure of DNA-bound human RAD17-RFC clamp loader and 9-1-1 checkpoint clamp |

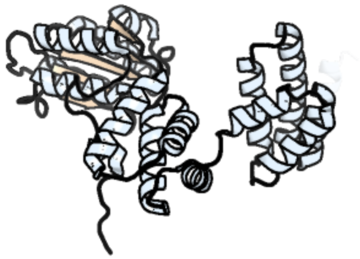

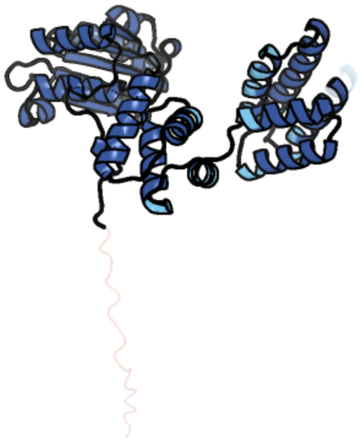

Figure 331: left: reference structure of 7tfh chain B. right: predicted structure of chlorv-1..331, unaligned sequences are shown as transparent

#### chlorv-1..332

- Sequence-based annotation for chlorv-1..332 is hypothetical protein
- No significant structural hit found

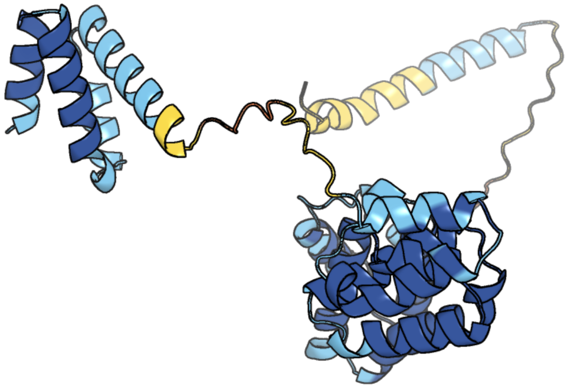

Figure 332: predicted structure of chlorv-1..332

#### chlorv-1..333

- Sequence-based annotation for chlorv-1..333 is putative Hsp70 protein
- Best hit was 5e84 chain F: 78 kDa glucose-regulated protein

| target | prob | fident | alnlen | evalue | theader |
|---|---|---|---|---|---|
| 5e84-assembly6.cif.gz_F | 1 | 0.541 | 615 | 1.933e-70 | ATP-bound state of BiP |
| 7n1r-assembly1.cif.gz_A | 1 | 0.544 | 613 | 3.322e-70 | A novel and unique ATP hydrolysis to AMP by a human Hsp70 BiP |
| 5tky-assembly1.cif.gz_A | 1 | 0.499 | 609 | 5.228e-69 | Crystal structure of the co-translational Hsp70 chaperone Ssb in the ATP-bound, open conformation |

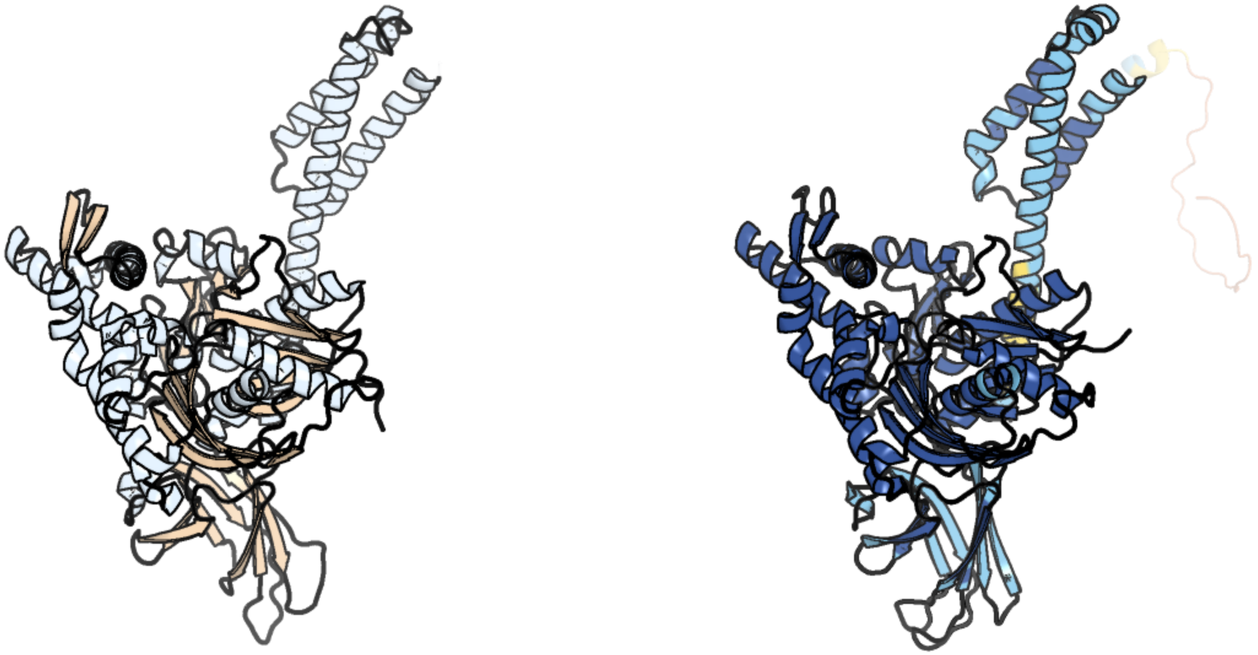

Figure 333: left: reference structure of 5e84 chain F. right: predicted structure of chlorv-1..333, unaligned sequences are shown as transparent

#### chlorv-1..334

- Sequence-based annotation for chlorv-1..334 is hypothetical protein
- No significant structural hit found

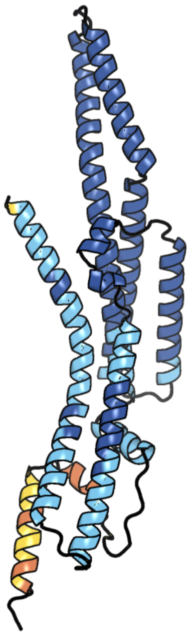

Figure 334: predicted structure of chlorv-1..334

### chlorv-1..335

- Sequence-based annotation for chlorv-1..335 is hypothetical protein
- No significant structural hit found

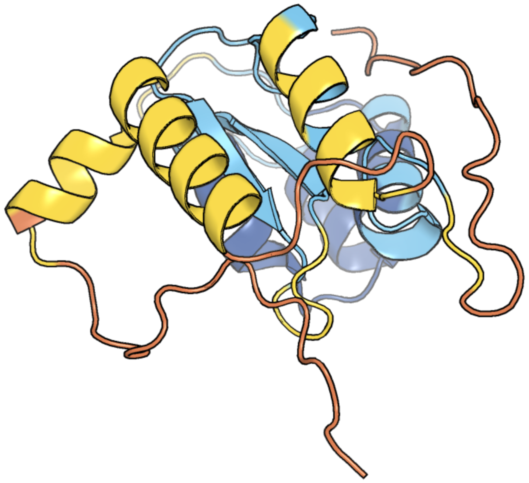

Figure 335: predicted structure of chlorv-1..335

#### chlorv-1..336

- Sequence-based annotation for chlorv-1..336 is putative DNA photolyase
- Best hit was 2e0i chain B: 432aa long hypothetical deoxyribodipyrimidine photolyase

| target | prob | fident | alnlen | evalue | theader |
|---|---|---|---|---|---|
| 2e0i-assembly2.cif.gz_B | 1 | 0.372 | 465 | 7.309e-34 | Crystal structure of archaeal photolyase from Sulfolobus tokodaii with two FAD molecules: Implication of a novel light-harvesting cofactor |
| 1owo-assembly1.cif.gz_A | 1 | 0.283 | 484 | 3.505e-33 | DATA4:photoreduced DNA photolyase / received X-rays dose 1.2 exp15 photons/mm2 |
| 4u63-assembly1.cif.gz_A | 1 | 0.289 | 487 | 3.866e-31 | Crystal structure of a bacterial class III photolyase from Agrobacterium tumefaciens at 1.67A resolution |

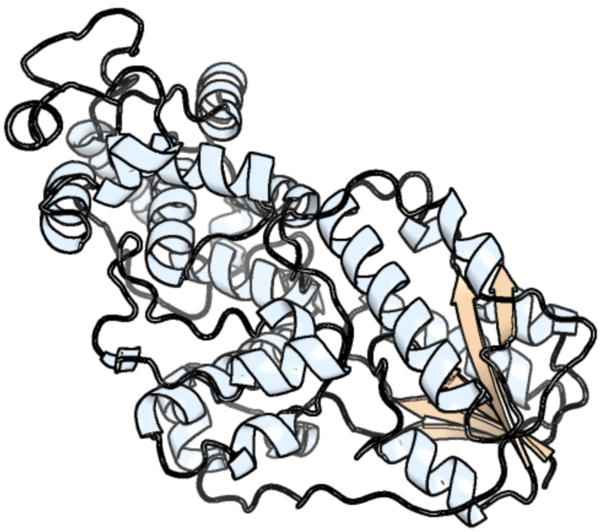

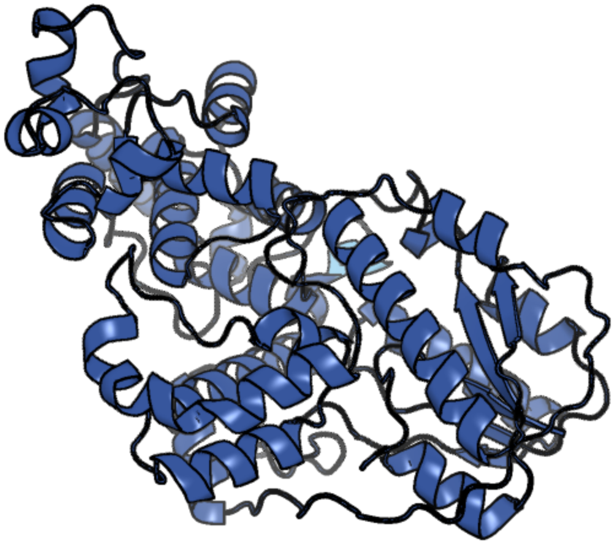

Figure 336: left: reference structure of 2e0i chain B. right: predicted structure of chlorv-1..336, unaligned sequences are shown as transparent

#### chlorv-1..337

- Sequence-based annotation for chlorv-1..337 is hypothetical protein
- No significant structural hit found

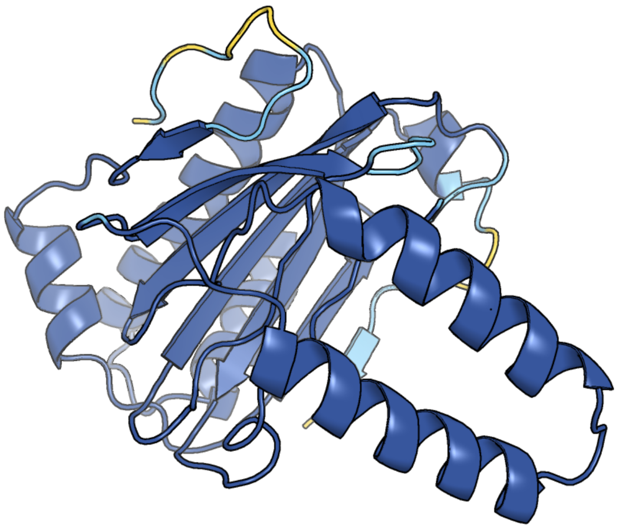

Figure 337: predicted structure of chlorv-1..337

#### chlorv-1..338

- Sequence-based annotation for chlorv-1..338 is hypothetical protein
- No significant structural hit found

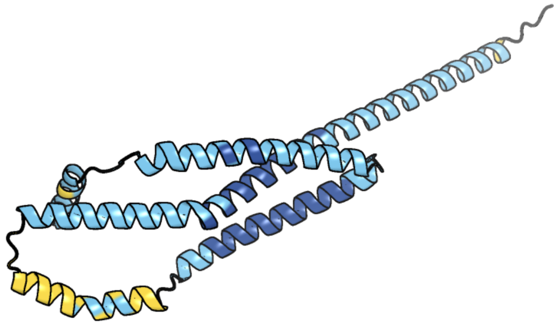

Figure 338: predicted structure of chlorv-1..338

#### chlorv-1..339

- Sequence-based annotation for chlorv-1..339 is putative Helicase/Zinc finger RING-type
- Best hit was 6l8n chain A: DNA repair protein RAD5

| target | prob | fident | alnlen | evalue | theader |
|---|---|---|---|---|---|
| 6l8n-assembly1.cif.gz_A | 1 | 0.171 | 934 | 3.466e-29 | Crystal structure of the K. lactis Rad5 |
| 7r78-assembly1.cif.gz_A | 1 | 0.153 | 962 | 3.875e-29 | cryo-EM structure of DNMT5 quaternary complex with hemimethylated DNA, AMP-PNP and SAH |
| 7r77-assembly1.cif.gz_A | 1 | 0.149 | 978 | 9.992e-29 | Cryo-EM structure of DNMT5 binary complex with hemimethylated DNA |

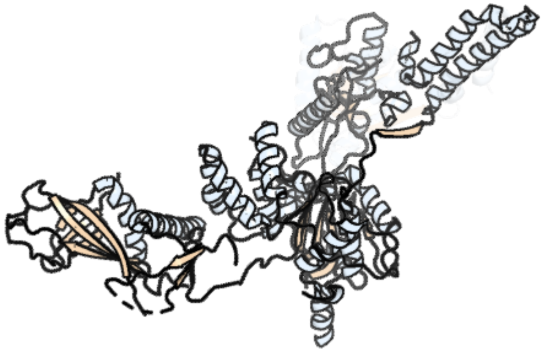

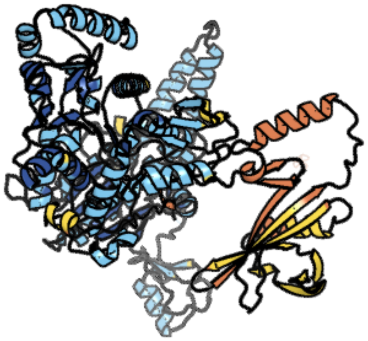

Figure 339: left: reference structure of 6l8n chain A. right: predicted structure of chlorv-1..339, unaligned sequences are shown as transparent

#### chlorv-1..340

- Sequence-based annotation for chlorv-1..340 is hypothetical protein
- Best hit was 2z0t chain D: Putative uncharacterized protein PH0355

| target | prob | fident | alnlen | evalue | theader |
|---|---|---|---|---|---|
| 2z0t-assembly4.cif.gz_D | 1 | 0.186 | 102 | 1.459e-05 | Crystal structure of hypothetical protein PH0355 |
| 1s04-assembly1.cif.gz_A | 1 | 0.186 | 102 | 0.0002177 | Solution NMR Structure of Protein PF0455 from Pyrococcus furiosus. Northeast Structural Genomics Consortium Target PfR13 |
| 1xne-assembly1.cif.gz_A | 1 | 0.137 | 102 | 0.0009626 | Solution Structure of Pyrococcus furiosus Protein PF0470: The Northeast Structural Genomics Consortium Target PfR14 |

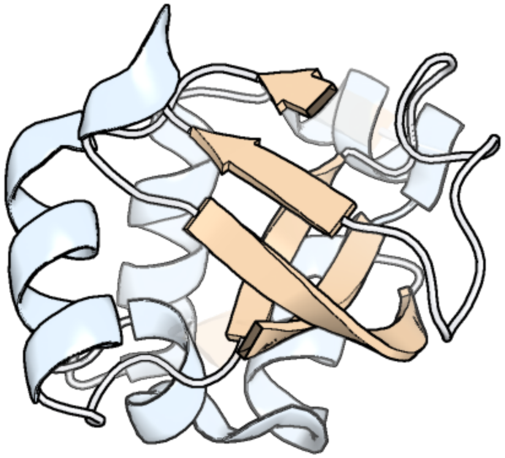

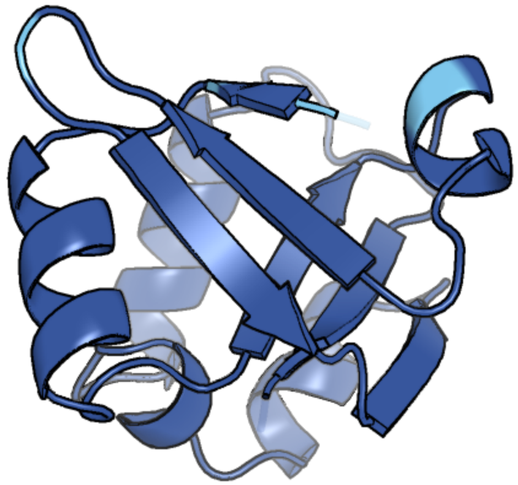

Figure 340: left: reference structure of 2z0t chain D. right: predicted structure of chlorv-1..340, unaligned sequences are shown as transparent

#### chlorv-1..341

- Sequence-based annotation for chlorv-1..341 is hypothetical protein
- Best hit was 2q3l chain A: Uncharacterized protein

| target | prob | fident | alnlen | evalue | theader |
|---|---|---|---|---|---|
| 2q3l-assembly2.cif.gz_A-2 | 1 | 0.097 | 123 | 1.407e-05 | CRYSTAL STRUCTURE OF AN UNCHARACTERIZED PROTEIN FROM DUF3478 FAMILY WITH A SPOIIAA-LIKE FOLD (SHEW_3102) FROM SHEWANELLA LOIHICA PV-4 AT 2.25 A RESOLUTION |
| 1vc1-assembly1.cif.gz_B | 1 | 0.177 | 118 | 4.868e-05 | Crystal structure of the TM1442 protein from Thermotoga maritima, a homolog of the Bacillus subtilis general stress response anti-anti-sigma factor RsbV |
| 6m36-assembly1.cif.gz_H | 1 | 0.155 | 109 | 8.211e-05 | The crystal structure of B. subtilis RsbV/RsbW complex in the monoclinic crystal form |

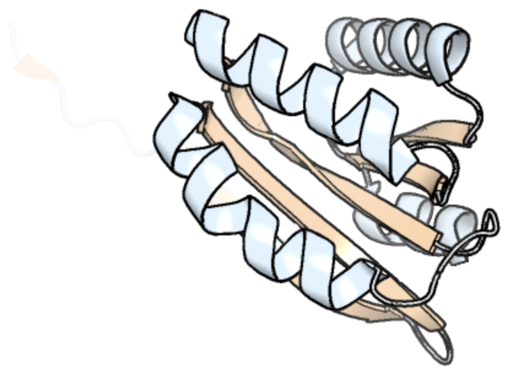

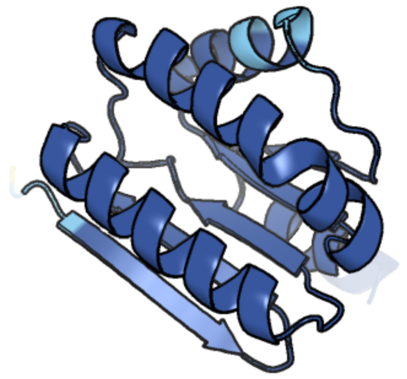

Figure 341: left: reference structure of 2q3l chain A. right: predicted structure of chlorv-1..341, unaligned sequences are shown as transparent

#### chlorv-1..342

- Sequence-based annotation for chlorv-1..342 is hypothetical protein
- No significant structural hit found

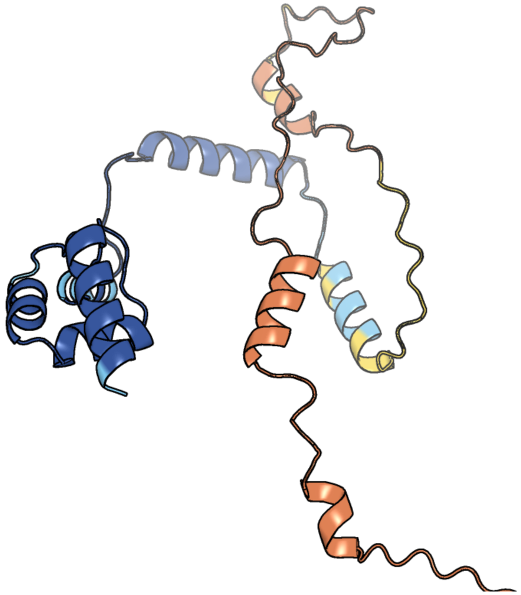

Figure 342: predicted structure of chlorv-1..342

#### chlorv-1..343

- Sequence-based annotation for chlorv-1..343 is putative Asparagine synthase
- Best hit was 6gq3 chain B: Asparagine synthetase [glutamine-hydrolyzing]

| target | prob | fident | alnlen | evalue | theader |
|---|---|---|---|---|---|
| 6gq3-assembly2.cif.gz_B | 1 | 0.333 | 548 | 6.214e-39 | Human asparagine synthetase (ASNS) in complex with 6-diazo-5-oxo-L-norleucine (DON) at 1.85 A resolution |
| 1ct9-assembly1.cif.gz_A | 1 | 0.314 | 535 | 2.494e-38 | CRYSTAL STRUCTURE OF ASPARAGINE SYNTHETASE B FROM ESCHERICHIA COLI |
| 1ct9-assembly1.cif.gz_D | 1 | 0.305 | 533 | 4.641e-37 | CRYSTAL STRUCTURE OF ASPARAGINE SYNTHETASE B FROM ESCHERICHIA COLI |

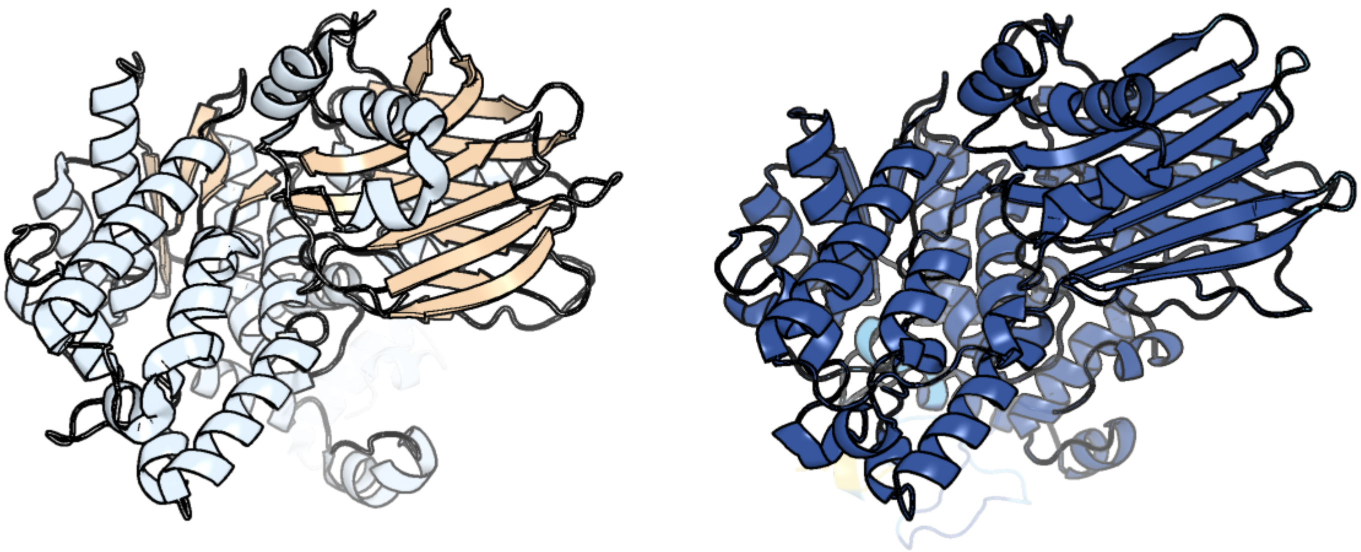

Figure 343: left: reference structure of 6gq3 chain B. right: predicted structure of chlorv-1..343, unaligned sequences are shown as transparent

#### chlorv-1..344

- Sequence-based annotation for chlorv-1..344 is hypothetical protein
- No significant structural hit found

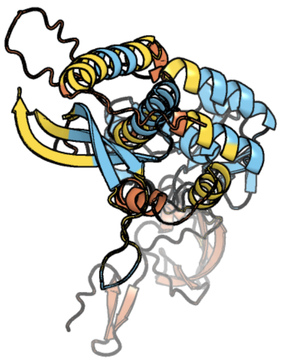

Figure 344: predicted structure of chlorv-1..344

#### chlorv-1..345

- Sequence-based annotation for chlorv-1..345 is hypothetical protein
- No significant structural hit found

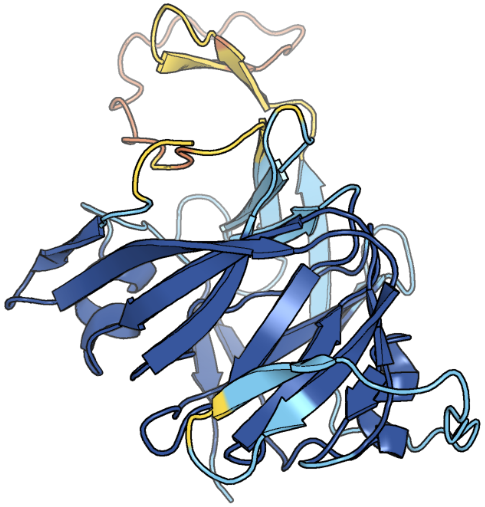

Figure 345: predicted structure of chlorv-1..345

#### chlorv-1..346

- Sequence-based annotation for chlorv-1..346 is hypothetical protein
- No significant structural hit found

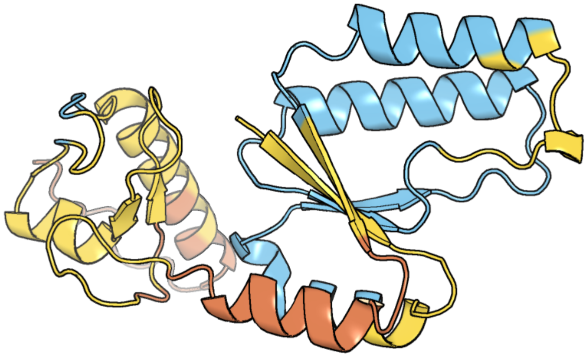

Figure 346: predicted structure of chlorv-1..346

### chlorv-1..347

- Sequence-based annotation for chlorv-1..347 is hypothetical protein
- No significant structural hit found

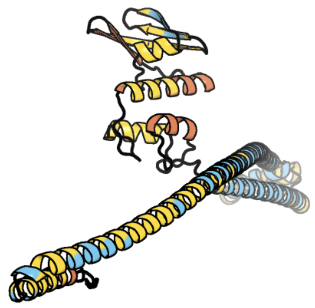

Figure 347: predicted structure of chlorv-1..347

### chlorv-1..348

- Sequence-based annotation for chlorv-1..348 is putative YqaJ-like viral recombinase
- Best hit was 5yet chain B: Uncharacterized protein R354

| target | prob | fident | alnlen | evalue | theader |
|---|---|---|---|---|---|
| 5yet-assembly1.cif.gz_B | 1 | 0.318 | 402 | 1.69e-29 | Structure of R354_WT |
| 5yeu-assembly1.cif.gz_A | 1 | 0.309 | 401 | 7.06e-28 | Structural and mechanistic analyses reveal a unique Cas4-like protein in the mimivirus virophage resistance element system |
| 5yeu-assembly1.cif.gz_B | 1 | 0.312 | 403 | 9.518e-27 | Structural and mechanistic analyses reveal a unique Cas4-like protein in the mimivirus virophage resistance element system |

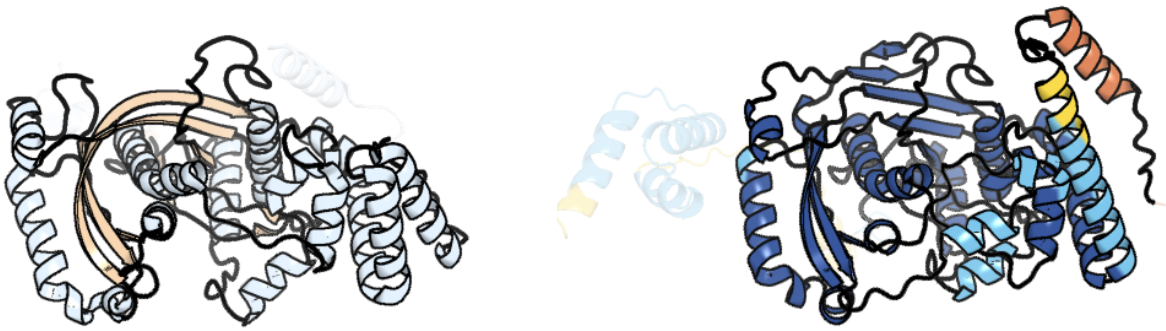

Figure 348: left: reference structure of 5yet chain B. right: predicted structure of chlorv-1..348, unaligned sequences are shown as transparent

#### chlorv-1..349

- Sequence-based annotation for chlorv-1..349 is hypothetical protein
- No significant structural hit found

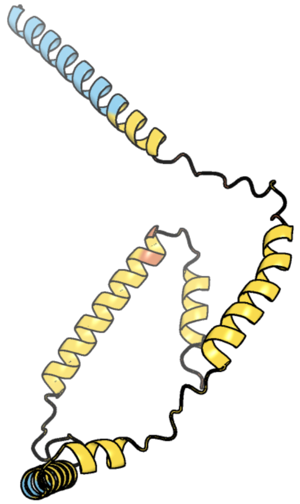

Figure 349: predicted structure of chlorv-1..349

#### chlorv-1..350

- Sequence-based annotation for chlorv-1..350 is putative DNA ligase/mRNA capping enzyme
- Best hit was 1ckm chain A: MRNA CAPPING ENZYME

| target | prob | fident | alnlen | evalue | theader |
|---|---|---|---|---|---|
| 1ckm-assembly1.cif.gz_A | 1 | 0.128 | 350 | 1.34e-12 | STRUCTURE OF TWO DIFFERENT CONFORMATIONS OF MRNA CAPPING ENZYME IN COMPLEX WITH GTP |
| 4pz8-assembly1.cif.gz_A | 1 | 0.144 | 361 | 1.092e-11 | PCE1 guanylyltransferase bound to SPT5 CTD |
| 3kyh-assembly1.cif.gz_D | 1 | 0.117 | 393 | 9.41e-11 | Saccharomyces cerevisiae Cet1-Ceg1 capping apparatus |

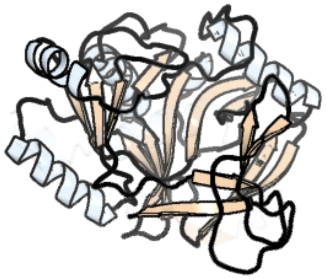

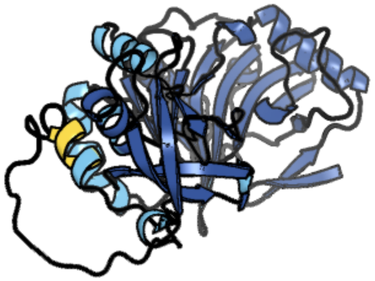

Figure 350: left: reference structure of 1ckm chain A. right: predicted structure of chlorv-1..350, unaligned sequences are shown as transparent

#### chlorv-1..351

- Sequence-based annotation for chlorv-1..351 is putative Aspartyl/Asparaginyl beta-hydroxylase
- Best hit was 5apa chain A: ASPARTYL/ASPARAGINYL BETA-HYDROXYLASE

| target | prob | fident | alnlen | evalue | theader |
|---|---|---|---|---|---|
| 5apa-assembly1.cif.gz_A | 1 | 0.211 | 175 | 5.779e-12 | Crystal structure of human aspartate beta-hydroxylase isoform a |
| 6q9i-assembly1.cif.gz_A | 1 | 0.194 | 180 | 1.734e-11 | Aspartyl/Asparaginyl beta-hydroxylase (AspH) H679A in complex with Factor X peptide fragment (39mer-4Ser) |
| 7bmj-assembly1.cif.gz_A | 1 | 0.202 | 178 | 4.605e-11 | Aspartyl/Asparaginyl beta-hydroxylase (AspH) oxygenase and TPR domains in complex with manganese, 5-fluoropyridine-2,4-dicarboxylic acid, and factor X substrate peptide fragment (39mer-4Ser) |

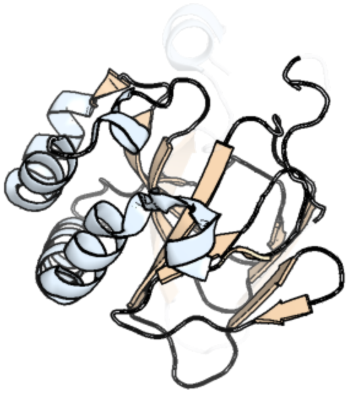

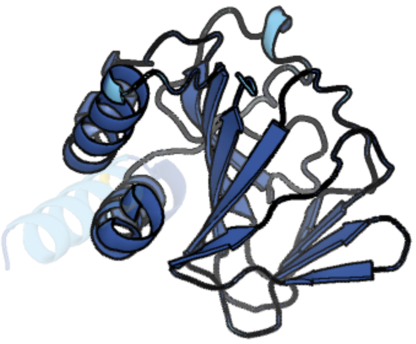

Figure 351: left: reference structure of 5apa chain A. right: predicted structure of chlorv-1..351, unaligned sequences are shown as transparent

#### chlorv-1..352

- Sequence-based annotation for chlorv-1..352 is putative Aspartyl/Asparaginyl beta-hydroxylase
- Best hit was 5apa chain A: ASPARTYL/ASPARAGINYL BETA-HYDROXYLASE

| target | prob | fident | alnlen | evalue | theader |
|---|---|---|---|---|---|
| 5apa-assembly1.cif.gz_A | 1 | 0.225 | 213 | 1.386e-13 | Crystal structure of human aspartate beta-hydroxylase isoform a |
| 6q9i-assembly1.cif.gz_A | 1 | 0.202 | 212 | 1.092e-12 | Aspartyl/Asparaginyl beta-hydroxylase (AspH) H679A in complex with Factor X peptide fragment (39mer-4Ser) |
| 7bmj-assembly1.cif.gz_A | 1 | 0.216 | 212 | 1.092e-12 | Aspartyl/Asparaginyl beta-hydroxylase (AspH) oxygenase and TPR domains in complex with manganese, 5-fluoropyridine-2,4-dicarboxylic acid, and factor X substrate peptide fragment (39mer-4Ser) |

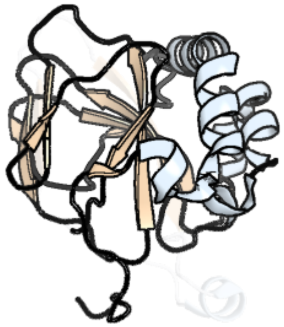
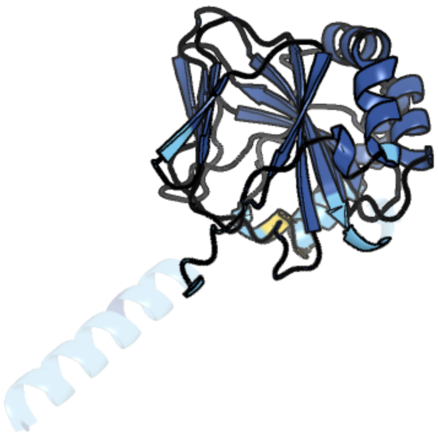

Figure 352: left: reference structure of 5apa chain A. right: predicted structure of chlorv-1..352, unaligned sequences are shown as transparent

### chlorv-1..353

- Sequence-based annotation for chlorv-1..353 is putative Protein phosphatase 2C
- Best hit was 4n0g chain B: Protein phosphatase 2C 37

| target | prob | fident | alnlen | evalue | theader |
|---|---|---|---|---|---|
| 4n0g-assembly2.cif.gz_B | 1 | 0.288 | 274 | 2.895e-24 | Crystal Structure of PYL13-PP2CA complex |
| 3nmv-assembly1.cif.gz_B | 1 | 0.258 | 282 | 6.575e-24 | Crystal structure of pyrabactin-bound abscisic acid receptor PYL2 mutant A93F in complex with type 2C protein phosphatase ABI2 |
| 5gwp-assembly1.cif.gz_A | 1 | 0.232 | 310 | 1.493e-23 | Crystal structure of RCAR3:PP2C wild-type with (+)-ABA |

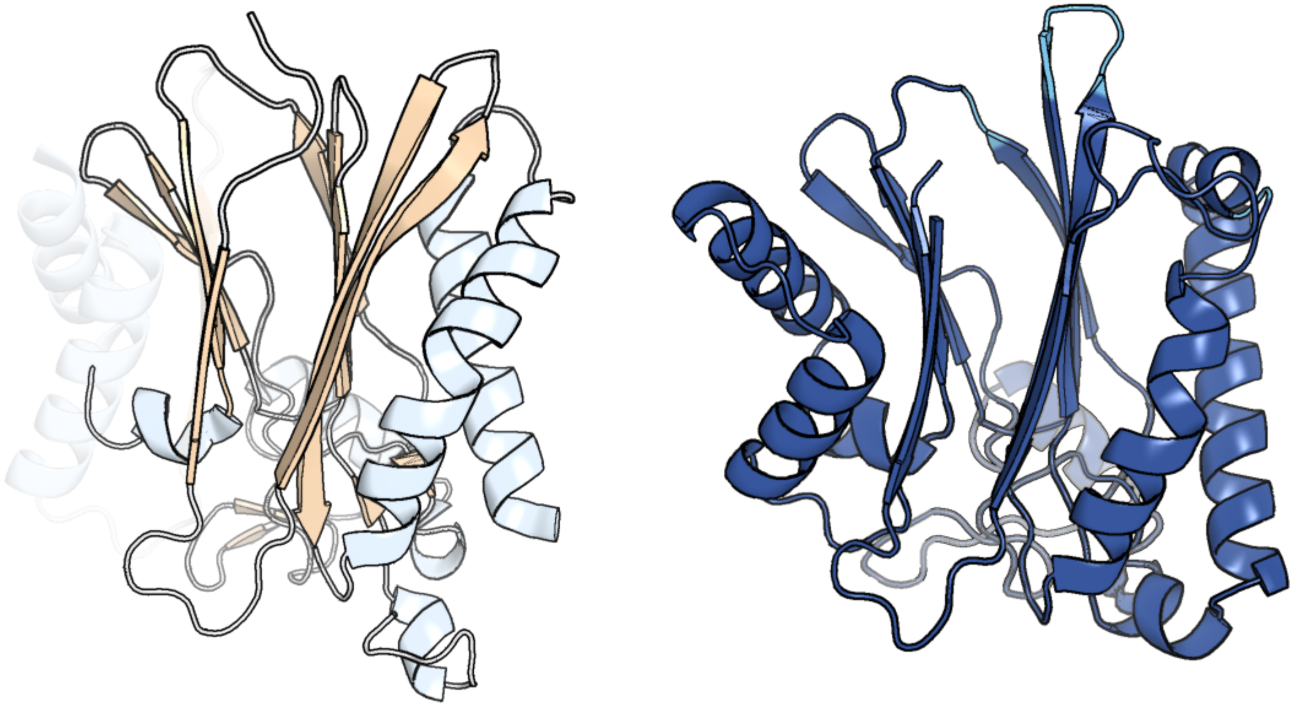

Figure 353: left: reference structure of 4n0g chain B. right: predicted structure of chlorv-1..353, unaligned sequences are shown as transparent

#### chlorv-1..354

- Sequence-based annotation for chlorv-1..354 is putative DNA mismatch repair protein MutS
- Best hit was 7ai6 chain B: DNA mismatch repair protein MutS

| target | prob | fident | alnlen | evalue | theader |
|---|---|---|---|---|---|
| 7ai6-assembly1.cif.gz_B | 1 | 0.191 | 877 | 1.021e-43 | MutS in mismatch bound state |
| 3k0s-assembly1.cif.gz_A | 1 | 0.191 | 867 | 1.124e-43 | Crystal structure of E.coli DNA mismatch repair protein MutS, D693N mutant, in complex with GT mismatched DNA |
| 1ng9-assembly1.cif.gz_A | 1 | 0.193 | 865 | 1.902e-43 | E.coli MutS R697A: an ATPase-asymmetry mutant |

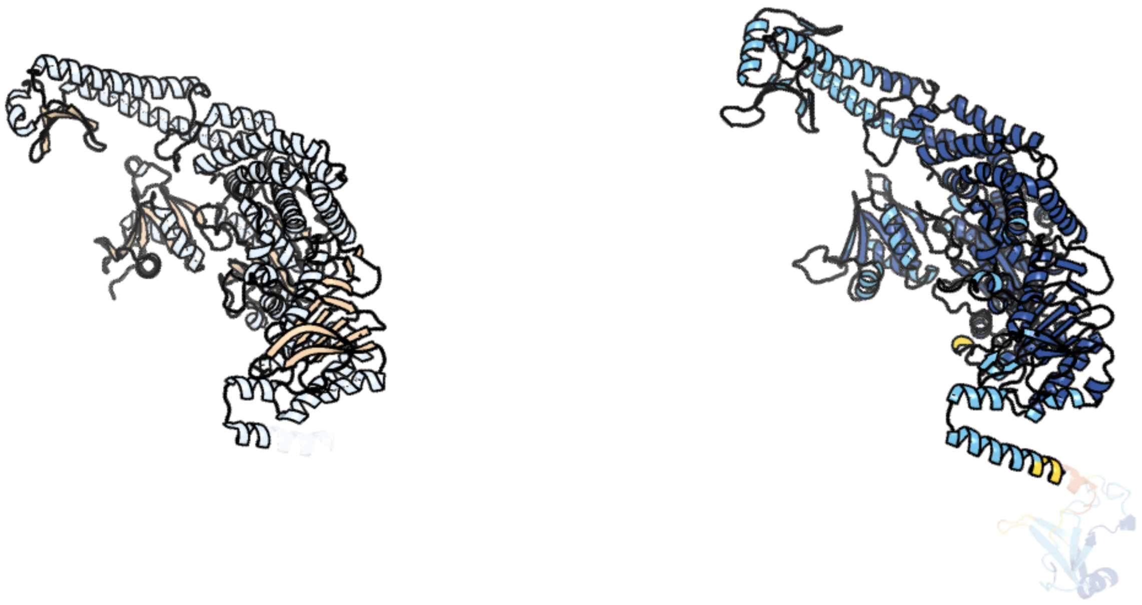

Figure 354: left: reference structure of 7ai6 chain B. right: predicted structure of chlorv-1..354, unaligned sequences are shown as transparent

#### chlorv-1..355

- Sequence-based annotation for chlorv-1..355 is hypothetical protein
- No significant structural hit found

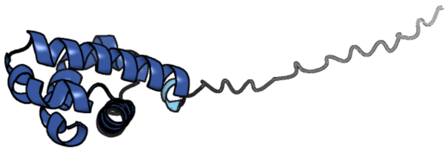

Figure 355: predicted structure of chlorv-1..355

#### chlorv-1..356

- Sequence-based annotation for chlorv-1..356 is hypothetical protein
- No significant structural hit found

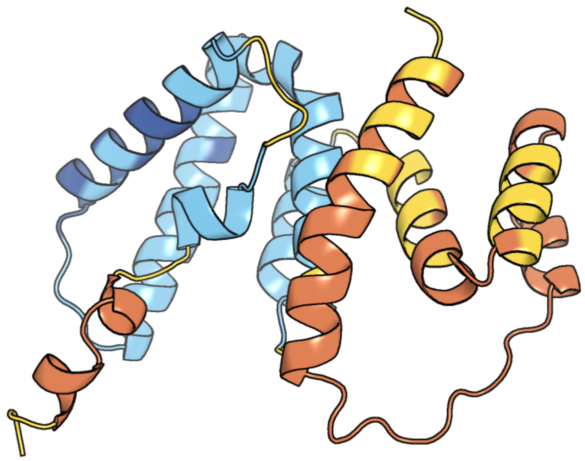

Figure 356: predicted structure of chlorv-1..356

#### chlorv-1..357

- Sequence-based annotation for chlorv-1..357 is putative Protein kinase
- Best hit was 5mxx chain A: SRPK1

| target | prob | fident | alnlen | evalue | theader |
|---|---|---|---|---|---|
| 5mxx-assembly1.cif.gz_A | 1 | 0.288 | 367 | 1.209e-23 | Crystal structure of human SR protein kinase 1 (SRPK1) in complex with compound 1 |
| 7zks-assembly1.cif.gz_A | 1 | 0.279 | 368 | 2.937e-23 | SRPK1 IN COMPLEX WITH INHIBITOR |
| 7pqs-assembly2.cif.gz_B | 1 | 0.287 | 369 | 3.435e-23 | SRPK1 in complex with MSC2711186 |

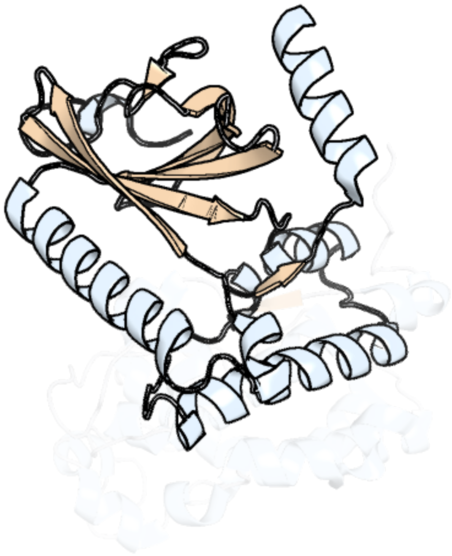

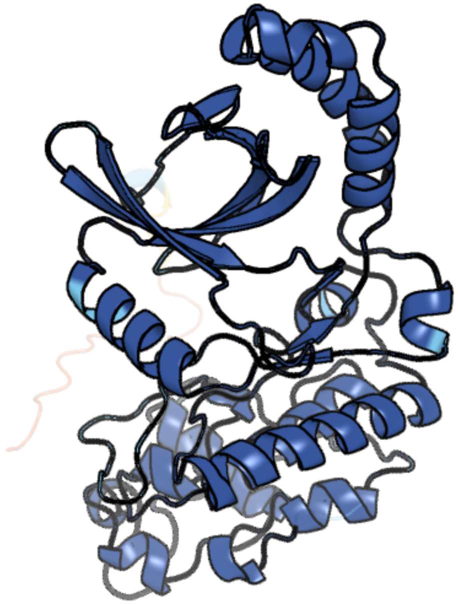

Figure 357: left: reference structure of 5mxx chain A. right: predicted structure of chlorv-1..357, unaligned sequences are shown as transparent

#### chlorv-1..358

- Sequence-based annotation for chlorv-1..358 is hypothetical protein
- No significant structural hit found

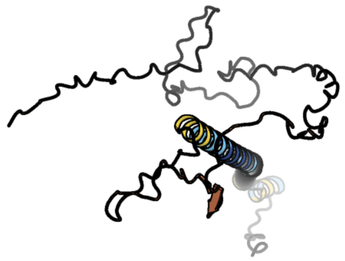

Figure 358: predicted structure of chlorv-1..358

#### chlorv-1..359

- Sequence-based annotation for chlorv-1..359 is putative RNA polymerase
- Best hit was 4a3c chain F: DNA-DIRECTED RNA POLYMERASES I, II, AND III SUBUNIT RPABC 2

| target | prob | fident | alnlen | evalue | theader |
|---|---|---|---|---|---|
| 4a3c-assembly1.cif.gz_F | 1 | 0.392 | 84 | 5.185e-08 | RNA Polymerase II initial transcribing complex with a 5nt DNA-RNA hybrid |
| 7zsb-assembly1.cif.gz_F | 1 | 0.308 | 120 | 1.806e-07 | Yeast RNA polymerase II transcription pre-initiation complex with the +1 nucleosome and NTP, complex C |
| 2nvx-assembly1.cif.gz_F | 1 | 0.373 | 83 | 2.059e-07 | RNA polymerase II elongation complex in 5 mM Mg+2 with 2'-dUTP |

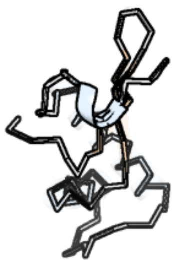

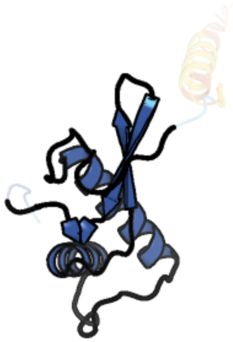

Figure 359: left: reference structure of 4a3c chain F. right: predicted structure of chlorv-1..359, unaligned sequences are shown as transparent

#### chlorv-1..360

- Sequence-based annotation for chlorv-1..360 is hypothetical protein
- No significant structural hit found

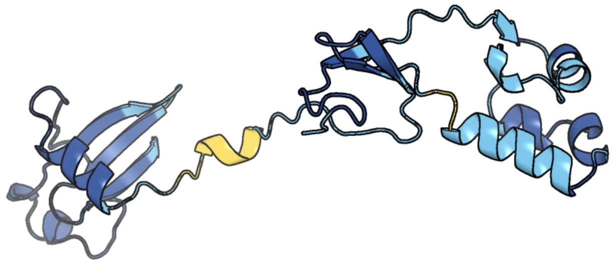

Figure 360: predicted structure of chlorv-1..360

#### chlorv-1..361

- Sequence-based annotation for chlorv-1..361 is putative D5-like helicase-primase
- Best hit was 8iqi chain B: Putative primase C962R

| target | prob | fident | alnlen | evalue | theader |
|---|---|---|---|---|---|
| 8iqi-assembly1.cif.gz_B | 1 | 0.191 | 951 | 1.364e-39 | Structure of Full-Length AsfvPrimPol in Complex-Form |
| 8iqi-assembly1.cif.gz_A | 1 | 0.193 | 948 | 2.121e-39 | Structure of Full-Length AsfvPrimPol in Complex-Form |
| 8iqi-assembly1.cif.gz_E | 1 | 0.193 | 954 | 5.811e-38 | Structure of Full-Length AsfvPrimPol in Complex-Form |

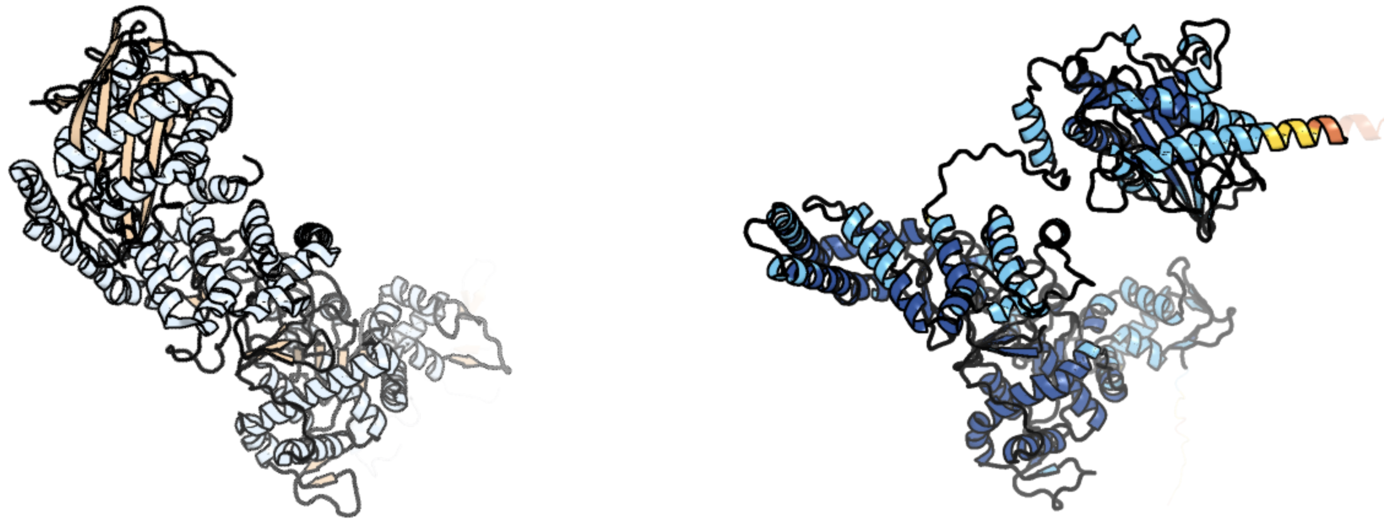

Figure 361: left: reference structure of 8iqi chain B. right: predicted structure of chlorv-1..361, unaligned sequences are shown as transparent

#### chlorv-1..362

- Sequence-based annotation for chlorv-1..362 is putative Zinc finger
- No significant structural hit found

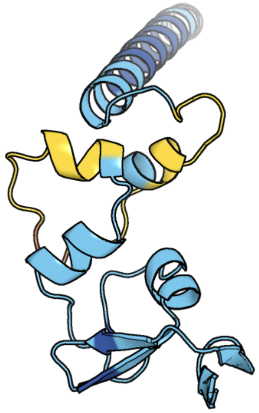

Figure 362: predicted structure of chlorv-1..362

#### chlorv-1..363

- Sequence-based annotation for chlorv-1..363 is hypothetical protein
- No significant structural hit found

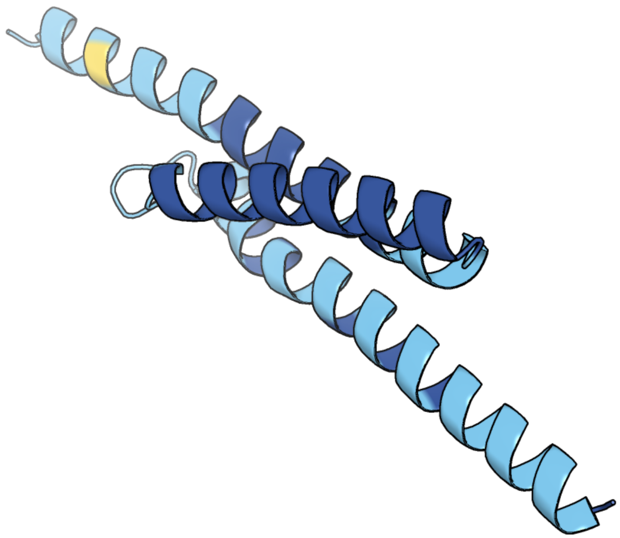

Figure 363: predicted structure of chlorv-1..363

#### chlorv-1..364

- Sequence-based annotation for chlorv-1..364 is putative HD phosphohydrolase
- Best hit was 5yhw chain C: Deoxynucleoside triphosphate triphosphohydrolase SAMHD1

| target | prob | fident | alnlen | evalue | theader |
|---|---|---|---|---|---|
| 5yhw-assembly2.cif.gz_C | 1 | 0.24 | 433 | 1.724e-21 | Crystal structure of Pig SAMHD1 |
| 7ltt-assembly1.cif.gz_D | 1 | 0.226 | 442 | 6.897e-21 | SAMHD1(113-626) H206R D207N R366C |
| 7a5y-assembly2.cif.gz_H | 1 | 0.236 | 436 | 1.06e-20 | Crystal structure of tetrameric human H215A-SAMHD1 (residues 109-626) with Rp-dGTP-alphaS (T8T) and Mg |

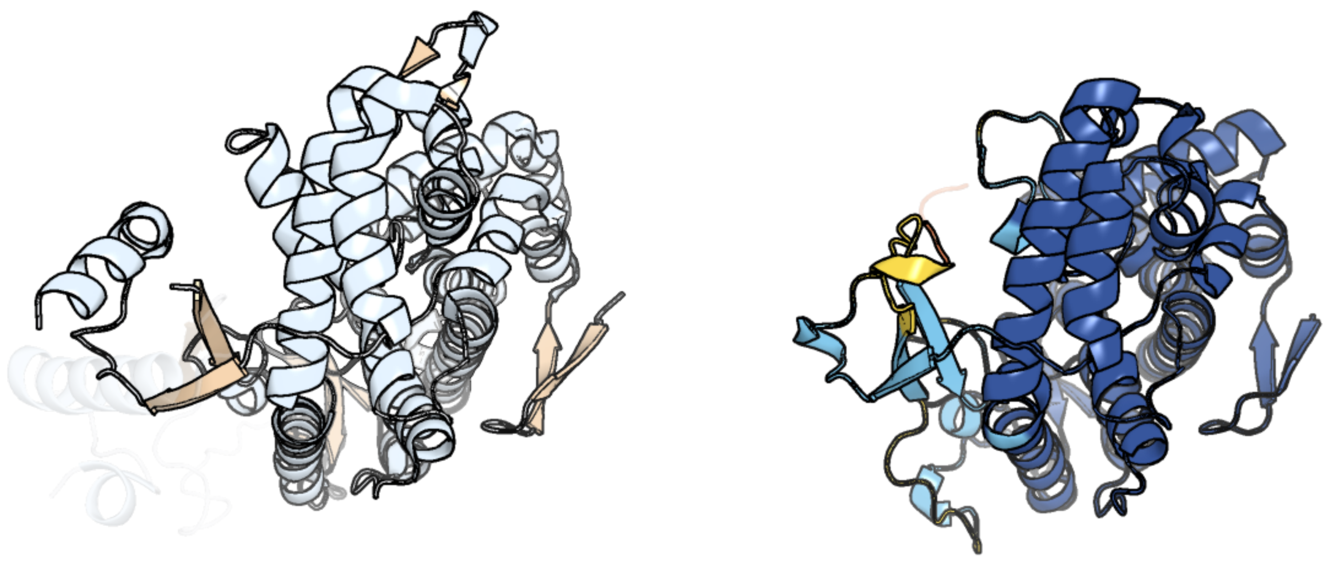

Figure 364: left: reference structure of 5yhw chain C. right: predicted structure of chlorv-1..364, unaligned sequences are shown as transparent

#### chlorv-1..365

- Sequence-based annotation for chlorv-1..365 is putative Nuclease
- Best hit was 2ihn chain A: Ribonuclease H

| target | prob | fident | alnlen | evalue | theader |
|---|---|---|---|---|---|
| 2ihn-assembly1.cif.gz_A | 1 | 0.212 | 287 | 1.612e-11 | Co-crystal of Bacteriophage T4 RNase H with a fork DNA substrate |
| 3h8s-assembly1.cif.gz_A | 1 | 0.217 | 289 | 8.087e-11 | Structure of D19N T4 RNase H in the presence of divalent magnesium |
| 3h7i-assembly1.cif.gz_A | 1 | 0.225 | 297 | 1.541e-10 | Structure of the metal-free D132N T4 RNase H |

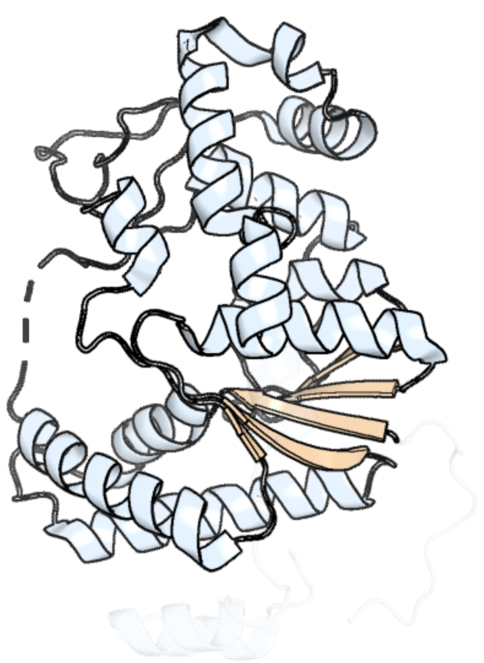

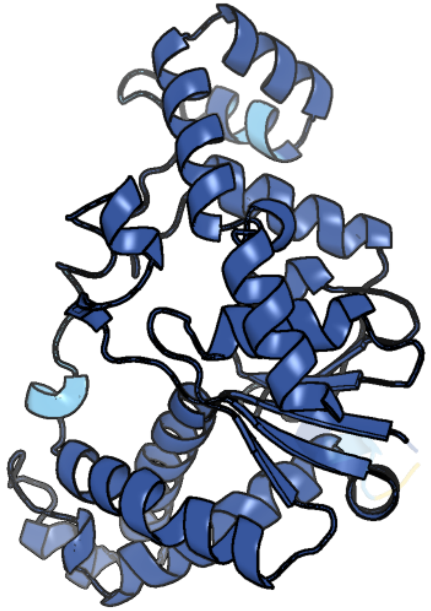

Figure 365: left: reference structure of 2ihn chain A. right: predicted structure of chlorv-1..365, unaligned sequences are shown as transparent

#### chlorv-1..366

- Sequence-based annotation for chlorv-1..366 is putative Peptidyl-tRNA hydrolase
- Best hit was 1xty chain D: Peptidyl-tRNA hydrolase

| target | prob | fident | alnlen | evalue | theader |
|---|---|---|---|---|---|
| 1xty-assembly2.cif.gz_D | 1 | 0.403 | 119 | 4.227e-12 | Crystal structure of Sulfolobus solfataricus peptidyl-tRNA hydrolase |
| 2zv3-assembly4.cif.gz_G | 1 | 0.377 | 114 | 6.339e-12 | Crystal structure of project MJ0051 from Methanocaldococcus jannaschii DSM 2661 |
| 2zv3-assembly2.cif.gz_C | 1 | 0.37 | 116 | 1.088e-11 | Crystal structure of project MJ0051 from Methanocaldococcus jannaschii DSM 2661 |

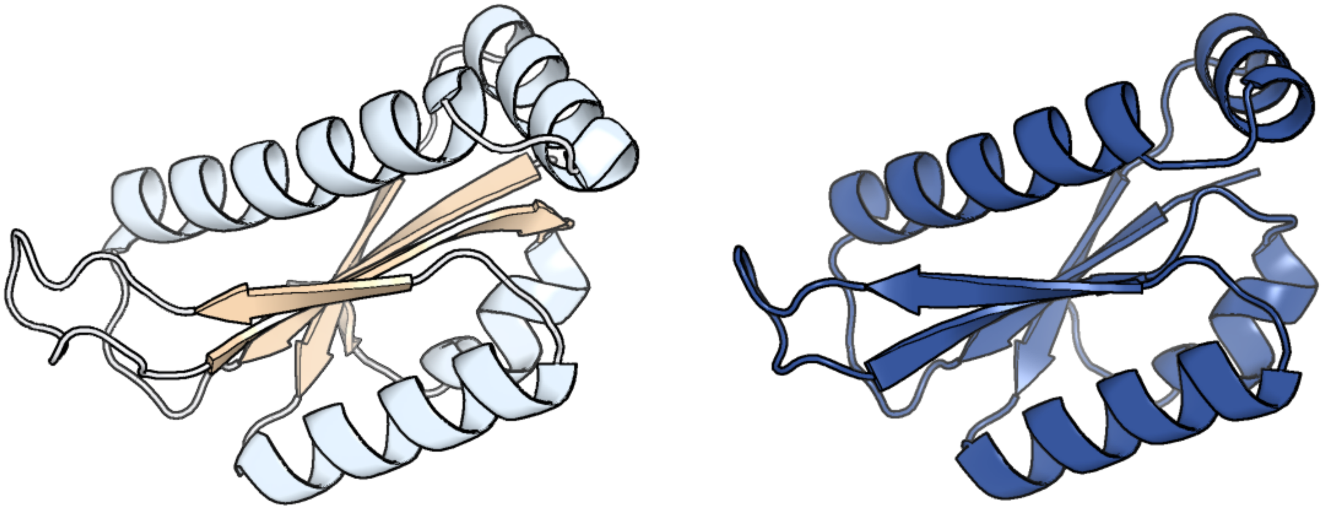

Figure 366: left: reference structure of 1xty chain D. right: predicted structure of chlorv-1..366, unaligned sequences are shown as transparent

#### chlorv-1..367

- Sequence-based annotation for chlorv-1..367 is hypothetical protein
- Best hit was 5uy6 chain A: Calcium/calmodulin-dependent protein kinase kinase 2

| target | prob | fident | alnlen | evalue | theader |
|---|---|---|---|---|---|
| 5uy6-assembly1.cif.gz_A | 1 | 0.122 | 302 | 1.988e-06 | Crystal Structure of the Human CAMKK2B |
| 8sam-assembly1.cif.gz_A | 1 | 0.114 | 384 | 7.441e-06 | Crystal structure of class III lanthipeptide synthetase LP-GS-ThurKC in complex with ATP |
| 8tuc-assembly1.cif.gz_A | 1 | 0.122 | 301 | 3.109e-05 | Unphosphorylated CaMKK2 in complex with CC-8977 |

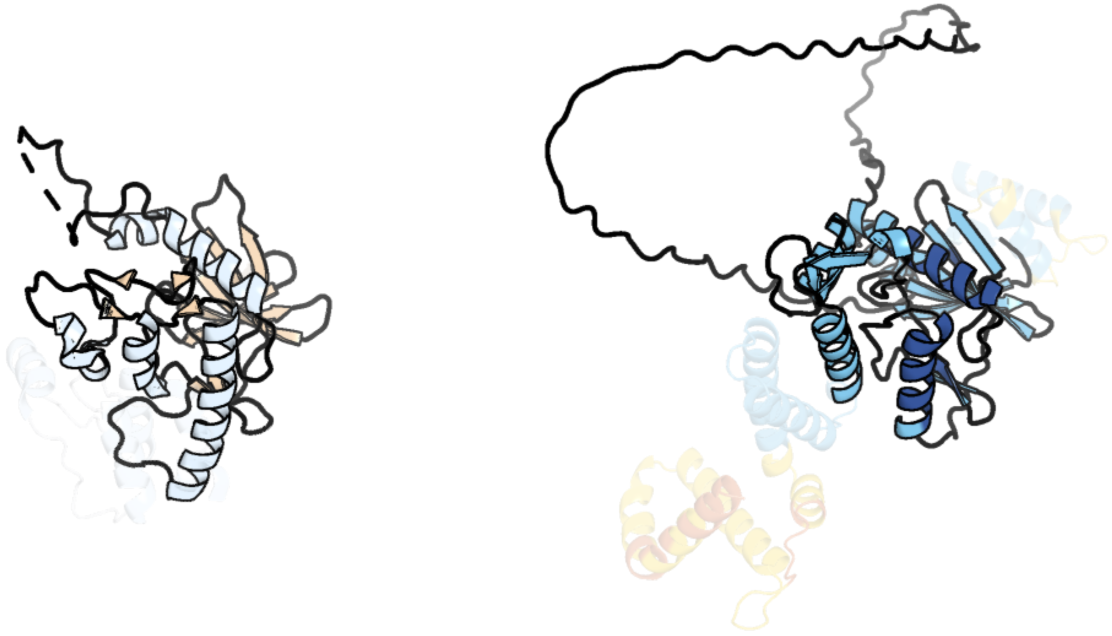

Figure 367: left: reference structure of 5uy6 chain A. right: predicted structure of chlorv-1..367, unaligned sequences are shown as transparent

#### chlorv-1..368

- Sequence-based annotation for chlorv-1..368 is putative MutM DNA repair protein formamidopyrimidine-DNA glycosylase H2TH domain
- Best hit was 3a46 chain B: Formamidopyrimidine-DNA glycosylase

| target | prob | fident | alnlen | evalue | theader |
|---|---|---|---|---|---|
| 3a46-assembly2.cif.gz_B | 1 | 0.332 | 304 | 2.284e-23 | Crystal structure of MvNei1/THF complex |
| 3a42-assembly1.cif.gz_A | 1 | 0.321 | 299 | 3.664e-23 | Crystal structure of MvNei1 |
| 3vk7-assembly2.cif.gz_B | 1 | 0.314 | 305 | 8.885e-23 | Crystal structure of DNA-glycosylase bound to DNA containing 5-Hydroxyuracil |

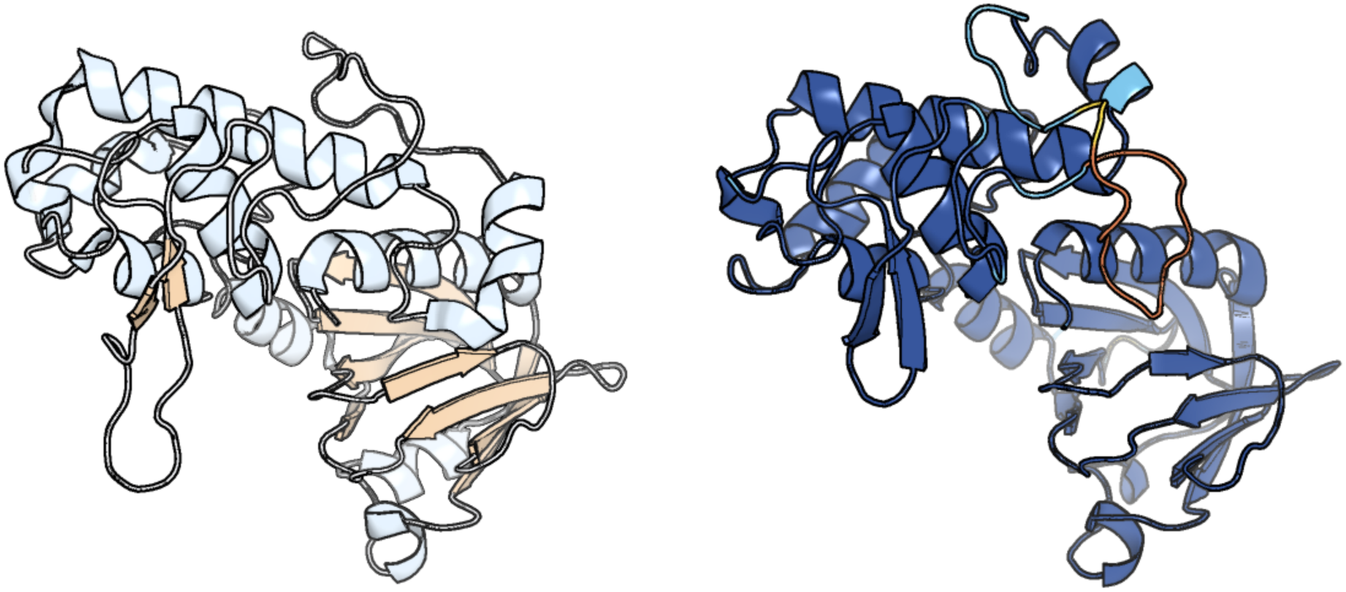

Figure 368: left: reference structure of 3a46 chain B. right: predicted structure of chlorv-1..368, unaligned sequences are shown as transparent

#### chlorv-1..369

- Sequence-based annotation for chlorv-1..369 is hypothetical protein
- No significant structural hit found

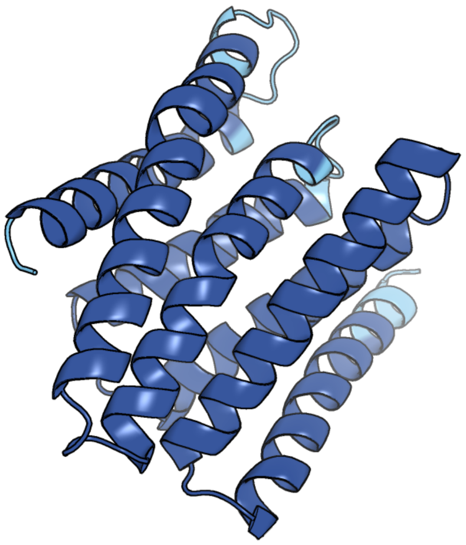

Figure 369: predicted structure of chlorv-1..369

#### chlorv-1..370

- Sequence-based annotation for chlorv-1..370 is putative Ribonucleotide reductase large chain
- Best hit was 3hnc chain B: Ribonucleoside-diphosphate reductase large subunit

| target | prob | fident | alnlen | evalue | theader |
|---|---|---|---|---|---|
| 3hnc-assembly1.cif.gz_B | 1 | 0.455 | 860 | 8.587e-81 | Crystal structure of human ribonucleotide reductase 1 bound to the effector TTP |
| 3hnf-assembly1.cif.gz_B | 1 | 0.453 | 860 | 1.151e-80 | Crystal structure of human ribonucleotide reductase 1 bound to the effectors TTP and dATP |
| 5tus-assembly3.cif.gz_B | 1 | 0.452 | 860 | 2.164e-79 | Potent competitive inhibition of human ribonucleotide reductase by a novel non-nucleoside small molecule |

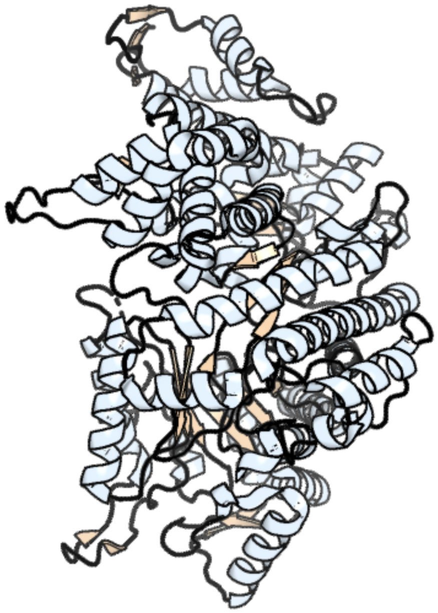

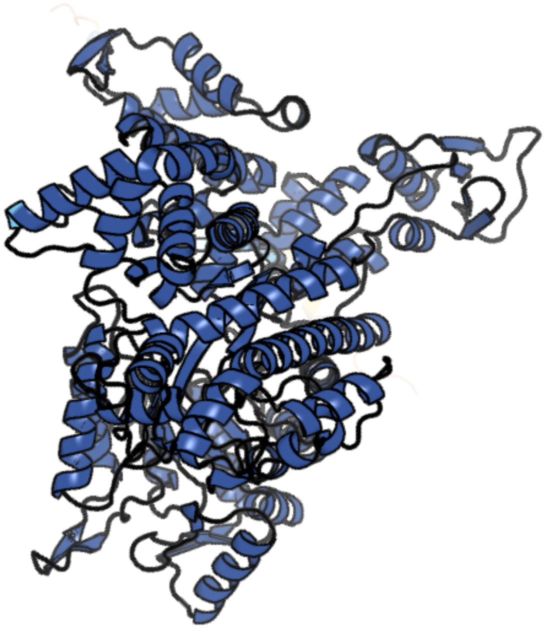

Figure 370: left: reference structure of 3hnc chain B. right: predicted structure of chlorv-1..370, unaligned sequences are shown as transparent

#### chlorv-1..371

- Sequence-based annotation for chlorv-1..371 is hypothetical protein
- No significant structural hit found

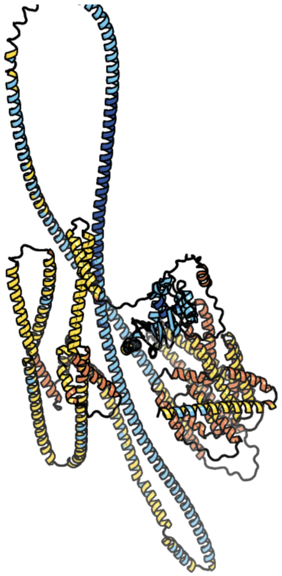

Figure 371: predicted structure of chlorv-1..371

#### chlorv-1..372

- Sequence-based annotation for chlorv-1..372 is putative DNA topoisomerase I
- Best hit was 4rul chain A: DNA topoisomerase 1

| target | prob | fident | alnlen | evalue | theader |
|---|---|---|---|---|---|
| 4rul-assembly1.cif.gz_A | 1 | 0.312 | 793 | 1.098e-55 | Crystal structure of full-length E.Coli topoisomerase I in complex with ssDNA |
| 6pcm-assembly1.cif.gz_A | 1 | 0.304 | 846 | 2.252e-53 | Crystal Structure of Mycobacterium smegmatis Topoisomerase I with ssDNA bound to both N- and C-terminal domains |
| 6ozw-assembly1.cif.gz_A | 1 | 0.368 | 605 | 4.138e-53 | Crystal structure of the 65-kilodalton amino-terminal fragment of DNA topoisomerase I from Streptococcus mutans |

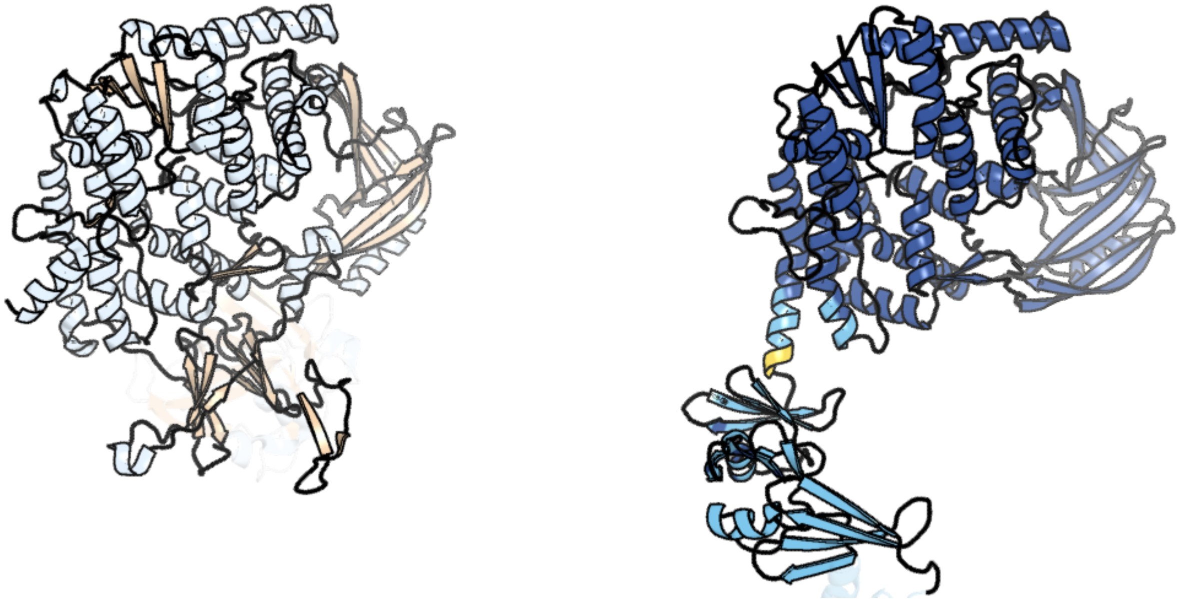

Figure 372: left: reference structure of 4rul chain A. right: predicted structure of chlorv-1..372, unaligned sequences are shown as transparent

### chlorv-1..373

- Sequence-based annotation for chlorv-1..373 is hypothetical protein
- No significant structural hit found

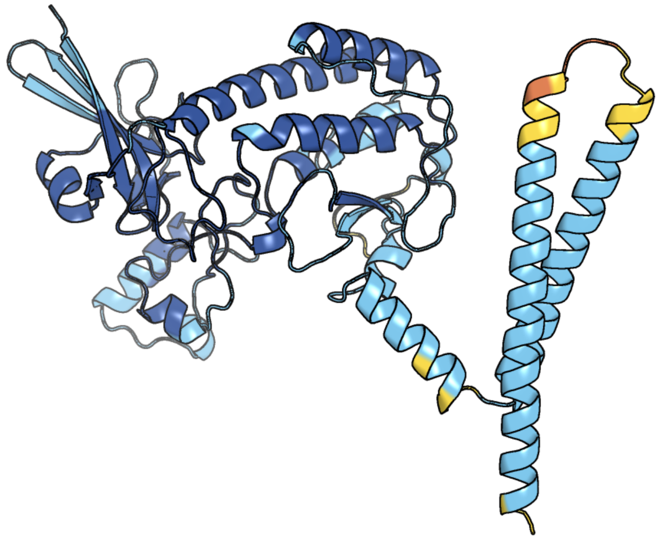

Figure 373: predicted structure of chlorv-1..373

#### chlorv-1..374

- Sequence-based annotation for chlorv-1..374 is putative Methyltransferase
- Best hit was 7eew chain A: Type I restriction-modification system methyltransferase subunit

| target | prob | fident | alnlen | evalue | theader |
|---|---|---|---|---|---|
| 7eew-assembly1.cif.gz_A-2 | 1 | 0.165 | 574 | 1.767e-18 | Crystal structure of the intact MTase from Vibrio vulnificus YJ016 in complex with the DNA-mimicking Ocr protein and the S-adenosyl-L-homocysteine (SAH) |
| 8cy2-assembly1.cif.gz_A | 1 | 0.167 | 579 | 2.574e-15 | CamA Adenine Methyltransferase Complexed to Cognate Substrate DNA and Inhibitor APNEA (Compound 9) |
| 8cxt-assembly1.cif.gz_A | 1 | 0.162 | 577 | 3.406e-15 | CamA Adenine Methyltransferase Complexed to Cognate Substrate DNA and Inhibitor N6-benzyladenosine (Compound 1) |

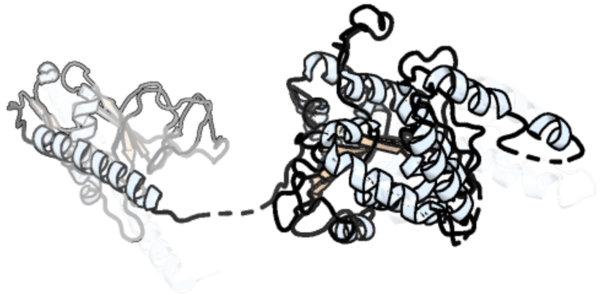

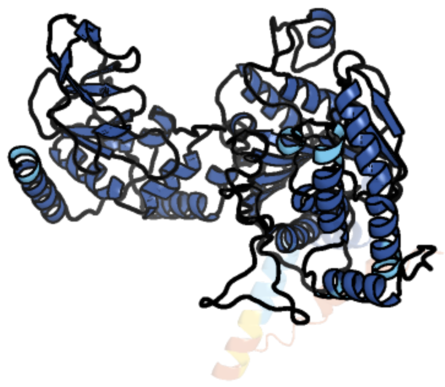

Figure 374: left: reference structure of 7eew chain A. right: predicted structure of chlorv-1..374, unaligned sequences are shown as transparent

#### chlorv-1..375

- Sequence-based annotation for chlorv-1..375 is hypothetical protein
- No significant structural hit found

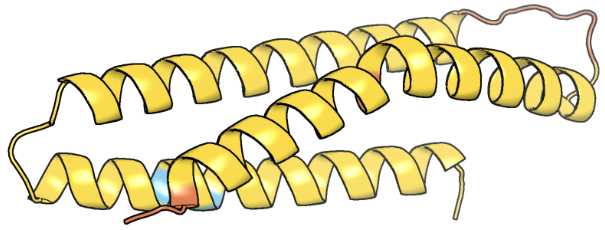

Figure 375: predicted structure of chlorv-1..375

#### chlorv-1..376

- Sequence-based annotation for chlorv-1..376 is hypothetical protein
- No significant structural hit found

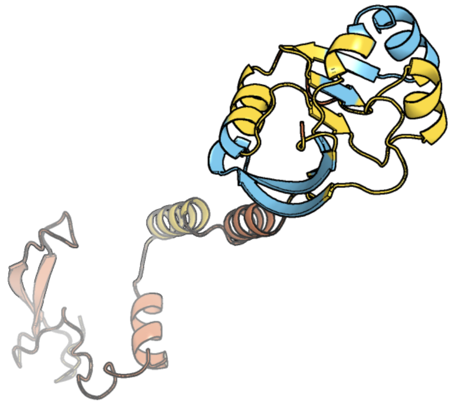

Figure 376: predicted structure of chlorv-1..376

#### chlorv-1..377

- Sequence-based annotation for chlorv-1..377 is hypothetical protein
- No significant structural hit found

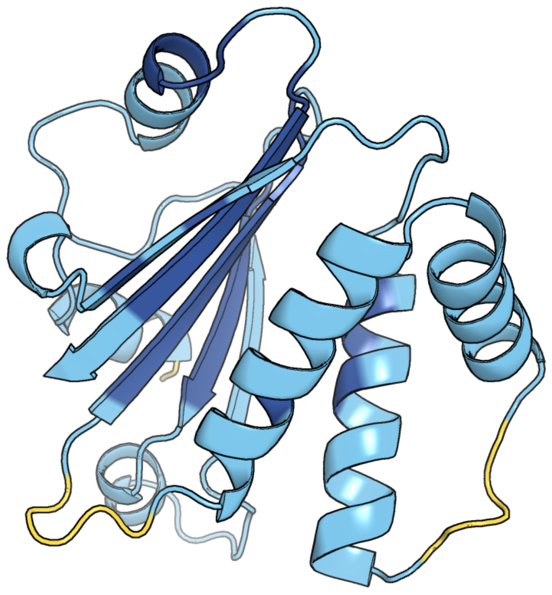

Figure 377: predicted structure of chlorv-1..377

#### chlorv-1..378

- Sequence-based annotation for chlorv-1..378 is putative Peptidase
- Best hit was 3mt6 chain T: ATP-dependent Clp protease proteolytic subunit

| target | prob | fident | alnlen | evalue | theader |
|---|---|---|---|---|---|
| 3mt6-assembly2.cif.gz_K | 1 | 0.206 | 165 | 2.459e-15 | Structure of ClpP from Escherichia coli in complex with ADEP1 |
| 3mt6-assembly1.cif.gz_T | 1 | 0.206 | 165 | 2.961e-15 | Structure of ClpP from Escherichia coli in complex with ADEP1 |
| 4emp-assembly1.cif.gz_F | 1 | 0.242 | 165 | 3.795e-15 | Crystal structure of the mutant of ClpP E137A from Staphylococcus aureus |

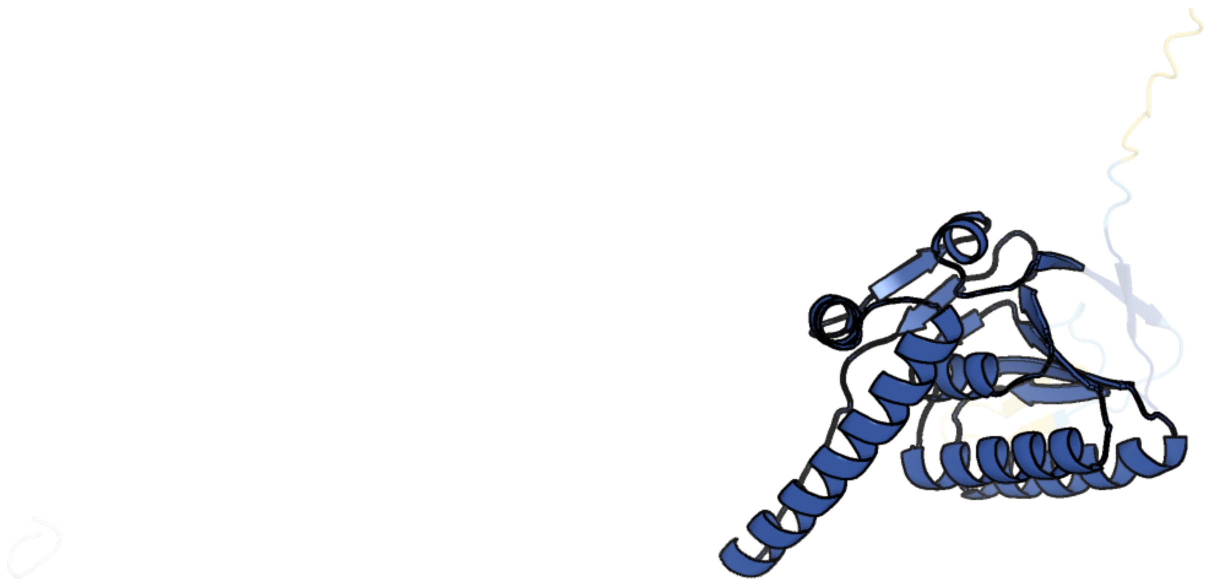

Figure 378: left: reference structure of 3mt6 chain T. right: predicted structure of chlorv-1..378, unaligned sequences are shown as transparent

#### chlorv-1..379

- Sequence-based annotation for chlorv-1..379 is putative Transcription factor/TATA-binding protein
- Best hit was 8cen chain O: TATA-binding protein

| target | prob | fident | alnlen | evalue | theader |
|---|---|---|---|---|---|
| 4b0a-assembly1.cif.gz_A | 1 | 0.17 | 258 | 2.95e-09 | The high-resolution structure of yTBP-yTAF1 identifies conserved and competing interaction surfaces in transcriptional activation |
| 1mp9-assembly1.cif.gz_B | 1 | 0.181 | 248 | 4.135e-09 | TBP from a mesothermophilic archaeon, Sulfolobus acidocaldarius |
| 8cen-assembly1.cif.gz_O | 1 | 0.158 | 246 | 5.179e-09 | Yeast RNA polymerase II transcription pre-initiation complex with core Mediator |

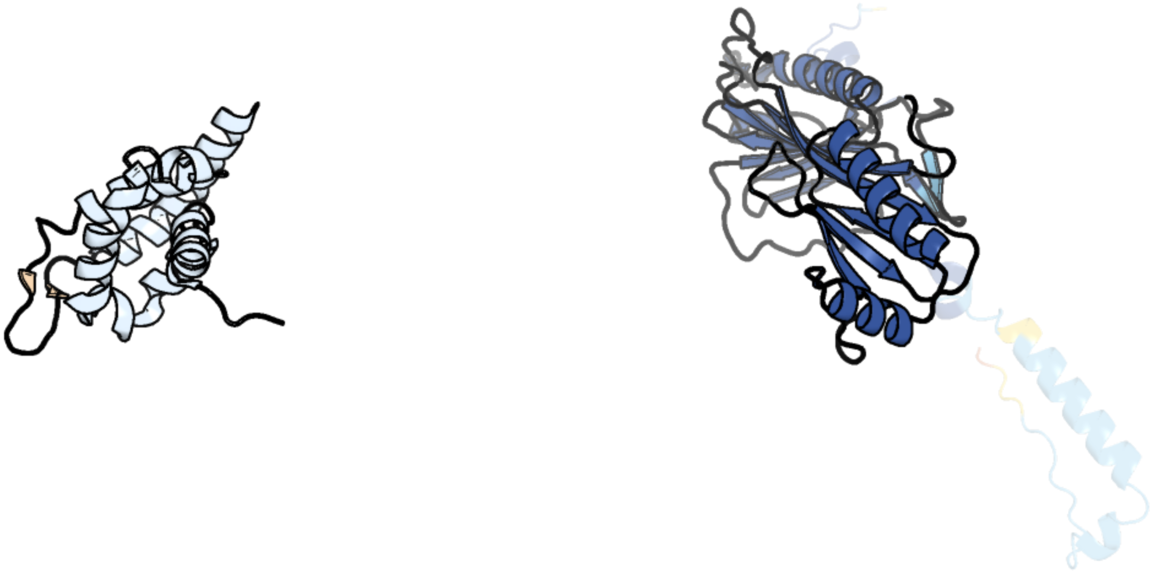

Figure 379: left: reference structure of 8cen chain O. right: predicted structure of chlorv-1..379, unaligned sequences are shown as transparent

#### chlorv-1..380

- Sequence-based annotation for chlorv-1..380 is putative Thymidine kinase
- Best hit was 4uxj chain E: THYMIDINE KINASE

| target | prob | fident | alnlen | evalue | theader |
|---|---|---|---|---|---|
| 4uxj-assembly2.cif.gz_E-3 | 1 | 0.3 | 180 | 1.51e-17 | Leishmania major Thymidine Kinase in complex with dTTP |
| 4uxi-assembly1.cif.gz_B-2 | 1 | 0.282 | 177 | 2.067e-17 | Leishmania major Thymidine Kinase in complex with thymidine |
| 2wvj-assembly1.cif.gz_D | 1 | 0.308 | 178 | 2.83e-17 | Mutation of Thr163 to Ser in Human Thymidine Kinase Shifts the Specificity from Thymidine towards the Nucleoside Analogue Azidothymidine |

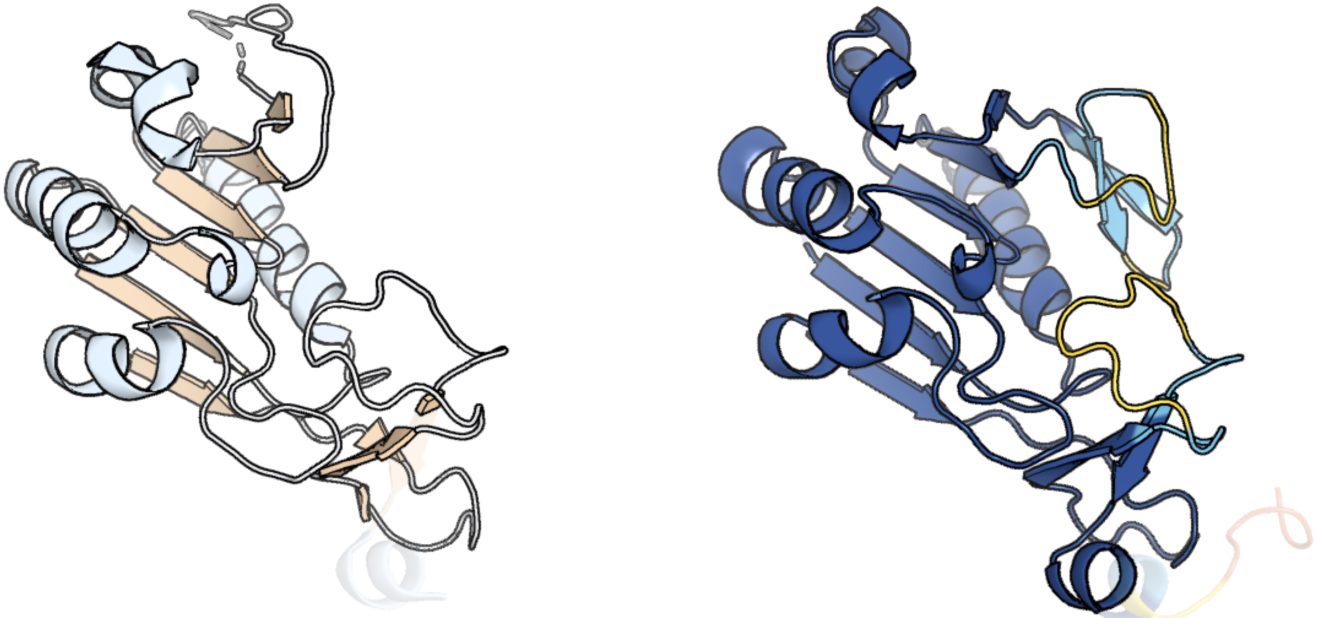

Figure 380: left: reference structure of 4uxj chain E. right: predicted structure of chlorv-1..380, unaligned sequences are shown as transparent

#### chlorv-1..381

- Sequence-based annotation for chlorv-1..381 is hypothetical protein
- Best hit was 1kcf chain B: HYPOTHETICAL 30.2 KD PROTEIN C25G10.02 IN CHROMOSOME I

| target | prob | fident | alnlen | evalue | theader |
|---|---|---|---|---|---|
| 1kcf-assembly1.cif.gz_B | 1 | 0.172 | 284 | 6.772e-06 | Crystal Structure of the Yeast Mitochondrial Holliday Junction Resolvase, Ydc2 |
| 1kcf-assembly1.cif.gz_A | 1 | 0.164 | 291 | 0.0001282 | Crystal Structure of the Yeast Mitochondrial Holliday Junction Resolvase, Ydc2 |
| 6p7a-assembly2.cif.gz_B-2 | 1 | 0.147 | 285 | 0.0002596 | CRYSTAL STRUCTURE OF THE FOWLPOX VIRUS HOLLIDAY JUNCTION RESOLVASE |

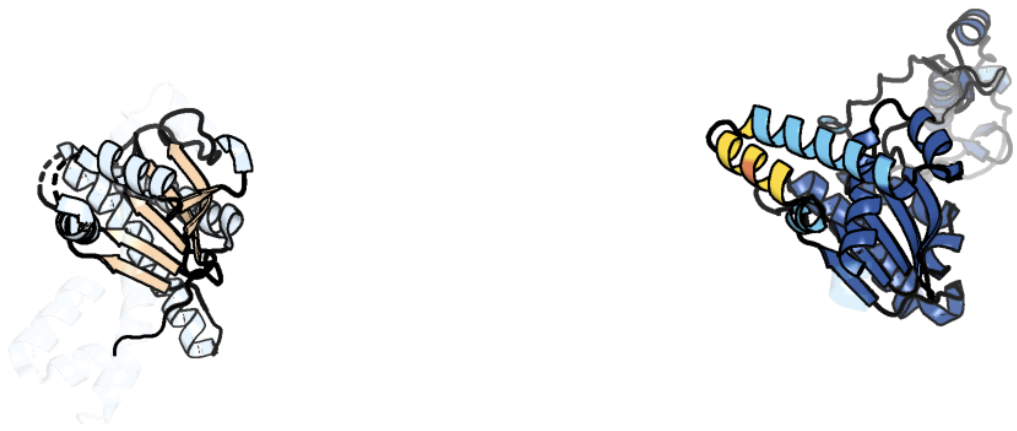

Figure 381: left: reference structure of 1kcf chain B. right: predicted structure of chlorv-1..381, unaligned sequences are shown as transparent

#### chlorv-1..382

- Sequence-based annotation for chlorv-1..382 is hypothetical protein
- No significant structural hit found

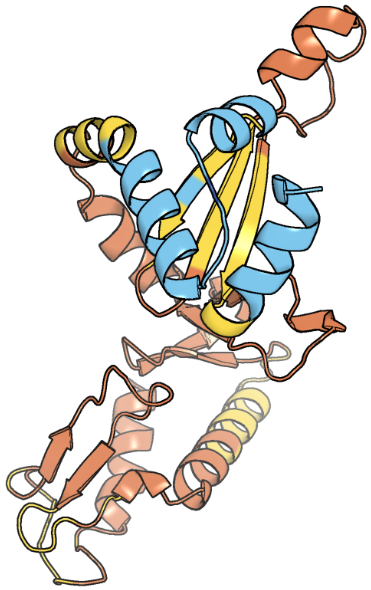

Figure 382: predicted structure of chlorv-1..382

#### chlorv-1..383

- Sequence-based annotation for chlorv-1..383 is hypothetical protein
- Best hit was 8ppl chain Ir: Eukaryotic translation initiation factor 2 subunit 1

| target | prob | fident | alnlen | evalue | theader |
|---|---|---|---|---|---|
| 8ppl-assembly1.cif.gz_Ir | 1 | 0.179 | 273 | 4.699e-13 | MERS-CoV Nsp1 bound to the human 43S pre-initiation complex |
| 6ybv-assembly1.cif.gz_r | 1 | 0.181 | 276 | 6.343e-13 | Structure of a human 48S translational initiation complex - eIF2-TC |
| 1q8k-assembly1.cif.gz_A | 1 | 0.198 | 272 | 2.614e-11 | Solution structure of alpha subunit of human eIF2 |

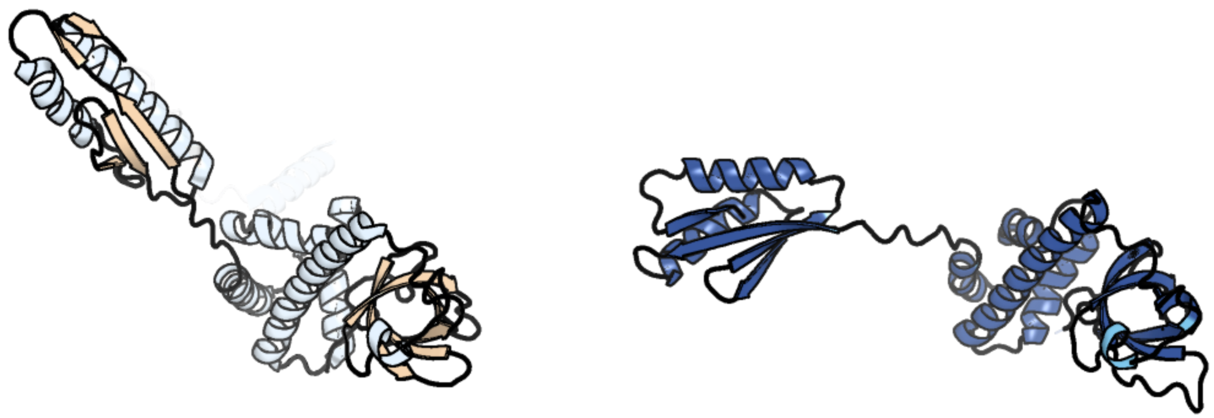

Figure 383: left: reference structure of 8ppl chain Ir. right: predicted structure of chlorv-1..383, unaligned sequences are shown as transparent

#### chlorv-1..384

- Sequence-based annotation for chlorv-1..384 is putative Poxvirus Late Transcription Factor VLTF2
- No significant structural hit found

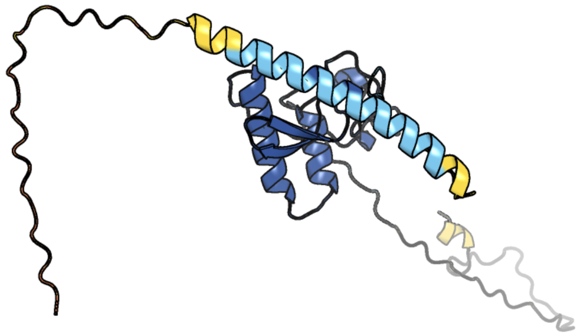

Figure 384: predicted structure of chlorv-1..384

#### chlorv-1..385

- Sequence-based annotation for chlorv-1..385 is hypothetical protein
- No significant structural hit found

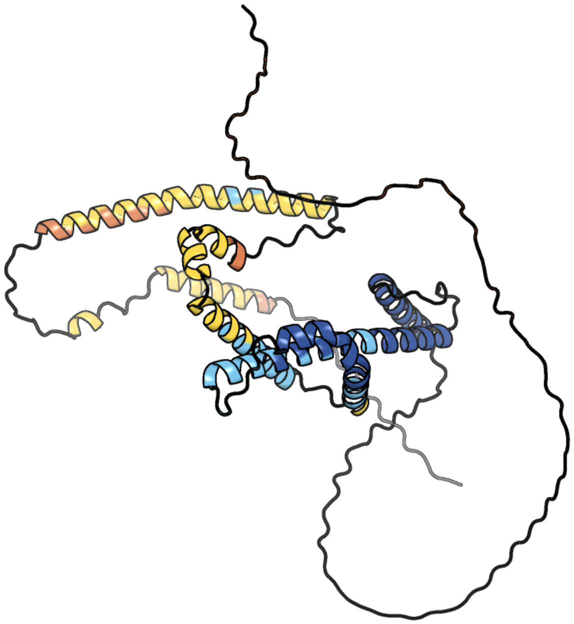

Figure 385: predicted structure of chlorv-1..385

### chlorv-1..386

- Sequence-based annotation for chlorv-1..386 is hypothetical protein
- No significant structural hit found

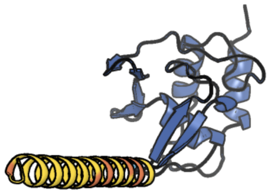

Figure 386: predicted structure of chlorv-1..386

#### chlorv-1..387

- Sequence-based annotation for chlorv-1..387 is hypothetical protein
- No significant structural hit found

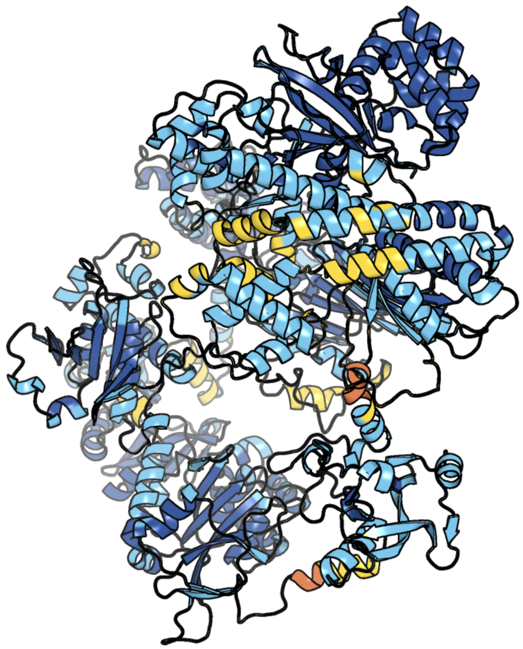

Figure 387: predicted structure of chlorv-1..387

#### chlorv-1..388

- Sequence-based annotation for chlorv-1..388 is hypothetical protein
- No significant structural hit found

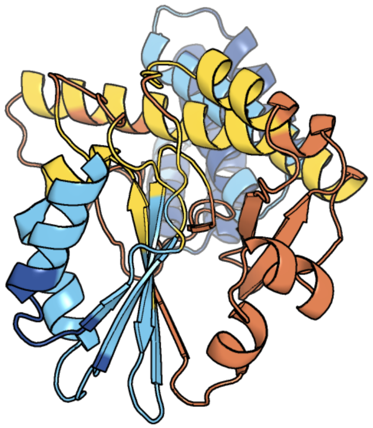

Figure 388: predicted structure of chlorv-1..388

#### chlorv-1..389

- Sequence-based annotation for chlorv-1..389 is hypothetical protein
- No significant structural hit found

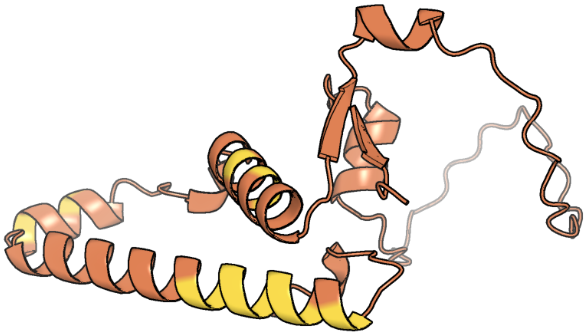

Figure 389: predicted structure of chlorv-1..389

#### chlorv-1..390

- Sequence-based annotation for chlorv-1..390 is hypothetical protein
- No significant structural hit found

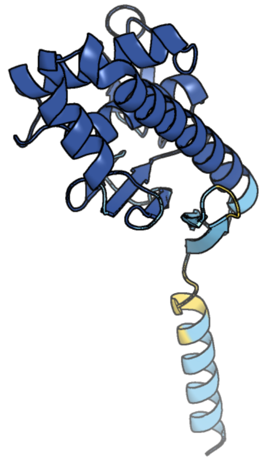

Figure 390: predicted structure of chlorv-1..390

#### chlorv-1..391

- Sequence-based annotation for chlorv-1..391 is hypothetical protein
- No significant structural hit found

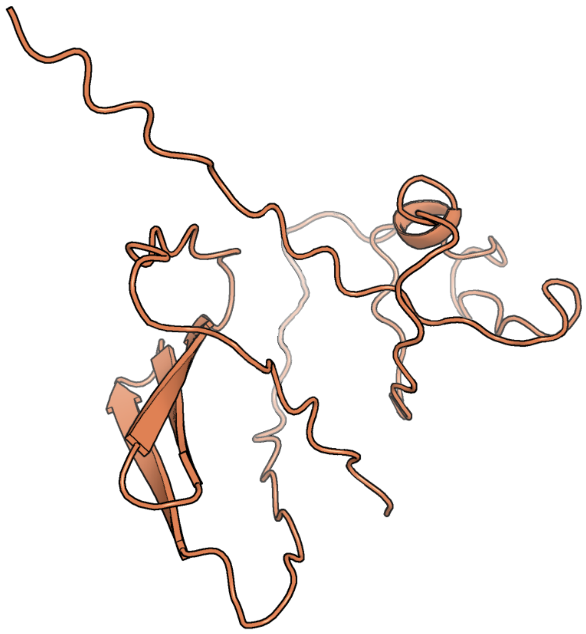

Figure 391: predicted structure of chlorv-1..391

#### chlorv-1..392

- Sequence-based annotation for chlorv-1..392 is putative Methyltransferase
- Best hit was 4n48 chain B: Cap-specific mRNA (nucleoside-2'-O-)-methyltransferase 1

| target | prob | fident | alnlen | evalue | theader |
|---|---|---|---|---|---|
| 4n48-assembly1.cif.gz__B | 1 | 0.133 | 412 | 1.485e-13 | Cap-specific mRNA (nucleoside-2'-O-)-methyltransferase 1 Protein in complex with capped RNA fragment |
| 4n48-assembly2.cif.gz__A | 1 | 0.142 | 406 | 1.839e-13 | Cap-specific mRNA (nucleoside-2'-O-)-methyltransferase 1 Protein in complex with capped RNA fragment |
| 4n49-assembly1.cif.gz__A | 1 | 0.141 | 397 | 2.047e-13 | Cap-specific mRNA (nucleoside-2'-O-)-methyltransferase 1 Protein in complex with m7GpppG and SAM |

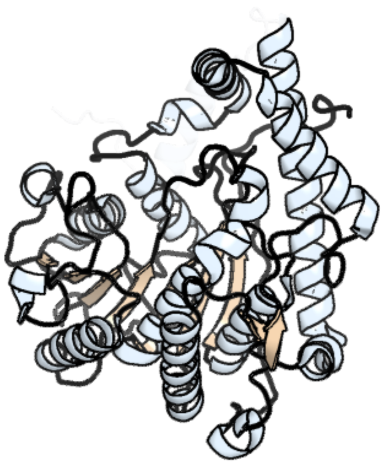

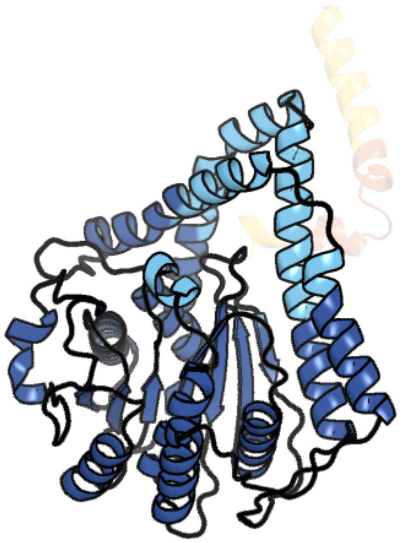

Figure 392: left: reference structure of 4n48 chain B. right: predicted structure of chlorv-1..392, unaligned sequences are shown as transparent

### chlorv-1..393

- Sequence-based annotation for chlorv-1..393 is hypothetical protein
- No significant structural hit found

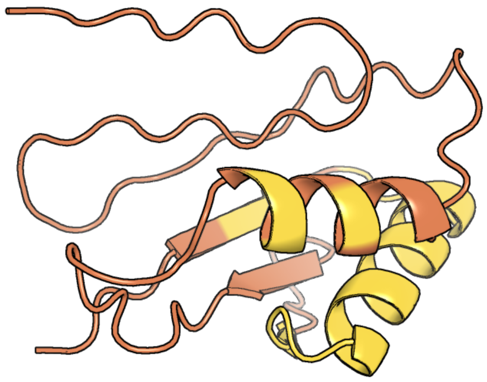

Figure 393: predicted structure of chlorv-1..393

#### chlorv-1..394

- Sequence-based annotation for chlorv-1..394 is putative Oxidoreductase/proline dehydrogenase
- Best hit was 2g37 chain A: proline dehydrogenase/delta-1-pyrroline-5-carboxylate dehydrogenase

| target | prob | fident | alnlen | evalue | theader |
|---|---|---|---|---|---|
| 2g37-assembly1.cif.gz_A | 1 | 0.167 | 257 | 8.573e-10 | Structure of Thermus thermophilus L-proline dehydrogenase |
| 5ur2-assembly1.cif.gz_A | 1 | 0.157 | 285 | 2.883e-09 | Crystal structure of proline utilization A (PutA) from Bdellovibrio bacteriovorus inactivated by N-propargylglycine |
| 5ur2-assembly2.cif.gz_D | 1 | 0.164 | 285 | 3.255e-09 | Crystal structure of proline utilization A (PutA) from Bdellovibrio bacteriovorus inactivated by N-propargylglycine |

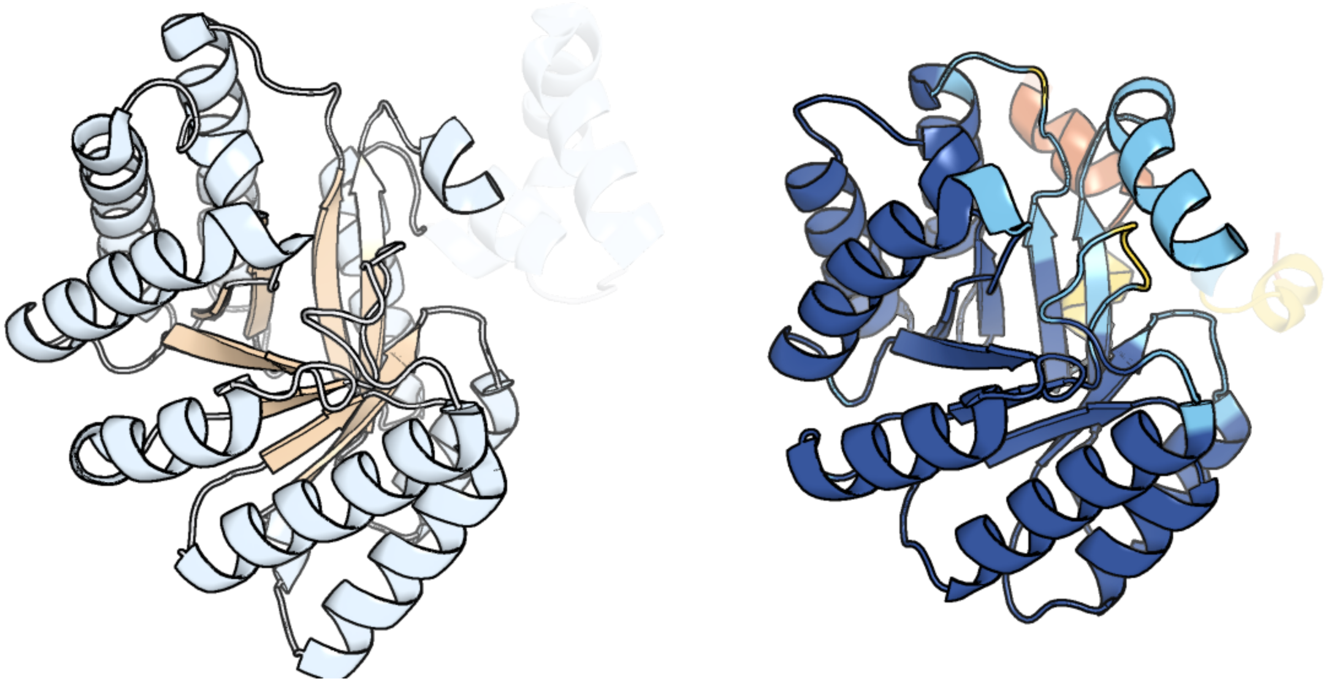

Figure 394: left: reference structure of 2g37 chain A. right: predicted structure of chlorv-1..394, unaligned sequences are shown as transparent

#### chlorv-1..395

- Sequence-based annotation for chlorv-1..395 is hypothetical protein
- No significant structural hit found

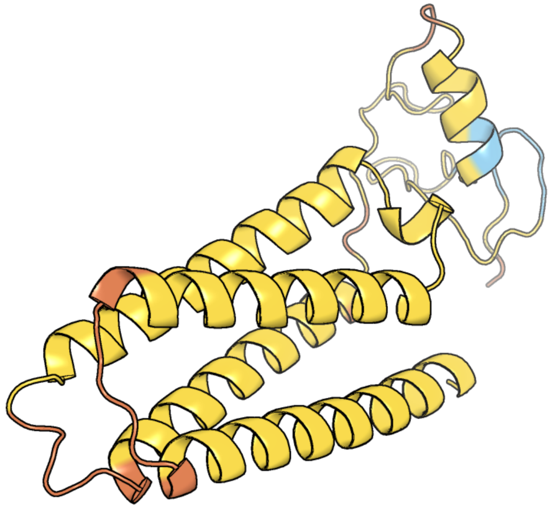

Figure 395: predicted structure of chlorv-1..395

#### chlorv-1..396

- Sequence-based annotation for chlorv-1..396 is hypothetical protein
- No significant structural hit found

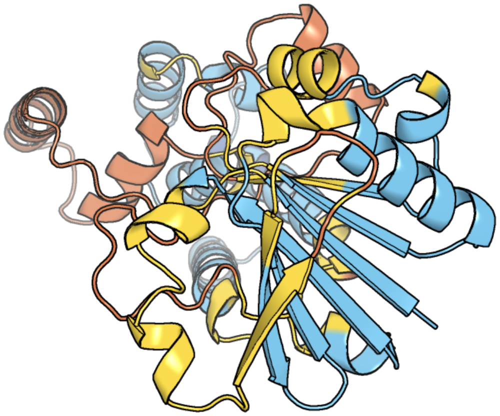

Figure 396: predicted structure of chlorv-1..396

#### chlorv-1..397

- Sequence-based annotation for chlorv-1..397 is putative Glycosidase
- Best hit was 4ac1 chain X: ENDO-N-ACETYL-BETA-D-GLUCOSAMINIDASE

| target | prob | fident | alnlen | evalue | theader |
|---|---|---|---|---|---|
| 4ac1-assembly1.cif.gz_X | 1 | 0.273 | 285 | 8.874e-21 | The structure of a fungal endo-beta-N-acetylglucosaminidase from glycosyl hydrolase family 18, at 1.3A resolution |
| 2y8v-assembly2.cif.gz_D | 1 | 0.27 | 281 | 4.763e-18 | Structure of chitinase, ChiC, from Aspergillus fumigatus. |
| 6k7z-assembly2.cif.gz_B | 1 | 0.18 | 293 | 6.603e-12 | Crystal structure of a GH18 chitinase from Pseudoalteromonas aurantia |

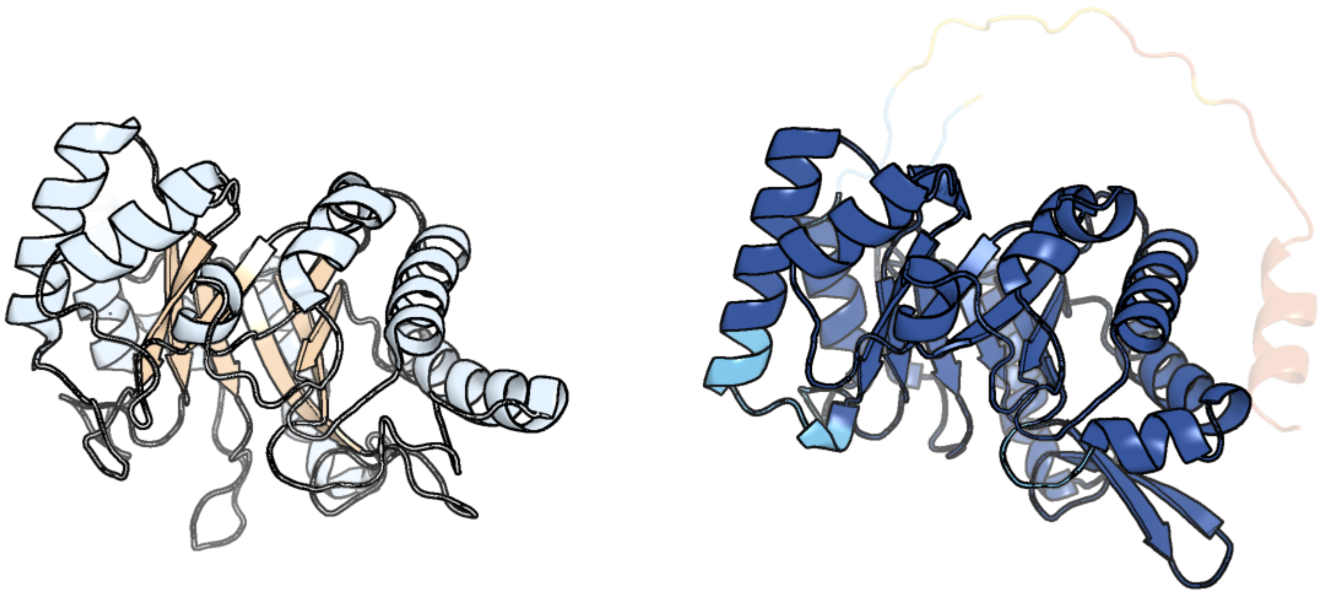

Figure 397: left: reference structure of 4ac1 chain X. right: predicted structure of chlorv-1..397, unaligned sequences are shown as transparent

#### chlorv-1..398

- Sequence-based annotation for chlorv-1..398 is putative NUDIX hydrolase
- Best hit was 2qjt chain A: Nicotinamide-nucleotide adenylyltransferase

| target | prob | fident | alnlen | evalue | theader |
|---|---|---|---|---|---|
| 2qjt-assembly1.cif.gz_A | 1 | 0.133 | 187 | 3.246e-07 | Crystal structure of a bifunctional NMN adenylyltransferase/ADP ribose pyrophosphatase complexed with AMP and MN ion from Francisella tularensis |
| 2r5w-assembly1.cif.gz_A | 1 | 0.162 | 160 | 8.197e-07 | Crystal structure of a bifunctional NMN adenylyltransferase/ADP ribose pyrophosphatase from Francisella tularensis |
| 3gz8-assembly2.cif.gz_D | 1 | 0.145 | 165 | 3.189e-06 | Cocrystal structure of NUDIX domain of Shewanella oneidensis NrtR complexed with ADP ribose |

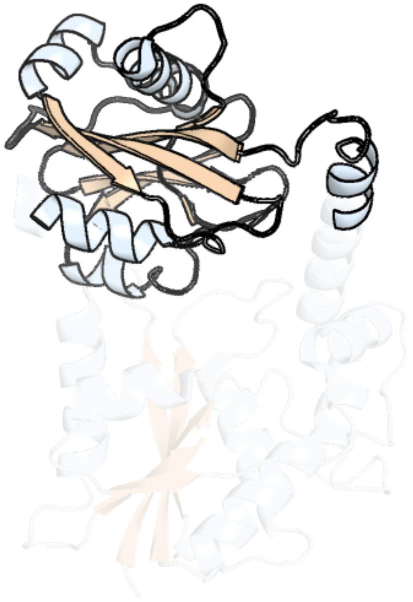

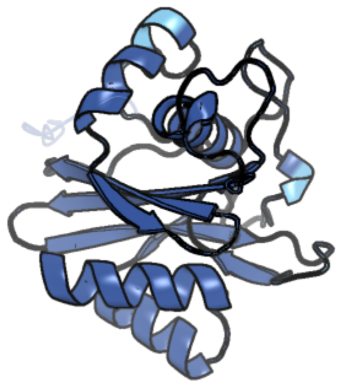

Figure 398: left: reference structure of 2qjt chain A. right: predicted structure of chlorv-1..398, unaligned sequences are shown as transparent

#### chlorv-1..399

- Sequence-based annotation for chlorv-1..399 is hypothetical protein
- No significant structural hit found

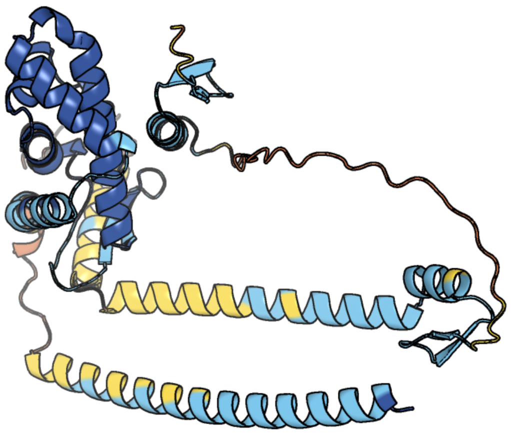

Figure 399: predicted structure of chlorv-1..399

#### chlorv-1..400

- Sequence-based annotation for chlorv-1..400 is putative Hydrolase
- Best hit was 6nkf chain B: Lip_vut4, C3L

| target | prob | fident | alnlen | evalue | theader |
|---|---|---|---|---|---|
| 6nkf-assembly2.cif.gz_B | 1 | 0.156 | 351 | 4.464e-11 | Crystal Structure of the Lipase Lip_vut4 from Goat Rumen metagenome. |
| 5aoa-assembly1.cif.gz_A | 1 | 0.16 | 343 | 1.354e-10 | The structure of a novel thermophilic esterase from the Planctomycetes species, Thermogutta terrifontis, Est2-Propionate bound |
| 5aob-assembly1.cif.gz_A | 1 | 0.153 | 345 | 1.599e-10 | The structure of a novel thermophilic esterase from the Planctomycetes species, Thermogutta terrifontis, Est2-butyrate bound |

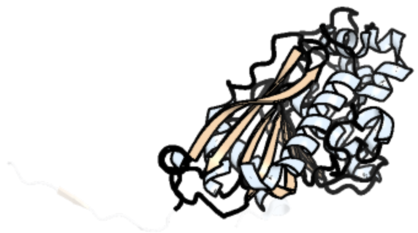

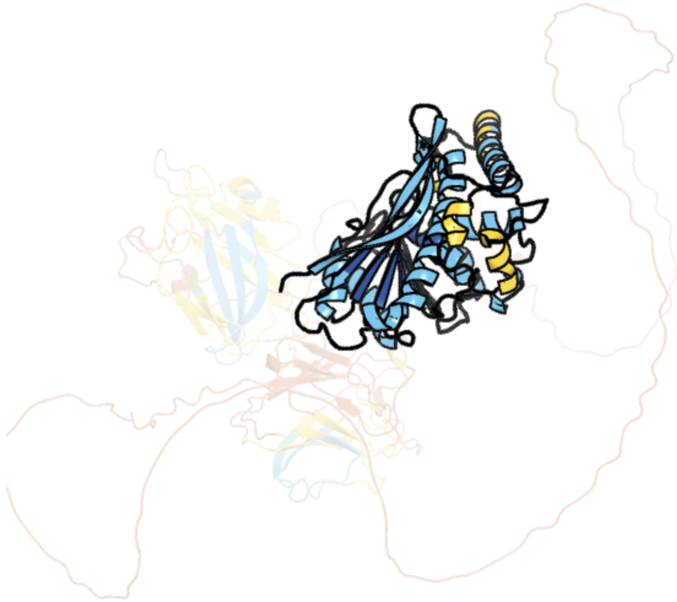

Figure 400: left: reference structure of 6nkf chain B. right: predicted structure of chlorv-1..400, unaligned sequences are shown as transparent

### chlorv-1..401

- Sequence-based annotation for chlorv-1..401 is putative Cytidine deaminase
- Best hit was 1tiy chain A: Guanine deaminase

| target | prob | fident | alnlen | evalue | theader |
|---|---|---|---|---|---|
| 1tiy-assembly1.cif.gz_A | 1 | 0.461 | 156 | 9.332e-22 | X-RAY STRUCTURE OF GUANINE DEAMINASE FROM BACILLUS SUBTILIS NORTHEAST STRUCTURAL GENOMICS CONSORTIUM TARGET SR160 |
| 7dbf-assembly1.cif.gz_D | 1 | 0.401 | 137 | 1.761e-15 | The structure of the Arabidopsis thaliana guanosine deaminase |
| 7w1q-assembly1.cif.gz_D | 1 | 0.367 | 155 | 3.407e-15 | The structure of the Arabidopsis thaliana guanosine deaminase mutant E82Q complexed with 2'-O-methylguanosine |

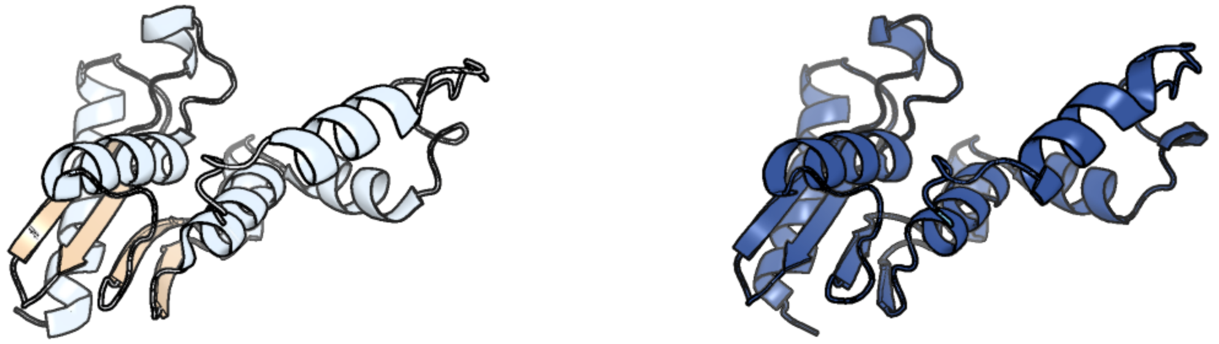

Figure 401: left: reference structure of 1tiy chain A. right: predicted structure of chlorv-1..401, unaligned sequences are shown as transparent

#### chlorv-1..402

- Sequence-based annotation for chlorv-1..402 is hypothetical protein
- No significant structural hit found

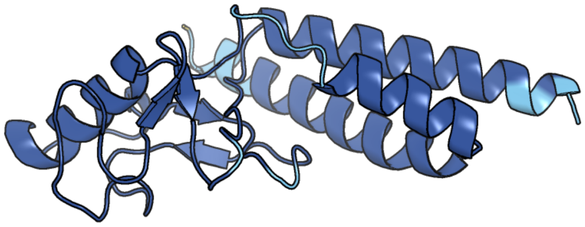

Figure 402: predicted structure of chlorv-1..402

#### chlorv-2..092

- Sequence-based annotation for chlorv-2..092 is
- No significant structural hit found

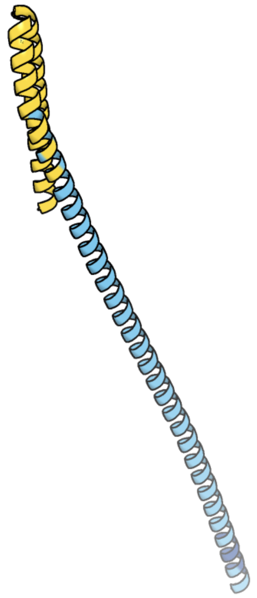

Figure 403: predicted structure of chlorv-2..092

#### chlorv-2..093

- Sequence-based annotation for chlorv-2..093 is
- No significant structural hit found

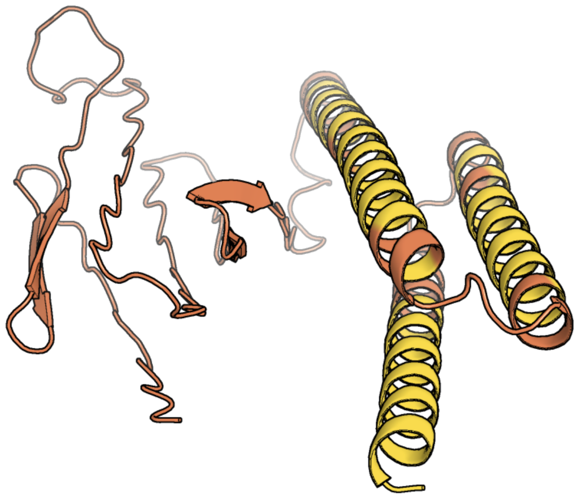

Figure 404: predicted structure of chlorv-2..093

#### chlorv-2..179

- Sequence-based annotation for chlorv-2..179 is
- No significant structural hit found

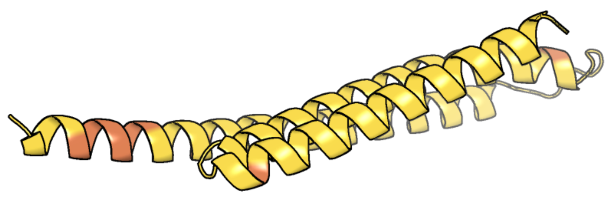

Figure 405: predicted structure of chlorv-2..179

#### chlorv-2..225

- Sequence-based annotation for chlorv-2..225 is
- No significant structural hit found

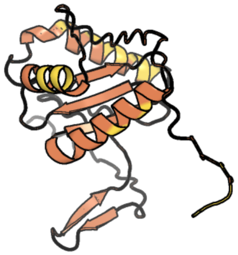

Figure 406: predicted structure of chlorv-2..225

#### chlorv-2..241

- Sequence-based annotation for chlorv-2..241 is
- No significant structural hit found

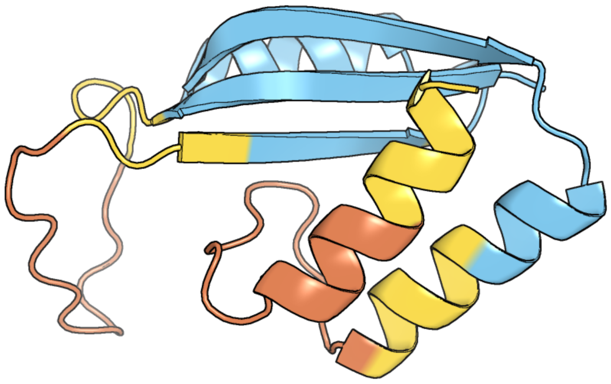

Figure 407: predicted structure of chlorv-2..241

#### chlorv-3..004

- Sequence-based annotation for chlorv-3..004 is
- Best hit was 8ftm chain B: 5'-3' RNA helicase-like protein

| target | prob | fident | alnlen | evalue | theader |
|---|---|---|---|---|---|
| 8ftm-assembly2.cif.gz_B | 1 | 0.167 | 691 | 6.478e-20 | Setx-ssRNA-ADP-SO4 complex |
| 8ftm-assembly1.cif.gz_A | 1 | 0.167 | 670 | 8.287e-20 | Setx-ssRNA-ADP-SO4 complex |
| 5ean-assembly1.cif.gz_A-2 | 1 | 0.174 | 584 | 2.516e-17 | Crystal structure of Dna2 in complex with a 5' overhang DNA |

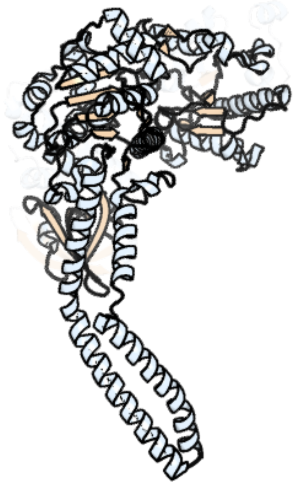

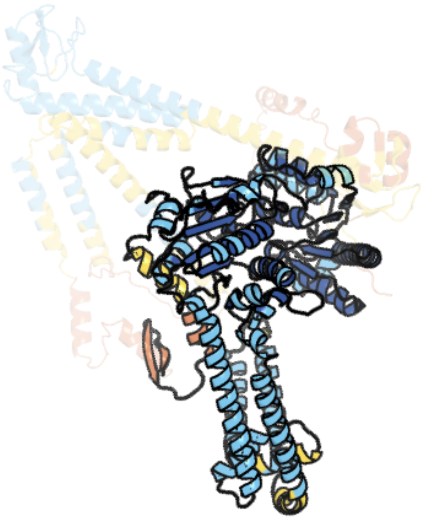

Figure 408: left: reference structure of 8ftm chain B. right: predicted structure of chlorv-3..004, unaligned sequences are shown as transparent

#### chlorv-3..006

- Sequence-based annotation for chlorv-3..006 is
- Best hit was 2ip2 chain B: Probable phenazine-specific methyltransferase

| target | prob | fident | alnlen | evalue | theader |
|---|---|---|---|---|---|
| 2ip2-assembly1.cif.gz_B | 1 | 0.084 | 237 | 1.325e-05 | Structure of the Pyocyanin Biosynthetic Protein PhzM |
| 3e05-assembly1.cif.gz_G | 1 | 0.118 | 227 | 1.325e-05 | CRYSTAL STRUCTURE OF Precorrin-6y C5,15-methyltransferase FROM Geobacter metallireducens GS-15 |
| 3dtn-assembly1.cif.gz_A | 1 | 0.094 | 244 | 1.41e-05 | Crystal structure of putative Methyltransferase-MM_2633 from Methanosarcina mazei . |

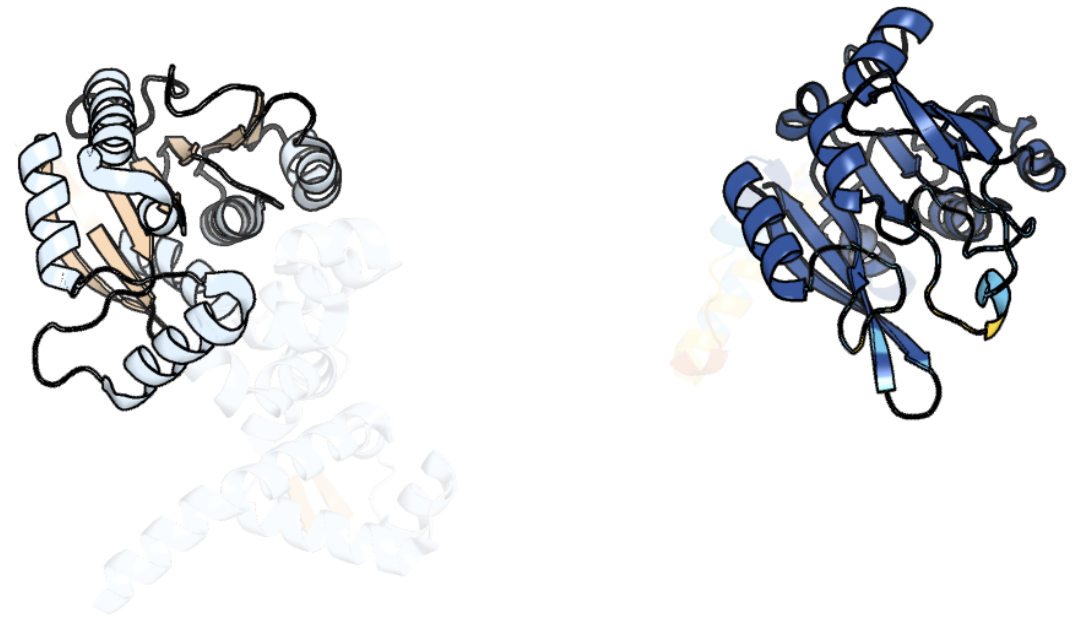

Figure 409: left: reference structure of 2ip2 chain B. right: predicted structure of chlorv-3..006, unaligned sequences are shown as transparent

#### chlorv-3..007

- Sequence-based annotation for chlorv-3..007 is
- No significant structural hit found

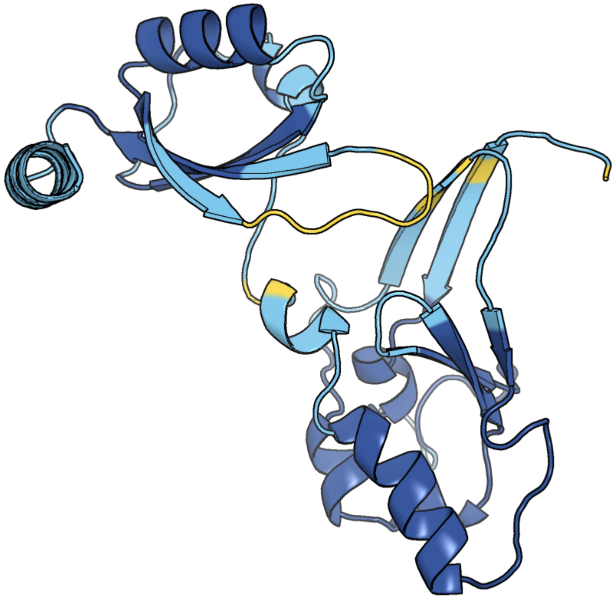

Figure 410: predicted structure of chlorv-3..007

#### chlorv-3..031

- Sequence-based annotation for chlorv-3..031 is
- No significant structural hit found

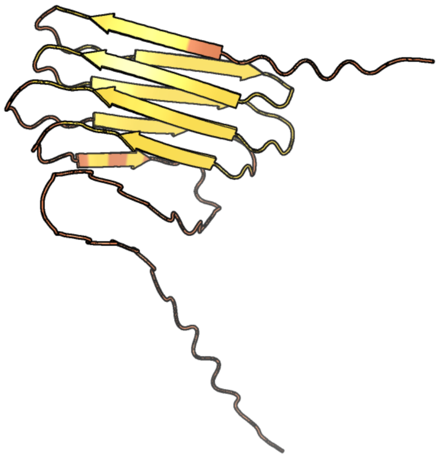

Figure 411: predicted structure of chlorv-3..031

#### chlorv-3..032

- Sequence-based annotation for chlorv-3..032 is
- No significant structural hit found

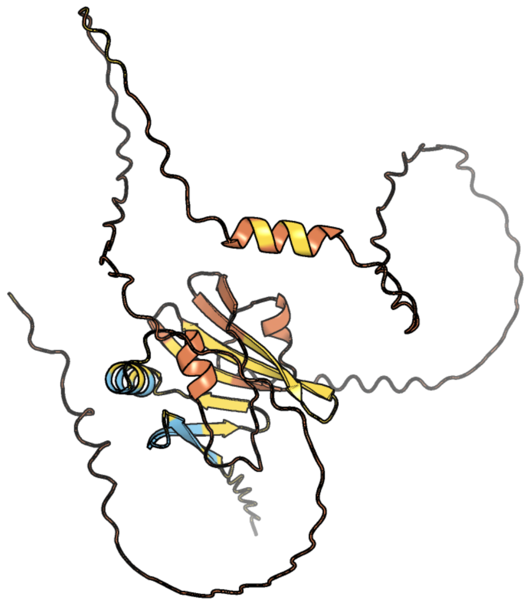

Figure 412: predicted structure of chlorv-3..032

#### chlorv-3..079

- Sequence-based annotation for chlorv-3..079 is
- No significant structural hit found

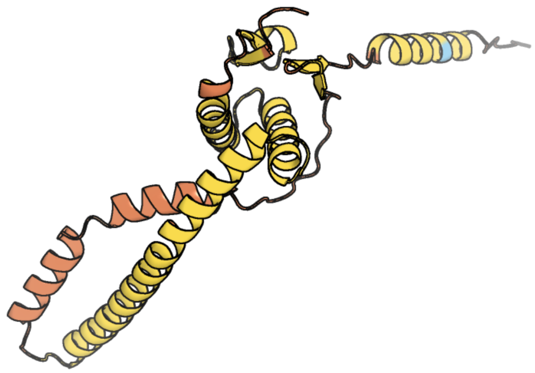

Figure 413: predicted structure of chlorv-3..079

#### chlorv-3..080

- Sequence-based annotation for chlorv-3..080 is
- No significant structural hit found

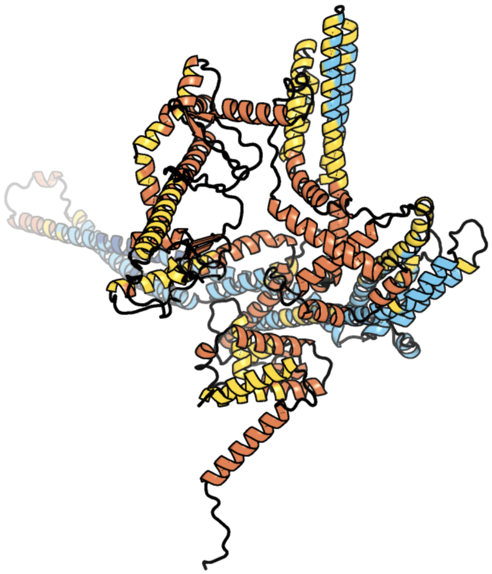

Figure 414: predicted structure of chlorv-3..080

#### chlorv-3..095

- Sequence-based annotation for chlorv-3..095 is
- No significant structural hit found

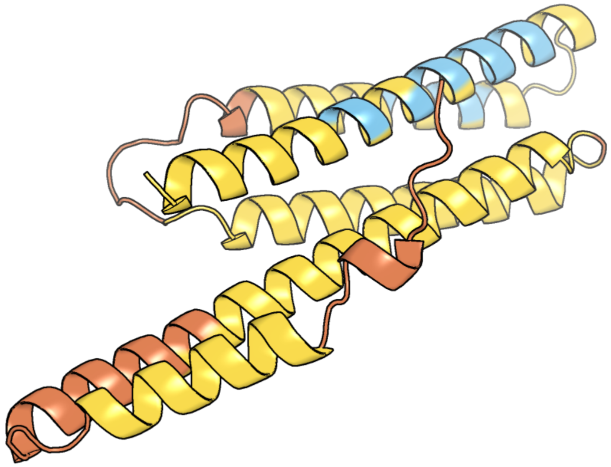

Figure 415: predicted structure of chlorv-3..095

#### chlorv-3..096

- Sequence-based annotation for chlorv-3..096 is
- No significant structural hit found

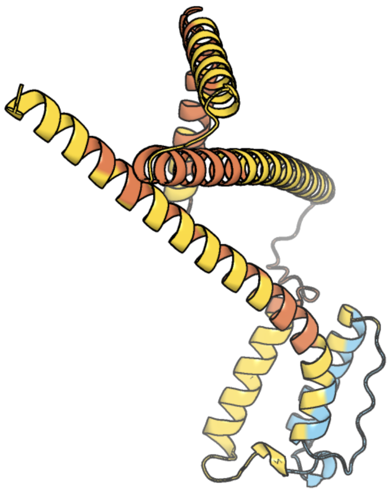

Figure 416: predicted structure of chlorv-3..096

### chlorv-3..102

- Sequence-based annotation for chlorv-3..102 is
- Best hit was 7eew chain A: Type I restriction-modification system methyltransferase subunit

| target | prob | fident | alnlen | evalue | theader |
|---|---|---|---|---|---|
| 7eew-assembly1.cif.gz_A-2 | 1 | 0.171 | 826 | 4.718e-25 | Crystal structure of the intact MTase from Vibrio vulnificus YJ016 in complex with the DNA-mimicking Ocr protein and the S-adenosyl-L-homocysteine (SAH) |
| 1ydx-assembly1.cif.gz_A | 1 | 0.217 | 377 | 3.143e-19 | Crystal structure of Type-I restriction-modification system S subunit from M. genitalium |
| 2okc-assembly1.cif.gz_A | 1 | 0.235 | 416 | 3.341e-17 | Crystal structure of Type I restriction enzyme StySJI M protein (NP_813429.1) from Bacteroides thetaiotaomicron VPI-5482 at 2.20 A resolution |

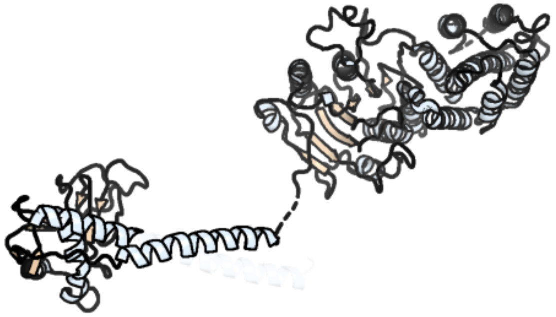

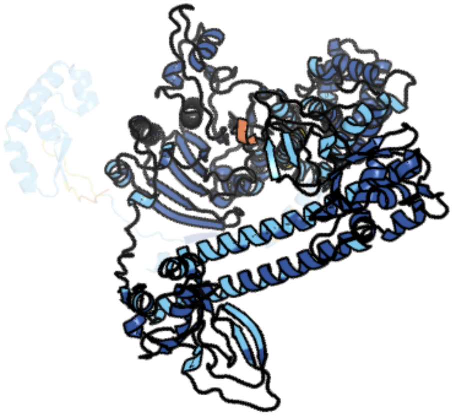

Figure 417: left: reference structure of 7eew chain A. right: predicted structure of chlorv-3..102, unaligned sequences are shown as transparent

#### chlorv-3..103

- Sequence-based annotation for chlorv-3..103 is
- Best hit was 4xqk chain B: LlaBIII

| target | prob | fident | alnlen | evalue | theader |
|---|---|---|---|---|---|
| 4xqk-assembly2.cif.gz_B | 1 | 0.174 | 424 | 3.565e-10 | ATP-dependent Type ISP restriction-modification enzyme LlaBIII bound to DNA |
| 4xqk-assembly1.cif.gz_A | 1 | 0.15 | 513 | 1.235e-09 | ATP-dependent Type ISP restriction-modification enzyme LlaBIII bound to DNA |
| 3h1t-assembly1.cif.gz_A | 1 | 0.146 | 246 | 1.751e-05 | The fragment structure of a putative HsdR subunit of a type I restriction enzyme from Vibrio vulnificus YJ016 |

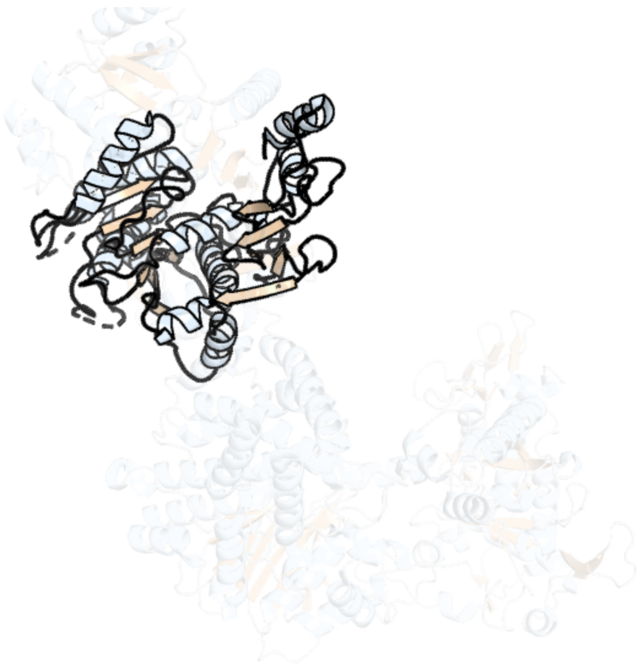

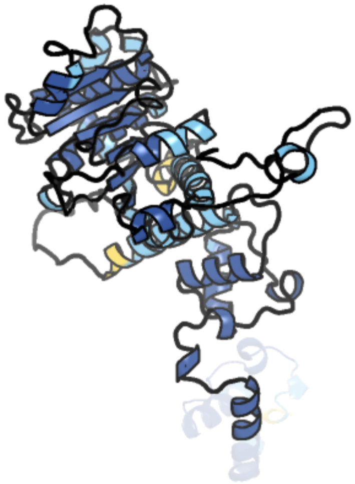

Figure 418: left: reference structure of 4xqk chain B. right: predicted structure of chlorv-3..103, unaligned sequences are shown as transparent

#### chlorv-3..104

- Sequence-based annotation for chlorv-3..104 is
- No significant structural hit found

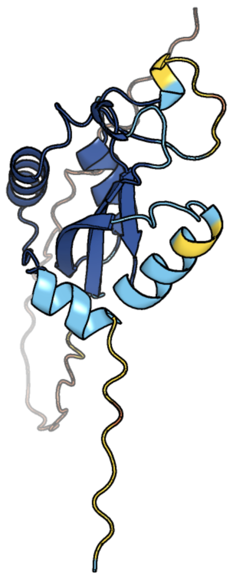

Figure 419: predicted structure of chlorv-3..104

#### chlorv-3..114

- Sequence-based annotation for chlorv-3..114 is
- No significant structural hit found

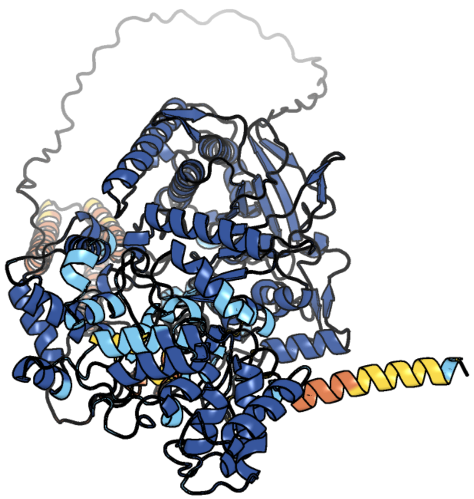

Figure 420: predicted structure of chlorv-3..114

#### chlorv-3..120

- Sequence-based annotation for chlorv-3..120 is
- No significant structural hit found

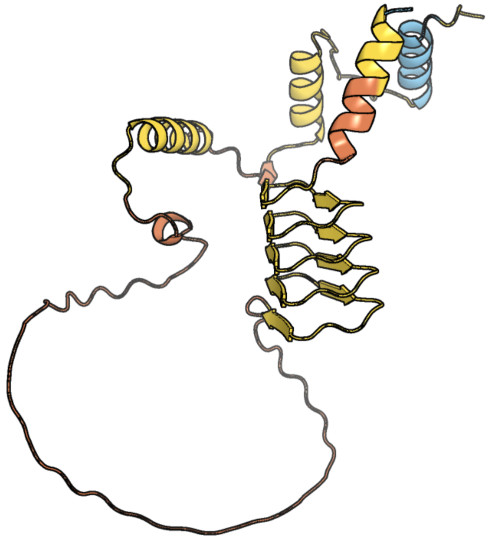

Figure 421: predicted structure of chlorv-3..120

#### chlorv-3..133

- Sequence-based annotation for chlorv-3..133 is
- Best hit was 7k6p chain K: Histone-lysine N-methyltransferase, H3 lysine-79 specific

| target | prob | fident | alnlen | evalue | theader |
|---|---|---|---|---|---|
| 7k6p-assembly1.cif.gz_K | 1 | 0.186 | 193 | 8.781e-09 | Active state Dot1 bound to the unacetylated H4 nucleosome |
| 5fa8-assembly1.cif.gz_A | 1 | 0.215 | 153 | 2.793e-08 | SAM complex with aKMT from the hyperthermophilic archaeon Sulfolobus islandicu |
| 1u2z-assembly2.cif.gz_B | 1 | 0.194 | 190 | 4.107e-08 | Crystal structure of histone K79 methyltransferase Dot1p from yeast |

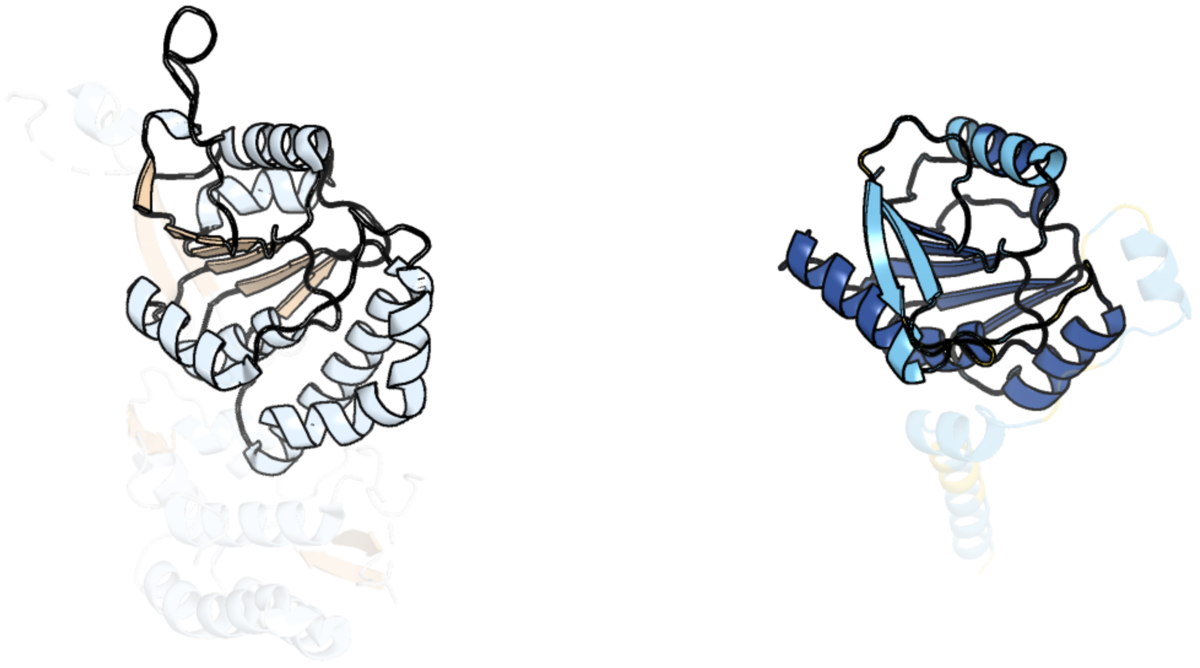

Figure 422: left: reference structure of 7k6p chain K. right: predicted structure of chlorv-3..133, unaligned sequences are shown as transparent

#### chlorv-3..150

- Sequence-based annotation for chlorv-3..150 is
- No significant structural hit found

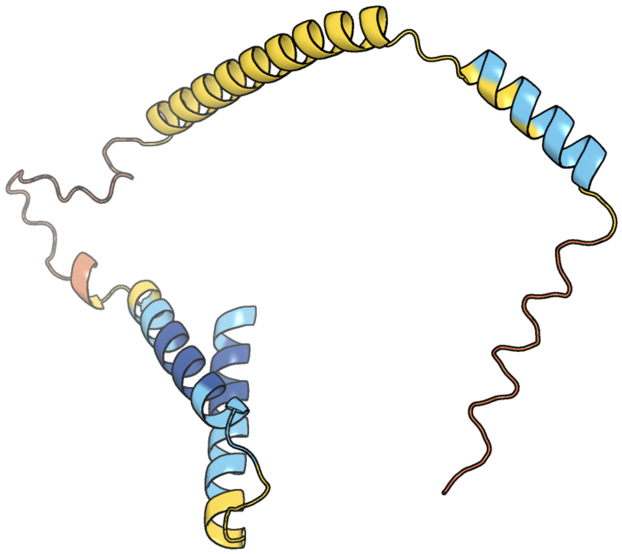

Figure 423: predicted structure of chlorv-3..150

#### chlorv-3..151

- Sequence-based annotation for chlorv-3..151 is
- No significant structural hit found

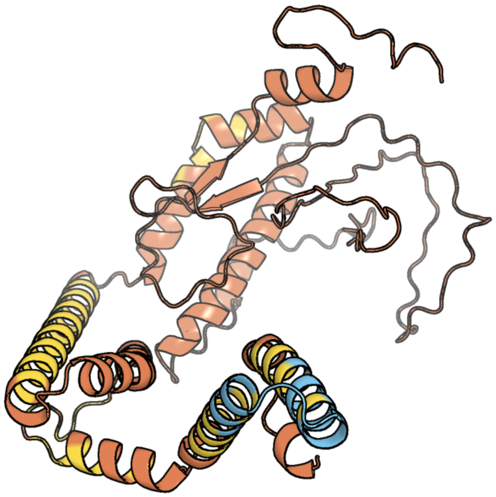

Figure 424: predicted structure of chlorv-3..151

#### chlorv-3..152

- Sequence-based annotation for chlorv-3..152 is
- No significant structural hit found

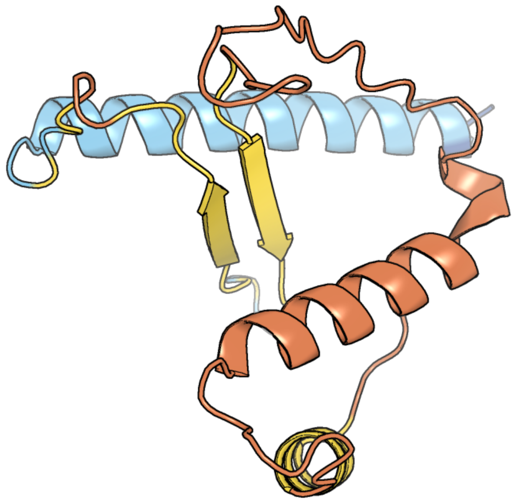

Figure 425: predicted structure of chlorv-3..152

#### chlorv-3..153

- Sequence-based annotation for chlorv-3..153 is
- No significant structural hit found

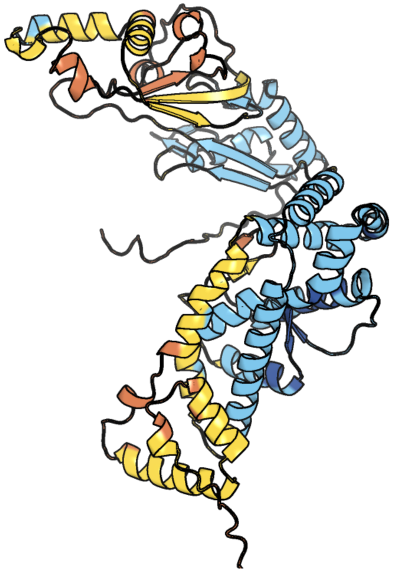

Figure 426: predicted structure of chlorv-3..153

#### chlorv-3..196

- Sequence-based annotation for chlorv-3..196 is
- No significant structural hit found

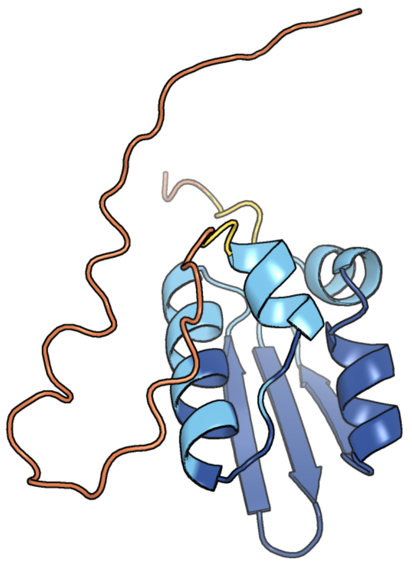

Figure 427: predicted structure of chlorv-3..196

#### chlorv-3..205

- Sequence-based annotation for chlorv-3..205 is
- No significant structural hit found

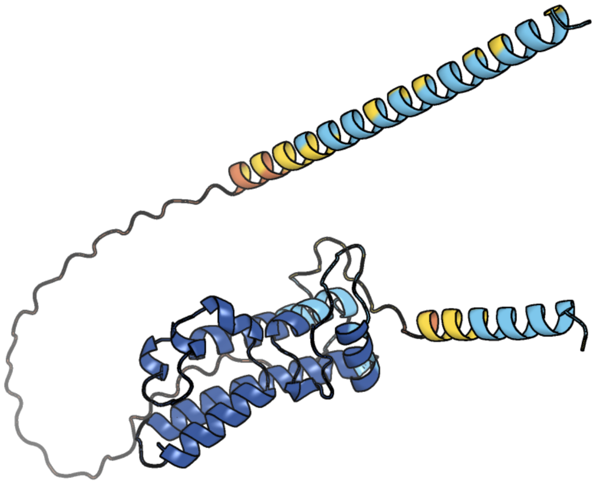

Figure 428: predicted structure of chlorv-3..205

#### chlorv-3..219

- Sequence-based annotation for chlorv-3..219 is
- No significant structural hit found

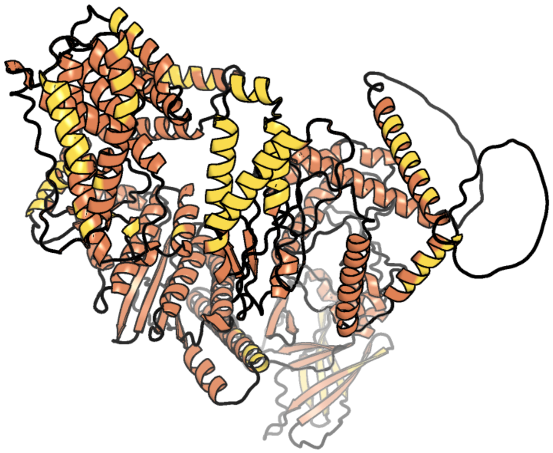

Figure 429: predicted structure of chlorv-3..219

#### chlorv-3..225

- Sequence-based annotation for chlorv-3..225 is
- No significant structural hit found

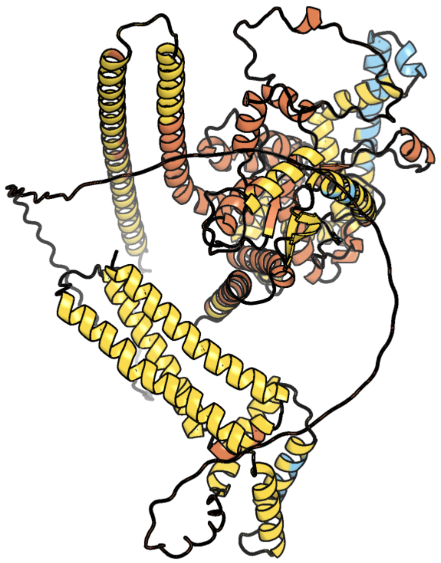

Figure 430: predicted structure of chlorv-3..225

#### chlorv-3..329

- Sequence-based annotation for chlorv-3..329 is
- No significant structural hit found

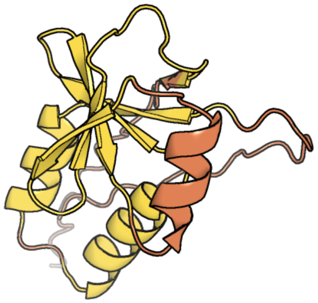

Figure 431: predicted structure of chlorv-3..329

#### chlorv-3..350

- Sequence-based annotation for chlorv-3..350 is
- No significant structural hit found

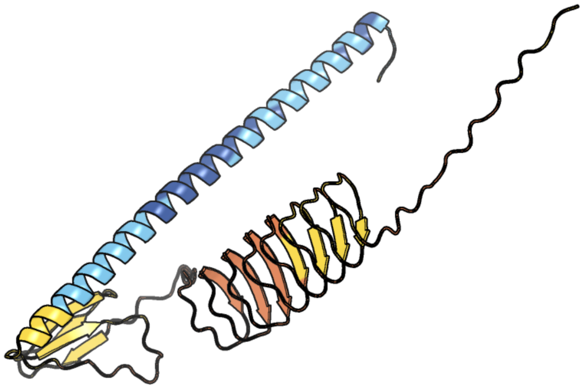

Figure 432: predicted structure of chlorv-3..350

#### chlorv-3..384

- Sequence-based annotation for chlorv-3..384 is
- No significant structural hit found

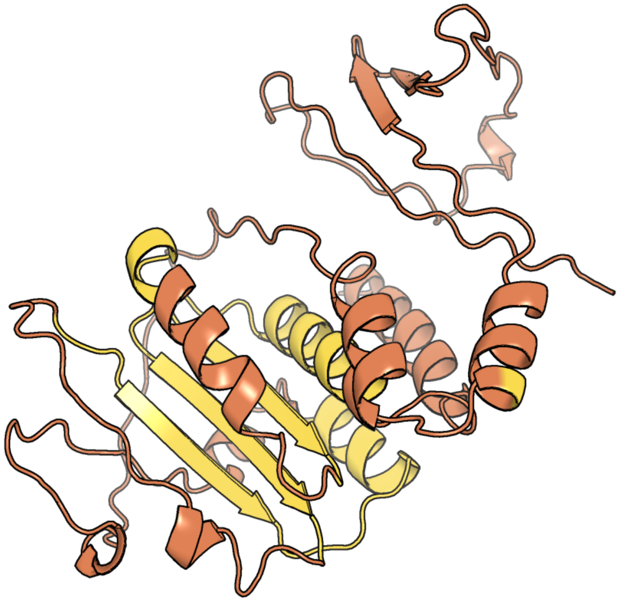

Figure 433: predicted structure of chlorv-3..384

#### chlorv-3..391

- Sequence-based annotation for chlorv-3..391 is
- Best hit was 6a0w chain A: Lipase

| target | prob | fident | alnlen | evalue | theader |
|---|---|---|---|---|---|
| 6a0w-assembly1.cif.gz_A | 1 | 0.204 | 171 | 3.713e-08 | Crystal structure of lipase from Rhizopus microsporus var. chinensis |
| 6qpr-assembly1.cif.gz_A | 1 | 0.208 | 187 | 4.175e-08 | Rhizomucor miehei lipase propeptide complex, Ser95/Ile96 deletion mutant |
| 5ap9-assembly1.cif.gz_A | 1 | 0.201 | 258 | 6.672e-08 | Controlled lid-opening in Thermomyces lanuginosus lipase - a switch for activity and binding |

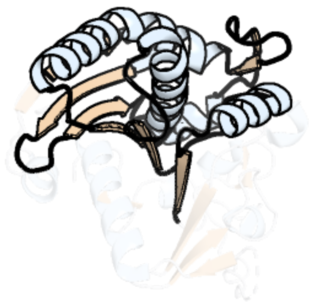

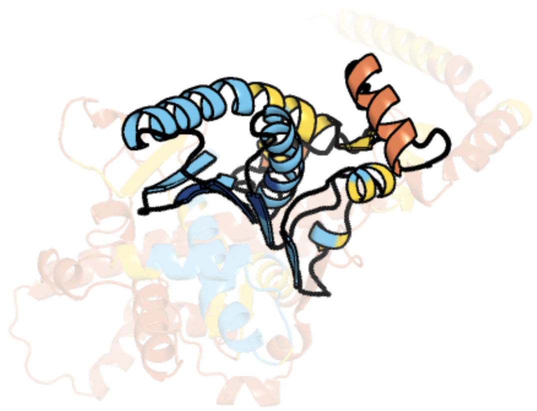

Figure 434: left: reference structure of 6a0w chain A. right: predicted structure of chlorv-3..391, unaligned sequences are shown as transparent

#### chlorv-4..001

- Sequence-based annotation for chlorv-4..001 is
- Best hit was 5l9b chain A: Egl nine homolog 1

| target | prob | fident | alnlen | evalue | theader |
|---|---|---|---|---|---|
| 5l9b-assembly1.cif.gz_A | 1 | 0.128 | 210 | 6.825e-07 | HIF PROLYL HYDROXYLASE 2 (PHD2/ EGLN1) IN COMPLEX WITH 2-OXOGLUTARATE (2OG) AND HIF-1ALPHA CODD (556-574) |
| 4j25-assembly8.cif.gz_H | 1 | 0.094 | 211 | 8.055e-07 | Crystal structure of a Pseudomonas putida prolyl-4-hydroxylase (P4H) |
| 4j25-assembly7.cif.gz_G | 1 | 0.095 | 209 | 1.062e-06 | Crystal structure of a Pseudomonas putida prolyl-4-hydroxylase (P4H) |

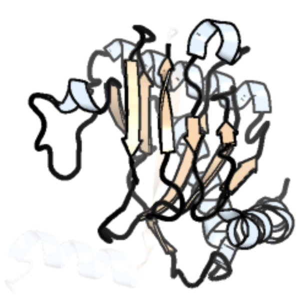

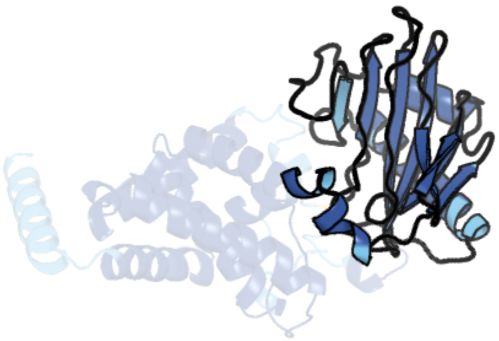

Figure 435: left: reference structure of 5l9b chain A. right: predicted structure of chlorv-4..001, unaligned sequences are shown as transparent

#### chlorv-4..002

- Sequence-based annotation for chlorv-4..002 is
- Best hit was 2rdq chain A: 1-deoxypentalenic acid 11-beta hydroxylase; Fe(II)/alpha-ketoglutarate dependent hydroxylase

| target | prob | fident | alnlen | evalue | theader |
|---|---|---|---|---|---|
| 2rdq-assembly1.cif.gz_A | 1 | 0.128 | 272 | 9.521e-10 | Crystal Structure of PtlH with Fe/alpha ketoglutarate bound |
| 7eys-assembly2.cif.gz_A | 1 | 0.147 | 264 | 4.767e-09 | Complex structure of SptF with Fe, alpha-ketoglutarate, and andiconin D |
| 7eyr-assembly2.cif.gz_A | 1 | 0.159 | 264 | 5.74e-09 | Fe(II)/(alpha)ketoglutarate-dependent dioxygenase SptF apo |

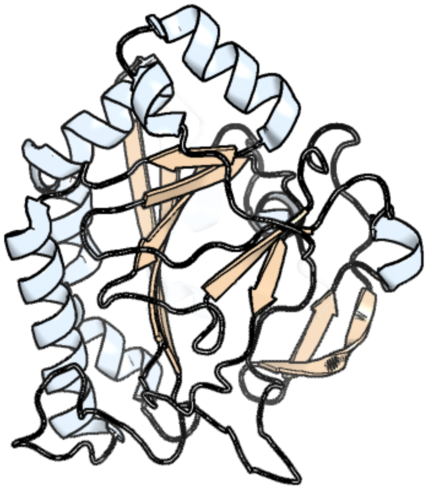

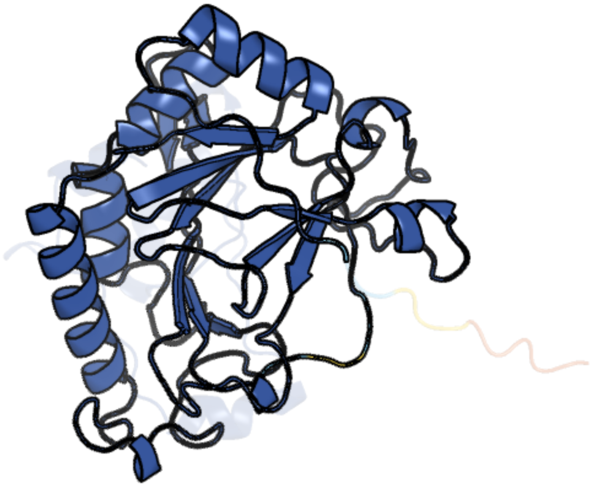

Figure 436: left: reference structure of 2rdq chain A. right: predicted structure of chlorv-4..002, unaligned sequences are shown as transparent

#### chlorv-4..004

- Sequence-based annotation for chlorv-4..004 is
- Best hit was 3ngm chain B: Extracellular lipase

| target | prob | fident | alnlen | evalue | theader |
|---|---|---|---|---|---|
| 3ngm-assembly2.cif.gz_B | 1 | 0.2 | 170 | 1.182e-07 | Crystal structure of lipase from Gibberella zeae |
| 6a0w-assembly1.cif.gz_A | 1 | 0.177 | 175 | 1.329e-07 | Crystal structure of lipase from Rhizopus microsporus var. chinensis |
| 2yij-assembly1.cif.gz_A | 1 | 0.143 | 306 | 1.68e-07 | Crystal Structure of phospholipase A1 |

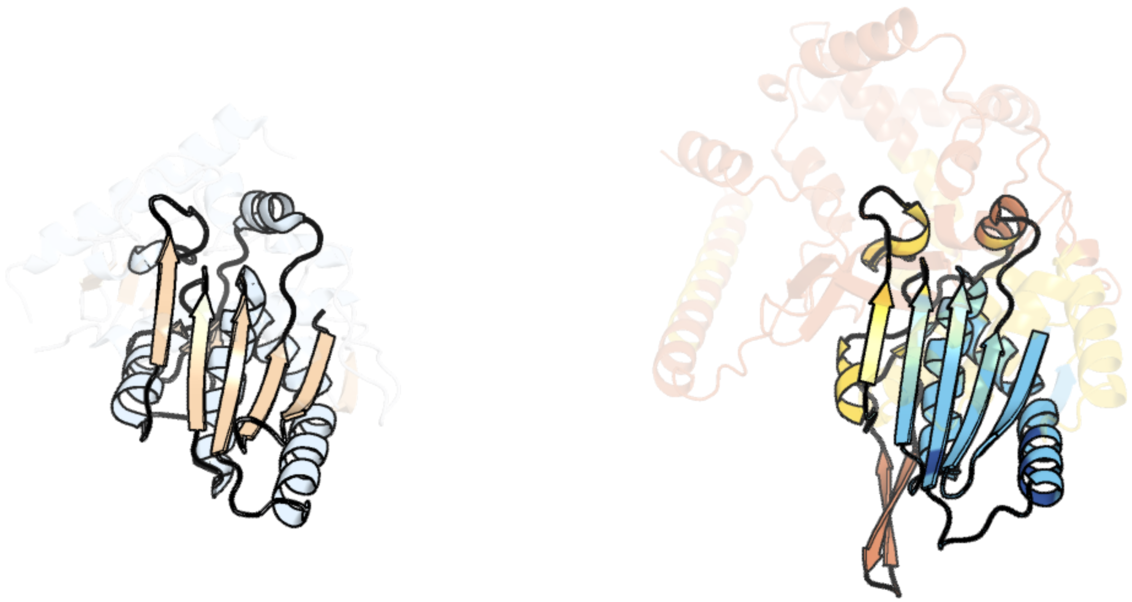

Figure 437: left: reference structure of 3ngm chain B. right: predicted structure of chlorv-4..004, unaligned sequences are shown as transparent

#### chlorv-4..011

- Sequence-based annotation for chlorv-4..011 is
- Best hit was 5h70 chain A: Probable thymidylate kinase

| target | prob | fident | alnlen | evalue | theader |
|---|---|---|---|---|---|
| 5h70-assembly1.cif.gz_A | 1 | 0.143 | 209 | 3.871e-06 | Crystal structure of ADP bound dTMP kinase (st1543) from Sulfolobus Tokodaii Strain7 |
| 7e9v-assembly1.cif.gz_A | 1 | 0.131 | 197 | 1.292e-05 | The Crystal Structure of human UMP-CMP kinase from Biortus. |
| 4ukd-assembly1.cif.gz_A | 1 | 0.154 | 188 | 1.377e-05 | UMP/CMP KINASE FROM SLIME MOLD COMPLEXED WITH ADP, UDP, BERYLLIUM FLUORIDE |

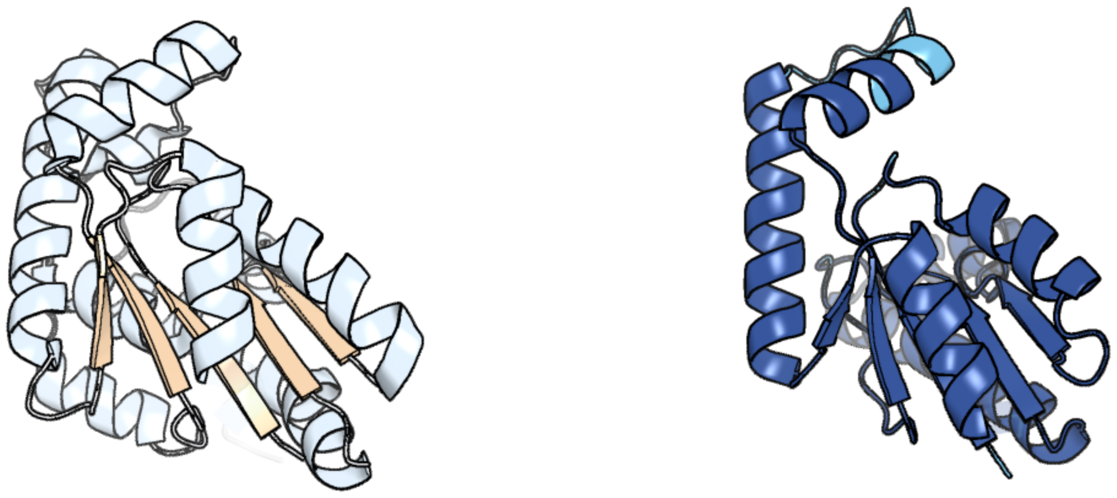

Figure 438: left: reference structure of 5h70 chain A. right: predicted structure of chlorv-4..011, unaligned sequences are shown as transparent

#### chlorv-4..012

- Sequence-based annotation for chlorv-4..012 is
- Best hit was 8gxn chain B: caffeyl-CoA-O-methyltransferase

| target | prob | fident | alnlen | evalue | theader |
|---|---|---|---|---|---|
| 8gxn-assembly1.cif.gz_B | 1 | 0.168 | 214 | 6.969e-08 | The crystal structure of CsFAOMT2 in complex with SAH |
| 5zw3-assembly1.cif.gz_B | 1 | 0.12 | 250 | 1.869e-07 | Crystal Structure of TrmR from B. subtilis |
| 5zw3-assembly1.cif.gz_A | 1 | 0.111 | 260 | 2.098e-07 | Crystal Structure of TrmR from B. subtilis |

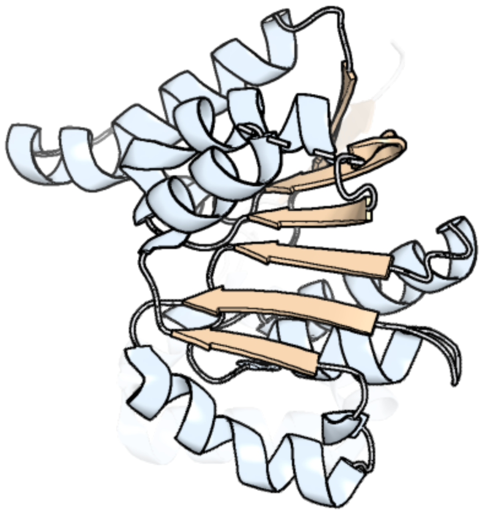

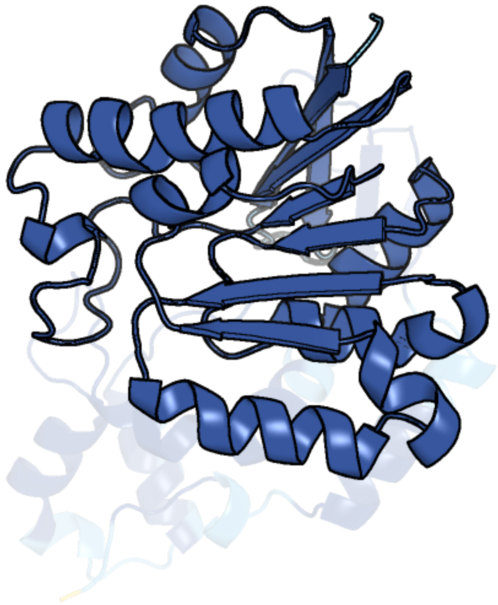

Figure 439: left: reference structure of 8gxn chain B. right: predicted structure of chlorv-4..012, unaligned sequences are shown as transparent

#### chlorv-4..013

- Sequence-based annotation for chlorv-4..013 is
- No significant structural hit found

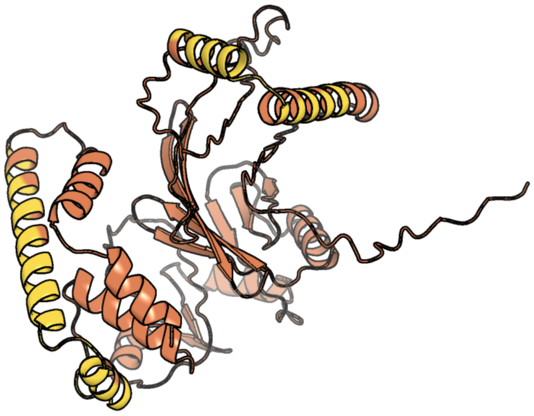

Figure 440: predicted structure of chlorv-4..013

#### chlorv-4..014

- Sequence-based annotation for chlorv-4..014 is
- Best hit was 1tf5 chain A: Preprotein translocase secA subunit

| target | prob | fident | alnlen | evalue | theader |
|---|---|---|---|---|---|
| 1tf5-assembly1.cif.gz_A | 1 | 0.1 | 744 | 1.421e-08 | Crystal structure of SecA in an open conformation from Bacillus Subtilis |
| 1tf2-assembly1.cif.gz_A | 1 | 0.103 | 761 | 3.061e-08 | Crystal structure of SecA:ADP in an open conformation from Bacillus Subtilis |
| 3jv2-assembly1.cif.gz_A | 1 | 0.094 | 787 | 5.262e-08 | Crystal Structure of B. subtilis SecA with bound peptide |

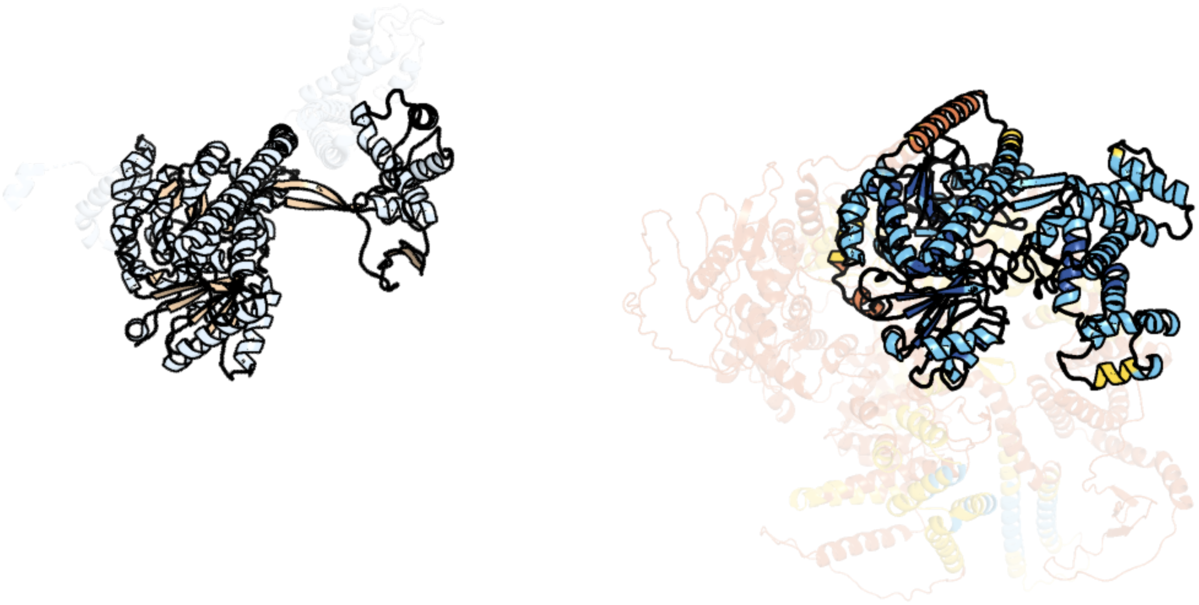

Figure 441: left: reference structure of 1tf5 chain A. right: predicted structure of chlorv-4..014, unaligned sequences are shown as transparent

#### chlorv-4..016

- Sequence-based annotation for chlorv-4..016 is
- No significant structural hit found

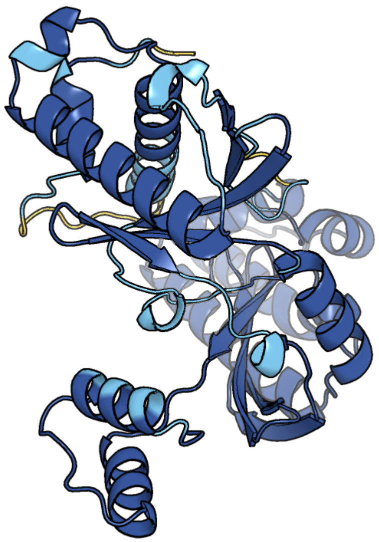

Figure 442: predicted structure of chlorv-4..016

### chlorv-4..017

- Sequence-based annotation for chlorv-4..017 is
- Best hit was 7lt5 chain C: Site-specific DNA-methyltransferase (adenine-specific)

| target | prob | fident | alnlen | evalue | theader |
|---|---|---|---|---|---|
| 7lt5-assembly3.cif.gz_C | 1 | 0.169 | 542 | 1.807e-18 | CamA Adenine Methyltransferase Complexed to Cognate Substrate DNA and Cofactor SAH |
| 7rfk-assembly3.cif.gz_C | 1 | 0.164 | 541 | 5.356e-18 | CamA Adenine Methyltransferase Complexed to Cognate Substrate DNA and Inhibitor Sinefungin |
| 7qw7-assembly1.cif.gz_A | 1 | 0.166 | 523 | 1.325e-17 | Adenine-specific DNA methyltransferase M.BseCI complexed with AdoHcy and cognate fully methylated DNA duplex |

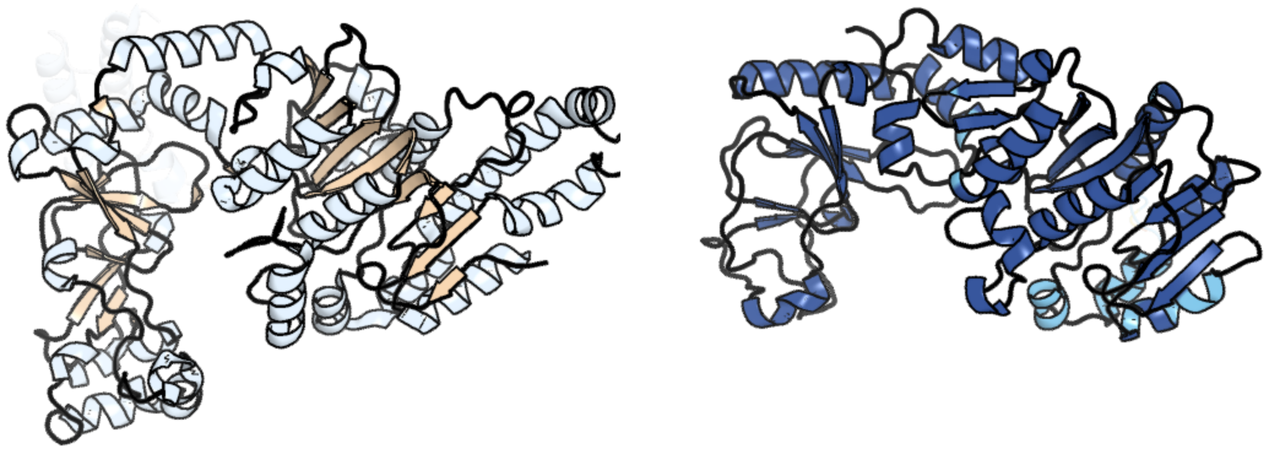

Figure 443: left: reference structure of 7lt5 chain C. right: predicted structure of chlorv-4..017, unaligned sequences are shown as transparent

#### chlorv-4..028

- Sequence-based annotation for chlorv-4..028 is
- No significant structural hit found

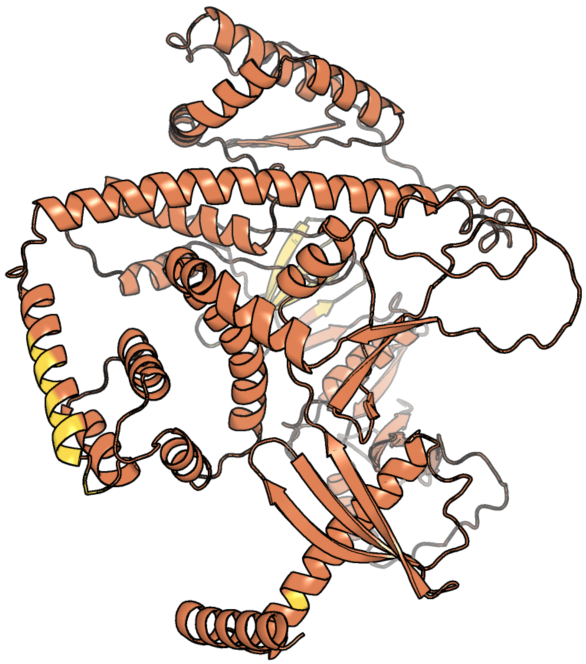

Figure 444: predicted structure of chlorv-4..028

#### chlorv-4..032

- Sequence-based annotation for chlorv-4..032 is
- No significant structural hit found

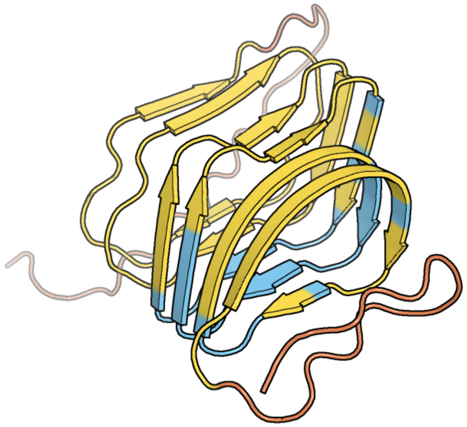

Figure 445: predicted structure of chlorv-4..032

#### chlorv-4..037

- Sequence-based annotation for chlorv-4..037 is
- No significant structural hit found

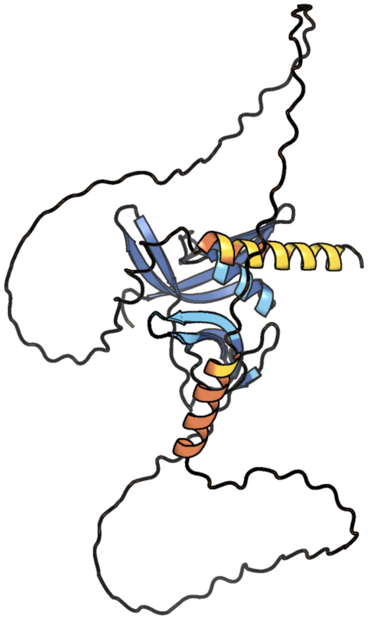

Figure 446: predicted structure of chlorv-4..037

#### chlorv-4..048

- Sequence-based annotation for chlorv-4..048 is
- No significant structural hit found

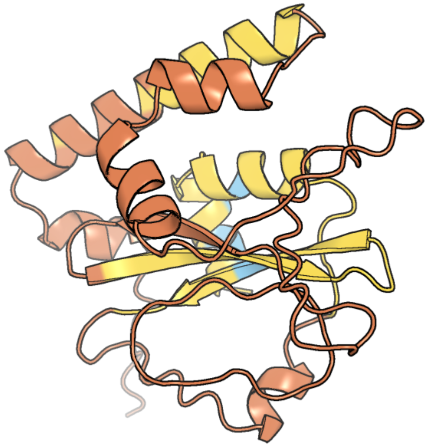

Figure 447: predicted structure of chlorv-4..048

#### chlorv-4..059

- Sequence-based annotation for chlorv-4..059 is
- Best hit was 8ssr chain A: Transcriptional repressor CTCF

| target | prob | fident | alnlen | evalue | theader |
|---|---|---|---|---|---|
| 8ssr-assembly1.cif.gz_A | 1 | 0.194 | 268 | 2.036e-08 | ZnFs 3-11 of CCCTC-binding factor (CTCF) Complexed with 35mer DNA 35-20 |
| 8ssu-assembly1.cif.gz_A | 1 | 0.212 | 250 | 3.951e-08 | ZnFs 3-11 of CCCTC-binding factor (CTCF) Complexed with 19mer DNA |
| 8ffz-assembly1.cif.gz_A | 1 | 0.222 | 193 | 4.458e-08 | TFIIIA-TFIIIC-Brf1-TBP complex bound to 5S rRNA gene |

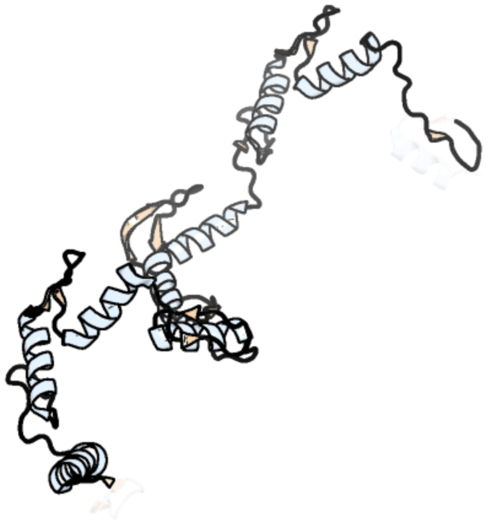

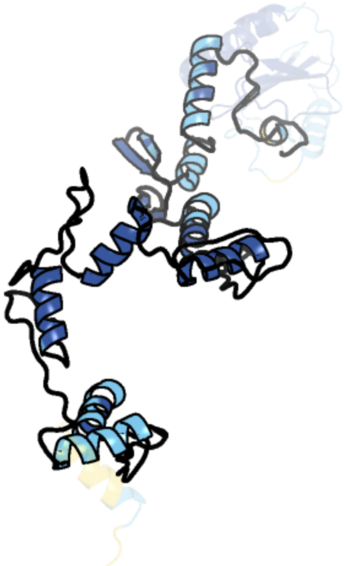

Figure 448: left: reference structure of 8ssr chain A. right: predicted structure of chlorv-4..059, unaligned sequences are shown as transparent

#### chlorv-4..061

- Sequence-based annotation for chlorv-4..061 is
- No significant structural hit found

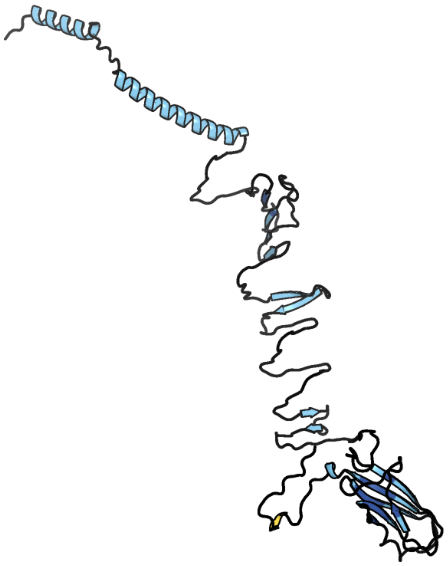

Figure 449: predicted structure of chlorv-4..061

#### chlorv-4..064

- Sequence-based annotation for chlorv-4..064 is
- No significant structural hit found

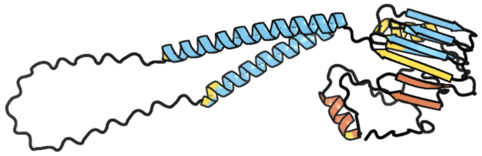

Figure 450: predicted structure of chlorv-4..064

#### chlorv-4..071

- Sequence-based annotation for chlorv-4..071 is
- Best hit was 2qip chain A: Protein of unknown function VPA0982

| target | prob | fident | alnlen | evalue | theader |
|---|---|---|---|---|---|
| 2qip-assembly1.cif.gz_A-2 | 1 | 0.201 | 159 | 4.608e-09 | Crystal structure of a protein of unknown function VPA0982 from Vibrio parahaemolyticus RIMD 2210633 |
| 5yaa-assembly4.cif.gz_D | 1 | 0.175 | 148 | 1.056e-06 | Crystal structure of Marf1 NYN domain from Mus musculus |
| 6fdl-assembly2.cif.gz_B | 1 | 0.164 | 152 | 1.612e-06 | Crystal structure of the NYN domain of human MARF1 |

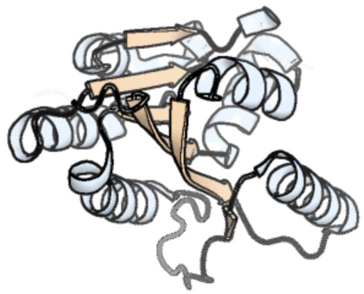

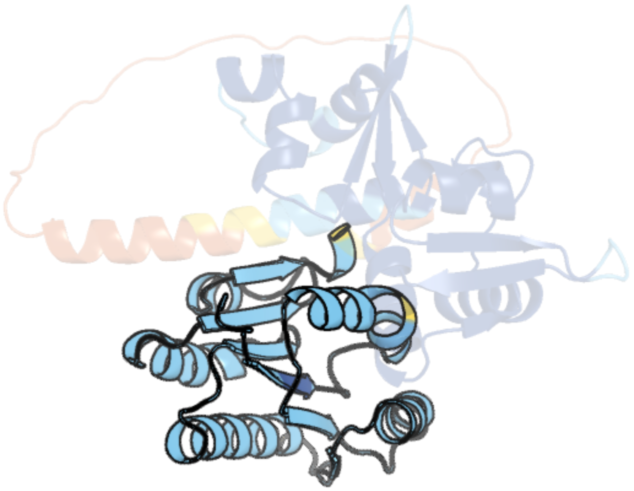

Figure 451: left: reference structure of 2qip chain A. right: predicted structure of chlorv-4..071, unaligned sequences are shown as transparent

#### chlorv-4..073

- Sequence-based annotation for chlorv-4..073 is
- No significant structural hit found

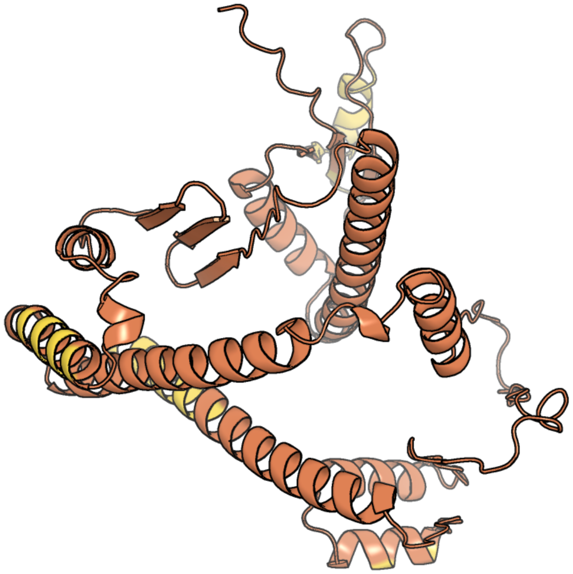

Figure 452: predicted structure of chlorv-4..073

#### chlorv-4..074

- Sequence-based annotation for chlorv-4..074 is
- No significant structural hit found

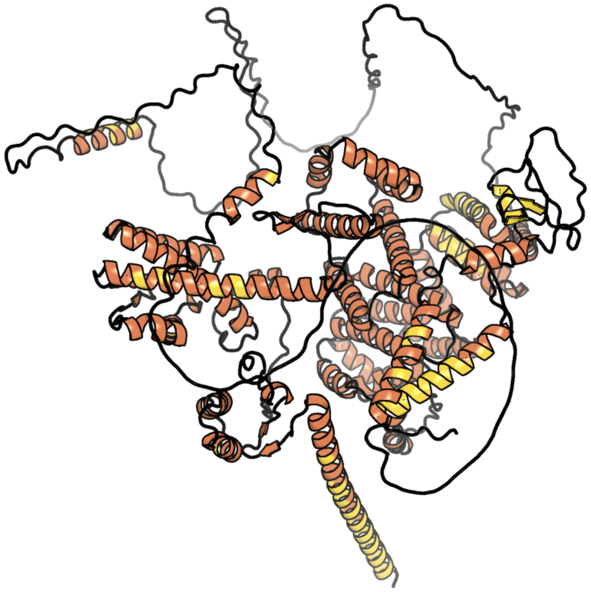

Figure 453: predicted structure of chlorv-4..074

#### chlorv-4..076

- Sequence-based annotation for chlorv-4..076 is
- Best hit was 3c48 chain B: Predicted glycosyltransferases

| target | prob | fident | alnlen | evalue | theader |
|---|---|---|---|---|---|
| 3c48-assembly1.cif.gz_B | 1 | 0.11 | 418 | 7.713e-10 | Structure of the retaining glycosyltransferase MshA: The first step in mycothiol biosynthesis. Organism: Corynebacterium glutamicum- APO (OPEN) structure. |
| 3c4q-assembly2.cif.gz_B | 1 | 0.115 | 426 | 8.149e-10 | Structure of the retaining glycosyltransferase MshA : The first step in mycothiol biosynthesis. Organism : Corynebacterium glutamicum- Complex with UDP |
| 5d00-assembly1.cif.gz_A | 1 | 0.107 | 401 | 8.61e-10 | Crystal structure of BshA from B. subtilis complexed with N-acetylglucosaminyl-malate and UMP |

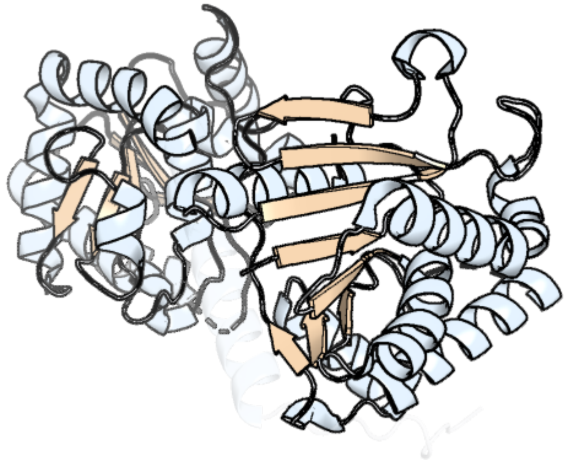

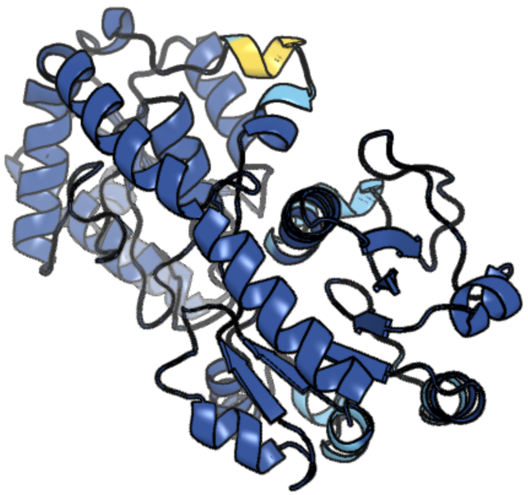

Figure 454: left: reference structure of 3c48 chain B. right: predicted structure of chlorv-4..076, unaligned sequences are shown as transparent

#### chlorv-4..080

- Sequence-based annotation for chlorv-4..080 is
- No significant structural hit found

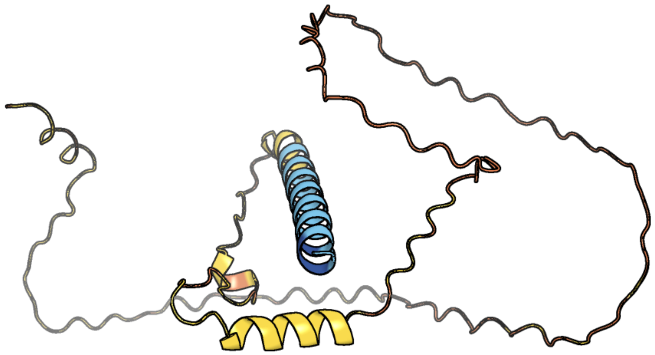

Figure 455: predicted structure of chlorv-4..080

#### chlorv-4..085

- Sequence-based annotation for chlorv-4..085 is
- No significant structural hit found

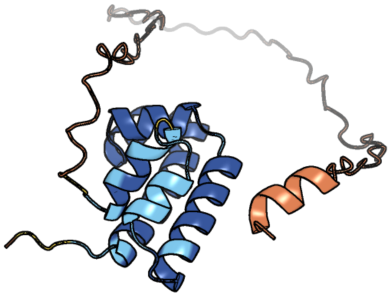

Figure 456: predicted structure of chlorv-4..085

#### chlorv-4..101

- Sequence-based annotation for chlorv-4..101 is
- No significant structural hit found

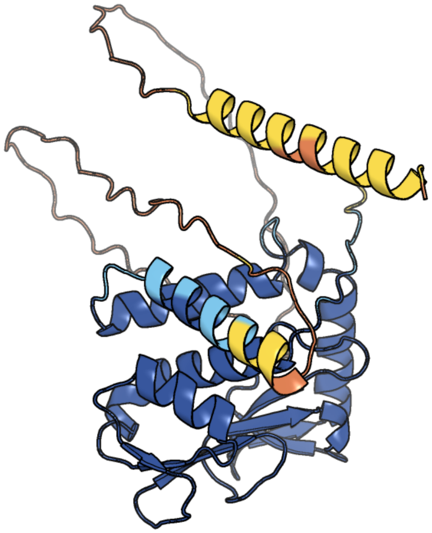

Figure 457: predicted structure of chlorv-4..101

#### chlorv-4..102

- Sequence-based annotation for chlorv-4..102 is
- No significant structural hit found

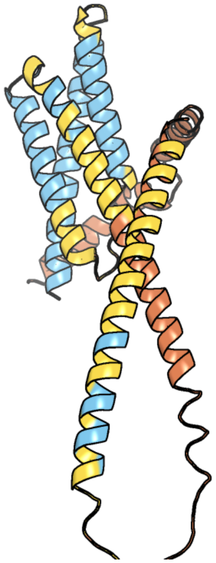

Figure 458: predicted structure of chlorv-4..102

#### chlorv-4..103

- Sequence-based annotation for chlorv-4..103 is
- No significant structural hit found

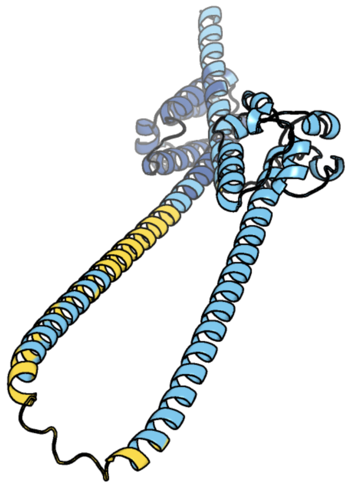

Figure 459: predicted structure of chlorv-4..103

#### chlorv-4..106

- Sequence-based annotation for chlorv-4..106 is
- No significant structural hit found

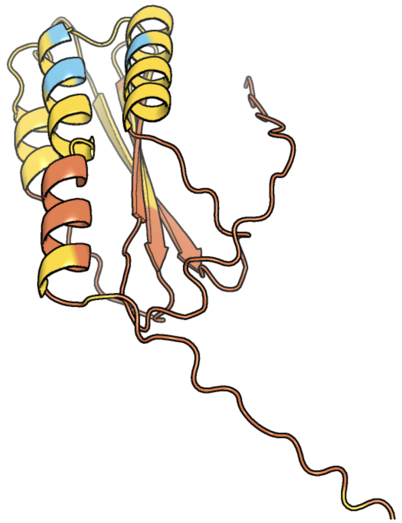

Figure 460: predicted structure of chlorv-4..106

#### chlorv-4..121

- Sequence-based annotation for chlorv-4..121 is
- No significant structural hit found

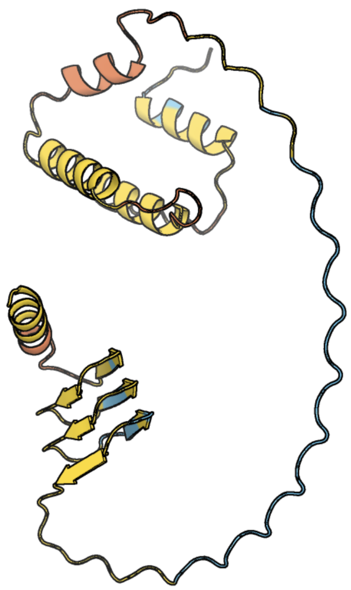

Figure 461: predicted structure of chlorv-4..121

#### chlorv-4..122

- Sequence-based annotation for chlorv-4..122 is
- No significant structural hit found

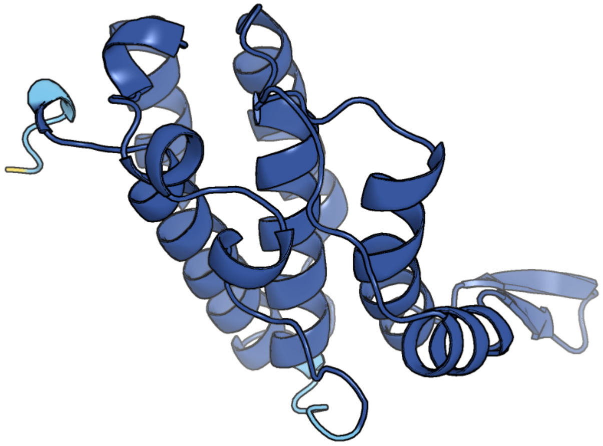

Figure 462: predicted structure of chlorv-4..122

#### chlorv-4..124

- Sequence-based annotation for chlorv-4..124 is
- No significant structural hit found

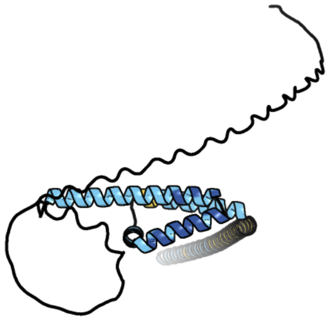

Figure 463: predicted structure of chlorv-4..124

#### chlorv-4..135

- Sequence-based annotation for chlorv-4..135 is
- No significant structural hit found

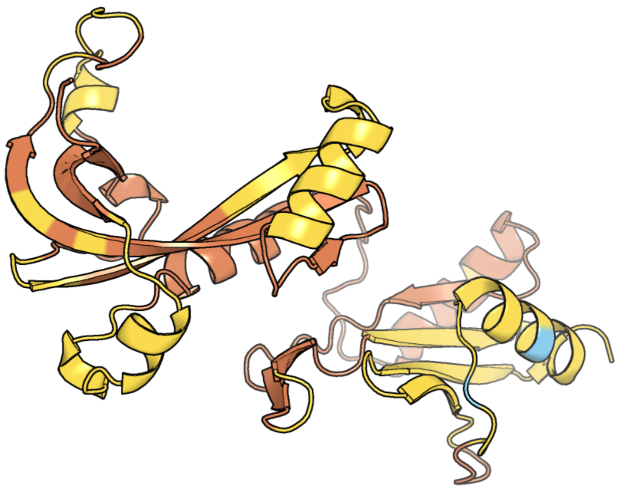

Figure 464: predicted structure of chlorv-4..135

### chlorv-4..148

- Sequence-based annotation for chlorv-4..148 is
- Best hit was 7z2c chain K: Kinesin-like protein

| target | prob | fident | alnlen | evalue | theader |
|---|---|---|---|---|---|
| 7z2c-assembly1.cif.gz_K | 1 | 0.157 | 363 | 1.324e-06 | P. falciparum kinesin-8B motor domain in no nucleotide bound to tubulin dimer |
| 7z2a-assembly1.cif.gz_K | 1 | 0.142 | 365 | 1.39e-06 | P. berghei kinesin-8B motor domain in no nucleotide state bound to tubulin dimer |
| 7tr0-assembly1.cif.gz_K | 1 | 0.144 | 332 | 3.171e-06 | CaKip3[2-436] - AMP-PNP in complex with a microtubule |

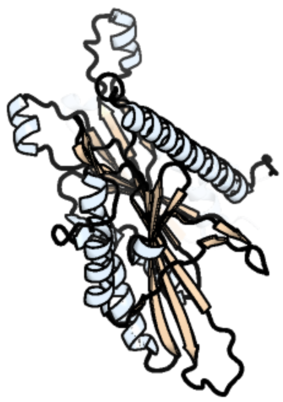

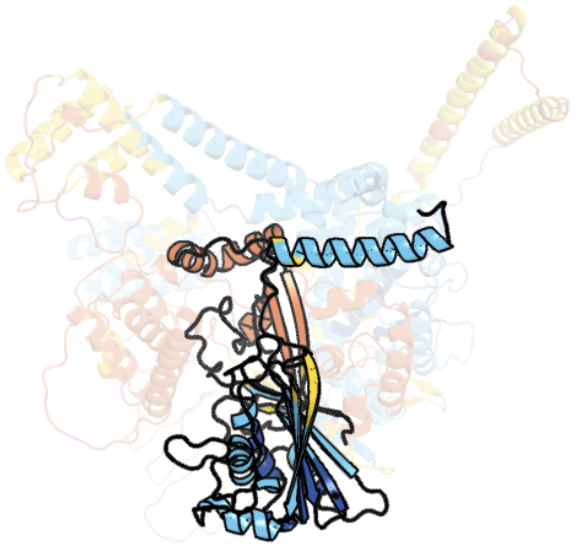

Figure 465: left: reference structure of 7z2c chain K. right: predicted structure of chlorv-4..148, unaligned sequences are shown as transparent

#### chlorv-4..151

- Sequence-based annotation for chlorv-4..151 is
- No significant structural hit found

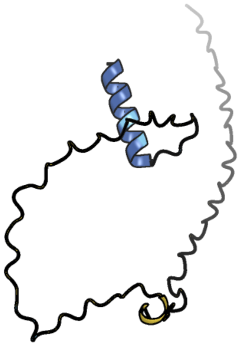

Figure 466: predicted structure of chlorv-4..151

#### chlorv-4..153

- Sequence-based annotation for chlorv-4..153 is
- No significant structural hit found

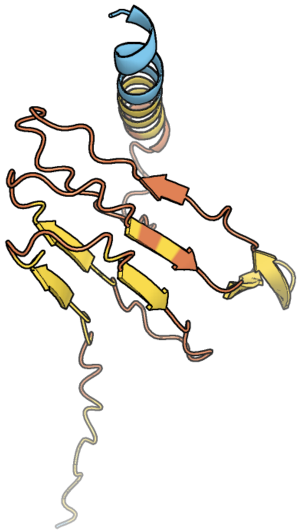

Figure 467: predicted structure of chlorv-4..153

#### chlorv-4..161

- Sequence-based annotation for chlorv-4..161 is
- No significant structural hit found

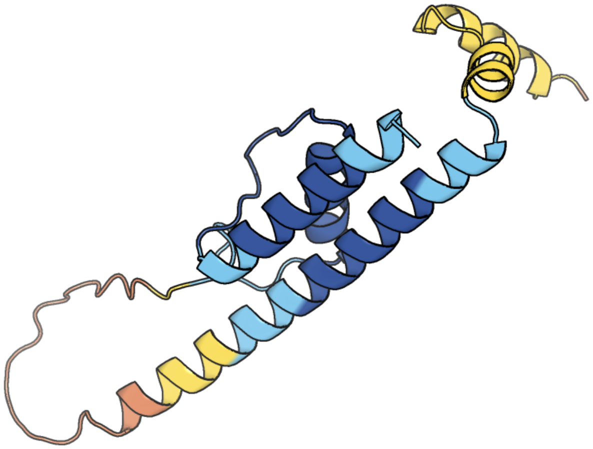

Figure 468: predicted structure of chlorv-4..161

#### chlorv-4..167

- Sequence-based annotation for chlorv-4..167 is
- No significant structural hit found

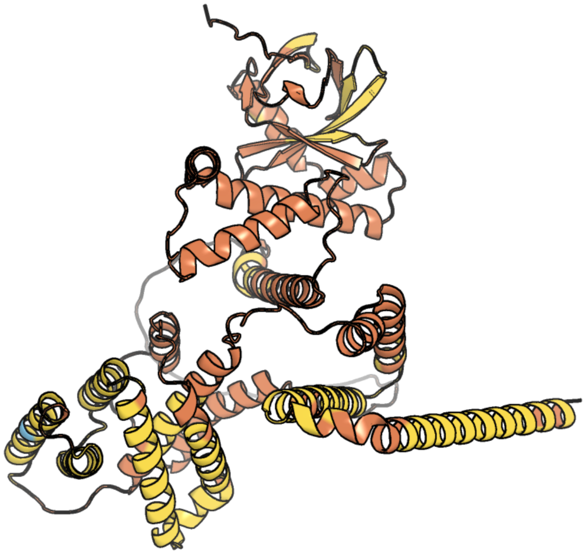

Figure 469: predicted structure of chlorv-4..167

#### chlorv-4..168

- Sequence-based annotation for chlorv-4..168 is
- No significant structural hit found

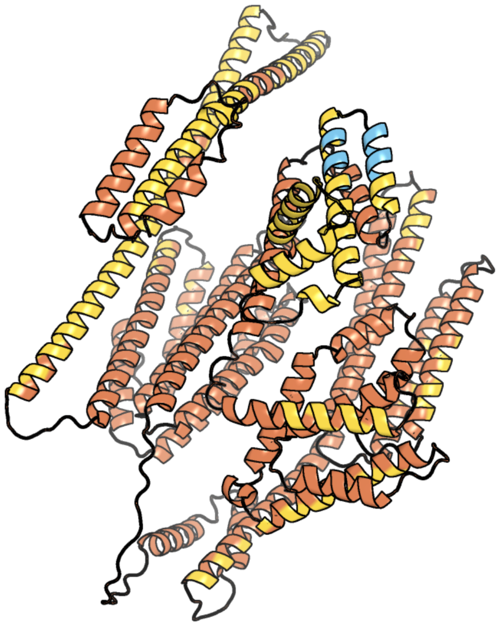

Figure 470: predicted structure of chlorv-4..168

#### chlorv-4..169

- Sequence-based annotation for chlorv-4..169 is
- No significant structural hit found

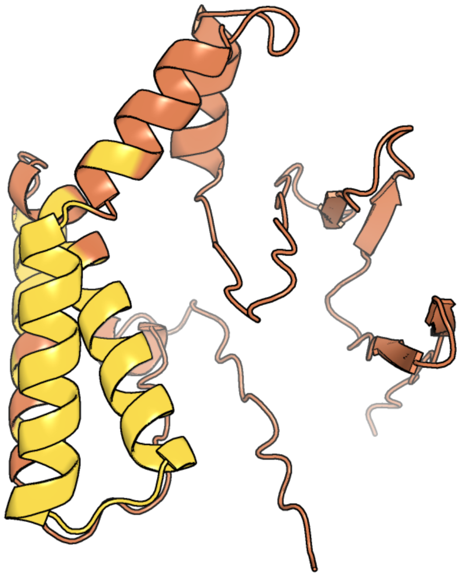

Figure 471: predicted structure of chlorv-4..169

#### chlorv-4..178

- Sequence-based annotation for chlorv-4..178 is
- No significant structural hit found

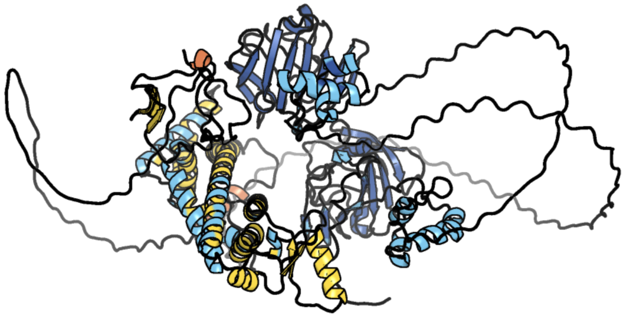

Figure 472: predicted structure of chlorv-4..178

#### chlorv-4..180

- Sequence-based annotation for chlorv-4..180 is
- No significant structural hit found

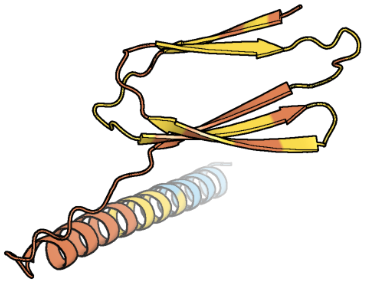

Figure 473: predicted structure of chlorv-4..180

#### chlorv-4..191

- Sequence-based annotation for chlorv-4..191 is
- No significant structural hit found

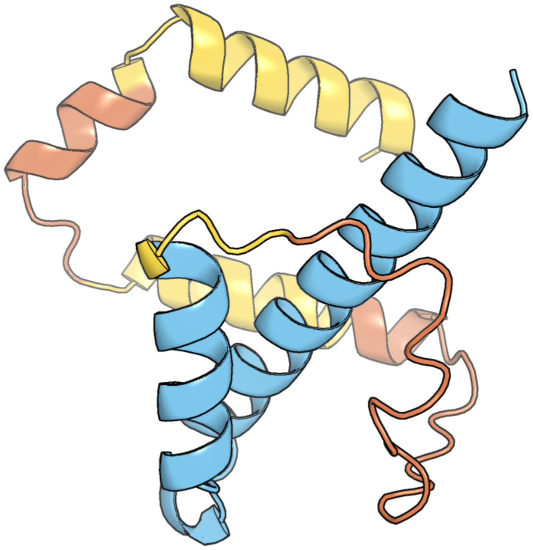

Figure 474: predicted structure of chlorv-4..191

#### chlorv-4..193

- Sequence-based annotation for chlorv-4..193 is
- No significant structural hit found

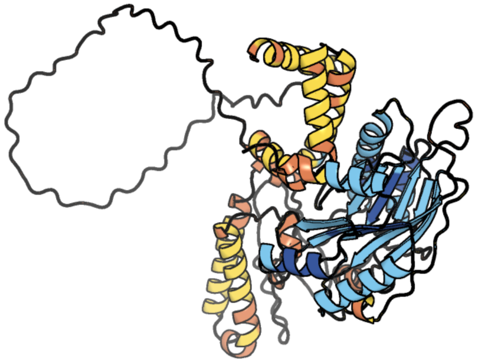

Figure 475: predicted structure of chlorv-4..193

#### chlorv-4..201

- Sequence-based annotation for chlorv-4..201 is
- Best hit was 1u7p chain D: magnesium-dependent phosphatase-1

| target | prob | fident | alnlen | evalue | theader |
|---|---|---|---|---|---|
| 1u7p-assembly4.cif.gz_D | 1 | 0.188 | 170 | 5.713e-05 | X-ray Crystal Structure of the Hypothetical Phosphotyrosine Phosphatase MDP-1 of the Haloacid Dehalogenase Superfamily |
| 2ah5-assembly1.cif.gz_A | 1 | 0.137 | 189 | 0.0002145 | Hydrolase, haloacid dehalogenase-like family protein SP0104 from Streptococcus pneumoniae |
| 3nrj-assembly3.cif.gz_I | 1 | 0.124 | 185 | 0.0002728 | Crystal structure of probable yrbi family phosphatase from pseudomonas syringae pv.phaseolica 1448a complexed with magnesium |

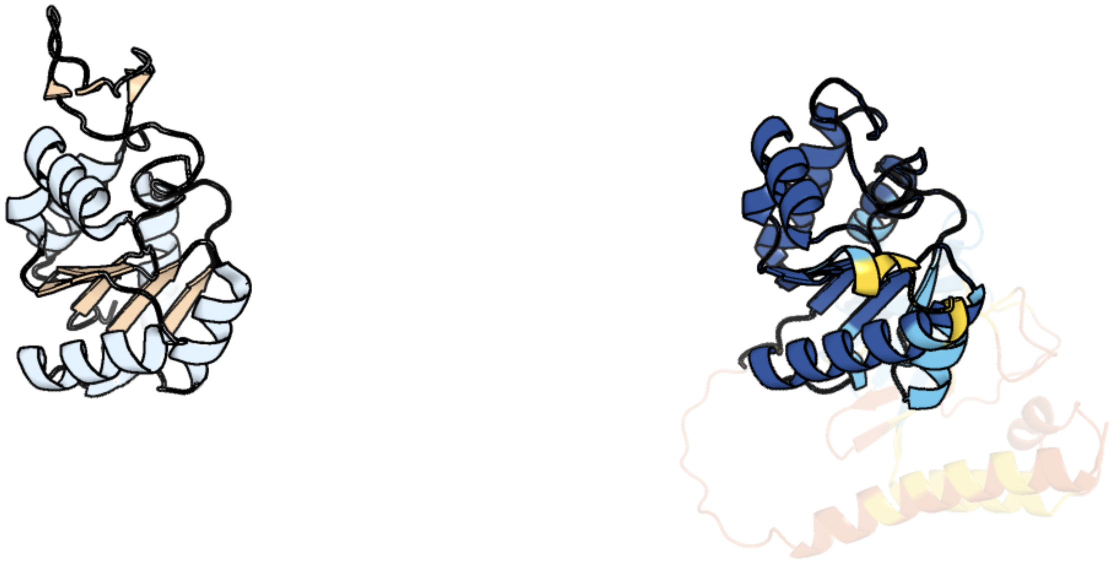

Figure 476: left: reference structure of 1u7p chain D. right: predicted structure of chlorv-4..201, unaligned sequences are shown as transparent

#### chlorv-4..203

- Sequence-based annotation for chlorv-4..203 is
- Best hit was 1k9e chain A: alpha-D-glucuronidase

| target | prob | fident | alnlen | evalue | theader |
|---|---|---|---|---|---|
| 1k9e-assembly1.cif.gz_A | 1 | 0.105 | 841 | 3.023e-14 | Crystal structure of a mutated family-67 alpha-D-glucuronidase (E285N) from Bacillus stearothermophilus T-6, complexed with 4-O-methyl-glucuronic acid |
| 6hze-assembly2.cif.gz_B | 1 | 0.095 | 1008 | 3.53e-14 | BP0997, GH138 enzyme targeting pectin rhamnogalacturonan II |
| 1mqr-assembly1.cif.gz_A-2 | 1 | 0.11 | 838 | 1.669e-13 | THE CRYSTAL STRUCTURE OF ALPHA-D-GLUCURONIDASE (E386Q) FROM BACILLUS STEAROTHERMOPHILUS T-6 |

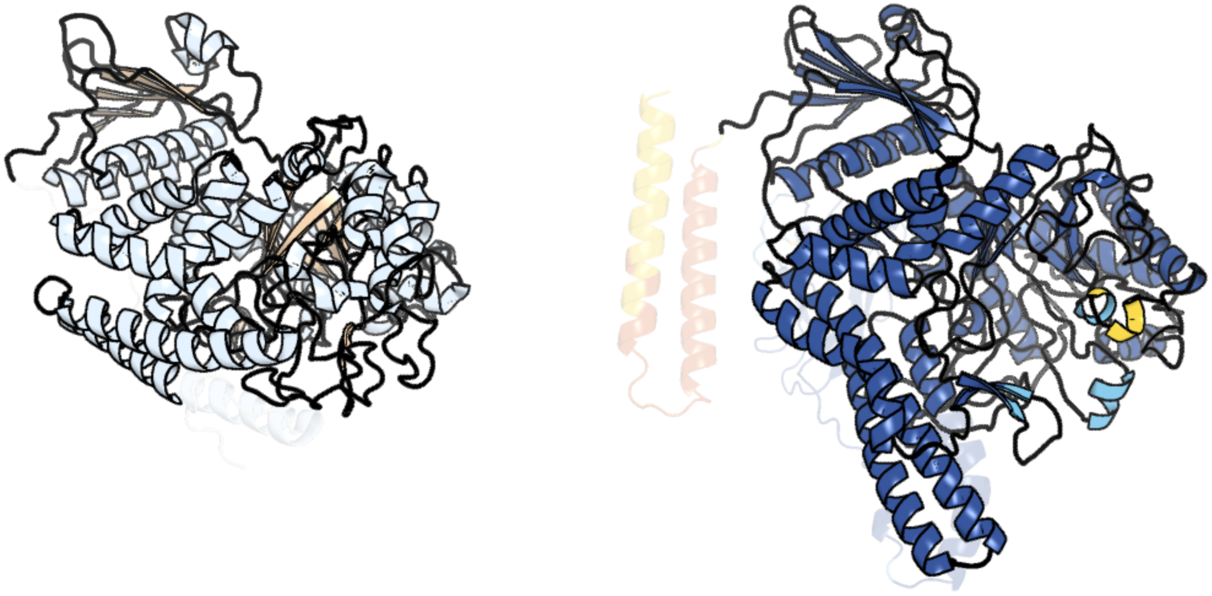

Figure 477: left: reference structure of 1k9e chain A. right: predicted structure of chlorv-4..203, unaligned sequences are shown as transparent

#### chlorv-4..204

- Sequence-based annotation for chlorv-4..204 is
- Best hit was 3kmj chain A: A612L protein

| target | prob | fident | alnlen | evalue | theader |
|---|---|---|---|---|---|
| 3kmj-assembly1.cif.gz_A | 1 | 0.31 | 116 | 3.202e-10 | Crystal structure of vSET under condition B |
| 4rz0-assembly1.cif.gz_A | 1 | 0.353 | 113 | 3.387e-10 | Crystal Structure of Plasmodium falciparum putative histone methyltransferase PFL0690c |
| 3kma-assembly1.cif.gz_B | 1 | 0.283 | 106 | 3.18e-09 | Crystal Structure of vSET under Condition A |

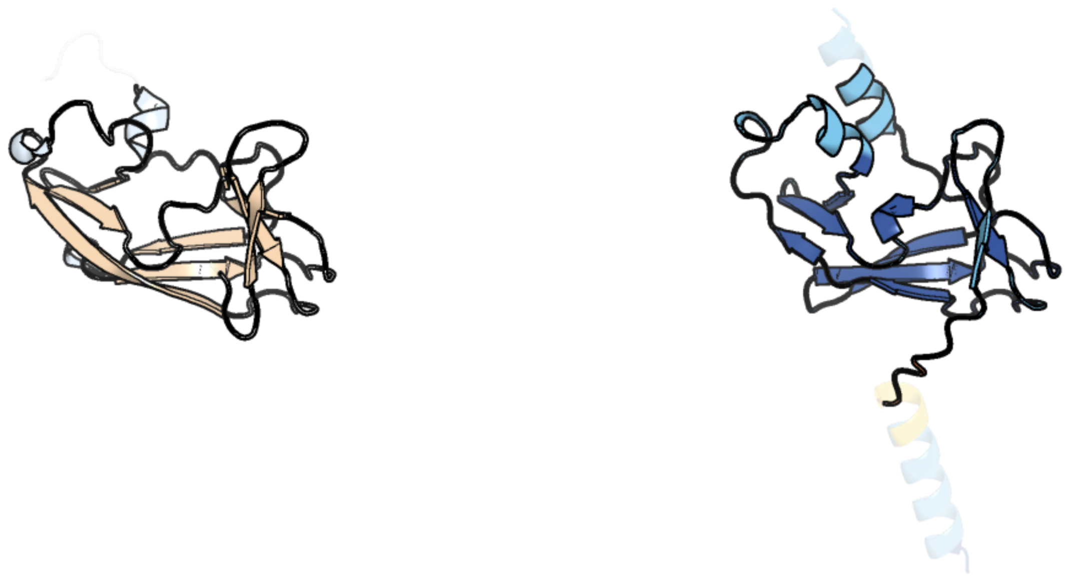

Figure 478: left: reference structure of 3kmj chain A. right: predicted structure of chlorv-4..204, unaligned sequences are shown as transparent

#### chlorv-4..212

- Sequence-based annotation for chlorv-4..212 is
- No significant structural hit found

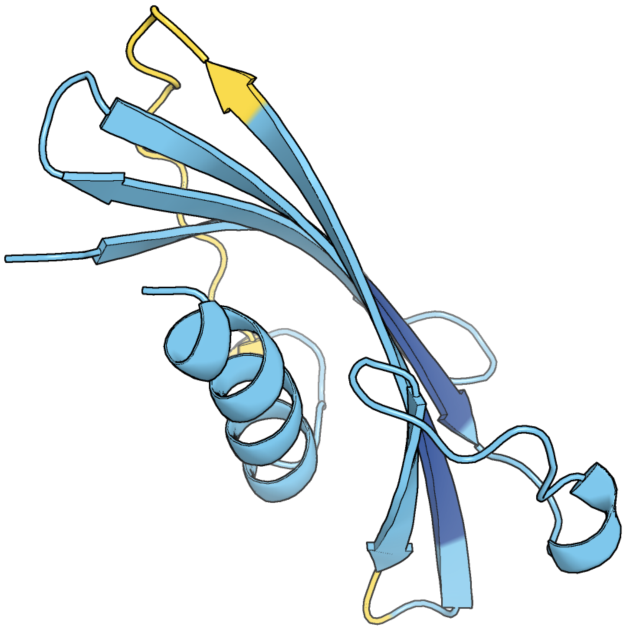

Figure 479: predicted structure of chlorv-4..212

#### chlorv-4..218

- Sequence-based annotation for chlorv-4..218 is
- No significant structural hit found

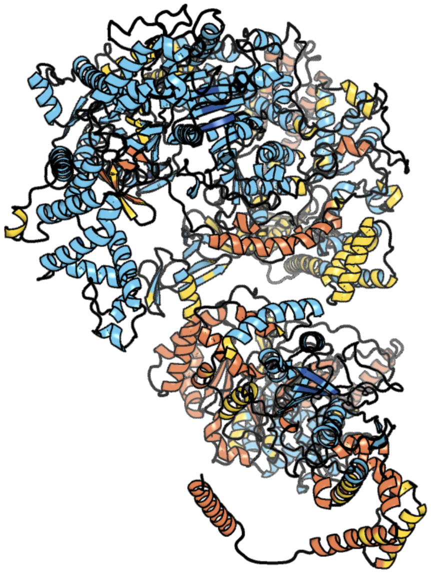

Figure 480: predicted structure of chlorv-4..218

#### chlorv-4..221

- Sequence-based annotation for chlorv-4..221 is
- Best hit was 7sqc chain 6B: FAP15

| target | prob | fident | alnlen | evalue | theader |
|---|---|---|---|---|---|
| 7sqc-assembly1.cif.gz_6B | 1 | 0.172 | 307 | 8.09e-13 | Ciliary C1 central pair apparatus isolated from Chlamydomonas reinhardtii |
| 1s95-assembly2.cif.gz_B | 1 | 0.179 | 317 | 1.009e-12 | Structure of serine/threonine protein phosphatase 5 |
| 7tvf-assembly2.cif.gz_C | 1 | 0.169 | 348 | 1.066e-12 | Crystal structure of the SHOC2-MRAS-PP1CA (SMP) complex to a resolution of 2.17 Angstrom |

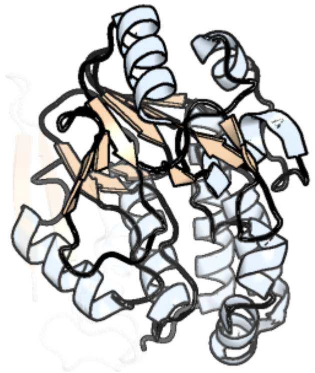

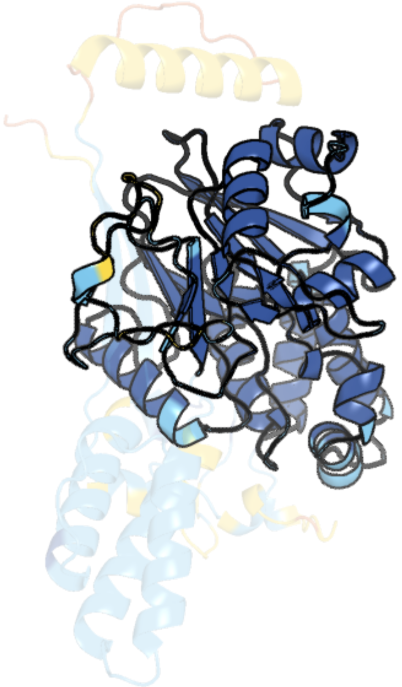

Figure 481: left: reference structure of 7sqc chain 6B. right: predicted structure of chlorv-4..221, unaligned sequences are shown as transparent

#### chlorv-4..244

- Sequence-based annotation for chlorv-4..244 is
- No significant structural hit found

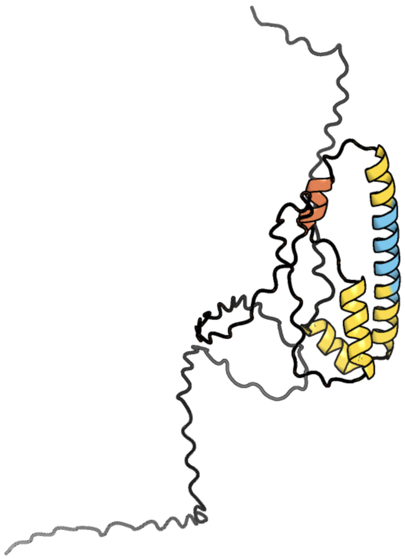

Figure 482: predicted structure of chlorv-4..244

#### chlorv-4..248

- Sequence-based annotation for chlorv-4..248 is
- No significant structural hit found

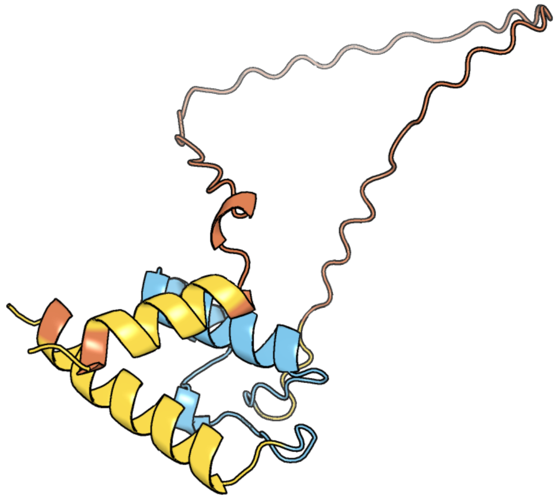

Figure 483: predicted structure of chlorv-4..248

#### chlorv-4..249

- Sequence-based annotation for chlorv-4..249 is
- No significant structural hit found

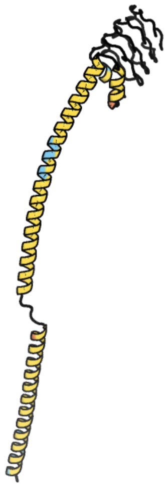

Figure 484: predicted structure of chlorv-4..249

#### chlorv-4..250

- Sequence-based annotation for chlorv-4..250 is
- No significant structural hit found

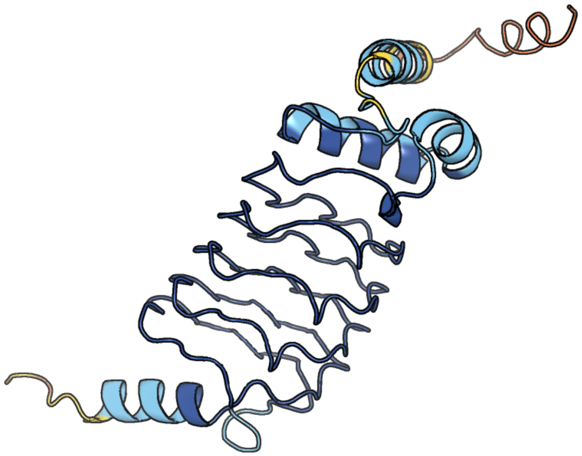

Figure 485: predicted structure of chlorv-4..250

#### chlorv-4..255

- Sequence-based annotation for chlorv-4..255 is
- No significant structural hit found

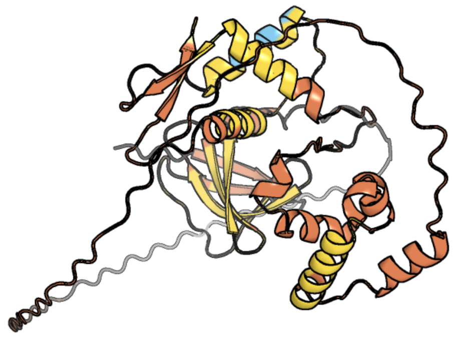

Figure 486: predicted structure of chlorv-4..255

#### chlorv-4..256

- Sequence-based annotation for chlorv-4..256 is
- Best hit was 5oxf chain A: GTP-binding protein

| target | prob | fident | alnlen | evalue | theader |
|---|---|---|---|---|---|
| 5oxf-assembly1.cif.gz_A | 1 | 0.179 | 567 | 7.922e-08 | An oligomerised bacterial dynamin pair provides a mechanism for the long range sensing and tethering of membranes |
| 5oxf-assembly1.cif.gz_B | 1 | 0.175 | 502 | 2.47e-06 | An oligomerised bacterial dynamin pair provides a mechanism for the long range sensing and tethering of membranes |
| 6jfm-assembly2.cif.gz_B | 1 | 0.095 | 440 | 3.089e-06 | Mitofusin2 (MFN2)_T111D |

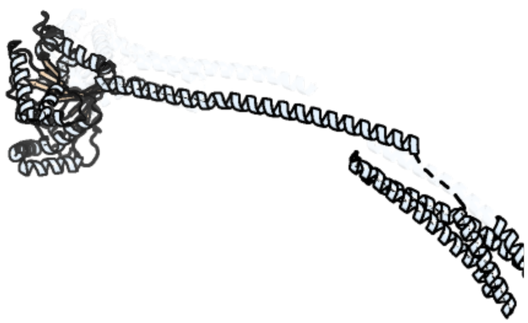

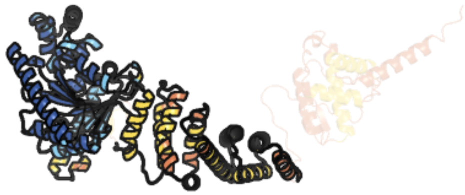

Figure 487: left: reference structure of 5oxf chain A. right: predicted structure of chlorv-4..256, unaligned sequences are shown as transparent

#### chlorv-4..263

- Sequence-based annotation for chlorv-4..263 is
- No significant structural hit found

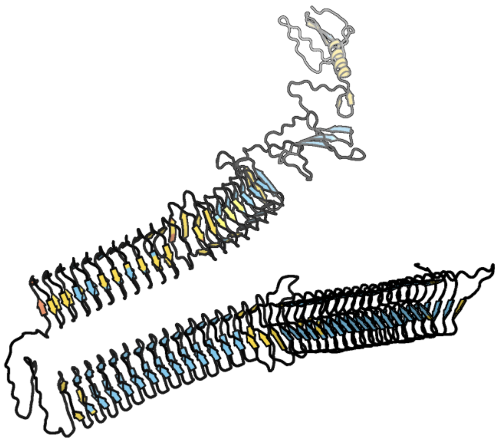

Figure 488: predicted structure of chlorv-4..263

### chlorv-4..272

- Sequence-based annotation for chlorv-4..272 is
- No significant structural hit found

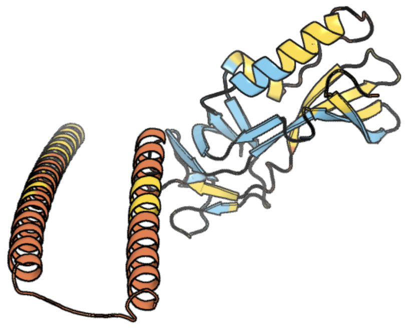

Figure 489: predicted structure of chlorv-4..272

#### chlorv-4..277

- Sequence-based annotation for chlorv-4..277 is
- Best hit was 2cdq chain B: ASPARTOKINASE

| target | prob | fident | alnlen | evalue | theader |
|---|---|---|---|---|---|
| 2cdq-assembly1.cif.gz_B | 1 | 0.202 | 475 | 2.093e-28 | Crystal structure of Arabidopsis thaliana aspartate kinase complexed with lysine and S-adenosylmethionine |
| 3c1n-assembly3.cif.gz_D | 1 | 0.255 | 470 | 2.093e-28 | Crystal Structure of Allosteric Inhibition Threonine-sensitive Aspartokinase from Methanococcus jannaschii with L-threonine |
| 3c1n-assembly2.cif.gz_B | 1 | 0.236 | 477 | 3.895e-28 | Crystal Structure of Allosteric Inhibition Threonine-sensitive Aspartokinase from Methanococcus jannaschii with L-threonine |

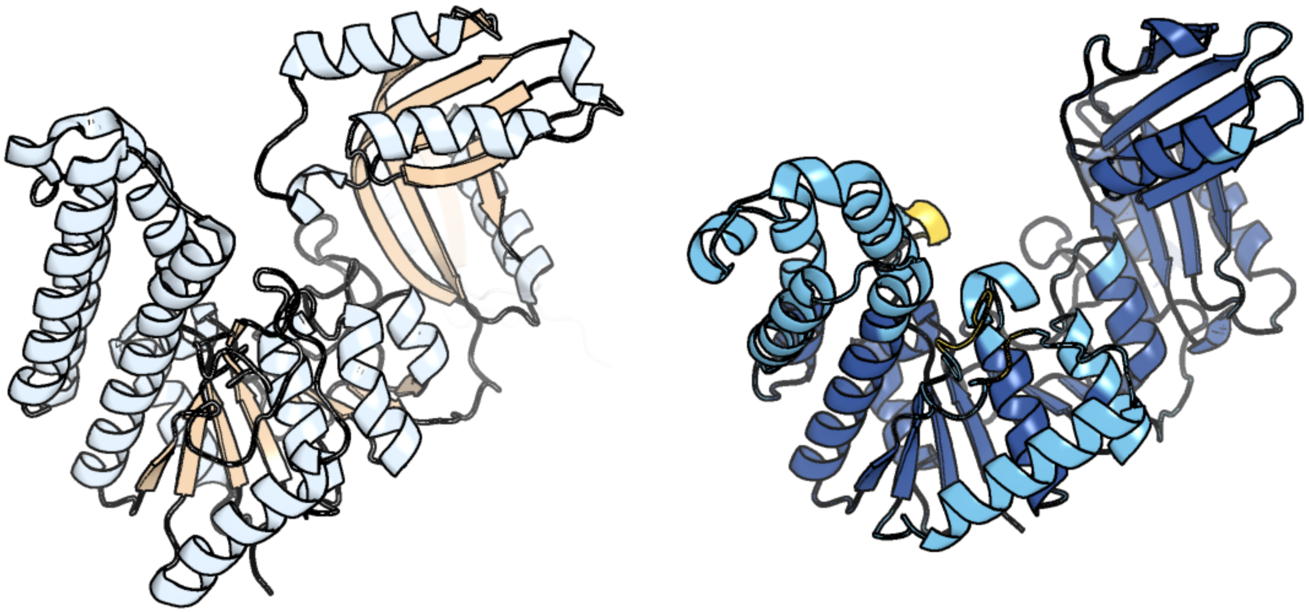

Figure 490: left: reference structure of 2cdq chain B. right: predicted structure of chlorv-4..277, unaligned sequences are shown as transparent

#### chlorv-4..282

- Sequence-based annotation for chlorv-4..282 is
- No significant structural hit found

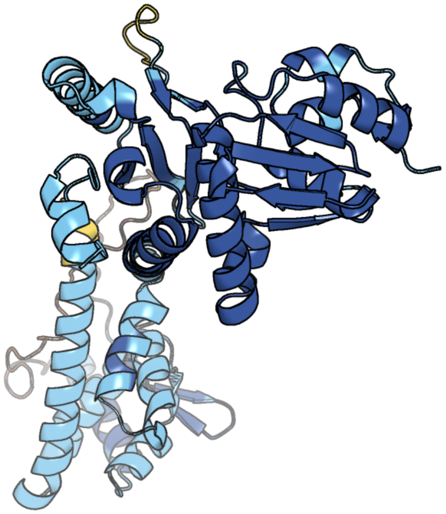

Figure 491: predicted structure of chlorv-4..282

#### chlorv-4..283

- Sequence-based annotation for chlorv-4..283 is
- No significant structural hit found

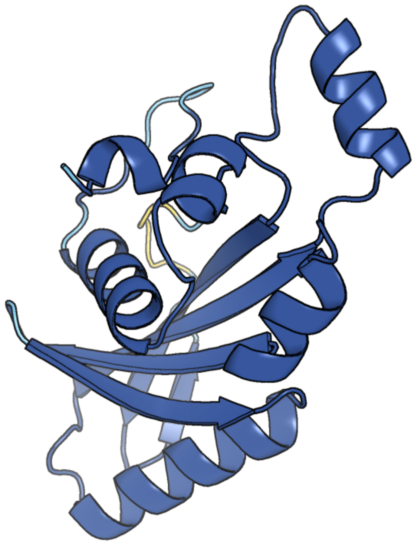

Figure 492: predicted structure of chlorv-4..283

### chlorv-4..292

- Sequence-based annotation for chlorv-4..292 is
- No significant structural hit found

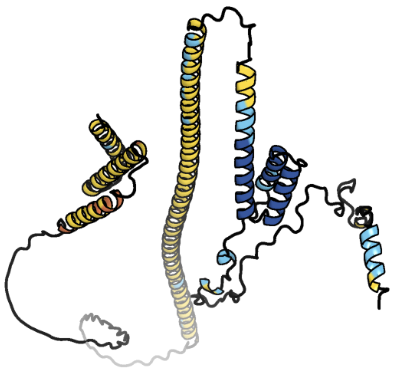

Figure 493: predicted structure of chlorv-4..292

### chlorv-4..310

- Sequence-based annotation for chlorv-4..310 is
- Best hit was 5djs chain A: Tetratricopeptide TPR_2 repeat protein

| target | prob | fident | alnlen | evalue | theader |
|---|---|---|---|---|---|
| 5djs-assembly1.cif.gz_A | 1 | 0.132 | 477 | 1.908e-17 | Thermobaculum terrenum O-GlcNAc transferase mutant - K341M |
| 8dth-assembly1.cif.gz_B | 1 | 0.164 | 469 | 2.19e-17 | Cryo-EM structure of Arabidopsis SPY alternative conformation 2 |
| 4gyw-assembly1.cif.gz_C | 1 | 0.126 | 600 | 7.921e-17 | Crystal structure of human O-GlcNAc Transferase in complex with UDP and a glycopeptide |

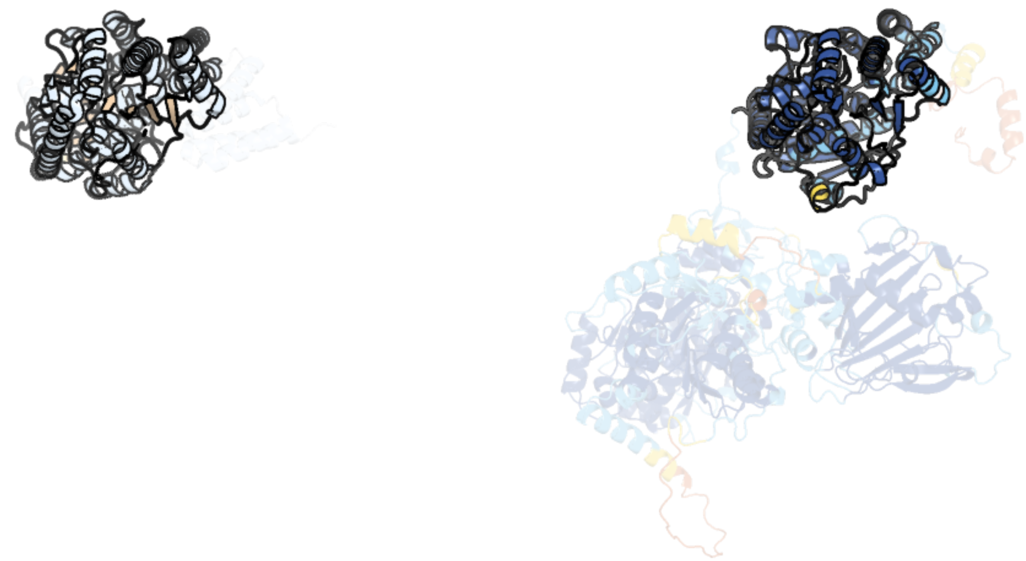

Figure 494: left: reference structure of 5djs chain A. right: predicted structure of chlorv-4..310, unaligned sequences are shown as transparent

#### chlorv-4..316

- Sequence-based annotation for chlorv-4..316 is
- No significant structural hit found

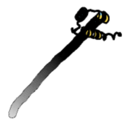

Figure 495: predicted structure of chlorv-4..316

#### chlorv-4..319

- Sequence-based annotation for chlorv-4..319 is
- No significant structural hit found

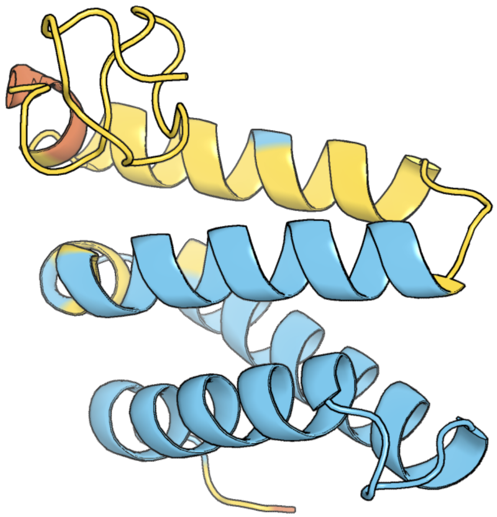

Figure 496: predicted structure of chlorv-4..319

#### chlorv-4..328

- Sequence-based annotation for chlorv-4..328 is
- No significant structural hit found

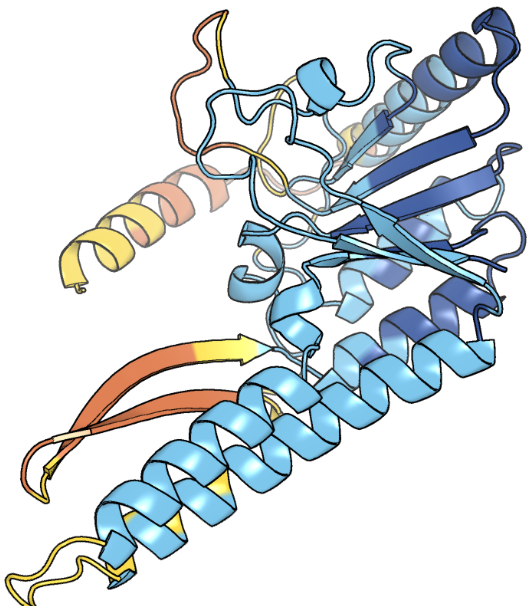

Figure 497: predicted structure of chlorv-4..328

#### chlorv-4..331

- Sequence-based annotation for chlorv-4..331 is
- No significant structural hit found

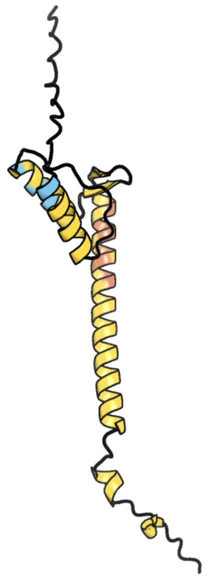

Figure 498: predicted structure of chlorv-4..331

#### chlorv-4..341

- Sequence-based annotation for chlorv-4..341 is
- No significant structural hit found

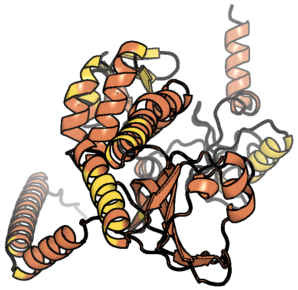

Figure 499: predicted structure of chlorv-4..341

### chlorv-4..354

- Sequence-based annotation for chlorv-4..354 is
- No significant structural hit found

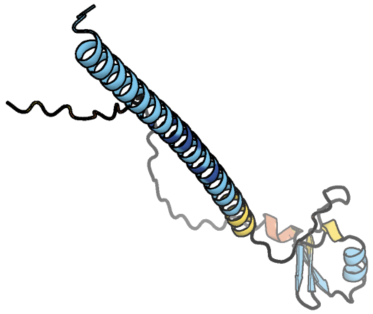

Figure 500: predicted structure of chlorv-4..354

#### chlorv-4..381

- Sequence-based annotation for chlorv-4..381 is
- No significant structural hit found

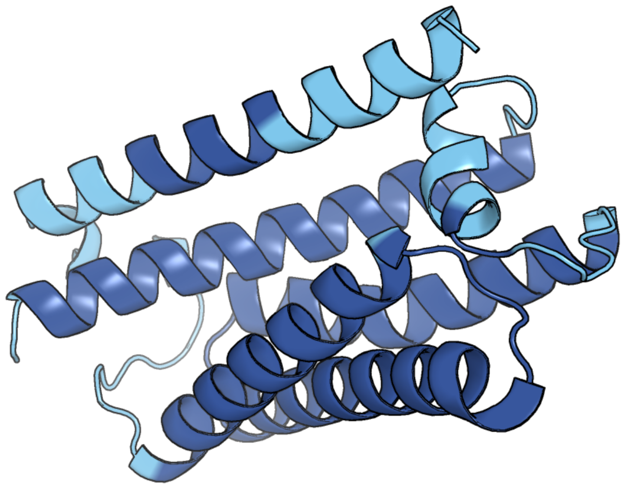

Figure 501: predicted structure of chlorv-4..381

#### chlorv-4..390

- Sequence-based annotation for chlorv-4..390 is
- No significant structural hit found

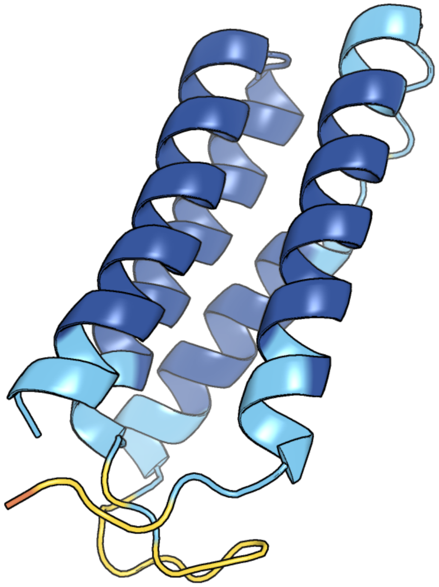

Figure 502: predicted structure of chlorv-4..390

### chlorv-4..391

- Sequence-based annotation for chlorv-4..391 is
- Best hit was 3oy7 chain B: Glycosyltransferase B736L

| target | prob | fident | alnlen | evalue | theader |
|---|---|---|---|---|---|
| 3oy7-assembly1.cif.gz_B | 1 | 0.129 | 531 | 5.98e-19 | Crystal structure of a virus encoded glycosyltransferase in complex with GDP-mannose |
| 7mi0-assembly1.cif.gz_A | 1 | 0.138 | 355 | 1.336e-12 | Crystal Structure of Glycosyltransferase from Rickettsia africae ESF-5 |
| 3l01-assembly2.cif.gz_B | 1 | 0.103 | 404 | 6.223e-12 | Crystal structure of monomeric glycogen synthase from Pyrococcus abyssi |

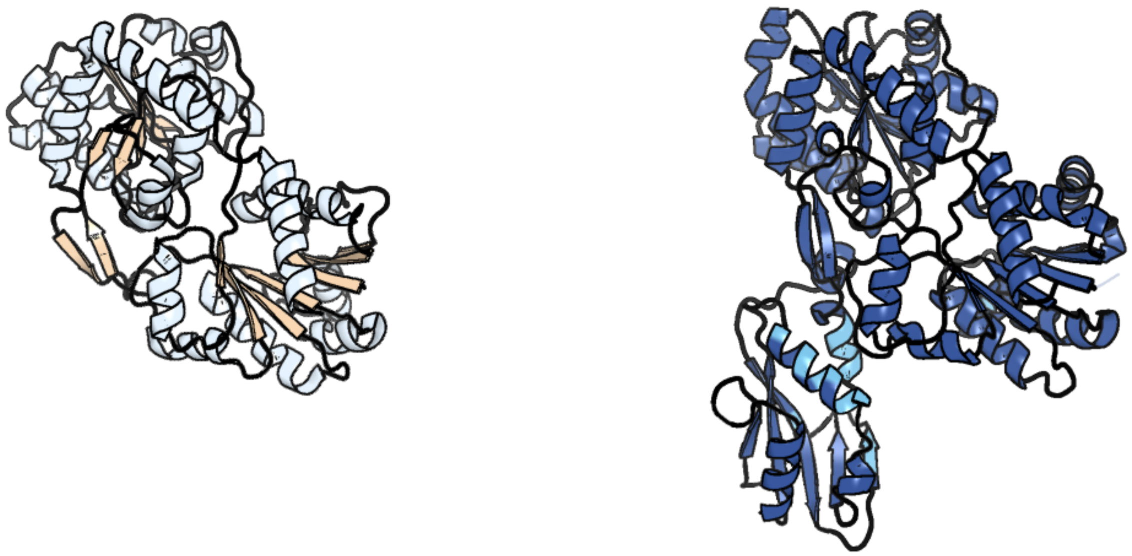

Figure 503: left: reference structure of 3oy7 chain B. right: predicted structure of chlorv-4..391, unaligned sequences are shown as transparent

#### chlorv-4..393

- Sequence-based annotation for chlorv-4..393 is
- No significant structural hit found

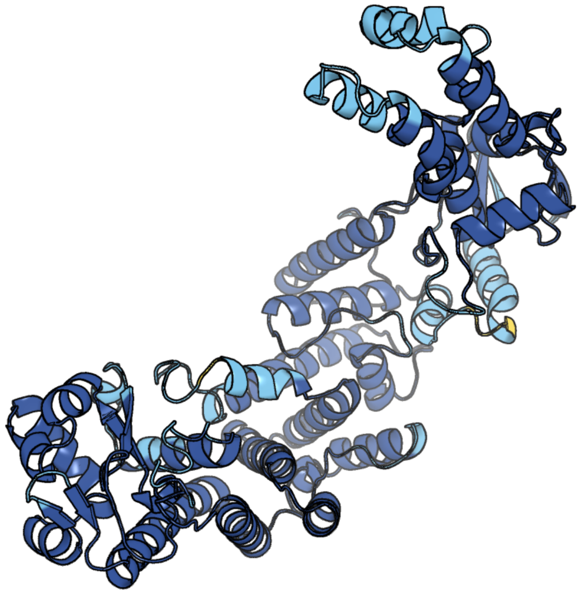

Figure 504: predicted structure of chlorv-4..393

#### chlorv-4..394

- Sequence-based annotation for chlorv-4..394 is
- Best hit was 5l9b chain A: Egl nine homolog 1

| target | prob | fident | alnlen | evalue | theader |
|---|---|---|---|---|---|
| 5l9b-assembly1.cif.gz_A | 1 | 0.128 | 210 | 1.335e-06 | HIF PROLYL HYDROXYLASE 2 (PHD2/ EGLN1) IN COMPLEX WITH 2-OXOGLUTARATE (2OG) AND HIF-1ALPHA CODD (556-574) |
| 6f0w-assembly1.cif.gz_A | 1 | 0.12 | 216 | 1.577e-06 | prolyl hydroxylase in complex with hypoxia inducible factor oxygen degradation domain peptide fragment from Trichoplax adhaerens |
| 4j25-assembly7.cif.gz_G | 1 | 0.095 | 209 | 1.667e-06 | Crystal structure of a Pseudomonas putida prolyl-4-hydroxylase (P4H) |

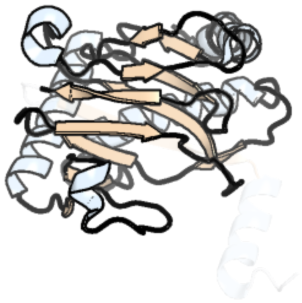

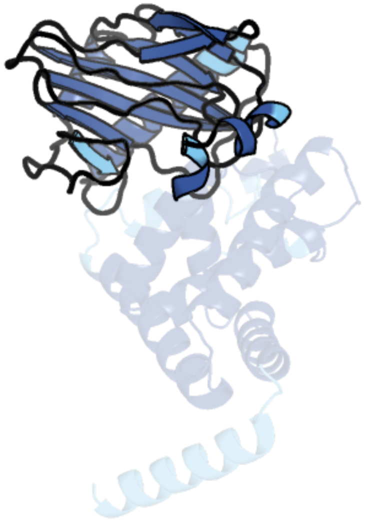

Figure 505: left: reference structure of 5l9b chain A. right: predicted structure of chlorv-4..394, unaligned sequences are shown as transparent
