## Supplementary figures and images for "Strain-level diversity of giant viruses infecting chlorarachniophyte algae in the subtropical North Pacific"

### Supplementary_File1_single_gene_trees.pdf

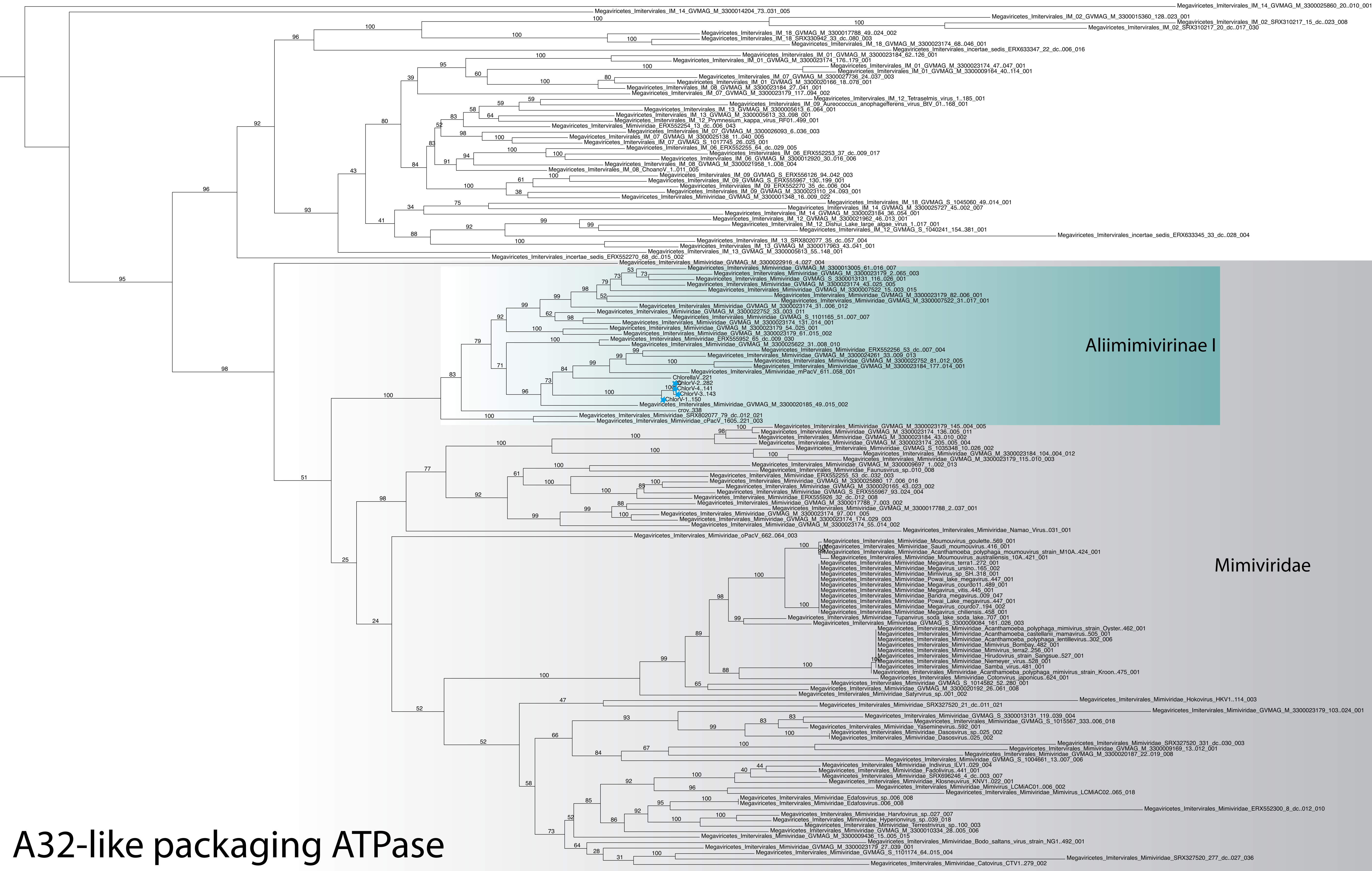


A32-like packaging ATPase

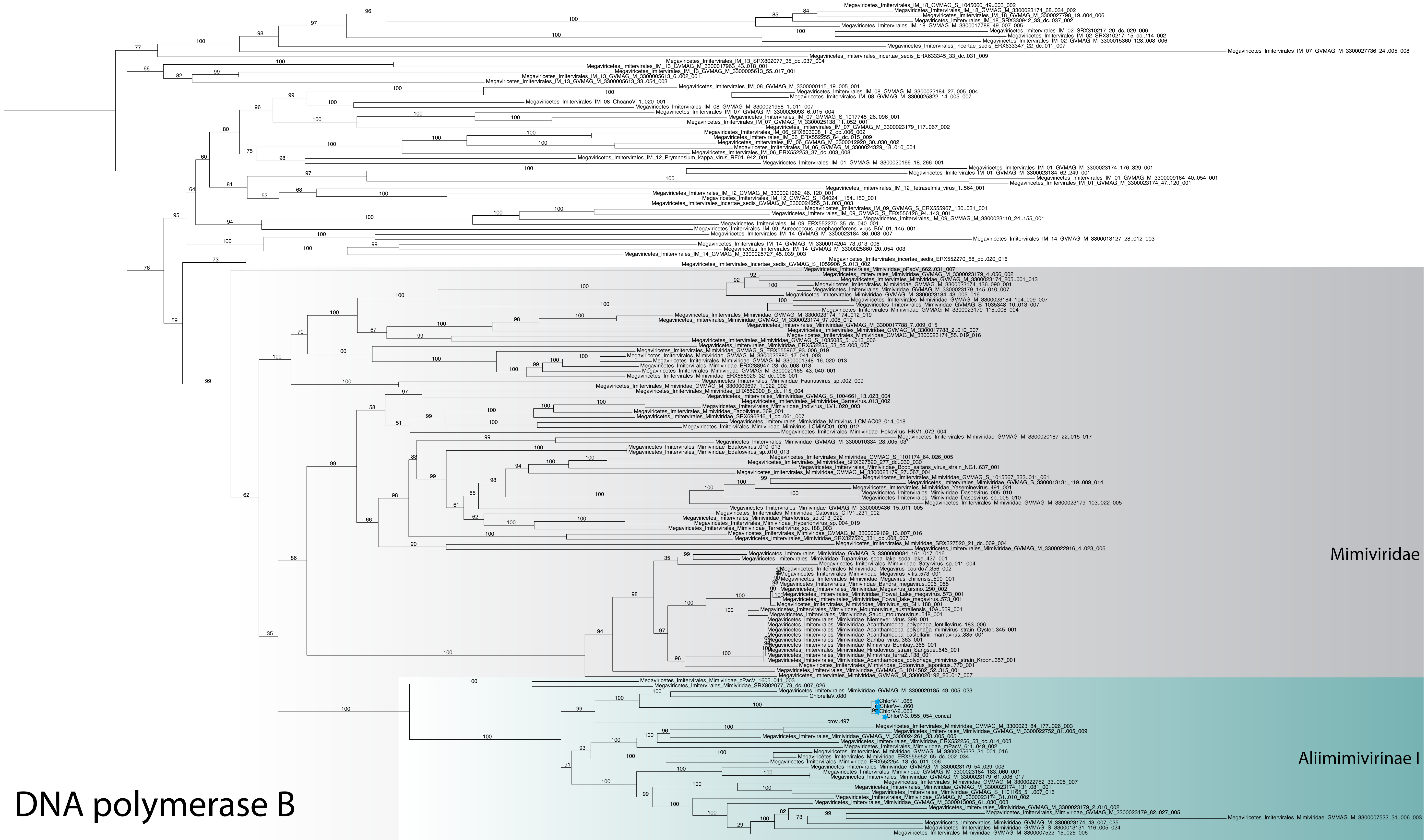

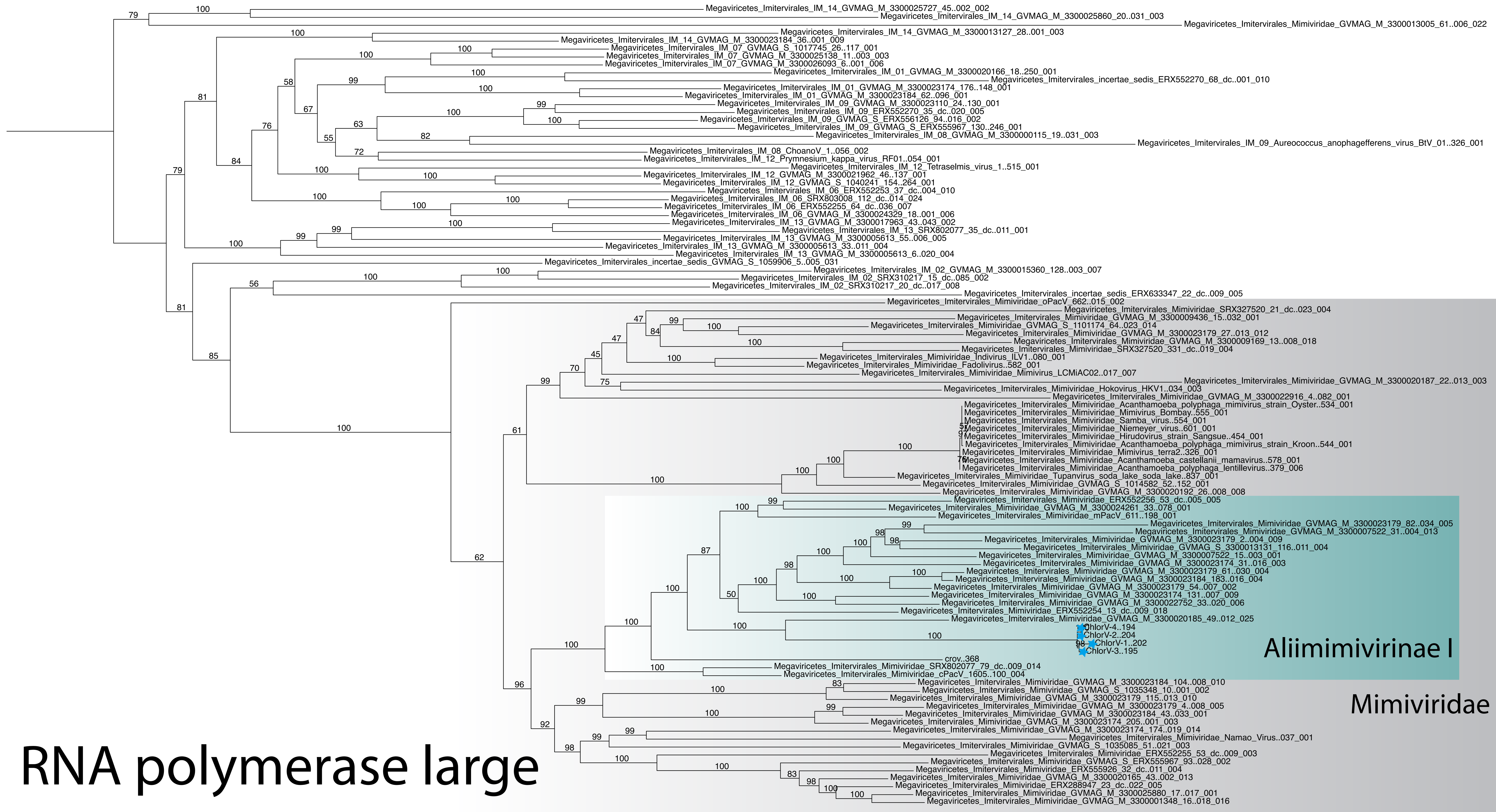

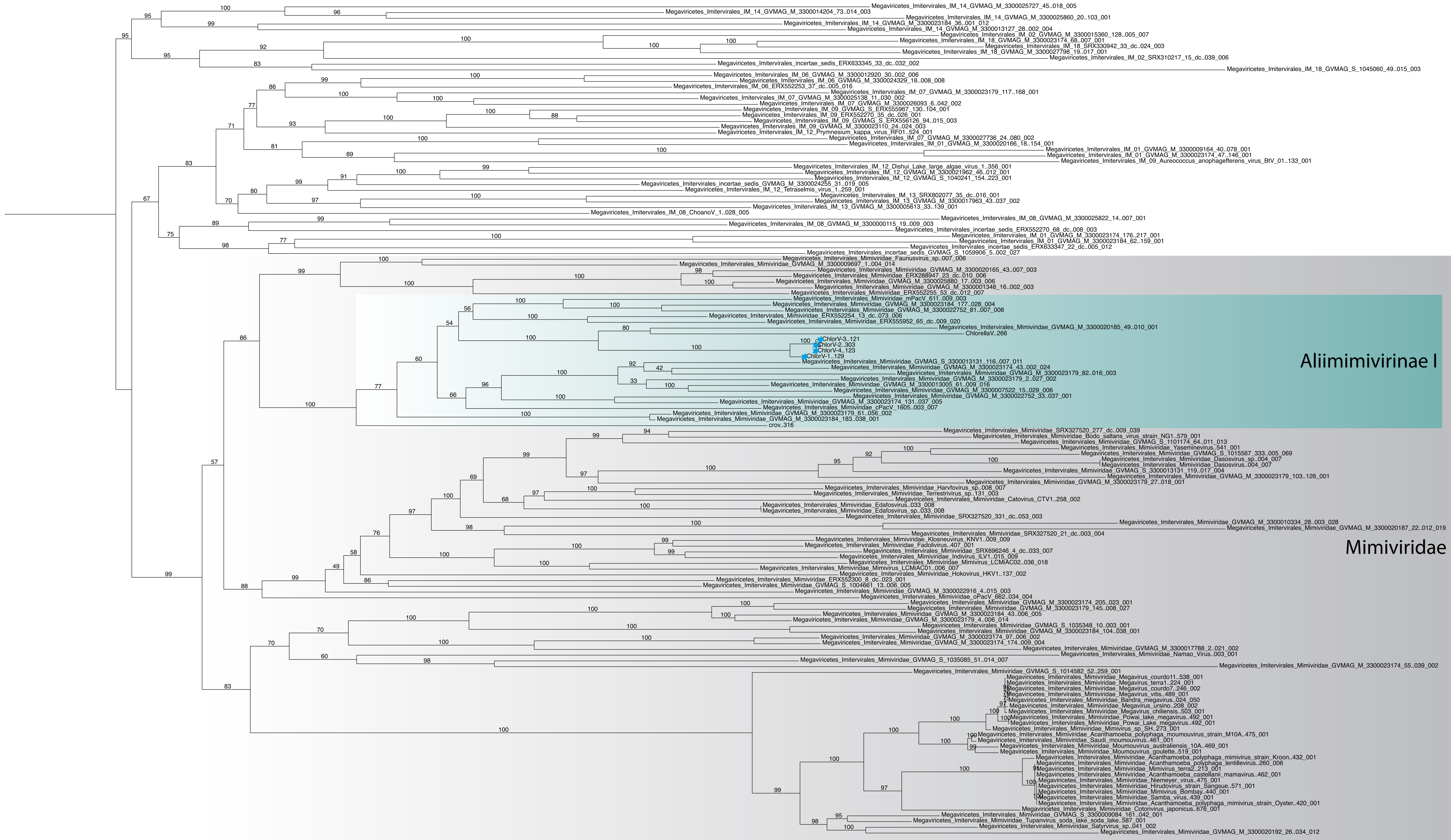

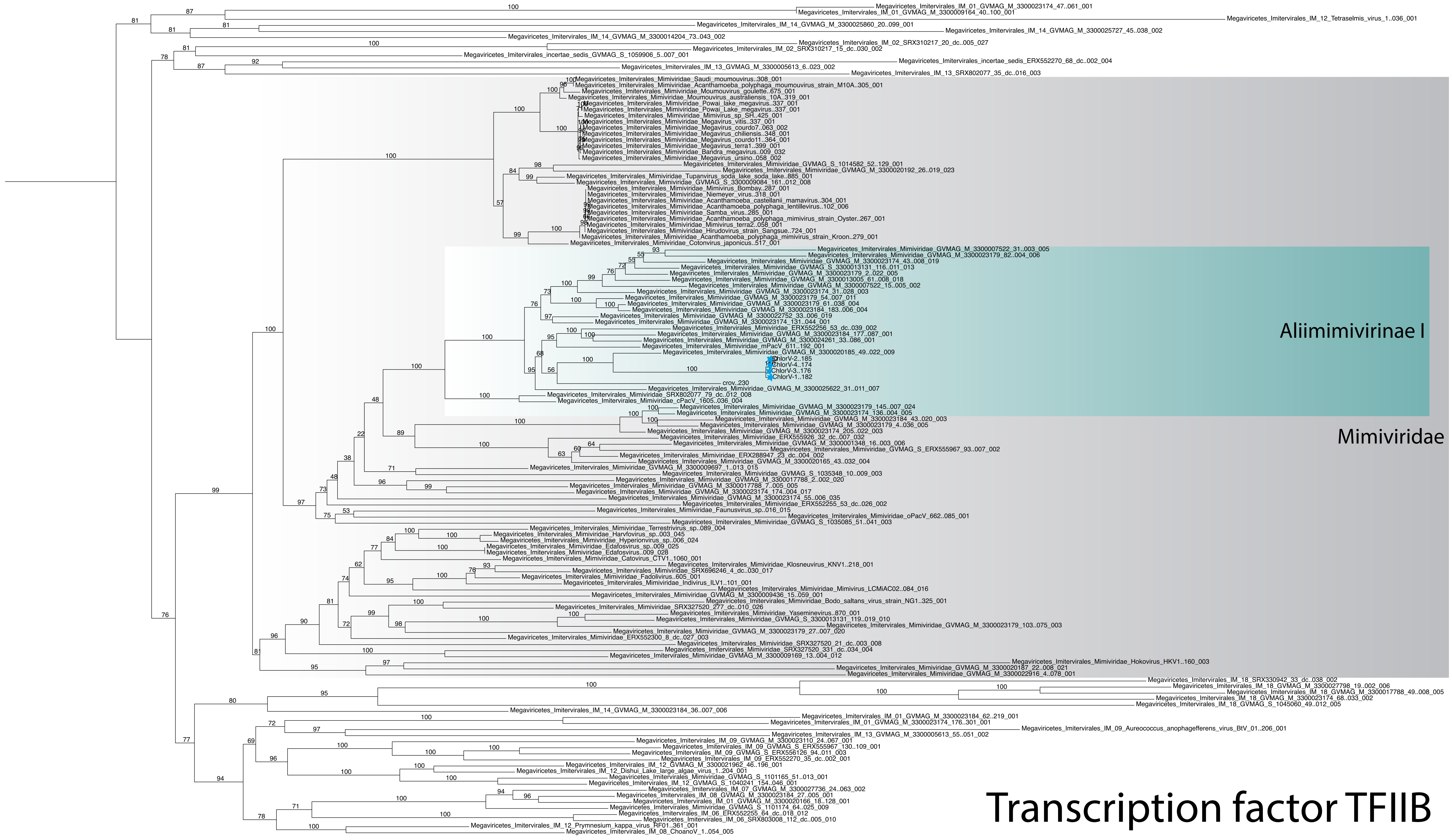

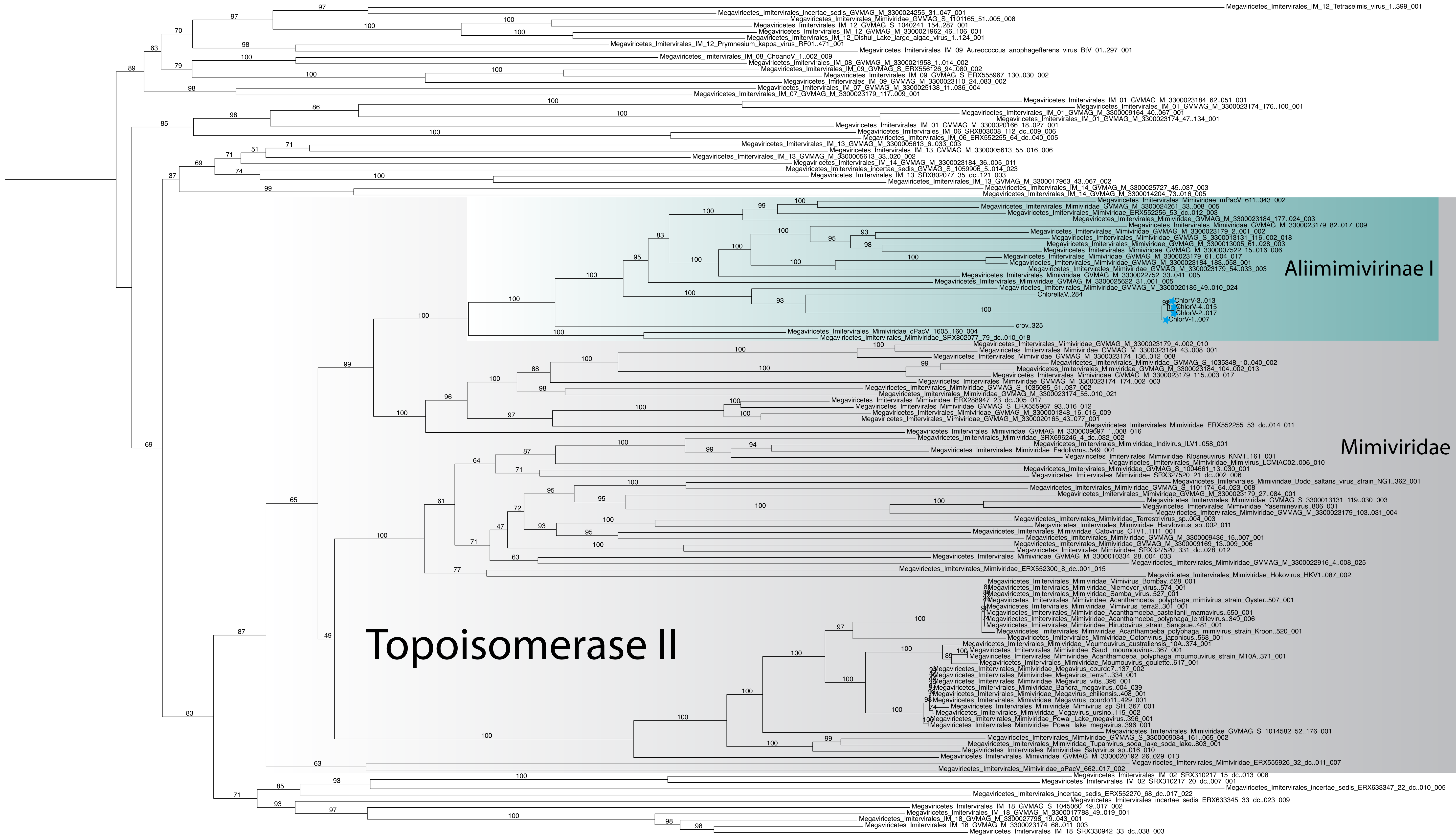

#### Virus late transcription factor 3

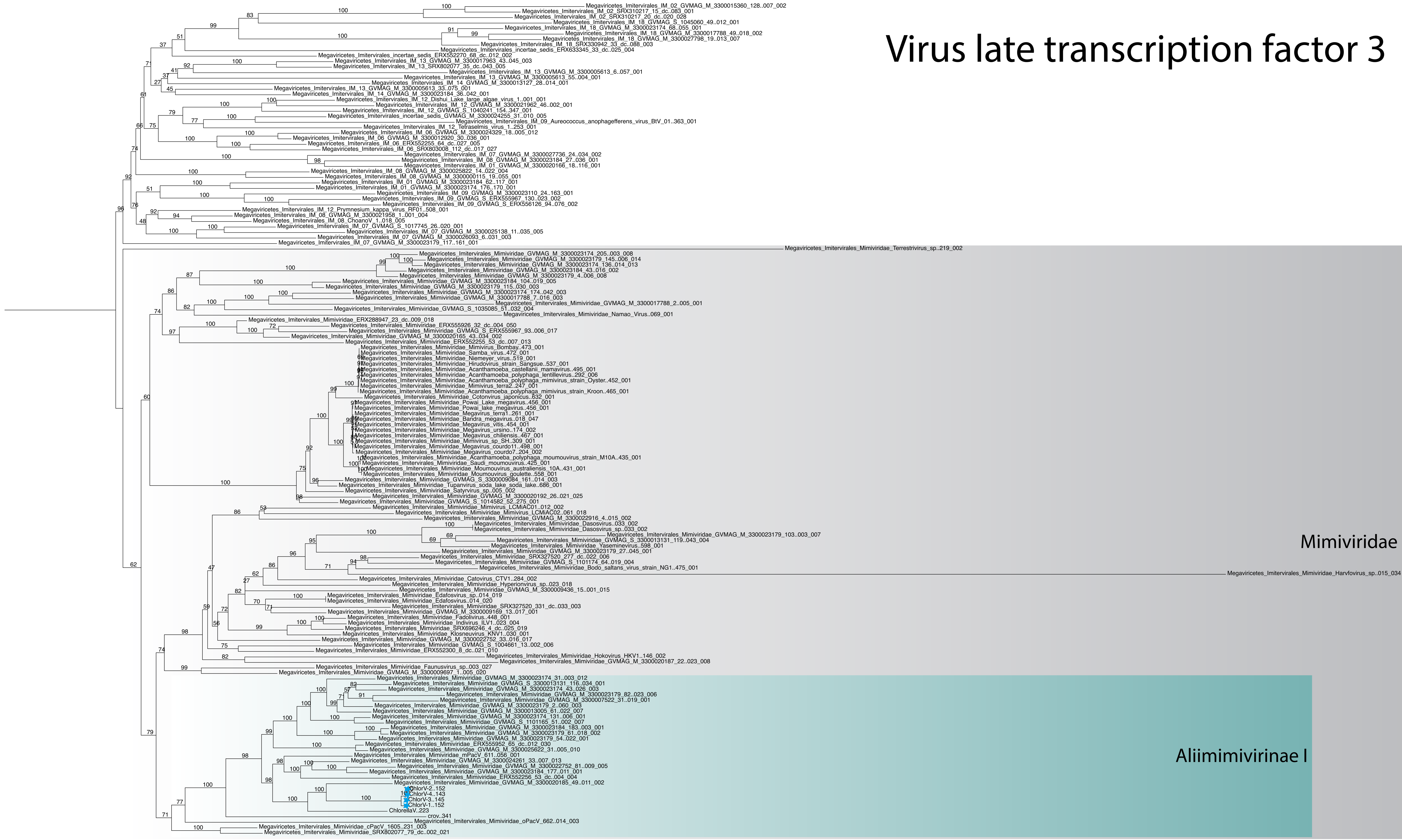
